# Disrupting linearity of PROTAC scaffolds prevents off-target complex I inhibition

**DOI:** 10.64898/2026.02.13.702718

**Authors:** Nick Richert, Hana Nůsková, Jana Samarin, Bozhidar S. Ivanov, Mohamad Saoud, Franziska Deis, Nicole de Vries, Sonja Sievers, Gernot Poschet, Slava Ziegler, Judy Hirst, Nikolas Gunkel, Aubry K. Miller

## Abstract

Chemical inducers of proximity have transformed small-molecule pharmacology, but the large, bifunctional architectures they often employ introduce new and poorly understood off-target risks. During a targeted protein degrader synthesis project, we identified a subset of compounds that cause rapid and unexpected ATP depletion in cells. Mechanistic studies traced this effect to inhibition of mitochondrial complex I, a central component of oxidative phosphorylation. This inhibition does not stem from off-target binding by either of the two target ligands, but from the overall long, linear architecture of the bifunctional molecules, which renders them effective ligands of the narrow ubiquinone binding tunnel of complex I. Strikingly, this liability extends to structurally unrelated bifunctional molecules, including six androgen receptor PROTACs including the clinical candidate ARV-110, which inhibits complex I at low nanomolar concentrations. To mitigate complex I inhibition, we established a generalizable design strategy to disrupt linear molecular geometry through the introduction of structural “bumps” or “kinks”. In a proof-of-concept study, we successfully apply this strategy to the ARV-110 scaffold, discovering potent AR-degrading ARV-110 analogs that do not inhibit complex I. These findings uncover a previously underappreciated structural determinant of off-target mitochondrial toxicity and establish new design principles for safer proximity-inducing therapeutics.

## Introduction

Targeted protein degradation has emerged over the past two decades as a particularly promising modality of chemically induced proximity in drug development^1–4^. Conventional small-molecule inhibitors block the function of their target protein only when physically occupying a binding site on the protein’s surface. Targeted protein degradation instead operates via a catalytic, event-driven mode of action that hijacks the cellular ubiquitin-proteasome system. Heterobifunctional degraders, also known as proteolysis targeting chimeras (PROTACs), consist of two chemically linked ligands that simultaneously bind the protein to be degraded (protein of interest) and an E3 ligase, generating a ternary complex. The induced proximity of the two proteins promotes ubiquitination of the protein of interest by the recruited E3 ligase complex, marking it for degradation by the proteasome even after the PROTAC has dissociated. The modular design of PROTACs allows for rapid adaptation to different targets. As a result, targeted protein degradation has generated immense interest in both academic and industrial settings^5^, with dozens of substances having entered clinical stages of development^2,6^.

We recently uncovered tetrathiolate zinc finger proteins as off-targets of proton pump inhibitors^7^, a class of prodrugs that covalently engage surface cysteine(s) of the gastric H^+^/K^+^-ATPase proton pump^8^, and are widely used in the treatment of peptic ulcers and gastroesophageal reflux. The low pH environment of parietal cell canaliculi in the stomach lining triggers prodrug activation via an acid-mediated Smiles rearrangement, thereby converting an otherwise inert sulfoxide functional group into a cysteine-reactive sulfenic acid moiety^9^. We found that coordination of proton pump inhibitors to the zinc ion in a tetrathiolate zinc finger can initiate non-canonical Lewis acid-mediated activation at neutral pH, resulting in covalent modification of one or more of the tetrathiolate cysteines.

The Ikaros zinc finger family are common targets and/or off-targets of immunomodulatory imide drug-based molecular glues and heterobifunctional degraders^10–13^. We aimed to target tetrathiolate zinc fingers as opposed to the Ikaros zinc finger family by converting the proton pump inhibitor rabeprazole into a heterobifunctional degrader. A library of rabeprazole-based bifunctional molecules did not degrade tetrathiolate zinc fingers, but a subset of them produced pronounced cytotoxicity. We traced this effect to inhibition of mitochondrial complex I, a multiprotein complex in the electron transport chain and an essential part of oxidative phosphorylation. Structure–activity relationship analysis revealed that a long, linear molecular core is essential for binding within the elongated, hydrophobic ubiquinone-binding tunnel of complex I, providing a structural basis for mitochondrial toxicity. Extending these findings to well-known degrader molecules, we found that six androgen receptor PROTACs—most notably the clinical candidate ARV-110—inhibit complex I, highlighting mitotoxicity as a general concern that should be addressed early in PROTAC development^14,15^. We explored multiple strategies to mitigate complex I binding and, thereby, mitotoxicity. Perturbing the linear geometry of the ARV-110 scaffold via linker and recruiter redesign preserved degrader potency while abolishing off-target complex I inhibition. Collectively, these findings establish structural design principles that enable the development of safer bifunctional degraders and provide a framework for mitigating mitochondrial toxicity in proximity-inducing therapeutics.

## Results

### Rabeprazole-thalidomide hybrids induce rapid loss of ATP

We began our investigation of the proton pump inhibitor rabeprazole as a PROTAC protein of interest recruiter by synthesizing a small library of heterobifunctional substances, selecting the methoxy group-bearing side chain of rabeprazole as exit vector^7,16^ (**Fig. 1a, Supplementary Fig. S1**). For the other terminus of the putative PROTACs, we chose the immunomodulatory imide drug thalidomide, which is commonly used to recruit cereblon, a substrate receptor of the Cullin-Ring E3 Ubiquitin Ligase 4 (CRL4) complex^17^. We employed flexible alkyl linkers of various lengths to produce **1**–**4**, as well as more rigid, cyclic structures^6^ to generate **5**–**10** (see **Supplementary Schemes S1**–**S5** for synthesis routes).

**Fig. 1.**
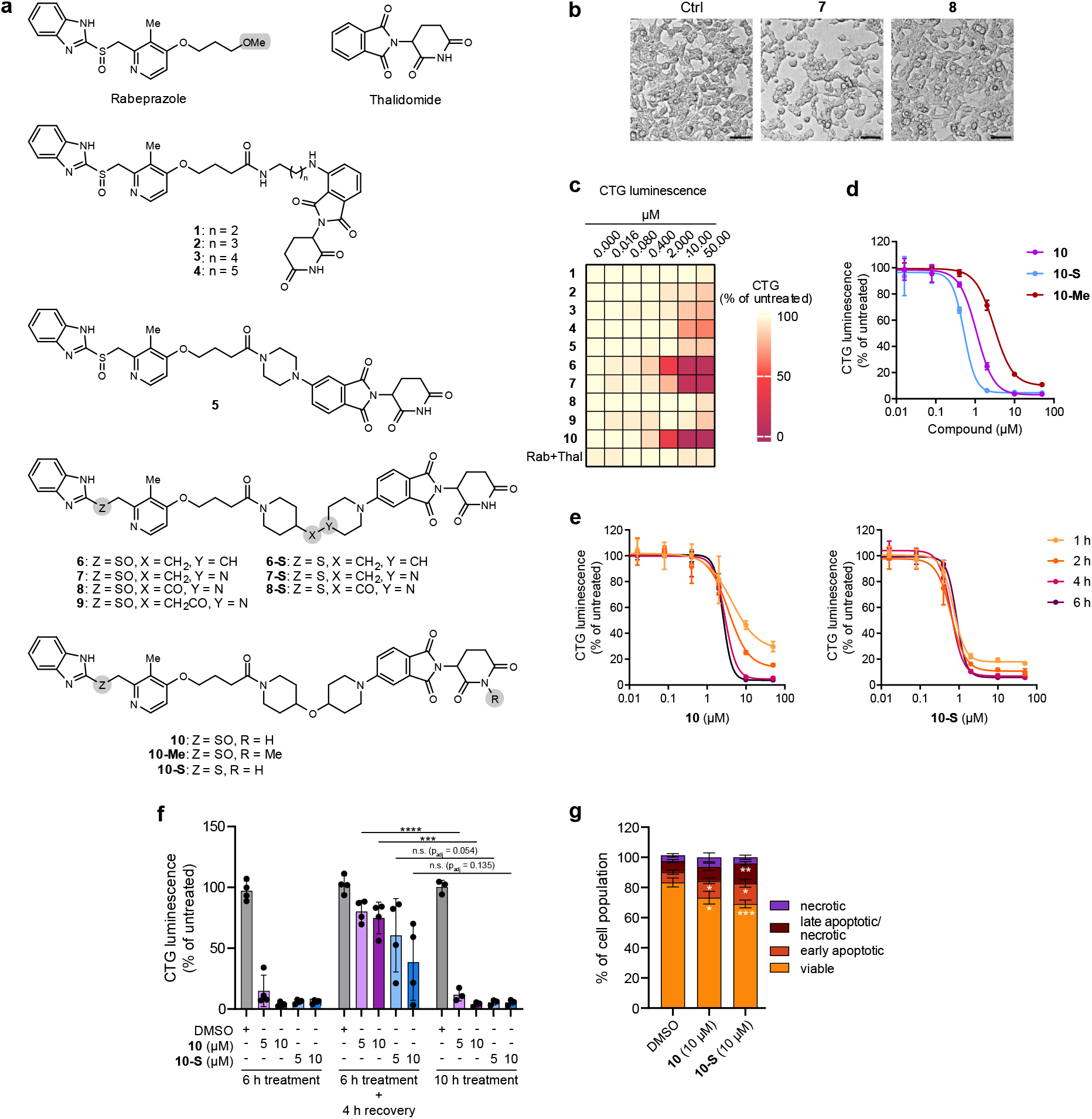
Rabeprazole-thalidomide hybrids induce rapid loss of ATP. **a**, Structures of rabeprazole-thalidomide hybrids. **b**, Representative (*n* = 3) bright-field images of HEK293T cells treated with **7** and **8** (10 µM each) for 6 h compared to DMSO control. Scale bar: 90 µm. **c**, CellTiter-Glo assay heatmap of HEK293T cells treated with rabeprazole-thalidomide hybrids for 6 h. Luminescence values were normalized to untreated control. Heatmap colors represent mean values (*n* ≥ 3; *m* = 3). Rab, rabeprazole; Thal, thalidomide. **d**, Representative CellTiter-Glo assay (*n* ≥ 3, *m* = 3) of **10, 10-S**, and **10-Me** in HEK293T cells after 6 h. Luminescence values were normalized to untreated control. Data points represent mean ± SD. **e**, Representative time-course CellTiter-Glo assay (*n* = 3, *m* = 3) of **10** and **10-S** in HEK293T cells. Luminescence values were normalized to untreated control. Data points represent mean ± SD. **f**, CellTiter-Glo washout and recovery assay (*n* = 4, *m* = 3) of **10** and **10-S** in HEK293T cells. Luminescence values were normalized to untreated control. Error bars represent mean ± SD. **g**, Flow cytometry analysis of cell death in HEK293T cells treated with 10 µM **10** and **10-S** for 6 h. Error bars represent mean ± SD. An example of gating strategy is shown in **Extended Data Fig. 1d**. The significant differences between DMSO control and treatments are indicated inside of the respective segments of the bar plot. All statistical analyses were determined by two-tailed unpaired *t*-tests with an adjustment for multiple comparisons using the Holm–Šídák method (\*\**p* < 0.01, \*\*\**p* < 0.001, \*\*\*\**p* < 0.0001). *n* = biological replicates, *m* = technical replicates.

We assessed the degrading activity of these compounds in HEK293T cells, as our study on non-canonical proton pump inhibitor activation was conducted primarily in this cell line^7^. Cells were treated with each substance for 6 h followed by immunoblotting of the rabeprazole-target proteins IRF2BP2, RNF156, ZYX, UBR7, and DENR (**Extended Data Fig. 1a**). None of the compounds induced reproducible protein degradation. Only slight decreases in short-lived proteins^18^ (e.g., IRF2BP2 and RNF156, **Extended Data Fig. 1b**) were induced at higher concentrations of some samples.

We noted, however, that some substances promoted cell rounding and well detachment at micromolar concentrations, providing an initial hint of a structure–activity relationship. For example, tertiary amine **7** induced cell rounding whereas its corresponding amide **8** had no effect (**Fig. 1b**). The substances **6** and **10** also promoted cell rounding and detachment at micromolar concentrations, indicating rapid cytotoxic effects (**Extended Data Fig. 1c**). We therefore treated HEK293T cells with the same set of ten compounds for 6 h in the CellTiter-Glo® (CTG) assay, which measures cellular ATP levels as a proxy for cell viability^19^. Consistent with our visual observations, the three substances that promoted cell-rounding (**6, 7**, and **10**) produced a dose-dependent reduction in CTG luminescence (**Fig. 1c**). Likewise, compounds that did not induce visible cellular stress, namely **1**–**4** and piperazine amides **5, 8**, and **9**, were largely inactive. Furthermore, a combination treatment of rabeprazole and thalidomide had no effect. Notably, both *N*-methyl glutarimide **10-Me** and sulfide **10-S** (**Supplementary Scheme S5**), monofunctional analogs of **10** that lack the ability to recruit cereblon and tetrathiolate zinc fingers, respectively, reduced ATP levels with potencies comparable to **10** (**Fig. 1d**). Sulfides **6-S, 7-S**, and **8-S** (**Supplementary Schemes S3** and **S4**), which, like **10-S**, do not recruit zinc fingers because they cannot undergo the prodrug Smiles rearrangement^9,20^, likewise mirror the activity of their respective parent sulfoxides **6**–**8** (**Extended Data Fig. 1d**). These findings suggest the observed cellular effects are neither a result of cereblon recruitment, nor of covalent modification of tetrathiolate zinc finger-containing proteins; however, the appropriate linking of both substances (or derivatives thereof) in one molecule is required.

We selected the two most potent and effective hybrids, sulfoxide **10** and sulfide **10-S**, for subsequent experiments, noting that rabeprazole-based sulfides (e.g., **10-S**) are preferable due to their greater chemical stability^9^ as the sulfoxide-dependent Smiles rearrangement appeared not to be not required for activity. A time-course CTG experiment with **10** and **10-S** showed a reduction in ATP levels even after 1 h of incubation (**Fig. 1e**). However, cells significantly recovered their ability to produce ATP within 4 h of removing **10** via medium change; a similar trend was observed with **10-S**, though the effect did not reach statistical significance (**Fig. 1f**). Cell staining with propidium iodide and annexin V showed small, statistically significant changes in apoptosis and necrosis when cells were treated for 6 h (**Fig. 1f, Extended Data Fig. 1g**). The small magnitude of the changes suggest that the cells are under stress, but that the almost complete loss of CTG signal within this short time frame is not a result of cell death. Taken together, our results suggest that these hybrid molecules induce rapid and reversible depletion of cellular ATP levels in HEK293T cells in a cereblon and proton pump inhibitor-target independent manner.

### Rabeprazole-thalidomide hybrids inhibit mitochondrial complex I

Based on the above findings, we postulated that a mechanism not involving ternary-complex formation was operative and asked whether mitochondrial respiration could be unexpectedly targeted by our substances. Mitochondrial respiration, the primary driver of aerobic ATP synthesis^21^, is mediated by a series of membrane-bound protein complexes (complexes I–V) in the inner mitochondrial membrane. We therefore measured the oxygen consumption rate (OCR) in a real-time extracellular flux analysis in HEK293T cells (**Fig. 2a**, left panel)^22^. Consistent with our hypothesis, **10-S** produced a rapid, dose-dependent decline in respiration, similar to the complex I inhibitor rotenone. In contrast, amide **8**, which was inactive in the CTG assay, had no effect up to 9 µM. We also monitored the extracellular acidification rate (ECAR) as a complementary readout of glycolytic activity (**Fig. 2a**, right panel)^23^. As expected, ECAR increased in HEK293T cells upon treatment with rotenone, whereas amide **8** had no detectable effect. In contrast, **10-S** reduced the ECAR, indicating concurrent inhibition of respiration and glycolysis, thereby severely limiting ATP production.

**Fig. 2.**
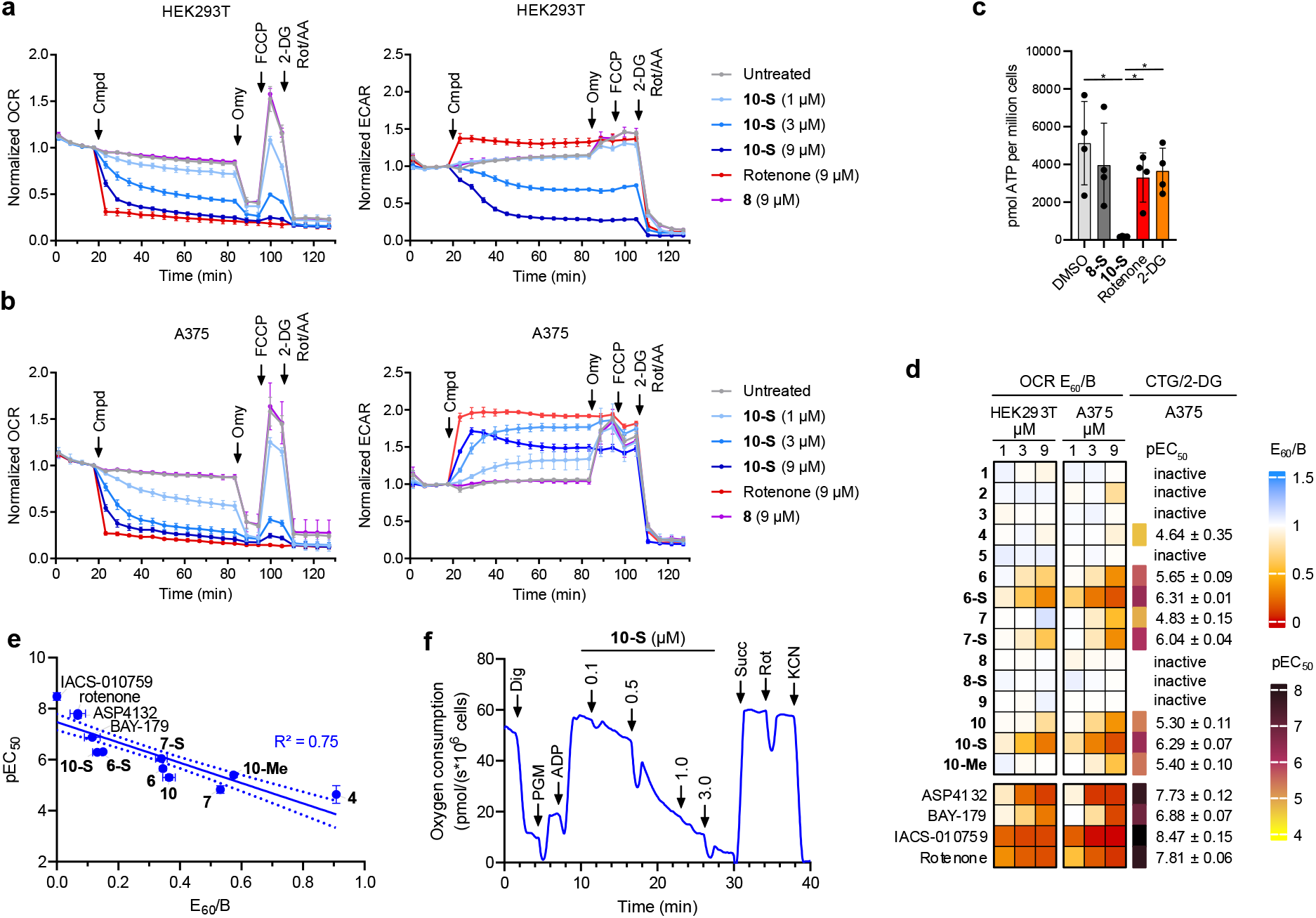
Rabeprazole-thalidomide hybrids inhibit mitochondrial complex I. **a, b**, Representative (*n* = 3) real-time extracellular flux experiments (*m* = 3–6, presented as mean ± SD) in HEK293T or A375 cells following the oxygen consumption rate (OCR) and extracellular acidification rate (ECAR). Rotenone was used as a positive control. Indicated injections: Cmpd, compound; Omy, 1 µM oligomycin, FCCP, 1 µM carbonyl cyanide-*p*-trifluoromethoxy-phenylhydrazone; 2-DG, 100 mM 2-deoxyglucose; Rot, 1 µM rotenone; AA, 0.5 µM antimycin A. **c**, Targeted metabolic quantification of ATP in HEK293T cells treated with 10 µM **8-S**, 10 µM **10-S**, 10 µM rotenone or 50 mM 2-deoxyglucose (2-DG) for 6 h. Data are presented as mean ± SD (*n* = 4). Statistical significance was determined by one-way ANOVA (ATP *p* = 0.0025) followed by two-tailed unpaired *t*-tests with an adjustment for multiple comparisons using the Holm–Šídák method to assess differences between conditions (\**p* < 0.05, \*\**p* < 0.01, \*\*\**p* < 0.001, \*\*\*\**p* < 0.0001). **d**, Heatmap of OCR E_60_/B values from HEK293T and A375 cells and results from the CTG/2-DG assay in A375 cells. E_60_/B values were determined as described in **Extended Data Fig. 3a**. OCR E_60_/B heatmap colors represent mean values (*n* = 1–3; *m* = 6). pEC_50_ values are mean ± SD (*n* = 3, *m* = 3). Inactive compounds are defined as reducing CTG luminescence < 50% at the highest tested concentration (50 µM). **e**, Correlation of OCR E_60_/B values from the extracellular flux assay (9 µM) and the CTG/2-DG pEC_50_ values from **Fig. 2d**. Simple linear regression: R^2^ = 0.75, *p* < 0.0001. **f**, A representative (*n* = 3) high resolution respirometry experiment in A375 cells. Concentrations of the injected compound **10-S** indicate the final concentration after each injection. Indicated injections: Dig, 24 µM digitonin; PGM, 10 mM pyruvate + 10 mM glutamate + 3 mM malate; ADP, 1 mM ADP; Succ, 10 mM succinate; Rot, 1 µM rotenone; KCN, 1 mM KCN. *n* = biological replicates, *m* = technical replicates.

To probe this finding with an orthogonal approach, we performed targeted metabolomic profiling of HEK293T cells treated with **10-S** as well as amide **8-S** as an inactive control. We included rotenone and 2-deoxy-D-glucose (2-DG), an inhibitor of the glycolysis ATP-synthesis pathway, as additional controls (**Fig. 2c**). While **8-S** had no statistically significant effect on cellular ATP levels, **10-S** treatment resulted in a complete loss of ATP. This is consistent with the performance of **10-S** in the CTG assay (see **Fig. 1d**) and its effect on both the OCR and ECAR (see **Fig. 2a**). Both rotenone and 2-DG appeared to lower ATP levels relative to DMSO control, though statistical significance was not achieved, indicating that HEK293T cells can adjust their metabolism in response to selective respiration and glycolysis inhibitors. In addition, **10-S** treatment significantly reduced guanosine-5′-triphosphate (GTP) levels, accompanied by significantly increased levels of the precursors AMP, GDP and, most strikingly, GMP (**Extended Data Fig. 2a**).

We wondered whether other cell lines would be similarly affected and tested **10** and **10-S** in the CTG assay against a panel of 16 human cell lines comprising lung and skin cancer entities, as well as immortalized, non-transformed cell lines (**Extended Data Fig. 2b**). HEK293T ATP levels were most strongly affected, with the other cell lines exhibiting a spectrum of responses. This was more apparent with sulfide **10-S** where A549, A375, LFX289, and H1693 cells appeared to be little affected, though the same cell lines were also the least affected by **10**. Further analysis of the A375 melanoma cell line surprisingly revealed that **10-S** more strongly reduced OCR in A375 cells than in HEK293T cells (**Fig. 2b**, left panel). Consistent with their limited response to **10-S** in the CTG assay (**Extended Data Fig. 2b**), A375 cells treated with **10-S** were able to compensate with increased glycolysis, thereby maintaining ATP production (**Fig. 2b**, right panel). This indicates that **10-S** inhibits respiration in both, and presumably all tested, cell lines in a specific manner, while the glycolysis inhibition observed in HEK293T cells is cell-line dependent.

We next measured the OCR of all fifteen rabeprazole-thalidomide hybrids alongside thalidomide, rabeprazole, and the mitochondrial respiration inhibitors rotenone, IACS-010759^24^, ASP4132^25^, and BAY-179^26^ (**Supplementary Fig. S2**), in both HEK293T and A375 cells. To quantify and compare activities between different substances with respect to respiration inhibition, we defined the metric E60/B as the ratio of the inhibitory effect 60 minutes after injection (E60) to the basal respiration rate just before injection (B) (**Extended Data Fig. 3a**; see **Extended Data Fig. 4** for complete extracellular flux data table). Only compounds that were active in the CTG assay (e.g., **6-S** and **10-S**) reduced respiration in both cell lines, whereas all CTG-inactive compounds (**1**–**5, 8, 8-S**, and **9**) had no effect on the OCR (**Fig. 2d**). This confirms that the observed cell-rounding and loss of ATP was connected to complex I inhibition. Activities across the two cell lines were consistent, with A375 cells showing overall higher sensitivity. We selected A375 cells for further respiration analyses owing to their higher OCR sensitivity and to separate respiration inhibition from the glycolytic impairment observed in HEK293T cells.

Real-time extracellular flux experiments provide valuable kinetic information, and the E60/B metric enables comparison of substances at individual concentrations. To establish an additional accurate, simple, and widely accessible method for comparing respiration inhibitors, we developed a CTG-based assay in A375 cells that enables precise pEC_50_ determinations. In this assay, A375 cells are treated concomitantly with the rapid-acting glycolysis inhibitor 2-DG and a candidate respiration inhibitor, and ATP levels are quantified after 2 h using CTG. As expected, the respiration inhibitors rotenone and BAY-179 reduced ATP levels in A375 cells only in the presence of 2-DG, whereas amides **8** and **8-S** had no effect under either condition, confirming assay specificity (**Extended Data Fig. 3b**). Literature-known inhibitors of mitochondrial respiration show the strongest effects in this CTG/2-DG assay (**Fig. 2d**) (note: potencies are hereafter reported as (pEC50 ± SD; EC50) where pEC50 = –log(EC50))^27^. Within our series, **6-S** and **10-S** are the most potent, exhibiting comparable activities (pEC50 = 6.29 ± 0.07; 0.51 µM and 6.31 ± 0.01; 0.49 µM, respectively). In contrast, compounds that were inactive in the extracellular flux assay (**1**–**5, 8, 8-S**, and **9**) showed minimal activity in the CTG/2-DG assay. Notably, pEC50 values determined from the CTG/2-DG assay correlate (*R*^2^ = 0.75) with the extracellular flux E60/B values (**Fig. 2e**), indicating that the CTG/2-DG assay can be used as a rapid screening method to discover respiration inhibitors.

We next sought to determine whether one of the five complexes within the electron transport chain was specifically targeted by these compounds, and performed high resolution respirometry in A375 cells^28^. A titration of compound **10-S** caused the expected decrease in oxygen consumption, showing an effect as low as 0.1 µM, while 3.0 µM resulted in complete suppression (**Fig. 2f**). Respiration was restored upon supplementation with the complex II substrate succinate, pinpointing **10-S** and this class of rabeprazole-thalidomide hybrids as mitochondrial complex I inhibitors. Complex I (CI), the primary entry point for electrons into the respiratory chain, thus emerges as the molecular origin of the observed mitotoxicity.

### Modified rabeprazole-thalidomide hybrids must be long and linear to inhibit complex I

We next set out to define the structural features of the rabeprazole–thalidomide hybrids that drive CI inhibition. To this end, we synthesized derivatives of **10-S**, systematically introducing truncations and modifications on both the rabeprazole (western) and thalidomide (eastern) moieties^29^ (**Fig. 3a, Supplementary Fig. S3, Supplementary Schemes S4** and **S5**). All substances were evaluated in the extracellular flux and CTG/2-DG assays in A375 cells (**Fig. 3a, Extended Data Fig. 4**). *N*-Methylation of the benzimidazole ring in **10-S** yielded **11** (pEC50 = 6.15 ± 0.12; 0.71 µM) with CI activity similar to **10-S**, while truncation to imidazole **12** reduced potency (pEC50 = 5.49 ± 0.12; 3.2 µM). Substitution of the imidazole in **12** with a phenyl ring (**13**), increased potency by one order of magnitude (pEC50 = 6.46 ± 0.10; 0.35 µM). Removal of the thioether moiety was tolerated, yielding simplified CI inhibitors **14** (pEC50 = 6.20 ± 0.06; 0.63 µM) and **15** (pEC50 = 5.74 ± 0.11; 1.8 µM). Further truncation to benzamide **16** (pEC50 = 4.80 ± 0.12; 16 µM) and acetamide **17** (inactive) diminished potency, identifying the limits of truncation on the western side. On the eastern side, exchanging the glutarimide ring of **10-S** for a lactam (**19**, pEC50 = 5.73 ± 0.05; 1.9 µM) or a cyclohexyl ring (**20**, pEC50 = 6.31 ± 0.07; 0.49 µM) was tolerated, while replacement with a phenyl group enhanced potency (**21**, pEC50 = 6.77 ± 0.03; 0.17 µM). Further truncation on the eastern part of the molecule to *N*-methylphthalimide **22** and aniline **23** reduced potencies (pEC50 = 6.38 ± 0.06; 0.42 µM and 5.94 ± 0.04; 1.1 µM, respectively) in the CTG/2-DG assay. *N*-methyl amine **24** was weakly active in both assays, identifying the limits of truncation on the eastern side.

**Fig. 3.**
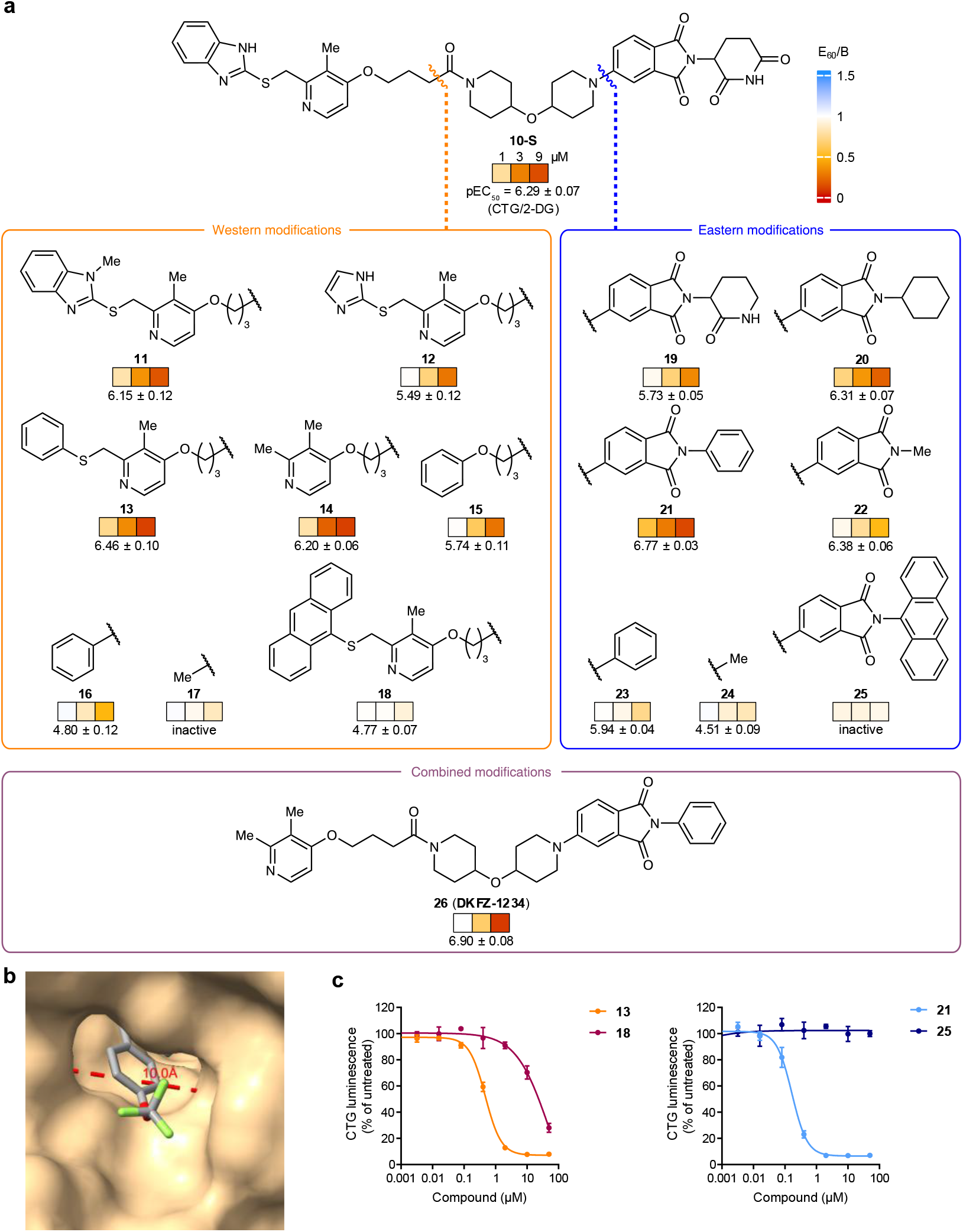
Modified rabeprazole-thalidomide hybrids must be long and linear to inhibit complex I. **a**, Truncations and modifications of **10-S**. Modifications in the orange box are linked to the substructure that is right of the orange intersecting line; modifications in the blue box are linked to the substructure that is left of the blue intersecting line. Heatmap of OCR E_60_/B values and pEC_50_ values from the CTG/2-DG assay in A375 cells are depicted under each structure. E_60_/B values were determined as described in **Extended Data Fig. 3a**. OCR E_60_/B heatmap colors represent mean values (*n* = 1–3; *m* = 6). pEC_50_ values are means ± SD (*n* = 3, *m* = 3). **b**, Reproduction of the cryo-EM structure of mammalian CI bound by IACS-2858 (PDB: 7B93). The indicated distance was measured from the −SMe methyl carbon of Met225 to C-3 of Pro48 in the ND1 subunit. **c**, Representative CTG/2-DG assay (*n* = 3, *m* = 3) of **13, 18, 21**, and **25**. Luminescence values were normalized to untreated control. Data points represent mean ± SD. *n* = biological replicates, *m* = technical replicates.

As several modifications on both the western and eastern sides retained or enhanced CI inhibition, we asked whether these contributions would be additive. We therefore combined the modifications of **14** and **21** to generate **26** (**DKFZ-1234**) (pEC50 = 6.90 ± 0.08; 0.13 µM), which is equipotent with the chemical probe BAY-179 (pEC50 = 6.88 ± 0.07; 0.13 µM, see **Fig. 2d**). Together, these data show that CI inhibition arises from distributed structural features across both the western and eastern moieties, rather than from either fragment alone. Combining modifications on both sides results in additive effects on potency, and only extensive truncation markedly reduces activity.

Upon cursory inspection, many reported CI inhibitors appear structurally unrelated to each other (see **Supplementary Fig. S2** for structures), as well as to our rabeprazole-thalidomide hybrids. Nevertheless, it is generally assumed that many CI inhibitors bind in the long and narrow ubiquinone-binding channel, where they must adopt extended linear conformations. This is highlighted by a recent cryo-EM structure of mammalian CI bound by IACS-2858 (analog of IACS-010759), which revealed IACS-2858 bound to the ubiquinone-binding tunnel of the protein complex in an elongated conformation^30^. In principle, many apparently unrelated scaffolds could fit this description. Using the IACS-2858 CI cryo-EM structure (PDB: 7B93), we estimated the width of the ubiquinone-binding tunnel opening to be ca. 10.0 Å (**Fig. 3b**). The trifluoromethoxy-substituted phenyl ring of IACS-2858, which protrudes from the tunnel entry, can be seen to fill this opening with little remaining free volume. We therefore designed two derivatives of compound **10-S** bearing 9-anthracenyl substitutions on the western (**18**) and eastern (**25**) moieties, respectively. These T-shaped molecules should be too wide to pass through the tunnel opening. Consistent with our hypothesis, **18** and **25** were inactive in the extracellular flux assay (**Fig. 3a**) and significantly less active (**18** vs. **13**) or completely inactive (**25** vs. **21**) in the CTG/2-DG assay (**Fig. 3c**). Taken together, our data suggest that rabeprazole-thalidomide hybrids must be relatively long and capable of adopting a linear conformation to inhibit CI. Because simplified Western truncations like **14** and **15** retain activity, it appears as if substituted thalidomide derivatives may have an inherent affinity for CI.

### Mitochondrial complex I is an off-target of clinically investigated degraders

With a structural principle for CI binding of rabeprazole-thalidomide hybrids and analogs in hand and given the widespread use of cereblon-recruiting moieties in bifunctional degraders and molecular glues, we asked whether other linear heterobifunctional molecules could inhibit mitochondrial respiration. We therefore tested a panel of thirteen well-characterized bifunctional degraders containing a variety of protein of interest- and cereblon-recruiting motifs (**Supplementary Fig. S4**). Notably, three clinically-investigated androgen receptor (AR) degraders inhibited respiration with E60B ≤ 0.40 at 9 µM: ARV-110 (E60/B = 0.12), ARV-766 (E60/B = 0.40), and BMS-986365 (E60/B = 0.17) (**Fig. 4a, Extended Data Fig. 4**)). In the CTG/2-DG assay, ARV-110 and BMS-986365 had nanomolar potencies (pEC50 = 6.58 ± 0.09; 0.26 µM and 6.22 ± 0.12; 0.60 µM, respectively). ARV-766 also showed sub-micromolar activity that plateaued at 34% maximum inhibition (**Extended Data Fig. 5a**). Notably, all three degraders harbor different AR-as well as cereblon-recruiting moieties (**Fig. 4b**). We therefore tested four additional AR-degraders (**Supplementary Fig. S4**) and found that all three cereblon-recruiting PROTACs (ARD-2051, AZ’3137, and BWA-522) were active in both assays (**Fig. 4c**); only the VHL-recruiting ARD-266 was inactive. The most potent of the six AR-degraders, ARV-110, rapidly decreased the OCR in a high resolution respirometry experiment at nanomolar concentrations, confirming CI as its primary target (**Fig. 4d**).

**Fig. 4.**
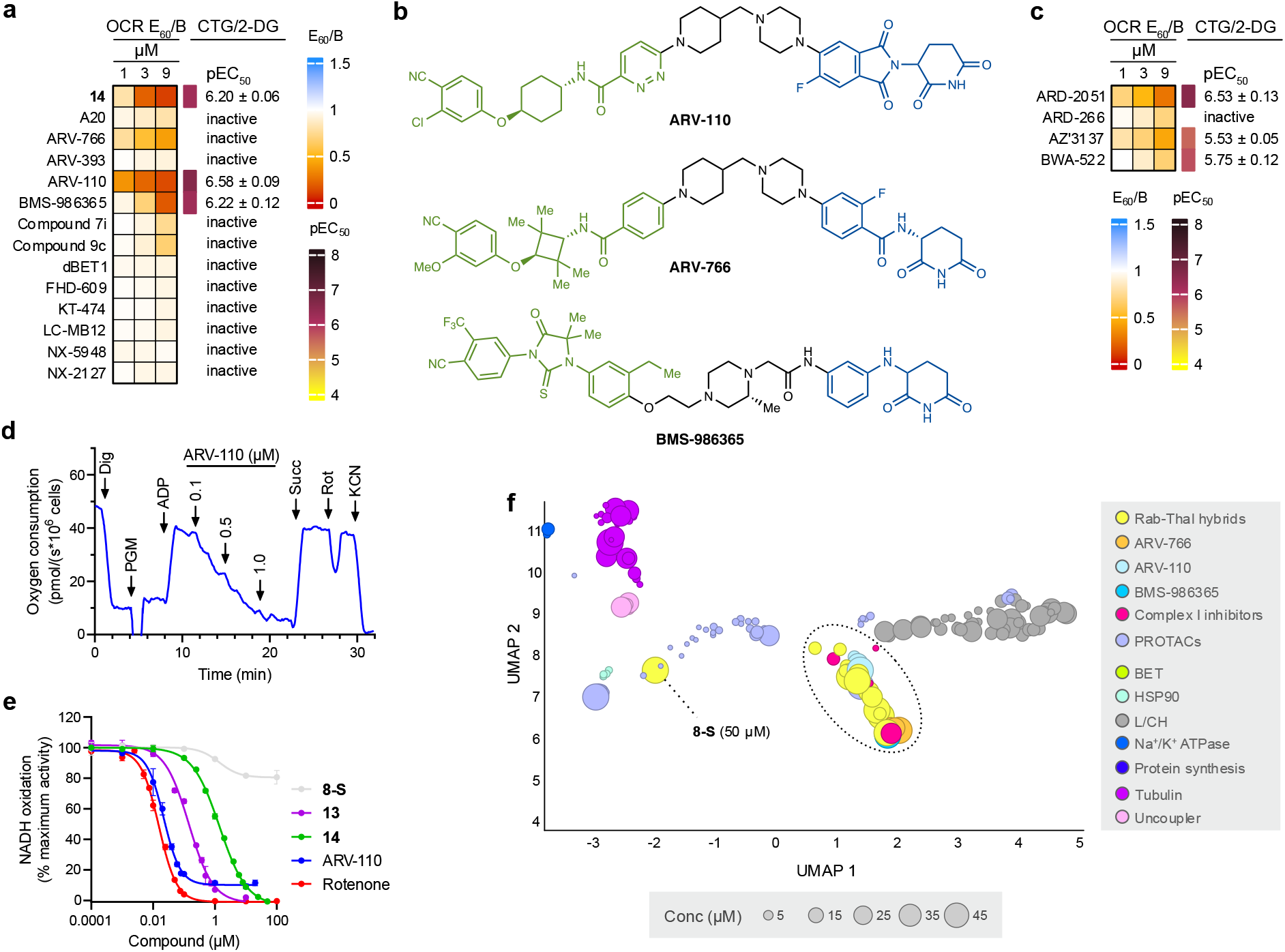
Mitochondrial complex I is an off-target of clinically investigated degraders. **a**, Heatmap of OCR E_60_/B values and pEC_50_ values from the CTG/2-DG assay in A375 cells. E_60_/B values were determined as indicated in **Extended Data Fig. 3a**. OCR E_60_/B heatmap colors represent mean values (*n* = 1–3; *m* = 6). pEC_50_ values are means ± SD (*n* = 3, *m* = 3). **b**, Structures of clinically investigated AR degraders ARV-110, ARV-766, and BMS-986365. The AR recruiter is highlighted in green, E3 ligase recruiter in blue. **c**, Heatmap of OCR E_60_/B values and pEC_50_ values from the CTG/2-DG assay in A375 cells. OCR E_60_/B heatmap colors represent mean values (*n* = 2; *m* = 6). pEC_50_ values are means ± SD (*n* = 3, *m* = 3). **d**, A representative (*n* = 3) high resolution respirometry experiment in A375 cells. Concentrations of the injected compound ARV-110 indicate the final concentration after each injection. Indicated injections: Dig, 30 µg·mL^-1^ digitonin; PGM, 10 mM pyruvate + 10 mM glutamate + 3 mM malate; ADP, 1 mM ADP; Succ, 10 mM succinate; Rot, 1 µM rotenone; KCN, 1 mM KCN. **e**, Dose–response curves of CI activity (*n* = 1; *m* = 4) measured as NADH:O_2_ oxidoreduction in isolated bovine mitochondrial membranes. Activities were normalized to the DMSO control (=100%). Data points represent mean ± SD. **f**, Zoomed-in view of the UMAP plot in **Extended Data Fig. 6a**. All 7 tested rabeprazole-thalidomide hybrids, 3 AR PROTACs, 16 other PROTACs, and 2 reference CI inhibitors (BAY-179 and ASP4132) are contained in the zoom along with 5 of 13 previously identified clusters. The size of the symbols corresponds to concentration. Not normalized, 10 neighbors. L/CH: lysosomotropism/cholesterol homeostasis. *n* = biological replicates, *m* = technical replicates.

We tested ARV-110 alongside two active truncated CI inhibitors (**13** and **14**) and inactive piperazine amide **8-S** in an in vitro, cell-free CI enzymatic assay, which uses isolated bovine (*Bos taurus*) mitochondrial membrane fragments and measures NADH consumption (**Fig. 4e**)^31^. ARV-110 (pIC50 = 7.66; 22 nM) was found to be roughly equipotent with rotenone (pIC50 = 7.81; 15 nM). We attribute incomplete maximum inhibition (90% maximum effect) from ARV-110 to its high lipophilicity (logD7.4 > 5.29) and low aqueous solubility (<0.03 µM) (**Extended Data Fig. 5b**), which results in precipitation at higher concentrations and limited exposure in the assay (**Extended Data Fig. 5c**). In comparison, compound **14** (pIC50 = 5.85; 1.4 µM), one of the more potent substances in the extracellular flux and CTG/2-DG assays (see **Fig. 3a**), showed complete inhibition, consistent with its lower lipophilicity (logD7.4 = 2.36) and higher aqueous solubility (105 µM) (**Extended Data Fig. 5b**). Such differences in physicochemical properties between ARV-110 and **14** may account for their divergent IC50 values in the cell-free assay (**Fig. 4e**) but comparable activities in the cell-based extracellular flux and CTG/2-DG assays (**Fig. 4a**), possibly reflecting the well-recognized permeability limitations of bifunctional degraders^6^. Phenyl-thioether **13** (logD7.4 = 4.63) showed intermediate potency (pIC50 = 6.84; 0.14 µM), consistent with its activity in the extracellular flux and CTG/2-DG assays (see **Fig. 3a**), while **8-S** (logD7.4 = 2.96) was confirmed to be inactive.

We next used Cell painting^32,33^, which can resolve a wide range of mode of actions, to morphologically profile 579 features of cells treated with a selection of active rabeprazole-thalidomide hybrids (**10-S, 13**–**15, 20**, and **21**), inactive control **8-S**, ARV-110, ARV-766, BMS-986365, and reference CI inhibitors BAY-179 and ASP4132. Substances were tested at 10 µM, 30 µM, and 50 µM, and most compounds produced significant morphological changes (>5% induction, for explanation see Supplementary Information) at all concentrations. Negative control amide **8-S** did so only at 50 µM. The multidimensional data was compared to profiles of substances previously measured in the Cell painting assay^33,34^, including sixteen unrelated PROTACs (**Fig. 4f, Extended Data Fig. 6a, Supplementary Fig. S5**). In this comparison, the six CI-inhibiting rabeprazole-thalidomide analogs **10-S, 13**–**15, 20**, and **21**, the three AR degraders, and the CI inhibitors BAY-179 and ASP4132 create a cluster that is distinct from any of the previously identified thirteen clusters^33,34^. Notably, amide **8-S**, which was inactive in the cellular flux, the CTG/2-DG, and the enzymatic assay, is not a member of the CI cluster. Some of the other tested PROTACs are found near the CI cluster, and MZ1 is found within the cluster, albeit only at 50 µM. A cross similarity heatmap of the profiles of the newly tested substances highlights the high biosimilarity of this set of substances to each other (**Extended Data Fig. 6b**). These profiles are very different from the profile of **8-S** and the clusters that include complex III or V inhibitors (pyrimidine biosynthesis^35^ and MitoStress cluster^34^, respectively, **Extended Data Fig. 6a** and **Supplementary Table S1**). Taken together, this set of data shows that clinically investigated AR degraders function as CI inhibitors and phenotypically cluster with established and newly identified CI inhibitors.

### Strategies to mitigate complex I inhibition while retaining degrader activity

Considering that numerous structurally distinct cereblon-recruiting molecules inhibit CI, we set out to develop a design strategy to attenuate off-target CI inhibition while preserving on-target degrader activity. The introduction of a wide T-shaped functional group on any bifunctional terminus (e.g., anthracenes **18** and **25**) should strongly reduce CI inhibition. Such a modification would, in most cases, abolish the primary biological activity of a bifunctional molecule, as the termini are usually highly optimized ligands. We therefore considered two conceptually related approaches based on perturbing the overall geometry and conformational landscape of a bifunctional scaffold. Both strategies rely on producing structures that should be unable to enter and occupy the narrow ubiquinone-binding tunnel of CI (**Fig. 5a**). One approach is to introduce a local bump within the linker region through branching or increased steric and conformational constraint^36^. Given the central role of the linker structure in tuning ternary complex formation as well as physicochemical, metabolic, and pharmacokinetic properties, we reasoned this region of a PROTAC might tolerate such geometric perturbations more readily than the recruiting termini. The second approach is to impose a pronounced kink in the molecular connectivity, thereby encoding a persistent L-shaped architecture into the scaffold. This could be accomplished via adjustment of the exit vector from, or the overall shape of, the E3 ligase- or protein of interest-recruiting moiety.

**Fig. 5.**
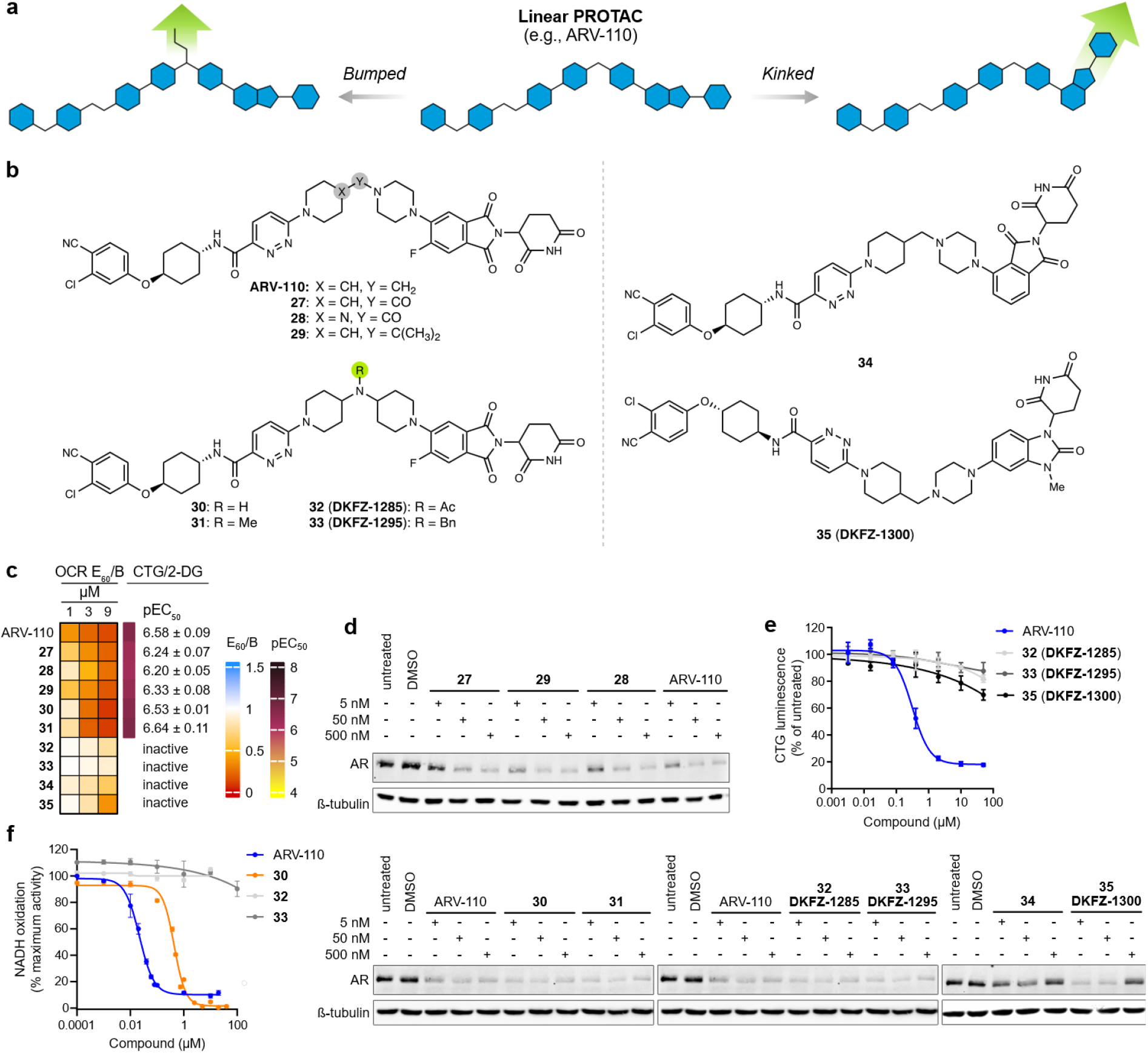
Strategies to mitigate complex I inhibition while retaining degrader activity. **a**, Design principle of introducing bumps and kinks into a bifunctional molecule to abolish CI inhibition. **b**, Structures of ARV-110 analogs. **c**, Heatmap of OCR E_60_/B values and pEC_50_ values from the CTG/2-DG assay in A375 cells. E_60_/B values were determined as indicated in **Extended Data Fig. 3a**. OCR E_60_/B heatmap colors represent mean values (*n* = 1–3; *m* = 6). pEC_50_ values are means ± SD (*n* = 3, *m* = 3). **d**, Representative immunoblots (*n* = 2–3) of AR after 6 h treatment with ARV-110 and derivatives in A375 cells. **e**, Representative CTG/2-DG assay (*n* = 3, *m* = 3) of ARV-110, **32, 33**, and **35**. Luminescence values were normalized to untreated control. Data points represent mean ± SD. **f**, Dose–response curves of CI activity (*n* = 1; *m* = 4) measured as NADH:O_2_ oxidoreduction in isolated bovine mitochondrial membranes. Activities were normalized to the DMSO control (=100%). Data points represent mean ± SD. *n* = biological replicates, *m* = technical replicates.

To test these design principles, we selected ARV-110 as a potent CI-inhibiting model PROTAC scaffold and prepared a selection of conformationally constrained, bumped, and kinked ARV-110 analogs (**Fig. 5b, Supplementary Fig. S6, Supplementary Schemes S6–S8**). We noted that amine **7-S** is an active CI inhibitor while its corresponding amide **8-S** is not, and that **7-S** is almost identical to ARV-110 with respect to the E3 ligase recruiter and linker (**Supplementary Fig. S7**). Reasoning that an analogous amine-to-amide conversion in ARV-110 might similarly decrease CI binding, we prepared the amide ARV-110 analog **27**. This substance showed slightly diminished CI activity relative to ARV-110 in both the CTG/2-DG (pEC_50_ = 6.24 ± 0.07; 0.58 µM) and extracellular flux assays (E60/B (9 µM) = 0.20) (**Fig. 5c, Extended Data Fig. 4**), while remaining an effective AR degrader (**Fig. 5d**); however, the CI inhibition-reducing effect from the amine-to-amide conversion was not as pronounced as in the rabeprazole-thalidomide series. Further attempts to introduce conformational restraint by introduction of a urea (**28**) or steric bulk via dimethyl-substitution (**29**) resulted in degradation (both effective) and CI inhibitory (pEC50 = 6.20 ± 0.05; 0.63 µM and 6.33 ± 0.08; 0.47 µM, respectively) profiles in the CTG/2-DG, as well as in the extracellular flux assays (E60/B (9 µM) = 0.24 and 0.17, respectively) that were similar to **27**.

Determined to improve on these results, we next investigated bumped ARV-110 analogs. To simplify late-stage introduction of bumps and avoid the introduction of a new stereocenter, we replaced the 1-(piperidin-4-ylmethyl)piperazine linker of ARV-110 with a di(piperidin-4-yl)amine linker, resulting in secondary amine **30** (**Fig. 5b**). This substance is a potent AR degrader (**Fig. 5d**) and CI inhibitor (pEC50 = 6.53 ± 0.01; 0.30 µM) (**Fig. 5c**), which encouraged us to synthesize bumped derivatives thereof. While *N-*methylation (**31**) did not provide a bump sufficient to abolish CI inhibition (pEC50 = 6.64 ± 0.11; 0.23 µM), acetylated (**32**; **DKFZ-1285**) and benzylated (**33**; **DKFZ-1295**) derivatives were inactive against CI (**Fig. 5c** and **5e**) and retained AR degradation activity (**Fig. 5d**). We confirmed these findings in the cell-free enzymatic assay, where **30** (pIC50 = 6.36; 0.44 µM) remained a nanomolar CI inhibitor, while **32** (**DKFZ-1285**) and **33** (**DKFZ-1295**) were inactive (**Fig. 5f**).

Encouraged by the bump approach, we next investigated the introduction of kinks into the ARV-110 scaffold. To do so, we shifted the linker attachment point of the E3 ligase recruiter thalidomide from the C-5 to the C-4 position. This redesign yielded **34**, which exhibited markedly reduced CI inhibition (**Fig. 5c**), but showed poor AR degradation despite incorporation of a commonly used C-4 exit vector (**Fig. 5d**)^37^. As an alternative strategy, we employed a distinct cereblon-recruiting moiety incorporating an intrinsic conformational kink. The resulting 1,3-substituted benzo[*d*]imidazol-2-one derivative, **35** (**DKFZ-1300**), displayed markedly reduced CI inhibition (**Fig. 5c** and **5e**) while maintaining AR degradation potency comparable to ARV-110 (**Fig. 5d**). A strong hook effect was apparent^2^, suggesting a difference in the cooperativity of the AR:PROTAC:cereblon ternary complex with this scaffold relative to ARV-110. In summary, our proof-of-concept study with ARV-110 shows that geometric perturbation of a potent CI inhibitory molecular scaffold by introducing bumps and kinks can mitigate off-target CI inhibition while retaining target activity.

## Discussion

Over the past two decades, chemically induced proximity has emerged as a highly promising pharmacological strategy in drug development. Among its various implementations, targeted protein degradation represents the most clinically advanced approach, with bifunctional degraders currently being evaluated in dozens of clinical trials, particularly for cancer therapy^2^. Despite this promise, the targeted protein degradation field faces multiple challenges: tumor resistance can arise through mutations in the recruited E3 ligase complex, altered ubiquitin-proteasome activity, or adaptive rewiring of downstream signaling pathways^2,4^. Residing in the “beyond-rule-of-5” space^38^, physicochemical properties such as high lipophilicity and poor aqueous solubility are challenges for pharmacokinetics optimization and, consequently, clinical application of PROTACs^10^. Finally, despite being considered highly target-specific, cereblon-recruiting PROTACs can result in off-target degradation of neosubstrates like the Ikaros zinc finger family^11,13^.

Our study reveals off-target inhibition of mitochondrial complex I as another potential hurdle in targeted protein degrader development. When incubating cells with rabeprazole-thalidomide hybrids, we noticed an unusually fast and pronounced exhaustion of the cellular ATP pool. Although we initially attributed the moderate degradation of IRF2BP2 and RNF156 to targeted protein degradation, this effect is more likely attributable to stalled protein synthesis resulting from ATP depletion^18^. Consistent with this hypothesis, IRF2BP2 and RNF156 have short half-lives and are naturally degraded within the 6 h incubation period of the experiment, whereas other target proteins (DENR, UBR7, and ZYX) remain unaffected (**Extended Data Fig. 1b**).

HEK293T cells treated with **10-S** and many of its analogs are unable to compensate for impaired mitochondrial respiration through glycolytic upregulation. Rather, glycolysis is concomitantly suppressed, leading to rapid ATP depletion. This is in contrast to most other cell lines (e.g., A375) that we tested. Differences in metabolic activity, glycolytic dependence, metabolic plasticity, or compensatory pathway activation likely contribute to this cell line-specific response^39^. However, HEK293T cells can upregulate glycolysis in response to the CI inhibitor rotenone, indicating that the effect is also compound-specific. Substances like **6-S, 7-S, 10-S**, and **13**, which affect both the OCR and ECAR in HEK293T cells (**Extended Data Fig. 4**), can therefore be considered as dual respiration and glycolysis inhibitors in HEK293T cells. Notably, simultaneous inhibition of mitochondrial respiration and glycolysis has been explored repeatedly to target tumor metabolism^40–43^, pointing to a potential application of such compounds. Interestingly, the three active truncated substances that lack a thioether (i.e., **14, 15**, and **26** (**DKFZ-1234**)) behave like rotenone and do not inhibit glycolysis in HEK293T cells (**Extended Data Fig. 4**). Compound **26** (**DKFZ-1234**) stands out from our study as a “pure” CI inhibitor in HEK293T cells with potency similar to the chemical probe BAY-179. Exploring the SAR of glycolysis inhibition (or the lack thereof) and uncovering the target causing this effect will be the focus of a future study.

We serendipitously discovered that compounds synthesized within a targeted degrader program act as CI inhibitors and subsequently identified additional bifunctional molecules exhibiting the same off-target behavior. These findings underscore the need to incorporate CI testing early in chemically induced proximity discovery efforts. Because extracellular flux analysis, high-resolution respirometry, Cell Painting, and cell-free CI inhibition assays are costly and not readily accessible in many laboratory settings, we developed a simple, widely accessible, and high-throughput-compatible CellTiter-Glo® assay that can be performed in 384-well format. In this assay, cells are co-treated with 2-DG, which rapidly blocks glycolysis, and a putative CI inhibitor, and ATP levels are quantified after two hours to enable rapid pEC_50_ determination. The Glu/Gal assay^44^, which compares cells cultured in high-glucose versus high-galactose media, is conceptually related but requires specialized media and is more time-intensive.

A systematic truncation strategy applied to **10-S** revealed that a long, linear molecular scaffold is required for effective CI inhibition, consistent with the structural features of many canonical CI inhibitors (**Supplementary Fig. S2**)^26,30,31^. We therefore hypothesize that these compounds inhibit CI by binding within the elongated (ca. 30 Å) and narrow ubiquinone-binding channel, thereby preventing substrate access. Consistent with this model, the T-shaped derivatives **18** and **25** exhibit little to no CI inhibition. Because only **25** was completely inactive in the CTG/2-DG assay (see **Fig. 3c**), we suggest that the thalidomide (Eastern) moiety of **10-S**, and potentially other PROTACs like ARV-110, enters the ubiquinone binding tunnel first. This would require future structural studies for confirmation. Nevertheless, the fact that many western truncations of **10-S** (see **Fig. 3**) do not abolish CI inhibition indicates that C-5-substituted thalidomide derivates may have an intrinsic affinity for complex I, that require only a simple hydrophobic residue beyond the linker (e.g., **14** or **15**). Notably, many molecular glue degraders based on the thalidomide scaffold have lengths and physicochemical properties similar to **14** and **15** and may, therefore, also have measurable CI-inhibitory profiles^12,45^.

To investigate whether other bifunctional molecules might inhibit CI, we assembled a collection of thirteen cereblon-recruiting PROTACs that are either well known (e.g., dBET1), have long, slender, hydrophobic structures (e.g., A20), have entered clinical evaluation (e.g., KT-474), or exhibit more than one of these criteria (e.g., ARV-110) (**Supplementary Fig. S4**). We identified ARV-110, ARV-766, and BMS-986365, three clinically investigated degraders, as potent CI inhibitors. Despite employing distinct protein-of-interest and E3 ligase recruiter scaffolds, all three inhibit complex I, indicating that CI off-target activity is not confined to a specific bifunctional chemotype. Focusing further on four additional AR PROTACs, we found that all three cereblon-recruiting degraders inhibit CI, indicating that CI inhibition appears to be a particular liability this PROTAC class. On the other hand, it raises the question whether this liability extends beyond AR-targeting degraders to the broader targeted protein degradation field^14,15,46^. As a counter example, our rabeprazole-thalidomide hybrids constitute a second, structurally distinct CI-inhibiting bifunctional scaffold that fulfills all major design criteria of contemporary degraders. While significantly more extensive screening of PROTAC libraries will be required to fully understand how often CI-inhibition arises in PROTAC scaffolds, this suggests that CI inhibition represents a broader structural liability rather than a target-specific phenomenon. Instead, long, slender, and hydrophobic molecular architectures, which are well represented by AR degrader scaffolds, appear predisposed to occupy the elongated ubiquinone-binding tunnel of CI. Consequently, this risk is likely relevant not only to degraders but to bifunctional proximity-inducing modalities more generally^47,48^.

In our study, ARV-110 emerged as the AR degrader with the highest potency in a cell-free CI activity assay (pEC50 = 7.66; 22 nM), a concentration that may fall within clinically relevant exposure ranges. Oral administration of ARV-110 at 5 mg·kg^-1^ in mice resulted in a maximal plasma concentration of 0.75 µM^49^. In clinical studies, 420 mg daily was established as the recommended phase II dose^50^, whereas 140 mg achieved plasma concentrations comparable to those associated with preclinical anti-tumor activity^51^. In preclinical models, ARV-110 exhibited a DC50 of 1 nM in LNCaP cells^52^. Taken together, these observations raise the possibility that off-target CI inhibition may have contributed to ARV-766 supplanting ARV-110 as the lead clinical candidate.

Mitotoxicity of an AR degrader scaffold developed at AstraZeneca was recently identified and mitigated through a tailored medicinal chemistry optimization process^15^. Our approach that strategically introduces steric bumps and conformational kinks goes beyond scaffold-specific optimization and offers a generalizable design principle for bifunctional molecules. This principle builds on established strategies in chemical biology. The addition of a bump to small-molecule inhibitors, as employed in the bump–hole strategy, is a well-established technique in chemical biology to confer selectivity for site-specifically mutated proteins^36^. The concept of introducing a molecular kink is reminiscent of the induced *E*/*Z* isomerization in photopharmacological switches such as azobenzenes, which confer light-dependent on/off selectivity^53^. In our study, the bumped PROTACs **32** (**DKFZ-1285**) and **33** (**DKFZ-1295**) showed the most diminished CI inhibition amongst all ARV-110 analogs, indicating that this approach may be particularly effective.

The need to add bumps and kinks can potentially be avoided by judicious selection of the two recruiting ligands as well as their exit vectors at the outset of a chemically induced proximity project. Thalidomide-based cereblon recruiters substituted at C-5 appear to be well suited for CI inhibition (e.g., ARV-110), whereas C-4 substitution (e.g., **34**) introduces a conformational kink that reduces CI binding. Unfortunately, C-5 substitution has been shown to abolish off-target neo-substrate activity that can arise for C-4 linked analogs^11^, creating a design trade-off between minimizing neo-substrate recruitment and preventing CI inhibition. Newer cereblon recruiters, such as the 1,3-substituted benzo[*d*]imidazol-2-one found in **35** (**DKFZ-1300**), incorporate an intrinsic conformational kink^54^, and may therefore offer promising starting points for synthesis. Novel modalities such as trivalent or macrocyclic degraders offer other possible strategies to avoid linearity^55^.

In summary, we have discovered a novel CI inhibitor scaffold and established structural principles and strategies to mitigate CI inhibition of bifunctional molecules including degraders and, more broadly, chemical inducers of proximity. We expect our study will spark new awareness for safer and more rational molecular design.

## Methods

**Chemical synthesis and characterization** can be found in the supporting information file.

### Cell culture

The human NSCLC cell lines (ATCC: A549, NCI-H23, NCI-H522, NCI-H661, NCI-H1573, NCI-H1693, NCI-H1793, NCI-H2023; DSMZ: LXF289, HCC827) were cultured in RPMI-1640, high- or low-glucose DMEM or DMEM/F12 media (Gibco, Sigma Aldrich). The human SCLC suspension cell line NCI-H82 (ATCC) and the NSCLC cell line NCI-H1944 (ATCC) were cultivated in RPMI-1640 (Gibco). The human cell line Beas-2B (Cytion) was grown in DMEM/F-12 (Gibco). HEK293T (ATCC), HaCaT (AddexBio), A375 (ATCC), and U2OS (DSMZ) cells were cultured in high-glucose DMEM (Sigma Aldrich). All the media contained 10% fetal bovine serum (FBS-12A, Capricorn Scientific), 100 U·mL^-1^ penicillin and 100 µg·mL^-1^ streptomycin (Gibco). Cells were maintained at 37 °C in a 5% CO2 atmosphere.

Prior to seeding for experiments, cells were harvested by centrifugation (4 min at 130 *g* at room temperature). Cells were detached by incubation with trypsin-EDTA (0.25%) for 5 min at 37 °C. All cell lines were regularly tested for *mycoplasma* contamination (Eurofins Genomics) and authenticated using Multiplex Cell Authentication by Multiplexion (Heidelberg) as described previously^56^.

### Reagents for biological evaluation

All reagents used in this study are listed in **Supplementary Table S2**.

### CellTiter-Glo® (CTG) assay

Cells were seeded in white 96-well plates at a density of 7,000 cells per well in 100 µL of culture medium except for H82, HaCaT, Beas-2b and A375 (10,000 cells per well) and HEK293T (30,000 cells per well). After 24 h, test compounds in 5-fold dilution series were added on top. Following incubation for a desired time period (1–24 h), viable cells were quantified using the CellTiter-Glo® (CTG) cell viability assay (Promega) according to the manufacturer’s instructions. CTG assays were performed in technical triplicate in three independent experiments. In each independent experiment, the luminescence values are normalized to untreated control (=100%). The heatmap values are the mean of the outcomes of the three independent experiments, which each were generated by taking the mean of the three technical replicate values. A pEC50 value was also generated for each independent experiment using a 4-parameter model in GraphPad Prism. Reported is the arithmetic mean ± SD of the three values. Compounds that don’t reach 50% inhibition at the highest concentration (50 µM) are considered inactive. In the washout experiment, cells were first incubated for 6 h with indicated compounds, followed by complete medium removal, rinsing with PBS, and addition of fresh medium without the compound. After 4 h of recovery, the CTG assay was performed.

### CTG/2-DG assay

A375 cells were seeded in a white 384-well plate at a density of 5,000 cells per well in 50 µL of culture medium containing 50 mM 2-deoxyglucose. This was quickly followed by compound injection using a D300e Digital Dispenser (Tecan) in seven 5-fold dilutions starting with 50 µM. Different DMSO volumes in the dilution series were normalized by injecting DMSO only. The CTG assay was performed 2 h after compound injections (5 µL of CTG reagent added per well). Each experiment was conducted in technical triplicate and repeated independently (*n* = 3).

### Real-time extracellular flux assay

The Seahorse XFe96 Analyzer (Agilent) was employed to analyze cell metabolism. A375 (13,000 cell per well) or HEK293T (30,000 cells per well) cells were seeded on XFe96 plates (Agilent) on the day prior to the assay. The plates for HEK293T cells were coated with poly-D-lysine (Gibco) according to the manufacturer’s instructions. The assay medium was prepared by supplementing the Seahorse DMEM medium (Agilent) with 2 mM glutamine (Gibco), 1 mM pyruvate (Sigma Aldrich), 10 mM glucose (Gibco) and 0.2% BSA (Merck). The culture medium was replaced with the assay medium approximately 1 h before starting the assay. The drug solutions for injections were diluted in the assay medium without BSA. All the compounds screened for inhibition of respiration were tested at three different concentrations (1, 3, and 9 µM) and injected first, followed by injection of oligomycin (1 µM), FCCP (1 µM), and a mixture of rotenone (1 µM), antimycin A (0.5 µM), and 2-deoxyglucose (100 mM) to completely inhibit both respiration and glycolysis, which reveals OCR and ECAR background, respectively.

For quantification, both OCR and ECAR are normalized to the basal value 20 min after start. The background is subtracted from B, E10, E60, and M. Finally, the ratios of E10/B, E60/B, and M/B are calculated to normalize for variability in the assay performance. The interplate variability manifests also with a random drift of the curve between E10 and E60 on some plates. To account for this, E10/B, E60/B, and M/B values calculated for tested compounds on each plate are normalized to the E10/B, E60/B, and M/B value of untreated control, respectively.

### High-resolution respirometry

Respiration was measured at 37 °C by the high-resolution respirometer Oxygraph-2k (Oroboros). Freshly harvested A375 cells were resuspended in an assay medium containing 80 mM KCl, 10 mM Tris-Cl, 3 mM MgCl2, 1 mM EDTA, and 5 mM potassium phosphate (pH 7.4). Three million cells were added to the 2 mL Oxygraph chamber and then permeabilized with digitonin (30 µg·mL^-1^). Substrates and inhibitors were injected into the chamber at the following concentrations: 10 mM pyruvate, 10 mM glutamate, 3 mM malate, 1 mM ADP, 1 μM rotenone, 10 mM succinate, 1 mM KCN. Suspected CI inhibitors were tested in the concentration range 0.1–3 µM. Oxygen consumption was expressed in pmol oxygen·s^-1^·cell^-1^.

### Complex I activity assay

Complex I activity was measured in bovine mitochondrial membranes^31^ by monitoring the rate of NADH:O2 oxidoreduction spectrophotometrically using a SpectraMax Plus 348 96-well plate reader (Molecular Devices). Membranes (10 µg·mL^-1^) were assayed in the presence of 200 µM NADH and 3 µM horse heart cytochrome c (Sigma Aldrich) in buffer containing 10 mM MOPS (pH 7.5) and 50 mM KCl. The assay was performed at 32 °C and initiated by addition of NADH. NADH absorbance was monitored for 20 min at 340–380 nm, and rates were determined by linear regression of the maximal slope (ε = 4.81 mM^−1^·cm^−1^). Activities were calculated as µmol NADH oxidized ·min^-1^·mg^-1^ and normalized to a DMSO control (100%). Compounds were tested from DMSO stock solutions over a concentration range of 0.0001–100 µM. Measurements were performed in technical quadruplicates using an initial 10-fold serial dilution, immediately followed by finer titrations to define the Hill slope for compounds exhibiting CI inhibition. Piericidin A was included as a positive inhibitory control. Dose-response curves were analyzed using GraphPad Prism with a four-parameter model to determine IC_50_ values.

### Targeted metabolomic analysis of nucleotide intermediates

To analyze nucleotide intermediates, HEK293T and A375 cells were quickly rinsed with ice-cold saline (0.9% NaCl) and scraped off into 1.2 mL of ice-cold saline. A small volume of each cell suspension (0.1–0.2 mL) was kept on ice and used for cell counting with the Luna-FL Automated Cell Counter (Logos). 1 mL of cell suspension was transferred into a microtube and spun down at 1,000 *g* for 1 min at 4 °C. After removal of supernatant, cell pellets were snap-frozen in liquid nitrogen and stored at –80 °C until further processing.

Frozen pellets containing approximately 3–8 million cells were processed using a modified energy carrier extraction protocol adapted from Rost et al.^57^, expanded to support polarity switching and the detection of additional metabolites. Samples were resuspended in 500 µL of ice-cold acetonitrile:methanol:water (3:1:1, *v*/*v*/*v*) containing 15 mM ammonium acetate (pH 9), vortexed briefly, and subjected to three freeze–thaw cycles (5 min in liquid nitrogen, followed by 10 min on dry ice). Extracts were centrifuged at 20,817 *g* for 10 min at –9 °C (Eppendorf 5417R), and supernatants were transferred to low-recovery LC-MS vials.

Chromatographic separation and detection were performed on an ACQUITY I-Class PLUS UPLC system (Waters) coupled to a QTRAP 6500+ triple quadrupole mass spectrometer (AB SCIEX) equipped with an electrospray ionization source. The ion source settings were as follow: curtain gas: 30 psi; collision gas: medium; ion spray: 5500 V/ −4500 V; with source temperature 400 °C. Metabolites were resolved on an ACQUITY Premier BEH Amide Vanguard Fit column (100 mm × 2.1 mm, 1.7 µm; Waters) maintained at 35 °C. NAD/NADH, NADP/NADPH, and related energy metabolites were separated using a gradient elution program (**Supplementary Table S3**) with mobile phase A (50:50 acetonitrile:water) and mobile phase B (90:10 acetonitrile:water), both containing 5 mM ammonium acetate and 0.05% ammonium hydroxide (pH 10).

Quantification was conducted in multiple-reaction monitoring (MRM) mode. MRM transitions, retention times, and instrument parameters are listed in **Supplementary Table S4**. Data were acquired using Analyst 1.7.2 and processed using OS 3.4.5.828 (AB SCIEX).

### SDS-PAGE and Western blotting

HEK293T were seeded in 6-well plates (one million cells per well) and A375 in 12-well plates (150,000 cells per well) on the day prior to treatment (6 h, compound concentrations as indicated in each figure legend). For total protein extraction, cells were harvested using Pierce RIPA buffer (Thermo Scientific, 89900) supplemented with a protease inhibitor cocktail (Serva) and 100 U·mL^-1^ benzonase (Merck). Protein concentrations were determined by Pierce BCA Protein Assay (Thermo Scientific) using BSA as a standard.

Cell lysates were denatured by incubation with Laemmli sample buffer (5.1% glycerol, 0.51% SDS, 0.051% bromophenol blue, 30 mM DTT, 10.6 mM Tris-HCl, pH 6.9) for 5 min at 95 °C. Proteins were separated with Tris-glycine SDS-PAGE, transferred onto a nitrocellulose membrane, and detected with primary antibodies diluted according to manufacturer’s instructions overnight at 4 °C, followed by incubation with secondary antibodies conjugated to fluorophores for 1 h at room temperature. Fluorescence was recorded with the Chemidoc Imager (Bio-Rad). The primary antibodies used in this study are: androgen receptor antibody (Cell Signaling 5153), ß-tubulin antibody (Sigma Aldrich T0198), IRF2BP2 antibody (Proteintech 18847-1-AP), RNF156 antibody (Proteintech 11285-1-AP), UBR7 antibody (Proteintech 16903-1-AP), DENR antibody (Proteintech 10656-1-AP), Zyxin antibody (Proteintech 10330-1-AP), GAPDH antibody (Santa Cruz sc-365062), c-MYC antibody (Cell Signaling 13987). As secondary antibodies, goat anti-rabbit and anti-mouse antibodies conjugated to either StarBright Blue 520 or StarBright Blue 700 were used (Bio-Rad).

### Cell painting assay

The Cell painting assay was performed as previously reported^33^ and adheres closely to the method described by Bray et al.^58^ See Supplementary Information for experimental details.

### Microscopy

HEK293T cells were seeded in a transparent flat-bottom 96-well plate (40,000 per well) on the day prior to the imaging. Immediately before imaging, cells were treated with 10 µM compounds for 6 h in triplicates. Brightfield images were recorded by the automated cell imager NYONE (Synentec) using the 10x objective with the resolution of 0.9 µm per pixel.

### Flow cytometry (cell death)

To assess cell death, the Annexin V-FITC Kit (Miltenyi Biotec) was used according to the manufacturer’s instructions in combination with flow cytometry analysis. After 6 h treatment, HEK293T cells were detached by pipetting up and down, washed with the binding buffer and stained for 15 min with annexin V-FITC at room temperature in the dark. Afterwards, cells were washed with PBS, resuspended in the binding buffer and briefly stained with propidium iodide immediately prior to analysis by the flow cytometer Guava easyCyte BGV HT (Cytek). Annexin V-FITC was detected in the Green-B channel (excitation 488 nm, emission 525/30 nm) and propidium iodide in the Red-G channel (excitation 532 nm, emission 695/50 nm). The FlowJo software was used to gate for single cells and calculate the percentage of apoptotic/necrotic cells.

### Aqueous solubility

The aqueous solubility was determined at WuXi AppTec Co., Ltd. (Shanghai) as follows: stock solutions of test and control compounds (10 μL) were added to individual wells of a 96-well plate, followed by 490 μL of pH 7.4 phosphate buffer (PB). Samples were mixed by vortexing (≥2 min) and incubated on an orbital shaker at 800 rpm at room temperature for 24 h. The plates were centrifuged at 25 °C for 10 min (approximately 4,000 rpm). Supernatants were filtered using MultiScreen® Solvinert PTFE filter plates (Millipore–Merck) and collected into a fresh 96-well plate by centrifugation (≥5 min). Analyte concentrations in the filtrates were quantified by LC-UV or LC-MS/MS using an ACQUITY UPLC BEH C18 (1.7 µm, 2.1 × 50 mm) or an XBridge® C18 column (3.5 µm, 4.6 × 100 mm) column with the following methods: Solvent A = H2O with 0.1% formic acid, solvent B = MeCN with 0.1% formic acid, or solvent A = H2O with 0.1% trifluoroacetic acid, solvent B = MeCN with 0.1% trifluoroacetic acid.

### Distribution coefficient (logD7.4)

The distribution coefficient (logD7.4) was determined at WuXi AppTec Co., Ltd. (Shanghai) as follows: Test and control compounds were aliquoted in duplicates (2 µL each) into microcentrifuge tubes, and PB-saturated 1-octanol (149 µL) and 1-octanol saturated PB (149 µL) were added to each tube to form biphasic systems. Samples were vigorously mixed for 2 min and then shaken at 800 rpm for 1 h at room temperature to allow phase equilibration. Samples were then centrifuged at 4000 rpm for 5 min to achieve phase separation. Aliquots from both the aqueous and organic layers were collected, diluted as appropriate, and analyzed by LC-MS/MS without the use of external calibration curves. Analytical LC/MS was performed on a Shimadzu LC-30AD system equipped with a SCIEX QTRAP 6500+ mass spectrometer using an ACQUITY UPLC BEH C18 column (1.7 µm, 2.1 × 50 mm) and the following method: Solvent A = H2O with 0.1% formic acid, solvent B = MeCN with 0.1% formic acid. The logD7.4 value was calculated according to the following equation:

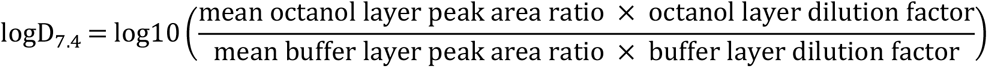

### Software and statistical analysis

Chemical structures were drawn using ChemDraw Professional 22.2.0 (Revvity Signals Software, Inc.). All figures were generated using Affinity Studio (Canva). Statistical analyses were performed in GraphPad Prism 10 software or in RStudio (Posit). Data is presented as mean ± SD in the bar graphs and dot plots. Inter-group differences were analyzed by one-way ANOVA, followed by two-tailed unpaired *t*-tests with Holm–Šídák correction for multiple comparisons (GraphPad Prism 10). The heatmaps were generated in RStudio using the ComplexHeatmap package^59^ and display the mean values of all the respective replicates. DataWarrior 6.1.0 (openmolecules.org) was used to generate UMAP plots.

## Acknowledgements

We thank the following colleagues for kindly providing us with a sample of their degraders: D. Kang (Nanjing University of Chinese Medicine) for **A20**^60^, G. Wang (Xavier University of Louisiana) for compound **9c**^61^, and Z. Chen (East China University of Science and Technology, Shanghai) for **7i**^62^. We thank the Metabolomics Core Technology Platform of the Heidelberg University supported by the CellNetworks Core Technology Platform CCTP for metabolite analyses.

## Author contributions

N.R., H.N., N.G., and A.K.M. designed the research and analyzed the data; N.R., with input from A.K.M., designed, synthesized, and characterized chemical compounds; H.N., with help from J.S., F.D, and N.d.V., and input from N.G., planned, performed, and analyzed most biological experiments; B.S.I., with input from J.H., designed, performed, and analyzed the complex I activity assay; M.S., with input from G.P., designed, performed, and analyzed the metabolomic profiling; S.S. supervised and planned the Cell painting assay and S.Z. analyzed the Cell painting profiles; N.R., H.N., N.G., and A.K.M. wrote the paper, with input from all other authors.

## Competing interests

The authors declare no competing interests.

**Extended Data Fig. 1.**
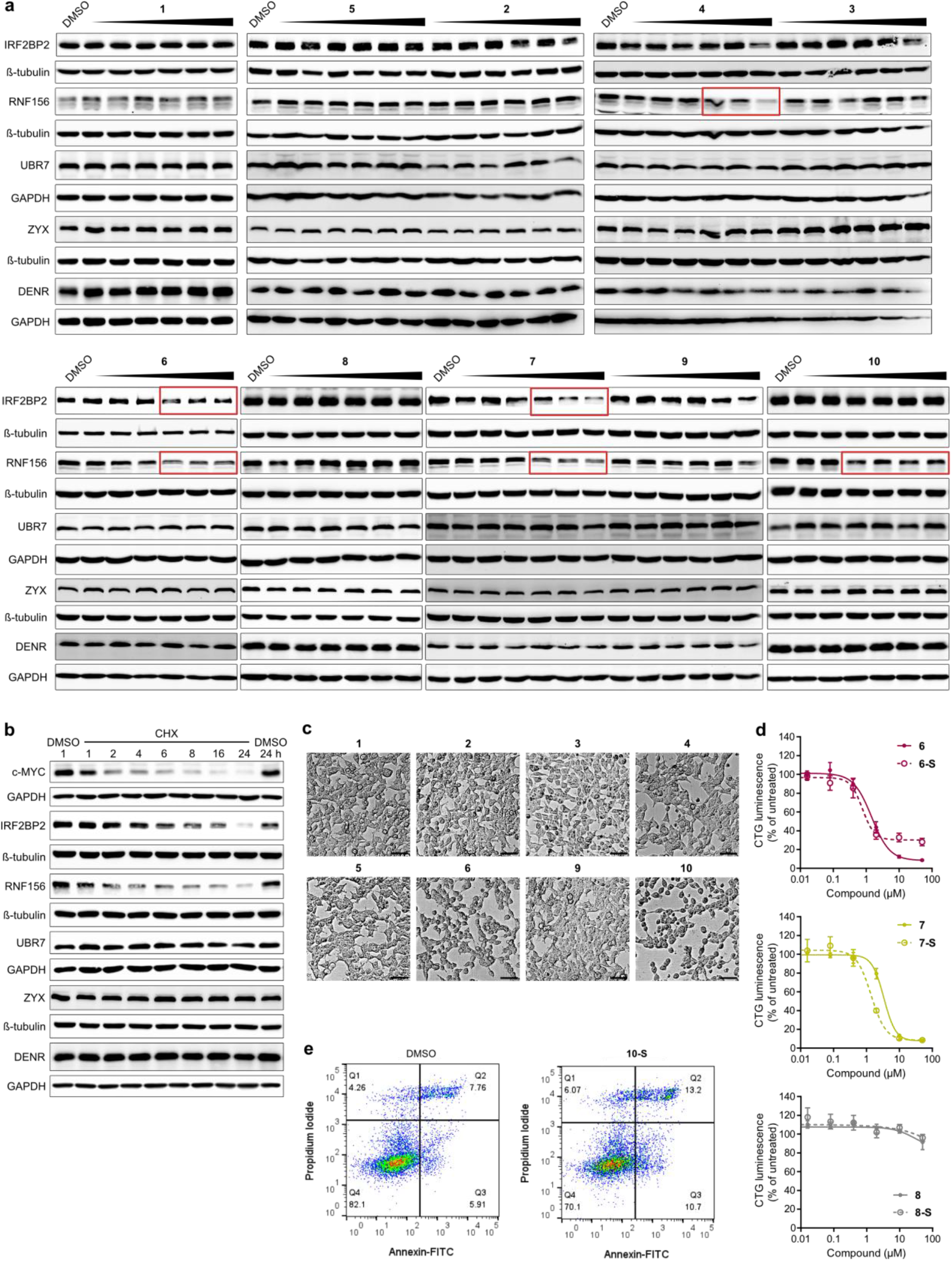
Rabeprazole-thalidomide hybrids induce stress in HEK293T cells. **a**, Representative Western blots (*n* = 1–2) of rabeprazole-target proteins (IRF2BP2, RNF156, ZYX, UBR7, and DENR) in HEK293T cells treated with rabeprazole-thalidomide hybrids for 6 h. **8** and **10** were tested from 50 µM in 3-fold dilutions; all other substances started at 50 µM with 5-fold dilutions. Red boxes highlight lower protein levels. **b**, Western blot analysis (*n* = 1) of HEK293T cells treated with 90 µM cycloheximide (CHX) to assess the turnover of proteins tested with c-MYC as a short half-life control protein. **c**, Bright-field images of HEK293T cells treated with 10 µM **1**–**6, 9**, and **10** for 6 h. Scale bar: 90 µm. **d**, Representative CellTiter-Glo assay (*n* ≥ 3, *m* = 3) of **10, 10-S**, and **10-Me** in HEK293T cells after 6 h. Luminescence values were normalized to untreated control. Data points represent means ± SD. **e**, An example of the gating for flow cytometry analysis of cell death in HEK293T cells (**Fig. 1f**) treated with 10 µM **10-S** for 6 h. Cells were stained with annexin V-FITC and propidium iodide (PI) and categorized as follows: viable = annexin-negative, PI-negative; early apoptotic = annexin-positive, PI-negative; late apoptotic/necrotic = annexin-positive, PI-positive; necrotic = annexin-negative, PI-positive. **f**, Representative CTG/2-DG assay (*n* = 3, *m* = 3) of **6, 6-S, 7, 7-S, 8**, and **8-S**. Luminescence values were normalized to untreated control. Data points represent mean ± SD. *n* = biological replicates, *m* = technical replicates.

**Extended Data Fig. 2.**
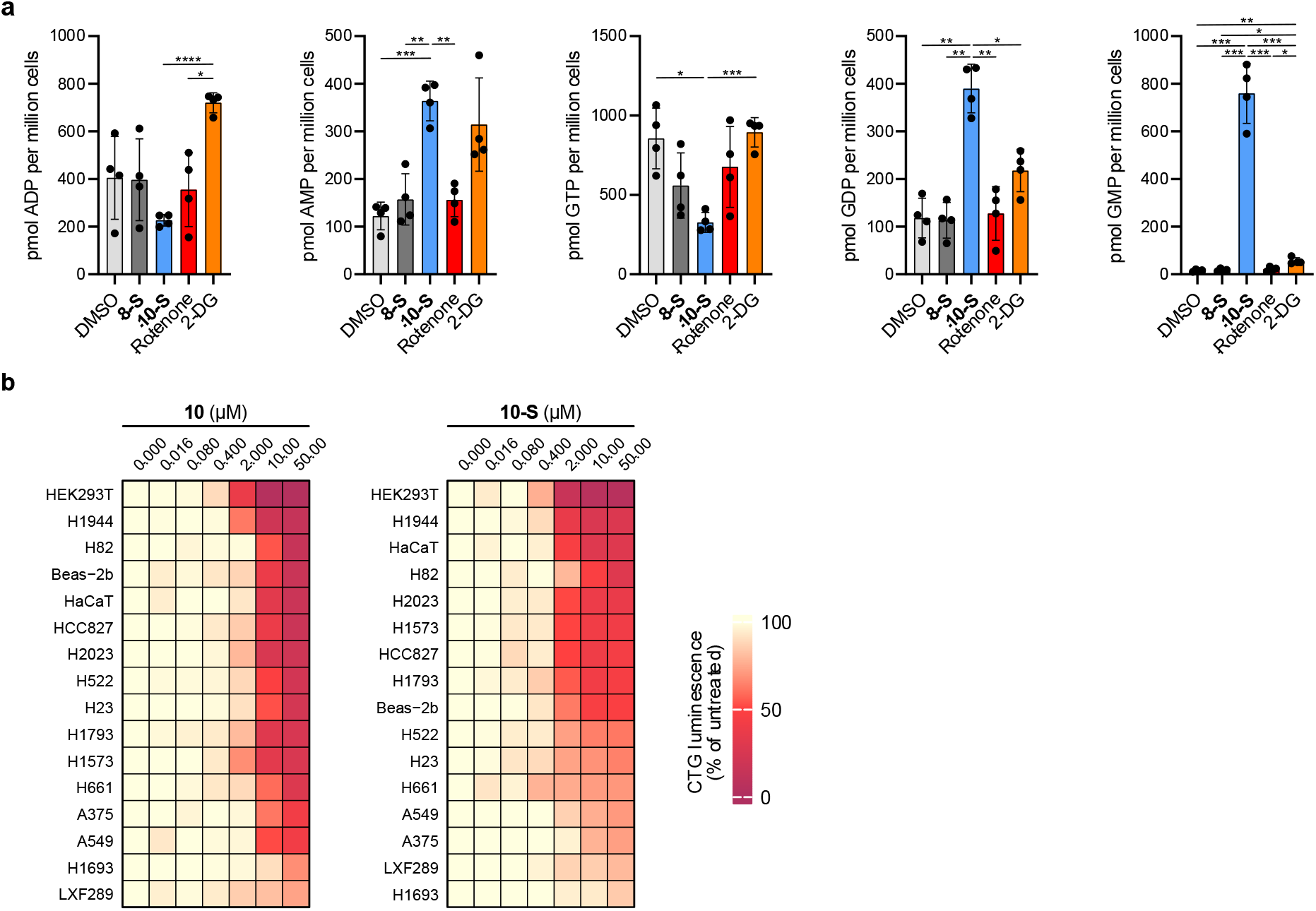
Rabeprazole-thalidomide hybrids induce rapid loss of ATP in different cell lines. **a**, Targeted metabolic quantification of cellular energy carriers in HEK293T cells treated with 10 µM **8-S**, 10 µM **10-S**, 10 µM rotenone or 50 mM 2-deoxyglucose (2-DG) for 6 h. Data are presented as mean ± SD (*n* = 4). Statistical significance was determined by one-way ANOVA (ADP *p* = 0.0006, AMP *p* = 0.0001, GTP *p* = 0.0018, GDP *p* < 0.0001, GMP *p* < 0.0001), followed by two-tailed unpaired *t*-tests with an adjustment for multiple comparisons using the Holm– Šídák method to assess differences between conditions (\**p* < 0.05, \*\**p* < 0.01, \*\*\**p* < 0.001, \*\*\*\**p* < 0.0001). **b**, Dose– response heatmaps of **10** and **10-S** in 16 different cell lines after 6 h incubation. Luminescence values were normalized to untreated control. Heatmap colors are the mean (*n* ≥ 2; *m* = 3). Cell lines are ranked based on maximum effect at 50 µM. *n* = biological replicates, *m* = technical replicates.

**Extended Data Fig. 3.**
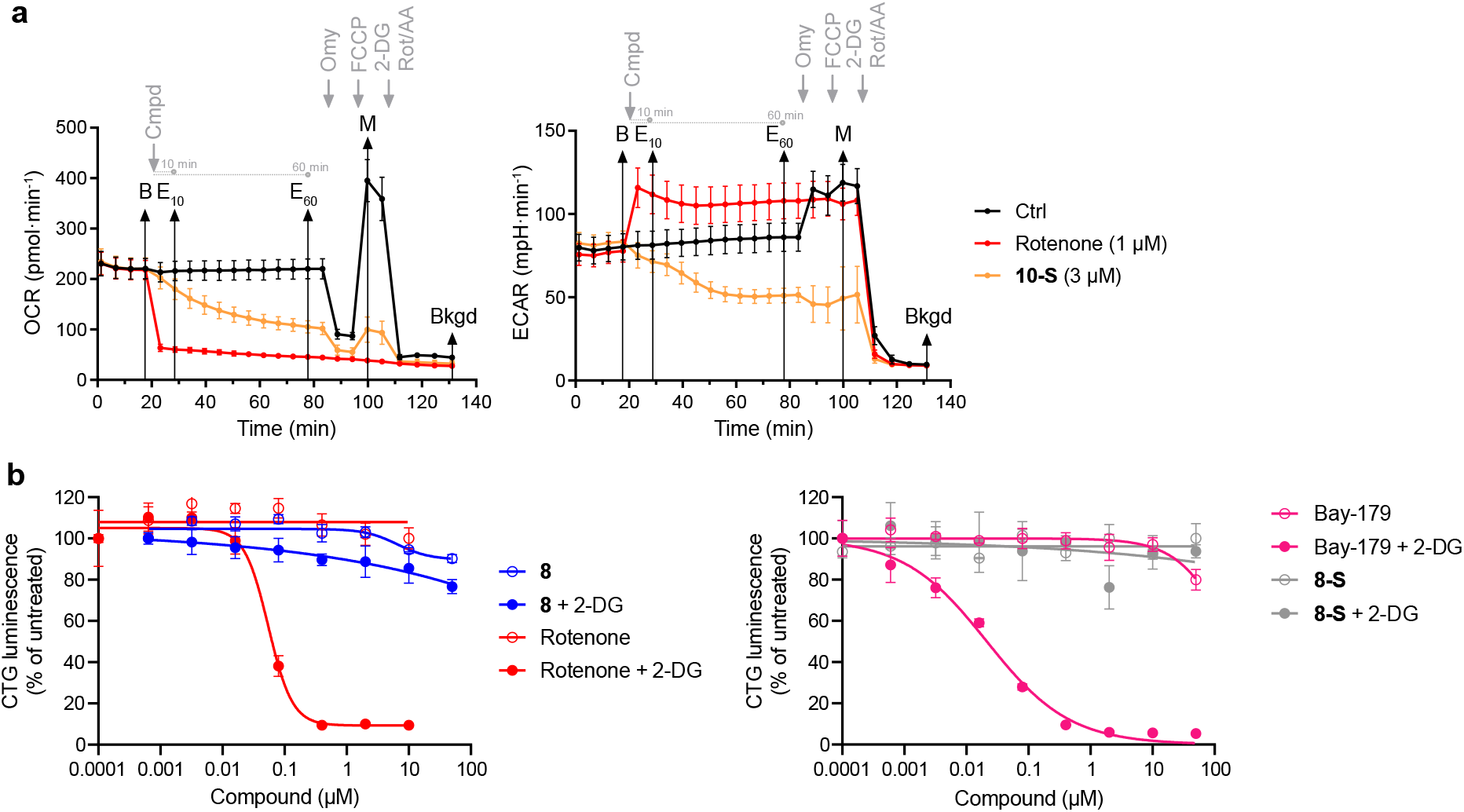
Extracellular flux quantification and CTG/2-DG assay validation. **a**, Defined values from the extracellular flux assay are depicted on a representative experiment in HEK293T. For both oxygen consumption rate (OCR) and extracellular acidification rate (ECAR), the basal rate (B) is defined as OCR or ECAR immediately before injection of the tested compound. E_10_ and E_60_ are defined as OCR or ECAR 10 min or 60 min after injection, respectively. M (maximum) is defined as the maximum OCR or ECAR that is reached after FCCP injection. Bkgd (background) is defined as OCR or ECAR at the end of the experiment. From these values, the ratios E_10_/B, E_60_/B, and M/B can be calculated (see methods section for detailed description of the data normalization process). **b**, Representative dose– response experiments (*n* = 1 for each experiment) comparing the standard CTG and CTG/2-DG assay. Luminescence values were normalized to untreated control and data points represent mean ± SD (*m* = 3). *n* = biological replicates, *m* = technical replicates.

**Extended Data Fig. 4.**
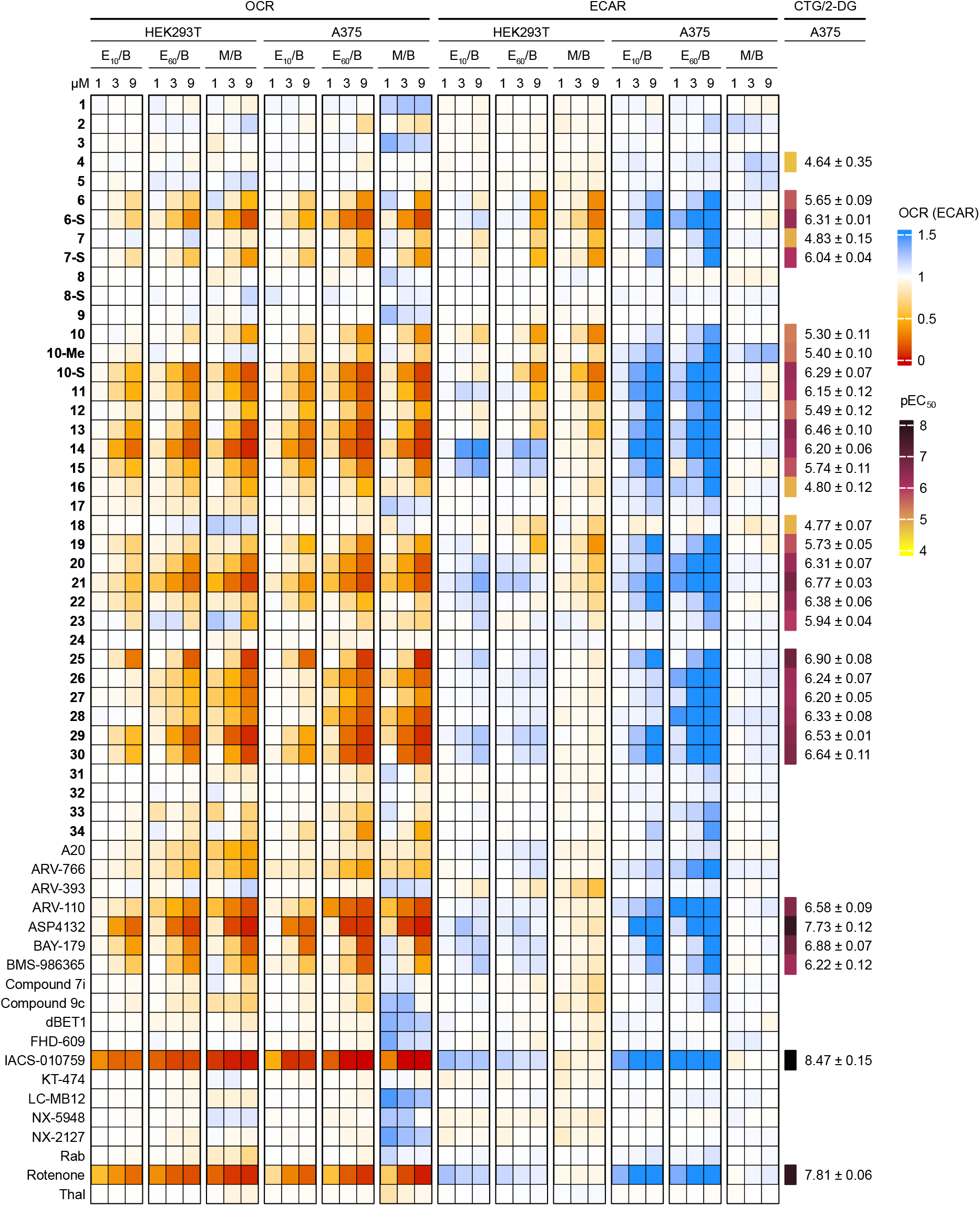
Extracellular flux data heatmap and CTG/2-DG screening data for every compound tested. Oxygen consumption rate (OCR) and extracellular acidification rate (ECAR) were recorded in HEK293T and A375 cells, while the CTG/2-DG assay was performed in A375 cells only. The E_10_/B, E_60_/B and M/B values were calculated as described in **Extended Data Fig. 3a**. The heatmap color represent mean values (*n* = 1–3; *m* = 6). The pEC_50_ values represent mean ± SD (*n* = 3, *m* = 3), the compounds with a missing pEC_50_ value were inactive in the CTG/2-DG assay.

**Extended Data Fig. 5.**
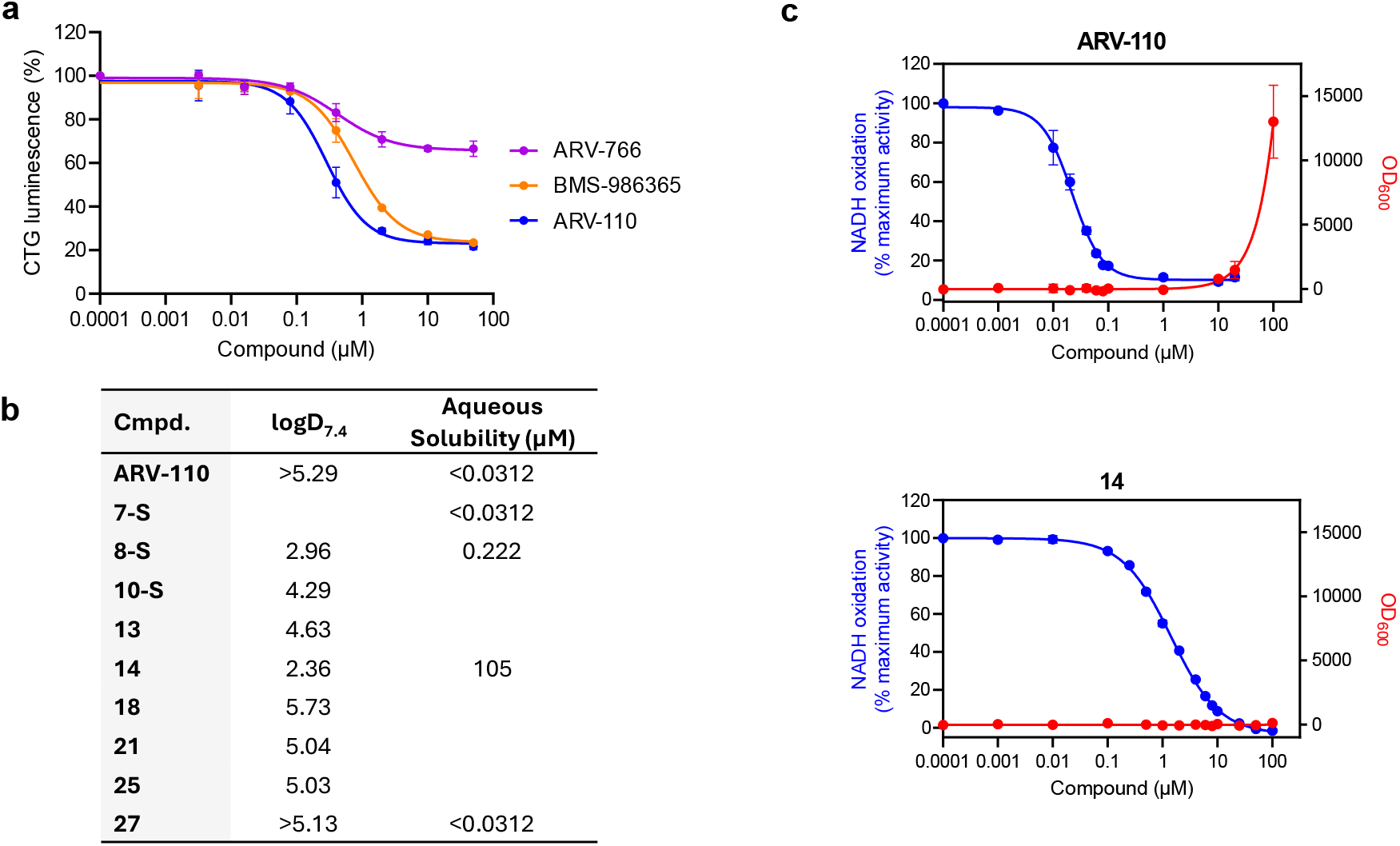
Representative CTG assay and physicochemical data. **a**, A representative CTG/2-DG assay for the AR degraders ARV-110, ARV-766, and BMS-986365 (*n* = 3; *m* = 3). Luminescence values were normalized to untreated control and data points represent mean ± SD. **b**, Table showing logD_7.4_ (distribution coefficient at pH 7.4, octanol/buffer) and aqueous solubility (µM). Data is presented as mean (*n* = 2). **c**, Blue graphs: dose–response curves of CI activity (*n* = 1; *m* = 4) measured as NADH:O_2_ oxidoreduction in isolated bovine mitochondrial membranes. Activities were normalized to the DMSO control (=100%). Data points represent mean ± SD. Red graphs: solubility of ARV-110 and **14** (*n* = 1; *m* = 4) was assessed by measuring light scattering at 600 nm (OD_600_). *n* = biological replicates, *m* = technical replicates.

**Extended Data Fig. 6.**
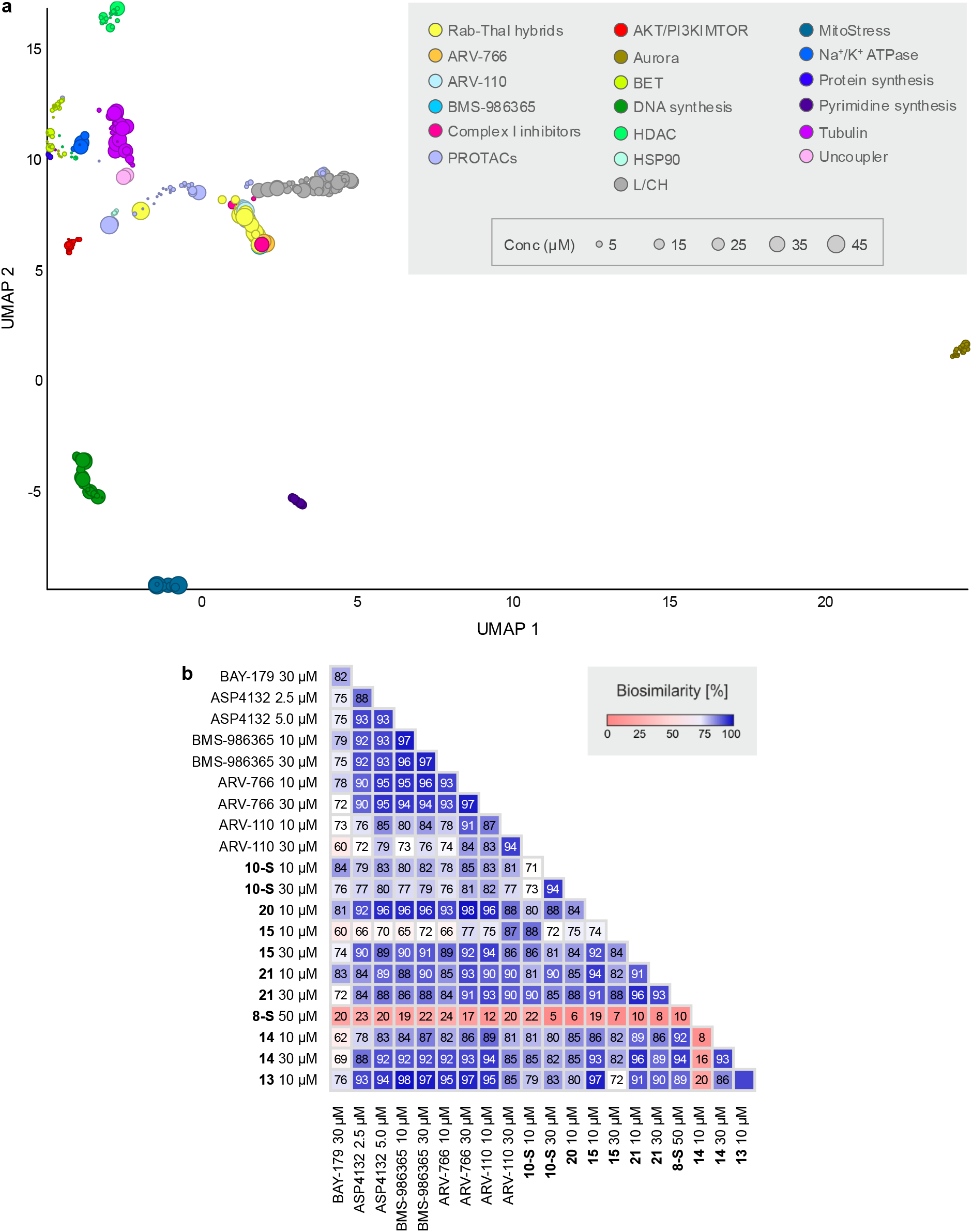
Analysis of the compound collection for bioactivity in the Cell painting assay. **a**, Entire UMAP plot. The size of each symbol corresponds to the concentration. Not normalized, 10 neighbors. L/CH: lysosomotropism/cholesterol homeostasis. **b**, Profile cross-correlation analysis of biosimilarities found in the Cell painting assay for the studied compounds.

## References

1. Gerry, C. J. & Schreiber, S. L. Unifying principles of bifunctional, proximity-inducing small molecules. Nat. Chem. Biol. 16, 369–378 (2020).

2. Hinterndorfer, M., Spiteri, V. A., Ciulli, A. & Winter, G. E. Targeted protein degradation for cancer therapy. Nat. Rev. Cancer 25, 493–516 (2025).

3. Nalawansha, D. A. & Crews, C. M. PROTACs: An Emerging Therapeutic Modality in Precision Medicine. Cell Chem. Biol. 27, 998–1014 (2020).

4. Tsai, J. M., Nowak, R. P., Ebert, B. L. & Fischer, E. S. Targeted protein degradation: from mechanisms to clinic. Nat. Rev. Mol. Cell Biol. 25, 740–757 (2024).

5. Dikic, I. et al. Opportunities in proximity modulation: Bridging academia and industry. Mol. Cell 85, 3012–3022 (2025).

6. Schade, M. et al. Structural and Physicochemical Features of Oral PROTACs. J. Med. Chem. 67, 13106–13116 (2024).

7. Marker, T. et al. Site-specific activation of the proton pump inhibitor rabeprazole by tetrathiolate zinc centres. Nat. Chem. Biol. 17, 507–517 (2025).

8. Shin, J. M., Cho, Y. M. & Sachs, G. Chemistry of covalent inhibition of the gastric (H+, K+)-ATPase by proton pump inhibitors. J. Am. Chem. Soc. 126, 7800–7811 (2004).

9. Lindberg, P. & Carlsson, E. in Analogue-based Drug Discovery Ch. 5, 81–113 (Wiley, 2006).

10. Bekes, M., Langley, D. R. & Crews, C. M. PROTAC targeted protein degraders: the past is prologue. Nat. Rev. Drug Discov. 21, 181–200 (2022).

11. Nguyen, T. M. et al. Proteolysis-targeting chimeras with reduced off-targets. Nat. Chem. 16, 218–228 (2024).

12. Petzold, G. et al. Mining the CRBN target space redefines rules for molecular glue-induced neosubstrate recognition. Science 389, eadt6736 (2025).

13. Garside, H. et al. Safety considerations for cereblon-recruiting targeted protein degraders. Nat. Rev. Drug Discov. (2026) Online: (10.1038/s41573-026-01426-2).

14. Trapotsi, M. A. et al. Cell Morphological Profiling Enables High-Throughput Screening for PROteolysis TArgeting Chimera (PROTAC) Phenotypic Signature. ACS Chem. Biol. 17, 1733–1744 (2022).

15. Scott, J. S. et al. Discovery of AZD9750, an Orally Bioavailable Androgen Receptor Degrader for the Treatment of Prostate Cancer. J. Med. Chem. 69, 3209–3232 (2026).

16. Paresi, C. J., Liu, Q. & Li, Y. M. Benzimidazole covalent probes and the gastric H(+)/K(+)-ATPase as a model system for protein labeling in a copper-free setting. Mol. Biosyst. 12, 1772–1780 (2016).

17. Winter, G. E. et al. Phthalimide conjugation as a strategy for in vivo target protein degradation. Science 348, 1376–1381 (2015).

18. Vetma, V. et al. Confounding Factors in Targeted Degradation of Short-Lived Proteins. ACS Chem. Biol. 19, 1484–1494 (2024).

19. Riss, T. L. et al. in Assay Guidance Manual Ch. Cell Viability Assays, 403–428 (Eli Lilly & Company and the National Center for Advancing Tanslational Sciences, 2004).

20. Olbe, L., Carlsson, E. & Lindberg, P. A proton-pump inhibitor expedition: the case histories of omeprazole and esomeprazole. Nat. Rev. Drug Discov. 2, 132–139 (2003).

21. Nolfi-Donegan, D., Braganza, A. & Shiva, S. Mitochondrial electron transport chain: Oxidative phosphorylation, oxidant production, and methods of measurement. Redox Biol. 37, 101674 (2020).

22. Caines, J. K., Barnes, D. A. & Berry, M. D. in Cancer Cell Biology: Methods and Protocols Ch. 17, 225–234 (Springer US, 2022).

23. Lee, M. & Yoon, J. H. Metabolic interplay between glycolysis and mitochondrial oxidation: The reverse Warburg effect and its therapeutic implication. World J. Biol. Chem. 6, 148–161 (2015).

24. Molina, J. R. et al. An inhibitor of oxidative phosphorylation exploits cancer vulnerability. Nat. Med. 24, 1036–1046 (2018).

25. Kuramoto, K. et al. Development of a potent and orally active activator of adenosine monophosphate-activated protein kinase (AMPK), ASP4132, as a clinical candidate for the treatment of human cancer. Bioorg. Med. Chem. 28, 115307 (2020).

26. Mowat, J. et al. Identification of the Highly Active, Species Cross-Reactive Complex I Inhibitor BAY-179. ACS Med. Chem. Lett. 13, 348–357 (2022).

27. Srinivasan, B. & Lloyd, M. D. Quantitation and Error Measurements in Dose–Response Curves. J. Med. Chem. 68, 2052–2056 (2025).

28. Pesta, D. & Gnaiger, E. in Mitochondrial Bioenergetics: Methods and Protocols Ch. 3, 25–58 (Humana Press, 2012).

29. Toriki, E. S. et al. Rational Chemical Design of Molecular Glue Degraders. ACS Cent. Sci. 9, 915–926 (2023).

30. Chung, I. et al. Cork-in-bottle mechanism of inhibitor binding to mammalian complex I. Sci. Adv. 7 (2021).

31. Bridges, H. R. et al. Structural basis of mammalian respiratory complex I inhibition by medicinal biguanides. Science 379, 351–357 (2023).

32. Gustafsdottir, S. M. et al. Multiplex cytological profiling assay to measure diverse cellular states. PLoS One 8, e80999 (2013).

33. Pahl, A. et al. Morphological subprofile analysis for bioactivity annotation of small molecules. Cell Chem. Biol. 30, 839–853.e837 (2023).

34. Rezaei Adariani, S. et al. Detection of a Mitochondrial Fragmentation and Integrated Stress Response Using the Cell Painting Assay. J. Med. Chem. 67, 13252–13270 (2024).

35. Schölermann, B. et al. Identification of Dihydroorotate Dehydrogenase Inhibitors Using the Cell Painting Assay. ChemBioChem 23, e202200475 (2022).

36. Islam, K. The Bump-and-Hole Tactic: Expanding the Scope of Chemical Genetics. Cell Chem. Biol. 25, 1171–1184 (2018).

37. Han, X. et al. Strategies toward Discovery of Potent and Orally Bioavailable Proteolysis Targeting Chimera Degraders of Androgen Receptor for the Treatment of Prostate Cancer. J. Med. Chem. 64, 12831–12854 (2021).

38. Price, E. et al. Beyond Rule of Five and PROTACs in Modern Drug Discovery: Polarity Reducers, Chameleonicity, and the Evolving Physicochemical Landscape. J. Med. Chem. 67, 5683–5698 (2024).

39. Zhang, J. et al. Measuring energy metabolism in cultured cells, including human pluripotent stem cells and differentiated cells. Nat. Protoc. 7, 1068–1085 (2012).

40. Noble, R. A. et al. Simultaneous targeting of glycolysis and oxidative phosphorylation as a therapeutic strategy to treat diffuse large B-cell lymphoma. Br. J. Cancer 127, 937–947 (2022).

41. Sheng, X. et al. Dual inhibition of oxidative phosphorylation and glycolysis to enhance cancer therapy. Bioorg. Chem. 147, 107325 (2024).

42. Wang, K. et al. Restraining Cancer Cells by Dual Metabolic Inhibition with a Mitochondrion-Targeted Platinum(II) Complex. Angew. Chem. Int. Ed. 58, 4638–4643 (2019).

43. Xu, Y. et al. Tumor chemical suffocation therapy by dual respiratory inhibitions. Chem. Sci. 12, 7763–7769 (2021).

44. Rana, P., Aleo, M. D., Gosink, M. & Will, Y. Evaluation of in Vitro Mitochondrial Toxicity Assays and Physicochemical Properties for Prediction of Organ Toxicity Using 228 Pharmaceutical Drugs. Chem. Res. Toxicol. 32, 156–167 (2019).

45. Won, H.-J. et al. Synthetic and Medicinal Chemistry Perspectives on Cereblon-Binding Molecular Glue Degraders: Recent Advances and Emerging Trends. European Journal of Organic Chemistry n/a, e202501208.

46. Ruffilli, C. Optimising PROTACs against integral membrane proteins PhD thesis, University of Amsterdam, (2025).

47. Gourisankar, S. et al. Rewiring cancer drivers to activate apoptosis. Nature 620, 417–425 (2023).

48. Sarott, R. C. et al. Relocalizing transcriptional kinases to activate apoptosis. Science 386, eadl5361 (2024).

49. Nguyen, T. T., Kim, J. W., Choi, H. I., Maeng, H. J. & Koo, T. S. Development of an LC-MS/MS Method for ARV-110, a PROTAC Molecule, and Applications to Pharmacokinetic Studies. Molecules 27 (2022).

50. Gao, X. et al. Phase 1/2 study of ARV-110, an androgen receptor (AR) PROTAC degrader, in metastatic castration-resistant prostate cancer (mCRPC). J. Clin. Oncol. 40, 17–17 (2022).

51. Petrylak, D. P. et al. First-in-human phase I study of ARV-110, an androgen receptor (AR) PROTAC degrader in patients (pts) with metastatic castrate-resistant prostate cancer (mCRPC) following enzalutamide (ENZ) and/or abiraterone (ABI). J. Clin. Oncol. 38, 3500–3500 (2020).

52. Snyder, L. B. et al. Preclinical Evaluation of Bavdegalutamide (ARV-110), a Novel PROteolysis TArgeting Chimera Androgen Receptor Degrader. Mol. Cancer Ther. 24, 511–522 (2025).

53. Broichhagen, J., Frank, J. A. & Trauner, D. A Roadmap to Success in Photopharmacology. Acc. Chem. Res. 48, 1947–1960 (2015).

54. Lejava, A. et al. Development of a Buchwald–Hartwig Amination for an Accelerated Library Synthesis of Cereblon Binders. ACS Med. Chem. Lett. 16, 89–95 (2025).

55. Cao, Y., Harris, A. L. & Ciulli, A. Branching beyond bifunctional linkers: synthesis of macrocyclic and trivalent PROTACs. Nat. Protoc. (2025).

56. Castro, F. et al. High-throughput SNP-based authentication of human cell lines. Int. J. Cancer 132, 308–314 (2013).

57. Rost, L. M., Shafaei, A., Fuchino, K. & Bruheim, P. Zwitterionic HILIC tandem mass spectrometry with isotope dilution for rapid, sensitive and robust quantification of pyridine nucleotides in biological extracts. J. Chromatogr. B Analyt. Technol. Biomed. Life Sci. 1144, 122078 (2020).

58. Bray, M.-A. et al. Cell Painting, a high-content image-based assay for morphological profiling using multiplexed fluorescent dyes. Nat. Protoc. 11, 1757–1774 (2016).

59. Gu, Z., Eils, R. & Schlesner, M. Complex heatmaps reveal patterns and correlations in multidimensional genomic data. Bioinformatics 32, 2847–2849 (2016).

60. Wang, J. et al. Discovery of a Novel Orally Bioavailable FLT3-PROTAC Degrader for Efficient Treatment of Acute Myeloid Leukemia and Overcoming Resistance of FLT3 Inhibitors. J. Med. Chem. 67, 7197–7223 (2024).

61. Peng, X. et al. Discovery of Oral Degraders of the ROS1 Fusion Protein with Potent Activity against Secondary Resistance Mutations. J. Med. Chem. 67, 18098–18123 (2024).

62. He, H. et al. Development of Degraders of Cyclin-Dependent Kinases 4 and 6 Based on Rational Drug Design. J. Med. Chem. 67, 11354–11364 (2024).

