## Supplementary Information for "Disrupting linearity of PROTAC scaffolds prevents off-target complex I inhibition"

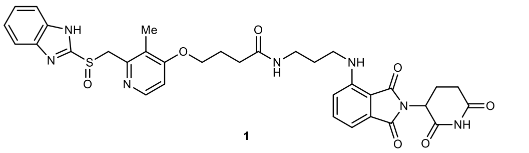

**1**

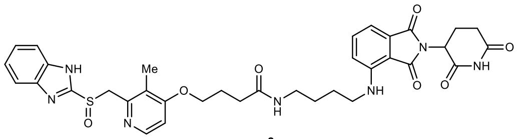

**2**

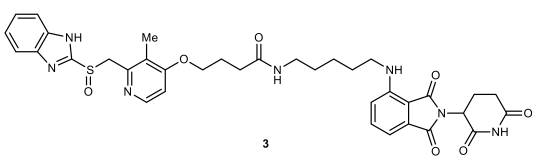

**3**

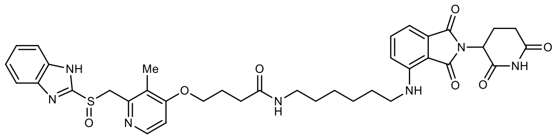

**4**

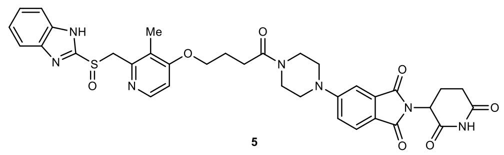

**5**

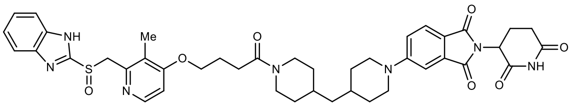

**6**

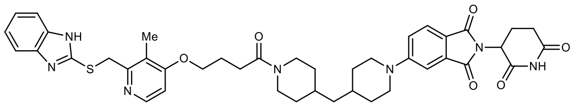

**6-S**

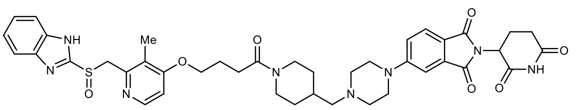

**7**

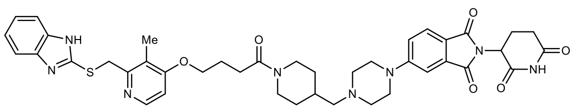

**7-S**

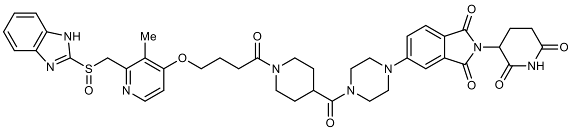

**8**

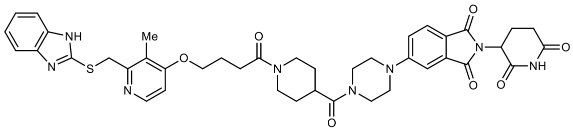

**8-S**

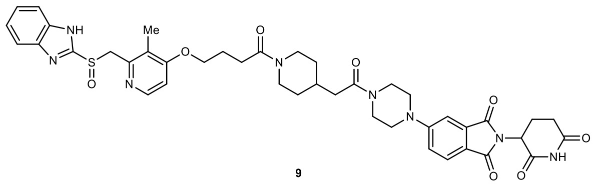

**9**

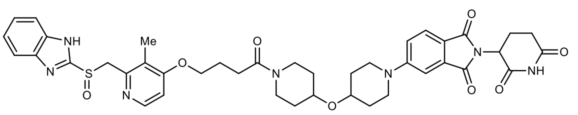

**10**

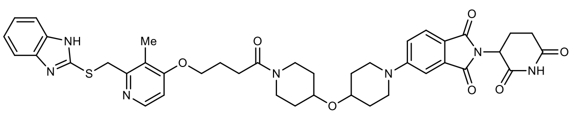

**10-S**

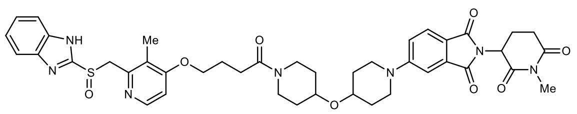

**10-Me**

**Supplementary Fig. S1**. Full structures of rabeprazole-thalidomide hybrids from **Fig. 1**.

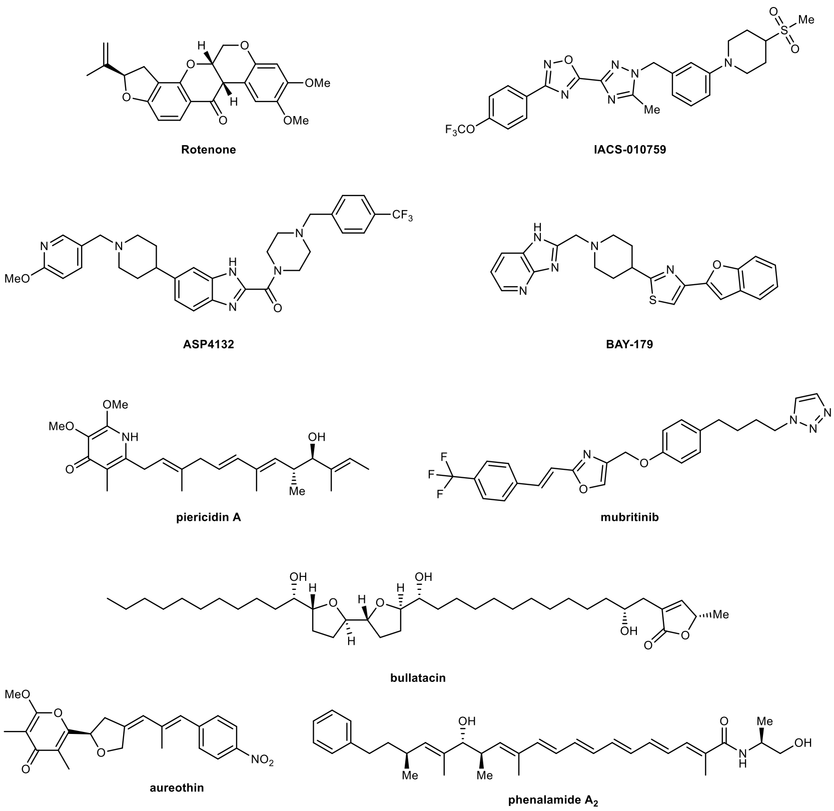

**Supplementary Fig. S2**. Structures of selected known inhibitors of mitochondrial C1.

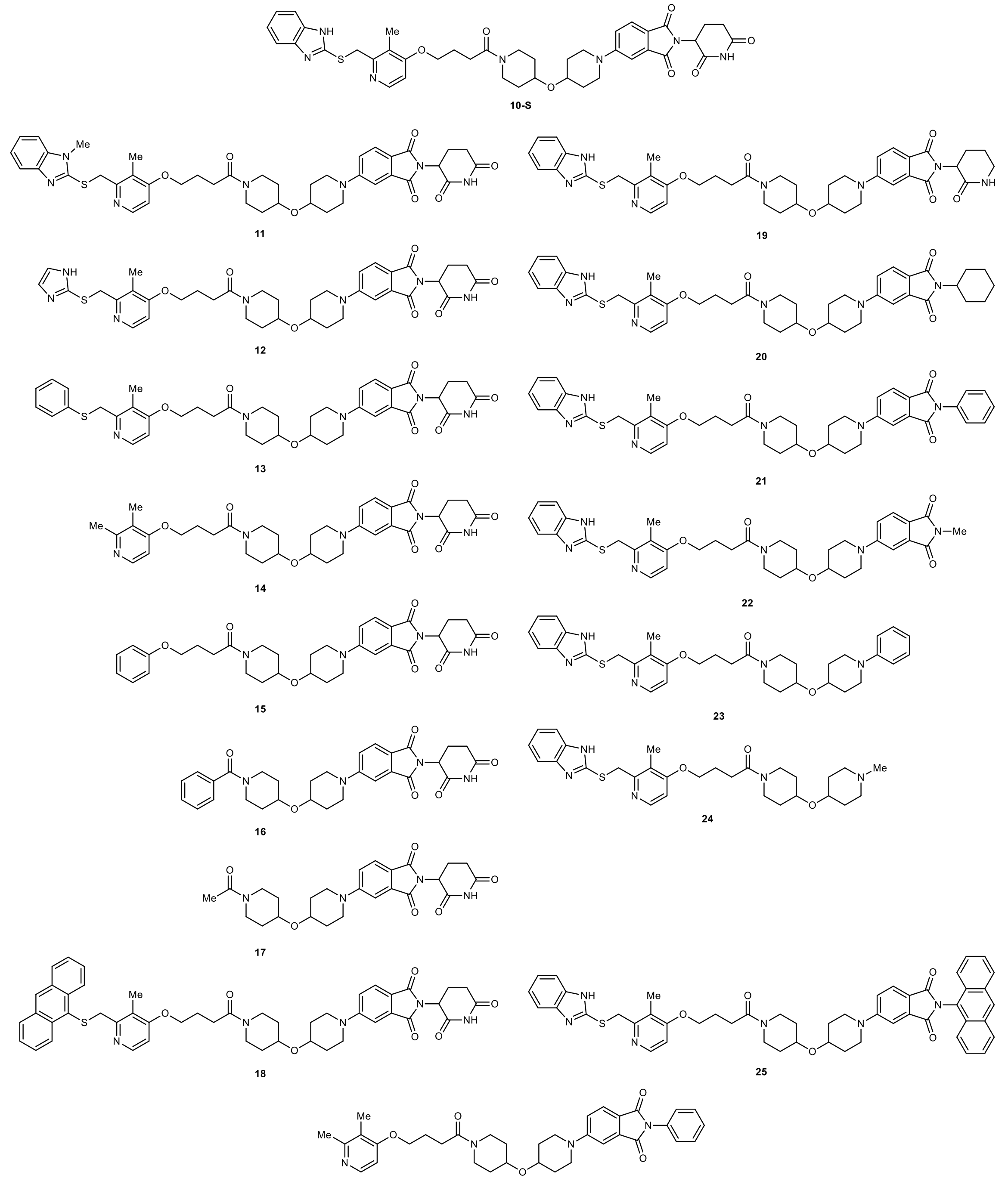

**Supplementary Fig. S3**. Structures of all truncations and modifications of **10-S** shown in **Fig. 3**.

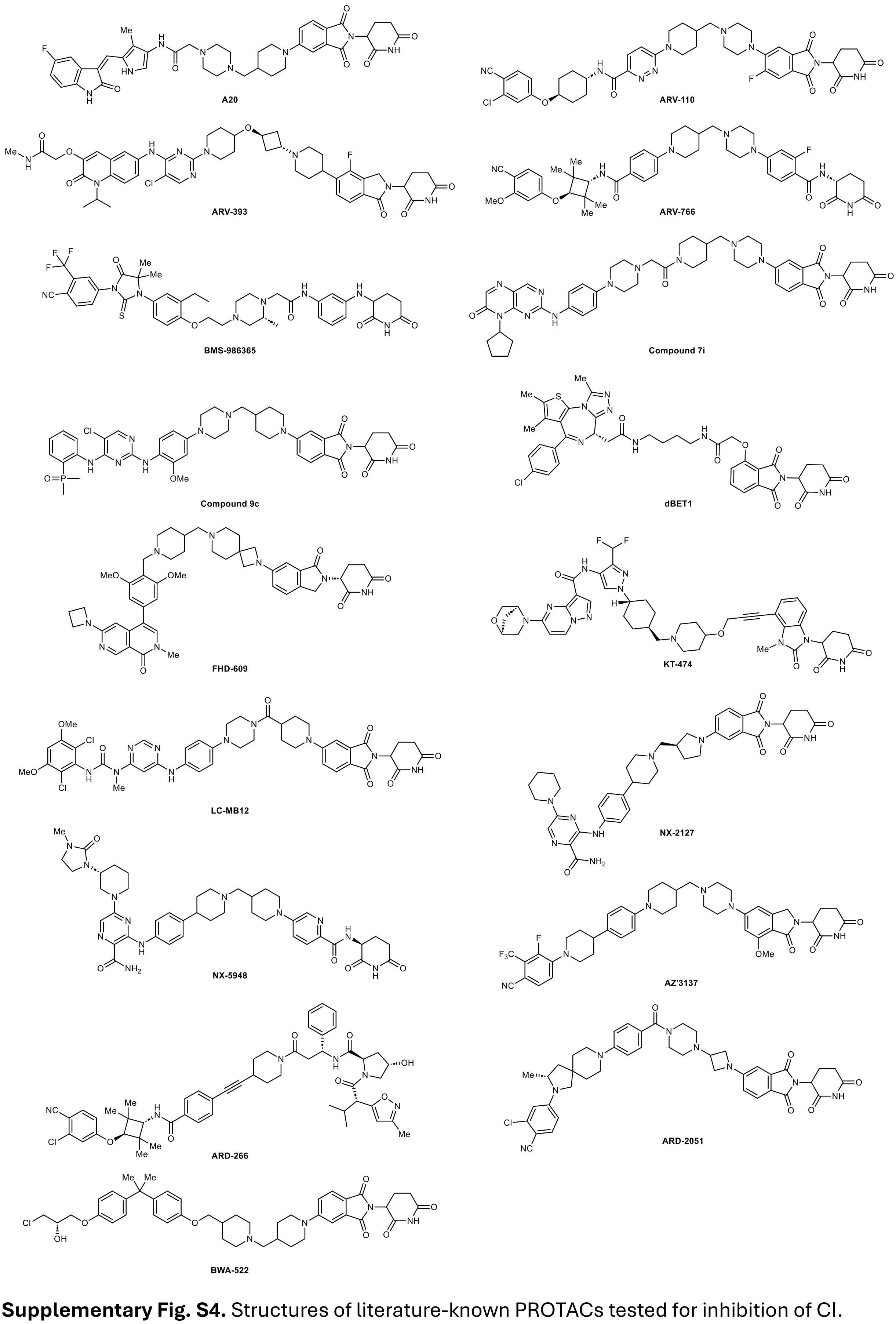

**Supplementary Fig. S4.** Structures of literature-known PROTACs tested for inhibition of CI.

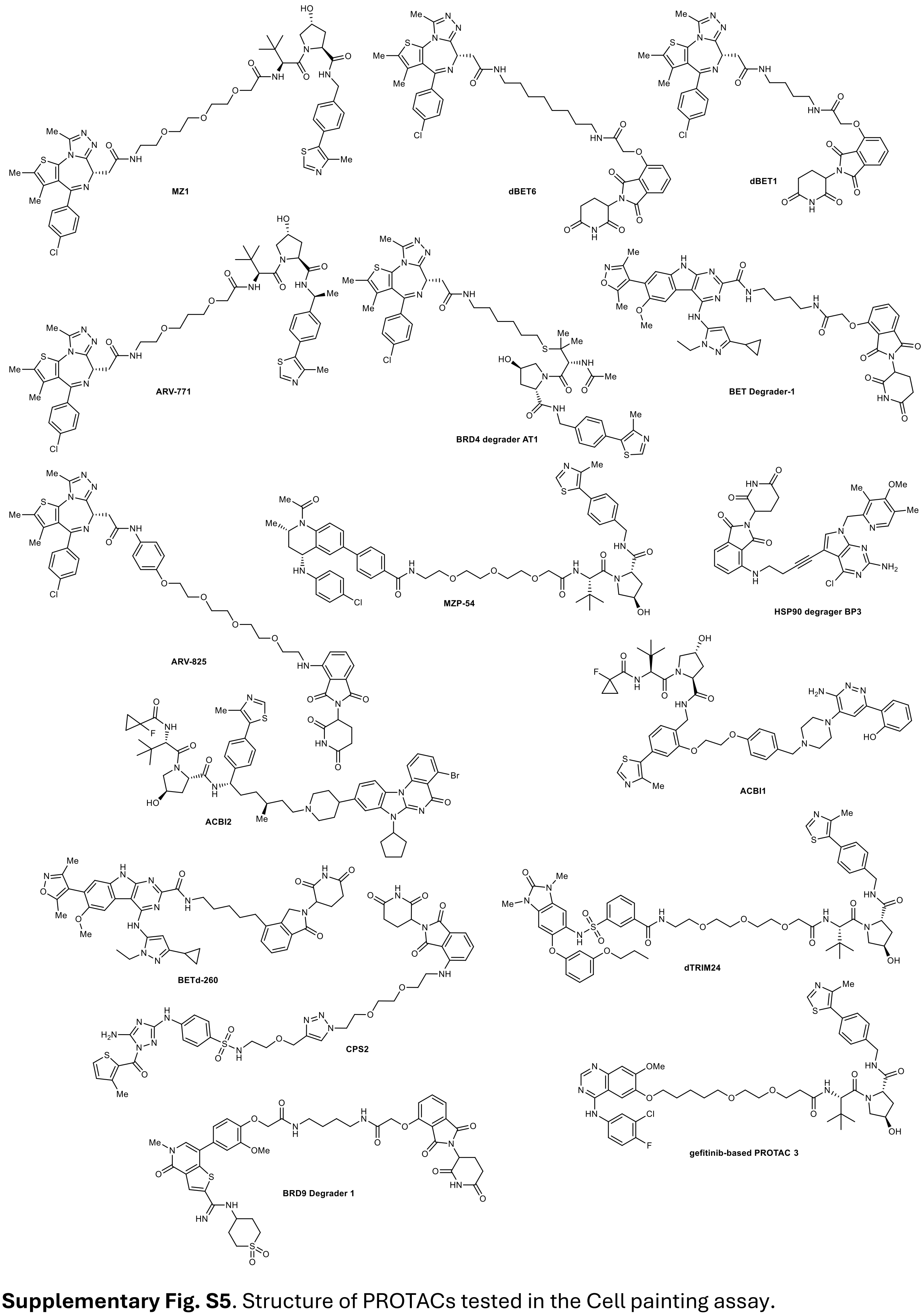

**Supplementary Fig. S5**. Structure of PROTACs tested in the Cell painting assay.

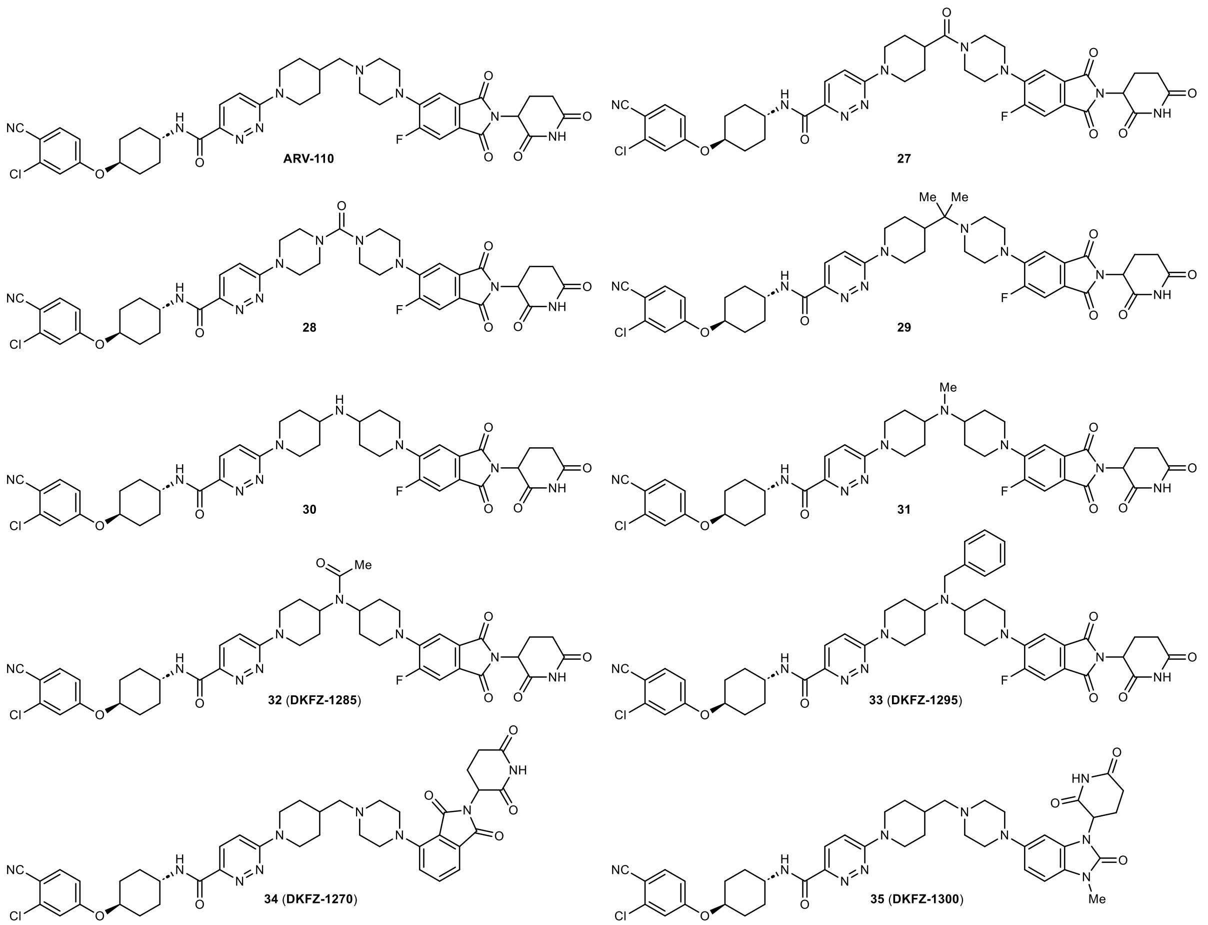

**Supplementary Fig. S6**. Structures of all ARV-110 analogs shown in **Fig. 5**.

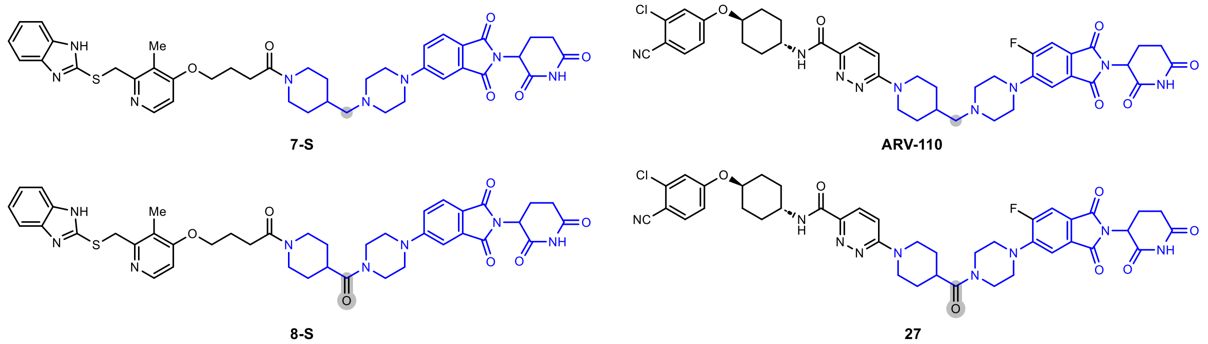

**Supplementary Fig. S7**. Structures of amine **7-S** and ARV-110 with the corresponding amides **8-S** and **26**.

**Supplementary Table S1**. Reagents for biological evaluation.

| Compound | Supplier | Catalog No. |
|---|---|---|
| 2-deoxyglucose | Thermo Scientific | 111980250 |
| ADP | Sigma-Aldrich | A5285 |
| Antimycin A | Sigma-Aldrich | A8674 |
| ARD-266 | BLDpharm | BD01448400 |
| ARD-2051 | BLDpharm | BD02208686 |
| ARV-110 | Selleck Chemicals | S6965 |
| ARV-393 | BLDpharm | BD02609354 |
| ARV-766 | BLDpharm | BD01911597 |
| ASP4132 | MedChemExpress | HY-136447 |
| AZ‘3137 | BLDpharm | BD02645304 |
| BAY-179 | MedChemExpress | HY-145707 |
| BMS-986365 | BLDpharm | BD02599175 |
| BWA-522 | BLDpharm | BD02312005 |
| Carbonyl cyanide 4-(trifluoromethoxy)phenylhydrazone (FCCP) | Sigma-Aldrich | C2920 |
| Cycloheximid (CHX) | Santa Cruz Biotechnology | SC-3508B |
| dBET1 | MedChemExpress | HY-101838 |
| Digitonin | Sigma-Aldrich | D141 |
| DL-Malic acid | Fluka | 02310 |
| FHD-609 | MedChemExpress | HY-153367 |
| IACS-010759 | MedChemExpress | HY-112037A |
| KCN | Sigma-Aldrich | 31252 |
| KT-474 | AmBeed | A1671259 |
| LC-MB12 | AmBeed | A2273159 |
| NX-2127 | AmBeed | A2020200 |
| NX-5948 | AmBeed | A2020219 |
| Oligomycin | Sigma-Aldrich | O4876 |
| Rabeprazole | Sigma-Aldrich | PHR1525 |
| Rotenone | Sigma-Aldrich | R8875 |
| Sodium glutamate | Thermo Scientific | J63424.09 |
| Sodium pyruvate | Fluka | 15990 |
| Sodium succinate | Sigma-Aldrich | 14170 |
| Thalidomide | Sigma-Aldrich | T144 |

**Supplementary Table S2**. LC gradient program used for separation of nucleotide and energy metabolites.

| Time (min) | Flow (mL·min$^{-1}$) | A (%) | B (%) |
|---|---|---|---|
| 0.00 | 0.400 | 5 | 95 |
| 0.50 | 0.400 | 5 | 95 |
| 0.51 | 0.350 | 5 | 95 |
| 4.00 | 0.350 | 60 | 40 |
| 6.00 | 0.300 | 77 | 23 |
| 9.00 | 0.300 | 95 | 5 |
| 11.90 | 0.300 | 100 | 0 |
| 12.00 | 0.350 | 5 | 95 |
| 15.00 | 0.400 | 5 | 95 |

**Supplementary Table S3.** Multiple-reaction monitoring (MRM) transitions, retention times, and mass spectrometry parameters for targeted metabolites.

| Compound ID | RT (min) | Q1 | Q3 | DP | EP | CE | CXP |
|---|---|---|---|---|---|---|---|
| ADP | 5.25 | 426.000 | 79.000 | -90.000 | -10.000 | -75.000 | -12.000 |
| ADP | 5.25 | 426.000 | 134.064 | -90.000 | -10.000 | -30.000 | -15.000 |
| AMP | 4.77 | 346.000 | 78.900 | -80.000 | -10.000 | -74.000 | -11.000 |
| AMP | 4.77 | 346.000 | 134.039 | -80.000 | -10.000 | -37.000 | -15.000 |
| ATP | 5.82 | 506.000 | 158.900 | -170.000 | -10.000 | -36.000 | -19.000 |
| ATP | 5.82 | 506.000 | 408.027 | -170.000 | -10.000 | -29.000 | -35.000 |
| GDP | 5.70 | 442.100 | 150.100 | -80.000 | -10.000 | -33.000 | -9.000 |
| GDP | 5.70 | 442.100 | 344.065 | -80.000 | -10.000 | -25.000 | -34.000 |
| GMP | 5.19 | 362.000 | 78.900 | -70.000 | -10.000 | -75.000 | -10.000 |
| GMP | 5.19 | 362.000 | 211.028 | -70.000 | -10.000 | -24.000 | -17.000 |
| GTP | 6.40 | 522.000 | 159.000 | -130.000 | -10.000 | -42.000 | -8.000 |
| GTP | 6.40 | 522.000 | 424.044 | -130.000 | -10.000 | -29.000 | -17.000 |

MRMs in positive and negative mode. For quantification the first transition was used, while the second is used as qualifier transition. ADP = Adenosine diphosphate; AMP = Adenosine monophosphate; ATP = Adenosine triphosphate; GDP = Guanosine diphosphate; GMP = Guanosine monophosphate; GTP = Guanosine triphosphate.

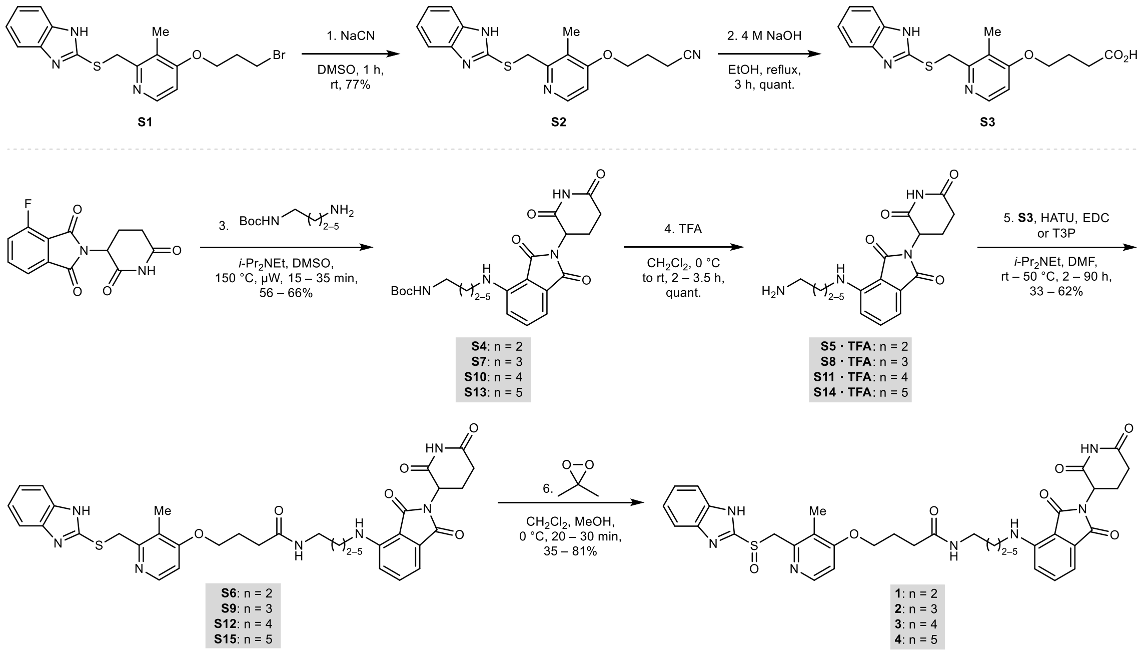

**Supplementary Scheme S1.** Synthesis route for rabeprazole-thalidomide hybrids **1–4**.

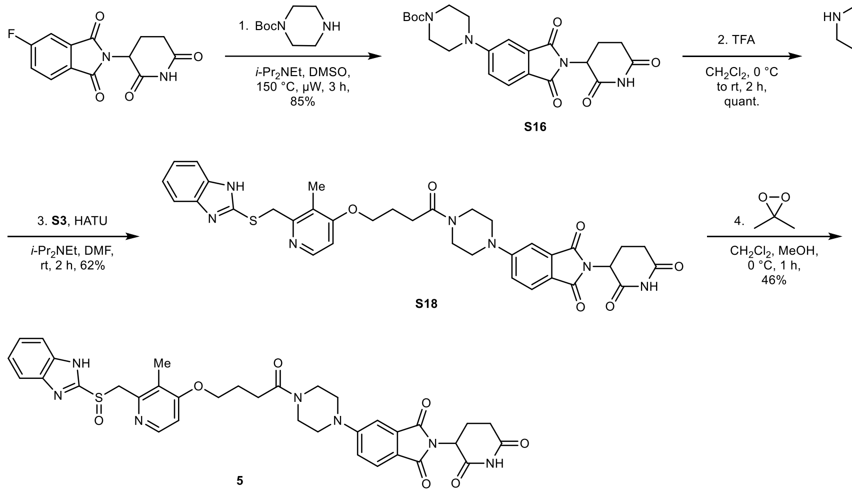

**Supplementary Scheme S2.** Synthesis route for rabeprazole-thalidomide hybrid **5**.

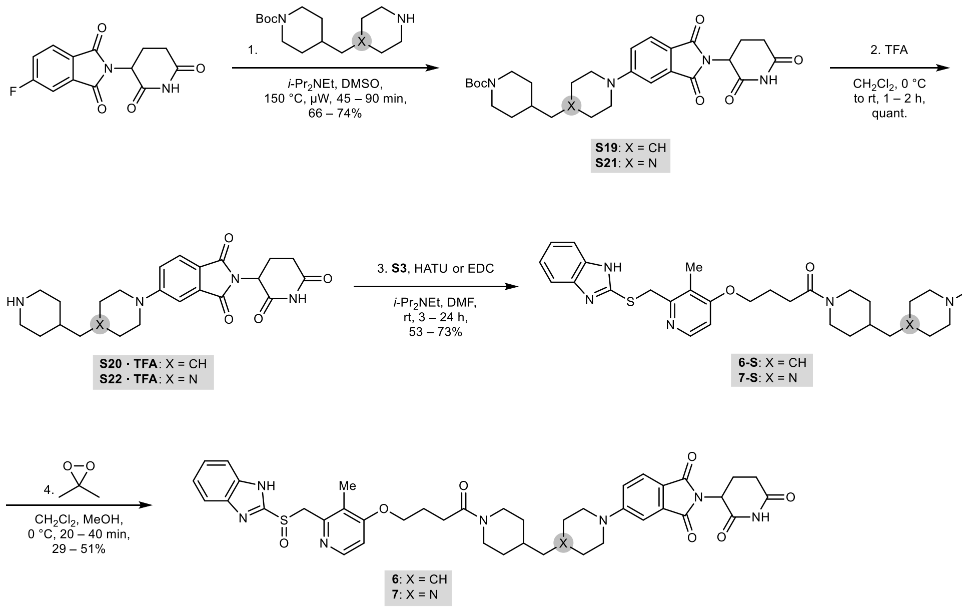

**Supplementary Scheme S3.** Synthesis route for rabeprazole-thalidomide hybrids **6-S**, **6**, **7-S**, and **7**.

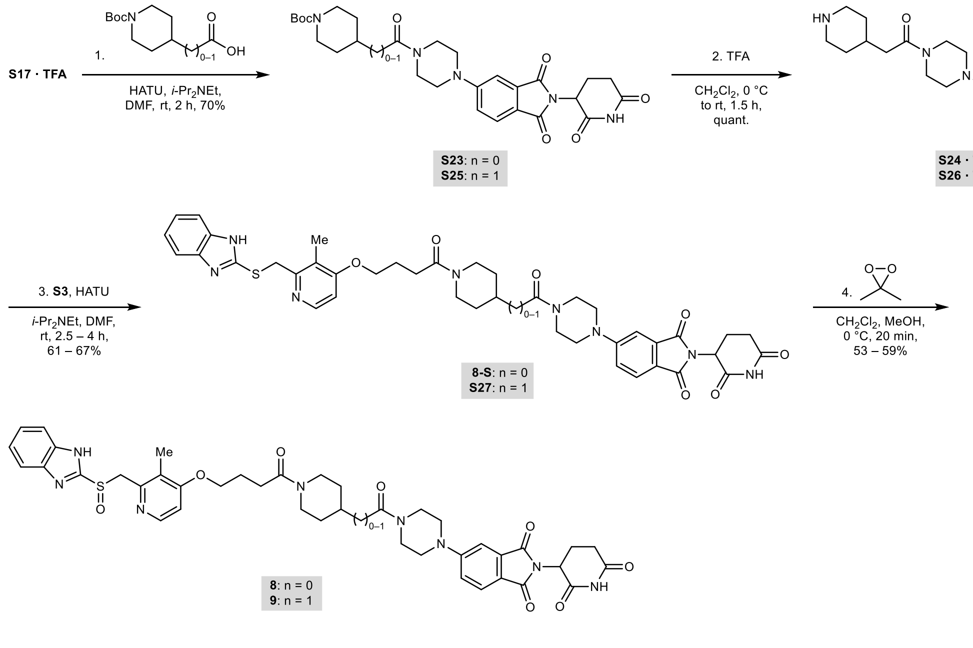

**Supplementary Scheme S4.** Synthesis route for rabeprazole-thalidomide hybrids **8-S**, **8**, and **9**.

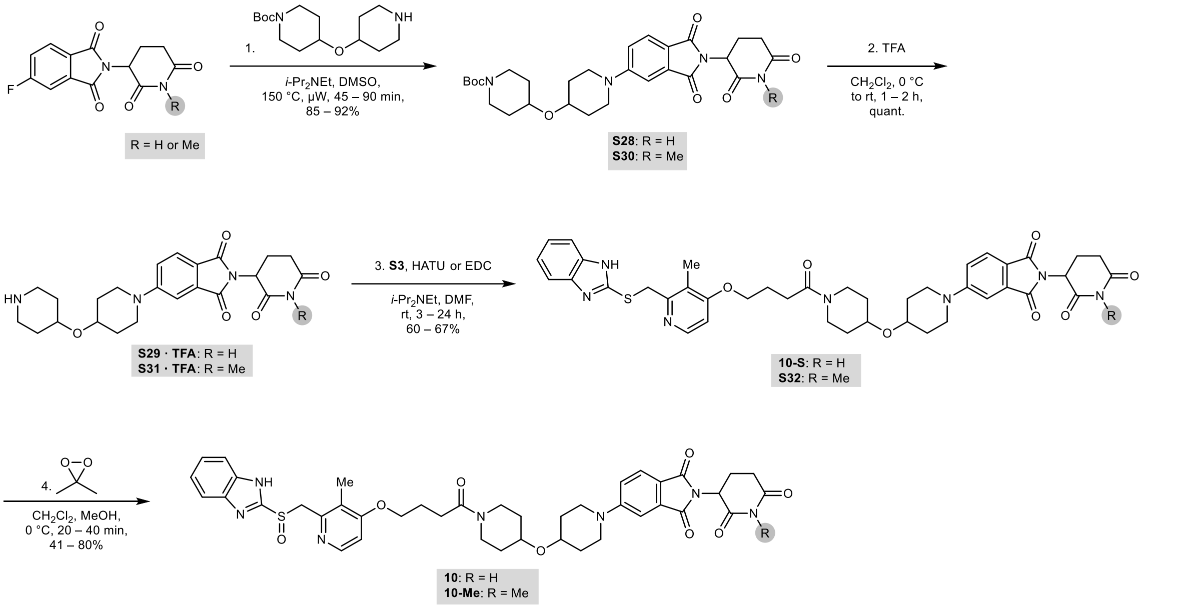

**Supplementary Scheme S5.** Synthesis route for rabeprazole-thalidomide hybrids **10-S**, **10**, and **10-Me**.

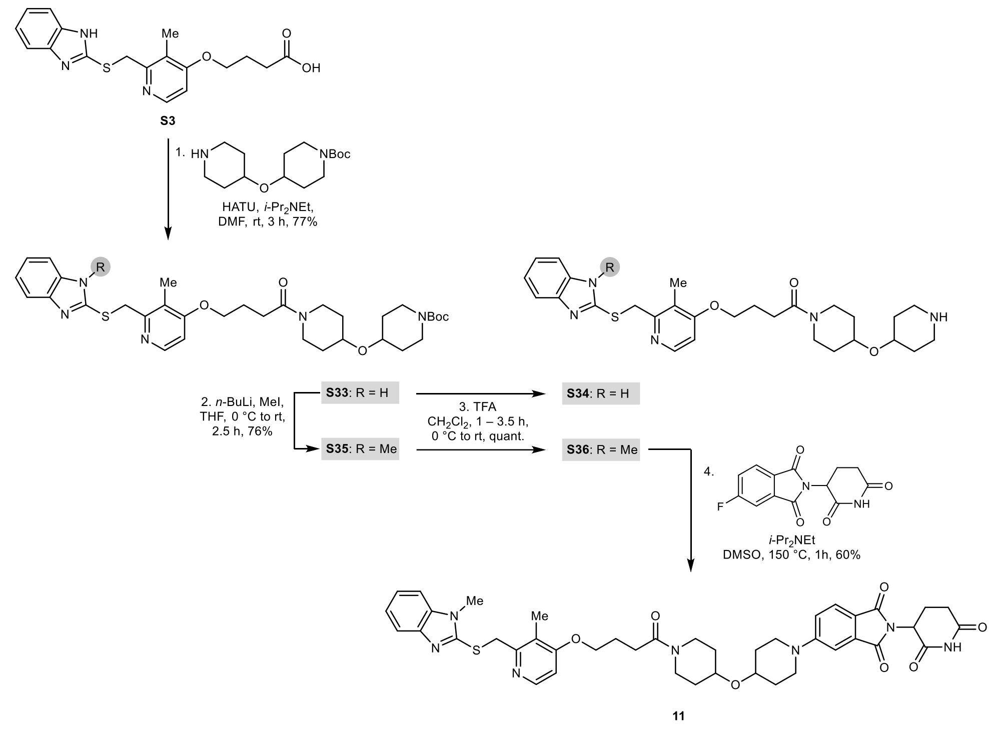

**Supplementary Scheme S6.** Synthesis route for truncated rabeprazole-thalidomide hybrid **11**.

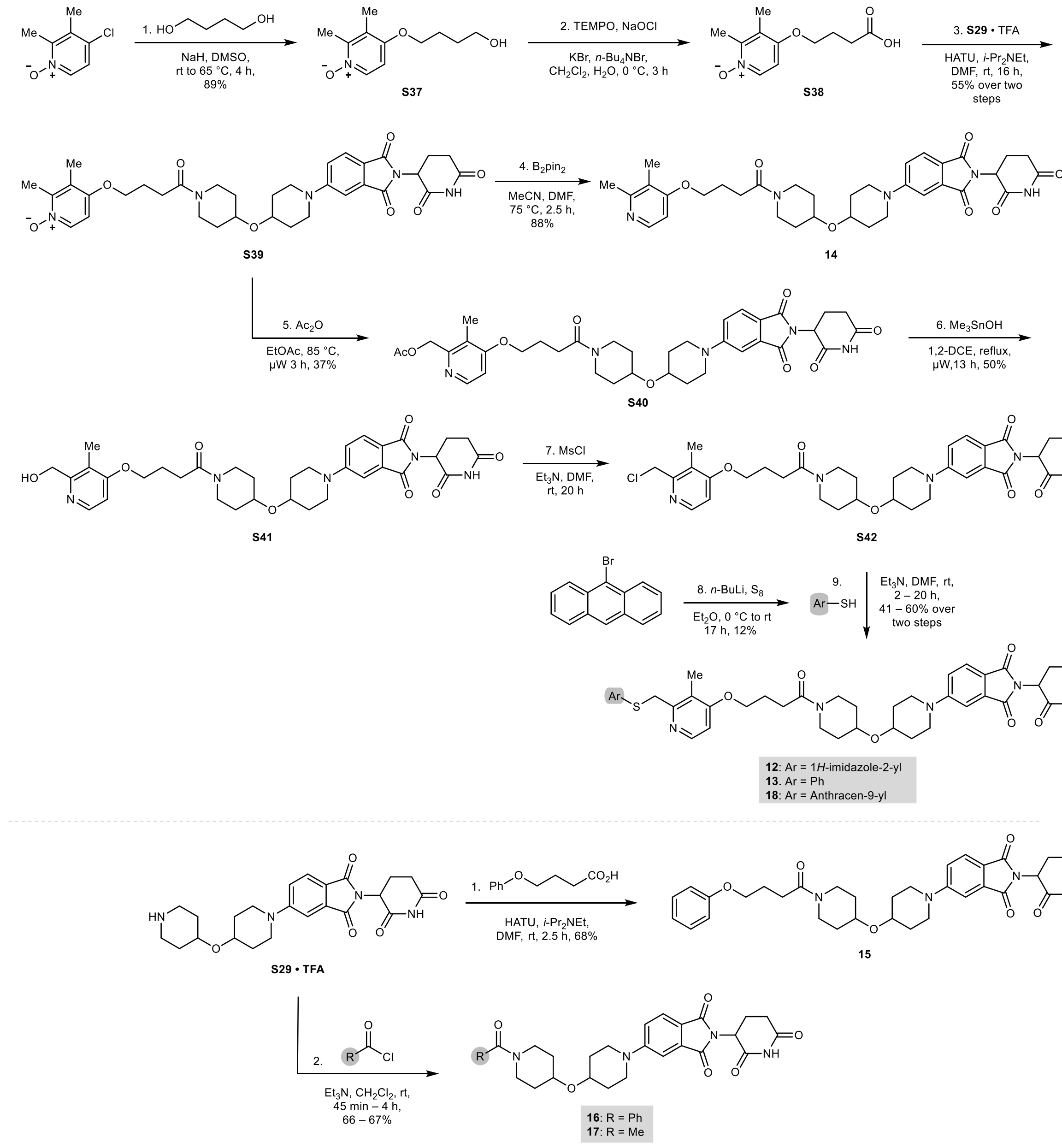

**Supplementary Scheme S7.** Synthesis route for truncated hybrids **12–18**.

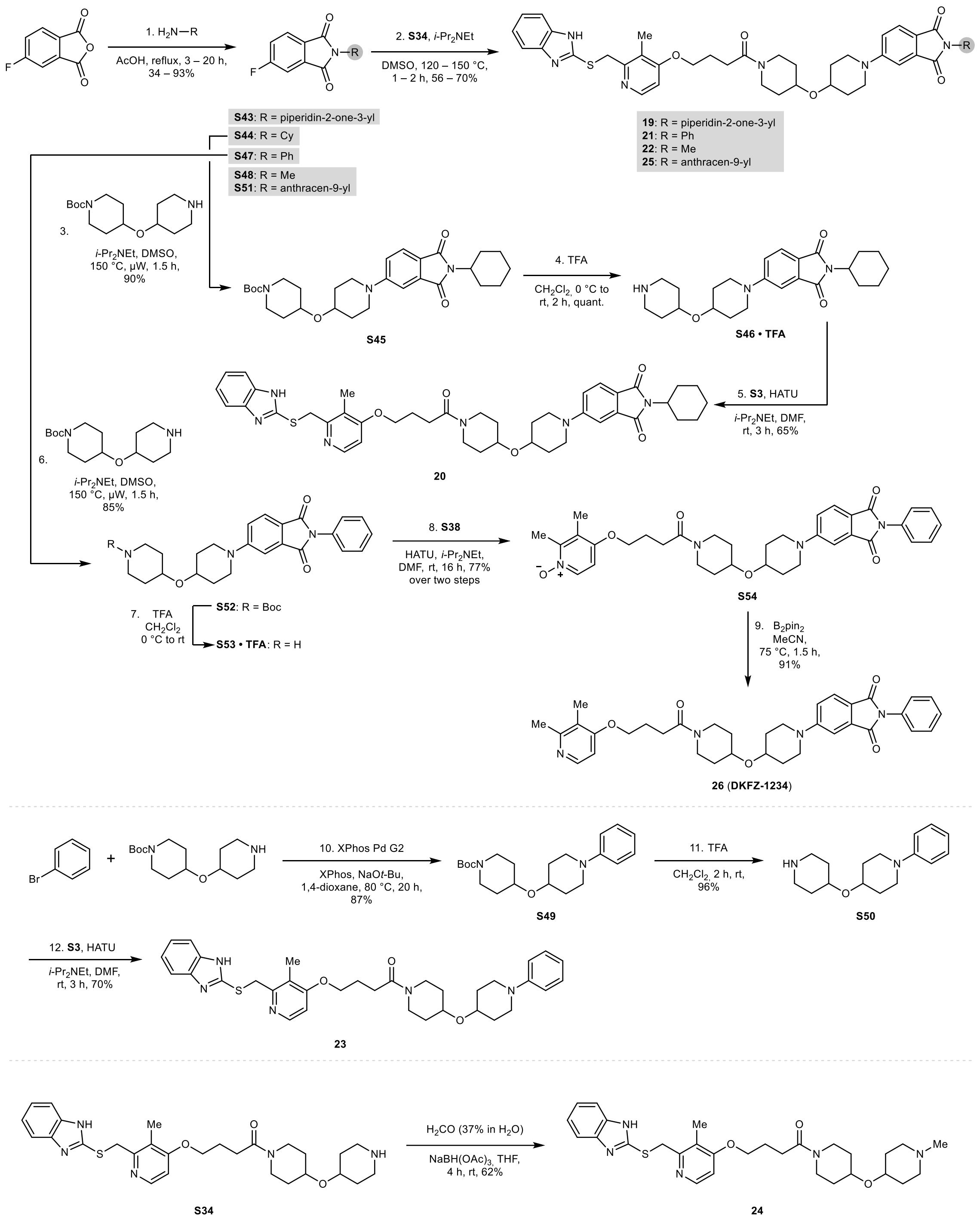

**Supplementary Scheme S8.** Synthesis route for truncated hybrids **19–26**.

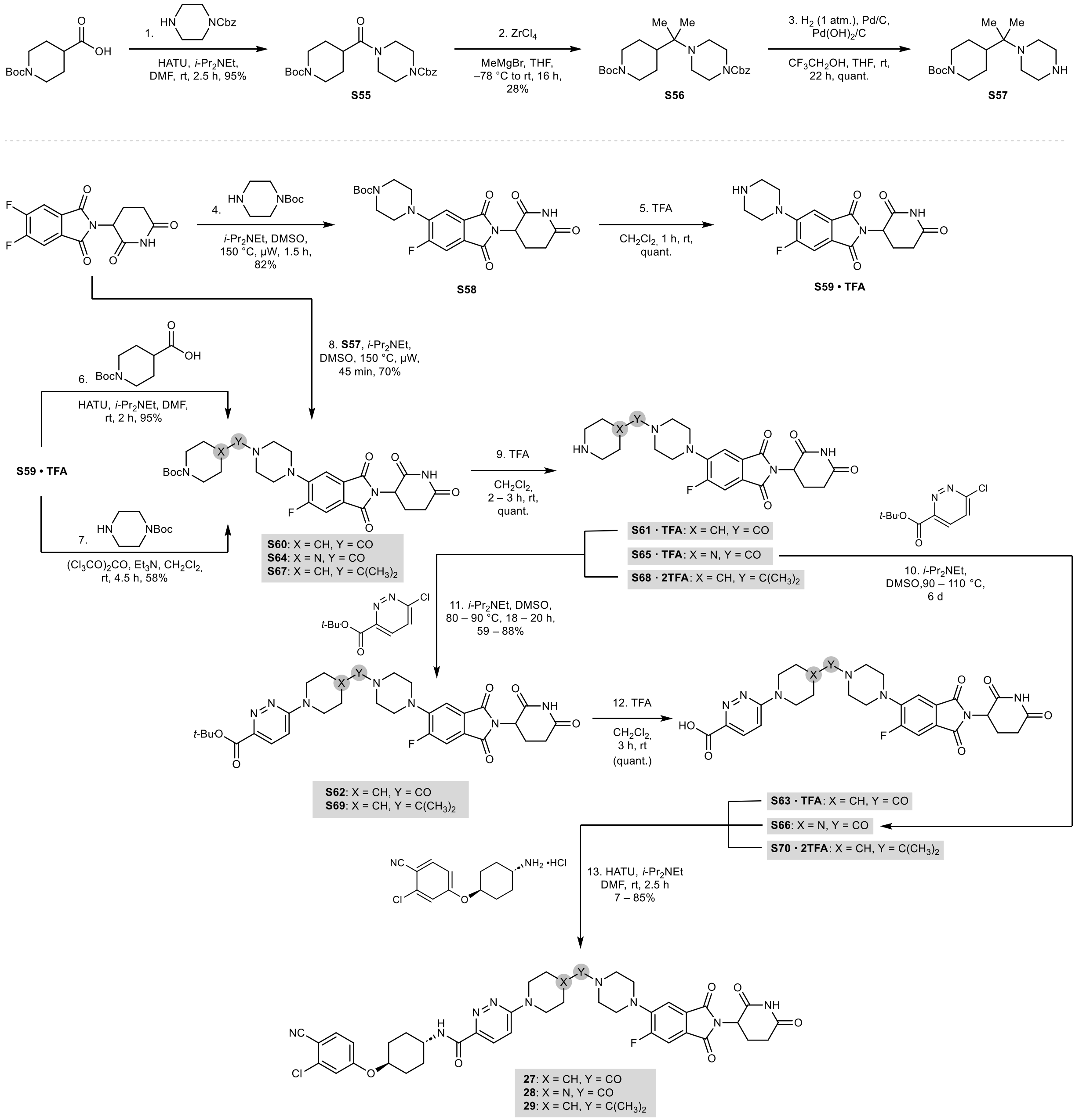

**Supplementary Scheme S9.** Synthesis route for conformationally constrained ARV-110 modifications **27–29**.

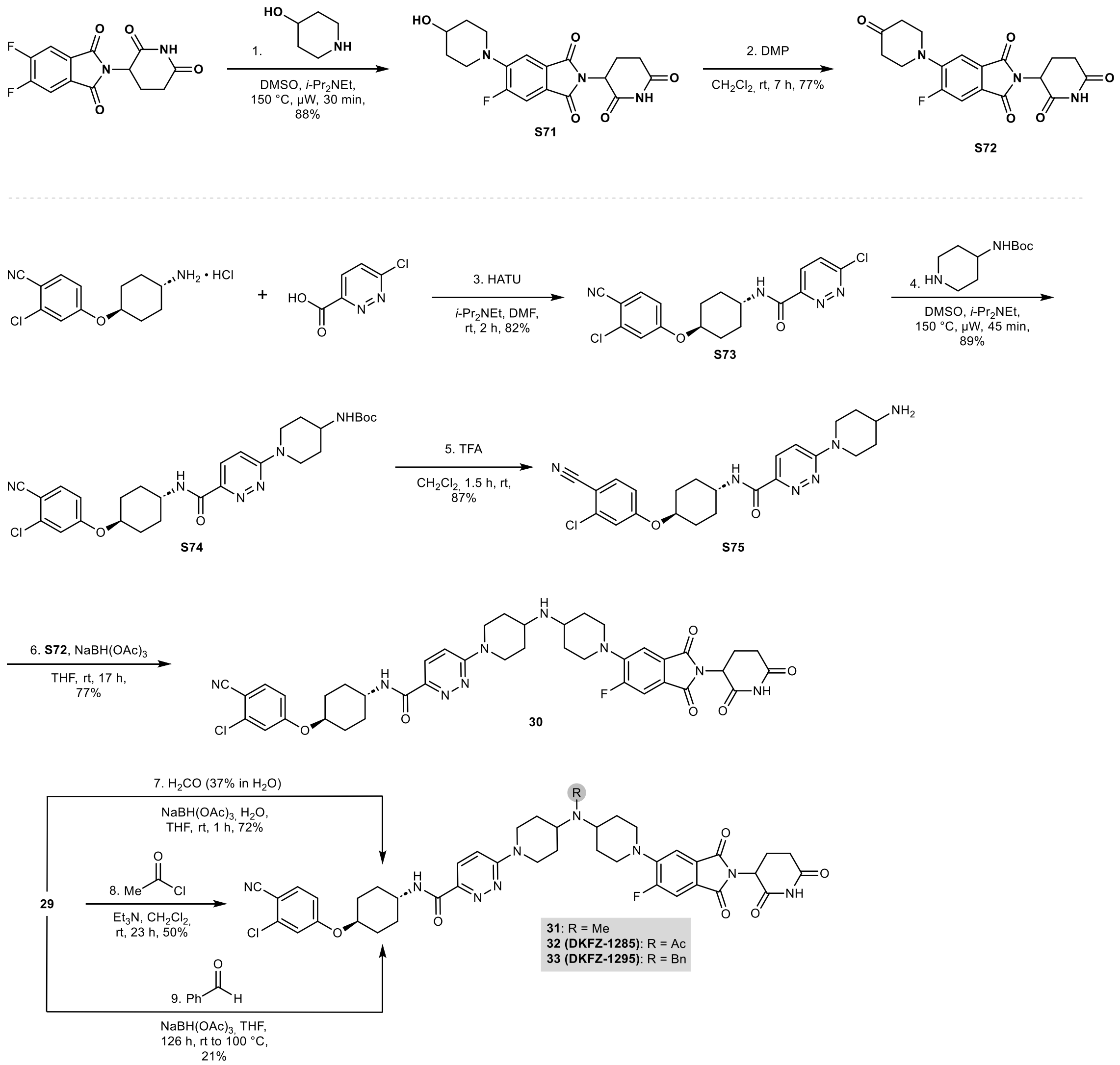

**Supplementary Scheme S10.** Synthesis route for “bumped” ARV-110 modifications **30–33**.

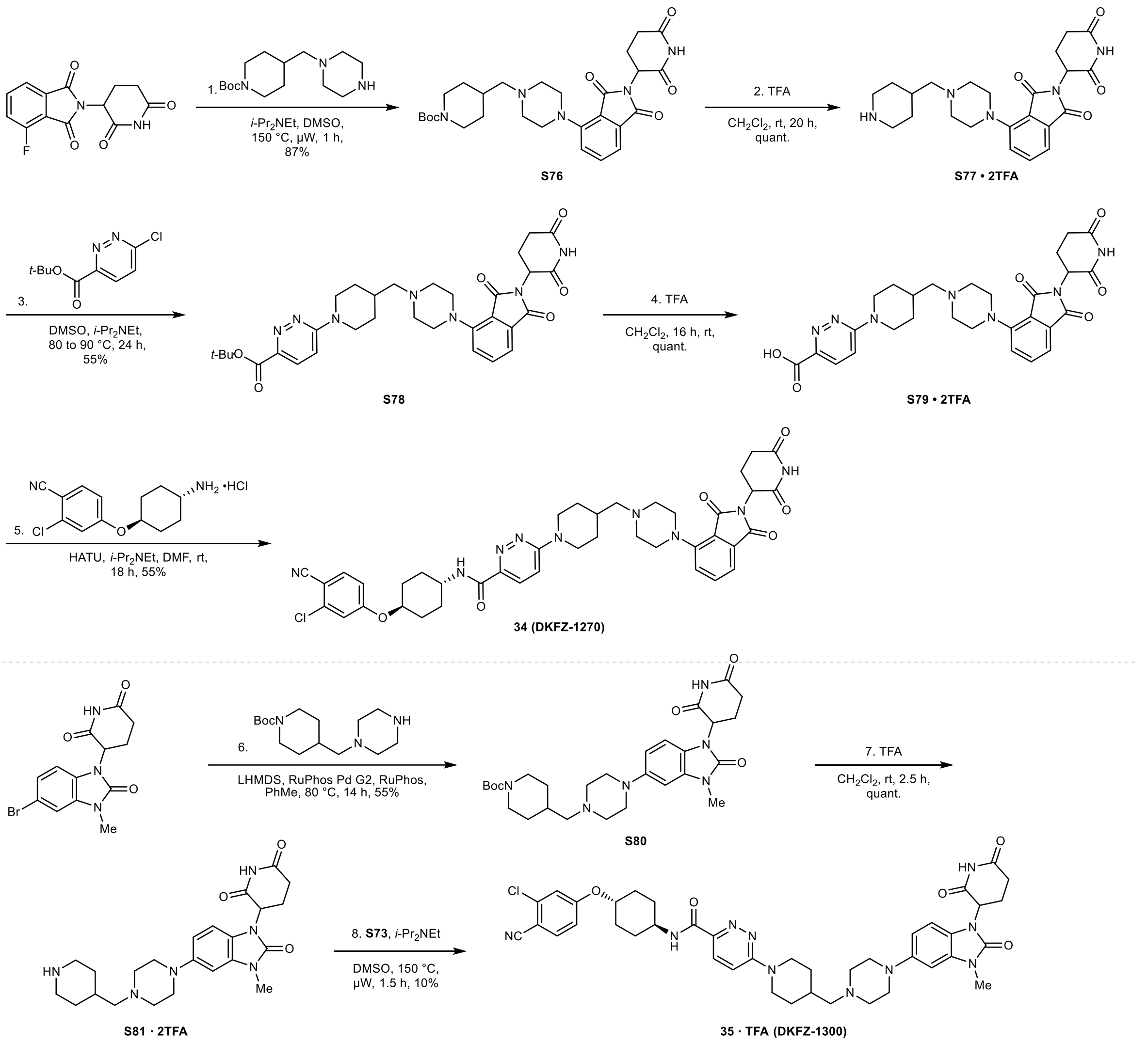

**Supplementary Scheme S11.** Synthesis route for “kinked” ARV-110 modifications **34** and **35** (**DKFZ-1300**).

### Cell Painting Assay and Analysis Protocol

The U-2OS female human bone osteosarcoma cell line was cultured in Dulbecco's Modified Eagle's medium (DMEM, high glucose) supplemented with 4 mM L-glutamine, 10% fetal bovine serum, 1 mM sodium pyruvate and non-essential amino acids. Cells were incubated at 37 °C and 5% $CO_2$ in humidified atmosphere. The MycoAlert® Mycoplasma Detection Kit was used according to the manufacturer's instruction to detect contamination with mycoplasma monthly. Cells were always tested free of mycoplasma.

Initially, 5 µL U-2OS medium were added to each well of a 384-well plate (PerkinElmer CellCarrier-384 Ultra). Subsequently, U-2OS cell were seeded with a density of 1,600 cells per well in 20 µL medium. The plate was incubated for 10 min at room temperature (rt), followed by an additional 4 h incubation (37 °C, 5% $CO_2$). Compound treatment was performed with the Echo 520 acoustic dispenser (Labcyte). Different concentrations of DMSO were used as controls dependent on the used compound concentration, e.g., 0.1% DMSO was used as a control for the profiling of compounds at 10 µM. Samples at a given compound concentration were compared to the DMSO sample of the same DMSO concentration. Incubation with compound was performed for 20 h (37 °C, 5% $CO_2$). Subsequently, mitochondria were stained with Mito Tracker Deep Red (Thermo Fisher Scientific, Cat. No. M22426). The Mito Tracker Deep Red stock solution (1 mM) was diluted to a final concentration of 100 nM in pre-warmed medium. The medium was removed from the plate leaving 10 µL residual volume and 25 µL of the Mito Tracker solution were added to each well. The plate was incubated for 30 min in darkness (37 °C, 5% $CO_2$). To fix the cells 7 µL of 18.5% formaldehyde in PBS was added, resulting in a final formaldehyde concentration of 3.7%. Subsequently, the plate was incubated for another 20 min in darkness (rt) and washed three times with 70 µL of PBS (Biotek Washer Elx405). Cells were permeabilized by addition of 25 µL 0.1% Triton X-100 to each well, followed by 15 min incubation (rt) in darkness. The cells were washed three times with PBS leaving a final volume of 10 µl. To each well, 25 µL of a staining solution was added, which contains 1% BSA, 5 $\mu L \cdot mL^{-1}$ Phalloidin (Alexa594 conjugate, Thermo Fisher Scientific, A12381), 25 $\mu g \cdot mL^{-1}$ Concanavalin A (Alexa488 conjugate, Thermo Fisher Scientific, Cat. No. C11252), 5 $\mu g \cdot mL^{-1}$ Hoechst-33342 (Sigma, Cat. No. B2261-25 mg), 1.5 $\mu g \cdot mL^{-1}$ WGA-Alexa594 conjugate (Thermo Fisher Scientific, Cat. No. W11262) and 1.5 µM SYTO 14 solution (Thermo Fisher Scientific, Cat. No. S7576). The plate is incubated for 30 min (rt) in darkness and washed three times with 70 µL PBS. After the final washing step, the PBS was not aspirated. The plates were sealed and centrifuged for 1 min at 500 rpm.

The plates were prepared in triplicates with shifted layouts to reduce plate effects and imaged using a Micro XL High-Content Screening System (Molecular Devices) in 5 channels (DAPI: Ex350-400/ Em410-480; FITC: Ex470-500/Em510-540; Spectrum Gold: Ex520-545/Em560-585; TxRed: Ex535-585/Em600-650; Cy5: Ex605-650/Em670-715) with 9 sites per well and 20x magnification (binning 2).

The generated images were processed with the *CellProfiler* package (https://cellprofiler.org/, version 3.0.0) (Carpenter, Jones et al. 2006) on a computing cluster of the Max Planck Society to extract 1,716 cell features per microscope site. The data was then further aggregated as medians per well (9 sites -> 1 well), then over the three replicates.

Further analysis was performed with custom *Python* (https://www.python.org/) scripts using the *Pandas* (https://pandas.pydata.org/) and *Dask* (https://dask.org/) data processing libraries as well as the *Scientific Python* (https://scipy.org/) package.

From the total set of 1,716 features, a subset of highly reproducible and robust features was determined using the procedure described by Woehrmann et al.[1] in the following way: two biological repeats of one plate containing reference compounds were analyzed. For every feature, its full profile over each whole plate was calculated. If the profiles from the two repeats showed a similarity ≥0.8 (see below), the feature was added to the set. This procedure was only performed once and resulted in a set of 579 robust features out of the total of 1,716 that was used for all further analyses.

The phenotypic profiles were compiled from the Z-scores of all individual cellular features, where the Z-score is a measure of how far away a data point is from a median value. Specifically, Z-scores of test compounds were calculated relative to the Median of DMSO controls. Thus, the Z-score of a test compound defines how many MADs (Median Absolute Deviations) the measured value is away from the Median of the controls as illustrated by the following formula:

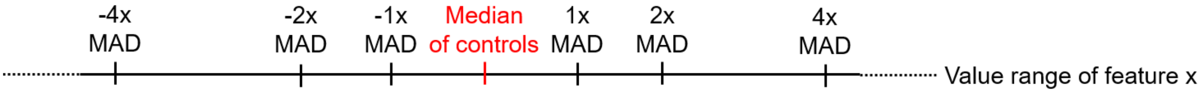

$$z\text{-}score = \frac{value_{meas.} - Median_{Controls}}{MAD_{Controls}}$$

The phenotypic compound profile is then determined as the list of Z-scores of all features for one compound. In addition to the phenotypic profile, an induction value was determined for each compound as the fraction of significantly changed features, in percent:

$$Induction\ [\%] = \frac{number\ of\ features\ with\ \text{abs.}\ values > 3}{total\ number\ of\ features}$$

Similarities of phenotypic profiles (termed Biosimilarity) were calculated from the correlation distances (CD) between two profiles (https://docs.scipy.org/doc/scipy/reference/generated/scipy.spatial.distance.correlation.html)[2]:

$$CD=1-\frac{(u-\overline{u}) \cdot (v-\overline{v})}{\|(u-\overline{u})\|_2\|(v-\overline{v})\|_2}$$

where $\overline{x}$ is the mean of the elements of $x$, $x \cdot y$ is the dot product of $x$ and $y$, and $\|x\|_2$ is the Euclidean norm of $x$:

$$\|x\|_2=\sqrt{x_1^2+x_2^2+\ldots+x_n^2}$$

The Biosimilarity is then defined as:

$$Biosimilarity=1-CD$$

Biosimilarity values smaller than 0 are set to 0 and the Biosimilarity is expressed in percent (0–100)."

Compounds are considered active in CPA for induction ≥5% as above this value the profiles become stable and reproducible. Two profiles are considered biosimilar for biosimilarity ≥75% as for biosimilarity values <75%, the number of biosimilar profiles increases drastically, because the comparisons become unspecific. Compounds are usually screened first at 10 µM or at 2 µM for reference compounds with low $IC_{50}$ values for the annotated target. If compounds were inactive or had low induction values, higher concentrations were tested. If the induction values were too high, lower concentrations were tested.

Cluster subprofiles were generated as recently described[3,4]. "For a set of cluster-defining profiles, dominating features were extracted as follows: for each profile, the sign for each of the 579 features value was assessed. The counter for positive or negative values was determined. For all cluster-defining compounds, the maximum of the two counters was determined and divided by the total number of defining profiles. A given feature was added to the cluster profile if its value has the same sign (i.e., positive or negative feature values) for 85% of the defining profiles. Afterwards, a representative median subprofile for the cluster was calculated by taking the median values over all cluster-defining profiles for every given feature and combining them into a new reduced profile. This median (*consensus*) subprofile was then used to calculate the biosimilarity of test compounds to the defined cluster subprofiles. Due to the shorter cluster subprofiles, the cluster biosimilarity threshold was set to 80%."

### Synthesis Notes

#### General Information for Chemical Synthesis

All reagents and solvents were purchased from commercial sources at the highest level of purity and used without purification. All reactions were stirred magnetically and performed under air unless otherwise noted. External bath temperatures are reported for heating. Anhydrous $CH_2Cl_2$, $Et_2O$, MeCN, THF, and toluene were prepared using an MBraun 800 Solvent Purification System. Degassed solvents were prepared by bubbling nitrogen through the solvent under sonication for at least 10 minutes. Standard inert techniques were employed for handling air- and moisture-sensitive reagents using oven-dried glassware under argon. Solutions of 3,3-dimethyldioxirane (DMDO) were prepared according to the procedure of Adam et al.[5] and stored over 4 Å activated molecular sieves at –80 °C. The concentration of DMDO was measured before each use via iodometric titration. Analytical thin-layer chromatography (TLC) was carried out on Merck silica gel 60 F254 glass plates. Detection was performed using UV light at 254 nm and 365 nm, iodine-saturated silica gel, or a basic $KMnO_4$ stain followed by heating. Analytical LC/MS was performed on an Agilent 1260 Infinity system equipped with a 6120 Series Single Quadrupole Electrospray and an evaporative light scattering detector (ELSD) using a Phenomenex Kinetex 2.6 µm or Gemini 5 µm C18 column (50 × 2.1 mm) and the following methods: T = 40 °C; solvent A = $H_2O$ with 0.01% formic acid or 0.1% $NH_4OH$; solvent B = MeCN; flow rate = 0.6 mL·min$^{-1}$; gradient: 99% to 10% A over 6 min, then 10% to 1% A over 2 min. Purification was performed by automated flash column chromatography using a Teledyne Isco CombiFlash NextGen 300+ or CombiFlash Rf 200 system with RediSep normal-phase silica flash columns and RediSep reversed-phase C18 flash columns, respectively, or by preparative HPLC on an Agilent 1260 Infinity as specified: Kinetex 5 µM C18 100 Å LC Column (250 × 30 mm); T = 40 °C; solvent A = $H_2O$ with 0.05% TFA, solvent B = MeCN; flow rate = 50 mL·min$^{-1}$. High-resolution electrospray ionization mass spectrometry (HRMS) was recorded on a Bruker ApexQe FT-ICR instrument. NMR spectra were recorded on Bruker Avance III 400 (400 MHz) or Avance 600 (600 MHz) spectrometers at 298.1 K unless otherwise noted. $^1$H and $^{13}$C spectra were referenced to (residual) solvent signals ($C\mathit{H}Cl_3$ = 7.26 ppm; $CDCl_3$ = 77.16 ppm; DMSO-$d_5$ = 2.50 ppm; DMSO-$d_6$ = 39.52 ppm; $CD_2\mathit{H}OD$ = 3.31 ppm; $CD_3OD$ = 49.00 ppm). NMR and LC/MS data were analyzed using MestReNova. Reported yields of intermediates take into account the most prominent impurities found in $^1$H NMR, $^{13}$C NMR, and $^{19}$F NMR spectra.

#### Synthesis Procedures

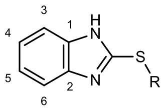

**General note 1:** Benzimidazole tautomerism can lead to signal broadening in the $^1$H and $^{13}$C NMR spectra. As a result, carbon signals for positions 1–6 may become undetectable. This is indicated in the characterization data for the respective compounds when this was observed.

**General note 2:** Rabeprazole-thalidomide hybrids **1–10** and **10-Me** are sensitive to acid due to the acid-mediated Smiles rearrangement of rabeprazole. Purification by column chromatography on silica gel using MeOH in $CH_2Cl_2$ and solvent removal at 40 °C resulted in partial decomposition. Filtration of $CH_2Cl_2$ through basic aluminum oxide prior to chromatography to remove traces of acid successfully resolved this problem. Alternative purification methods like column chromatography on silica gel using $CH_2Cl_2$ stored over KOH, or by reversed-phase column chromatography on C18 using MeCN in $H_2O$ with 0.1% $NH_4OH$ resulted in partial hydrolysis of the glutarimide.

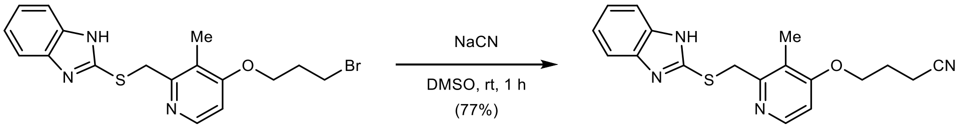

**4-((2-(((1*H*-Benzo[*d*]imidazol-2-yl)thio)methyl)-3-methylpyridin-4-yl)oxy)butanenitrile (S2):** To a solution of **S1** (5.84 g, 14.1 mmol, 1.00 equiv.)[6] in DMSO (70 mL) was added NaCN (6.93 g, 141 mmol, 10.0 equiv.) at rt. The reaction was stirred at rt for 1 h. The mixture was poured into saturated, aqueous $NaHCO_3$ (500 mL) and extracted with EtOAc (3 × 150 mL). The combined organic layers were washed with water (2 × 150 mL) and brine (150 mL), dried with $MgSO_4$, filtered, and concentrated *in vacuo*. Purification by column chromatography (220 g silica, 0 to 9% MeOH in $CH_2Cl_2$ in 20 min) yielded 3.70 g (10.9 mmol, 77%) of **S2** as an off-white solid. **Note:** This reaction can also be conducted with fewer equivalents of NaCN and longer reaction times.

**TLC** $R_f$ = 0.61 (10% MeOH with 0.5% $NH_4OH$ in $CH_2Cl_2$).

**LC/MS** (ESI) *m/z*: $(M+H)^+$ 339.1.

**$^1$H NMR** (400 MHz, $CDCl_3$) δ 8.37 (dd, *J* = 5.8, 0.6 Hz, 1H), 7.60 – 7.48 (m, 2H), 7.22 – 7.14 (m, 2H), 6.75 (d, *J* = 5.7 Hz, 1H), 4.39 (s, 2H), 4.16 (t, *J* = 5.7 Hz, 2H), 2.61 (t, *J* = 7.0 Hz, 2H), 2.27 (s, 3H), 2.26 – 2.17 (m, 2H) ppm. **Note:** The NH peak is in exchange with residual water in the spectrum.

**$^{13}$C NMR** (101 MHz, $CDCl_3$) δ 163.7, 157.1, 151.7, 147.5, 122.0, 121.2, 118.8, 106.1, 66.0, 35.0, 25.3, 14.4, 10.9 ppm. **Note:** C1, C2, C3, and C6 are not detected due to benzimidazole tautomerism.

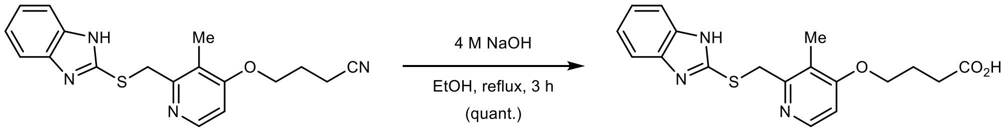

**4-((2-(((1*H*-Benzo[*d*]imidazol-2-yl)thio)methyl)-3-methylpyridin-4-yl)oxy)butanoic acid (S3):** To a solution of **S2** (3.70 g, 10.9 mmol, 1.00 equiv.) in EtOH (18 mL) was added 4 M aqueous NaOH (35.5 mL, 142 mmol, 13.0 equiv.) at rt. The reaction was heated to reflux for 3 h. The mixture was cooled to rt and concentrated *in vacuo*. The pH was adjusted to 6 by the dropwise addition of concentrated HCl at 0 °C. A colorless precipitate formed which was collected by filtration using a sintered glass funnel, washed with aqueous HCl (pH = 6; 50 mL), and dried *in vacuo*, yielding 3.92 g (10.9 mmol, quantitative) of **S3** as a colorless powder. **Note:** Further acidification below pH = 6 resulted in the formation of **S3 · HCl** (see procedure below).

**TLC** $R_f$ = 0.46, streaky (20% MeOH with 0.5% AcOH in $CH_2Cl_2$).

**LC/MS** (ESI) *m/z*: $(M+H)^+$ 358.1.

**$^{1}$H NMR** (400 MHz, DMSO-$d_6$) δ 8.20 (dd, *J* = 5.7, 0.6 Hz, 1H), 7.51 – 7.39 (m, 2H), 7.15 – 7.05 (m, 2H), 6.96 (d, *J* = 5.7 Hz, 1H), 4.69 (s, 2H), 4.07 (t, *J* = 6.6 Hz, 2H), 2.20 (s, 3H), 2.11 (t, *J* = 7.1 Hz, 2H), 1.91 (p, *J* = 6.9 Hz, 2H) ppm. **Note:** The NH peak is in exchange with residual water in the spectrum.

**$^{13}$C NMR** (101 MHz, DMSO-$d_6$) δ 176.3, 162.8, 154.6, 150.4, 147.7, 139.9, 121.2, 119.7, 113.7, 106.3, 68.2, 36.3, 33.2, 25.5, 10.4 ppm.

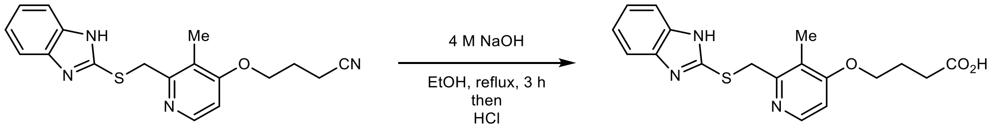

**S2** **S3 · HCl**

**4-((2-(((1*H*-Benzo[*d*]imidazol-2-yl)thio)methyl)-3-methylpyridin-4-yl)oxy)butanoic acid (S3 · HCl):** To a solution of **S2** (334 mg, 0.99 mmol, 1.00 equiv.) in EtOH (5.0 mL) was added 4 M aqueous NaOH (10.0 mL, 40.0 mmol, 40.5 equiv.) at rt. The reaction was heated to reflux for 3 h. The mixture was cooled to rt and concentrated *in vacuo*. The pH was adjusted to 4 by the dropwise addition of 3 M aqueous HCl at 0 °C. A colorless precipitate formed which was collected by filtration using a sintered glass funnel, washed with ice-cold water (10 mL), and dried *in vacuo*, yielding 240 mg (0.67 mmol, 68%) of **S3 · HCl** as a colorless powder.

**TLC** $R_f$ = 0.46, streaky (20% MeOH with 0.5% AcOH in $CH_2Cl_2$).

**LC/MS** (ESI) *m/z*: $(M+H)^+$ 358.1.

**$^{1}$H NMR** (400 MHz, DMSO-$d_6$) δ 8.56 (d, *J* = 6.5 Hz, 1H), 7.54 – 7.45 (m, 2H), 7.37 (d, *J* = 6.6 Hz, 1H), 7.22 – 7.13 (m, 2H), 4.85 (s, 2H), 4.27 (t, *J* = 6.2 Hz, 2H), 2.42 (t, *J* = 7.2 Hz, 2H), 2.27 (s, 3H), 2.02 (p, *J* = 6.7 Hz, 2H) ppm. **Note:** The NH peak is in exchange with residual water in the spectrum.

**$^{13}$C NMR** (101 MHz, DMSO-$d_6$) δ 177.2, 170.4, 154.1, 151.8, 146.3, 142.1, 126.5, 125.3, 117.3, 111.1, 72.5, 35.6, 33.3, 27.0, 13.7 ppm.

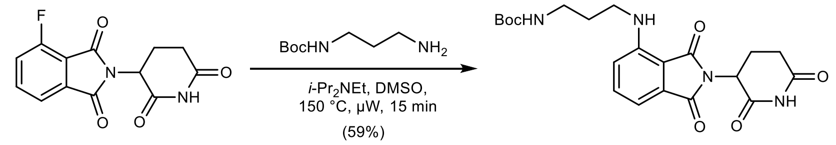

**S4**

***tert*-Butyl (3-((2-(2,6-dioxopiperidin-3-yl)-1,3-dioxoisoindolin-4-yl)amino)propyl)carbamate (S4):** To a solution of 2-(2,6-dioxopiperidin-3-yl)-4-fluoroisoindoline-1,3-dione (100 mg, 0.36 mmol, 1.00 equiv.) in anhydrous DMSO (1.8 mL) was added *tert*-butyl (3-aminopropyl) carbamate (75 μL, 0.42 mmol, 1.15 equiv.) and *i*-$Pr_2NEt$ (189 μL, 1.09 mmol, 3.00 equiv.) at rt under argon. The reaction was stirred at 150 °C for 15 min under microwave irradiation. The mixture was diluted with water (100 mL) and extracted with EtOAc (3 × 25 mL). The combined organic layers were washed with water (50 mL) and brine (50 mL), dried with $MgSO_4$, filtered, and concentrated *in vacuo*. Purification by column chromatography (24 g silica, 30 to 70% EtOAc in *n*-heptane in 12 min) yielded 92 mg (0.21 mmol, 59%) of **S4** as a bright-yellow solid.

**TLC** $R_f$ = 0.52 (10% MeOH in $CH_2Cl_2$).

**LC/MS** (ESI) *m/z*: $(M+Na)^+$ 453.1.

**$^{1}$H NMR** (400 MHz, $CDCl_3$) δ 8.08 (s, 1H), 7.49 (dd, *J* = 8.5, 7.1 Hz, 1H), 7.10 (dd, *J* = 7.1, 0.6 Hz, 1H), 6.89 (d, *J* = 8.5 Hz, 1H), 6.31 (s, 1H), 4.96 – 4.87 (m, 1H), 4.64 (s, 1H), 3.33 (q, *J* = 7.0 Hz, 2H),

3.25 (q, $J$ = 6.5 Hz, 2H), 2.94 – 2.66 (m, 3H), 2.18 – 2.06 (m, 1H), 1.84 (p, $J$ = 6.8 Hz, 2H), 1.45 (s, 9H) ppm.

**$^{13}$C NMR** (101 MHz, $CDCl_3$) δ 169.6, 168.4, 167.7, 156.2, 146.9, 136.3, 132.7, 116.7, 113.5, 111.8, 110.3, 79.7, 49.0, 40.2, 38.2, 31.6, 30.1, 28.5, 22.9 ppm.

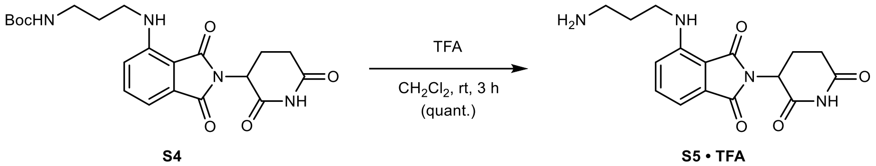

**4-((3-Aminopropyl)amino)-2-(2,6-dioxopiperidin-3-yl)isoindoline-1,3-dione (S5 · TFA):** To a solution of **S4** (85 mg, 0.20 mmol, 1.00 equiv.) in $CH_2Cl_2$ (1.0 mL) was added trifluoroacetic acid (1.0 mL, 13.1 mmol, 66.0 equiv.) slowly at 0 °C. After 5 min, the reaction was allowed to warm to rt and stirred for 3 h. The mixture was concentrated *in vacuo* – residual trifluoroacetic acid was removed by co-evaporation with MeOH (3 × 5 mL) and drying *in vacuo*, yielding 88 mg (0.20 mmol, quantitative) of **S5 · TFA** as a yellow, sticky solid.

**TLC** $R_f$ = 0.10, streaky (20% MeOH with 0.5% $NH_4OH$ in $CH_2Cl_2$).

**LC/MS** (ESI) *m/z*: $(M+H)^+$ 331.1.

**$^1$H NMR** (400 MHz, DMSO-$d_6$) δ 11.10 (s, 1H), 7.81 (bs, 3H), 7.60 (dd, $J$ = 8.6, 7.1 Hz, 1H), 7.14 (d, $J$ = 8.6 Hz, 1H), 7.05 (d, $J$ = 7.0 Hz, 1H), 6.75 (t, $J$ = 6.3 Hz, 1H), 5.06 (dd, $J$ = 12.8, 5.4 Hz, 1H), 3.45 – 3.36 (m, 2H), 2.97 – 2.80 (m, 3H), 2.66 – 2.52 (m, 2H), 2.03 (dtd, $J$ = 13.0, 6.3, 3.2 Hz, 1H), 1.84 (p, $J$ = 6.9 Hz, 2H) ppm.

**$^{13}$C NMR** (101 MHz, DMSO-$d_6$) δ 172.8, 170.1, 168.8, 167.3, 146.1, 136.3, 132.3, 117.2, 110.6, 109.4, 48.6, 38.7, 36.6, 31.0, 26.6, 22.2 ppm.

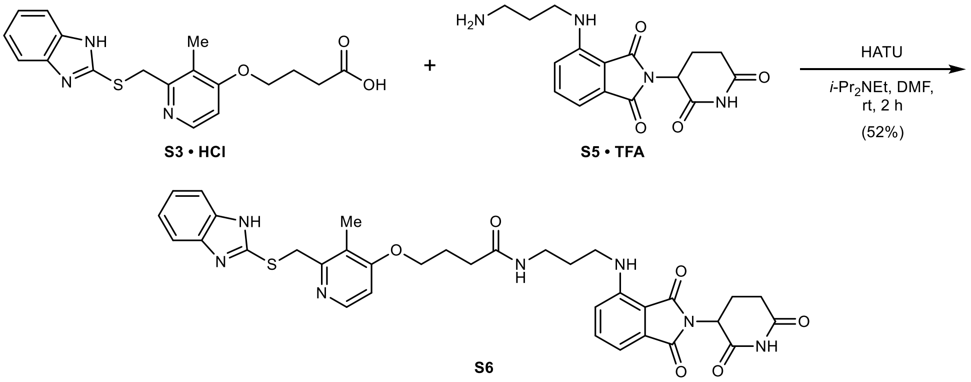

**4-((2-(((1*H*-Benzo[*d*]imidazol-2-yl)thio)methyl)-3-methylpyridin-4-yl)oxy)-*N*-(3-((2-(2,6-dioxopiperidin-3-yl)-1,3-dioxoisoindolin-4-yl)amino)propyl)butanamide (S6):** To a suspension of **S3 · HCl** (40 mg, 0.11 mmol, 1.00 equiv.) in anhydrous DMF (1.0 mL) was added **S5 · TFA** (88 mg, 0.20 mmol, 1.15 equiv.), HATU (51 mg, 0.13 mmol, 1.20 equiv.), and $i$-$Pr_2NEt$ (78 µL, 0.45 mmol, 4.40 equiv.) at rt under argon. The reaction was stirred at rt for 2 h. The mixture was poured into water (100 mL) and extracted with EtOAc (3 × 25 mL). The combined organic layers were washed with saturated aqueous $NaHCO_3$ (25 mL), saturated aqueous $NH_4Cl$ (25 mL), water (25 mL), and brine (25 mL), dried with $MgSO_4$, filtered, and concentrated *in vacuo*. Purification by column chromatography (12 g silica, 0 to 20% MeOH in $CH_2Cl_2$ in 12 min) yielded 35 mg (0.05 mmol, 52%) of **S6** as a bright-yellow solid.

**TLC** $R_f$ = 0.41 (10% MeOH in $CH_2Cl_2$).

**LC/MS** (ESI) *m/z*: $(M+H)^+$ 670.2.

**$^1$H NMR** (400 MHz, $CDCl_3$) δ 8.62 (s, 1H), 8.29 (d, *J* = 5.7 Hz, 1H), 7.57 – 7.46 (m, 2H), 7.42 (dd, *J* = 8.5, 7.1 Hz, 1H), 7.21 – 7.12 (m, 2H), 7.06 (d, *J* = 6.9 Hz, 1H), 6.80 (d, *J* = 8.5 Hz, 1H), 6.72 (d, *J* = 5.8 Hz, 1H), 6.34 (t, *J* = 5.8 Hz, 1H), 6.24 (t, *J* = 6.0 Hz, 1H), 4.89 (dd, *J* = 12.3, 5.4 Hz, 1H), 4.34 (s, 2H), 4.05 (t, *J* = 6.0 Hz, 2H), 3.37 (q, *J* = 6.5 Hz, 2H), 3.27 (q, *J* = 6.4 Hz, 2H), 2.88 – 2.63 (m, 3H), 2.39 (t, *J* = 7.1 Hz, 2H), 2.27 – 2.00 (m, 6H), 1.81 (p, *J* = 6.7 Hz, 2H) ppm. **Note:** The benzimidazole NH peak is in exchange with residual water in the spectrum.

**$^{13}$C NMR** (101 MHz, $CDCl_3$) δ 172.7, 171.9, 169.5, 169.1, 167.7, 164.2, 156.1, 151.5, 147.4, 146.7, 139.7, 136.2, 132.5, 122.0, 121.1, 116.7, 114.1, 111.5, 110.1, 106.3, 67.7, 49.0, 40.1, 37.0, 35.0, 32.5, 31.5, 29.2, 24.9, 22.8, 10.7 ppm.

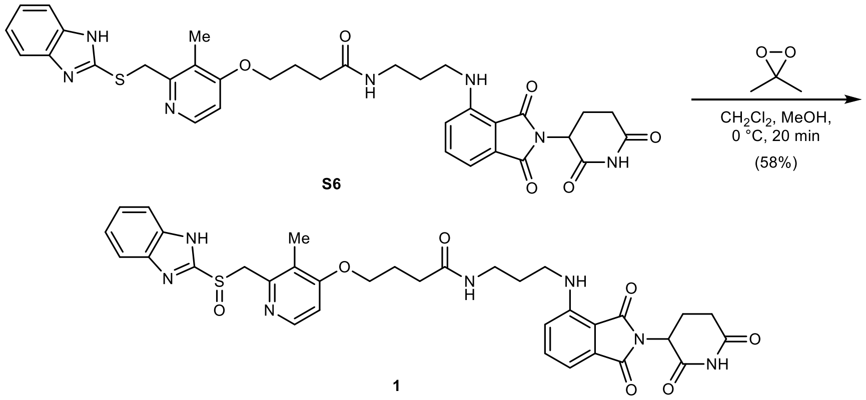

**4-((2-(((1*H*-Benzo[*d*]imidazol-2-yl)sulfinyl)methyl)-3-methylpyridin-4-yl)oxy)-*N*-(3-((2-(2,6-dioxopiperidin-3-yl)-1,3-dioxoisoindolin-4-yl)amino)propyl)butanamide (1):** To a solution of **S6** (35 mg, 0.05 mmol, 1.00 equiv.) in anhydrous $CH_2Cl_2$ (1.0 mL) and anhydrous MeOH (0.4 mL) was added a solution of 3,3-dimethyldioxirane (1.25 mL, 0.042 M in acetone by iodometric titration, 0.05 mmol, 1.00 equiv.) dropwise at 0 °C under argon. The reaction was stirred for 20 min at 0 °C, then concentrated *in vacuo*. Purification by column chromatography (12 g silica, 0 to 15% MeOH in $CH_2Cl_2$ in 13 min) yielded 21 mg (0.03 mmol, 58%) of **1** as a brown-yellow solid after lyophilization from MeCN/$H_2O$.

**TLC** $R_f$ = 0.40 (10% MeOH in $CH_2Cl_2$).

**HRMS** (ESI) *m/z*: $(M+H)^+$ calc. for $C_{34}H_{36}N_7O_7S$: 686.2391; found: 686.2395.

**$^1$H NMR** (400 MHz, $CD_3OD$) δ 8.11 (d, *J* = 5.7 Hz, 1H), 7.69 – 7.60 (m, 2H), 7.57 – 7.47 (m, 1H), 7.39 – 7.30 (m, 2H), 7.05 – 6.98 (m, 2H), 6.90 (d, *J* = 5.8 Hz, 1H), 5.03 (dd, *J* = 12.9, 5.1 Hz, 1H), 4.79 (d, *J* = 13.3 Hz, 1H), 4.74 (d, *J* = 13.3 Hz, 1H), 4.10 (t, *J* = 6.1 Hz, 2H), 3.35 (t, *J* = 6.7 Hz, 2H), 3.29 – 3.26 (m, 2H), 2.90 – 2.61 (m, 3H), 2.39 (t, *J* = 7.3 Hz, 2H), 2.17 (s, 3H), 2.15 – 2.03 (m, 3H), 1.82 (p, *J* = 6.7 Hz, 2H) ppm. **Note:** The NH peaks are in exchange with $CD_3OD$ in the spectrum.

**$^{13}$C NMR** (101 MHz, $CD_3OD$) δ 175.3, 174.7, 171.6, 170.7, 169.3, 165.4, 154.3, 150.1, 149.3, 148.0, 137.2, 134.0, 125.2, 124.8, 117.9, 111.9, 111.2, 107.7, 69.1, 61.4, 50.2, 41.1, 38.1, 33.5, 32.2, 30.0, 26.1, 23.8, 11.3 ppm. **Note:** C1, C2, C3, and C6 are not detected due to benzimidazole tautomerism.

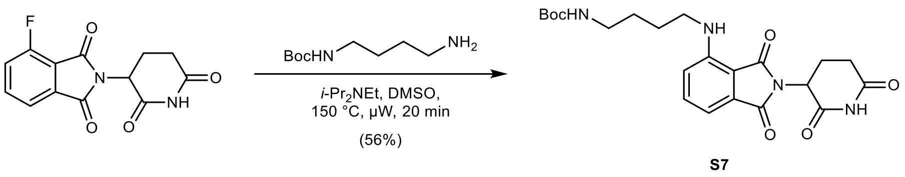

***tert*-Butyl (3-((2-(2,6-dioxopiperidin-3-yl)-1,3-dioxoisoindolin-4-yl)amino)propyl)carbamate (S7):** To a solution of 2-(2,6-dioxopiperidin-3-yl)-4-fluoroisoindoline-1,3-dione (100 mg, 0.36 mmol, 1.00 equiv.) in anhydrous DMSO (1.8 mL) was added *tert*-butyl (4-aminobutyl) carbamate (79 µL, 0.40 mmol, 1.10 equiv.) and *i*-$Pr_2NEt$ (189 µL, 1.09 mmol, 3.00 equiv.) at rt under argon. The reaction was stirred at 150 °C for 20 min under microwave irradiation. The mixture was diluted with water (100 mL) and extracted with EtOAc (3 × 25 mL). The combined organic layers were washed with water (50 mL) and brine (50 mL), dried with $MgSO_4$, filtered, and concentrated *in vacuo*. Purification by column chromatography (24 g silica, 30 to 70% EtOAc in *n*-heptane in 13 min) yielded 94 mg (0.21 mmol, 56%) of **S7** as a bright-yellow solid.

**TLC** $R_f$ = 0.60 (75% EtOAc in *n*-heptane).

**LC/MS** (ESI) *m/z*: $(M+Na)^+$ 467.2.

**$^1$H NMR** (400 MHz, $CDCl_3$) δ 8.06 (s, 1H), 7.49 (ddd, *J* = 8.5, 7.1, 0.6 Hz, 1H), 7.09 (dd, *J* = 7.1, 0.6 Hz, 1H), 6.89 (d, *J* = 8.5 Hz, 1H), 6.23 (t, *J* = 5.7 Hz, 1H), 4.91 (dd, *J* = 12.1, 5.3 Hz, 1H), 4.57 (s, 1H), 3.30 (q, *J* = 6.9 Hz, 2H), 3.17 (q, *J* = 6.6 Hz, 2H), 2.97 – 2.66 (m, 3H), 2.19 – 2.08 (m, 1H), 1.77 – 1.65 (m, 2H), 1.65 – 1.60 (m, 2H), 1.44 (s, 9H) ppm.

**$^{13}$C NMR** (101 MHz, $CDCl_3$) δ 170.9, 169.5, 168.3, 167.6, 156.0, 146.9, 136.2, 132.5, 116.7, 111.6, 110.0, 79.4, 48.9, 42.3, 40.1, 31.4, 28.4, 27.6, 26.5, 22.8 ppm.

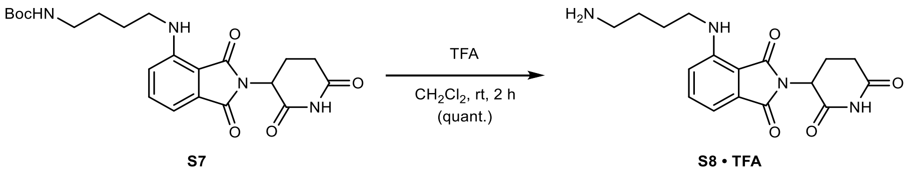

**4-((4-Aminobutyl)amino)-2-(2,6-dioxopiperidin-3-yl)isoindoline-1,3-dione (S8 · TFA):** To a solution of **S7** (90 mg, 0.19 mmol, 1.00 equiv.) in $CH_2Cl_2$ (1.0 mL) was added trifluoroacetic acid (1.0 mL, 13.2 mmol, 68.0 equiv.) slowly at 0 °C. After 5 min, the reaction was allowed to warm to rt and stirred for 2 h. The mixture was concentrated *in vacuo* – residual trifluoroacetic acid was removed by co-evaporation with MeOH (3 × 5 mL) and drying *in vacuo*, yielding 89 mg (0.19 mmol, quantitative) of **S8 · TFA** as a yellow, sticky solid.

**TLC** $R_f$ = 0.09, streaky (20% MeOH with 0.5% $NH_4OH$ in $CH_2Cl_2$).

**LC/MS** (ESI) *m/z*: $(M+H)^+$ 345.2.

**$^1$H NMR** (400 MHz, DMSO-$d_6$) δ 11.10 (s, 1H), 7.74 (s, 3H), 7.59 (dd, *J* = 8.6, 7.1 Hz, 1H), 7.12 (d, *J* = 8.6 Hz, 1H), 7.04 (d, *J* = 7.0 Hz, 1H), 6.61 (t, *J* = 6.1 Hz, 1H), 5.05 (dd, *J* = 12.7, 5.4 Hz, 1H), 3.37 – 3.26 (m, 2H), 2.96 – 2.77 (m, 3H), 2.65 – 2.52 (m, 2H), 2.09 – 1.97 (m, 1H), 1.66 – 1.56 (m, 4H) ppm.

**$^{13}$C NMR** (101 MHz, DMSO-$d_6$) δ 172.8, 170.1, 168.9, 167.3, 146.3, 136.3, 132.2, 117.3, 110.5, 109.1, 48.6, 41.2, 38.6, 31.0, 25.7, 24.5, 22.2 ppm.

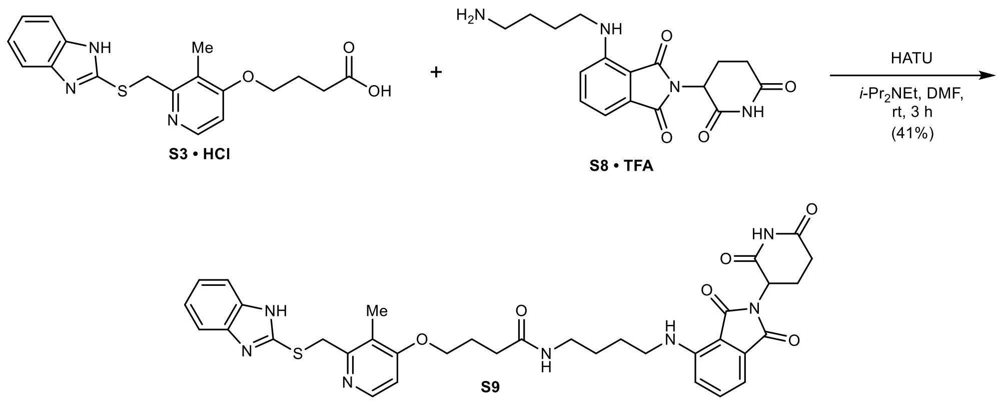

4-((2-(((1*H*-Benzo[*d*]imidazol-2-yl)thio)methyl)-3-methylpyridin-4-yl)oxy)-*N*-(4-((2-(2,6-dioxopiperidin-3-yl)-1,3-dioxoisoindolin-4-yl)amino)butyl)butanamide (S9): To a suspension of **S3 · HCl** (40 mg, 0.10 mmol, 1.00 equiv.) in anhydrous DMF (1.0 mL) was added HATU (46 mg, 0.12 mmol, 1.20 equiv.) and *i*-$Pr_2NEt$ (71 µL, 0.41 mmol, 4.00 equiv.) at rt under argon. After 5 min, **S8 · TFA** (47 mg, 0.10 mmol, 1.00 equiv.) was added and the reaction was stirred at rt for 3 h. The mixture was poured into water (100 mL) and extracted with EtOAc (3 × 25 mL). The combined organic layers were washed with saturated aqueous $NaHCO_3$ (25 mL), saturated aqueous $NH_4Cl$ (25 mL), water (25 mL), and brine (25 mL), dried with $MgSO_4$, filtered, and concentrated *in vacuo*. Purification by column chromatography (12 g silica, 0 to 15% MeOH in $CH_2Cl_2$ in 13 min) yielded 29 mg (0.04 mmol, 41%) of **S9** as a bright-yellow solid.

**TLC** $R_f$ = 0.38 (10% MeOH in $CH_2Cl_2$).

**LC/MS** (ESI) *m/z*: $(M+H)^+$ 684.2.

**$^1$H NMR** (400 MHz, $CD_3OD$) δ 8.17 (dd, *J* = 5.8, 0.6 Hz, 1H), 7.54 – 7.44 (m, 3H), 7.23 – 7.12 (m, 2H), 7.04 – 6.93 (m, 2H), 6.88 (d, *J* = 5.8 Hz, 1H), 5.03 (dd, *J* = 12.5, 5.5 Hz, 1H), 4.59 (s, 2H), 4.09 (t, *J* = 6.1 Hz, 2H), 3.25 (t, *J* = 6.5 Hz, 2H), 3.22 (t, *J* = 6.4 Hz, 2H), 2.90 – 2.77 (m, 1H), 2.77 – 2.61 (m, 2H), 2.39 (t, *J* = 7.2 Hz, 2H), 2.23 (s, 3H), 2.17 – 2.01 (m, 3H), 1.68 – 1.51 (m, 4H) ppm. **Note:** The NH peaks are in exchange with $CD_3OD$ in the spectrum.

**$^{13}$C NMR** (101 MHz, $CD_3OD$) δ 175.1, 174.7, 171.6, 170.8, 169.3, 165.5, 155.9, 151.2, 148.8, 148.2, 137.2, 133.9, 123.5, 122.8, 118.0, 111.8, 111.1, 107.5, 68.9, 50.2, 43.0, 39.9, 37.5, 33.4, 32.2, 27.8, 27.7, 26.2, 23.8, 10.9 ppm. **Note:** C1, C2, C3, and C6 are not detected due to benzimidazole tautomerism.

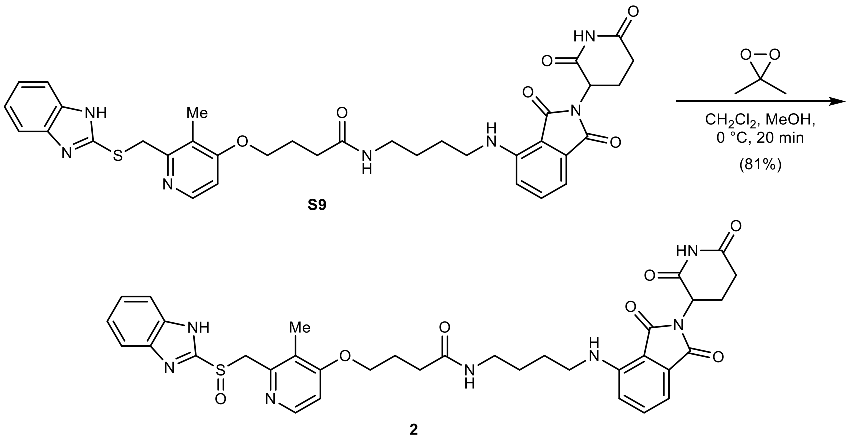

**4-((2-(((1*H*-Benzo[*d*]imidazol-2-yl)sulfinyl)methyl)-3-methylpyridin-4-yl)oxy)-*N*-(4-((2-(2,6-dioxopiperidin-3-yl)-1,3-dioxoisoindolin-4-yl)amino)butyl)butanamide** (**2**): To a solution of **S9** (28 mg, 0.04 mmol, 1.00 equiv.) in anhydrous $CH_2Cl_2$ (1.0 mL, stored over KOH) and anhydrous MeOH (0.5 mL) was added a solution of 3,3-dimethyldioxirane (573 µL, 0.065 M in acetone by iodometric titration, 0.04 mmol, 1.00 equiv.) dropwise at 0 °C under argon. The reaction was stirred for 20 min at 0 °C, then concentrated *in vacuo*. Purification by column chromatography (4 g silica, 0 to 25% MeOH in $CH_2Cl_2$ in 12 min) yielded 21 mg (0.03 mmol, 81%) of **2** as a yellow solid.

**TLC** $R_f$ = 0.36 (10% MeOH with 0.5% $NH_4OH$ in $CH_2Cl_2$).

**HRMS** (ESI) *m/z*: $(M+H)^+$ calc. for $C_{35}H_{38}N_7O_7S$: 700.2548; found: 700.2551.

**$^1$H NMR** (400 MHz, DMSO-$d_6$) δ 13.56 (s, 1H), 11.09 (s, 1H), 8.20 (dd, *J* = 5.7, 0.6 Hz, 1H), 7.88 (t, *J* = 5.6 Hz, 1H), 7.68 – 7.59 (m, 2H), 7.56 (dd, *J* = 8.6, 7.1 Hz, 1H), 7.33 – 7.24 (m, 2H), 7.09 (d, *J* = 8.6 Hz, 1H), 7.01 (d, *J* = 6.7 Hz, 1H), 6.93 (d, *J* = 5.7 Hz, 1H), 6.54 (t, *J* = 6.0 Hz, 1H), 5.04 (dd, *J* = 12.9, 5.4 Hz, 1H), 4.79 (d, *J* = 13.6 Hz, 1H), 4.69 (d, *J* = 13.7 Hz, 1H), 4.04 (t, *J* = 6.2 Hz, 2H), 3.29 – 3.23 (m, 2H), 3.08 (q, *J* = 6.5 Hz, 2H), 2.88 (ddd, *J* = 17.4, 14.0, 5.4 Hz, 1H), 2.62 – 2.51 (m, 2H), 2.25 (t, *J* = 7.3 Hz, 2H), 2.15 (s, 3H), 2.02 (ddd, *J* = 10.8, 5.5, 3.0 Hz, 1H), 1.95 (p, *J* = 6.9 Hz, 2H), 1.54 (p, *J* = 7.1 Hz, 2H), 1.50 – 1.40 (m, 2H) ppm.

**$^{13}$C NMR** (101 MHz, DMSO-$d_6$) δ 176.1, 174.5, 173.4, 172.2, 170.6, 166.1, 157.9, 153.5, 151.4, 149.7, 139.5, 135.5, 126.3, 125.2, 120.5, 119.8, 113.7, 112.3, 109.7, 70.8, 63.4, 51.8, 44.8, 41.4, 34.9, 34.3, 29.8, 29.5, 27.9, 25.4, 14.0 ppm. **Note:** C1 and C2 are not detected due to benz-imidazole tautomerism.

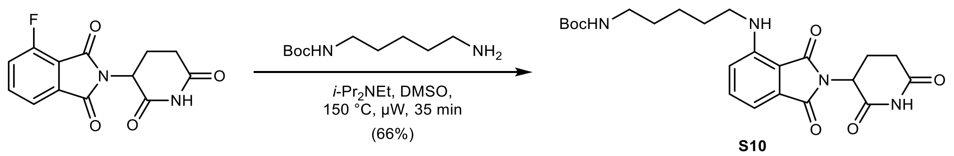

**S10**

***tert*-Butyl (5-((2-(2,6-dioxopiperidin-3-yl)-1,3-dioxoisoindolin-4-yl)amino)pentyl)carbamate (S10):** To a solution of 2-(2,6-dioxopiperidin-3-yl)-4-fluoroisoindoline-1,3-dione (400 mg, 1.45 mmol, 1.00 equiv.) in anhydrous DMSO (7.5 mL) was added *tert*-butyl (5-aminopentyl) carbamate (380 mg, 1.88 mmol, 1.30 equiv.) and *i*-$Pr_2NEt$ (0.74 mL, 4.34 mmol, 3.00 equiv.) at rt under argon. The reaction was stirred at 150 °C for 35 min under microwave irradiation. The mixture was diluted with water (200 mL) and extracted with EtOAc (3 × 50 mL). The combined organic layers were washed with water (50 mL) and brine (50 mL), dried with $MgSO_4$, filtered, and concentrated *in vacuo*. Purification by column chromatography (40 g silica, 30 to 70% EtOAc in *n*-heptane in 14 min) yielded 439 mg (0.96 mmol, 66%) of **S10** as a yellow solid.

**TLC** $R_f$ = 0.05 (30% EtOAc in *n*-heptane).

**LC/MS** (ESI) *m/z*: $(M+Na)^+$ 481.2.

**$^1$H NMR** (400 MHz, $CDCl_3$) δ 7.97 (bs, 1H), 7.49 (ddd, *J* = 8.5, 7.1, 0.6 Hz, 1H), 7.09 (dd, *J* = 7.1, 0.6 Hz, 1H), 6.88 (d, *J* = 8.5 Hz, 1H), 6.23 (t, *J* = 5.6 Hz, 1H), 4.91 (dd, *J* = 12.1, 5.4 Hz, 1H), 4.53 (bs, 1H), 3.27 (q, *J* = 6.7 Hz, 2H), 3.19 – 3.06 (m, 2H), 2.94 – 2.66 (m, 3H), 2.19 – 2.09 (m, 1H), 1.69 (p, *J* = 7.2 Hz, 2H), 1.55 – 1.50 (m, 2H), 1.44 (s, 11H) ppm.

**$^{13}$C NMR** (101 MHz, $CDCl_3$) δ 171.0, 169.6, 168.4, 167.7, 156.1, 147.1, 136.3, 132.6, 116.8, 111.6, 110.1, 79.3, 49.0, 42.7, 40.5, 31.6, 30.0, 29.1, 28.6, 24.3, 23.0 ppm.

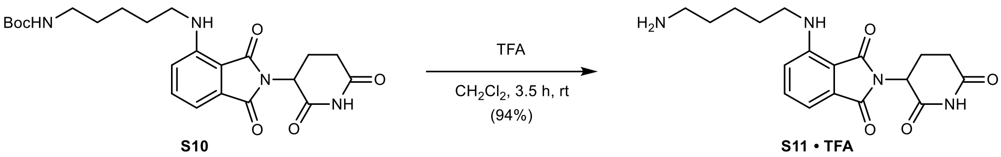

**4-((5-Aminopentyl)amino)-2-(2,6-dioxopiperidin-3-yl)isoindoline-1,3-dione (S11 · TFA):** To a solution of **S10** (440 mg, 0.96 mmol, 1.00 equiv.) in $CH_2Cl_2$ (9.0 mL) was added trifluoroacetic acid (4.4 mL, 57.4 mmol, 60.0 equiv.) slowly at 0 °C. After 5 min, the reaction was allowed to warm to rt and stirred for 3.5 h. The mixture was concentrated *in vacuo* – residual trifluoroacetic acid was removed by co-evaporation with MeOH (3 × 5 mL) and drying *in vacuo*, yielding 434 mg (0.90 mmol, 94%) of **S11 · TFA** as a yellow solid.

**TLC** $R_f$ = 0.63, streaky (30% MeOH in $CH_2Cl_2$).

**LC/MS** (ESI) *m/z*: $(M+H)^+$ 358.2.

$^1$**H NMR** (400 MHz, DMSO-$d_6$) δ 11.10 (s, 1H), 7.78 (bs, 3H), 7.58 (dd, *J* = 8.6, 7.1 Hz, 1H), 7.10 (d, *J* = 8.5 Hz, 1H), 7.03 (dd, *J* = 7.1, 0.6 Hz, 1H), 6.55 (t, *J* = 5.9 Hz, 1H), 5.05 (dd, *J* = 12.8, 5.4 Hz, 1H), 3.30 (q, *J* = 6.7 Hz, 2H), 2.95 – 2.73 (m, 3H), 2.66 – 2.53 (m, 2H), 2.09 – 1.97 (m, 1H), 1.65 – 1.52 (m, 4H), 1.45 – 1.32 (m, 2H) ppm.

$^{13}$**C NMR** (101 MHz, DMSO-$d_6$) δ 172.8, 170.1, 168.9, 167.3, 146.4, 136.3, 132.2, 117.2, 110.5, 109.1, 48.5, 41.6, 38.7, 31.0, 28.2, 26.7, 23.2, 22.2 ppm.

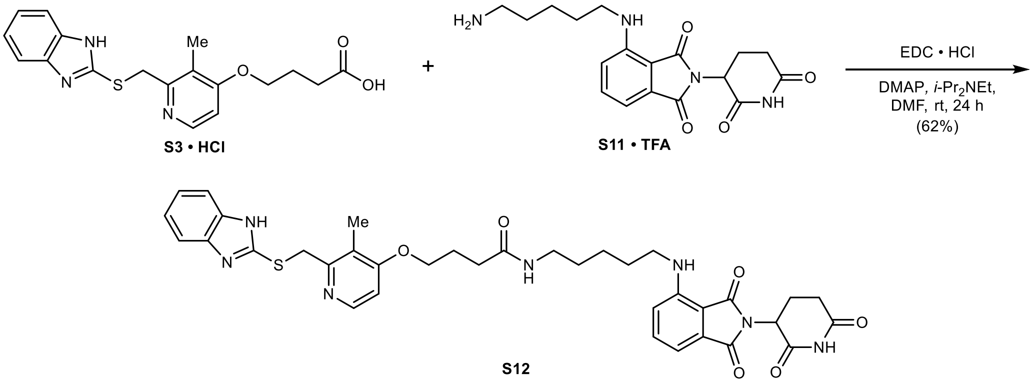

**4-((2-(((1*H*-Benzo[*d*]imidazol-2-yl)thio)methyl)-3-methylpyridin-4-yl)oxy)-*N*-(5-((2-(2,6-dioxopiperidin-3-yl)-1,3-dioxoisoindolin-4-yl)amino)pentyl)butanamide (S12):** To a suspension of **S3 · HCl** (60 mg, 0.15 mmol, 1.00 equiv.) in anhydrous DMF (1.5 mL) was added EDC · HCl (44 mg, 0.23 mmol, 1.50 equiv.), DMAP (4.0 mg, 0.03 mmol, 0.20 equiv.), and *i*-$Pr_2NEt$ (133 µL, 0.76 mmol, 5.00 equiv.) at rt under argon. After 5 min, **S11 · TFA** (72 mg, 0.15 mmol, 1.00 equiv.) was added and the reaction was stirred at rt for 24 h. The mixture was poured into water (100 mL) and extracted with EtOAc (2 × 25 mL), then with 10% MeOH in $CH_2Cl_2$ (2 × 50 mL). The combined organic layers were washed with saturated aqueous $NaHCO_3$ (25 mL), water (25 mL), and brine (25 mL), dried with $MgSO_4$, filtered, and concentrated *in vacuo*. Purification by column chromatography (24 g silica, 0 to 20% MeOH in $CH_2Cl_2$ in 13 min) yielded 66 mg (0.09 mmol, 62%) of **S12** as a bright-yellow solid.

**TLC** $R_f$ = 0.43 (10% MeOH in $CH_2Cl_2$).

**LC/MS** (ESI) *m/z*: $(M+H)^+$ 698.3.

$^1$**H NMR** (400 MHz, $CDCl_3$) δ 8.67 (s, 1H), 8.31 (d, *J* = 5.8 Hz, 1H), 7.56 – 7.48 (m, 2H), 7.44 (ddd, *J* = 8.5, 7.1, 0.6 Hz, 1H), 7.20 – 7.11 (m, 2H), 7.06 (dd, *J* = 7.1, 0.6 Hz, 1H), 6.81 (d, *J* = 8.5 Hz, 1H), 6.72 (d, *J* = 5.8 Hz, 1H), 6.16 (t, *J* = 5.6 Hz, 1H), 5.93 (t, *J* = 5.8 Hz, 1H), 4.89 (dd, *J* = 12.4, 5.2 Hz,

1H), 4.35 (s, 2H), 4.05 (t, $J$ = 6.0 Hz, 2H), 3.24 (q, $J$ = 6.7 Hz, 2H), 3.18 (q, $J$ = 6.6 Hz, 2H), 2.90 – 2.64 (m, 3H), 2.36 (t, $J$ = 7.1 Hz, 2H), 2.20 (s, 3H), 2.18 – 2.05 (m, 3H), 1.61 (p, $J$ = 7.1 Hz, 2H), 1.50 (p, $J$ = 7.2 Hz, 2H), 1.44 – 1.34 (m, 2H) ppm. **Note:** The benzimidazole NH peak is in exchange with residual water in the spectrum.

**$^{13}$C NMR** (101 MHz, $CDCl_3$) δ 174.7, 174.0, 172.4, 171.4, 170.4, 167.0, 159.3, 154.3, 150.1, 149.7, 139.0, 135.3, 124.7, 123.8, 119.4, 114.3, 112.7, 109.0, 70.4, 51.7, 45.2, 42.1, 37.7, 35.3, 34.2, 32.2, 31.6, 27.5, 27.0, 25.6, 13.5 ppm. **Note:** C1, C2, C3, and C6 are not detected due to benzimidazole tautomerism.

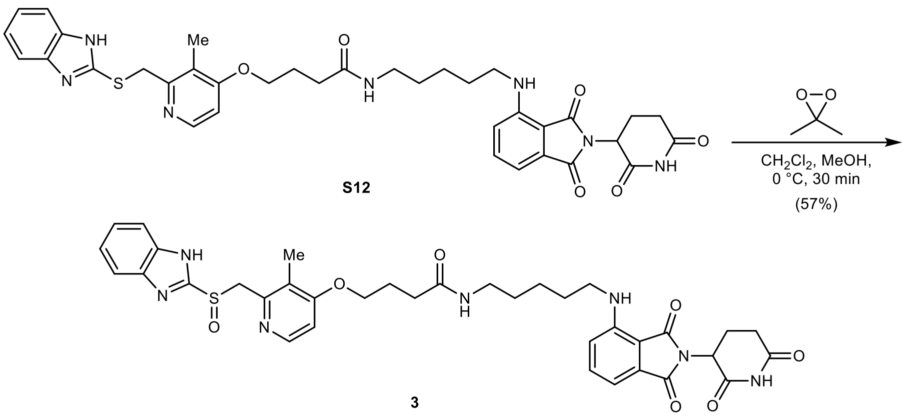

**4-((2-(((1*H*-Benzo[*d*]imidazol-2-yl)sulfinyl)methyl)-3-methylpyridin-4-yl)oxy)-*N*-(5-((2-(2,6-dioxopiperidin-3-yl)-1,3-dioxoisoindolin-4-yl)amino)pentyl)butanamide** (**3**): To a solution of **S12** (37 mg, 0.05 mmol, 1.00 equiv.) in anhydrous $CH_2Cl_2$ (0.5 mL, stored over basic aluminum oxide) and anhydrous MeOH (0.5 mL) was added a solution of 3,3-dimethyldioxirane (0.82 mL, 0.063 M in acetone by iodometric titration, 0.05 mmol, 1.00 equiv.) dropwise at 0 °C under argon. The reaction was stirred for 30 min at 0 °C, then concentrated *in vacuo*. Purification by column chromatography (12 g silica, 0 to 20% MeOH in $CH_2Cl_2$, filtered through basic aluminum oxide, in 10 min) yielded 21 mg (0.03 mmol, 57%) of **3** as a yellow solid.

**TLC** $R_f$ = 0.40 (10% MeOH in $CH_2Cl_2$).

**HRMS** (ESI) *m/z*: $(M+Na)^+$ calc. for $C_{36}H_{39}N_7NaO_7S$: 736.2524; found: 736.2527.

**$^1$H NMR** (400 MHz, DMSO-$d_6$) δ 13.56 (s, 1H), 11.09 (s, 1H), 8.20 (dd, $J$ = 5.6, 0.6 Hz, 1H), 7.84 (t, $J$ = 5.6 Hz, 1H), 7.77 – 7.50 (m, 3H), 7.39 – 7.22 (m, 2H), 7.08 (d, $J$ = 8.6 Hz, 1H), 7.01 (dd, $J$ = 7.1, 0.5 Hz, 1H), 6.94 (d, $J$ = 5.7 Hz, 1H), 6.52 (t, $J$ = 5.9 Hz, 1H), 5.04 (dd, $J$ = 12.9, 5.4 Hz, 1H), 4.79 (d, $J$ = 13.6 Hz, 1H), 4.70 (d, $J$ = 13.7 Hz, 1H), 4.04 (t, $J$ = 6.2 Hz, 2H), 3.26 (q, $J$ = 6.7 Hz, 2H), 3.05 (q, $J$ = 6.5 Hz, 2H), 2.87 (ddd, $J$ = 17.4, 14.1, 5.4 Hz, 1H), 2.63 – 2.52 (m, 2H), 2.24 (t, $J$ = 7.3 Hz, 2H), 2.15 (s, 3H), 2.07 – 1.90 (m, 3H), 1.56 (p, $J$ = 7.3 Hz, 2H), 1.43 (p, $J$ = 7.0 Hz, 2H), 1.32 (q, $J$ = 8.1 Hz, 2H) ppm.

**$^{13}$C NMR** (101 MHz, DMSO-$d_6$) δ 172.8, 171.2, 170.1, 168.9, 167.3, 162.8, 154.3, 150.1, 148.1, 146.4, 136.3, 132.2, 121.9, 117.2, 110.4, 109.0, 106.4, 67.5, 60.2, 48.5, 41.8, 38.3, 31.6, 31.0, 28.8, 28.4, 24.6, 23.7, 22.1, 10.7 ppm. **Note:** C1, C2, C3, C4, C5, and C6 are not detected due to benzimidazole tautomerism.

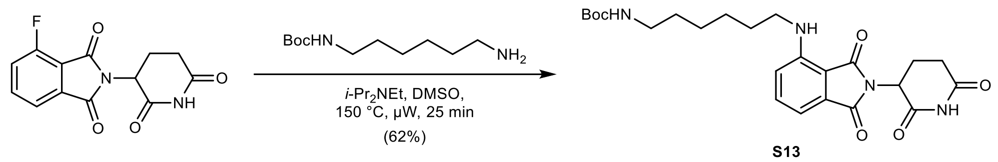

***tert*-Butyl (6-((2-(2,6-dioxopiperidin-3-yl)-1,3-dioxoisoindolin-4-yl)amino)hexyl)carbamate (S13):** To a solution of 2-(2,6-dioxopiperidin-3-yl)-4-fluoroisoindoline-1,3-dione (400 mg, 1.45 mmol, 1.00 equiv.) in anhydrous DMSO (7.5 mL) was added *tert*-butyl (6-amino-hexyl)carbamate (360 mg, 1.67 mmol, 1.15 equiv.) and *i*-$Pr_2NEt$ (757 µL, 4.34 mmol, 3.00 equiv.) at rt under argon. The reaction was stirred at 150 °C for 25 min under microwave irradiation. The mixture was diluted with water (200 mL) and extracted with EtOAc (3 × 50 mL). The combined organic layers were washed with water (50 mL) and brine (50 mL), dried with $MgSO_4$, filtered, and concentrated *in vacuo*. Purification by column chromatography (40 g silica, 30 to 70% EtOAc in *n*-heptane in 14 min) yielded 425 mg (0.90 mmol, 62%) of **S13** as a yellow solid.

**TLC** $R_f$ = 0.34 (5% MeOH in $CH_2Cl_2$).

**LC/MS** (ESI) *m/z*: $(M+Na)^+$ 495.2.

**$^1$H NMR** (400 MHz, $CDCl_3$) δ 7.98 (s, 1H), 7.49 (dd, *J* = 8.5, 7.1 Hz, 1H), 7.09 (dd, *J* = 7.1, 0.6 Hz, 1H), 6.88 (dd, *J* = 8.5, 0.7 Hz, 1H), 6.22 (s, 1H), 4.96 – 4.87 (m, 1H), 4.51 (s, 1H), 3.26 (t, *J* = 7.0 Hz, 2H), 3.11 (d, *J* = 6.4 Hz, 2H), 2.94 – 2.68 (m, 3H), 2.19 – 2.09 (m, 1H), 1.67 (p, *J* = 7.2 Hz, 2H), 1.56 (m, 2H), 1.50 (m, 2H), 1.44 (s, 9H), 1.41 – 1.32 (m, 2H) ppm.

**$^{13}$C NMR** (101 MHz, $CDCl_3$) δ 171.0, 169.7, 168.4, 167.8, 156.1, 147.1, 136.3, 132.6, 116.8, 111.6, 110.0, 79.3, 49.0, 42.7, 40.6, 31.6, 30.2, 29.3, 28.6, 26.8, 26.6, 23.0 ppm.

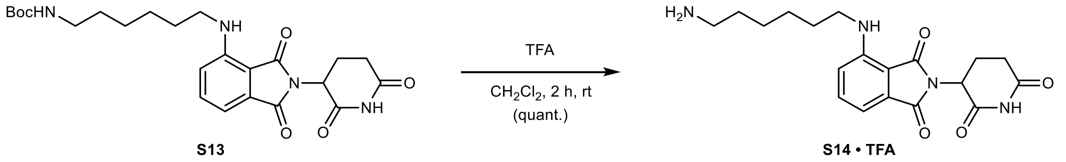

**4-((6-Aminohexyl)amino)-2-(2,6-dioxopiperidin-3-yl)isoindoline-1,3-dione (S14 · TFA):** To a solution of **S13** (425 mg, 0.90 mmol, 1.00 equiv.) in $CH_2Cl_2$ (8.0 mL) was added trifluoroacetic acid (4.1 mL, 54.0 mmol, 60.0 equiv.) slowly at 0 °C. After 5 min, the reaction was allowed to warm to rt and stirred for 2 h. The mixture was concentrated *in vacuo* – residual trifluoroacetic acid was removed by co-evaporation with MeOH (3 × 5 mL) and drying *in vacuo*, yielding 441 mg (0.90 mmol, quantitative) of **S14 · TFA** as a yellow solid.

**TLC** $R_f$ = 0.50, streaky (20% MeOH with 0.5% $NH_4OH$ in $CH_2Cl_2$).

**LC/MS** (ESI) *m/z*: $(M+H)^+$ 373.1.

**$^1$H NMR** (400 MHz, $CD_3OD$) δ 7.55 (dd, *J* = 8.5, 7.2 Hz, 1H), 7.08 – 7.02 (m, 2H), 5.06 (dd, *J* = 12.5, 5.5 Hz, 1H), 3.35 (t, *J* = 6.9 Hz, 2H), 2.96 – 2.80 (m, 3H), 2.80 – 2.64 (m, 2H), 2.16 – 2.05 (m, 1H), 1.76 – 1.62 (m, 4H), 1.49 (m, 4H) ppm. **Note:** The NH peaks are in exchange with $CD_3OD$ in the spectrum.

**$^{13}$C NMR** (101 MHz, $CD_3OD$) δ 174.6, 171.7, 170.9, 169.3, 148.3, 137.3, 134.0, 118.0, 111.8, 111.1, 50.2, 43.2, 40.7, 32.2, 30.0, 28.5, 27.4, 27.2, 23.8 ppm.

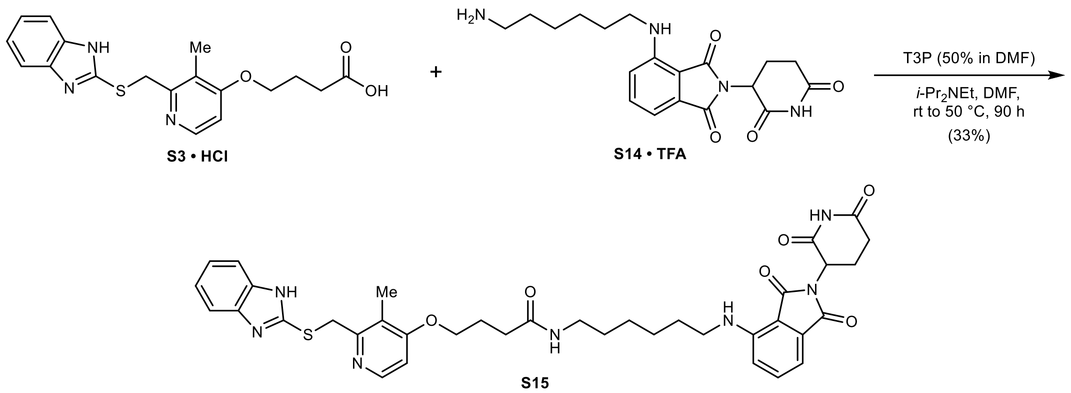

**4-((2-(((1*H*-Benzo[*d*]imidazol-2-yl)thio)methyl)-3-methylpyridin-4-yl)oxy)-*N*-(6-((2-(2,6-dioxopiperidin-3-yl)-1,3-dioxoisoindolin-4-yl)amino)hexyl)butanamide (S15):** To a suspension of **S3 · HCl** (60 mg, 0.15 mmol, 1.00 equiv.) in anhydrous DMF (1.5 mL) was added **S14 · TFA** (78 mg, 0.16 mmol, 1.05 equiv.), *i*-$Pr_2$NEt (106 µL, 0.61 mmol, 4.00 equiv.), and T3P (0.13 mL, 50% in DMF, 0.23 mmol, 1.50 equiv.) at 0 °C under argon. After 5 min, the mixture was allowed to warm to rt and stirred for 20 h. To drive the reaction to completion, T3P (45 µL, 50% in DMF, 0.08 mmol, 0.50 equiv.) and *i*-$Pr_2$NEt (27 µL, 0.15 mmol, 1.00 equiv.) were added and the reaction was stirred at rt for 2 h. The reaction was warmed to 50 °C and stirred for 68 h. The mixture was poured into water (100 mL) and extracted with EtOAc (2 × 25 mL), then with 10% MeOH in $CH_2Cl_2$ (2 × 50 mL). The combined organic layers were washed with saturated aqueous $NaHCO_3$ (25 mL), water (25 mL), and brine (25 mL), dried with $MgSO_4$, filtered, and concentrated *in vacuo*. Purification by column chromatography (12 g silica, 0 to 20% MeOH in $CH_2Cl_2$ in 10 min) yielded 36 mg (0.05 mmol, 33%) of **S15** as a bright-yellow solid. **Note:** HATU appears to be superior to T3P-mediated amide coupling.

**TLC** $R_f$ = 0.41 (10% MeOH in $CH_2Cl_2$).

**LC/MS** (ESI) *m/z*: $(M+H)^+$ 712.2.

**$^1$H NMR** (400 MHz, $CDCl_3$) δ 8.79 (s, 1H), 8.29 (d, *J* = 5.8 Hz, 1H), 7.55 – 7.48 (m, 2H), 7.44 (dd, *J* = 8.5, 7.1 Hz, 1H), 7.20 – 7.11 (m, 2H), 7.05 (d, *J* = 7.1 Hz, 1H), 6.81 (d, *J* = 8.5 Hz, 1H), 6.72 (d, *J* = 5.8 Hz, 1H), 6.17 (t, *J* = 5.6 Hz, 1H), 5.93 (t, *J* = 5.9 Hz, 1H), 4.90 (dd, *J* = 11.9, 5.3 Hz, 1H), 4.37 (s, 2H), 4.05 (t, *J* = 6.0 Hz, 2H), 3.27 – 3.13 (m, 4H), 2.90 – 2.64 (m, 3H), 2.36 (t, *J* = 7.1 Hz, 2H), 2.20 (s, 3H), 2.17 – 2.06 (m, 3H), 1.56 (q, *J* = 7.2 Hz, 2H), 1.46 (p, *J* = 7.2 Hz, 2H), 1.42 – 1.27 (m, 4H) ppm. **Note:** The benzimidazole NH peak is in exchange with residual water in the spectrum.

**$^{13}$C NMR** (101 MHz, $CDCl_3$) δ 174.7, 174.1, 172.4, 171.5, 170.4, 167.1, 159.0, 154.1, 149.9, 149.7, 142.3, 138.9, 135.3, 124.7, 124.0, 119.5, 117.1, 114.2, 112.7, 109.0, 70.5, 51.7, 45.3, 42.3, 37.6, 35.2, 34.3, 32.3, 31.8, 29.4, 27.5, 25.6, 13.5 ppm.

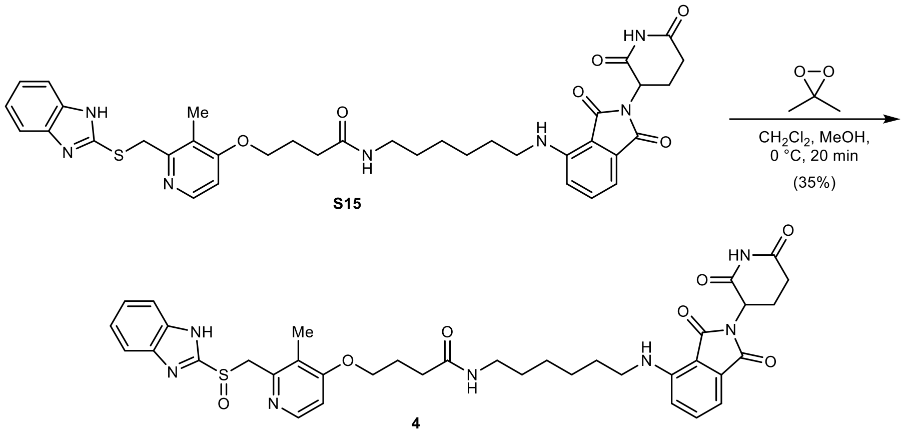

**4-((2-(((1*H*-Benzo[*d*]imidazol-2-yl)sulfinyl)methyl)-3-methylpyridin-4-yl)oxy)-*N*-(6-((2-(2,6-dioxopiperidin-3-yl)-1,3-dioxoisoindolin-4-yl)amino)hexyl)butanamide** (**4**): To a solution of **S15** (36 mg, 0.05 mmol, 1.00 equiv.) in anhydrous $CH_2Cl_2$ (0.5 mL) and anhydrous MeOH (0.5 mL) was added a solution of 3,3-dimethyldioxirane (0.8 mL, 0.063 M in acetone by iodometric titration, 0.05 mmol, 1.00 equiv.) dropwise at 0 °C under argon. The reaction was stirred for 20 min at 0 °C, then concentrated *in vacuo*. Purification by column chromatography (4 g silica, 0 to 20% MeOH in $CH_2Cl_2$, stored over KOH, in 10 min), followed by column chromatography (4 g silica, 5 to 15% MeOH in $CH_2Cl_2$ in 10 min) yielded 13 mg (0.02 mmol, 35%) of **4** as a yellow solid.

**TLC** $R_f$ = 0.46 (10% MeOH in $CH_2Cl_2$).

**HRMS** (ESI) *m/z*: $(M+H)^+$ calc. for $C_{37}H_{42}N_7O_7S$: 728.2861; found: 728.2850.

**$^1$H NMR** (400 MHz, DMSO-$d_6$) δ 11.09 (s, 1H), 8.22 (dd, *J* = 5.7, 0.7 Hz, 1H), 7.83 (t, *J* = 5.6 Hz, 1H), 7.63 – 7.54 (m, 3H), 7.20 – 7.14 (m, 2H), 7.08 (d, *J* = 8.6 Hz, 1H), 7.01 (dd, *J* = 7.1, 0.6 Hz, 1H), 6.92 (d, *J* = 5.7 Hz, 1H), 6.52 (t, *J* = 5.9 Hz, 1H), 5.05 (dd, *J* = 12.9, 5.4 Hz, 1H), 4.80 (d, *J* = 13.4 Hz, 1H), 4.61 (d, *J* = 13.4 Hz, 1H), 4.04 (t, *J* = 6.2 Hz, 2H), 3.28 (p, *J* = 6.7 Hz, 2H), 3.03 (q, *J* = 6.5 Hz, 2H), 2.88 (ddd, *J* = 17.4, 14.1, 5.5 Hz, 1H), 2.63 – 2.52 (m, 2H), 2.24 (t, *J* = 7.3 Hz, 2H), 2.16 (s, 3H), 2.07 – 1.91 (m, 3H), 1.59 – 1.49 (m, 2H), 1.45 – 1.25 (m, 6H) ppm. **Note:** The benzimidazole NH peak is in exchange with residual water in the spectrum.

**$^{13}$C NMR** (101 MHz, DMSO-$d_6$) δ 172.8, 171.2, 170.1, 168.9, 167.3, 162.7, 156.7, 150.9, 148.0, 146.4, 141.1, 136.3, 132.2, 121.9, 121.7, 117.2, 116.5, 110.4, 109.0, 106.2, 67.5, 59.8, 48.5, 41.8, 38.4, 31.6, 31.0, 29.1, 28.6, 26.1, 26.0, 24.6, 22.1, 10.8 ppm.

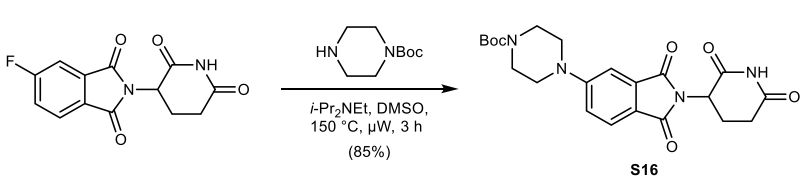

***tert*-Butyl 4-(2-(2,6-dioxopiperidin-3-yl)-1,3-dioxoisoindolin-5-yl)piperazine-1-carboxylate (S16):** To a solution of 2-(2,6-dioxopiperidin-3-yl)-5-fluoroisoindoline-1,3-dione (750 mg, 2.72 mmol, 1.00 equiv.) in anhydrous DMSO (10 mL) was added *tert*-butyl piperazine-1-carboxylate (556 mg, 2.99 mmol, 1.10 equiv.) and *i*-$Pr_2NEt$ (0.95 mL, 5.43 mmol, 2.00 equiv.) at rt under argon. The reaction was stirred at 150 °C for 1.5 h under microwave irradiation. To drive the reaction to completion, *tert*-butyl piperazine-1-carboxylate (50.5 mg, 0.27 mmol, 0.10 equiv.) was added. The reaction was stirred at 150 °C for another 1.5 h under microwave irradiation. The mixture was diluted with water (200 mL) and extracted with EtOAc (3 × 50 mL). The combined organic layers were washed with water (100 mL) and brine (100 mL), dried with $MgSO_4$, filtered, and concentrated *in vacuo*. Purification by column chromatography (80 g silica, 0 to 6% MeOH in

$CH_2Cl_2$ in 12 min) yielded 1.01 g (2.29 mmol, 85%) of **S16** as a brown-yellow solid. **Note**: Ensuring consumption of the imide starting material greatly facilitates purification of the product due to their having similar polarities. This was found to be the case for most reactions of this type.

**TLC** $R_f$ = 0.51 (10% MeOH in $CH_2Cl_2$).

**LC/MS** (ESI) *m/z*: $(M+Na)^+$ 465.1.

**$^1$H NMR** (600 MHz, $CDCl_3$) δ 8.21 (s, 1H), 7.71 (d, *J* = 8.5 Hz, 1H), 7.28 (d, *J* = 2.3 Hz, 1H), 7.05 (dd, *J* = 8.5, 2.4 Hz, 1H), 4.94 (dd, *J* = 12.5, 5.4 Hz, 1H), 3.60 (t, *J* = 5.3 Hz, 4H), 3.41 (t, *J* = 5.3 Hz, 4H), 2.92 – 2.68 (m, 3H), 2.17 – 2.09 (m, 1H), 1.48 (s, 9H) ppm.

**$^{13}$C NMR** (101 MHz, $CDCl_3$) δ 171.1, 168.4, 167.9, 167.3, 155.4, 154.7, 134.4, 125.6, 120.1, 118.3, 109.0, 80.5, 49.3, 47.5, 43.2, 31.6, 28.5, 22.9 ppm.

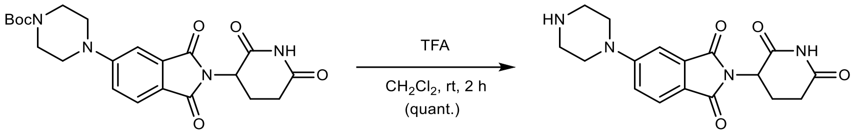

16 S17 • TFA

**2-(2,6-Dioxopiperidin-3-yl)-5-(piperazin-1-yl)isoindoline-1,3-dione (S17 · TFA):** To a solution of **S16** (1.18 g, 2.29 mmol, 1.00 equiv.) in $CH_2Cl_2$ (10 mL) was added trifluoroacetic acid (5.0 mL, 65.4 mmol, 28.5 equiv.) slowly at 0 °C. After 5 min, the reaction was allowed to warm to rt and stirred for 2 h. The mixture was concentrated *in vacuo* – residual trifluoroacetic acid was removed by co-evaporation with MeOH (2 × 5 mL) and toluene (2 × 5 mL) and drying *in vacuo*, yielding 1.04 g (2.29 mmol, quantitative) of **S17 · TFA** as a yellow powder.

**TLC** $R_f$ = 0.21 (10% MeOH in $CH_2Cl_2$).

**LC/MS** (ESI) *m/z*: $(M+H)^+$ 343.1.

**$^1$H NMR** (600 MHz, DMSO-$d_6$) δ 11.10 (s, 1H), 8.99 (s, 2H), 7.75 (d, *J* = 8.5 Hz, 1H), 7.46 (d, *J* = 2.3 Hz, 1H), 7.33 (dd, *J* = 8.6, 2.4 Hz, 1H), 5.09 (dd, *J* = 12.9, 5.5 Hz, 1H), 3.67 (t, *J* = 5.2 Hz, 4H), 3.32 (s, 4H), 2.98 – 2.80 (m, 1H), 2.66 – 2.51 (m, 2H), 2.14 – 1.95 (m, 1H) ppm.

**$^{13}$C NMR** (151 MHz, $CD_3OD$) δ 174.6, 171.6, 169.0, 168.7, 156.2, 135.5, 126.1, 122.8, 120.6, 110.5, 50.5, 46.1, 44.3, 32.2, 23.7 ppm.

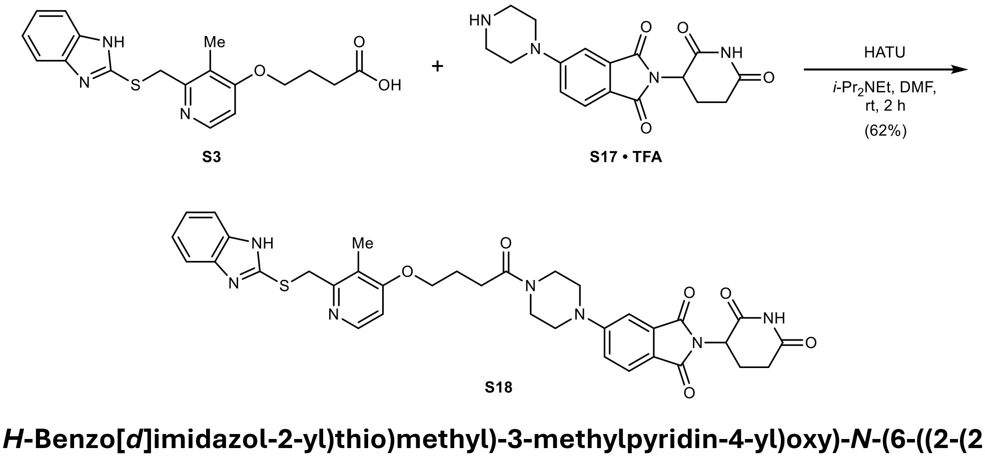

**4-((2-(((1*H*-Benzo[*d*]imidazol-2-yl)thio)methyl)-3-methylpyridin-4-yl)oxy)-*N*-(6-((2-(2,6-dioxopiperidin-3-yl)-1,3-dioxoisoindolin-5-yl)amino)hexyl)butanamide (S18):** To a suspension of **S3** (39 mg, 0.11 mmol, 1.00 equiv.) in anhydrous DMF (1.0 mL) was added **S17 · TFA** (52 mg, 0.11 mmol, 1.05 equiv.), HATU (50 mg, 0.13 mmol, 1.20 equiv.), and *i*-$Pr_2NEt$ (57 µL, 0.33 mmol,

3.00 equiv.) at rt under argon. The reaction was stirred at rt for 2 h. The mixture was poured into water (100 mL) and extracted with EtOAc (3 × 25 mL). The combined organic layers were washed with saturated aqueous $NaHCO_3$ (25 mL), saturated aqueous $NH_4Cl$ (25 mL), water (25 mL), and brine (25 mL), dried with $MgSO_4$, filtered, and concentrated *in vacuo*. Purification by column chromatography (12 g silica, 0 to 10% MeOH in $CH_2Cl_2$ in 10 min) yielded 49 mg (0.07 mmol, 62%) of **S18** as a bright-yellow solid.

**TLC** $R_f$ = 0.46 (10% MeOH in $CH_2Cl_2$).

**LC/MS** (ESI) *m/z*: $(M+H)^+$ 682.2.

**$^1$H NMR** (400 MHz, DMSO-$d_6$) δ 12.61 (s, 1H), 11.08 (s, 1H), 8.24 (d, *J* = 5.6 Hz, 1H), 7.70 (d, *J* = 8.5 Hz, 1H), 7.45 (s, 2H), 7.34 (d, *J* = 2.3 Hz, 1H), 7.23 (dd, *J* = 8.6, 2.3 Hz, 1H), 7.16 – 7.08 (m, 2H), 6.97 (d, *J* = 5.7 Hz, 1H), 5.07 (dd, *J* = 12.9, 5.4 Hz, 1H), 4.70 (s, 2H), 4.11 (t, *J* = 6.3 Hz, 2H), 3.66 – 3.59 (m, 4H), 3.55 – 3.44 (m, 4H), 2.95 – 2.81 (m, 1H), 2.63 – 2.52 (m, 4H), 2.23 (s, 3H), 2.05 – 1.97 (m, 3H) ppm.

**$^{13}$C NMR** (101 MHz, DMSO-$d_6$) δ 172.8, 170.3, 170.1, 167.5, 167.0, 162.7, 154.8, 154.7, 150.2, 147.7, 133.8, 124.9, 121.4, 119.8, 118.4, 117.7, 107.9, 106.3, 67.5, 54.9, 48.8, 46.6, 46.5, 44.0, 36.2, 31.0, 28.5, 24.1, 22.2, 10.5 ppm. **Note:** C1, C2, C3, and C6 are not detected due to benzimidazole tautomerism.

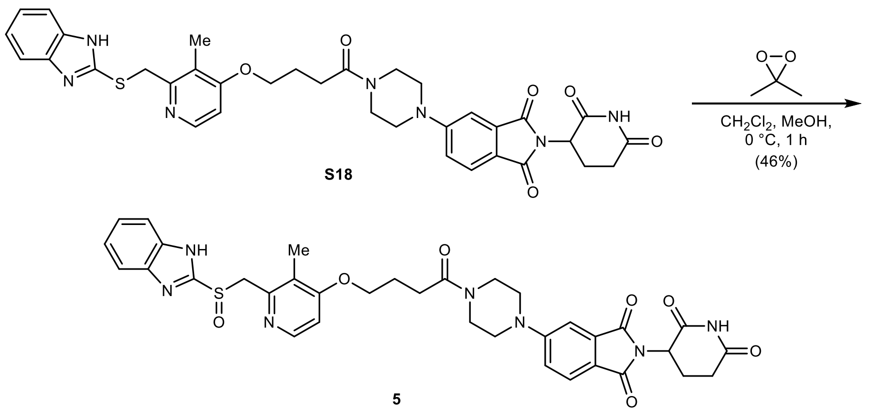

**5-(4-(4-((2-(((1*H*-Benzo[*d*]imidazol-2-yl)sulfinyl)methyl)-3-methylpyridin-4-yl)oxy)butanoyl) piperazin-1-yl)-2-(2,6-dioxopiperidin-3-yl)isoindoline-1,3-dione (5):** To a solution of **S18** (38 mg, 0.06 mmol, 1.00 equiv.) in anhydrous $CH_2Cl_2$ (1.0 mL) and anhydrous MeOH (0.5 mL) was added a solution of 3,3-dimethyldioxirane (1.10 mL, 0.053 M in acetone by iodometric titration, 0.06 mmol, 1.00 equiv.) dropwise at 0 °C under argon. The reaction was stirred for 1 h at 0 °C, then concentrated *in vacuo*. Purification by reversed-phase column chromatography (15.5 g C18, 0 to 100% MeCN in $H_2O$ with 0.1% $NH_4OH$ in 12 min) yielded 18 mg (0.03 mmol, 46%) of **5** as a yellow solid.

**TLC** $R_f$ = 0.62 (10% MeOH in $CH_2Cl_2$).

**HRMS** (ESI) *m/z*: $(M+H)^+$ calc. for $C_{35}H_{36}N_7O_7S$: 698.2391; found: 698.2392.

**$^1$H NMR** (400 MHz, DMSO-$d_6$) δ 13.56 (s, 1H), 11.08 (s, 1H), 8.21 (d, *J* = 5.6 Hz, 1H), 7.80 – 7.53 (m, 3H), 7.34 (d, *J* = 2.3 Hz, 1H), 7.33 – 7.26 (m, 2H), 7.24 (dd, *J* = 8.7, 2.3 Hz, 1H), 6.97 (d, *J* = 5.7 Hz, 1H), 5.07 (dd, *J* = 12.9, 5.4 Hz, 1H), 4.80 (d, *J* = 13.6 Hz, 1H), 4.70 (d, *J* = 13.7 Hz, 1H), 4.11 (t, *J* = 6.3 Hz, 2H), 3.69 – 3.59 (m, 4H), 3.55 – 3.43 (m, 4H), 2.88 (ddd, *J* = 17.3, 14.0, 5.5 Hz, 1H), 2.63 – 2.52 (m, 4H), 2.15 (s, 3H), 2.05 – 1.92 (m, 3H) ppm.

**$^{13}$C NMR** (101 MHz, DMSO-$d_6$) δ 176.1, 173.6, 173.3, 170.8, 170.2, 166.1, 158.1, 157.6, 153.4, 151.4, 137.1, 128.2, 126.6, 125.1, 121.7, 121.0, 115.7, 111.2, 109.7, 70.8, 63.5, 52.1, 49.9, 49.8,

47.3, 43.7, 34.2, 31.8, 27.4, 25.4, 14.0 ppm. **Note:** C1 and C2 are not detected due to benzimidazole tautomerism.

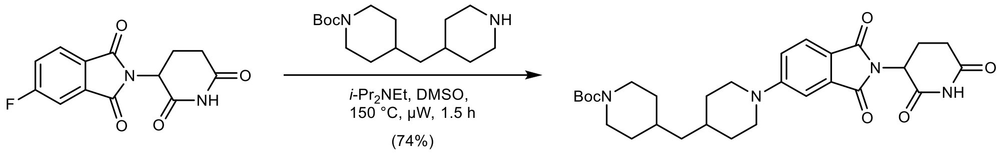

S19

***tert*-Butyl 4-((1-(2-(2,6-dioxopiperidin-3-yl)-1,3-dioxoisoindolin-5-yl)piperidin-4-yl)methyl) piperidine-1-carboxylate (S19):** To a solution of 2-(2,6-dioxopiperidin-3-yl)-5-fluoroisoindoline-1,3-dione (100 mg, 0.36 mmol, 1.00 equiv.) in anhydrous DMSO (3.0 mL) was added *tert*-butyl 4-(piperidin-4-ylmethyl)piperidine-1-carboxylate (112 mg, 0.40 mmol, 1.10 equiv.) and *i*-$Pr_2NEt$ (94 µL, 1.09 mmol, 3.00 equiv.) at rt under argon. The reaction was stirred at 150 °C for 40 min under microwave irradiation. To drive the reaction to completion, *i*-$Pr_2NEt$ (31 µL, 0.36 mmol, 1.00 equiv.) was added. The reaction was stirred at 150 °C for another 50 min under microwave irradiation. The mixture was diluted with water (100 mL) and extracted with EtOAc (3 × 25 mL). The combined organic layers were washed with water (50 mL) and brine (50 mL), dried with $MgSO_4$, filtered, and concentrated *in vacuo*. Purification by column chromatography (24 g silica, 40 to 70% EtOAc in *n*-heptane in 14 min), followed by reversed-phase column chromatography (15.5 g C18, 30 to 100% MeCN in $H_2O$ in 12 min) yielded 144 mg (0.27 mmol, 74%) of **S19** as a bright-yellow solid.

**TLC** $R_f$ = 0.50 (10% MeOH in $CH_2Cl_2$).

**LC/MS** (ESI) *m/z*: $(M+Na)^+$ 561.1.

**$^1$H NMR** (400 MHz, $CDCl_3$) δ 7.98 (s, 1H), 7.66 (d, *J* = 8.6 Hz, 1H), 7.29 – 7.27 (m, 1H), 7.03 (dd, *J* = 8.6, 2.4 Hz, 1H), 4.93 (dd, *J* = 12.3, 5.3 Hz, 1H), 4.24 – 4.00 (m, 2H), 3.94 (d, *J* = 12.9 Hz, 2H), 3.06 – 2.62 (m, 7H), 2.19 – 2.07 (m, 1H), 1.83 – 1.75 (m, 2H), 1.70 – 1.59 (m, 3H), 1.57 – 1.49 (m, 1H), 1.46 (s, 9H), 1.34 – 1.14 (m, 4H), 1.08 (qd, *J* = 12.4, 4.3 Hz, 2H) ppm.

**$^{13}$C NMR** (151 MHz, $CDCl_3$) δ 168.4, 168.2, 167.4, 155.5, 155.0, 134.5, 125.6, 118.6, 117.8, 108.7, 79.4, 49.2, 48.3, 44.4, 43.5, 32.7, 32.5, 32.5, 31.9, 31.6, 28.6, 28.6, 22.9 ppm.

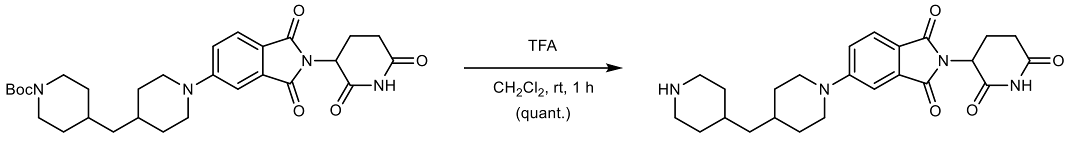

S19 S20 • TFA

**2-(2,6-Dioxopiperidin-3-yl)-5-(4-(piperidin-4-ylmethyl)piperidin-1-yl)isoindoline-1,3-dione (S20 · TFA):** To a solution of **S19** (140 mg, 0.26 mmol, 1.00 equiv.) in $CH_2Cl_2$ (1.0 mL) was added trifluoroacetic acid (1.0 mL, 13.3 mmol, 51.0 equiv.) slowly at 0 °C. After 5 min, the reaction was allowed to warm to rt and stirred for 1 h. The mixture was concentrated *in vacuo* – residual trifluoroacetic acid was removed by co-evaporation with MeOH (3 × 5 mL) and drying *in vacuo*, yielding 144 mg (0.26 mmol, quantitative) of **S20 · TFA** as a yellow, sticky solid.

**TLC** $R_f$ = 0.17, streaky (40% MeOH with 0.5% $NH_4OH$ in $CH_2Cl_2$).

**LC/MS** (ESI) *m/z*: $(M+H)^+$ 439.2.

**$^1$H NMR** (400 MHz, $CD_3OD$) δ 7.66 (d, *J* = 8.6 Hz, 1H), 7.33 (d, *J* = 2.3 Hz, 1H), 7.20 (dd, *J* = 8.7, 2.4 Hz, 1H), 5.06 (dd, *J* = 12.5, 5.5 Hz, 1H), 4.12 – 4.00 (m, 2H), 3.42 – 3.33 (m, 2H), 3.04 – 2.93 (m,

4H), 2.92 – 2.63 (m, 3H), 2.15 – 2.05 (m, 1H), 2.00 – 1.92 (m, 2H), 1.90 – 1.80 (m, 2H), 1.80 – 1.64 (m, 2H), 1.45 – 1.20 (m, 6H) ppm. **Note:** The NH peaks are in exchange with $CD_3OD$ in the spectrum.

**$^{13}$C NMR** (101 MHz, DMSO-$d_6$) δ 172.8, 170.1, 167.6, 167.0, 155.0, 134.0, 125.0, 117.6, 117.4, 107.7, 48.7, 47.4, 43.3, 42.3, 31.7, 31.2, 31.0, 29.7, 28.7, 22.2 ppm.

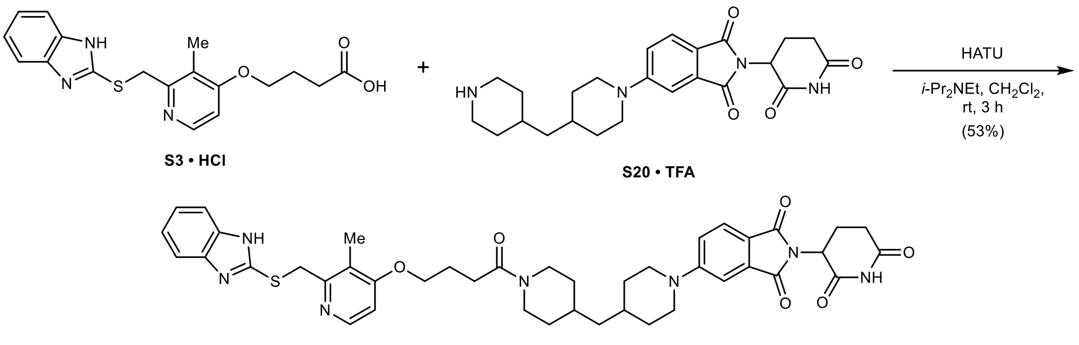

**5-(4-((1-(4-((2-(((1*H*-Benzo[*d*]imidazol-2-yl)thio)methyl)-3-methylpyridin-4-yl)oxy)butanoyl) piperidin-4-yl)methyl)piperidin-1-yl)-2-(2,6-dioxopiperidin-3-yl)isoindoline-1,3-dione (6-S):** To a suspension of **S3 · HCl** (40 mg, 0.10 mmol, 1.00 equiv.) in anhydrous $CH_2Cl_2$ (1.5 mL) was added **S20 · TFA** (56 mg, 0.10 mmol, 1.00 equiv.), HATU (46 mg, 0.12 mmol, 1.20 equiv.), and *i*-$Pr_2NEt$ (71 μL, 0.41 mmol, 4.00 equiv.) at rt under argon. The reaction was stirred at rt for 3 h. The mixture was poured into water (100 mL) and extracted with EtOAc (3 × 25 mL), then with 10% MeOH in $CH_2Cl_2$ (2 × 50 mL). The combined organic layers were washed with saturated aqueous $NaHCO_3$ (25 mL), saturated aqueous $NH_4Cl$ (3 × 25 mL), water (25 mL), and brine (25 mL), dried with $MgSO_4$, filtered, and concentrated *in vacuo*. Purification by column chromatography (12 g silica, 0 to 10% MeOH in $CH_2Cl_2$ in 13 min) yielded 42 mg (0.05 mmol, 53%) of **6-S** as a yellow solid.

**TLC** $R_f$ = 0.43 (10% MeOH in $CH_2Cl_2$).

**HRMS** (ESI) *m/z*: $(M+H)^+$ calc. for $C_{42}H_{48}N_7O_6S$: 778.3381; found: 778.3382.

**$^1$H NMR** (400 MHz, $CDCl_3$) δ 8.72 (s, 1H), 8.32 (d, *J* = 5.7 Hz, 1H), 7.63 (d, *J* = 8.5 Hz, 1H), 7.54 – 7.47 (m, 2H), 7.23 (d, *J* = 2.3 Hz, 1H), 7.19 – 7.11 (m, 2H), 6.99 (dd, *J* = 8.7, 2.4 Hz, 1H), 6.76 (d, *J* = 5.8 Hz, 1H), 4.92 (dd, *J* = 12.3, 5.5 Hz, 1H), 4.64 – 4.56 (m, 1H), 4.38 (s, 2H), 4.10 (t, *J* = 6.0 Hz, 2H), 3.94 – 3.79 (m, 3H), 3.04 – 2.65 (m, 6H), 2.59 – 2.47 (m, 3H), 2.24 (s, 3H), 2.17 (p, *J* = 6.8 Hz, 2H), 2.13 – 2.05 (m, 1H), 1.77 – 1.65 (m, 4H), 1.58 (ddh, *J* = 10.6, 6.9, 3.4 Hz, 2H), 1.27 – 1.17 (m, 2H), 1.14 (t, *J* = 7.0 Hz, 2H), 1.10 – 0.97 (m, 2H) ppm. **Note:** The benzimidazole NH peak is in exchange with residual water in the spectrum.

**$^{13}$C NMR** (101 MHz, $CDCl_3$) δ 174.2, 172.9, 171.4, 170.9, 170.1, 167.0, 159.2, 158.1, 154.3, 150.2, 137.2, 128.2, 124.6, 123.8, 121.3, 120.5, 116.9, 111.4, 109.1, 70.6, 52.0, 50.9, 48.5, 46.0, 44.9, 37.8, 35.9, 35.5, 35.2, 34.9, 34.6, 34.4, 34.3, 31.9, 27.4, 25.6, 13.6 ppm. **Note:** C1 and C2 are not detected due to benzimidazole tautomerism.

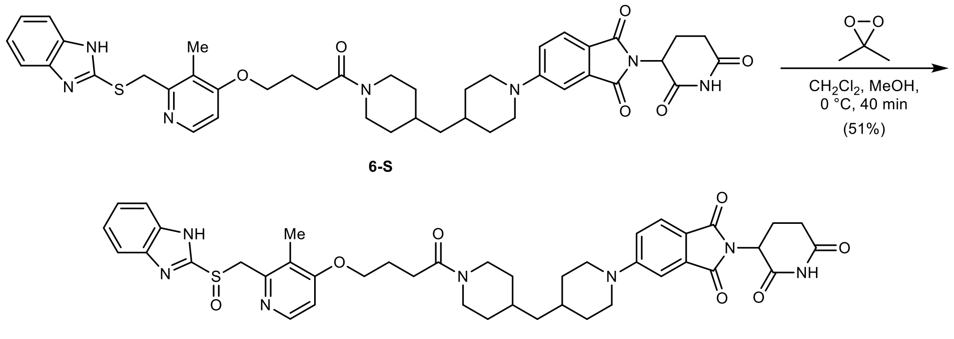

**5-(4-((1-(4-((2-(((1*H*-Benzo[*d*]imidazol-2-yl)thio)methyl)-3-methylpyridin-4-yl)oxy)butanoyl) piperidin-4-yl)methyl)piperidin-1-yl)-2-(2,6-dioxopiperidin-3-yl)isoindoline-1,3-dione (6):** To a solution of **6-S** (42 mg, 0.05 mmol, 1.00 equiv.) in anhydrous $CH_2Cl_2$ (1.0 mL, stored over KOH) and anhydrous MeOH (0.5 mL) was added a solution of 3,3-dimethyldioxirane (0.85 mL, 0.065 M in acetone by iodometric titration, 0.05 mmol, 1.00 equiv.) dropwise at 0 °C under argon. The reaction was stirred for 40 min at 0 °C, then concentrated *in vacuo*. Purification by column chromatography (12 g silica, 0 to 15% MeOH in $CH_2Cl_2$ in 13 min) yielded 22 mg (0.03 mmol, 51%) of **6** as a bright-yellow solid.

**TLC** $R_f$ = 0.45 (10% MeOH in $CH_2Cl_2$).

**HRMS** (ESI) *m/z*: $(M+H)^+$ calc. for $C_{42}H_{48}N_7O_7S$: 794.3330; found: 794.3341.

**$^1$H NMR** (400 MHz, DMSO-*d*$_6$) δ 13.56 (s, 1H), 11.07 (s, 1H), 8.21 (d, *J* = 5.6 Hz, 1H), 7.71 – 7.59 (m, 3H), 7.34 – 7.25 (m, 3H), 7.21 (dd, *J* = 8.8, 2.3 Hz, 1H), 6.96 (d, *J* = 5.7 Hz, 1H), 5.06 (dd, *J* = 12.9, 5.4 Hz, 1H), 4.80 (d, *J* = 13.7 Hz, 1H), 4.70 (d, *J* = 13.7 Hz, 1H), 4.38 (d, *J* = 12.8 Hz, 1H), 4.08 (t, *J* = 6.3 Hz, 2H), 4.03 (d, *J* = 13.0 Hz, 2H), 3.84 (d, *J* = 13.3 Hz, 1H), 3.02 – 2.81 (m, 4H), 2.63 – 2.51 (m, 3H), 2.48 – 2.43 (m, 2H), 2.14 (s, 3H), 2.06 – 1.91 (m, 3H), 1.77 – 1.53 (m, 6H), 1.21 – 1.05 (m, 4H), 1.03 – 0.83 (m, 2H) ppm.

**$^{13}$C NMR** (101 MHz, DMSO-*d*$_6$) δ 172.8, 170.1, 169.5, 167.6, 166.9, 162.8, 155.0, 154.4, 150.1, 148.1, 134.0, 125.0, 123.1, 121.8, 117.5, 117.3, 107.7, 106.4, 67.5, 60.2, 48.7, 47.4, 45.1, 42.8, 41.3, 32.6, 32.0, 31.9, 31.8, 31.3, 31.0, 28.5, 24.3, 22.2, 10.7 ppm. **Note:** C1, C2, C3, and C6 are not detected due to benzimidazole tautomerism.

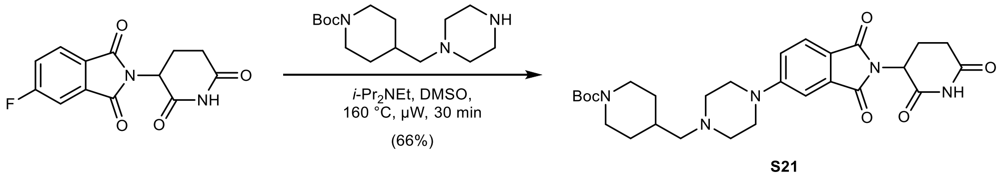

***tert*-Butyl 4-((4-(2-(2,6-dioxopiperidin-3-yl)-1,3-dioxoisoindolin-5-yl)piperazin-1-yl)methyl) piperidine-1-carboxylate (S21):** To a solution of 2-(2,6-dioxopiperidin-3-yl)-5-fluoroisoindoline-1,3-dione (200 mg, 0.72 mmol, 1.00 equiv.) in anhydrous DMSO (5.0 mL) was added *tert*-butyl 4-(piperazin-1-ylmethyl)piperidine-1-carboxylate (226 mg, 0.80 mmol, 1.10 equiv.) and *i*-$Pr_2NEt$ (378 µL, 2.17 mmol, 3.00 equiv.) at rt under argon. The reaction was stirred at 160 °C for 30 min under microwave irradiation. The mixture was diluted with water (200 mL) and extracted with EtOAc (3 × 50 mL). The combined organic layers were washed with water (50 mL) and brine (50 mL), dried with $MgSO_4$, filtered, and concentrated *in vacuo*. Purification by column chromatography (24 g silica, 0 to 15% MeOH in $CH_2Cl_2$ in 14 min) yielded 265 mg (0.48 mmol, 66%) of **S21** as a yellow solid.

**TLC** $R_f$ = 0.40 (10% MeOH in $CH_2Cl_2$).

**LC/MS** (ESI) *m/z*: (M+Na)$^+$ 562.2.

**$^1$H NMR** (400 MHz, $CDCl_3$) δ 8.07 (s, 1H), 7.68 (d, *J* = 8.5 Hz, 1H), 7.28 (d, *J* = 2.3 Hz, 1H), 7.05 (dd, *J* = 8.6, 2.4 Hz, 1H), 4.95 (dd, *J* = 12.2, 5.5 Hz, 1H), 4.30 – 3.94 (m, 2H), 3.46 – 3.38 (m, 4H), 2.94 – 2.65 (m, 5H), 2.60 – 2.52 (m, 4H), 2.23 (d, *J* = 7.1 Hz, 2H), 2.16 – 2.08 (m, 1H), 1.75 (d, *J* = 13.4 Hz, 2H), 1.71 – 1.64 (m, 1H), 1.46 (s, 9H), 1.10 (qd, *J* = 12.3, 4.3 Hz, 2H) ppm.

**$^{13}$C NMR** (101 MHz, $CDCl_3$) δ 173.7, 171.0, 170.8, 170.0, 158.4, 157.7, 137.1, 128.2, 122.2, 120.6, 111.4, 82.1, 67.2, 55.9, 52.0, 50.3, 46.7, 36.4, 34.3, 33.5, 31.3, 25.6 ppm.

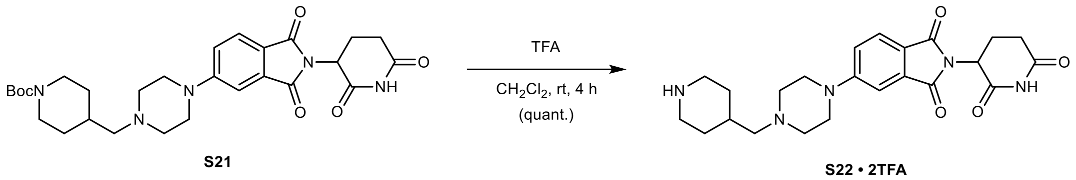

**2-(2,6-Dioxopiperidin-3-yl)-5-(4-(piperidin-4-ylmethyl)piperazin-1-yl)isoindoline-1,3-dione (S22 · 2TFA):** To a solution of **S21** (260 mg, 0.47 mmol, 1.00 equiv.) in $CH_2Cl_2$ (3.0 mL) was added trifluoroacetic acid (1.0 mL, 13.1 mmol, 28.0 equiv.) slowly at 0 °C. After 5 min, the reaction was allowed to warm to rt and stirred for 4 h. The mixture was concentrated *in vacuo* – residual trifluoroacetic acid was removed by co-evaporation with MeOH (3 × 5 mL) and drying *in vacuo*, yielding 313 mg (0.47 mmol, quantitative) of **S22 · 2TFA** as a yellow solid.

**TLC** $R_f$ = 0.09, streaky (50% MeOH with 0.5% $NH_4OH$ in $CH_2Cl_2$).

**LC/MS** (ESI) *m/z*: (M+H)$^+$ 440.2.

**$^1$H NMR** (400 MHz, $CD_3OD$) δ 7.77 (d, *J* = 8.5 Hz, 1H), 7.48 (d, *J* = 2.1 Hz, 1H), 7.36 (dd, *J* = 8.5, 2.4 Hz, 1H), 5.10 (dd, *J* = 12.5, 5.4 Hz, 1H), 3.98 – 3.37 (m, 10H), 3.22 (d, *J* = 6.9 Hz, 2H), 3.06 (td, *J* = 13.0, 2.9 Hz, 2H), 2.94 – 2.80 (m, 1H), 2.80 – 2.64 (m, 2H), 2.37 – 2.25 (m, 1H), 2.17 – 2.06 (m, 3H), 1.64 – 1.48 (m, 2H) ppm. **Note:** The NH peaks are in exchange with $CD_3OD$ in the spectrum.

**$^{13}$C NMR** (101 MHz, $CD_3OD$) δ 174.6, 171.6, 169.0, 168.7, 155.8, 135.6, 126.1, 123.1, 120.6, 110.5, 62.2, 53.1, 50.6, 46.0, 44.4, 32.2, 30.2, 27.7, 23.7 ppm.

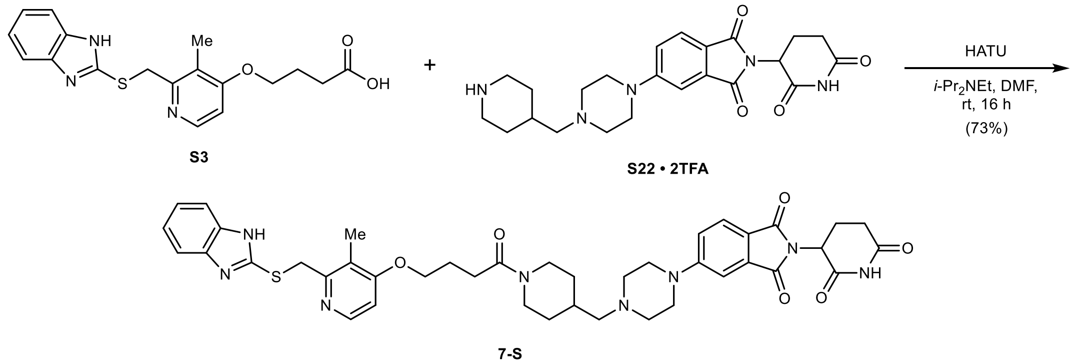

**5-(4-((1-(4-((2-(((1*H*-Benzo[*d*]imidazol-2-yl)thio)methyl)-3-methylpyridin-4-yl)oxy)butanoyl) piperidin-4-yl)methyl)piperazin-1-yl)-2-(2,6-dioxopiperidin-3-yl)isoindoline-1,3-dione (7-S):** To a solution of **S3** (29 mg, 0.08 mmol, 1.00 equiv.) in anhydrous DMF (0.7 mL) was added a solution of **S22 · 2TFA** (57 mg, 0.09 mmol, 1.05 equiv.) in anhydrous DMF (0.5 mL), HATU (37 mg, 0.10 mmol, 1.20 equiv.), and *i*-$Pr_2$NEt (57 µL, 0.33 mmol, 4.00 equiv.) at rt under argon. The reaction was stirred at rt for 16 h. The mixture was poured into water (200 mL) and extracted with 10% MeOH in $CH_2Cl_2$ (3 × 50 mL). The combined organic layers were washed with water (50 mL)

and brine (50 mL), dried with $MgSO_4$, filtered, and concentrated *in vacuo*. Purification by column chromatography (12 g silica, 0 to 12% MeOH in $CH_2Cl_2$ in 12 min) yielded 47 mg (0.05 mmol, 73%) of **7-S** as a yellow powder after lyophilization from MeCN/$H_2O$.

**TLC** $R_f$ = 0.39 (10% MeOH in $CH_2Cl_2$).

**HRMS** (ESI) *m/z*: $(M+H)^+$ calc. for $C_{41}H_{47}N_8O_6S$: 779.3334; found: 779.3335.

**1H NMR** (400 MHz, DMSO-*d*$_6$) δ 12.62 (s, 1H), 11.08 (s, 1H), 8.23 (dd, *J* = 5.7, 0.6 Hz, 1H), 7.66 (d, *J* = 8.5 Hz, 1H), 7.52 (s, 1H), 7.38 (s, 1H), 7.33 (d, *J* = 2.3 Hz, 1H), 7.23 (dd, *J* = 8.7, 2.3 Hz, 1H), 7.16 – 7.08 (m, 2H), 6.96 (d, *J* = 5.8 Hz, 1H), 5.07 (dd, *J* = 12.9, 5.4 Hz, 1H), 4.69 (s, 2H), 4.38 (d, *J* = 12.8 Hz, 1H), 4.08 (t, *J* = 6.3 Hz, 2H), 3.85 (d, *J* = 13.4 Hz, 1H), 3.50 – 3.36 (m, 4H), 2.98 (t, *J* = 12.6 Hz, 1H), 2.88 (ddd, *J* = 17.3, 14.0, 5.5 Hz, 1H), 2.66 – 2.53 (m, 3H), 2.49 – 2.36 (m, 6H), 2.22 (s, 3H), 2.13 (d, *J* = 6.9 Hz, 2H), 2.07 – 1.93 (m, 3H), 1.86 – 1.64 (m, 3H), 1.05 – 0.80 (m, 2H) ppm.

$^{13}$**C NMR** (101 MHz, DMSO-*d*$_6$) δ 172.8, 170.1, 169.6, 167.6, 167.0, 162.7, 155.2, 154.7, 150.3, 147.8, 143.7, 135.4, 133.8, 124.9, 121.6, 121.1, 119.7, 118.3, 117.7, 117.4, 110.4, 107.9, 106.3, 67.5, 63.6, 52.7, 48.8, 46.9, 44.9, 41.1, 36.3, 32.7, 31.0, 30.9, 30.2, 28.5, 24.3, 22.2, 10.5 ppm. **Note:** C1, C2, C3, C4, C5 and C6 each appear as distinct signals in the spectrum.

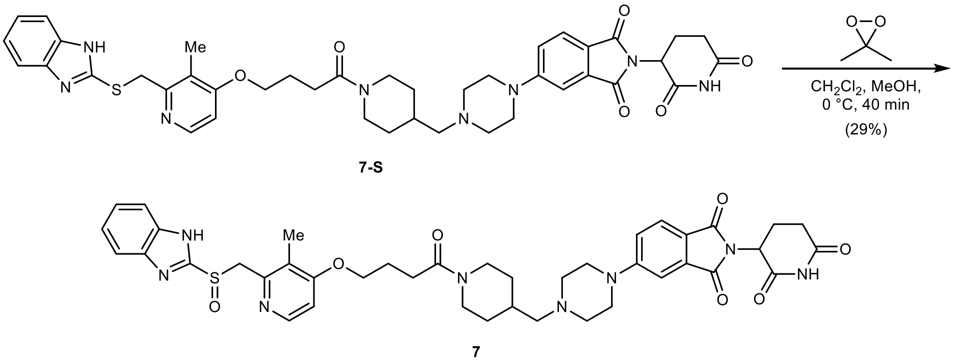

**5-(4-((1-(4-((2-(((1*H*-Benzo[*d*]imidazol-2-yl)sulfinyl)methyl)-3-methylpyridin-4-yl)oxy)buta-noyl)piperidin-4-yl)methyl)piperazin-1-yl)-2-(2,6-dioxopiperidin-3-yl)iso-indoline-1,3-dione (7):** To a solution of **7-S** (37 mg, 0.05 mmol, 1.00 equiv.) in anhydrous $CH_2Cl_2$ (1.0 mL, stored over KOH) and anhydrous MeOH (0.5 mL) was added a solution of 3,3-dimethyldioxirane (0.73 mL, 0.065 M in acetone by iodometric titration, 0.05 mmol, 1.00 equiv.) dropwise at 0 °C under argon. The reaction was stirred for 20 min at 0 °C. To drive the reaction to completion, a solution of 3,3-dimethyldioxirane (0.33 mL, 0.065 M in acetone by iodometric titration, 0.02 mmol, 0.45 equiv.) was added dropwise in two portions over 10 min at 0 °C. The mixture was stirred for 10 min at 0 °C, then concentrated *in vacuo*. Purification by column chromatography (4 g silica, 0 to 20% MeOH in $CH_2Cl_2$ in 12 min) yielded 11 mg (0.01 mmol, 29%) of **7** as a yellow solid. **Note:** *N*-Oxide formation was observed as a major side reaction which consumed 3,3-dimethyldioxirane.

**TLC** $R_f$ = 0.37 (10% MeOH in $CH_2Cl_2$).

**HRMS** (ESI) *m/z*: $(M+H)^+$ calc. for $C_{41}H_{47}N_8O_7S$: 795.3283; found: 795.3290.

$^1$**H NMR** (400 MHz, DMSO-*d*$_6$) δ 11.08 (s, 1H), 8.23 (d, *J* = 5.6 Hz, 1H), 7.67 (d, *J* = 8.5 Hz, 1H), 7.64 – 7.54 (m, 2H), 7.33 (d, *J* = 2.2 Hz, 1H), 7.24 (dd, *J* = 8.7, 2.3 Hz, 1H), 7.20 – 7.14 (m, 2H), 6.95 (d, *J* = 5.7 Hz, 1H), 5.07 (dd, *J* = 12.9, 5.4 Hz, 1H), 4.81 (d, *J* = 13.3 Hz, 1H), 4.61 (d, *J* = 13.4 Hz, 1H), 4.42 – 4.35 (m, 1H), 4.08 (t, *J* = 6.3 Hz, 2H), 3.89 – 3.81 (m, 1H), 3.47 – 3.39 (m, 4H), 2.98 (t, *J* = 12.7 Hz, 1H), 2.88 (ddd, *J* = 17.3, 14.0, 5.5 Hz, 1H), 2.64 – 2.53 (m, 4H), 2.49 – 2.43 (m, 5H), 2.19 – 2.12 (m, 5H), 2.07 – 1.92 (m, 3H), 1.86 – 1.67 (m, 3H), 1.07 – 0.84 (m, 2H) ppm. **Note:** The benzimidazole NH peak is in exchange with residual water in the spectrum.

$^{13}$**C NMR** (101 MHz, DMSO-$d_6$) δ 172.8, 170.1, 169.6, 167.6, 167.0, 162.7, 157.0, 155.2, 150.9, 148.0, 141.3, 133.8, 124.9, 121.8, 121.6, 118.3, 117.7, 116.5, 107.9, 106.3, 67.4, 63.6, 59.8, 52.7, 48.8, 46.9, 44.9, 41.1, 32.7, 31.0, 30.1, 28.5, 24.3, 22.2, 10.8 ppm.

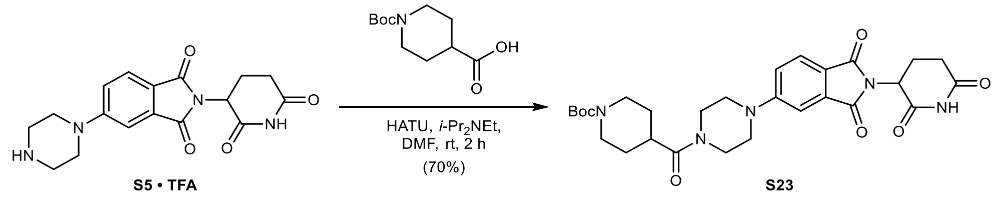

***tert*-Butyl 4-(4-(2-(2,6-dioxopiperidin-3-yl)-1,3-dioxoisoindolin-5-yl)piperazine-1-carbonyl) piperidine-1-carboxylate (S23):** To a solution of 1-(*tert*-butoxycarbonyl) piperidine-4-carboxylic acid (50 mg, 0.22 mmol, 1.00 equiv.) in anhydrous DMF (2.0 mL) was added HATU (10 mg, 0.26 mmol, 1.20 equiv.) and *i*-$Pr_2NEt$ (115 µL, 0.66 mmol, 3.00 equiv.) at rt under argon. After 5 min, **S5 · TFA** (100 mg, 0.22 mmol, 1.00 equiv.) was added and the reaction was stirred at rt for 2 h. The mixture was poured into water (100 mL) and extracted with EtOAc (3 × 25 mL). The combined organic layers were washed with saturated aqueous $NaHCO_3$ (25 mL), saturated aqueous $NH_4Cl$ (25 mL), water (25 mL), and brine (25 mL), dried with $MgSO_4$, filtered, and concentrated *in vacuo*. Purification by column chromatography (12 g silica, 0 to 10% MeOH in $CH_2Cl_2$ in 12 min) yielded 85 mg (0.15 mmol, 70%) of **S23** as a bright-yellow solid.

**TLC** $R_f$ = 0.52 (10% MeOH in $CH_2Cl_2$).

**LC/MS** (ESI) *m/z*: $(M+Na)^+$ 576.1.

$^{1}$**H NMR** (400 MHz, $CDCl_3$) δ 8.05 (s, 1H), 7.73 (d, *J* = 8.5 Hz, 1H), 7.29 (d, *J* = 2.3 Hz, 1H), 7.07 (dd, *J* = 8.5, 2.4 Hz, 1H), 4.95 (dd, *J* = 12.4, 5.3 Hz, 1H), 4.17 (s, 2H), 3.89 – 3.63 (m, 4H), 3.54 – 3.36 (m, 4H), 2.97 – 2.71 (m, 5H), 2.70 – 2.59 (m, 1H), 2.20 – 2.09 (m, 1H), 1.83 – 1.65 (m, 4H), 1.46 (s, 9H) ppm.

$^{13}$**C NMR** (151 MHz, $CDCl_3$) δ 173.3, 171.1, 168.4, 167.8, 167.2, 155.2, 154.8, 134.4, 125.6, 120.7, 118.5, 109.1, 79.9, 49.3, 48.0, 47.6, 44.8, 41.2, 38.7, 31.6, 28.6, 28.6, 28.5, 28.0, 22.8 ppm.

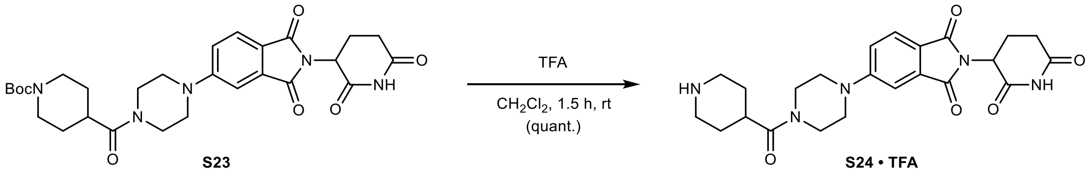

**2-(2,6-Dioxopiperidin-3-yl)-5-(4-(piperidine-4-carbonyl)piperazin-1-yl)isoindoline-1,3-dione (S24 · TFA):** To a solution of **S23** (80 mg, 0.14 mmol, 1.00 equiv.) in $CH_2Cl_2$ (1.0 mL) was added trifluoroacetic acid (1.0 mL, 13.2 mmol, 91.0 equiv.) slowly at 0 °C. After 5 min, the reaction was allowed to warm to rt and stirred for 1.5 h. The mixture was concentrated *in vacuo* – residual trifluoroacetic acid was removed by co-evaporation with MeOH (3 × 5 mL) and drying *in vacuo*, yielding 83 mg (0.15 mmol, quantitative) of **S24 · TFA** as a yellow, sticky solid.

**TLC** $R_f$ = 0.13, streaky (20% MeOH with 0.5% $NH_4OH$ in $CH_2Cl_2$).

**LC/MS** (ESI) *m/z*: $(M+H)^+$ 454.2.

$^{1}$**H NMR** (400 MHz, $CD_3OD$) δ 7.72 (dd, *J* = 8.5, 0.4 Hz, 1H), 7.39 (d, *J* = 2.3 Hz, 1H), 7.26 (dd, *J* = 8.6, 2.4 Hz, 1H), 5.08 (dd, *J* = 12.5, 5.5 Hz, 1H), 3.81 (dt, *J* = 16.6, 5.4 Hz, 4H), 3.53 (dt, *J* = 18.5, 5.3 Hz, 4H), 3.46 (dt, *J* = 12.9, 3.7 Hz, 2H), 3.18 – 3.04 (m, 3H), 2.94 – 2.80 (m, 1H), 2.80 – 2.64 (m, 2H),

2.18 – 2.06 (m, 1H), 2.05 – 1.84 (m, 4H) ppm. **Note:** The NH peaks are in exchange with $CD_3OD$ in the spectrum.

**$^{13}$C NMR** (101 MHz, DMSO-$d_6$) δ 172.8, 171.7, 170.1, 167.5, 167.0, 154.9, 133.8, 124.9, 118.7, 117.9, 108.1, 48.8, 47.1, 46.5, 43.9, 42.5, 40.7, 34.6, 31.0, 25.1, 22.2 ppm.

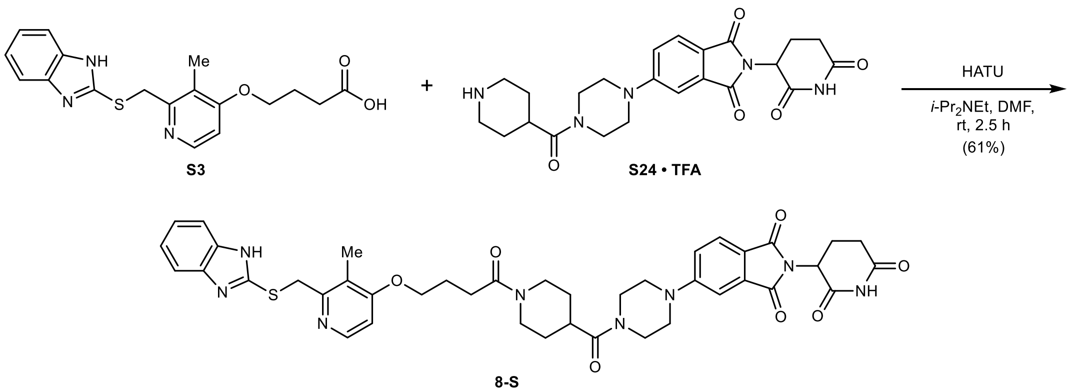

**5-(4-(1-(4-((2-(((1*H*-Benzo[*d*]imidazol-2-yl)thio)methyl)-3-methylpyridin-4-yl)oxy)butanoyl) piperidine-4-carbonyl)piperazin-1-yl)-2-(2,6-dioxopiperidin-3-yl)isoindoline-1,3-dione (8-S):** To a solution of **S3** (33 mg, 0.09 mmol, 1.00 equiv.) in anhydrous DMF (1.0 mL) was added HATU (42 mg, 0.11 mmol, 1.20 equiv.), **S24 · TFA** (55 mg, 0.10 mmol, 1.05 equiv.), and *i*-$Pr_2NEt$ (48.5 µL, 0.28 mmol, 3.00 equiv.) at rt under argon. The reaction was stirred at rt for 2.5 h. The mixture was poured into water (200 mL) and extracted with 5% MeOH in $CH_2Cl_2$ (3 × 50 mL). The combined organic layers were washed with 5% aqueous LiCl (50 mL), water (50 mL), and brine (50 mL), dried with $MgSO_4$, filtered, and concentrated *in vacuo*. Purification by column chromatography (12 g silica, 0 to 11% MeOH with 2.5% $NH_4OH$ in $CH_2Cl_2$ in 12 min) yielded 45 mg (0.06 mmol, 61%) of **8-S** as a yellow powder after lyophilization from MeCN/$H_2O$.

**TLC** $R_f$ = 0.68 (10% MeOH in $CH_2Cl_2$).

**HRMS** (ESI) *m/z*: $(M+H)^+$ calc. for $C_{41}H_{45}N_8O_7S$: 793.3126; found: 793.3126.

**$^1$H NMR** (400 MHz, $CDCl_3$) δ 8.76 (s, 1H), 8.34 (d, *J* = 5.8 Hz, 1H), 7.70 (d, *J* = 8.5 Hz, 1H), 7.57 – 7.48 (m, 2H), 7.26 (d, *J* = 2.3 Hz, 1H), 7.23 – 7.11 (m, 2H), 7.03 (dd, *J* = 8.6, 2.3 Hz, 1H), 6.78 (d, *J* = 5.8 Hz, 1H), 4.95 (dd, *J* = 12.2, 5.4 Hz, 1H), 4.60 (d, *J* = 13.3 Hz, 1H), 4.40 (s, 2H), 4.12 (t, *J* = 6.0 Hz, 2H), 3.94 (d, *J* = 13.5 Hz, 1H), 3.85 – 3.61 (m, 4H), 3.50 – 3.33 (m, 4H), 3.18 – 3.02 (m, 1H), 2.92 – 2.66 (m, 5H), 2.54 (t, *J* = 7.0 Hz, 2H), 2.26 (s, 3H), 2.24 – 2.04 (m, 3H), 1.88 – 1.62 (m, 4H) ppm. **Note:** The benzimidazole NH peak is in exchange with residual water in the spectrum.

**$^{13}$C NMR** (101 MHz, $CDCl_3$) δ 175.6, 174.1, 173.0, 171.3, 170.5, 169.9, 167.0, 159.2, 157.8, 154.4, 150.1, 142.4, 137.1, 128.2, 124.6, 123.8, 123.3, 121.1, 117.0, 111.7, 109.1, 70.6, 52.1, 50.5, 50.2, 47.5, 43.9, 40.9, 37.8, 34.3, 31.9, 31.4, 31.1, 27.3, 25.5, 13.6 ppm.

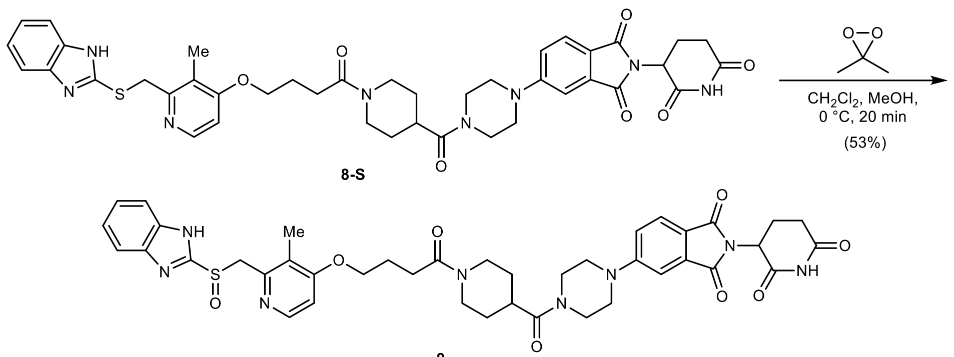

8

**5-(4-(1-(4-((2-(((1*H*-Benzo[*d*]imidazol-2-yl)sulfinyl)methyl)-3-methylpyridin-4-yl)oxy)butanoyl)piperidine-4-carbonyl)piperazin-1-yl)-2-(2,6-dioxopiperidin-3-yl)isoindoline-1,3-dione** (**8**): To a solution of **8-S** (13 mg, 0.02 mmol, 1.00 equiv.) in anhydrous $CH_2Cl_2$ (0.5 mL, stored over KOH) and anhydrous MeOH (0.5 mL) was added a solution of 3,3-dimethyl-dioxirane (0.26 mL, 0.064 M in acetone by iodometric titration, 0.02 mmol, 1.00 equiv.) dropwise at 0 °C under argon. The reaction was stirred for 20 min at 0 °C, then concentrated *in vacuo*. Purification by column chromatography (4 g silica, 0 to 20% MeOH in $CH_2Cl_2$ in 10 min) yielded 7 mg (0.01 mmol, 53%) of **8** as a yellow solid.

**TLC** $R_f$ = 0.36 (10% MeOH in $CH_2Cl_2$).

**HRMS** (ESI) *m/z*: $(M+Na)^+$ calc. for $C_{41}H_{44}N_8NaO_8S$: 831.2895; found: 831.2900.

**$^1$H NMR** (400 MHz, DMSO-$d_6$) δ 11.09 (s, 1H), 8.27 (d, *J* = 5.5 Hz, 1H), 7.70 (dd, *J* = 8.6, 1.2 Hz, 1H), 7.49 – 7.40 (m, 2H), 7.36 (d, *J* = 2.3 Hz, 1H), 7.25 (d, *J* = 8.7 Hz, 1H), 6.93 (d, *J* = 5.7 Hz, 1H), 6.90 – 6.84 (m, 2H), 5.08 (dd, *J* = 12.9, 5.4 Hz, 1H), 4.90 – 4.78 (m, 1H), 4.44 – 4.35 (m, 2H), 4.08 (t, *J* = 6.3 Hz, 2H), 3.92 – 3.85 (m, 1H), 3.77 – 3.67 (m, 2H), 3.67 – 3.57 (m, 2H), 3.57 – 3.43 (m, 5H), 3.08 (t, *J* = 12.7 Hz, 1H), 3.01 – 2.81 (m, 2H), 2.72 – 2.59 (m, 2H), 2.59 – 2.52 (m, 2H), 2.19 (s, 3H), 2.06 – 1.94 (m, 3H), 1.75 – 1.63 (m, 2H), 1.59 – 1.46 (m, 1H), 1.46 – 1.33 (m, 1H) ppm. **Note:** The benzimidazole NH peak is in exchange with residual water in the spectrum.

**$^{13}$C NMR** (101 MHz, DMSO-$d_6$) δ 172.8, 172.5, 170.1, 169.8, 167.5, 167.0, 162.6, 154.9, 152.6, 147.9, 146.5, 133.8, 124.9, 121.8, 118.6, 118.3, 117.9, 117.4, 108.0, 105.9, 67.3, 59.0, 48.8, 47.1, 46.5, 44.2, 43.9, 40.6, 40.5, 37.0, 31.0, 28.7, 28.6, 28.1, 24.3, 22.2, 10.9 ppm. **Note:** C1 and C2 are not detected due to benzimidazole tautomerism.

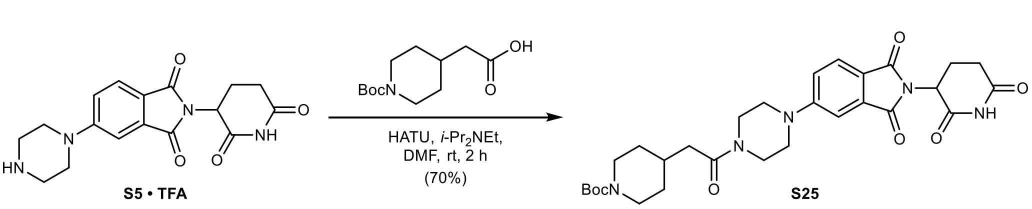

***tert*-Butyl 4-(2-(4-(2-(2,6-dioxopiperidin-3-yl)-1,3-dioxoisoindolin-5-yl)piperazin-1-yl)-2-oxoethyl)piperidine-1-carboxylate (S25):** To a solution of 2-(1-(*tert*-butoxy-carbonyl)piperidin-4-yl) acetic acid (64 mg, 0.16 mmol, 1.00 equiv.) in anhydrous DMF (2.0 mL) was added HATU (120 mg, 0.32 mmol, 1.20 equiv.) and *i*-$Pr_2NEt$ (137 µL, 0.79 mmol, 3.00 equiv.) at rt under argon. After 5 min, **S5 · TFA** (120 mg, 0.26 mmol, 1.00 equiv.) was added and the reaction was stirred at rt for 2 h. The mixture was poured into water (100 mL) and extracted with EtOAc (3 × 25 mL). The combined organic layers were washed with saturated aqueous $NaHCO_3$ (25 mL), saturated aqueous $NH_4Cl$ (25 mL), water (25 mL), and brine (25 mL), dried with $MgSO_4$, filtered, and concentrated *in vacuo*. Purification by column chromatography (12 g silica,

0 to 10% MeOH in $CH_2Cl_2$ in 12 min) yielded 104 mg (0.18 mmol, 70%) of **S25** as a bright-yellow solid.

**TLC** $R_f$ = 0.69 (10% MeOH in $CH_2Cl_2$).

**LC/MS** (ESI) *m/z*: $(M+Na)^+$ 590.3.

**$^1$H NMR** (400 MHz, $CDCl_3$) δ 8.05 (s, 1H), 7.73 (dd, *J* = 8.5, 0.4 Hz, 1H), 7.28 (d, *J* = 2.2 Hz, 1H), 7.06 (dd, *J* = 8.5, 2.4 Hz, 1H), 4.95 (dd, *J* = 12.3, 5.3 Hz, 1H), 4.26 – 3.97 (m, 2H), 3.86 – 3.79 (m, 2H), 3.71 – 3.64 (m, 2H), 3.48 – 3.38 (m, 4H), 2.94 – 2.80 (m, 2H), 2.79 – 2.65 (m, 3H), 2.29 (d, *J* = 6.8 Hz, 2H), 2.19 – 2.10 (m, 1H), 2.10 – 1.96 (m, 1H), 1.79 – 1.71 (m, 2H), 1.45 (s, 9H), 1.15 (ddd, *J* = 12.6, 12.6, 4.3 Hz, 2H) ppm.

**$^{13}$C NMR** (151 MHz, $CDCl_3$) δ 171.1, 170.5, 168.3, 167.8, 167.2, 155.1, 155.0, 134.4, 125.6, 120.6, 118.4, 109.0, 79.5, 53.6, 49.3, 47.7, 47.6, 45.0, 41.0, 39.7, 33.3, 32.4, 31.6, 28.6, 28.6, 22.8 ppm.

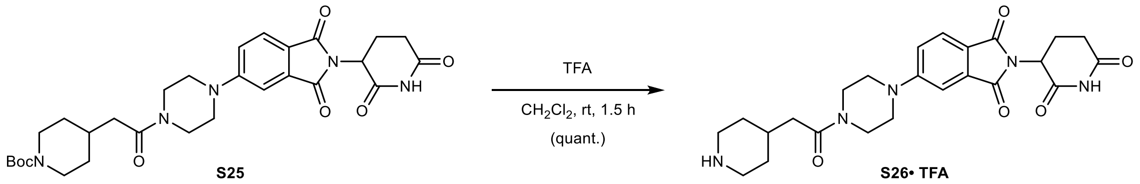

**2-(2,6-Dioxopiperidin-3-yl)-5-(4-(2-(piperidin-4-yl)acetyl)piperazin-1-yl)isoindoline-1,3-dione (S26 · TFA):** To a solution of **S25** (100 mg, 0.18 mmol, 1.00 equiv.) in $CH_2Cl_2$ (1.0 mL) was added trifluoroacetic acid (1.0 mL, 13.2 mmol, 75.0 equiv.) slowly at 0 °C. After 5 min, the reaction was allowed to warm to rt and stirred for 1.5 h. The mixture was concentrated *in vacuo* – residual trifluoroacetic acid was removed by co-evaporation with MeOH (3 × 5 mL) and drying *in vacuo*, yielding 103 mg (0.18 mmol, quantitative) of **S26 · TFA** as a yellow, sticky solid.

**TLC** $R_f$ = 0.04, streaky (20% MeOH with 0.5% $NH_4OH$ in $CH_2Cl_2$).

**LC/MS** (ESI) *m/z*: $(M+H)^+$ 468.2.

**$^1$H NMR** (400 MHz, $CD_3OD$) δ 7.71 (d, *J* = 8.5 Hz, 1H), 7.38 (d, *J* = 2.3 Hz, 1H), 7.25 (dd, *J* = 8.6, 2.4 Hz, 1H), 5.08 (dd, *J* = 12.6, 5.5 Hz, 1H), 3.77 (dt, *J* = 11.6, 5.2 Hz, 4H), 3.58 – 3.47 (m, 4H), 3.43 – 3.36 (m, 2H), 3.08 – 2.96 (m, 2H), 2.93 – 2.82 (m, 1H), 2.79 – 2.64 (m, 2H), 2.47 (d, *J* = 6.8 Hz, 2H), 2.20 – 2.07 (m, 2H), 2.02 (d, *J* = 14.3 Hz, 2H), 1.54 – 1.39 (m, 2H) ppm. **Note:** The NH peaks are in exchange with $CD_3OD$ in the spectrum.

**$^{13}$C NMR** (101 MHz, $CD_3OD$) δ 174.6, 172.0, 171.6, 169.3, 168.9, 156.7, 135.6, 126.0, 121.1, 119.4, 109.4, 50.5, 48.4, 48.1, 46.0, 45.2, 42.3, 39.6, 32.2, 32.0, 29.9, 23.8 ppm.

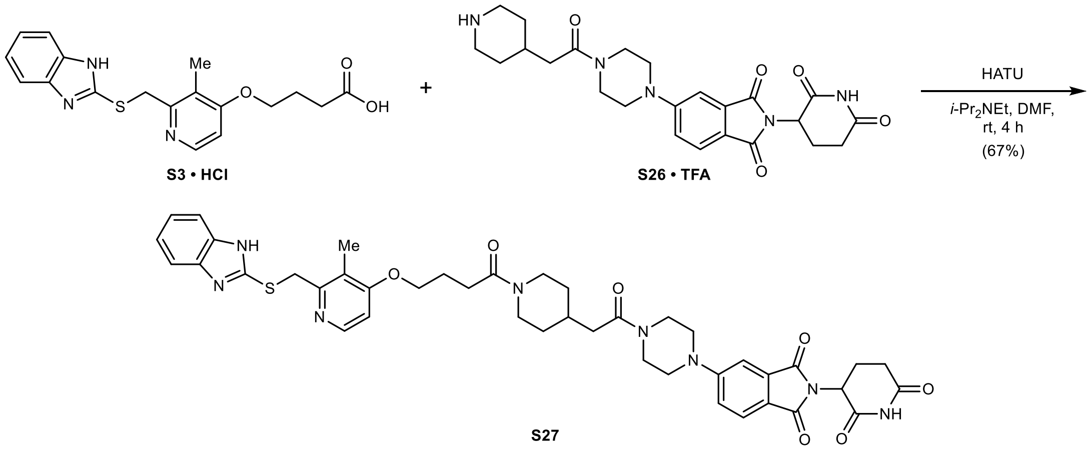

**5-(4-(2-(1-(4-((2-(((1*H*-Benzo[*d*]imidazol-2-yl)thio)methyl)-3-methylpyridin-4-yl)oxy)butanoyl)piperidin-4-yl)acetyl)piperazin-1-yl)-2-(2,6-dioxopiperidin-3-yl)isoindoline-1,3-dione (S27):** To a suspension of **S3 · HCl** (40 mg, 0.10 mmol, 1.00 equiv.) in anhydrous DMF (1.0 mL) was added HATU (46 mg, 0.12 mmol, 1.20 equiv.) and *i*-$Pr_2NEt$ (80 µL, 0.46 mmol, 4.50 equiv.) at rt under argon. After 10 min, **S26 · TFA** (59 mg, 0.10 mmol, 1.00 equiv.) and DMF (0.5 mL) were added and the reaction was stirred at rt for 4 h. The mixture was poured into water (100 mL) and extracted with EtOAc (3 × 25 mL). The combined organic layers were washed with saturated aqueous $NaHCO_3$ (25 mL), saturated aqueous $NH_4Cl$ (25 mL), water (25 mL), and brine (25 mL), dried with $MgSO_4$, filtered, and concentrated *in vacuo*. Purification by column chromatography (12 g silica, 0 to 15% MeOH in $CH_2Cl_2$ in 13 min) yielded 55 mg (0.07 mmol, 67%) of **S27** as a yellow solid.

**TLC** $R_f$ = 0.32 (10% MeOH in $CH_2Cl_2$).

**LC/MS** (ESI) *m/z*: $(M+H)^+$ 807.3.

**$^1$H NMR** (400 MHz, $CDCl_3$) δ 8.34 (dd, *J* = 5.8, 0.6 Hz, 1H), 8.24 (s, 1H), 7.71 (d, *J* = 8.5 Hz, 1H), 7.57 – 7.48 (m, 2H), 7.26 – 7.24 (m, 1H), 7.22 – 7.13 (m, 2H), 7.04 (dd, *J* = 8.5, 2.4 Hz, 1H), 6.80 (d, *J* = 5.9 Hz, 1H), 4.94 (dd, *J* = 12.4, 5.3 Hz, 1H), 4.63 (d, *J* = 13.2 Hz, 1H), 4.38 (s, 2H), 4.13 (t, *J* = 6.0 Hz, 2H), 3.86 (d, *J* = 14.0 Hz, 1H), 3.82 – 3.71 (m, 2H), 3.68 – 3.56 (m, 2H), 3.45 – 3.38 (m, 4H), 3.05 (dt, *J* = 12.4, 2.6 Hz, 1H), 2.96 – 2.67 (m, 3H), 2.64 – 2.46 (m, 3H), 2.34 – 2.09 (m, 9H), 1.87 (d, *J* = 11.3 Hz, 1H), 1.79 (d, *J* = 13.2 Hz, 1H), 1.19 – 1.01 (m, 2H) ppm. **Note:** The benzimidazole NH peak is in exchange with residual water in the spectrum.

**$^{13}$C NMR** (101 MHz, $CDCl_3$) δ 173.7, 172.9, 172.8, 171.0, 170.5, 169.9, 167.2, 159.3, 157.8, 154.4, 150.1, 137.1, 128.3, 124.7, 123.8, 123.3, 121.1, 117.1, 111.6, 109.1, 70.6, 52.1, 50.3, 48.5, 47.6, 44.8, 43.7, 42.1, 37.6, 35.8, 35.6, 34.9, 34.3, 31.8, 27.3, 25.5, 13.6 ppm. **Note:** C1 and C2 are not detected due to benzimidazole tautomerism.

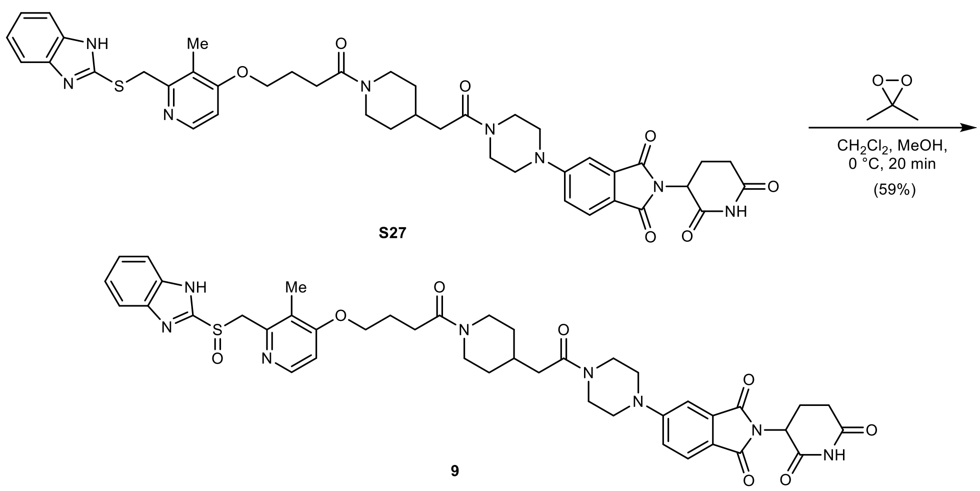

**5-(4-(2-(1-(4-((2-(((1*H*-Benzo[*d*]imidazol-2-yl)sulfinyl)methyl)-3-methylpyridin-4-yl)oxy)butanoyl)piperidin-4-yl)acetyl)piperazin-1-yl)-2-(2,6-dioxopiperidin-3-yl)isoindoline-1,3-dione** (**9**): To a solution of **S27** (29 mg, 0.03 mmol, 1.00 equiv.) in anhydrous $CH_2Cl_2$ (1.0 mL, stored over KOH) and anhydrous MeOH (0.5 mL) was added a solution of 3,3-dimethyldioxirane (0.51 mL, 0.065 M in acetone by iodometric titration, 0.03 mmol, 1.00 equiv.) dropwise at 0 °C under argon. The reaction was stirred for 20 min at 0 °C, then concentrated *in vacuo*. Purification by column chromatography (4 g silica, 0 to 20% MeOH in $CH_2Cl_2$ in 10 min) yielded 16 mg (0.02 mmol, 59%) of **9** as a yellow solid.

**TLC** $R_f$ = 0.27 (10% MeOH in $CH_2Cl_2$).

**HRMS** (ESI) *m/z*: $(M+H)^+$ calc. for $C_{42}H_{46}N_8O_8S$: 823.3232; found: 823.3236.

**$^1$H NMR** (400 MHz, DMSO-*d*$_6$) δ 11.08 (s, 1H), 8.22 (d, *J* = 5.6 Hz, 1H), 7.69 (d, *J* = 8.5 Hz, 1H), 7.66 – 7.57 (m, 2H), 7.34 (d, *J* = 2.3 Hz, 1H), 7.28 – 7.20 (m, 3H), 6.96 (d, *J* = 5.7 Hz, 1H), 5.07 (dd, *J* = 12.9, 5.4 Hz, 1H), 4.80 (d, *J* = 13.5 Hz, 1H), 4.66 (d, *J* = 13.5 Hz, 1H), 4.40 – 4.32 (m, 1H), 4.08 (t, *J* = 6.3 Hz, 2H), 3.88 – 3.80 (m, 1H), 3.65 – 3.58 (m, 4H), 3.52 – 3.43 (m, 4H), 2.99 (t, *J* = 12.3 Hz, 1H), 2.88 (ddd, *J* = 17.3, 14.1, 5.6 Hz, 1H), 2.63 – 2.51 (m, 3H), 2.48 – 2.44 (m, 2H), 2.29 (d, *J* = 6.8 Hz, 2H), 2.15 (s, 3H), 2.06 – 1.91 (m, 4H), 1.69 (t, *J* = 12.7 Hz, 2H), 1.17 – 0.93 (m, 2H) ppm. **Note:** The benzimidazole NH peak is in exchange with residual water in the spectrum.

**$^{13}$C NMR** (101 MHz, DMSO-*d*$_6$) δ 172.8, 170.0, 169.7, 169.5, 167.5, 167.0, 162.8, 155.4, 154.9, 150.5, 148.1, 133.8, 124.9, 122.5, 121.9, 118.5, 117.8, 107.9, 106.4, 67.5, 60.0, 48.8, 46.8, 46.5, 44.9, 44.2, 41.2, 40.1, 38.4, 32.5, 32.2, 31.5, 31.0, 28.5, 24.3, 22.2, 10.7 ppm. **Note:** C1, C2, C3, and C6 are not detected due to benzimidazole tautomerism.

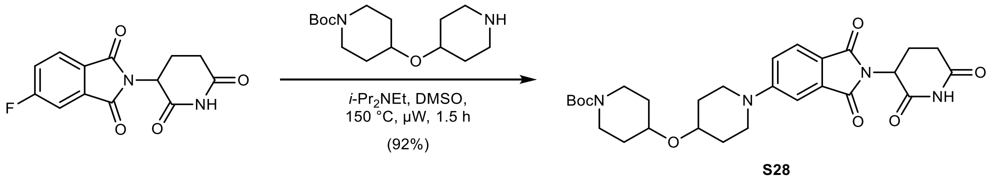

***tert*-Butyl 4-((1-(2-(2,6-dioxopiperidin-3-yl)-1,3-dioxoisoindolin-5-yl)piperidin-4-yl)oxy)piperidine-1-carboxylate (S28):** To a solution of 2-(2,6-dioxopiperidin-3-yl)-5-fluoroisoindoline-1,3-dione (750 mg, 2.72 mmol, 1.00 equiv.) in anhydrous DMSO (13 mL) was added *tert*-butyl 4-(piperidin-4-yloxy)piperidine-1-carboxylate (849 mg, 2.99 mmol, 1.10 equiv.) and *i*-$Pr_2NEt$ (1.18 mL, 6.79 mmol, 2.50 equiv.) at rt under argon. The reaction was stirred at 150 °C for 1.5 h under microwave irradiation. The mixture was diluted with water (300 mL) and extracted with EtOAc (3 × 100 mL). The combined organic layers were washed with water (150 mL) and brine (150 mL), dried with $MgSO_4$, filtered, and concentrated *in vacuo*. Purification by column chromatography (80 g silica, 40 to 80% EtOAc in *n*-heptane in 15 min) yielded 1.35 g (2.50 mmol, 92%) of **S28** as a yellow solid.

**TLC** $R_f$ = 0.56 (75% EtOAc in *n*-heptane).

**LC/MS** (ESI) *m/z*: $(M+Na)^+$ 563.2.

**$^1$H NMR** (400 MHz, $CDCl_3$) δ 8.09 (s, 1H), 7.67 (d, *J* = 8.6 Hz, 1H), 7.28 (d, *J* = 2.4 Hz, 1H), 7.04 (dd, *J* = 8.6, 2.4 Hz, 1H), 4.93 (dd, *J* = 12.3, 5.3 Hz, 1H), 3.86 – 3.66 (m, 5H), 3.61 (tt, *J* = 7.9, 3.7 Hz, 1H), 3.28 (ddd, *J* = 12.7, 8.2, 3.5 Hz, 2H), 3.12 (ddd, *J* = 13.0, 9.0, 3.5 Hz, 2H), 2.94 – 2.65 (m, 3H), 2.18 – 2.07 (m, 1H), 1.99 – 1.86 (m, 2H), 1.86 – 1.75 (m, 2H), 1.75 – 1.65 (m, 2H), 1.59 – 1.48 (m, 2H), 1.45 (s, 9H) ppm.

**$^{13}$C NMR** (101 MHz, $CDCl_3$) δ 171.0, 168.3, 168.0, 167.2, 155.2, 154.9, 134.4, 125.5, 118.8, 117.8, 108.6, 79.5, 71.9, 70.7, 49.1, 45.3, 41.3, 31.8, 31.5, 31.1, 28.5, 22.8 ppm.

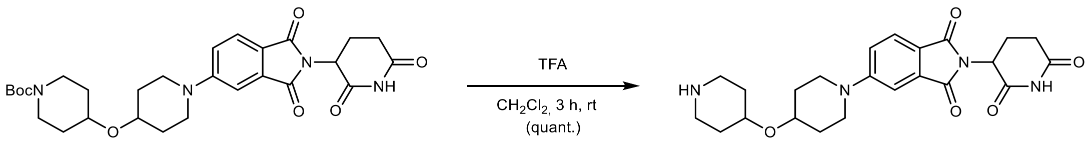

**2-(2,6-Dioxopiperidin-3-yl)-5-(4-(piperidin-4-yloxy)piperidin-1-yl)isoindoline-1,3-dione (S29 · TFA):** To a solution of **S28** (1.35 g, 2.50 mmol, 1.00 equiv.) in $CH_2Cl_2$ (12 mL) was added trifluoroacetic acid (4.0 mL, 52.4 mmol, 20.0 equiv.) slowly at 0 °C. After 5 min, the reaction was allowed to warm to rt and stirred for 3 h. The mixture was concentrated *in vacuo* – residual trifluoroacetic acid was removed by co-evaporation with MeOH (3 × 5 mL), benzene (2 × 5 mL), and drying *in vacuo*, yielding 1.38 g (2.50 mmol, quantitative) of **S29 · TFA** as a yellow solid.

**TLC** $R_f$ = 0.34, streaky (20% MeOH with 0.5% $NH_4OH$ in $CH_2Cl_2$).

**LC/MS** (ESI) *m/z*: $(M+H)^+$ 441.2.

**$^1$H NMR** (400 MHz, $CD_3OD$) δ 7.66 (d, *J* = 8.6 Hz, 1H), 7.34 (d, *J* = 2.3 Hz, 1H), 7.22 (dd, *J* = 8.6, 2.4 Hz, 1H), 5.06 (dd, *J* = 12.5, 5.4 Hz, 1H), 3.88 (tt, *J* = 6.5, 3.2 Hz, 1H), 3.84 – 3.73 (m, 3H), 3.40 – 3.26 (m, 4H), 3.13 (ddd, *J* = 12.7, 6.9, 4.0 Hz, 2H), 2.93 – 2.79 (m, 1H), 2.79 – 2.63 (m, 2H), 2.16 – 2.06 (m, 1H), 2.06 – 1.94 (m, 4H), 1.91 – 1.79 (m, 2H), 1.73 – 1.60 (m, 2H) ppm.

**$^{13}$C NMR** (101 MHz, $CD_3OD$) δ 174.6, 171.7, 169.5, 169.0, 156.8, 135.7, 126.1, 119.9, 119.1, 109.2, 73.2, 68.8, 50.4, 46.3, 42.1, 32.2, 32.1, 29.5, 23.8 ppm.

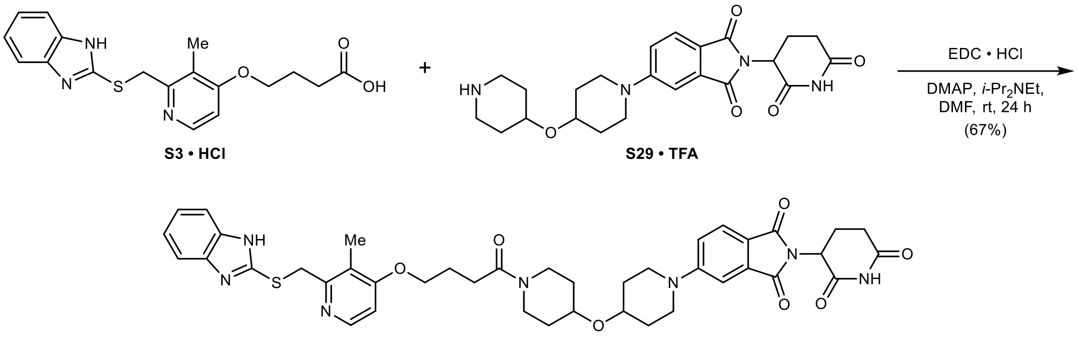

**5-(4-((1-(4-((2-(((1*H*-Benzo[*d*]imidazol-2-yl)thio)methyl)-3-methylpyridin-4-yl)oxy)butanoyl) piperidin-4-yl)oxy)piperidin-1-yl)-2-(2,6-dioxopiperidin-3-yl)isoindoline-1,3-dione (10-S):** To a suspension of **S3 · HCl** (39 mg, 0.10 mmol, 1.00 equiv.) in anhydrous DMF (1.0 mL) was added EDC · HCl (28 mg, 0.15 mmol, 1.50 equiv.), DMAP (2.5 mg, 0.02 mmol, 0.20 equiv.), and *i*-$Pr_2NEt$ (77 µL, 0.44 mmol, 4.50 equiv.) at rt under argon. After 10 min, **S29 · TFA** (55 mg, 0.10 mmol, 1.00 equiv.) was added and the reaction was stirred at rt for 24 h. The mixture was poured into water (100 mL) and extracted with 10% MeOH in $CH_2Cl_2$ (3 × 25 mL). The combined organic layers were washed with saturated aqueous $NaHCO_3$ (25 mL), water (25 mL), and brine (25 mL), dried with $MgSO_4$, filtered, and concentrated *in vacuo*. Purification by column chromatography (12 g silica, 0 to 10% MeOH in $CH_2Cl_2$ in 13 min) yielded 51 mg (0.07 mmol, 67%) of **10-S** as a bright-yellow solid.

**TLC** $R_f$ = 0.49 (10% MeOH in $CH_2Cl_2$).

**HRMS** (ESI) *m/z*: $(M+H)^+$ calc. for $C_{41}H_{46}N_7O_7S$: 780.3174; found: 780.3183.

**$^1$H NMR** (400 MHz, $CDCl_3$) δ 8.34 (d, *J* = 5.7 Hz, 1H), 8.30 (s, 1H), 7.66 (d, *J* = 8.6 Hz, 1H), 7.60 – 7.47 (m, 2H), 7.28 – 7.26 (m, 1H), 7.21 – 7.13 (m, 2H), 7.03 (dd, *J* = 8.6, 2.4 Hz, 1H), 6.78 (d, *J* = 5.8 Hz, 1H), 4.93 (dd, *J* = 12.3, 5.6 Hz, 1H), 4.38 (s, 2H), 4.12 (t, *J* = 6.0 Hz, 2H), 3.89 (ddd, *J* = 12.0, 7.3, 3.8 Hz, 1H), 3.73 – 3.63 (m, 5H), 3.38 (ddd, *J* = 12.6, 8.0, 3.6 Hz, 1H), 3.33 – 3.20 (m, 3H), 2.93 – 2.65 (m, 3H), 2.53 (t, *J* = 7.0 Hz, 2H), 2.26 (s, 3H), 2.19 (p, *J* = 6.3 Hz, 2H), 2.15 – 2.07 (m, 1H), 1.96 – 1.86 (m, 2H), 1.85 – 1.74 (m, 2H), 1.73 – 1.63 (m, 2H), 1.63 – 1.51 (m, 2H) ppm. **Note:** The benzimidazole NH peak is in exchange with residual water in the spectrum.

**$^{13}$C NMR** (101 MHz, $CDCl_3$) δ 171.2, 170.2, 168.5, 168.1, 167.3, 164.3, 156.7, 155.2, 151.7, 147.5, 134.5, 125.6, 121.9, 121.1, 119.0, 118.0, 108.7, 106.4, 71.1, 71.1, 67.9, 49.3, 45.3, 42.7, 39.1, 35.0, 32.4, 31.6, 31.4, 31.2, 31.0, 29.1, 24.6, 22.9, 10.9 ppm. **Note:** C1, C2, C3, and C6 are not detected due to benzimidazole tautomerism.

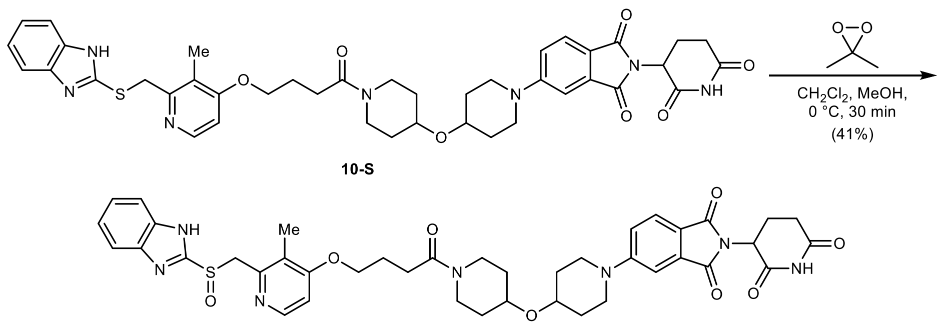

10-S

10

**5-(4-((1-(4-((2-(((1*H*-Benzo[*d*]imidazol-2-yl)sulfinyl)methyl)-3-methylpyridin-4-yl)oxy)butanoyl)piperidin-4-yl)oxy)piperidin-1-yl)-2-(2,6-dioxopiperidin-3-yl)isoindoline-1,3-dione** (**10**): To a solution of **10-S** (51 mg, 0.07 mmol, 1.00 equiv.) in anhydrous $CH_2Cl_2$ (0.5 mL, stored over basic aluminum oxide) and anhydrous MeOH (0.5 mL) was added a solution of 3,3-dimethyldioxirane (1.04 mL, 0.063 M in acetone by iodometric titration, 0.05 mmol, 1.00 equiv.) dropwise at 0 °C under argon. The reaction was stirred for 30 min at 0 °C, then concentrated *in vacuo*. Purification by column chromatography (4 g silica, 0 to 20% MeOH in $CH_2Cl_2$, filtered through basic aluminum oxide, in 12 min) yielded 22 mg (0.03 mmol, 41%) of **10** as a yellow solid.

**TLC** $R_f$ = 0.37 (10% MeOH in $CH_2Cl_2$).

**HRMS** (ESI) *m/z*: $(M+Na)^+$ calc. for $C_{41}H_{45}N_7NaO_8S$: 818.2943; found: 818.2946.

**$^1$H NMR** (400 MHz, DMSO-*d*$_6$) δ 13.56 (s, 1H), 11.07 (s, 1H), 8.21 (dd, *J* = 5.6, 0.6 Hz, 1H), 7.82 – 7.50 (m, 3H), 7.41 – 7.20 (m, 4H), 6.96 (d, *J* = 5.7 Hz, 1H), 5.06 (dd, *J* = 12.9, 5.4 Hz, 1H), 4.79 (d, *J* = 13.6 Hz, 1H), 4.70 (d, *J* = 13.6 Hz, 1H), 4.08 (t, *J* = 6.3 Hz, 2H), 3.92 – 3.84 (m, 1H), 3.84 – 3.60 (m, 5H), 3.29 – 3.14 (m, 3H), 3.14 – 3.04 (m, 1H), 2.88 (ddd, *J* = 17.2, 14.0, 5.5 Hz, 1H), 2.63 – 2.52 (m, 2H), 2.49 – 2.45 (m, 2H), 2.14 (s, 3H), 2.06 – 1.92 (m, 3H), 1.92 – 1.70 (m, 4H), 1.54 – 1.42 (m, 2H), 1.42 – 1.25 (m, 2H) ppm.

**$^{13}$C NMR** (101 MHz, DMSO-*d*$_6$) δ 172.8, 170.1, 169.7, 167.6, 166.9, 162.8, 154.7, 154.3, 150.1, 148.1, 134.0, 125.0, 121.8, 117.6, 117.6, 107.8, 106.4, 70.8, 70.6, 67.5, 60.2, 48.7, 44.9, 42.4, 38.7, 32.1, 31.4, 31.0, 30.8, 28.4, 24.3, 22.2, 10.7 ppm. **Note:** C1, C2, C3, C4, C5, and C6 are not detected due to benzimidazole tautomerism.

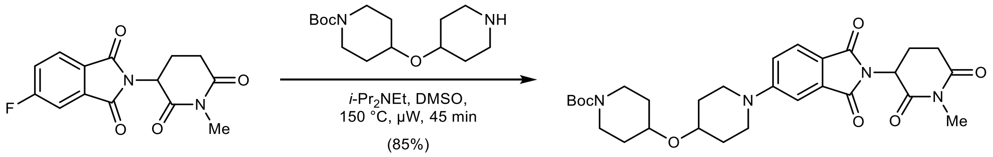

S30

***tert*-Butyl 4-((1-(2-(1-methyl-2,6-dioxopiperidin-3-yl)-1,3-dioxoisoindolin-5-yl)piperidin-4-yl) oxy)piperidine-1-carboxylate (S30):** To a solution of 5-fluoro-2-(1-methyl-2,6-dioxopiperidin-3-yl)isoindoline-1,3-dione (76 mg, 0.26 mmol, 1.00 equiv.)[7] in anhydrous DMSO (2.0 mL) was added *tert*-butyl 4-(piperidin-4-yloxy)piperidine-1-carboxylate (82 mg, 0.29 mmol, 1.10 equiv.) and *i*-$Pr_2NEt$ (137 µL, 0.79 mmol, 3.00 equiv.) at rt under argon. The reaction was stirred at 150 °C for 45 min under microwave irradiation. The mixture was diluted with water (150 mL) and extracted with EtOAc (3 × 50 mL). The combined organic layers were washed with water (100 mL) and brine (100 mL), dried with $MgSO_4$, filtered, and concentrated *in vacuo*. Purification by column chromatography (12 g silica, 40 to 75% EtOAc in *n*-heptane in 12 min) yielded 126 mg (0.22 mmol, 85%) of **S30** as a bright-yellow solid.

**TLC** $R_f$ = 0.52 (75% EtOAc in *n*-heptane).

**LC/MS** (ESI) *m/z*: $(M+Na)^+$ 577.3.

**$^1$H NMR** (400 MHz, $CDCl_3$) δ 7.67 (d, *J* = 8.5 Hz, 1H), 7.28 (d, *J* = 2.3 Hz, 1H), 7.05 (dd, *J* = 8.6, 2.4 Hz, 1H), 4.93 (dd, *J* = 12.6, 5.4 Hz, 1H), 3.82 – 3.66 (m, 5H), 3.61 (tt, *J* = 7.9, 3.6 Hz, 1H), 3.28 (ddd, *J* = 12.6, 8.2, 3.4 Hz, 2H), 3.20 (s, 3H), 3.12 (ddd, *J* = 13.0, 9.0, 3.5 Hz, 2H), 3.04 – 2.88 (m, 1H), 2.87 – 2.67 (m, 2H), 2.14 – 2.06 (m, 1H), 1.97 – 1.86 (m, 2H), 1.85 – 1.75 (m, 2H), 1.70 (dtd, *J* = 11.8, 7.8, 3.6 Hz, 2H), 1.57 – 1.48 (m, 2H), 1.46 (s, 9H) ppm.

**$^{13}$C NMR** (101 MHz, $CDCl_3$) δ 171.4, 169.1, 168.3, 167.6, 155.3, 155.0, 134.6, 125.5, 119.0, 117.9, 108.7, 79.6, 72.0, 70.9, 50.0, 45.4, 41.4, 32.1, 31.9, 31.2, 28.6, 27.4, 22.2 ppm.

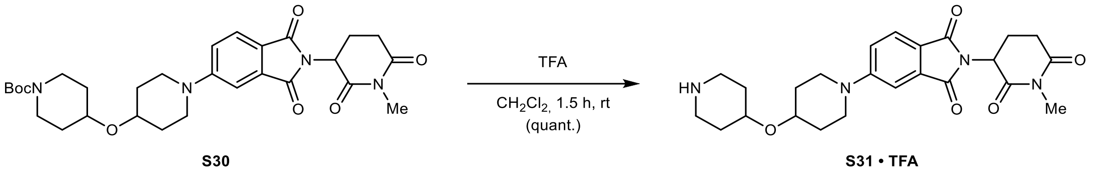

**2-(1-Methyl-2,6-dioxopiperidin-3-yl)-5-(4-(piperidin-4-yloxy)piperidin-1-yl)isoindoline-1,3-dione (S31 · TFA):** To a solution of **S30** (126 mg, 0.23 mmol, 1.00 equiv.) in $CH_2Cl_2$ (2.0 mL) was added trifluoroacetic acid (1.0 mL, 13.1 mmol, 57.5 equiv.) slowly at 0 °C. After 5 min, the reaction was allowed to warm to rt and stirred for 1.5 h. The mixture was concentrated *in vacuo* – residual trifluoroacetic acid was removed by co-evaporation with MeOH (3 × 5 mL), toluene (3 × 5 mL), and drying *in vacuo*, yielding 129 mg (0.23 mmol, quantitative) of **S31 · TFA** as a yellow solid.

**TLC** $R_f$ = 0.52, streaky (20% MeOH with 0.5% $NH_4OH$ in $CH_2Cl_2$).

**LC/MS** (ESI) *m/z*: $(M+H)^+$ 455.2.

**$^1$H NMR** (400 MHz, $CD_3OD$) δ 7.65 (d, *J* = 8.5 Hz, 1H), 7.33 (d, *J* = 2.4 Hz, 1H), 7.21 (dd, *J* = 8.6, 2.4 Hz, 1H), 5.09 (dd, *J* = 12.9, 5.4 Hz, 1H), 3.88 (dt, *J* = 6.6, 3.3 Hz, 1H), 3.84 – 3.69 (m, 3H), 3.42 – 3.33 (m, 2H), 3.29 – 3.25 (m, 2H), 3.21 – 3.08 (m, 5H), 2.99 – 2.81 (m, 2H), 2.76 – 2.61 (m, 1H), 2.14 – 1.92 (m, 5H), 1.91 – 1.79 (m, 2H), 1.73 – 1.60 (m, 2H) ppm. **Note:** The NH peaks are in exchange with $CD_3OD$ in the spectrum.

**$^{13}$C NMR** (101 MHz, $CD_3OD$) δ 173.7, 171.5, 169.5, 169.0, 156.7, 135.7, 126.1, 119.8, 119.1, 109.2, 73.2, 68.8, 51.1, 46.3, 42.1, 32.5, 32.1, 29.4, 27.3, 23.0 ppm.

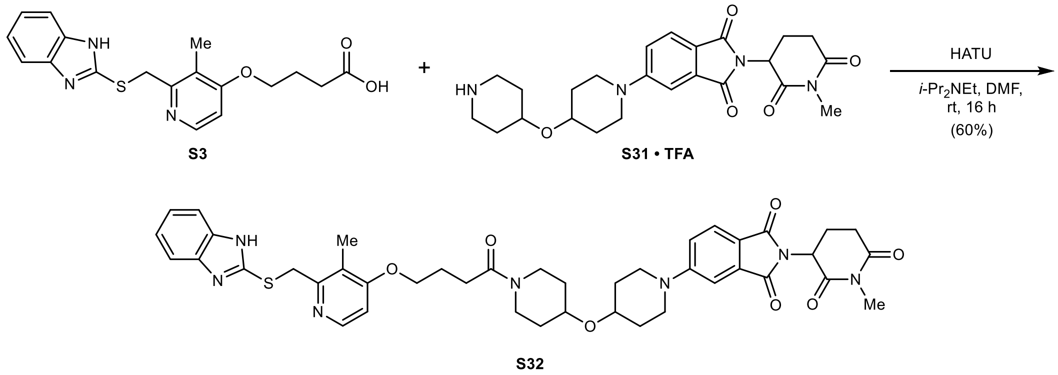

**5-(4-((1-(4-((2-(((1*H*-Benzo[*d*]imidazol-2-yl)thio)methyl)-3-methylpyridin-4-yl)oxy)butanoyl) piperidin-4-yl)oxy)piperidin-1-yl)-2-(1-methyl-2,6-dioxopiperidin-3-yl)isoindoline-1,3-dione (S32):** To a solution of **S31 · TFA** (72 mg, 0.12 mmol, 1.05 equiv.) in anhydrous DMF (1.0 mL) was added **S3** (42 mg, 0.12 mmol, 1.00 equiv.), HATU (54 mg, 0.14 mmol, 1.20 equiv.), and *i*-$Pr_2NEt$

(61 µL, 0.35 mmol, 3.00 equiv.) at rt under argon. The reaction was stirred at rt for 16 h. The mixture was poured into water (100 mL) and extracted with EtOAc (3 × 25 mL). The combined organic layers were washed with 5% aqueous LiCl (25 mL), water (25 mL), and brine (25 mL), dried with $MgSO_4$, filtered, and concentrated *in vacuo*. Purification by column chromatography (12 g silica, 0 to 10% MeOH in $CH_2Cl_2$ in 11 min) yielded 56 mg (0.07 mmol, 60%) of **S32** as a yellow solid.

**TLC** $R_f$ = 0.52 (10% MeOH in $CH_2Cl_2$).

**LC/MS** (ESI) *m/z*: $(M+H)^+$ 794.3.

**$^1$H NMR** (400 MHz, $CDCl_3$) δ 8.35 (dd, *J* = 5.8, 0.6 Hz, 1H), 7.67 (dd, *J* = 8.5, 0.4 Hz, 1H), 7.57 – 7.49 (m, 2H), 7.29 – 7.26 (m, 1H), 7.22 – 7.14 (m, 2H), 7.04 (dd, *J* = 8.6, 2.4 Hz, 1H), 6.80 (d, *J* = 5.8 Hz, 1H), 4.93 (dd, *J* = 12.5, 5.4 Hz, 1H), 4.37 (s, 2H), 4.14 (t, *J* = 6.0 Hz, 2H), 3.95 – 3.86 (m, 1H), 3.77 – 3.63 (m, 5H), 3.44 – 3.33 (m, 1H), 3.34 – 3.23 (m, 3H), 3.21 (s, 3H), 3.02 – 2.91 (m, 1H), 2.87 – 2.67 (m, 2H), 2.54 (t, *J* = 7.0 Hz, 2H), 2.27 (s, 3H), 2.20 (p, *J* = 6.5 Hz, 2H), 2.13 – 2.06 (m, 1H), 1.97 – 1.86 (m, 2H), 1.86 – 1.76 (m, 2H), 1.76 – 1.63 (m, 2H), 1.63 – 1.51 (m, 2H) ppm. **Note:** The benzimidazole NH peak is in exchange with residual water in the spectrum.

**$^{13}$C NMR** (101 MHz, $CDCl_3$) δ 171.4, 170.2, 169.1, 168.3, 167.5, 164.4, 156.7, 155.2, 151.8, 147.4, 134.6, 125.5, 122.0, 121.1, 119.1, 118.0, 114.3, 108.7, 106.4, 71.1, 71.1, 67.9, 50.0, 45.4, 42.7, 39.1, 34.9, 32.5, 32.1, 31.5, 31.2, 31.0, 29.1, 27.4, 24.6, 22.2, 10.9 ppm. **Note:** C1 and C2 are not detected due to benzimidazole tautomerism.

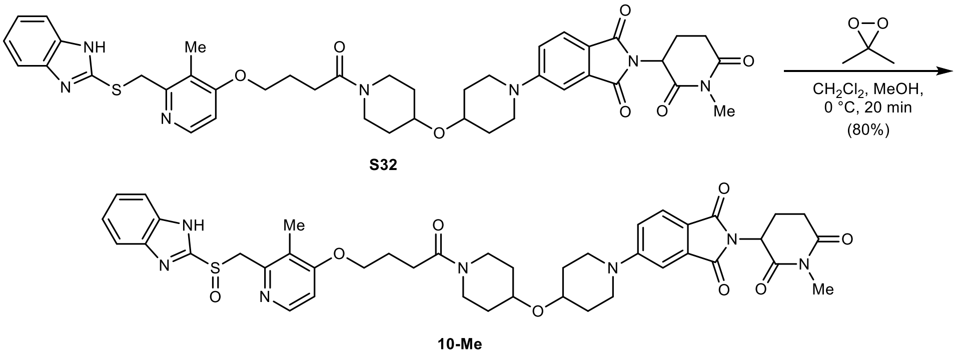

**5-(4-((1-(4-((2-(((1*H*-Benzo[*d*]imidazol-2-yl)sulfinyl)methyl)-3-methylpyridin-4-yl)oxy)butanoyl)piperidin-4-yl)oxy)piperidin-1-yl)-2-(1-methyl-2,6-dioxopiperidin-3-yl)isoindoline-1,3-dione (10-Me):** To a solution of **S32** (44 mg, 0.06 mmol, 1.00 equiv.) in anhydrous $CH_2Cl_2$ (0.75 mL, stored over basic aluminum oxide) was added a solution of 3,3-dimethyldioxirane (0.78 mL, 0.071 M in acetone by iodometric titration, 0.06 mmol, 1.00 equiv.) dropwise at 0 °C under argon. The reaction was stirred for 20 min at 0 °C, then concentrated *in vacuo*. Purification by column chromatography (4 g silica, 0 to 13% MeOH in $CH_2Cl_2$, filtered through basic aluminum oxide, in 16 min) yielded 36 mg (0.04 mmol, 80%) of **10-Me** as a yellow solid after lyophilization from MeCN/$H_2O$.

**TLC** $R_f$ = 0.49 (10% MeOH with 0.5% $NH_4OH$ in $CH_2Cl_2$).

**HRMS** (ESI) *m/z*: $(M+Na)^+$ calc. for $C_{42}H_{47}N_7NaO_8S$: 832.3099; found: 832.3107.

**$^1$H NMR** (400 MHz, DMSO-$d_6$) δ 8.21 (d, *J* = 5.6 Hz, 1H), 7.82 – 7.48 (m, 3H), 7.41 – 7.28 (m, 3H), 7.25 (dd, *J* = 8.7, 2.4 Hz, 1H), 6.96 (d, *J* = 5.7 Hz, 1H), 5.13 (dd, *J* = 13.0, 5.3 Hz, 1H), 4.79 (d, *J* = 13.6 Hz, 1H), 4.70 (d, *J* = 13.6 Hz, 1H), 4.08 (t, *J* = 6.3 Hz, 2H), 3.94 – 3.84 (m, 1H), 3.84 – 3.60 (m, 5H), 3.30 – 3.14 (m, 3H), 3.14 – 3.04 (m, 1H), 3.01 (s, 3H), 2.99 – 2.88 (m, 1H), 2.82 – 2.66 (m, 1H), 2.63 – 2.52 (m, 3H), 2.14 (s, 3H), 2.06 – 1.92 (m, 3H), 1.92 – 1.68 (m, 4H), 1.56 – 1.25 (m, 4H) ppm. **Note:** The benzimidazole NH peak is in exchange with residual water in the spectrum.

**$^{13}$C NMR** (101 MHz, DMSO-$d_6$) δ 171.8, 169.9, 169.8, 167.6, 167.0, 162.8, 154.8, 154.4, 150.1, 148.1, 134.0, 125.0, 123.2, 121.9, 117.7, 117.6, 107.8, 106.5, 70.8, 70.7, 67.5, 60.2, 49.3, 44.9, 42.4, 38.7, 32.1, 31.5, 31.2, 30.8, 28.5, 26.6, 24.3, 21.4, 10.7 ppm. **Note:** C1, C2, C3, and C6 are not detected due to benzimidazole tautomerism.

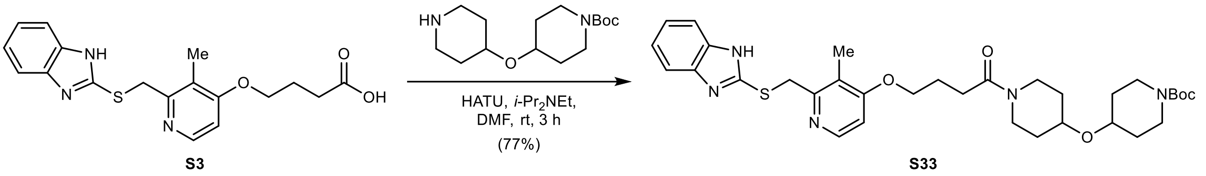

***tert*-Butyl 4-((1-(4-((2-(((1*H*-benzo[*d*]imidazol-2-yl)thio)methyl)-3-methylpyridin-4-yl)oxy)butanoyl)piperidin-4-yl)oxy)piperidine-1-carboxylate (S33):** To a suspension of **S3** (249 mg, 0.70 mmol, 1.00 equiv.) in anhydrous DMF (3.5 mL) was added *tert*-butyl 4-(piperidin-4-yloxy)piperidine-1-carboxylate (208 mg, 0.73 mmol, 1.05 equiv.) HATU (318 mg, 0.84 mmol, 1.20 equiv.), and *i*-$Pr_2NEt$ (364 µL, 2.09 mmol, 3.00 equiv.) at rt under argon. The reaction was stirred at rt for 3 h. The mixture was poured into water (200 mL) and extracted with EtOAc (3 × 50 mL). The combined organic layers were washed with 5% aqueous LiCl (25 mL), water (25 mL), and brine (25 mL), dried with $MgSO_4$, filtered, and concentrated *in vacuo*. Purification by column chromatography (40 g silica, 0 to 7.5% MeOH with 2.5% $NH_4OH$ in $CH_2Cl_2$ in 15 min) yielded 336 mg (0.54 mmol, 77%) of **S33** as an off-white solid.

**TLC** $R_f$ = 0.43 (10% MeOH in $CH_2Cl_2$).

**LC/MS** (ESI) *m/z*: $(M+H)^+$ 624.3.

**$^1$H NMR** (400 MHz, $CDCl_3$) δ 8.34 (dd, *J* = 5.8, 0.6 Hz, 1H), 7.58 – 7.49 (m, 2H), 7.22 – 7.13 (m, 2H), 6.80 (d, *J* = 5.8 Hz, 1H), 4.38 (s, 2H), 4.13 (t, *J* = 6.1 Hz, 2H), 3.89 (ddd, *J* = 12.2, 7.3, 3.8 Hz, 1H), 3.80 – 3.69 (m, 2H), 3.65 (tt, *J* = 7.4, 3.6 Hz, 2H), 3.56 (tt, *J* = 7.9, 3.7 Hz, 1H), 3.37 (ddd, *J* = 12.6, 8.1, 3.7 Hz, 1H), 3.27 (ddd, *J* = 12.9, 8.1, 3.5 Hz, 1H), 3.10 (ddd, *J* = 13.1, 9.0, 3.4 Hz, 2H), 2.53 (t, *J* = 7.0 Hz, 2H), 2.26 (s, 3H), 2.20 (p, *J* = 6.3 Hz, 2H), 1.84 – 1.71 (m, 4H), 1.62 – 1.48 (m, 4H), 1.45 (s, 9H) ppm. **Note:** The benzimidazole NH peak is in exchange with residual water in the spectrum.

**$^{13}$C NMR** (101 MHz, $CDCl_3$) δ 170.2, 164.4, 156.7, 155.0, 151.8, 147.4, 122.0, 121.1, 114.4, 106.4, 79.7, 72.0, 71.0, 68.0, 42.8, 41.3, 39.1, 34.9, 32.5, 31.9, 31.5, 29.2, 28.6, 24.6, 10.9 ppm. **Note:** C1 and C2 are not detected due to benzimidazole tautomerism.

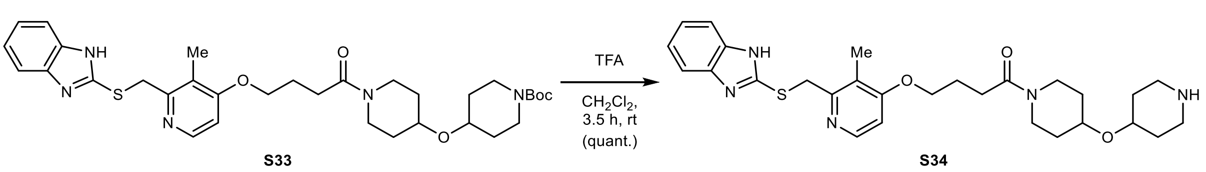

**4-((2-(((1*H*-Benzo[*d*]imidazol-2-yl)thio)methyl)-3-methylpyridin-4-yl)oxy)-1-(4-(piperidin-4-yloxy)piperidin-1-yl)butan-1-one (S34):** To a solution of **S33** (219 mg, 0.35 mmol, 1.00 equiv.) in $CH_2Cl_2$ (1.5 mL) was added trifluoroacetic acid (1.5 mL, 19.7 mmol, 56.0 equiv.) slowly at 0 °C. After 5 min, the reaction was allowed to warm to rt and stirred for 3.5 h. The mixture was poured into saturated aqueous $Na_2CO_3$ (100 mL) and extracted with 10% MeOH in $CH_2Cl_2$ (4 × 50 mL). The combined organic layers were dried with $MgSO_4$, filtered, and concentrated *in vacuo*, yielding 182 mg (0.35 mmol, quantitative) of **S34** as an off-white solid.

**TLC** $R_f$ = 0.20, streaky (50% MeOH with 0.5% $NH_4OH$ in $CH_2Cl_2$).

**LC/MS** (ESI) *m/z*: $(M+H)^+$ 524.2.

**$^1$H NMR** (400 MHz, $CD_3OD$) δ 8.20 (dd, *J* = 5.8, 0.6 Hz, 1H), 7.53 – 7.44 (m, 2H), 7.25 – 7.16 (m, 2H), 6.94 (d, *J* = 5.9 Hz, 1H), 4.61 (s, 2H), 4.14 (t, *J* = 6.1 Hz, 2H), 3.96 – 3.87 (m, 1H), 3.74 (m, 2H), 3.60 (tt, *J* = 8.4, 3.8 Hz, 1H), 3.38 – 3.23 (m, 2H), 3.10 – 3.00 (m, 2H), 2.72 – 2.54 (m, 4H), 2.24 (s, 3H), 2.13 (p, *J* = 6.7 Hz, 2H), 1.95 – 1.75 (m, 4H), 1.48 (m, 4H) ppm. **Note:** The NH peaks are in exchange with $CD_3OD$ in the spectrum.

**$^{13}$C NMR** (101 MHz, $CD_3OD$) δ 172.9, 165.5, 155.), 151.2, 148.8, 123.5, 122.7, 115.1, 107.5, 72.9, 72.3, 68.9, 44.3, 44.2, 40.4, 37.5, 33.5, 33.2, 32.7, 30.2, 25.9, 10.9 ppm. **Note:** C1 and C2 are not detected due to benzimidazole tautomerism.

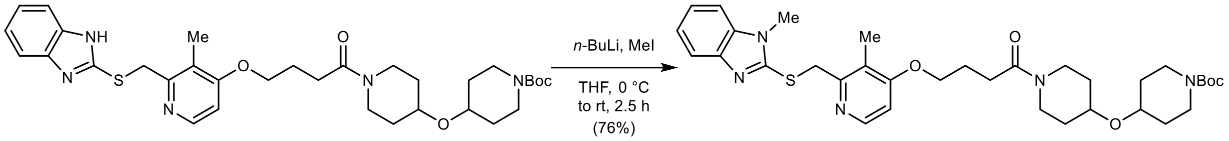

**S33** **S35**

***tert*-Butyl 4-((1-(4-((3-methyl-2-(((1-methyl-1*H*-benzo[*d*]imidazol-2-yl)thio)methyl)pyridin-4-yl)oxy)butanoyl)piperidin-4-yl)oxy)piperidine-1-carboxylate (S35):** To a solution of **S33** (50 mg, 0.08 mmol, 1.00 equiv.) in anhydrous THF (0.8 mL) was added *n*-butyllithium (0.06 mL, 1.6 M in hexanes, 0.10 mmol, 1.20 equiv.) at 0 °C under argon. After 15 min, methyl iodide (10 µL, 0.16 mmol, 2.00 equiv.) was added at 0 °C. The reaction was allowed to warm to rt and stirred for 2 h. The mixture was poured into saturated aqueous $NH_4Cl$ (100 mL) and extracted with EtOAc (3 × 25 mL). The combined organic layers were washed with water (25 mL) and brine (25 mL), dried with $MgSO_4$, filtered, and concentrated *in vacuo*. Purification by column chromatography (12 g silica, 0 to 8% MeOH with 2.5% $NH_4OH$ in $CH_2Cl_2$ in 12 min) yielded 39 mg (0.06 mmol, 76%) of **S35** as an off-white solid.

**TLC** $R_f$ = 0.68 (10% MeOH with 0.5% $NH_4OH$ in $CH_2Cl_2$).

**LC/MS** (ESI) *m/z*: $(M+H)^+$ 638.3.

**$^1$H NMR** (400 MHz, $CDCl_3$) δ 8.29 (d, *J* = 5.7 Hz, 1H), 7.72 – 7.63 (m, 1H), 7.25 – 7.16 (m, 3H), 6.71 (d, *J* = 5.7 Hz, 1H), 4.80 (s, 2H), 4.07 (t, *J* = 6.0 Hz, 2H), 3.88 (ddd, *J* = 15.2, 7.5, 3.9 Hz, 1H), 3.79 – 3.61 (m, 7H), 3.55 (tt, *J* = 7.9, 3.7 Hz, 1H), 3.35 (ddd, *J* = 12.6, 8.0, 3.6 Hz, 1H), 3.25 (ddd, *J* = 13.0, 8.1, 3.5 Hz, 1H), 3.09 (ddd, *J* = 12.9, 8.9, 3.4 Hz, 2H), 2.51 (t, *J* = 7.0 Hz, 2H), 2.27 (s, 3H), 2.16 (p, *J* = 6.6 Hz, 2H), 1.85 – 1.70 (m, 4H), 1.61 – 1.46 (m, 4H), 1.43 (s, 9H) ppm.

**$^{13}$C NMR** (101 MHz, $CDCl_3$) δ 170.3, 163.4, 154.9, 154.5, 152.4, 148.1, 143.5, 136.9, 122.0, 121.8, 120.9, 118.3, 108.6, 105.9, 79.6, 71.9, 70.9, 67.6, 42.7, 41.3, 39.0, 37.5, 32.4, 31.7, 31.4, 30.1, 29.2, 28.5, 24.7, 11.0 ppm.

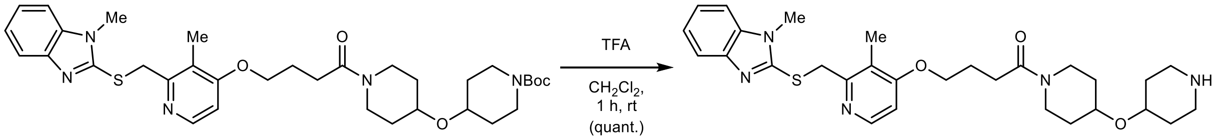

**S35** **S36**

**4-((3-Methyl-2-(((1-methyl-1*H*-benzo[*d*]imidazol-2-yl)thio)methyl)pyridin-4-yl)oxy)-1-(4-(piperidin-4-yloxy)piperidin-1-yl)butan-1-one (S36):** To a solution of **S35** (39 mg, 0.06 mmol, 1.00 equiv.) in $CH_2Cl_2$ (0.6 mL) was added trifluoroacetic acid (0.3 mL, 3.91 mmol, 64.0 equiv.) slowly at 0 °C. After 5 min, the reaction was allowed to warm to rt and stirred for 1 h. The mixture was poured into saturated aqueous $Na_2CO_3$ (100 mL) and extracted with 10% MeOH in $CH_2Cl_2$ (4 × 50 mL). The combined organic layers were dried with $MgSO_4$, filtered, and concentrated *in vacuo*, yielding 33 mg (0.06 mmol, quantitative) of **S36** as an off-white solid.

**TLC** $R_f$ = 0.19, streaky (50% MeOH with 0.5% $NH_4OH$ in $CH_2Cl_2$).

**LC/MS** (ESI) *m/z*: $(M+H)^+$ 538.3.

**$^1$H NMR** (400 MHz, $CD_3OD$) δ 8.15 (dd, *J* = 5.8, 0.6 Hz, 1H), 7.63 – 7.54 (m, 1H), 7.45 – 7.35 (m, 1H), 7.31 – 7.19 (m, 2H), 6.92 (d, *J* = 5.8 Hz, 1H), 4.60 (s, 2H), 4.12 (t, *J* = 6.2 Hz, 2H), 3.97 – 3.86 (m, 1H), 3.81 – 3.69 (m, 2H), 3.67 (s, 3H), 3.59 (tt, *J* = 8.4, 3.8 Hz, 1H), 3.37 – 3.23 (m, 2H), 3.04 (dt, *J* = 12.6, 4.6 Hz, 2H), 2.64 (ddd, *J* = 12.8, 9.8, 3.2 Hz, 2H), 2.61 – 2.54 (m, 2H), 2.19 (s, 3H), 2.17 – 2.06 (m, 2H), 1.94 – 1.75 (m, 4H), 1.48 (tt, *J* = 12.5, 4.0 Hz, 4H) ppm. **Note:** The NH peak is in exchange with $CD_3OD$ in the spectrum.

**$^{13}$C NMR** (101 MHz, $CD_3OD$) δ 172.8, 165.3, 155.5, 151.9, 148.9, 143.9, 137.7, 123.9, 123.5, 122.7, 118.8, 110.6, 107.5, 73.1, 72.3, 68.9, 44.4, 44.2, 40.4, 38.3, 33.5, 33.4, 33.4, 32.7, 30.7, 30.2, 25.9, 10.9 ppm.

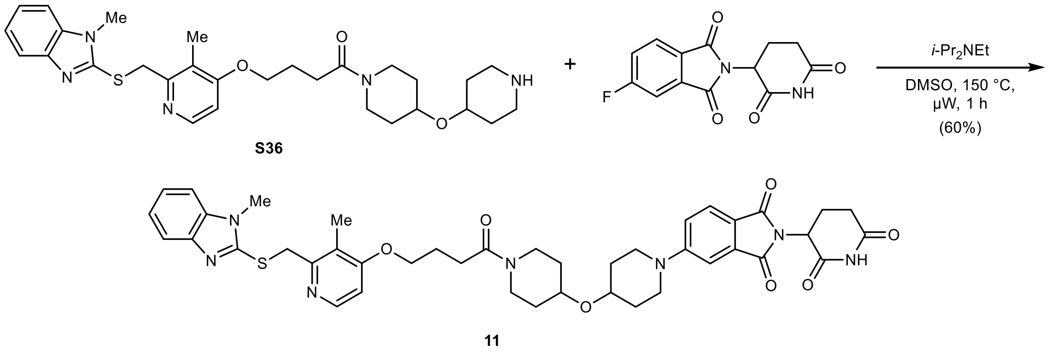

**2-(2,6-Dioxopiperidin-3-yl)-5-(4-((1-(4-((3-methyl-2-(((1-methyl-1*H*-benzo[*d*]imidazol-2-yl) thio)methyl)pyridin-4-yl)oxy)butanoyl)piperidin-4-yl)oxy)piperidin-1-yl)isoindoline-1,3-dione (11):** To a solution of **S36** (33 mg, 0.06 mmol, 1.00 equiv.) in anhydrous DMSO (1.2 mL) was added 2-(2,6-dioxopiperidin-3-yl)-5-fluoroisoindoline-1,3-dione (17 mg, 0.06 mmol, 1.00 equiv.) and *i*-$Pr_2NEt$ (21 µL, 0.12 mmol, 2.00 equiv.) at rt under argon. The reaction was stirred at 150 °C for 1 h under microwave irradiation. The mixture was diluted with water (100 mL) and extracted with EtOAc (3 × 25 mL). The combined organic layers were washed with water (50 mL) and brine (50 mL), dried with $MgSO_4$, filtered, and concentrated *in vacuo*. Purification by column chromatography (12 g silica, 0 to 9% MeOH with 2.5% $NH_4OH$ in $CH_2Cl_2$ in 14 min) yielded 29 mg (0.04 mmol, 60%) of **11** as a yellow powder after lyophilization from $MeCN/H_2O$.

**TLC** $R_f$ = 0.69 (10% MeOH with 0.5% $NH_4OH$ in $CH_2Cl_2$).

**HRMS** (ESI) *m/z*: $(M+H)^+$ calc. for $C_{42}H_{48}N_7O_7S$: 794.3330; found: 794.3356.

**$^1$H NMR** (400 MHz, $CDCl_3$) δ 8.31 (dd, *J* = 5.7, 0.6 Hz, 1H), 8.18 (bs, 1H), 7.72 – 7.64 (m, 2H), 7.27 (d, *J* = 2.4 Hz, 1H), 7.26 – 7.19 (m, 3H), 7.04 (dd, *J* = 8.6, 2.4 Hz, 1H), 6.72 (d, *J* = 5.7 Hz, 1H), 4.93 (dd, *J* = 12.3, 5.3 Hz, 1H), 4.82 (s, 2H), 4.09 (t, *J* = 5.9 Hz, 2H), 3.90 (ddd, *J* = 18.5, 8.6, 5.1 Hz, 1H), 3.75 – 3.64 (m, 8H), 3.38 (ddd, *J* = 12.6, 8.1, 3.6 Hz, 1H), 3.34 – 3.23 (m, 3H), 2.93 – 2.67 (m, 3H), 2.54 (t, *J* = 7.0 Hz, 2H), 2.29 (s, 3H), 2.24 – 2.08 (m, 3H), 1.98 – 1.87 (m, 2H), 1.87 – 1.75 (m, 2H), 1.75 – 1.63 (m, 2H), 1.63 – 1.50 (m, 2H) ppm.

**$^{13}$C NMR** (101 MHz, $CDCl_3$) δ 171.1, 170.4, 168.4, 168.1, 167.3, 163.4, 155.2, 154.6, 152.4, 148.2, 143.6, 136.9, 134.5, 125.6, 122.0, 121.9, 120.9, 119.0, 118.4, 118.0, 108.7, 108.6, 105.9, 71.2, 71.0, 67.6, 49.3, 45.4, 45.3, 42.8, 39.1, 37.5, 32.5, 31.6, 31.4, 31.2, 31.0, 30.2, 29.3, 24.8, 22.9, 11.0 ppm.

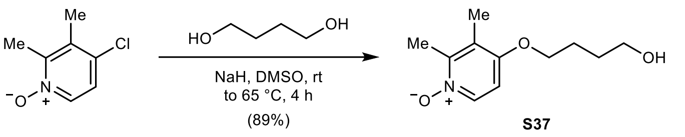

**4-(4-Hydroxybutoxy)-2,3-dimethylpyridine 1-oxide (S37):** To a solution of butane-1,4-diol (1.68 mL, 19.0 mmol, 6.00 equiv.) in anhydrous DMSO (6.0 mL) was added NaH (254 mg, 60 w% dispersion in mineral oil, 6.35 mmol, 2.00 equiv.) at rt under argon. The reaction was stirred at rt for 2 h. 4-Chloro-2,3-dimethylpyridine 1-oxide (500 mg, 3.17 mmol, 1.00 equiv.)[8] was added at 65 °C. The reaction was stirred at 65 °C for 2 h. The mixture was poured into MeOH (150 mL), washed with *n*-heptane (3 × 50 mL), and concentrated *in vacuo*. Purification by reversed-phase column chromatography (130 g C18, 5 to 40% MeCN in $H_2O$ with 0.05% TFA in 20 min), followed by another reversed-phase column chromatography (130 g C18, 5 to 30% MeCN in $H_2O$ with 0.05% TFA in 15 min) to remove residual butane-1,4-diol yielded 596 mg (2.82 mmol, 89%) of **S37** as a light-brown oil. **Note:** The high polarity of **S37** precludes an aqueous work-up procedure.

**TLC** $R_f$ = 0.46 (20% MeOH in $CH_2Cl_2$).

**LC/MS** (ESI) *m/z*: $(M+H)^+$ 212.2.

**$^1$H NMR** (400 MHz, DMSO-$d_6$) δ 8.07 (dt, *J* = 7.3, 0.6 Hz, 1H), 6.92 (d, *J* = 7.2 Hz, 1H), 4.47 (t, *J* = 5.2 Hz, 1H), 4.03 (t, *J* = 6.4 Hz, 2H), 3.45 (td, *J* = 6.4, 5.1 Hz, 2H), 2.34 (q, *J* = 0.6 Hz, 3H), 2.12 (q, *J* = 0.5 Hz, 3H), 1.82 – 1.70 (m, 2H), 1.62 – 1.50 (m, 2H) ppm.

**$^{13}$C NMR** (101 MHz, DMSO-$d_6$) δ 153.4, 147.4, 136.4, 122.4, 106.7, 68.4, 60.3, 28.9, 25.2, 13.9, 11.6 ppm.

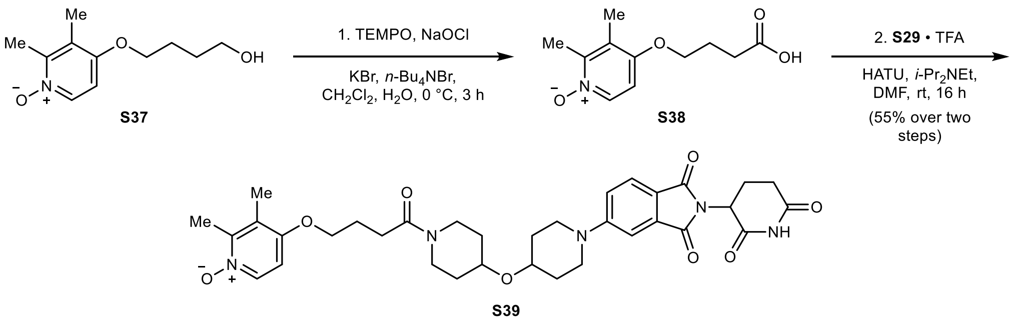

**4-(4-(4-((1-(2-(2,6-Dioxopiperidin-3-yl)-1,3-dioxoisoindolin-5-yl)piperidin-4-yl)oxy)piperidin-1-yl)-4-oxobutoxy)-2,3-dimethylpyridine 1-oxide (S39):** To a suspension of **S37** (254 mg, 1.12 mmol, 1.00 equiv.) in $CH_2Cl_2$ (7.5 mL) and $H_2O$ (7.5 mL) was added KBr (67 mg, 0.56 mmol, 0.50 equiv.), *n*-$Bu_4NBr$ (23 mg, 0.07 mmol, 0.07 equiv.), and TEMPO (11 mg, 0.07 mmol, 0.07 equiv.), followed by the dropwise addition of NaOCl (4.02 mL, 5.9% in water by iodometric titration, 3.91 mmol, 3.50 equiv.) at 0 °C. The reaction was stirred for 3 h at 0 °C, then concentrated *in vacuo*. **Note:** The high polarity of **S38** precludes an aqueous work-up procedure. To a suspension of crude **S38** in anhydrous DMF (5.0 mL) was added **S29 · TFA** (611 mg, 1.10 mmol, 1.04 equiv.), HATU (483 mg, 1.27 mmol, 1.20 equiv.), and *i*-$Pr_2NEt$ (645 µL, 3.71 mmol, 3.50 equiv.) at rt under argon. The reaction was stirred at rt for 16 h. The mixture was poured into water (300 mL) and extracted with 10% MeOH in $CH_2Cl_2$ (5 × 50 mL). The combined organic layers were washed with 5% aqueous LiCl (2 × 50 mL), water (2 × 50 mL), and brine (50 mL), dried with $MgSO_4$, filtered, and concentrated *in vacuo*. Residual DMF was removed by co-evaporation with toluene (2 × 10 mL). Purification by column chromatography (40 g silica, 3 to 25% MeOH with 2.5% $NH_4OH$ in $CH_2Cl_2$ in 20 min) yielded 376 mg (0.58 mmol, 55% over two steps) of **S39** as a yellow solid.

**TLC** $R_f$ = 0.44 (10% MeOH with 0.5% $NH_4OH$ in $CH_2Cl_2$).

**HRMS** (ESI) *m/z*: $(M+Na)^+$ calc. for $C_{34}H_{41}N_5NaO_8$: 670.2847; found: 670.2855.

**$^1$H NMR** (400 MHz, DMSO-$d_6$) δ 11.08 (s, 1H), 8.07 (dt, *J* = 7.2, 0.5 Hz, 1H), 7.65 (d, *J* = 8.5 Hz, 1H), 7.33 (d, *J* = 2.3 Hz, 1H), 7.25 (dd, *J* = 8.7, 2.4 Hz, 1H), 6.93 (d, *J* = 7.3 Hz, 1H), 5.06 (dd, *J* = 12.9, 5.4 Hz, 1H), 4.05 (t, *J* = 6.4 Hz, 2H), 3.94 – 3.84 (m, 1H), 3.84 – 3.76 (m, 2H), 3.76 – 3.62 (m, 3H), 3.29 – 3.15 (m, 3H), 3.14 – 3.04 (m, 1H), 2.88 (ddd, *J* = 17.3, 14.0, 5.4 Hz, 1H), 2.64 – 2.53 (m, 2H), 2.47 (d, *J* = 7.2 Hz, 2H), 2.34 (d, *J* = 0.6 Hz, 3H), 2.13 (s, 3H), 2.08 – 1.92 (m, 3H), 1.92 – 1.70 (m, 4H), 1.56 – 1.26 (m, 4H) ppm.

**$^{13}$C NMR** (101 MHz, DMSO-$d_6$) δ 172.8, 170.1, 169.7, 167.6, 166.9, 154.7, 153.3, 147.5, 136.5, 134.0, 125.0, 122.5, 117.6, 117.6, 107.8, 106.7, 70.8, 70.7, 67.9, 48.7, 44.9, 42.4, 38.7, 32.1, 31.4, 31.0, 30.8, 30.8, 28.4, 24.3, 22.2, 13.9, 11.6 ppm.

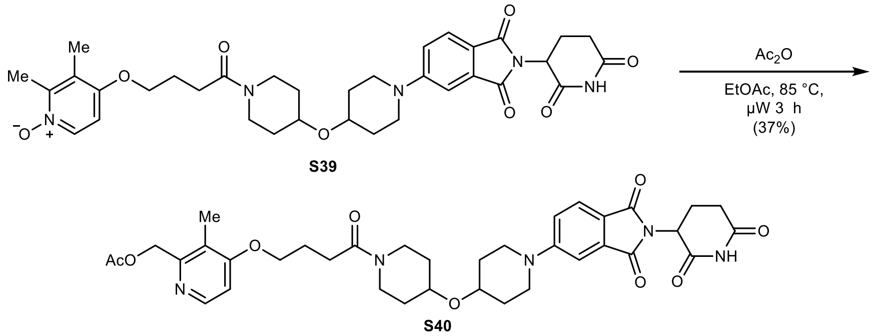

**(4-(4-(4-((1-(2-(2,6-Dioxopiperidin-3-yl)-1,3-dioxoisoindolin-5-yl)piperidin-4-yl)oxy)piperidin-1-yl)-4-oxobutoxy)-3-methylpyridin-2-yl)methyl acetate (S40):** To a suspension of **S39** (298 mg, 0.46 mmol, 1.00 equiv.) in EtOAc (4.0 mL, dried over $MgSO_4$) was added acetic acid anhydride (0.28 mL, 2.90 mmol, 6.35 equiv.) at rt under argon. The reaction was heated to 85 °C and stirred for 3 h under microwave irradiation. The mixture was poured into water (100 mL) and extracted with 10% MeOH in $CH_2Cl_2$ (4 × 50 mL). The combined organic layers were washed with brine (50 mL), dried with $MgSO_4$, filtered, and concentrated *in vacuo*. Purification by column chromatography (40 g silica, 0 to 7.5% MeOH with 2.5% $NH_4OH$ in $CH_2Cl_2$ in 25 min) yielded 116 mg (0.17 mmol, 37%) of **S40** as a yellow solid.

**TLC** $R_f$ = 0.56 (10% MeOH with 0.5% $NH_4OH$ in $CH_2Cl_2$).

**LC/MS** (ESI) *m/z*: $(M+H)^+$ 690.3.

**$^1$H NMR** (400 MHz, $CDCl_3$) δ 8.76 (d, *J* = 6.7 Hz, 1H), 8.31 (s, 1H), 7.66 (d, *J* = 8.5 Hz, 1H), 7.29 (d, *J* = 6.8 Hz, 1H), 7.27 (d, *J* = 2.4 Hz, 1H), 7.05 (dd, *J* = 8.6, 2.4 Hz, 1H), 5.36 (s, 2H), 4.93 (dd, *J* = 12.3, 5.3 Hz, 1H), 4.36 (t, *J* = 6.6 Hz, 2H), 3.88 (m, 1H), 3.71 (m, 5H), 3.44 (td, *J* = 8.4, 4.0 Hz, 1H), 3.28 (m, 3H), 2.78 (m, 3H), 2.53 (t, *J* = 6.6 Hz, 2H), 2.25 (m, 5H), 2.18 (s, 3H), 2.12 (m, 1H), 1.92 (m, 2H), 1.83 (m, 2H), 1.69 (m, 2H), 1.61 (m, 2H) ppm.

**$^{13}$C NMR** (101 MHz, $CDCl_3$) δ 171.2, 170.4, 169.7, 168.8, 168.5, 168.1, 167.3, 155.2, 148.3, 142.7, 134.5, 125.6, 124.2, 119.0, 118.0, 108.7, 108.0, 71.2, 70.9, 70.3, 59.9, 49.3, 45.4, 45.3, 42.7, 39.1, 32.4, 31.6, 31.3, 31.2, 31.0, 28.6, 24.0, 22.9, 20.5, 10.4 ppm.

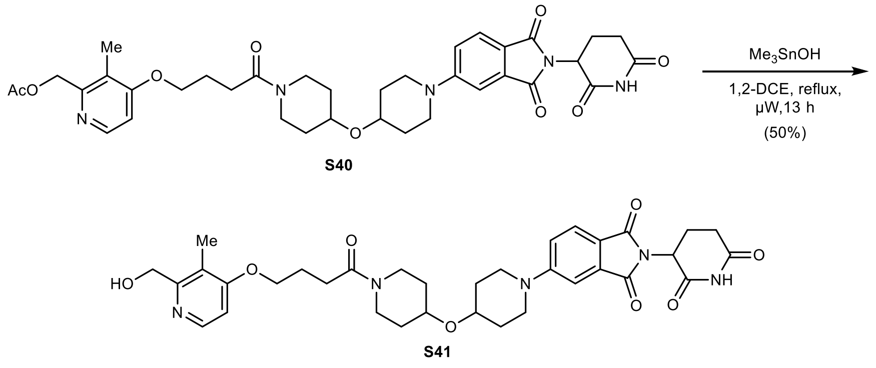

**2-(2,6-Dioxopiperidin-3-yl)-5-(4-((1-(4-((2-(hydroxymethyl)-3-methylpyridin-4-yl)oxy)butanoyl)piperidin-4-yl)oxy)piperidin-1-yl)isoindoline-1,3-dione (S41):** To a suspension of **S40** (81 mg, 0.12 mmol, 1.00 equiv.) in anhydrous 1,2-DCE (1.5 mL) was added trimethyltin hydroxide (212 mg, 1.17 mmol, 10.0 equiv.) at rt under argon. The reaction was heated to reflux under microwave irradiation and stirred for 12.5 h. To drive the reaction to completion, trimethyltin hydroxide (212 mg, 1.17 mmol, 10.0 equiv.) was added. The reaction was heated to reflux under microwave irradiation and stirred for 30 min. The mixture was concentrated *in vacuo*. Purification by reversed-phase column chromatography (15.5 g C18, 5 to 45% MeCN in $H_2O$ with 0.05% TFA in 15 min) yielded 38 mg (0.06 mmol, 50%) of **S41** as a yellow solid.

**TLC** $R_f$ = 0.66, streaky (20% MeOH in $CH_2Cl_2$).

**LC/MS** (ESI) *m/z*: $(M+H)^+$ 648.3.

**$^1$H NMR** (400 MHz, $CD_3CN$) δ 8.93 (bs, 1H), 8.40 (d, *J* = 6.9 Hz, 1H), 7.63 (d, *J* = 8.5 Hz, 1H), 7.34 (d, *J* = 6.9 Hz, 1H), 7.29 (d, *J* = 2.4 Hz, 1H), 7.16 (dd, *J* = 8.7, 2.4 Hz, 1H), 4.98 – 4.89 (m, 1H), 4.86 (s, 2H), 4.34 (t, *J* = 6.4 Hz, 2H), 3.98 – 3.90 (m, 1H), 3.81 – 3.65 (m, 5H), 3.31 – 3.21 (m, 3H), 3.21 – 3.10 (m, 1H), 2.83 – 2.61 (m, 3H), 2.53 (t, *J* = 7.0 Hz, 2H), 2.15 (s, 3H), 2.14 – 2.04 (m, 3H), 1.92 – 1.88 (m, 2H), 1.88 – 1.74 (m, 2H), 1.57 (dtd, *J* = 12.6, 8.6, 3.7 Hz, 2H), 1.52 – 1.34 (m, 2H) ppm.

**$^{13}$C NMR** (101 MHz, $CD_3CN$) δ 172.1, 169.9, 169.7, 169.0, 168.0, 167.3, 155.3, 153.6, 140.1, 134.5, 124.9, 122.2, 118.3, 117.8, 108.0, 107.9, 71.3, 71.1, 70.3, 58.3, 49.1, 45.1, 45.1, 42.6, 38.9, 32.2, 31.6, 31.1, 31.0, 31.0, 28.4, 24.0, 22.3, 8.7 ppm.

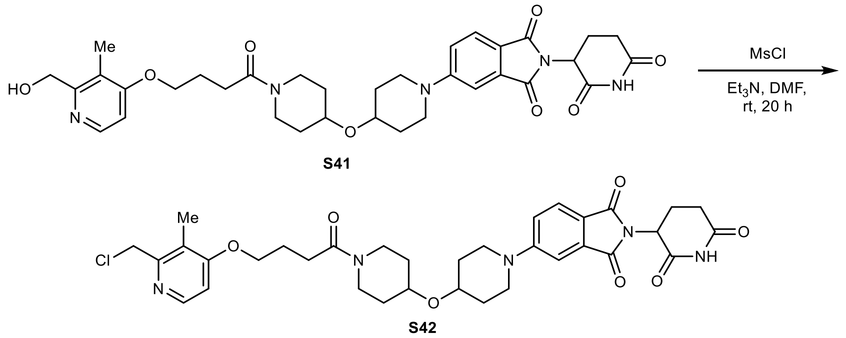

**5-(4-((1-(4-((2-(Chloromethyl)-3-methylpyridin-4-yl)oxy)butanoyl)piperidin-4-yl)oxy)piperidin-1-yl)-2-(2,6-dioxopiperidin-3-yl)isoindoline-1,3-dione (S42):** To a suspension of **S41** (34 mg, 0.04 mmol, 1.00 equiv.) in anhydrous $CH_2Cl_2$ (0.7 mL) and anhydrous DMF (0.1 mL) was added methanesulfonyl chloride (15.5 µL, 0.20 mmol, 5.00 equiv.) and $Et_3N$ (33.6 µL, 0.24 mmol, 6.00 equiv.) at 0 °C under argon. After 5 min, the mixture was allowed to warm to rt and stirred for 20 h. The mixture was concentrated *in vacuo* and used in the next step without further purification or characterization. Based on LC/MS, 75% purity was determined. **Note:** The mesylated alcohol is identified as a reaction intermediate via LC/MS and undergoes substitution to the chloride **S42**.

**TLC** $R_f$ = 0.60 (10% MeOH in $CH_2Cl_2$).

**LC/MS** (ESI) *m/z*: $(M+H)^+$ 666.2.

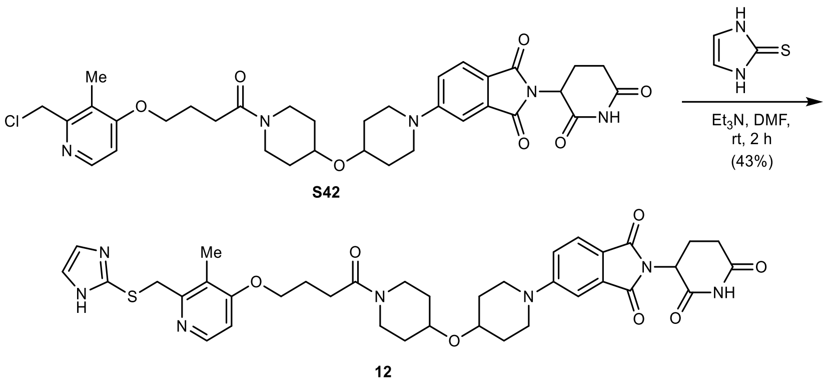

**5-(4-((1-(4-((2-(((1*H*-Imidazol-2-yl)thio)methyl)-3-methylpyridin-4-yl)oxy)butanoyl)piperidin-4-yl)oxy)piperidin-1-yl)-2-(2,6-dioxopiperidin-3-yl)isoindoline-1,3-dione (12):** To a suspension of **S42** (18 mg, approx. 75% purity according to LC/MS, 0.02 mmol, 1.00 equiv.) in anhydrous DMF (0.3 mL) was added $Et_3N$ (6.6 µL, 0.05 mmol, 2.10 equiv.) and 1,3-dihydro-2*H*-imidazole-2-thione (2.4 mg, 0.02 mmol, 1.05 equiv.) at rt under argon. The reaction was stirred at rt for 2 h. The mixture was poured into water (50 mL) and extracted with 10% MeOH in $CH_2Cl_2$ (4 × 10 mL). The combined organic layers were washed with 5% aqueous LiCl (2 × 10 mL), water (10 mL), and brine (10 mL), dried with $MgSO_4$, filtered, and concentrated *in vacuo*. Purification by reversed-phase column chromatography (15.5 g C18, 10 to 60% MeCN in $H_2O$ in 15 min) yielded 7 mg (0.01 mmol, 43% over two steps) of **12** as a yellow powder.

**TLC** $R_f$ = 0.39 (10% MeOH with 0.5% $NH_4OH$ in $CH_2Cl_2$).

**HRMS** (ESI) *m/z*: $(M+H)^+$ calc. for $C_{37}H_{44}N_7O_7S$: 730.3017; found: 730.3019.

**$^1$H NMR** (400 MHz, $CDCl_3$) δ 8.29 (d, *J* = 5.7 Hz, 1H), 8.11 (bs, 1H), 7.67 (d, *J* = 8.5 Hz, 1H), 7.28 (d, *J* = 2.4 Hz, 1H), 7.08 – 7.01 (m, 3H), 6.76 (d, *J* = 5.8 Hz, 1H), 4.93 (dd, *J* = 12.3, 5.4 Hz, 1H), 4.23 (s, 2H), 4.12 (t, *J* = 6.0 Hz, 2H), 3.98 – 3.86 (m, 1H), 3.81 – 3.64 (m, 5H), 3.39 (td, *J* = 8.7, 4.3 Hz, 1H), 3.36 – 3.23 (m, 3H), 2.98 – 2.66 (m, 3H), 2.54 (t, *J* = 6.9 Hz, 2H), 2.25 – 2.16 (m, 5H), 2.16 – 2.09 (m, 1H), 1.92 (t, *J* = 10.2 Hz, 2H), 1.86 – 1.76 (m, 2H), 1.75 – 1.64 (m, 2H), 1.62 – 1.52 (m, 2H) ppm. **Note:** The imidazole NH peak is in exchange with residual water in the spectrum.

**$^{13}$C NMR** (101 MHz, $CDCl_3$) δ 171.1, 170.3, 168.4, 168.1, 167.4, 164.2, 157.4, 155.3, 147.6, 142.0, 134.6, 125.6, 120.9, 119.0, 118.0, 108.8, 106.1, 71.2, 71.1, 67.8, 49.3, 45.4, 45.4, 42.7, 39.1, 36.0, 32.5, 31.6, 31.5, 31.2, 31.0, 29.2, 24.6, 22.9, 10.9 ppm. **Note:** C1 and C2 are not detected due to imidazole tautomerism.

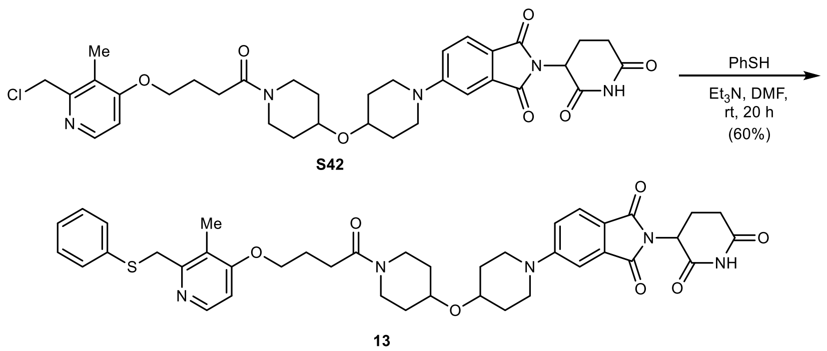

**2-(2,6-Dioxopiperidin-3-yl)-5-(4-((1-(4-((3-methyl-2-((phenylthio)methyl)pyridin-4-yl)oxy)butanoyl)piperidin-4-yl)oxy)piperidin-1-yl)isoindoline-1,3-dione (13):** To a suspension of **S42** (18 mg, approx. 75% purity according to LC/MS, 0.02 mmol, 1.00 equiv.) in anhydrous DMF (0.3 mL) was added $Et_3N$ (6.6 µL, 0.05 mmol, 2.10 equiv.) and benzenethiol (2.4 µL, 0.02 mmol, 1.05 equiv.) at rt under argon. The reaction was stirred at rt for 16 h. To drive the reaction to completion, $Et_3N$ (3.3 µL, 0.02 mmol, 1.05 equiv.) and benzenethiol (2.4 µL, 0.02 mmol, 1.05 equiv.) were added. The reaction was stirred at rt for another 4 h. The mixture was poured into saturated aqueous $NaHCO_3$ (50 mL) and extracted with 10% MeOH in $CH_2Cl_2$ (3 × 10 mL). The combined organic layers were washed with 5% aqueous LiCl (2 × 10 mL), water (10 mL), and brine (10 mL), dried with $MgSO_4$, filtered, and concentrated *in vacuo*. Purification by column chromatography (12 g silica, 0 to 10% MeOH in $CH_2Cl_2$ in 13 min) yielded 10 mg (0.01 mmol, 60% over two steps) of **13** as a yellow powder after lyophilization from MeCN/$H_2O$.

**TLC** $R_f$ = 0.57 (10% MeOH with 0.5% $NH_4OH$ in $CH_2Cl_2$).

**HRMS** (ESI) *m/z*: $(M+H)^+$ calc. for $C_{40}H_{46}N_5O_7S$: 740.3112; found: 740.3115.

**$^1$H NMR** (400 MHz, $CDCl_3$) δ 8.30 (d, *J* = 5.7 Hz, 1H), 8.13 (s, 1H), 7.72 (d, *J* = 8.6 Hz, 1H), 7.50 – 7.42 (m, 2H), 7.34 – 7.31 (m, 2H), 7.30 – 7.28 (m, 1H), 7.25 – 7.21 (m, 1H), 7.09 (dd, *J* = 8.6, 2.4 Hz, 1H), 6.72 (d, *J* = 5.7 Hz, 1H), 4.98 (dd, *J* = 12.3, 5.3 Hz, 1H), 4.34 (s, 2H), 4.12 (t, *J* = 6.0 Hz, 2H), 4.01 – 3.91 (m, 1H), 3.81 – 3.68 (m, 5H), 3.49 – 3.38 (m, 1H), 3.39 – 3.28 (m, 3H), 2.98 – 2.70 (m, 3H), 2.58 (t, *J* = 7.1 Hz, 2H), 2.27 – 2.14 (m, 6H), 2.01 – 1.92 (m, 2H), 1.90 – 1.81 (m, 2H), 1.80 – 1.69 (m, 2H), 1.68 – 1.57 (m, 2H) ppm.

**$^{13}$C NMR** (101 MHz, $CDCl_3$) δ 171.1, 170.4, 168.4, 168.1, 167.3, 163.5, 155.9, 155.2, 147.9, 136.5, 134.6, 130.2, 129.0, 126.6, 125.6, 121.0, 119.0, 118.0, 108.8, 105.7, 71.2, 71.0, 67.5, 49.3, 45.4, 45.4, 42.8, 39.4, 39.1, 32.5, 31.6, 31.4, 31.2, 31.0, 29.3, 24.8, 22.9, 11.0 ppm.

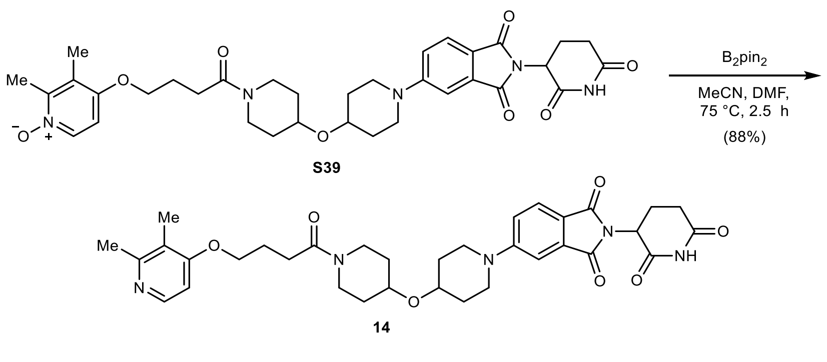

**5-(4-((1-(4-((2,3-Dimethylpyridin-4-yl)oxy)butanoyl)piperidin-4-yl)oxy)piperidin-1-yl)-2-(2,6-dioxopiperidin-3-yl)isoindoline-1,3-dione (14):** To a suspension of **S39** (50 mg, 0.07 mmol, 1.00 equiv.) in anhydrous MeCN (0.7 mL) and anhydrous DMF (0.1 mL) was added bis(pinacolato)diboron (19 mg, 0.08 mmol, 1.10 equiv.) at rt under argon. The reaction was heated to 75 °C and stirred for 2.5 h. The mixture was poured into water (100 mL) and extracted with 10% MeOH in $CH_2Cl_2$ (5 × 30 mL). The combined organic layers were washed with water (50 mL), and brine (50 mL), dried with $MgSO_4$, filtered, and concentrated *in vacuo*. Purification by column chromatography (12 g silica, 1 to 10% MeOH with 2.5% $NH_4OH$ in $CH_2Cl_2$ in 10 min) yielded 39 mg (0.06 mmol, 88%) of **14** as a yellow powder after lyophilization from MeCN/$H_2O$.

**TLC** $R_f$ = 0.46 (10% MeOH with 0.5% $NH_4OH$ in $CH_2Cl_2$).

**HRMS** (ESI) *m/z*: $(M+H)^+$ calc. for $C_{34}H_{42}N_5O_7$: 632.3079; found: 632.3077.

**$^1$H NMR** (400 MHz, $CDCl_3$) δ 8.30 – 8.15 (m, 2H), 7.67 (d, *J* = 8.5 Hz, 1H), 7.28 (d, *J* = 2.4 Hz, 1H), 7.05 (dd, *J* = 8.6, 2.4 Hz, 1H), 6.62 (d, *J* = 5.7 Hz, 1H), 4.93 (dd, *J* = 12.2, 5.3 Hz, 1H), 4.06 (t, *J* = 6.0 Hz, 2H), 3.99 – 3.85 (m, 1H), 3.83 – 3.63 (m, 5H), 3.38 (ddd, *J* = 12.6, 8.1, 3.5 Hz, 1H), 3.34 – 3.23 (m, 3H), 2.93 – 2.65 (m, 3H), 2.54 (t, *J* = 7.1 Hz, 2H), 2.47 (s, 3H), 2.22 – 2.08 (m, 6H), 1.97 – 1.87 (m, 2H), 1.87 – 1.77 (m, 2H), 1.77 – 1.64 (m, 2H), 1.64 – 1.51 (m, 2H) ppm.

**$^{13}$C NMR** (101 MHz, $CDCl_3$) δ 171.1, 170.5, 168.4, 168.1, 167.3, 162.8, 157.8, 155.2, 147.6, 134.6, 125.6, 119.8, 119.0, 118.0, 108.8, 104.7, 71.2, 71.0, 67.3, 49.3, 45.4, 45.4, 42.8, 39.1, 32.5, 31.6, 31.4, 31.2, 31.0, 29.4, 24.9, 22.9, 22.8, 11.1 ppm.

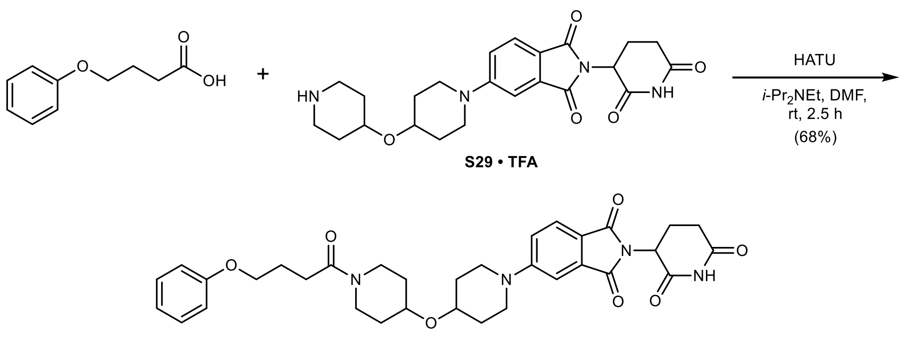

**2-(2,6-Dioxopiperidin-3-yl)-5-(4-((1-(4-phenoxybutanoyl)piperidin-4-yl)oxy)piperidin-1-yl) isoindoline-1,3-dione (15):** To a solution of **S29 · TFA** (55 mg, 0.10 mmol, 1.00 equiv.) in anhydrous DMF (1.0 mL) was added 4-phenoxybutanoic acid (22 mg, 0.12 mmol, 1.20 equiv.), HATU (45 mg, 0.12 mmol, 1.20 equiv.), and *i*-$Pr_2NEt$ (52 µL, 0.30 mmol, 3.00 equiv.) at rt under argon. The reaction was stirred at rt for 2.5 h. The mixture was poured into water (100 mL) and extracted with EtOAc (3 × 25 mL). The combined organic layers were washed with 5% aqueous LiCl (25 mL), water (25 mL), and brine (25 mL), dried with $MgSO_4$, filtered, and concentrated *in vacuo*. The crude reaction mixture was subjected to column chromatography (12 g silica, 0 to 5% MeOH in $CH_2Cl_2$ in 10 min), but $^1$H NMR spectroscopy of the apparently pure material revealed 14% of the active ester of 4-phenoxybutanoic acid. To drive the reaction to completion, the crude was dissolved in $CH_2Cl_2$ (0.5 mL), then **S29 · TFA** (12 mg, 0.02 mmol, 0.20 equiv.) and *i*-$Pr_2NEt$ (5.8 µL, 0.03 mmol, 0.33 equiv.) were added. The reaction was stirred at rt for 2 h. The mixture was poured into water (100 mL) and extracted with $CH_2Cl_2$ (3 × 25 mL). The combined organic layers were washed with brine (25 mL), dried with $MgSO_4$, filtered, and concentrated *in vacuo*. Purification by column chromatography (4 g silica, 0 to 6% MeOH in $CH_2Cl_2$ in 10 min) yielded 41 mg (0.07 mmol, 68%) of **15** as a yellow solid.

**TLC** $R_f$ = 0.46 (10% MeOH in $CH_2Cl_2$).

**HRMS** (ESI) *m/z*: $(M+H)^+$ calc. for $C_{33}H_{39}N_4O_7$: 603.2813; found: 603.2819.

**$^1$H NMR** (400 MHz, $CDCl_3$) δ 8.06 (bs, 1H), 7.67 (d, *J* = 8.6 Hz, 1H), 7.33 – 7.23 (m, 3H), 7.05 (dd, *J* = 8.6, 2.4 Hz, 1H), 6.98 – 6.85 (m, 3H), 4.94 (dd, *J* = 12.3, 5.3 Hz, 1H), 4.03 (t, *J* = 5.9 Hz, 2H), 3.92 (ddd, *J* = 12.1, 7.2, 3.9 Hz, 1H), 3.79 – 3.64 (m, 5H), 3.37 (ddd, *J* = 12.7, 8.2, 3.6 Hz, 1H), 3.33 – 3.24 (m, 3H), 2.94 – 2.67 (m, 3H), 2.54 (t, *J* = 7.3 Hz, 2H), 2.19 – 2.09 (m, 3H), 1.99 – 1.87 (m, 2H), 1.87 – 1.76 (m, 2H), 1.76 – 1.65 (m, 2H), 1.60 – 1.50 (m, 2H) ppm.

**$^{13}$C NMR** (101 MHz, $CDCl_3$) δ 171.1, 170.8, 168.4, 168.1, 167.3, 159.0, 155.3, 134.6, 129.6, 125.6, 120.8, 119.0, 118.0, 114.6, 108.8, 71.4, 71.0, 67.0, 49.3, 45.4, 42.8, 39.0, 32.5, 31.6, 31.5, 31.2, 31.0, 29.6, 25.1, 22.9 ppm.

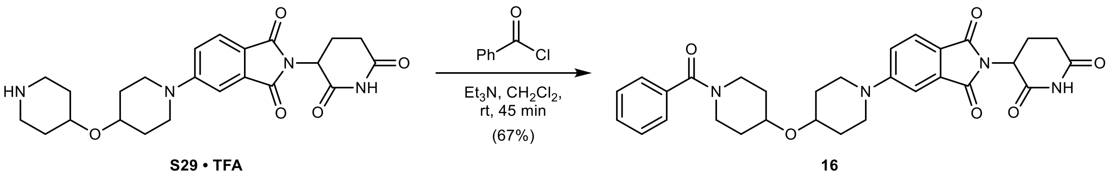

**5-(4-((1-Benzoylpiperidin-4-yl)oxy)piperidin-1-yl)-2-(2,6-dioxopiperidin-3-yl)isoindoline-1,3-dione (16):** To a suspension of **S29 · TFA** (35 mg, 0.06 mmol, 1.00 equiv.) in anhydrous $CH_2Cl_2$ (0.6 mL) was added $Et_3N$ (35 µL, 0.25 mmol, 4.00 equiv.) and benzoyl chloride (18 µL, 0.16 mmol, 2.50 equiv.) at rt under argon. The reaction was stirred at rt for 45 min. The mixture was poured into saturated aqueous $NaHCO_3$ (100 mL) and extracted with EtOAc (3 × 25 mL). The combined organic layers were washed with water (25 mL) and brine (25 mL), dried with $MgSO_4$, filtered, and concentrated *in vacuo*. Purification by column chromatography (12 g silica, 0 to 4.5% MeOH in $CH_2Cl_2$ in 9 min), followed by reversed-phase column chromatography (15.5 g C18, 10 to 50% MeCN in $H_2O$ in 12 min) yielded 23 mg (0.04 mmol, 67%) of **16** as a yellow powder after lyophilization from MeCN/$H_2O$.

**TLC** $R_f$ = 0.27 (5% MeOH in $CH_2Cl_2$).

**HRMS** (ESI) *m/z*: $(M+Na)^+$ calc. for $C_{30}H_{32}N_4NaO_6$: 567.2214; found: 567.2226.

**$^1$H NMR** (400 MHz, $CDCl_3$) δ 8.14 (s, 1H), 7.67 (d, *J* = 8.5 Hz, 1H), 7.46 – 7.34 (m, 5H), 7.28 (d, *J* = 2.3 Hz, 1H), 7.05 (dd, *J* = 8.6, 2.4 Hz, 1H), 4.93 (dd, *J* = 12.3, 5.3 Hz, 1H), 4.06 (bs, 1H), 3.81 – 3.40 (m, 6H), 3.40 – 3.11 (m, 3H), 2.93 – 2.65 (m, 3H), 2.18 – 2.07 (m, 1H), 1.99 – 1.43 (m, 8H) ppm.

**$^{13}$C NMR** (101 MHz, $CDCl_3$) δ 171.1, 170.5, 168.4, 168.1, 167.3, 155.2, 136.3, 134.5, 129.7, 128.6, 126.9, 125.6, 119.0, 118.0, 108.8, 71.4, 71.0, 49.3, 45.4, 45.1, 39.5, 32.6, 31.6, 31.1, 22.9 ppm.

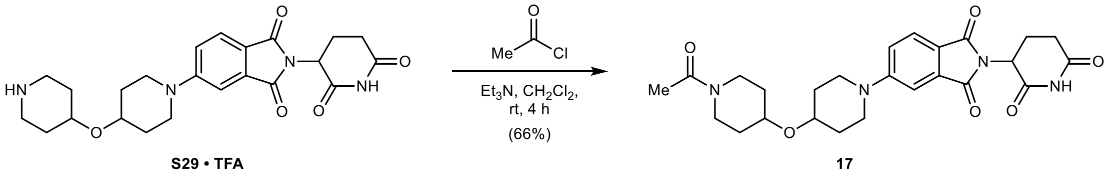

**5-(4-((1-Acetylpiperidin-4-yl)oxy)piperidin-1-yl)-2-(2,6-dioxopiperidin-3-yl)isoindoline-1,3-dione (17):** To a suspension of **S29 · TFA** (35 mg, 0.06 mmol, 1.00 equiv.) in anhydrous $CH_2Cl_2$ (0.6 mL) was added $Et_3N$ (22 µL, 0.16 mmol, 2.50 equiv.) and acetyl chloride (9.0 µL, 0.13 mmol, 2.00 equiv.) at rt under argon. The reaction was stirred at rt for 4 h. The mixture was poured into saturated aqueous $NaHCO_3$ (100 mL) and extracted with EtOAc (3 × 25 mL). The combined organic layers were washed with water (25 mL) and brine (25 mL), dried with $MgSO_4$, filtered, and concentrated *in vacuo*. Purification by column chromatography (12 g silica, 0 to 9% MeOH in $CH_2Cl_2$ in 9 min) yielded 20 mg (0.04 mmol, 66%) of **17** as a yellow powder after lyophilization from MeCN/$H_2O$.

**TLC** $R_f$ = 0.41 (10% MeOH in $CH_2Cl_2$).

**HRMS** (ESI) *m/z*: $(M+Na)^+$ calc. for $C_{25}H_{30}N_4NaO_6$: 505.2058; found: 505.2062.

**$^1$H NMR** (400 MHz, $CDCl_3$) δ 8.09 – 8.01 (m, 1H), 7.67 (d, *J* = 8.5 Hz, 1H), 7.28 (d, *J* = 2.3 Hz, 1H), 7.05 (dd, *J* = 8.6, 2.4 Hz, 1H), 4.94 (dd, *J* = 12.3, 5.3 Hz, 1H), 3.91 (ddd, *J* = 12.0, 7.2, 3.9 Hz, 1H), 3.76 – 3.62 (m, 5H), 3.41 – 3.22 (m, 4H), 2.94 – 2.65 (m, 3H), 2.18 – 2.10 (m, 1H), 2.10 (s, 3H), 1.99 – 1.88 (m, 2H), 1.88 – 1.76 (m, 2H), 1.76 – 1.65 (m, 2H), 1.65 – 1.50 (m, 2H) ppm.

**$^{13}$C NMR** (101 MHz, $CDCl_3$) δ 171.0, 168.9, 168.3, 168.0, 167.2, 155.1, 134.5, 125.5, 118.9, 117.9, 108.6, 71.2, 70.9, 49.2, 45.3, 45.3, 43.6, 38.7, 32.4, 31.5, 31.3, 31.1, 30.9, 22.8, 21.5 ppm.

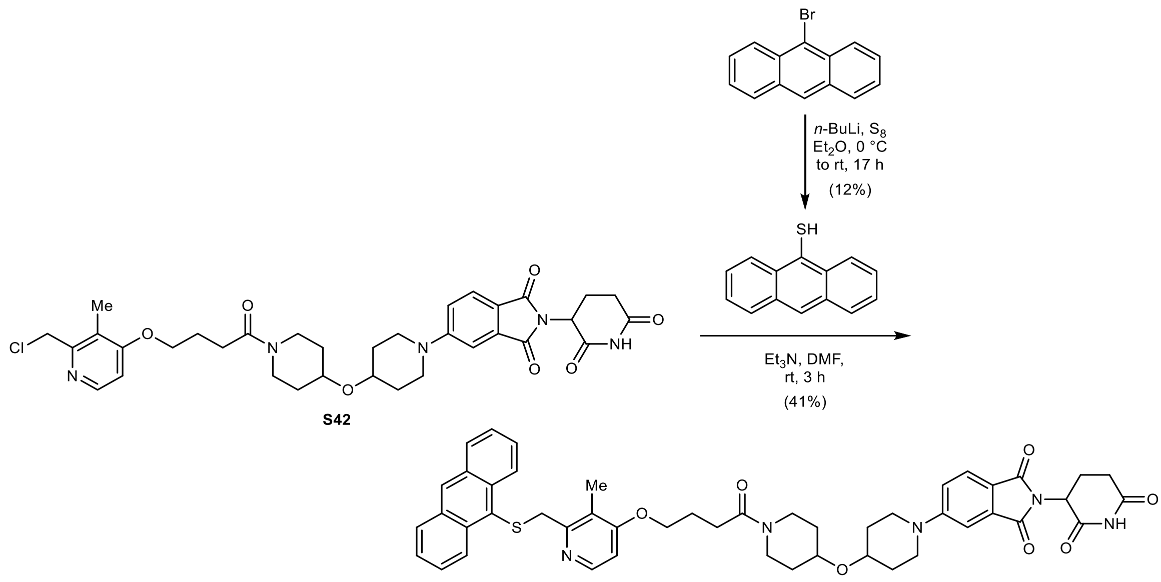

**5-(4-((1-(4-((2-((Anthracen-9-ylthio)methyl)-3-methylpyridin-4-yl)oxy)butanoyl)piperidin-4-yl)oxy)piperidin-1-yl)-2-(2,6-dioxopiperidin-3-yl)isoindoline-1,3-dione (18):** To a suspension of 9-bromoanthracene (1.00 g, 3.89 mmol, 1.00 equiv.) in anhydrous $Et_2O$ (10 mL) was added *n*-butyllithium (1.87 mL, 2.5 M in hexanes, 4.67 mmol, 1.20 equiv.) dropwise at 0 °C under argon. The reaction was stirred for 40 min at 0 °C. Sulfur (0.15 g, 4.67 mmol, 1.20 equiv.) was added. The reaction was allowed to warm to rt and stirred for 16 h. The mixture was diluted with $Et_2O$ (100 mL) and extracted with 2 M aqueous NaOH (3 × 75 mL). The combined aqueous layers were acidified to pH = 1 using 3 M aqueous HCl at 0 °C, then re-extracted with $Et_2O$ (3 × 75 mL). The combined organic layers were dried with $MgSO_4$, filtered, and concentrated *in vacuo*. **Note:** Recrystallization from 5% EtOAc in *n*-heptane at reflux was unsuccessful and likely resulted in the presence of oxygen and elevated temperatures in the formation of polysulfides. Purification by column chromatography (40 g silica, 0 to 30% $CH_2Cl_2$ in *n*-heptane in 15 min) yielded 100 mg (0.47 mmol, 12%) of 9-thioanthracene as a yellow solid.

**TLC** $R_f$ = 0.58 (66% $CH_2Cl_2$ in *n*-heptane).

**LC/MS** (ESI) *m/z*: $(M+H)^+$ 211.0.

**$^1$H NMR** (400 MHz, $CDCl_3$) δ 8.63 (dq, *J* = 8.9, 1.0 Hz, 2H), 8.40 (s, 1H), 8.01 (ddt, *J* = 8.4, 1.4, 0.7 Hz, 2H), 7.59 (ddd, *J* = 8.9, 6.5, 1.4 Hz, 2H), 7.51 (ddd, *J* = 8.5, 6.5, 1.1 Hz, 2H), 3.69 (d, *J* = 0.9 Hz, 1H) ppm.

**$^{13}$C NMR** (101 MHz, $CDCl_3$) δ 132.3, 131.8, 129.2, 127.2, 126.5, 126.3, 125.5, 123.8 ppm. The spectroscopic data is in accordance with the literature.[9]

To a suspension of 9-thioanthracene (27 mg, approx. 62% according to LC/MS, 0.03 mmol, 1.00 equiv.) in anhydrous DMF (0.3 mL) was added $Et_3N$ (6.2 µL, 0.05 mmol, 1.77 equiv.) and **S42** (9 mg, 0.04 mmol, 1.70 equiv.) at rt under argon. The reaction was stirred at rt for 3 h. The mixture was poured into water (50 mL) and extracted with 10% MeOH in $CH_2Cl_2$ (5 × 10 mL). The combined organic layers were washed with 5% aqueous LiCl (2 × 10 mL), water (10 mL), and brine (10 mL), dried with $MgSO_4$, filtered, and concentrated *in vacuo*. Purification by column chromatography (12 g silica, 0 to 9% MeOH with 2.5% $NH_4OH$ in $CH_2Cl_2$ in 12 min) yielded 14 mg (0.02 mmol, 41% over two steps) of **18** as a yellow powder after lyophilization from MeCN/$H_2O$.

**TLC** $R_f$ = 0.31 (5% MeOH with 0.5% $NH_4OH$ in $CH_2Cl_2$).

**HRMS** (ESI) *m/z*: $(M+H)^+$ calc. for $C_{48}H_{50}N_5O_7S$: 840.3425; found: 840.3430.

**$^1$H NMR** (400 MHz, $CDCl_3$) δ 8.90 – 8.83 (m, 2H), 8.48 (s, 1H), 8.19 (dd, *J* = 5.7, 0.6 Hz, 1H), 8.15 (bs, 1H), 8.03 – 7.96 (m, 2H), 7.66 (d, *J* = 8.5 Hz, 1H), 7.54 – 7.42 (m, 4H), 7.26 – 7.25 (m, 1H), 7.02 (dd, *J* = 8.6, 2.4 Hz, 1H), 6.58 (d, *J* = 5.7 Hz, 1H), 4.93 (dd, *J* = 12.3, 5.4 Hz, 1H), 4.15 (s, 2H), 4.00 (t, *J* = 6.0 Hz, 2H), 3.97 – 3.86 (m, 1H), 3.77 – 3.61 (m, 5H), 3.39 (ddd, *J* = 12.6, 8.1, 3.6 Hz, 1H), 3.33 – 3.21 (m, 3H), 2.92 – 2.64 (m, 3H), 2.48 (t, *J* = 7.1 Hz, 2H), 2.19 – 2.07 (m, 3H), 1.98 – 1.87 (m, 2H), 1.87 – 1.77 (m, 5H), 1.74 – 1.63 (m, 2H), 1.63 – 1.52 (m, 2H) ppm.

**$^{13}$C NMR** (101 MHz, $CDCl_3$) δ 171.1, 170.4, 168.4, 168.1, 167.3, 163.2, 156.4, 155.2, 148.1, 135.1, 134.5, 131.9, 129.3, 128.9, 128.9, 127.1, 126.7, 125.6, 125.4, 120.8, 119.0, 118.0, 108.7, 105.3, 71.2, 71.0, 67.4, 49.3, 45.4, 45.3, 42.8, 41.1, 39.1, 32.5, 31.6, 31.4, 31.2, 31.0, 29.3, 24.8, 22.9, 10.7 ppm.

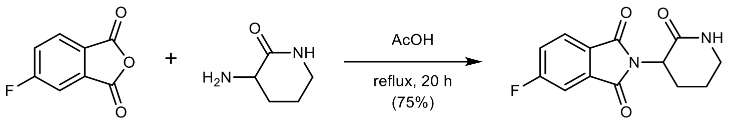

**S43**

**5-Fluoro-2-(2-oxopiperidin-3-yl)isoindoline-1,3-dione (S43):** To a solution of 5-fluoroisobenzofuran-1,3-dione (200 mg, 1.20 mmol, 1.00 equiv.) in acetic acid (6.0 mL) was added 3-aminopiperidin-2-one (144 mg, 1.26 mmol, 1.05 equiv.) at rt. The reaction was heated to reflux and stirred for 20 h. The mixture was concentrated *in vacuo*. The crude product was suspended in ice-cold water (5 mL), collected by filtration using a sintered glass funnel, washed with ice-cold water (5 mL), and dried *in vacuo*, yielding 237 mg (0.90 mmol, 75%) of **S43** as a light-brown solid.

**TLC** $R_f$ = 0.34 (5% MeOH in $CH_2Cl_2$).

**LC/MS** (ESI) *m/z*: $(M+H)^+$ 263.1.

**$^1$H NMR** (400 MHz, $CDCl_3$) δ 7.85 (ddd, *J* = 8.3, 4.5, 0.5 Hz, 1H), 7.52 (ddd, *J* = 7.0, 2.3, 0.5 Hz, 1H), 7.38 (ddd, *J* = 8.9, 8.2, 2.3 Hz, 1H), 6.17 (bs, 1H), 4.75 (dd, *J* = 12.0, 6.2 Hz, 1H), 3.58 – 3.46 (m, 1H), 3.39 (dddt, *J* = 11.3, 5.5, 3.8, 2.1 Hz, 1H), 2.47 – 2.32 (m, 1H), 2.19 – 2.03 (m, 2H), 2.03 – 1.88 (m, 1H) ppm.

**$^{19}$F{$^1$H} NMR** (377 MHz, $CDCl_3$) δ –101.8 ppm.

**$^{13}$C NMR** (101 MHz, $CDCl_3$) δ 168.0, 166.8, 166.6 (d, *J* = 257.1 Hz), 166.5 (d, *J* = 2.8 Hz), 135.0 (d, *J* = 9.5 Hz), 128.0 (d, *J* = 2.8 Hz), 126.0 (d, *J* = 9.3 Hz), 121.2 (d, *J* = 23.7 Hz), 111.4 (d, *J* = 24.8 Hz), 49.6, 42.6, 26.4, 22.4 ppm.

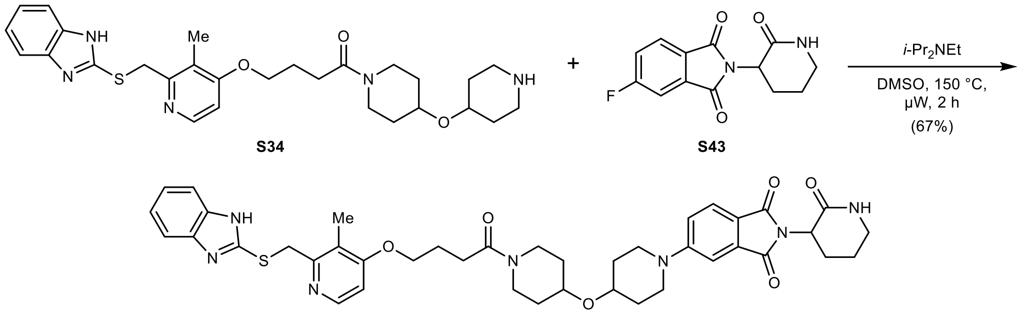

**5-(4-((1-(4-((2-(((1*H*-Benzo[*d*]imidazol-2-yl)thio)methyl)-3-methylpyridin-4-yl)oxy)butanoyl) piperidin-4-yl)oxy)piperidin-1-yl)-2-(2-oxopiperidin-3-yl)isoindoline-1,3-dione (19):** To a solution of **S34** (45 mg, 0.09 mmol, 1.00 equiv.) in anhydrous DMSO (0.6 mL) was added **S43** (24 mg, 0.09 mmol, 1.05 equiv.) and *i*-$Pr_2NEt$ (30 μL, 0.17 mmol, 2.00 equiv.) at rt under argon. The reaction was stirred at 150 °C for 2 h under microwave irradiation. The mixture was diluted with water (100 mL) and extracted with EtOAc (3 × 25 mL). The combined organic layers were washed with water (50 mL) and brine (50 mL), dried with $MgSO_4$, filtered, and concentrated *in vacuo*. Purification by column chromatography (12 g silica, 0 to 9% MeOH with 2.5% $NH_4OH$ in $CH_2Cl_2$ in 15 min) yielded 46 mg (0.06 mmol, 67%) of **19** as a yellow powder after lyophilization from MeCN/$H_2O$.

**TLC** $R_f$ = 0.51 (10% MeOH with 0.5% $NH_4OH$ in $CH_2Cl_2$).

**HRMS** (ESI) *m/z*: $(M+H)^+$ calc. for $C_{41}H_{48}N_7O_6S$: 766.3381; found: 766.3400.

**$^1$H NMR** (400 MHz, $CDCl_3$) δ 8.34 (d, *J* = 5.7 Hz, 1H), 7.63 (d, *J* = 8.5 Hz, 1H), 7.59 – 7.47 (m, 2H), 7.24 (d, *J* = 2.3 Hz, 1H), 7.22 – 7.13 (m, 2H), 7.00 (dd, *J* = 8.6, 2.4 Hz, 1H), 6.79 (d, *J* = 5.8 Hz, 1H), 5.96 (bs, 1H), 4.72 (dd, *J* = 12.0, 6.2 Hz, 1H), 4.38 (s, 2H), 4.13 (t, *J* = 6.0 Hz, 2H), 3.89 (ddd, *J* = 11.8, 7.2, 3.8 Hz, 1H), 3.75 – 3.61 (m, 5H), 3.52 (td, *J* = 11.7, 4.2 Hz, 1H), 3.45 – 3.33 (m, 2H), 3.33 – 3.16 (m, 3H), 2.53 (t, *J* = 7.0 Hz, 2H), 2.47 – 2.32 (m, 1H), 2.26 (s, 3H), 2.19 (p, *J* = 6.6 Hz, 2H), 2.14 – 2.01 (m, 2H), 2.01 – 1.85 (m, 3H), 1.85 – 1.73 (m, 2H), 1.73 – 1.61 (m, 2H), 1.61 – 1.50 (m, 2H) ppm. **Note:** The benzimidazole NH peak is in exchange with residual water in the spectrum.

**$^{13}$C NMR** (101 MHz, $CDCl_3$) δ 170.1, 168.4, 168.4, 167.8, 164.3, 156.6, 155.0, 151.6, 147.3, 134.7, 125.1, 121.8, 121.0, 119.7, 117.8, 108.7, 106.3, 71.1, 71.0, 67.8, 49.1, 45.4, 45.4, 42.6, 42.5, 39.0, 34.9, 32.4, 31.3, 31.1, 30.9, 29.0, 26.5, 24.5, 22.4, 10.8 ppm. **Note:** C1, C2, C3, and C6 are not detected due to benzimidazole tautomerism.

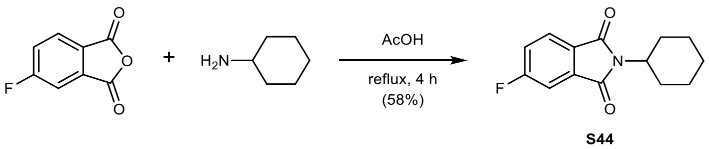

**2-Cyclohexyl-5-fluoroisoindoline-1,3-dione (S44):** To a solution of 5-fluoroisobenzofuran-1,3-dione (200 mg, 1.20 mmol, 1.00 equiv.) in acetic acid (6.0 mL) was added cyclohexylamine (138 μL, 1.20 mmol, 1.00 equiv.) at rt. The mixture was heated to reflux and stirred for 4 h. The mixture was diluted with water and cooled to 0 °C. An off-white precipitate formed which was collected by filtration using a sintered glass funnel, washed with ice-cold water (5 mL), and dried *in vacuo*, yielding 173 mg (0.70 mmol, 58%) of **S44** as an off-white powder.

**TLC** $R_f$ = 0.83 (30% EtOAc in *n*-heptane).

**LC/MS** (ESI) *m/z*: $(M+H)^+$ 248.1.

**$^1$H NMR** (400 MHz, $CDCl_3$) δ 7.81 (ddd, *J* = 8.2, 4.5, 0.5 Hz, 1H), 7.48 (ddd, *J* = 7.0, 2.3, 0.5 Hz, 1H), 7.35 (ddd, *J* = 8.9, 8.2, 2.3 Hz, 1H), 4.09 (tt, *J* = 12.3, 3.9 Hz, 1H), 2.19 (qd, *J* = 12.5, 3.5 Hz, 2H), 1.94 – 1.82 (m, 2H), 1.79 – 1.66 (m, 3H), 1.45 – 1.19 (m, 3H) ppm.

**$^{19}$F{$^1$H} NMR** (377 MHz, $CDCl_3$) δ –102.5 ppm.

**$^{13}$C NMR** (101 MHz, $CDCl_3$) δ 167.5, 167.2 (d, *J* = 2.6 Hz), 166.5 (d, *J* = 256.4 Hz), 135.1 (d, *J* = 9.2 Hz), 128.0, 125.5 (d, *J* = 9.3 Hz), 120.9 (d, *J* = 23.6 Hz), 111.0 (d, *J* = 24.8 Hz), 51.4, 30.0, 26.1, 25.2 ppm.

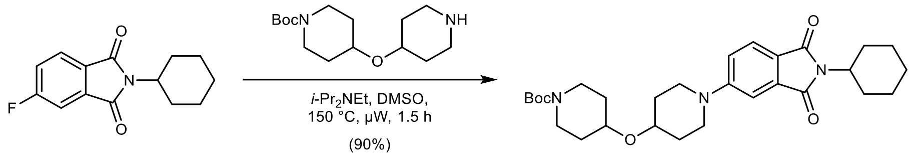

***tert*-Butyl 4-((1-(2-cyclohexyl-1,3-dioxoisoindolin-5-yl)piperidin-4-yl)oxy)piperidine-1-carboxylate (S45):** To a solution of **S44** (73 mg, 0.30 mmol, 1.00 equiv.) in anhydrous DMSO (2.0 mL) was added *tert*-butyl 4-(piperidin-4-yloxy)piperidine-1-carboxylate (88 mg, 0.31 mmol, 1.05 equiv.) and *i*-$Pr_2NEt$ (154 µL, 0.89 mmol, 3.00 equiv.) at rt under argon. The reaction was stirred at 150 °C for 1.5 h under microwave irradiation. The mixture was diluted with water (100 mL) and extracted with EtOAc (3 × 25 mL). The combined organic layers were washed with brine (50 mL), dried with $MgSO_4$, filtered, and concentrated *in vacuo*. Purification by column chromatography (12 g silica, 5 to 45% EtOAc in *n*-heptane in 12 min) yielded 136 mg (0.27 mmol, 90%) of **S45** as a yellow solid.

**TLC** $R_f$ = 0.50 (50% EtOAc in *n*-heptane).

**LC/MS** (ESI) *m/z*: $(M+H)^+$ 512.3.

**$^1$H NMR** (400 MHz, $CDCl_3$) δ 7.60 (dd, *J* = 8.4, 0.4 Hz, 1H), 7.23 (d, *J* = 2.1 Hz, 1H), 7.00 (dd, *J* = 8.5, 2.4 Hz, 1H), 4.04 (tt, *J* = 12.3, 3.8 Hz, 1H), 3.83 – 3.65 (m, 5H), 3.60 (tt, *J* = 7.9, 3.7 Hz, 1H), 3.22 (ddd, *J* = 12.6, 8.5, 3.4 Hz, 2H), 3.11 (ddd, *J* = 13.0, 9.0, 3.5 Hz, 2H), 2.18 (qd, *J* = 12.4, 3.4 Hz, 2H), 1.99 – 1.88 (m, 2H), 1.88 – 1.75 (m, 4H), 1.75 – 1.64 (m, 5H), 1.59 – 1.48 (m, 2H), 1.45 (s, 9H), 1.43 – 1.20 (m, 3H) ppm.

**$^{13}$C NMR** (101 MHz, $CDCl_3$) δ 169.2, 168.8, 155.0, 155.0, 134.8, 124.7, 119.9, 117.7, 108.4, 79.6, 71.9, 71.1, 50.7, 45.7, 41.4, 31.9, 31.2, 30.1, 28.6, 26.2, 25.3 ppm.

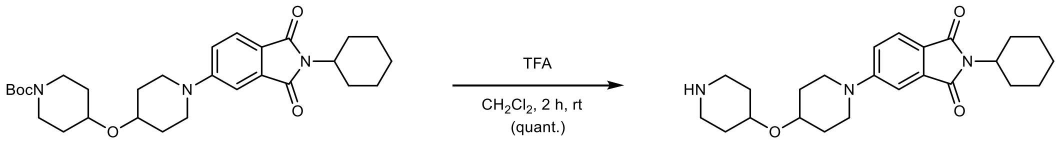

**2-Cyclohexyl-5-(4-(piperidin-4-yloxy)piperidin-1-yl)isoindoline-1,3-dione (S46 · TFA):** To a solution of **S45** (136 mg, 0.27 mmol, 1.00 equiv.) in $CH_2Cl_2$ (2.0 mL) was added trifluoroacetic acid (1.0 mL, 13.3 mmol, 50.0 equiv.) slowly at 0 °C. After 5 min, the reaction was allowed to warm to rt and stirred for 2 h. The mixture was concentrated *in vacuo* – residual trifluoroacetic acid was removed by co-evaporation with MeOH (3 × 5 mL), toluene (3 × 5 mL), and drying *in vacuo*, yielding 140 mg (0.27 mmol, quantitative) of **S46 · TFA** as a yellow solid.

**TLC** $R_f$ = 0.36, streaky (20% MeOH with 0.5% $NH_4OH$ in $CH_2Cl_2$).

**LC/MS** (ESI) *m/z*: $(M+H)^+$ 412.2.

**$^{1}$H NMR** (400 MHz, $CD_3OD$) δ 7.59 (dd, $J$ = 8.5, 0.4 Hz, 1H), 7.28 (d, $J$ = 2.3 Hz, 1H), 7.17 (dd, $J$ = 8.5, 2.4 Hz, 1H), 4.07 – 3.97 (m, 1H), 3.88 (tt, $J$ = 6.5, 3.2 Hz, 1H), 3.84 – 3.71 (m, 3H), 3.40 – 3.22 (m, 4H), 3.13 (ddd, $J$ = 12.5, 7.0, 4.0 Hz, 2H), 2.18 (qd, $J$ = 12.5, 3.6 Hz, 2H), 2.08 – 1.93 (m, 4H), 1.93 – 1.80 (m, 4H), 1.77 – 1.61 (m, 5H), 1.46 – 1.17 (m, 3H) ppm. **Note:** The NH peaks are in exchange with $CD_3OD$ in the spectrum.

**$^{13}$C NMR** (101 MHz, $CD_3OD$) δ 170.3, 170.1, 156.6, 135.8, 125.6, 120.4, 119.0, 109.0, 73.2, 68.8, 51.9, 46.4, 42.1, 32.2, 31.0, 29.5, 27.2, 26.4 ppm.

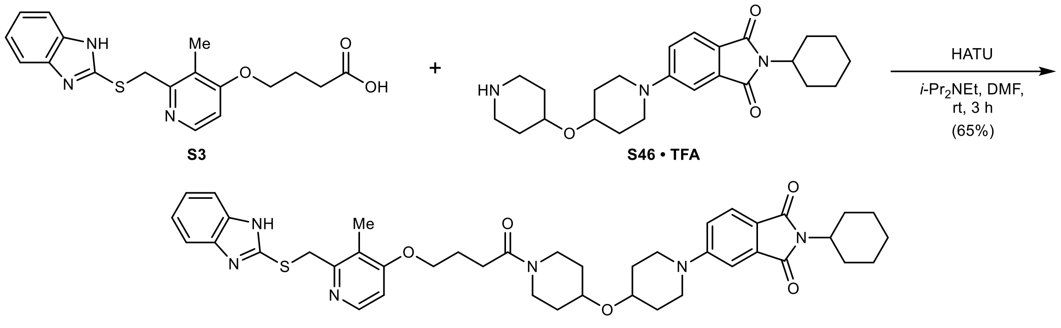

**5-(4-((1-(4-((2-(((1*H*-Benzo[*d*]imidazol-2-yl)thio)methyl)-3-methylpyridin-4-yl)oxy)butanoyl) piperidin-4-yl)oxy)piperidin-1-yl)-2-cyclohexylisoindoline-1,3-dione (20):** To a solution of **S46 · TFA** (49 mg, 0.09 mmol, 1.00 equiv.) in anhydrous DMF (1.0 mL) was added **S3** (35 mg, 0.10 mmol, 1.05 equiv.), HATU (42 mg, 0.11 mmol, 1.20 equiv.), and *i*-$Pr_2NEt$ (64 µL, 0.37 mmol, 4.00 equiv.) at rt under argon. The reaction was stirred at rt for 3 h. The mixture was poured into water (100 mL) and extracted with EtOAc (3 × 25 mL). The combined organic layers were washed with 5% aqueous LiCl (25 mL), water (25 mL), and brine (25 mL), dried with $MgSO_4$, filtered, and concentrated *in vacuo*. Purification by column chromatography (12 g silica, 0.5 to 8.5% MeOH with 2.5% $NH_4OH$ in $CH_2Cl_2$ in 10 min) yielded 45 mg (0.06 mmol, 65%) of **20** as a yellow solid.

**TLC** $R_f$ = 0.57 (10% MeOH with 0.5% $NH_4OH$ in $CH_2Cl_2$).

**HRMS** (ESI) *m/z*: $(M+H)^+$ calc. for $C_{42}H_{51}N_6O_5S$: 751.3636; found: 751.3643.

**$^{1}$H NMR** (400 MHz, $CDCl_3$) δ 8.34 (dd, $J$ = 5.8, 0.6 Hz, 1H), 7.60 (d, $J$ = 8.4 Hz, 1H), 7.56 – 7.49 (m, 2H), 7.22 (d, $J$ = 2.3 Hz, 1H), 7.20 – 7.14 (m, 2H), 6.99 (dd, $J$ = 8.5, 2.4 Hz, 1H), 6.79 (d, $J$ = 5.8 Hz, 1H), 4.37 (s, 2H), 4.12 (t, $J$ = 6.0 Hz, 2H), 4.04 (tt, $J$ = 12.3, 3.8 Hz, 1H), 3.89 (ddd, $J$ = 11.9, 7.3, 3.8 Hz, 1H), 3.75 – 3.61 (m, 5H), 3.38 (ddd, $J$ = 12.6, 8.0, 3.6 Hz, 1H), 3.33 – 3.15 (m, 3H), 2.53 (t, $J$ = 7.0 Hz, 2H), 2.25 (s, 3H), 2.24 – 2.11 (m, 4H), 1.96 – 1.75 (m, 6H), 1.75 – 1.61 (m, 5H), 1.61 – 1.50 (m, 2H), 1.43 – 1.21 (m, 3H) ppm. **Note:** The benzimidazole NH peak is in exchange with residual water in the spectrum.

**$^{13}$C NMR** (101 MHz, $CDCl_3$) δ 170.1, 169.1, 168.6, 164.3, 156.6, 154.9, 151.7, 147.3, 139.4, 134.7, 124.6, 121.8, 121.0, 119.9, 117.7, 114.3, 108.3, 106.3, 71.2, 71.0, 67.8, 50.6, 45.5, 45.5, 42.6, 39.0, 34.9, 32.4, 31.3, 31.2, 31.0, 30.0, 29.0, 26.1, 25.2, 24.5, 10.8 ppm.

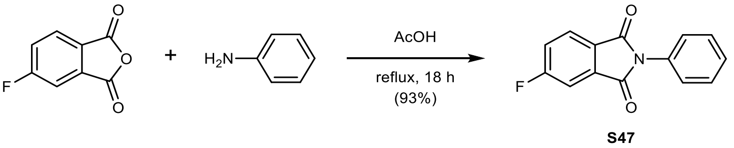

**5-Fluoro-2-phenylisoindoline-1,3-dione (S47):** To a solution of 5-fluoroisobenzofuran-1,3-dione (200 mg, 1.20 mmol, 1.00 equiv.) in acetic acid (4.0 mL) was added aniline (115 µL,

1.26 mmol, 1.05 equiv.) at rt. The reaction was heated to reflux and stirred for 18 h. The mixture was diluted with water and cooled to 0 °C. An off-white precipitate formed which was collected by filtration using a sintered glass funnel, washed with ice-cold water (5 mL), and dried *in vacuo*, yielding 270 mg (1.12 mmol, 93%) of **S47** as an off-white powder.

**TLC** $R_f$ = 0.51 (30% EtOAc in *n*-heptane).

**LC/MS** (ESI) *m/z*: $(M+H)^+$ 242.1.

**$^1$H NMR** (400 MHz, $CDCl_3$) δ 7.97 (ddd, *J* = 8.2, 4.5, 0.5 Hz, 1H), 7.63 (ddd, *J* = 7.0, 2.3, 0.5 Hz, 1H), 7.57 – 7.37 (m, 6H) ppm.

**$^{19}$F{$^1$H} NMR** (377 MHz, $CDCl_3$) δ –101.1 ppm.

**$^{13}$C NMR** (101 MHz, $CDCl_3$) δ 166.8 (d, *J* = 257.5 Hz), 166.4, 166.1 (d, *J* = 2.9 Hz), 134.8 (d, *J* = 9.4 Hz), 131.6, 129.3, 128.4, 127.7 (d, *J* = 2.8 Hz), 126.6, 126.3 (d, *J* = 9.3 Hz), 121.7 (d, *J* = 23.7 Hz), 111.7 (d, *J* = 24.9 Hz) ppm.

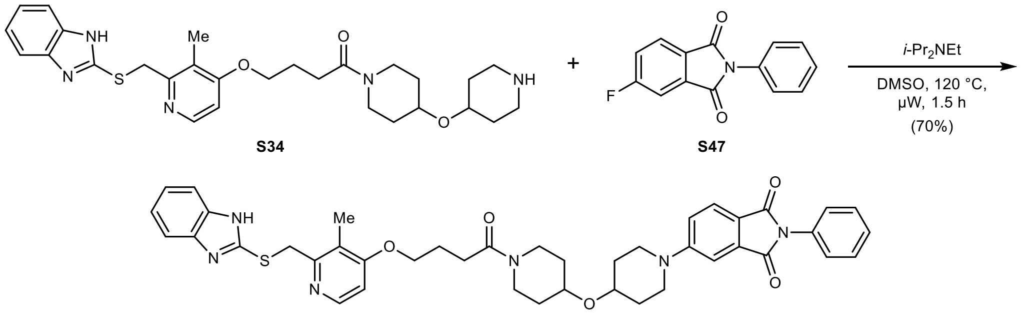

**5-(4-((1-(4-((2-(((1*H*-Benzo[*d*]imidazol-2-yl)thio)methyl)-3-methylpyridin-4-yl)oxy)butanoyl) piperidin-4-yl)oxy)piperidin-1-yl)-2-phenylisoindoline-1,3-dione (21):** To a solution of **S34** (40 mg, 0.08 mmol, 1.00 equiv.) in anhydrous DMSO (0.5 mL) was added **S47** (19 mg, 0.08 mmol, 1.00 equiv.) and *i*-$Pr_2NEt$ (40 μL, 0.23 mmol, 3.00 equiv.) at rt under argon. The reaction was stirred at 120 °C for 30 min under microwave irradiation. To drive the reaction to completion, **S47** (4.0 mg, 0.02 mmol, 0.20 equiv.) was added. The reaction was stirred at 120 °C for another 1 h under microwave irradiation. The mixture was diluted with water (100 mL) and extracted with EtOAc (3 × 25 mL). The combined organic layers were washed with water (50 mL) and brine (50 mL), dried with $MgSO_4$, filtered, and concentrated *in vacuo*. Purification by column chromatography (12 g silica, 0 to 6% MeOH with 2.5% $NH_4OH$ in $CH_2Cl_2$ in 10 min) yielded 40 mg (0.05 mmol, 70%) of **21** as a yellow powder after lyophilization from MeCN/$H_2O$.

**TLC** $R_f$ = 0.50 (10% MeOH in $CH_2Cl_2$).

**HRMS** (ESI) *m/z*: $(M+H)^+$ calc. for $C_{42}H_{45}N_6O_5S$: 745.3167; found: 745.3184.

**$^1$H NMR** (400 MHz, $CDCl_3$) δ 8.35 (dd, *J* = 5.8, 0.6 Hz, 1H), 7.75 (d, *J* = 8.5 Hz, 1H), 7.58 – 7.46 (m, 4H), 7.46 – 7.33 (m, 4H), 7.22 – 7.13 (m, 2H), 7.08 (dd, *J* = 8.6, 2.4 Hz, 1H), 6.79 (d, *J* = 5.8 Hz, 1H), 4.38 (s, 2H), 4.13 (t, *J* = 6.0 Hz, 2H), 3.91 (ddd, *J* = 11.5, 7.1, 3.8 Hz, 1H), 3.77 – 3.64 (m, 5H), 3.39 (ddd, *J* = 12.6, 8.1, 3.6 Hz, 1H), 3.34 – 3.23 (m, 3H), 2.54 (t, *J* = 7.0 Hz, 2H), 2.26 (s, 3H), 2.20 (p, *J* = 6.5 Hz, 2H), 2.00 – 1.87 (m, 2H), 1.87 – 1.76 (m, 2H), 1.76 – 1.64 (m, 2H), 1.64 – 1.51 (m, 2H) ppm. **Note:** The benzimidazole NH peak is in exchange with residual water in the spectrum.

**$^{13}$C NMR** (101 MHz, $CDCl_3$) δ 170.2, 168.1, 167.5, 164.4, 156.8, 155.3, 151.8, 147.5, 134.6, 132.3, 129.1, 127.9, 126.7, 125.5, 122.0, 121.1, 119.2, 118.2, 108.7, 106.4, 71.2, 71.2, 67.9, 45.5, 45.4, 42.8, 39.1, 35.0, 32.5, 31.5, 31.3, 31.1, 29.1, 24.6, 10.9 ppm. **Note:** C1, C2, C3, and C6 are not detected due to benzimidazole tautomerism.

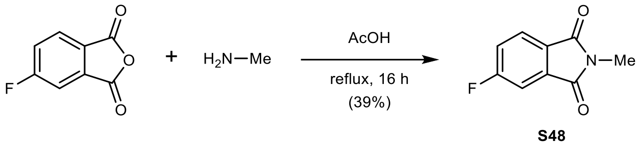

**5-Fluoro-2-methylisoindoline-1,3-dione (S48)**: To a solution of 5-fluoroisobenzofuran-1,3-dione (200 mg, 1.20 mmol, 1.00 equiv.) in acetic acid (5.0 mL) was added molecular sieves (4 Å) and methanamine (0.125 mL, 40% in $H_2O$, 1.44 mmol, 1.20 equiv.) at rt. The reaction was sealed and heated to reflux for 16 h. The mixture was concentrated *in vacuo*. The crude product was suspended in water (5 mL), collected by filtration using a sintered glass funnel, washed with water (5 mL), and dried *in vacuo*, yielding 84 mg (0.47 mmol, 39%) of **S48** as a colorless solid. **Note:** The reaction was difficult to monitor by TLC or LC/MS and might not have gone to completion. Methanamine is volatile.

**TLC** $R_f$ = 0.38 (20% EtOAc in *n*-heptane).

**LC/MS** (ESI) *m/z*: $(M+H)^+$ 180.1.

**$^1$H NMR** (400 MHz, $CDCl_3$) δ 7.85 (ddd, *J* = 8.2, 4.5, 0.5 Hz, 1H), 7.52 (ddd, *J* = 7.0, 2.3, 0.5 Hz, 1H), 7.37 (ddd, *J* = 8.9, 8.2, 2.3 Hz, 1H), 3.18 (s, 3H) ppm.

**$^{19}$F{$^1$H} NMR** (377 MHz, $CDCl_3$) δ –102.1 ppm.

**$^{13}$C NMR** (101 MHz, $CDCl_3$) δ 167.5, 167.2 (d, *J* = 2.8 Hz), 166.5 (d, *J* = 256.5 Hz), 135.2 (d, *J* = 9.4 Hz), 128.2 (d, *J* = 3.0 Hz), 125.7 (d, *J* = 9.4 Hz), 121.0 (d, *J* = 23.6 Hz), 111.3 (d, *J* = 25.0 Hz), 24.3 ppm.

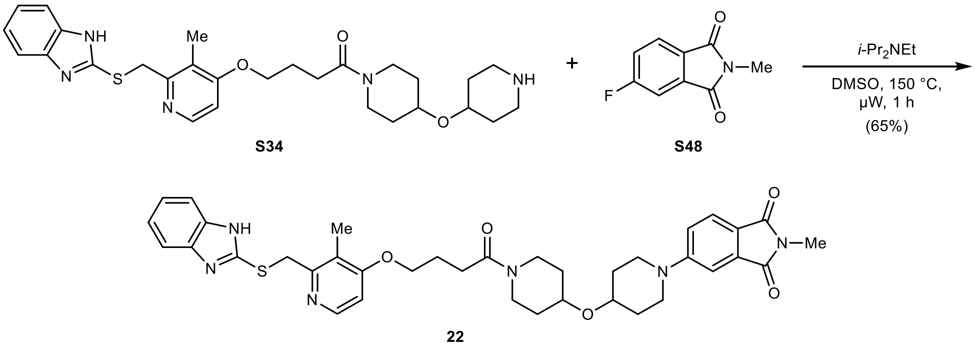

**5-(4-((1-(4-((2-(((1*H*-Benzo[*d*]imidazol-2-yl)thio)methyl)-3-methylpyridin-4-yl)oxy)butanoyl) piperidin-4-yl)oxy)piperidin-1-yl)-2-methylisoindoline-1,3-dione** (**22**): To a solution of **S34** (45 mg, 0.09 mmol, 1.00 equiv.) in anhydrous DMSO (0.6 mL) was added **S48** (16 mg, 0.09 mmol, 1.05 equiv.) and *i*-$Pr_2NEt$ (30 µL, 0.17 mmol, 2.00 equiv.) at rt under argon. The reaction was stirred at 150 °C for 1 h under microwave irradiation. The mixture was diluted with water (100 mL) and extracted with EtOAc (3 × 25 mL). The combined organic layers were washed with water (50 mL) and brine (50 mL), dried with $MgSO_4$, filtered, and concentrated *in vacuo*. Purification by column chromatography (12 g silica, 1 to 9% MeOH with 2.5% $NH_4OH$ in $CH_2Cl_2$ in 12 min) yielded 38 mg (0.06 mmol, 65%) of **22** as a yellow powder after lyophilization from MeCN/$H_2O$.

**TLC** $R_f$ = 0.65 (10% MeOH with 0.5% $NH_4OH$ in $CH_2Cl_2$).

**HRMS** (ESI) *m/z*: $(M+H)^+$ calc. for $C_{37}H_{43}N_6O_5S$: 683.3010; found: 683.3021.

**$^1$H NMR** (400 MHz, $CDCl_3$) δ 8.34 (d, *J* = 5.7 Hz, 1H), 7.63 (d, *J* = 8.4 Hz, 1H), 7.59 – 7.48 (m, 2H), 7.26 – 7.25 (m, 1H), 7.22 – 7.13 (m, 2H), 6.99 (dd, *J* = 8.5, 2.4 Hz, 1H), 6.79 (d, *J* = 5.8 Hz, 1H), 4.37 (s, 2H), 4.13 (t, *J* = 6.0 Hz, 2H), 3.90 (ddd, *J* = 11.4, 7.1, 3.8 Hz, 1H), 3.76 – 3.60 (m, 5H), 3.38 (ddd, *J* = 12.6, 8.0, 3.6 Hz, 1H), 3.33 – 3.18 (m, 3H), 3.12 (s, 3H), 2.53 (t, *J* = 6.9 Hz, 2H), 2.26 (s,

3H), 2.19 (p, *J* = 6.5 Hz, 2H), 1.96 – 1.86 (m, 2H), 1.86 – 1.75 (m, 2H), 1.75 – 1.61 (m, 2H), 1.61 – 1.51 (m, 2H) ppm. **Note:** The benzimidazole NH peak is in exchange with residual water in the spectrum.

**$^{13}$C NMR** (101 MHz, $CDCl_3$) δ 170.2, 169.3, 168.8, 164.4, 156.7, 155.0, 151.8, 147.4, 135.0, 124.9, 121.9, 121.1, 119.9, 117.6, 108.6, 106.4, 71.2, 71.1, 67.9, 45.5, 45.5, 42.7, 39.1, 35.0, 32.5, 31.4, 31.3, 31.1, 29.1, 24.6, 23.9, 10.9 ppm. **Note:** C1, C2, C3, and C6 are not detected due to benzimidazole tautomerism.

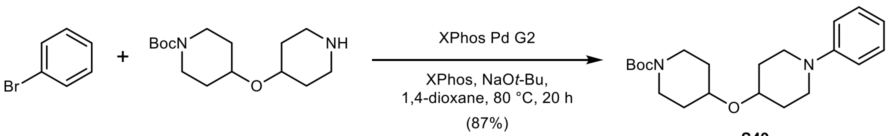

**S49**

***tert*-Butyl 4-((1-phenylpiperidin-4-yl)oxy)piperidine-1-carboxylate (S49):** To a mixture of *tert*-butyl 4-(piperidin-4-yloxy)piperidine-1-carboxylate (132 mg, 0.46 mmol, 1.00 equiv.), XPhos Pd G2 (18 mg, 0.02 mmol, 0.05 equiv.), XPhos (11 mg, 0.02 mmol, 0.05 equiv.), and NaO*t*-Bu (134 mg, 1.39 mmol, 3.00 equiv.) in anhydrous, degassed 1,4-dioxane (2.0 mL) was added bromobenzene (53 µL, 0.51 mmol, 1.10 equiv.) at rt under argon. The mixture was heated to 80 °C for 1.5 h. To drive the reaction to completion, bromobenzene (53 µL, 0.51 mmol, 1.10 equiv.) was added and the reaction was stirred at 80 °C for 18 h. The mixture was filtered through Celite on a sintered glass funnel, rinsed with EtOAc (20 mL), and concentrated *in vacuo*. Purification by column chromatography (12 g silica, 5 to 35% EtOAc in *n*-heptane in 10 min) yielded 146 mg (0.41 mmol, 87%) of **S49** as a colorless, waxy solid.

**TLC** $R_f$ = 0.68 (50% EtOAc in *n*-heptane).

**LC/MS** (ESI) *m/z*: $(M+H)^+$ 361.2.

**$^1$H NMR** (400 MHz, $CDCl_3$) δ 7.28 – 7.21 (m, 2H), 6.98 – 6.90 (m, 2H), 6.83 (tt, *J* = 7.3, 1.1 Hz, 1H), 3.87 – 3.69 (m, 2H), 3.67 – 3.48 (m, 4H), 3.09 (ddd, *J* = 13.1, 9.2, 3.5 Hz, 2H), 2.92 (ddd, *J* = 12.6, 9.6, 3.2 Hz, 2H), 1.95 (ddd, *J* = 11.4, 5.9, 3.3 Hz, 2H), 1.87 – 1.76 (m, 2H), 1.71 (dtd, *J* = 12.9, 9.1, 3.8 Hz, 2H), 1.56 – 1.49 (m, 2H), 1.46 (s, 9H) ppm.

**$^{13}$C NMR** (101 MHz, $CDCl_3$) δ 155.0, 151.5, 129.2, 119.5, 116.6, 79.6, 72.0, 71.8, 47.5, 41.5, 31.9, 31.9, 28.6 ppm.

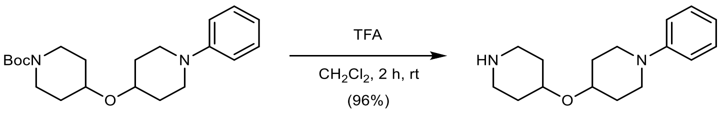

**S49** **S50**

**1-Phenyl-4-(piperidin-4-yloxy)piperidine (S50):** To a solution of **S49** (146 mg, 0.41 mmol, 1.00 equiv.) in $CH_2Cl_2$ (2.0 mL) was added trifluoroacetic acid (1.0 mL, 13.4 mmol, 33.0 equiv.) slowly at 0 °C. After 5 min, the reaction was allowed to warm to rt and stirred for 2 h. The mixture was poured into saturated aqueous $Na_2CO_3$ (50 mL) and extracted with 10% MeOH in $CH_2Cl_2$ (3 × 25 mL). The combined organic layers were dried with $MgSO_4$, filtered, and concentrated *in vacuo*, yielding 101 mg (0.39 mmol, 96%) of **S50** as a light-yellow solid.

**TLC** $R_f$ = 0.08, streaky (20% MeOH with 0.5% $NH_4OH$ in $CH_2Cl_2$).

**LC/MS** (ESI) *m/z*: $(M+H)^+$ 261.2.

**$^1$H NMR** (400 MHz, $CD_3OD$) δ 7.26 – 7.17 (m, 2H), 7.01 – 6.94 (m, 2H), 6.82 (tt, *J* = 7.4, 1.0 Hz, 1H), 3.69 – 3.56 (m, 2H), 3.55 – 3.45 (m, 2H), 3.08 – 2.98 (m, 2H), 2.89 (ddd, *J* = 12.7, 9.8, 3.1 Hz, 2H), 2.61 (ddd, *J* = 13.0, 10.1, 3.0 Hz, 2H), 2.02 – 1.93 (m, 2H), 1.93 – 1.85 (m, 2H), 1.72 – 1.60 (m, 2H), 1.52 – 1.40 (m, 2H) ppm. **Note:** The NH peak is in exchange with $CD_3OD$ in the spectrum.

**$^{13}$C NMR** (101 MHz, $CD_3OD$) δ 152.3, 129.4, 120.3, 117.5, 72.8, 72.6, 48.4, 44.1, 33.3, 32.3 ppm.

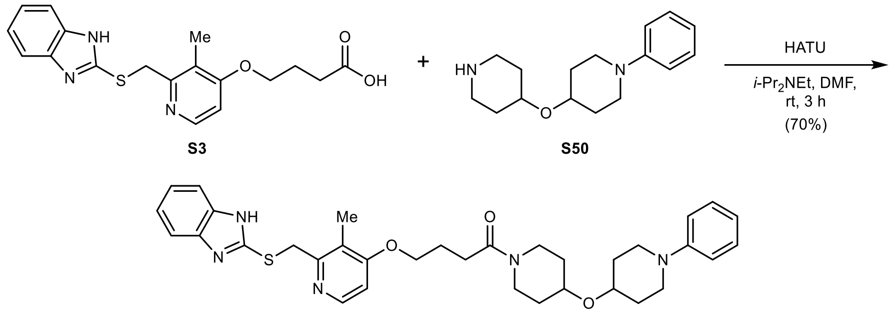

23

**4-((2-(((1*H*-Benzo[*d*]imidazol-2-yl)thio)methyl)-3-methylpyridin-4-yl)oxy)-1-(4-((1-phenylpiperidin-4-yl)oxy)piperidin-1-yl)butan-1-one (23):** To a solution of **S50** (33 mg, 0.13 mmol, 1.00 equiv.) in anhydrous DMF (1.0 mL) was added **S3** (47 mg, 0.13 mmol, 1.05 equiv.), HATU (58 mg, 0.15 mmol, 1.20 equiv.), and *i*-$Pr_2NEt$ (66 µL, 0.38 mmol, 3.00 equiv.) at rt under argon. The reaction was stirred at rt for 3 h. The mixture was poured into water (150 mL) and extracted with EtOAc (3 × 50 mL). The combined organic layers were washed with 5% aqueous LiCl (50 mL), water (50 mL), and brine (50 mL), dried with $MgSO_4$, filtered, and concentrated *in vacuo*. Purification by column chromatography (12 g silica, 0 to 7.5% MeOH with 2.5% $NH_4OH$ in $CH_2Cl_2$ in 10 min) yielded 53 mg (0.09 mmol, 70%) of **23** as an off-white solid.

**TLC** $R_f$ = 0.45 (10% MeOH in $CH_2Cl_2$).

**HRMS** (ESI) *m/z*: $(M+H)^+$ calc. for $C_{34}H_{42}N_5O_3S$: 600.3003; found: 600.3003.

**$^1$H NMR** (400 MHz, $CDCl_3$) δ 8.34 (dd, *J* = 5.7, 0.6 Hz, 1H), 7.59 – 7.49 (m, 2H), 7.29 – 7.22 (m, 2H), 7.21 – 7.15 (m, 2H), 6.97 – 6.89 (m, 2H), 6.83 (tt, *J* = 7.3, 1.1 Hz, 1H), 6.79 (d, *J* = 5.8 Hz, 1H), 4.38 (s, 2H), 4.13 (t, *J* = 6.1 Hz, 2H), 3.92 (ddd, *J* = 12.1, 7.2, 3.8 Hz, 1H), 3.75 – 3.65 (m, 2H), 3.62 – 3.46 (m, 3H), 3.37 (ddd, *J* = 13.0, 8.1, 3.6 Hz, 1H), 3.27 (ddd, *J* = 12.9, 8.2, 3.5 Hz, 1H), 2.98 – 2.87 (m, 2H), 2.53 (t, *J* = 7.0 Hz, 2H), 2.26 (s, 3H), 2.19 (p, *J* = 6.6 Hz, 2H), 1.99 – 1.89 (m, 2H), 1.86 – 1.76 (m, 2H), 1.70 (dtt, *J* = 13.1, 8.7, 4.2 Hz, 2H), 1.64 – 1.51 (m, 2H) ppm. **Note:** The benzimidazole NH peak is in exchange with residual water in the spectrum.

**$^{13}$C NMR** (101 MHz, $CDCl_3$) δ 170.1, 164.3, 156.6, 151.7, 151.3, 147.3, 139.7, 129.1, 121.8, 121.0, 119.5, 116.5, 114.3, 106.3, 72.1, 70.8, 67.9, 47.3, 42.7, 39.1, 34.8, 32.4, 31.9, 31.7, 31.4, 29.1, 24.5, 10.8 ppm.

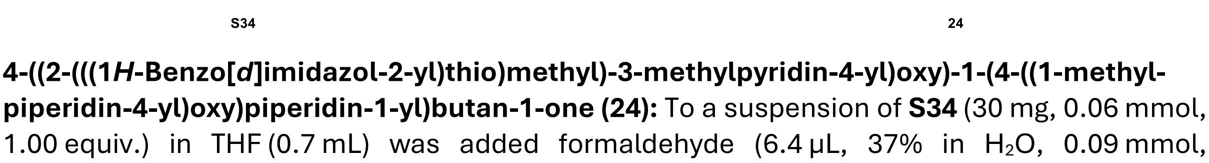

S34 24

**4-((2-(((1*H*-Benzo[*d*]imidazol-2-yl)thio)methyl)-3-methylpyridin-4-yl)oxy)-1-(4-((1-methylpiperidin-4-yl)oxy)piperidin-1-yl)butan-1-one (24):** To a suspension of **S34** (30 mg, 0.06 mmol, 1.00 equiv.) in THF (0.7 mL) was added formaldehyde (6.4 µL, 37% in $H_2O$, 0.09 mmol, 1.50 equiv.) and $NaBH(OAc)_3$ (24 mg, 0.11 mmol, 2.00 equiv.) at rt. The reaction was stirred at rt for 4 h. The mixture was poured into aqueous $NaHCO_3$ (50 mL) and extracted with EtOAc (3 × 25 mL). The combined organic layers were washed with brine (25 mL), dried with $MgSO_4$, filtered, and concentrated *in vacuo*. Purification by column chromatography (12 g silica, 0 to 20% MeOH with 2.5% $NH_4OH$ in $CH_2Cl_2$ in 16 min) yielded 19 mg (0.04 mmol, 62%) of **24** as a colorless solid.

**TLC** $R_f$ = 0.20 (10% MeOH with 0.5% $NH_4OH$ in $CH_2Cl_2$).

**HRMS** (ESI) *m/z*: $(M+H)^+$ calc. for $C_{29}H_{40}N_5O_3S$: 538.2846, found: 538.2846.

**$^1$H NMR** (600 MHz, DMSO-*d*$_6$) δ 12.61 (s, 1H), 8.23 (d, *J* = 5.6 Hz, 1H), 7.57 – 7.31 (m, 2H), 7.15 – 7.09 (m, 2H), 6.96 (d, *J* = 5.7 Hz, 1H), 4.69 (s, 2H), 4.08 (t, *J* = 6.4 Hz, 2H), 3.87 (dt, *J* = 11.4, 5.1 Hz, 1H), 3.68 – 3.58 (m, 2H), 3.44 – 3.36 (m, 1H), 3.17 (ddd, *J* = 13.2, 9.4, 3.3 Hz, 1H), 3.04 (ddd, *J* = 13.0, 9.4, 3.5 Hz, 1H), 2.64 – 2.55 (m, 2H), 2.48 (t, *J* = 7.4 Hz, 2H), 2.21 (s, 3H), 2.13 (s, 3H), 2.05 – 1.91 (m, 4H), 1.82 – 1.66 (m, 4H), 1.46 – 1.37 (m, 2H), 1.37 – 1.31 (m, 1H), 1.30 – 1.24 (m, 1H) ppm.

**$^{13}$C NMR** (101 MHz, DMSO-*d*$_6$) δ 169.7, 162.7, 154.7, 150.2, 147.8, 121.6, 119.7, 117.4, 110.4, 106.3, 71.0, 70.6, 67.4, 53.0, 45.7, 42.4, 38.7, 36.3, 32.1, 31.7, 31.5, 28.4, 24.3, 10.4 ppm.

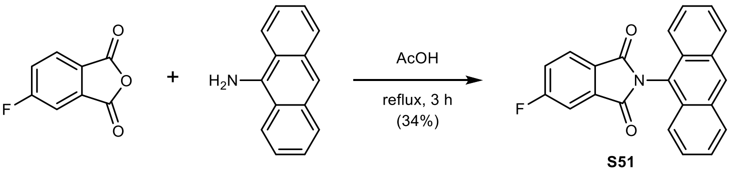

**2-(Anthracen-9-yl)-5-fluoroisoindoline-1,3-dione (S51):** To a solution of 5-fluoroisobenzo-furan-1,3-dione (80 mg, 0.48 mmol, 1.00 equiv.) in acetic acid (1.5 mL) was added anthracen-9-amine (98 mg, 0.51 mmol, 1.05 equiv.) at rt. The mixture was heated to reflux and stirred for 3 h. The mixture was diluted with water and cooled to 0 °C. A light-yellow precipitate formed which was collected by filtration using a sintered glass funnel, washed with ice-cold water (5 mL), and dried *in vacuo*. Residual water was removed by co-evaporation with benzene (2 × 5 mL). Purification by column chromatography (12 g silica, 0 to 30% EtOAc in *n*-heptane in 10 min) yielded 56 mg (0.16 mmol, 34%) of **S51** as a light-yellow, crystalline solid. **Note:** TLC suggested low purity of purchased anthracen-9-amine.

**TLC** $R_f$ = 0.30 (20% EtOAc in *n*-heptane).

**LC/MS** (ESI) *m/z*: $(M+H)^+$ 342.0.

**$^1$H NMR** (400 MHz, DMSO-*d*$_6$) δ 8.89 (s, 1H), 8.31 – 8.22 (m, 2H), 8.18 (dd, *J* = 8.3, 4.6 Hz, 1H), 8.02 (dd, *J* = 7.5, 2.3 Hz, 1H), 7.98 – 7.92 (m, 2H), 7.85 (ddd, *J* = 9.5, 8.3, 2.4 Hz, 1H), 7.65 – 7.53 (m, 4H) ppm.

**$^{19}$F{$^1$H} NMR** (377 MHz, DMSO-*d*$_6$) δ –102.5 ppm.

**$^{13}$C NMR** (101 MHz, DMSO-*d*$_6$) δ 167.0, 166.7 (d, *J* = 2.9 Hz), 166.2 (d, *J* = 253.8 Hz), 134.8 (d, *J* = 10.0 Hz), 131.1, 129.0, 128.9, 128.7, 128.0 (d, *J* = 2.5 Hz), 127.6, 126.7 (d, *J* = 9.8 Hz), 125.9, 123.3, 122.7, 121.8 (d, *J* = 23.8 Hz), 111.8 (d, *J* = 25.4 Hz) ppm.

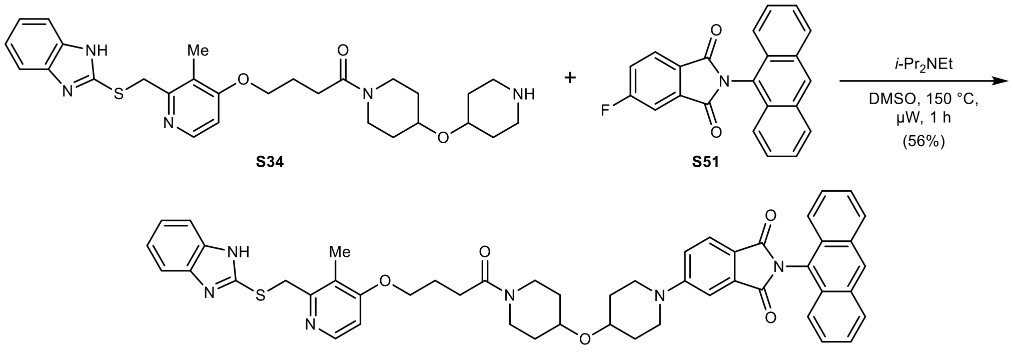

**25**

**5-(4-((1-(4-((2-(((1*H*-Benzo[*d*]imidazol-2-yl)thio)methyl)-3-methylpyridin-4-yl)oxy)butanoyl) piperidin-4-yl)oxy)piperidin-1-yl)-2-(anthracen-9-yl)isoindoline-1,3-dione** (**25**): To a solution of **S34** (45 mg, 0.09 mmol, 1.00 equiv.) in anhydrous DMSO (0.7 mL) was added **S51** (26 mg, 0.08 mmol, 1.05 equiv.) and *i*-$Pr_2NEt$ (25 µL, 0.15 mmol, 2.00 equiv.) at rt under argon. The reaction was stirred at 150 °C for 1 h under microwave irradiation. The mixture was diluted with water (150 mL) and extracted with EtOAc (3 × 50 mL). The combined organic layers were washed with water (50 mL) and brine (50 mL), dried with $MgSO_4$, filtered, and concentrated *in vacuo*. Purification by column chromatography (12 g silica, 0 to 4% MeOH with 2.5% $NH_4OH$ in $CH_2Cl_2$ in 11 min) yielded 34 mg (0.04 mmol, 56%) of **25** as a yellow powder after lyophilization from $MeCN/H_2O$.

**TLC** $R_f$ = 0.38 (5% MeOH in $CH_2Cl_2$).

**HRMS** (ESI) *m/z*: $(M+H)^+$ calc. for $C_{50}H_{49}N_6O_5S$: 845.3480; found: 845.3480.

**$^1$H NMR** (400 MHz, $CDCl_3$) δ 8.60 (s, 1H), 8.34 (d, *J* = 5.7 Hz, 1H), 8.14 – 8.04 (m, 2H), 7.86 (d, *J* = 8.5 Hz, 1H), 7.81 – 7.73 (m, 2H), 7.60 – 7.50 (m, 2H), 7.50 – 7.43 (m, 5H), 7.23 – 7.14 (m, 3H), 6.79 (d, *J* = 5.8 Hz, 1H), 4.38 (s, 2H), 4.13 (t, *J* = 6.0 Hz, 2H), 3.92 (ddd, *J* = 12.3, 7.2, 3.7 Hz, 1H), 3.83 – 3.64 (m, 5H), 3.45 – 3.23 (m, 4H), 2.54 (t, *J* = 7.0 Hz, 2H), 2.26 (s, 3H), 2.20 (p, *J* = 6.4 Hz, 2H), 2.01 – 1.90 (m, 2H), 1.88 – 1.77 (m, 2H), 1.77 – 1.66 (m, 2H), 1.66 – 1.52 (m, 2H) ppm. **Note:** The benzimidazole NH peak is in exchange with residual water in the spectrum.

**$^{13}$C NMR** (101 MHz, $CDCl_3$) δ 170.2, 168.9, 168.3, 164.3, 156.7, 155.4, 151.8, 147.5, 134.9, 131.9, 129.7, 129.1, 129.0, 127.3, 125.9, 125.6, 124.0, 122.7, 121.9, 121.1, 119.4, 118.2, 108.9, 106.4, 71.2, 71.2, 67.9, 45.5, 45.4, 42.7, 39.1, 35.0, 32.5, 31.5, 31.3, 31.1, 29.1, 24.6, 10.9 ppm. **Note:** C1, C2, C3, and C6 are not detected due to benzimidazole tautomerism.

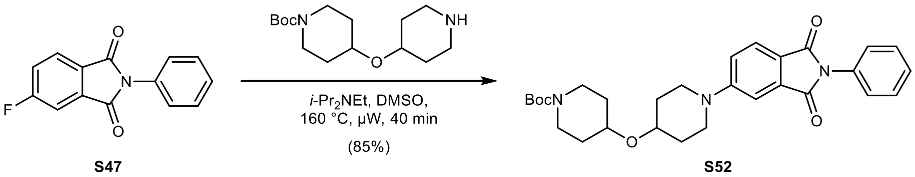

**S47** **S52**

***tert*-Butyl 4-((1-(1,3-dioxo-2-phenylisoindolin-5-yl)piperidin-4-yl)oxy)piperidine-1-carboxylate (S52):** To a solution of **S47** (80 mg, 0.33 mmol, 1.00 equiv.) in anhydrous DMSO (2.0 mL) was added *tert*-butyl 4-(piperidin-4-yloxy)piperidine-1-carboxylate (96 mg, 0.34 mmol, 1.02 equiv.) and *i*-$Pr_2NEt$ (144 µL, 0.83 mmol, 2.50 equiv.) at rt under argon. The reaction was stirred at 160 °C for 40 min under microwave irradiation. The mixture was diluted with water (100 mL) and extracted with EtOAc (3 × 25 mL). The combined organic layers were washed with water (50 mL) and brine (50 mL), dried with $MgSO_4$, filtered, and concentrated *in vacuo*. Purification by column chromatography (12 g silica, 35 to 50% EtOAc in *n*-heptane in 8 min) yielded 142 mg (0.28 mmol, 85%) of **S52** as a yellow solid.

**TLC** $R_f$ = 0.63 (75% EtOAc in *n*-heptane).

**LC/MS** (ESI) *m/z*: $(M+H)^+$ 506.2.

**$^1$H NMR** (400 MHz, $CDCl_3$) δ 7.75 (dd, *J* = 8.5, 0.4 Hz, 1H), 7.49 (m, 2H), 7.42 (m, 2H), 7.37 (m, 2H), 7.10 (dd, *J* = 8.5, 2.4 Hz, 1H), 3.74 (m, 5H), 3.62 (tt, *J* = 7.9, 3.7 Hz, 1H), 3.30 (ddd, *J* = 13.1, 8.3, 3.5 Hz, 2H), 3.12 (ddd, *J* = 13.3, 9.0, 3.5 Hz, 2H), 1.94 (m, 2H), 1.80 (m, 2H), 1.71 (m, 2H), 1.54 (m, 2H), 1.46 (s, 9H) ppm.

**$^{13}$C NMR** (101 MHz, $CDCl_3$) δ 168.1, 167.5, 155.3, 155.0, 134.6, 132.3, 129.1, 127.8, 126.7, 125.5, 119.2, 118.1, 108.7, 79.6, 72.0, 70.9, 45.5, 41.4, 31.9, 31.2, 28.6 ppm.

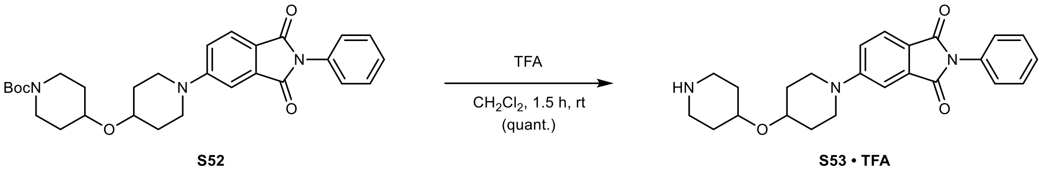

**2-Phenyl-5-(4-(piperidin-4-yloxy)piperidin-1-yl)isoindoline-1,3-dione (S53 · TFA):** To a solution of **S52** (132 mg, 0.26 mmol, 1.00 equiv.) in $CH_2Cl_2$ (2.0 mL) was added trifluoroacetic acid (1.0 mL, 13.1 mmol, 50.0 equiv.) slowly at 0 °C. After 5 min, the reaction was allowed to warm to rt and stirred for 1.5 h. The mixture was concentrated *in vacuo* – residual trifluoroacetic acid was removed by co-evaporation with MeOH (2 × 5 mL) and drying *in vacuo*, yielding 136 mg (0.26 mmol, quantitative) of **S53 · TFA** as a yellow solid.

**TLC** $R_f$ = 0.14, streaky (20% MeOH with 0.5% $NH_4OH$ in $CH_2Cl_2$).

**LC/MS** (ESI) *m/z*: $(M+H)^+$ 406.2.

**$^1$H NMR** (400 MHz, DMSO-$d_6$) δ 8.48 (m, 2H), 7.72 (d, *J* = 8.5 Hz, 1H), 7.51 (m, 2H), 7.41 (m, 4H), 7.29 (dd, *J* = 8.6, 2.4 Hz, 1H), 3.77 (m, 4H), 3.26 (ddd, *J* = 12.9, 9.3, 3.1 Hz, 2H), 3.18 (m, 2H), 2.99 (m, 2H), 1.91 (m, 4H), 1.64 (m, 2H), 1.50 (m, 2H) ppm.

**$^{13}$C NMR** (101 MHz, DMSO-$d_6$) δ 167.3, 166.7, 154.8, 134.2, 132.2, 128.8, 127.7, 127.2, 125.1, 117.9, 117.9, 107.8, 70.9, 67.6, 44.9, 40.8, 30.7, 28.3 ppm.

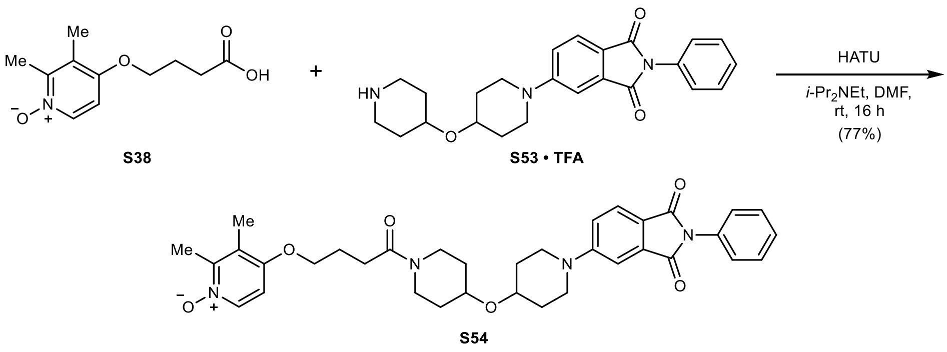

**4-(4-(4-((1-(1,3-Dioxo-2-phenylisoindolin-5-yl)piperidin-4-yl)oxy)piperidin-1-yl)-4-oxobutoxy)-2,3-dimethylpyridine 1-oxide (S54):** To a suspension of **S38** (48 mg, 0.21 mmol, 1.05 equiv.) in anhydrous DMF (1.0 mL) was added HATU (92 mg, 0.24 mmol, 1.20 equiv.), *i*-$Pr_2NEt$ (105 µL, 0.60 mmol, 3.00 equiv.), and **S53 · TFA** (105 mg, 0.20 mmol, 1.00 equiv.) at rt under argon. The reaction was stirred at rt for 16 h. The mixture was poured into water (150 mL) and extracted with EtOAc (25 mL), then with 10% MeOH in $CH_2Cl_2$ (3 × 25 mL). The combined organic layers were washed with 5% aqueous LiCl (2 × 25 mL), water (25 mL), and brine (25 mL), dried with $MgSO_4$, filtered, and concentrated *in vacuo*. Purification by column chromatography (12 g silica, 1 to 15% MeOH in $CH_2Cl_2$ in 12 min) yielded 95 mg (0.16 mmol, 77%) of **S54** as a yellow solid.

**TLC** $R_f$ = 0.41 (10% MeOH in $CH_2Cl_2$).

**LC/MS** (ESI) *m/z*: $(M+H)^+$ 613.3.

**$^1$H NMR** (400 MHz, $CDCl_3$) δ 8.13 (d, *J* = 7.2 Hz, 1H), 7.75 (d, *J* = 8.5 Hz, 1H), 7.49 (m, 2H), 7.39 (m, 4H), 7.10 (dd, *J* = 8.6, 2.4 Hz, 1H), 6.67 (d, *J* = 7.3 Hz, 1H), 4.08 (t, *J* = 6.2 Hz, 2H), 3.92 (m, 1H), 3.73 (m, 5H), 3.40 (ddd, *J* = 12.7, 8.1, 3.7 Hz, 1H), 3.30 (m, 3H), 2.53 (m, 5H), 2.18 (m, 5H), 1.95 (m, 2H), 1.83 (m, 2H), 1.72 (m, 2H), 1.61 (m, 2H) ppm.

**$^{13}$C NMR** (101 MHz, $CDCl_3$) δ 170.2, 168.1, 167.5, 155.3, 155.2, 149.1, 137.4, 134.6, 132.3, 129.2, 127.9, 126.7, 125.6, 123.2, 119.3, 118.2, 108.7, 105.8, 71.2, 71.2, 68.2, 45.5, 45.5, 42.8, 39.1, 32.5, 31.4, 31.3, 31.1, 29.2, 24.7, 14.5, 12.1 ppm.

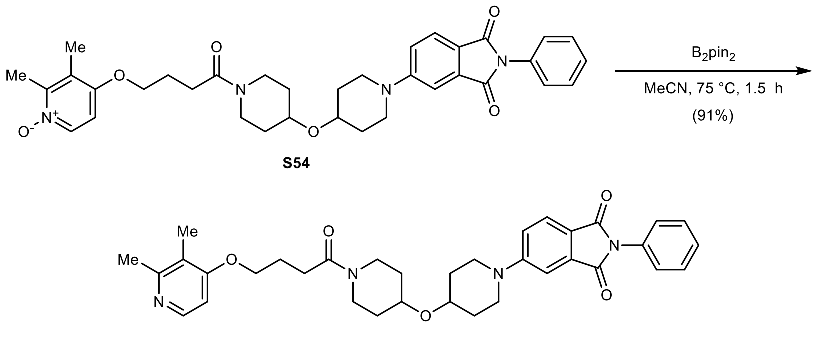

26 (DKFZ-1234)

**5-(4-((1-(4-((2,3-Dimethylpyridin-4-yl)oxy)butanoyl)piperidin-4-yl)oxy)piperidin-1-yl)-2-phenylisoindoline-1,3-dione (26; DKFZ-1234):** To a solution of **S54** (93 mg, 0.15 mmol, 1.00 equiv.) in anhydrous MeCN (1.5 mL) was added bis(pinacolato)diboron (41 mg, 0.16 mmol, 1.10 equiv.) at rt under argon. The reaction was heated to 75 °C and stirred for 1.5 h. The mixture was poured into water (100 mL) and extracted with 10% MeOH in $CH_2Cl_2$ (5 × 30 mL). The combined organic layers were washed with water (50 mL) and brine (50 mL), dried with $MgSO_4$, filtered, and concentrated *in vacuo*. Purification by column chromatography (12 g silica, 0 to 8% MeOH with 2.5% $NH_4OH$ in $CH_2Cl_2$ in 15 min) yielded 80 mg (0.13 mmol, 91%) of **26** (**DKFZ-1234**) as a yellow powder.

**TLC** $R_f$ = 0.58 (10% MeOH with 0.5% $NH_4OH$ in $CH_2Cl_2$).

**HRMS** (ESI) *m/z*: $(M+H)^+$ calc. for $C_{35}H_{41}N_4O_5$: 597.3071; found: 597.3070.

**$^1$H NMR** (400 MHz, $CDCl_3$) δ 8.20 (d, *J* = 5.7 Hz, 1H), 7.75 (d, *J* = 8.5 Hz, 1H), 7.49 (m, 2H), 7.42 (m, 2H), 7.37 (m, 2H), 7.09 (dd, *J* = 8.6, 2.4 Hz, 1H), 6.63 (d, *J* = 5.7 Hz, 1H), 4.07 (t, *J* = 6.0 Hz, 2H), 3.92 (ddd, *J* = 12.0, 7.2, 3.8 Hz, 1H), 3.72 (m, 5H), 3.39 (ddd, *J* = 12.7, 8.1, 3.6 Hz, 1H), 3.30 (t, *J* = 10.8 Hz, 3H), 2.54 (t, *J* = 7.1 Hz, 2H), 2.48 (s, 3H), 2.18 (p, *J* = 6.3 Hz, 2H), 2.13 (s, 3H), 1.94 (m, 2H), 1.81 (m, 2H), 1.71 (m, 2H), 1.57 (m, 2H) ppm.

**$^{13}$C NMR** (101 MHz, $CDCl_3$) δ 170.5, 168.1, 167.4, 162.9, 157.7, 155.3, 147.6, 134.6, 132.3, 129.1, 127.9, 126.7, 125.5, 119.8, 119.3, 118.2, 108.7, 104.7, 71.2, 71.1, 67.3, 45.5, 45.4, 42.8, 39.1, 32.5, 31.4, 31.3, 31.1, 29.4, 24.9, 22.8, 11.1 ppm.

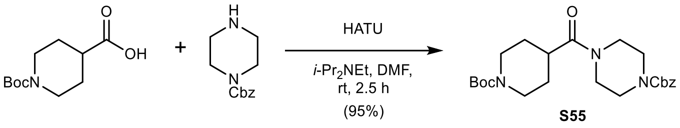

**Benzyl 4-(1-(*tert*-butoxycarbonyl)piperidine-4-carbonyl)piperazine-1-carboxylate (S55):** To a solution of 1-(*tert*-butoxycarbonyl)piperidine-4-carboxylic acid (1.00 g, 4.36 mmol, 1.00 equiv.) in anhydrous DMF (10 mL) was added HATU (1.82 g, 4.80 mmol, 1.20 equiv.), benzyl piperazine-1-carboxylate (0.84 mL, 4.36 mmol, 1.00 equiv.), and *i*-$Pr_2NEt$ (1.52 mL, 8.72 mmol, 2.00 equiv.) at rt under argon. The reaction was stirred at rt for 2.5 h. The mixture was concentrated *in vacuo*, then poured into water (300 mL) and extracted with EtOAc (3 × 100 mL). The combined organic layers were washed with 5% aqueous LiCl (3 × 100 mL), water (3 × 100 mL), and brine (2 × 100 mL), dried with $MgSO_4$, filtered, and concentrated *in vacuo*. Purification by column chromatography (120 g silica, 30 to 100% EtOAc in *n*-heptane in 25 min) yielded 1.80 g (4.17 mmol, 95%) of **S55** as a colorless solid.

**TLC** $R_f$ = 0.39 (75% EtOAc in *n*-heptane).

**LC/MS** (ESI) *m/z*: $(M+Na)^+$ 452.2.

**$^1$H NMR** (400 MHz, $CDCl_3$) δ 7.35 (m, 5H), 5.15 (s, 2H), 4.15 (m, 2H), 3.54 (m, 8H), 2.78 (m, 2H), 2.59 (m, 1H), 1.69 (m, 4H), 1.45 (s, 9H) ppm.

**$^{13}$C NMR** (101 MHz, $CDCl_3$) δ 173.2, 155.1, 154.7, 136.3, 128.6, 128.3, 128.1, 79.7, 67.5, 45.2, 43.9, 43.9, 41.5, 38.5, 28.5, 28.4 ppm.

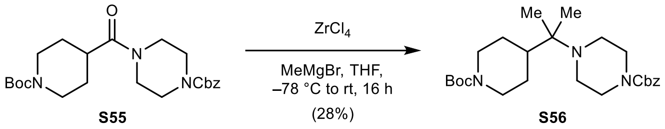

**Benzyl 4-(2-(1-(*tert*-butoxycarbonyl)piperidin-4-yl)propan-2-yl)piperazine-1-carboxylate (S56):** To a solution of $ZrCl_4$ (549 mg, 2.35 mmol, 1.60 equiv.) in anhydrous THF (20 mL) was added a solution of **S55** (635 mg, 1.47 mmol, 1.00 equiv.) in anhydrous THF (29 mL) dropwise over 15 min at -78 °C under argon. MeMgBr (2.00 mL, 3.74 M in $Et_2O$ determined by NMR, 7.36 mmol, 5.00 equiv.) was added dropwise over 5 min at -78 °C. After 30 min, the reaction was allowed to warm to rt and stirred for 16 h. The mixture was quenched with saturated aqueous $NH_4Cl$ (200 mL) and extracted with EtOAc (4 × 75 mL). The combined organic layers were stirred with Celite for 10 min, then filtered through Celite on a sintered glass funnel, and rinsed with EtOAc (50 mL). The combined organic layers were washed with brine (2 × 50 mL), dried with $MgSO_4$, filtered, and concentrated *in vacuo*. Purification by column chromatography (40 g silica, 5 to 40% EtOAc in *n*-heptane in 17 min) yielded 182 mg (0.41 mmol, 28%) of **S56** as a colorless oil.

**TLC** $R_f$ = 0.60 (50% EtOAc in *n*-heptane).

**LC/MS** (ESI) *m/z*: $(M+H)^+$ 446.3.

**$^1$H NMR** (400 MHz, $CDCl_3$) δ 7.39 – 7.28 (m, 5H), 5.13 (s, 2H), 4.26 – 4.06 (m, 2H), 3.53 – 3.37 (m, 4H), 2.59 (t, *J* = 12.8 Hz, 2H), 2.54 – 2.41 (m, 4H), 1.76 – 1.66 (m, 2H), 1.64 – 1.50 (m, 1H), 1.45 (s, 9H), 1.16 (tt, *J* = 12.8, 6.4 Hz, 2H), 0.88 (s, 6H) ppm.

**$^{13}$C NMR** (101 MHz, $CDCl_3$) δ 155.2, 154.9, 136.9, 128.5, 128.0, 127.9, 79.3, 67.0, 57.8, 45.1, 44.8, 44.8, 43.2, 28.5, 26.9, 19.3 ppm.

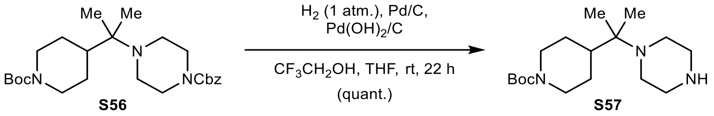

***tert*-Butyl 4-(2-(piperazin-1-yl)propan-2-yl)piperidine-1-carboxylate (S57):** To a solution of **S56** (121 mg, 0.27 mmol, 1.00 equiv.) in anhydrous THF (2.0 mL) and 2,2,2-trifluoroethanol (0.7 mL, dried over $MgSO_4$) was added Pd/C (14.5 mg, 10w%, 0.01 mmol, 0.05 equiv.) and $Pd(OH)_2/C$ (19 mg, 20w%, 0.03 mmol, 0.10 equiv.) at rt under argon. The mixture was evacuated and refilled with hydrogen three times. The reaction was stirred at rt for 22 h. The atmosphere was exchanged with argon and the mixture was filtered through Celite on a sintered glass funnel, rinsed with EtOH (15 mL), and concentrated *in vacuo*, yielding 85 mg (0.27 mmol, quantitative) of **S57** as a light-brown, waxy solid.

**TLC** $R_f$ = 0.13, streaky (20% MeOH with 0.5% NH4OH in $CH_2Cl_2$).

**LC/MS** (ESI) *m/z*: $(M+H)^+$ 312.2.

**$^1$H NMR** (400 MHz, $CD_3OD$) δ 4.19 – 4.09 (m, 2H), 2.89 – 2.75 (m, 4H), 2.75 – 2.61 (m, 2H), 2.61 – 2.49 (m, 4H), 1.80 – 1.65 (m, 3H), 1.45 (s, 9H), 1.13 (qd, *J* = 12.6, 4.3 Hz, 2H), 0.91 (s, 6H) ppm. **Note:** The NH peaks are in exchange with $CD_3OD$ in the spectrum.

**$^{13}$C NMR** (101 MHz, $CD_3OD$) δ 156.5, 80.8, 58.8, 47.4, 46.7, 45.9, 44.1, 28.7, 28.1, 19.4 ppm.

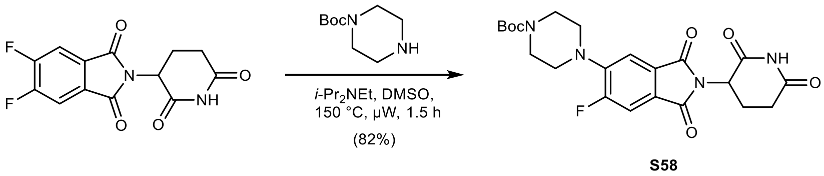

***tert*-Butyl 4-(2-(2,6-dioxopiperidin-3-yl)-6-fluoro-1,3-dioxoisoindolin-5-yl)piperazine-1-carboxylate (S58):** To a solution of 2-(2,6-dioxopiperidin-3-yl)-5,6-difluoroisoindoline-1,3-dione (500 mg, 1.70 mmol, 1.00 equiv.) in anhydrous DMSO (8.0 mL) was added *tert*-butyl piperazine-1-carboxylate (323 mg, 1.73 mmol, 1.02 equiv.) and *i*-$Pr_2NEt$ (0.74 mL, 4.25 mmol, 2.50 equiv.) at rt under argon. The reaction was stirred at 150 °C for 1.5 h under microwave irradiation. The mixture was diluted with water (200 mL) and extracted with EtOAc (3 × 50 mL). The combined organic layers were washed with water (100 mL) and brine (100 mL), dried with $MgSO_4$, filtered, and concentrated *in vacuo*. Purification by column chromatography (40 g silica, 40 to 65% EtOAc in *n*-heptane in 12 min) yielded 645 mg (1.40 mmol, 82%) of **S58** as a yellow powder.

**TLC** $R_f$ = 0.53 (75% EtOAc in *n*-heptane).

**LC/MS** (ESI) *m/z*: $(M+Na)^+$ 483.1.

**$^1$H NMR** (400 MHz, $CDCl_3$) δ 8.35 (s, 1H), 7.48 (d, *J* = 10.9 Hz, 1H), 7.38 (d, *J* = 7.2 Hz, 1H), 4.93 (dd, *J* = 12.2, 5.3 Hz, 1H), 3.61 (m, 4H), 3.20 (m, 4H), 2.81 (m, 3H), 2.13 (m, 1H), 1.48 (s, 9H) ppm.

**$^{19}$F{$^1$H} NMR** (377 MHz, $CDCl_3$) δ –110.9 ppm.

**$^{13}$C NMR** (101 MHz, $CDCl_3$) δ 171.1, 168.2, 167.0, 166.4 (d, *J* = 2.4 Hz), 158.4 (d, *J* = 256.0 Hz), 154.7, 145.8 (d, *J* = 9.0 Hz), 129.1 (d, *J* = 2.8 Hz), 124.7 (d, *J* = 9.6 Hz), 114.0 (d, *J* = 4.6 Hz), 112.2, 80.4, 50.1, 50.0, 49.6, 43.5, 31.5, 28.5, 22.8 ppm.

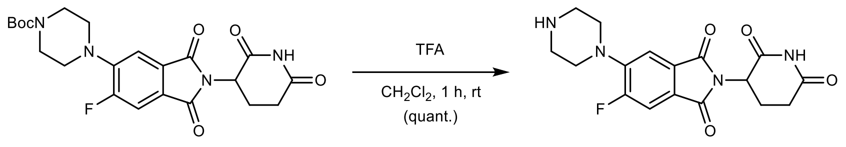

**2-(2,6-Dioxopiperidin-3-yl)-5-fluoro-6-(piperazin-1-yl)isoindoline-1,3-dione (S59 · TFA):** To a solution of **S58** (622 mg, 1.35 mmol, 1.00 equiv.) in $CH_2Cl_2$ (9.0 mL) was added trifluoroacetic acid (3.0 mL, 39.1 mmol, 29.0 equiv.) slowly at 0 °C. After 5 min, the reaction was allowed to warm to rt and stirred for 1 h. The mixture was concentrated *in vacuo* – residual trifluoroacetic acid was removed by co-evaporation with MeOH (2 × 5 mL), toluene (2 × 5 mL), and drying *in vacuo*, yielding 645 mg (1.36 mmol, quantitative) of **S59 · TFA** as a pale-yellow powder.

**TLC** $R_f$ = 0.23, streaky (10% MeOH with 0.5% $NH_4OH$ in $CH_2Cl_2$).

**LC/MS** (ESI) *m/z*: $(M+H)^+$ 361.1.

**$^1$H NMR** (400 MHz, DMSO-$d_6$) δ 11.13 (s, 1H), 9.03 (bs, 2H), 7.82 (d, *J* = 11.2 Hz, 1H), 7.60 (d, *J* = 7.4 Hz, 1H), 5.14 (dd, *J* = 12.8, 5.3 Hz, 1H), 3.48 (m, 4H), 3.30 (m, 4H), 2.91 (ddd, *J* = 16.6, 13.7, 5.3 Hz, 1H), 2.63 (m, 2H), 2.06 (m, 1H) ppm.

**$^{19}$F{$^1$H} NMR** (377 MHz, DMSO-$d_6$) δ –74.3 (TFA), –112.0 ppm.

**$^{13}$C NMR** (101 MHz, DMSO-$d_6$) δ 172.8, 169.9, 166.5, 166.1 (d, *J* = 1.7 Hz), 158.3 (q, *J* = 33.8 Hz, TFA), 157.5 (d, *J* = 253.7 Hz), 144.2 (d, *J* = 9.0 Hz), 128.7 (d, *J* = 2.5 Hz), 124.6 (d, *J* = 9.8 Hz), 116.4 (q, *J* = 295.2 Hz, TFA), 114.5 (d, *J* = 3.7 Hz), 112.2 (d, *J* = 25.1 Hz), 49.1, 46.6, 46.6, 42.6, 42.6, 31.0, 22.0 ppm.

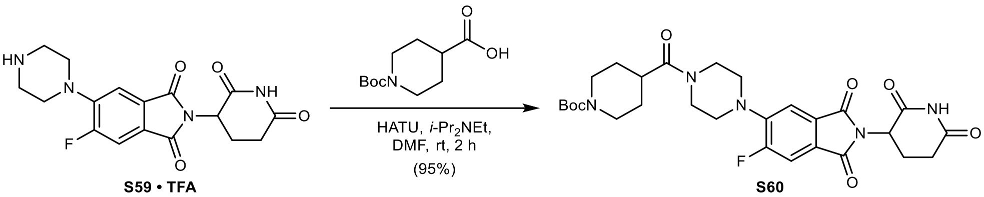

***tert*-Butyl 4-(4-(2-(2,6-dioxopiperidin-3-yl)-6-fluoro-1,3-dioxoisoindolin-5-yl)piperazine-1-carbonyl)piperidine-1-carboxylate (S60):** To a solution of **S59 · TFA** (387 mg, 0.82 mmol, 1.00 equiv.) in anhydrous DMF (4.0 mL) was added 1-(*tert*-butoxycarbonyl)piperidine-4-carboxylic acid (196 mg, 0.86 mmol, 1.05 equiv.), HATU (372 mg, 0.98 mmol, 1.20 equiv.), and *i*-$Pr_2NEt$ (426 µL, 2.45 mmol, 3.00 equiv.) at rt under argon. The reaction was stirred at rt for 2 h. The mixture was poured into water (200 mL), extracted with EtOAc (2 × 30 mL) and $CH_2Cl_2$ (3 × 50 mL). The combined organic layers were washed with 5% aqueous LiCl (2 × 50 mL), water (50 mL), and brine (50 mL), dried with $MgSO_4$, filtered, and concentrated *in vacuo*. Purification by column chromatography (40 g silica, 10 to 75% EtOAc in *n*-heptane, then 0.5 to 7% MeOH in $CH_2Cl_2$ in 18 min) yielded 443 mg (0.77 mmol, 95%) of **S60** as a yellow powder.

**TLC** $R_f$ = 0.65 (10% MeOH in $CH_2Cl_2$).

**LC/MS** (ESI) *m/z*: $(M+Na)^+$ 594.2.

**$^1$H NMR** (400 MHz, DMSO-$d_6$) δ 11.11 (s, 1H), 7.77 (d, *J* = 11.3 Hz, 1H), 7.49 (d, *J* = 7.3 Hz, 1H), 5.11 (dd, *J* = 12.8, 5.4 Hz, 1H), 3.94 (m, 2H), 3.67 (m, 4H), 3.24 (m, 4H), 2.88 (m, 4H), 2.58 (m, 2H), 2.04 (m, 1H), 1.64 (m, 2H), 1.40 (m, 11H) ppm.

**$^{19}$F{$^1$H} NMR** (377 MHz, DMSO-$d_6$) δ -112.1 ppm.

**$^{13}$C NMR** (101 MHz, DMSO-$d_6$) δ 172.7, 172.4, 169.9, 166.6, 166.1, 157.4 (d, *J* = 253.5 Hz), 153.9, 145.1 (d, *J* = 8.8 Hz), 128.7, 123.9 (d, *J* = 10.1 Hz), 114.1 (d, *J* = 3.5 Hz), 112.0 (d, *J* = 25.5 Hz), 78.6, 54.9, 50.0, 49.4, 49.1, 44.5, 42.8, 42.8, 40.9, 36.9, 30.9, 28.2, 28.1, 22.0 ppm.

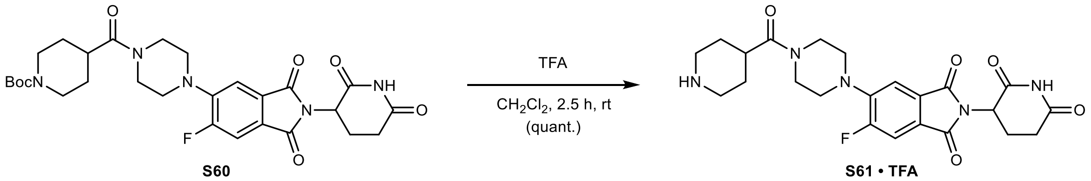

S60 S61 • TFA

**2-(2,6-Dioxopiperidin-3-yl)-5-fluoro-6-(4-(piperidine-4-carbonyl)piperazin-1-yl)isoindoline-1,3-dione (S61 · TFA):** To a solution of **S60** (411 mg, 0.72 mmol, 1.00 equiv.) in $CH_2Cl_2$ (4.5 mL) was added trifluoroacetic acid (1.5 mL, 19.4 mmol, 27.0 equiv.) slowly at 0 °C. After 5 min, the reaction was allowed to warm to rt and stirred for 2.5 h. The mixture was concentrated *in vacuo* – residual trifluoroacetic acid was removed by co-evaporation with MeOH (3 × 5 mL) and drying *in vacuo*, yielding 414 mg (0.72 mmol, quantitative) of **S61 · TFA** as a yellow powder.

**TLC** $R_f$ = 0.07, streaky (20% MeOH with 0.5% $NH_4OH$ in $CH_2Cl_2$).

**LC/MS** (ESI) *m/z*: $(M+H)^+$ 472.2.

**$^1$H NMR** (400 MHz, DMSO-$d_6$) δ 11.11 (s, 1H), 8.62 (m, 1H), 8.36 (m, 1H), 7.78 (d, *J* = 11.2 Hz, 1H), 7.49 (d, *J* = 7.3 Hz, 1H), 5.11 (dd, *J* = 12.8, 5.4 Hz, 1H), 3.67 (m, 4H), 3.25 (m, 5H), 2.93 (m, 4H), 2.56 (m, 2H), 2.04 (m, 1H), 1.77 (m, 4H) ppm.

**$^{19}$F{$^1$H} NMR** (377 MHz, DMSO-$d_6$) δ –73.7 (TFA), –112.1 ppm.

**$^{13}$C NMR** (101 MHz, DMSO-$d_6$) δ 172.8, 171.6, 169.9, 166.6, 166.1, 157.4 (d, *J* = 253.3 Hz), 145.0 (d, *J* = 8.8 Hz), 128.7, 124.0 (d, *J* = 9.5 Hz), 114.1 (d, *J* = 4.6 Hz), 112.1 (d, *J* = 25.1 Hz), 49.9, 49.4, 49.1, 44.5, 42.6, 42.6, 40.9, 34.6, 30.9, 25.1, 25.1, 22.1 ppm.

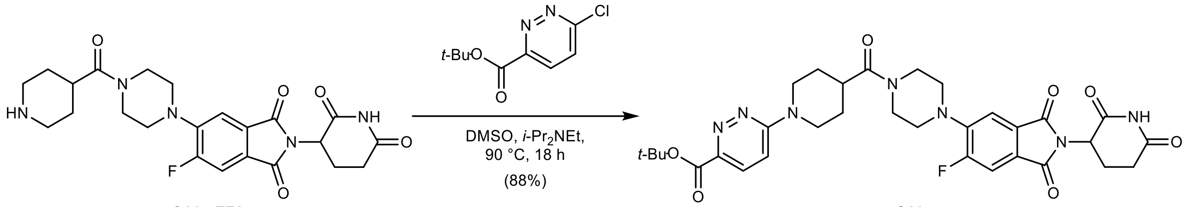

S61 • TFA S62

***tert*-Butyl 6-(4-(4-(2-(2,6-dioxopiperidin-3-yl)-6-fluoro-1,3-dioxoisoindolin-5-yl)piperazine-1-carbonyl)piperidin-1-yl)pyridazine-3-carboxylate (S62):** To a solution of **S61 · TFA** (100 mg, 0.17 mmol, 1.00 equiv.) in anhydrous DMSO (1.2 mL) was added *tert*-butyl 6-chloropyridazine-3-carboxylate (110 mg, 0.51 mmol, 3.00 equiv.) and *i*-$Pr_2NEt$ (89 µL, 0.51 mmol, 3.00 equiv.) at rt under argon. The reaction was stirred at 90 °C for 18 h. The mixture was diluted with water (150 mL) and extracted with EtOAc (3 × 50 mL). The combined organic layers were washed with water (50 mL) and brine (50 mL), dried with $MgSO_4$, filtered, and concentrated *in vacuo*. Purification by column chromatography (12 g silica, 0 to 6.5% MeOH with 2.5% $NH_4OH$ in $CH_2Cl_2$ in 10 min) yielded 98 mg (0.15 mmol, 88%) of **S62** as a yellow powder.

**TLC** $R_f$ = 0.57 (10% MeOH with 2.5% $NH_4OH$ in $CH_2Cl_2$).

**LC/MS** (ESI) *m/z*: $(M+H)^+$ 650.3.

**$^1$H NMR** (400 MHz, $CDCl_3$) δ 8.18 (s, 1H), 7.83 (d, *J* = 9.6 Hz, 1H), 7.52 (d, *J* = 10.8 Hz, 1H), 7.39 (d, *J* = 7.2 Hz, 1H), 6.88 (d, *J* = 9.7 Hz, 1H), 4.94 (dd, *J* = 12.3, 5.3 Hz, 1H), 4.57 (m, 2H), 3.79 (m, 4H),

3.27 (d, $J$ = 21.7 Hz, 4H), 3.17 (ddd, $J$ = 13.4, 11.0, 3.8 Hz, 2H), 2.82 (m, 4H), 2.14 (m, 1H), 1.91 (m, 4H), 1.63 (s, 9H) ppm.

**$^{19}$F{$^1$H} NMR** (377 MHz, $CDCl_3$) δ –110.8 ppm.

**$^{13}$C NMR** (101 MHz, $CDCl_3$) δ 172.9, 170.9, 168.1, 166.8, 166.3, 163.9, 159.6, 158.5 (d, $J$ = 255.9 Hz), 145.4 (d, $J$ = 9.2 Hz), 144.5, 129.1, 128.7, 125.3 (d, $J$ = 9.8 Hz), 114.1 (d, $J$ = 4.4 Hz), 112.5 (d, $J$ = 25.0 Hz), 110.8, 82.2, 50.5, 50.0, 49.6, 45.4, 44.4, 44.4, 41.6, 38.3, 31.5, 28.3, 28.1, 28.1, 22.8 ppm.

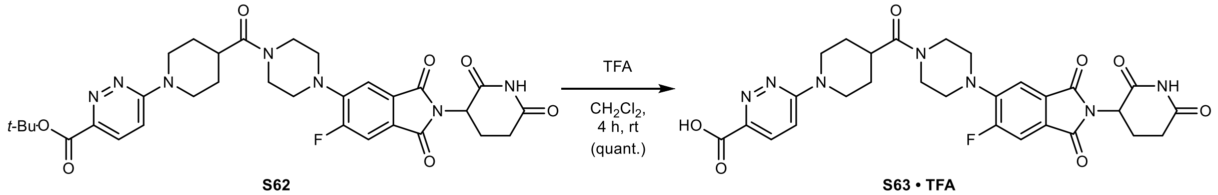

**6-(4-(4-(2-(2,6-Dioxopiperidin-3-yl)-6-fluoro-1,3-dioxoisoindolin-5-yl)piperazine-1-carbonyl) piperidin-1-yl)pyridazine-3-carboxylic acid (S63 · TFA):** To a solution of **S62** (94 mg, 0.14 mmol, 1.00 equiv.) in $CH_2Cl_2$ (1.5 mL) was added trifluoroacetic acid (0.5 mL, 6.51 mmol, 45.0 equiv.) slowly at 0 °C. After 5 min, the reaction was allowed to warm to rt and stirred for 4 h. The mixture was concentrated *in vacuo* – residual trifluoroacetic acid was removed by co-evaporation with MeOH (2 × 5 mL), toluene (2 × 5 mL), and drying *in vacuo*, yielding 87 mg (0.14 mmol, quantitative) of **S63 · TFA** as a yellow solid.

**TLC** $R_f$ = 0.15, streaky (20% MeOH in $CH_2Cl_2$).

**LC/MS** (ESI) *m/z*: $(M+H)^+$ 594.2.

**$^1$H NMR** (400 MHz, DMSO-$d_6$) δ 11.11 (s, 1H), 7.85 (d, $J$ = 9.6 Hz, 1H), 7.77 (d, $J$ = 11.3 Hz, 1H), 7.50 (d, $J$ = 7.3 Hz, 1H), 7.40 (d, $J$ = 9.7 Hz, 1H), 5.11 (dd, $J$ = 12.8, 5.4 Hz, 1H), 4.51 (m, 2H), 3.77 (m, 2H), 3.65 (m, 2H), 3.30 (m, 2H), 3.21 (m, 5H), 2.89 (ddd, $J$ = 16.3, 13.5, 5.2 Hz, 1H), 2.58 (m, 2H), 2.05 (m, 1H), 1.80 (m, 2H), 1.61 (m, 2H) ppm. **Note:** The proton corresponding to the carboxylic acid is in exchange with residual water in the spectrum.

**$^{19}$F{$^1$H} NMR** (377 MHz, DMSO-$d_6$) δ –74.8 (TFA), –112.1 ppm.

**$^{13}$C NMR** (101 MHz, DMSO-$d_6$) δ 172.8, 172.3, 169.9, 166.6, 166.1, 165.1, 158.9, 157.4 (d, $J$ = 253.7 Hz), 145.0, 142.5, 128.8, 128.7 (d, $J$ = 2.1 Hz), 123.9 (d, $J$ = 10.1 Hz), 114.1, 112.7, 112.0 (d, $J$ = 24.7 Hz), 50.0, 49.4, 49.1, 44.6, 44.0, 44.0, 40.9, 36.8, 31.0, 27.7, 27.7, 22.1 ppm.

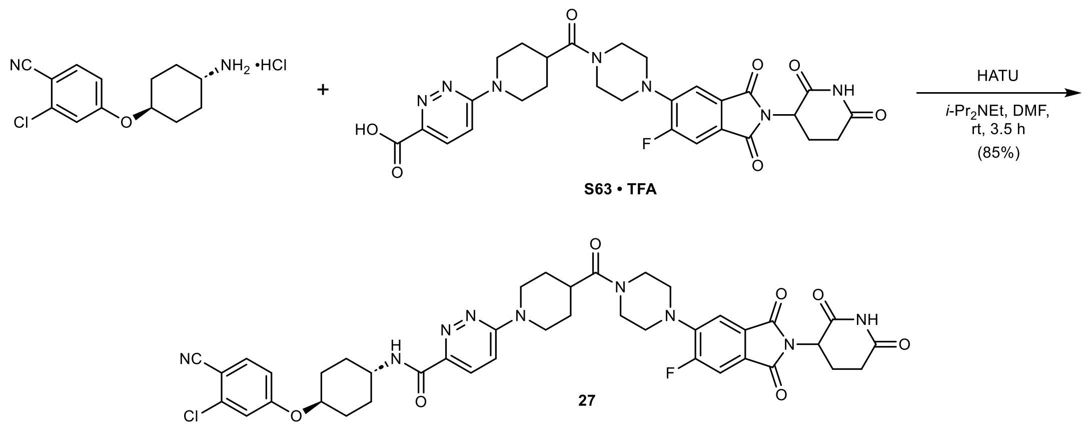

***N*-((1*r*,4*r*)-4-(3-Chloro-4-cyanophenoxy)cyclohexyl)-6-(4-(4-(2-(2,6-dioxopiperidin-3-yl)-6-fluoro-1,3-dioxoisoindolin-5-yl)piperazine-1-carbonyl)piperidin-1-yl)pyridazine-3-carboxamide (27):** To a solution of **S63 · TFA** (39 mg, 0.07 mmol, 1.00 equiv.) in anhydrous DMF (0.7 mL) was added HATU (30 mg, 0.08 mmol, 1.20 equiv.), 4-([trans-4-aminocyclohexyl]oxy)-2-chlorobenzonitrile hydrochloride (20 mg, 0.07 mmol, 1.05 equiv.), and *i*-$Pr_2NEt$ (57 µL, 0.33 mmol, 5.00 equiv.) at rt under argon. The reaction was stirred at rt for 3.5 h. The mixture was poured into water (200 mL) and extracted with EtOAc (3 × 50 mL). The combined organic layers were washed with 5% aqueous LiCl (2 × 50 mL), water (50 mL), and brine (50 mL), dried with $MgSO_4$, filtered, and concentrated *in vacuo*. Purification by column chromatography (12 g silica, 0 to 7.5% MeOH in $CH_2Cl_2$ in 11 min) yielded 46 mg (0.06 mmol, 85%) of **27** as a yellow powder after lyophilization from MeCN/$H_2O$.

**TLC** $R_f$ = 0.72 (10% MeOH in $CH_2Cl_2$).

**HRMS** (ESI) *m/z*: $(M+Na)^+$ calc. for $C_{41}H_{41}FN_9NaO_7Cl$: 848.2694; found: 848.2693.

**$^1$H NMR** (400 MHz, $CDCl_3$) δ 8.26 (s, 1H), 7.99 (d, *J* = 9.6 Hz, 1H), 7.87 (d, *J* = 8.2 Hz, 1H), 7.55 (d, *J* = 8.7 Hz, 1H), 7.52 (d, *J* = 10.7 Hz, 1H), 7.39 (d, *J* = 7.2 Hz, 1H), 7.00 (m, 2H), 6.85 (dd, *J* = 8.8, 2.4 Hz, 1H), 4.94 (dd, *J* = 12.2, 5.3 Hz, 1H), 4.53 (m, 2H), 4.32 (td, *J* = 10.1, 5.1 Hz, 1H), 4.05 (m, 1H), 3.80 (m, 4H), 3.20 (m, 6H), 2.82 (m, 4H), 2.16 (m, 5H), 1.94 (m, 4H), 1.67 (m, 2H), 1.46 (m, 2H) ppm.

**$^{19}$F{$^1$H} NMR** (377 MHz, $CDCl_3$) δ –110.8 ppm.

**$^{13}$C NMR** (101 MHz, $CDCl_3$) δ 172.8, 170.9, 168.1, 166.8, 166.3 (d, *J* = 2.5 Hz), 162.8, 161.8, 160.1, 158.5 (d, *J* = 256.2 Hz), 145.3 (d, *J* = 9.2 Hz), 144.5, 138.4, 135.2, 129.1 (d, *J* = 2.8 Hz), 127.0, 125.3 (d, *J* = 9.8 Hz), 117.0, 116.6, 114.8, 114.0 (d, *J* = 4.3 Hz), 112.6, 112.4, 104.9, 75.8, 50.5, 50.0, 49.7, 47.2, 45.4, 44.5, 44.5, 41.6, 38.2, 31.6, 30.2, 29.8, 29.8, 28.1, 28.1, 22.8 ppm.

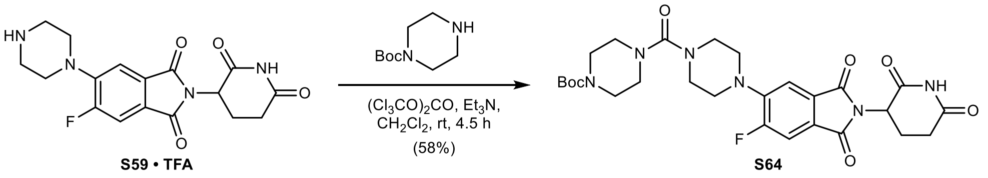

***tert*-Butyl 4-(4-(2-(2,6-dioxopiperidin-3-yl)-6-fluoro-1,3-dioxoisoindolin-5-yl)piperazine-1-carbonyl)piperazine-1-carboxylate (S64):** To a solution of triphosgene (19 mg, 0.06 mmol, 0.50 equiv.) in anhydrous $CH_2Cl_2$ (0.8 mL) was added a solution of **S59 · TFA** (60 mg, 0.13 mmol, 1.00 equiv.) and $Et_3N$ (53 µL, 0.38 mmol, 3.00 equiv.) in anhydrous $CH_2Cl_2$ (0.8 mL) dropwise at 0 °C under argon. The reaction was allowed to warm to rt and stirred for 3 h, then *tert*-butyl piperazine-1-carboxylate (236 mg, 1.27 mmol, 10.0 equiv.) was added. The reaction was stirred at rt for 1.5 h. The mixture was quenched with saturated, aqueous $NaHCO_3$ (75 mL) and extracted with EtOAc (3 × 20 mL). The combined organic layers were washed with saturated, aqueous $NH_4Cl$ (2 × 20 mL), water (20 mL), and brine (20 mL), dried with $MgSO_4$, filtered, and concentrated *in vacuo*. Purification by column chromatography (12 g silica, 0 to 5% MeOH in $CH_2Cl_2$ in 11 min) yielded 42 mg (0.07 mmol, 58%) of **S64** as a yellow powder.

**TLC** $R_f$ = 0.44 (5% MeOH in $CH_2Cl_2$).

**LC/MS** (ESI) *m/z*: $(M+Na)^+$ 595.2.

**$^1$H NMR** (400 MHz, DMSO-$d_6$) δ 11.11 (s, 1H), 7.74 (d, *J* = 11.3 Hz, 1H), 7.47 (d, *J* = 7.4 Hz, 1H), 5.11 (dd, *J* = 12.8, 5.4 Hz, 1H), 3.46 – 3.29 (m, 8H), 3.29 – 3.21 (m, 4H), 3.21 – 3.10 (m, 4H), 2.89 (ddd, *J* = 16.9, 13.9, 5.4 Hz, 1H), 2.67 – 2.53 (m, 2H), 2.10 – 1.98 (m, 1H), 1.41 (s, 9H) ppm.

**$^{19}$F{$^1$H} NMR** (377 MHz, DMSO-$d_6$) δ –112.0 ppm.

**$^{13}$C NMR** (101 MHz, DMSO-$d_6$) δ 172.7, 169.9, 166.6, 166.1, 162.7, 157.4 (d, *J* = 253.6 Hz), 153.9, 145.2 (d, *J* = 8.9 Hz), 128.7, 123.8 (d, *J* = 9.7 Hz), 113.9 (d, *J* = 4.6 Hz), 112.0 (d, *J* = 25.0 Hz), 79.0, 49.3, 49.1, 46.2, 46.0, 43.0, 30.9, 28.0, 22.1 ppm.

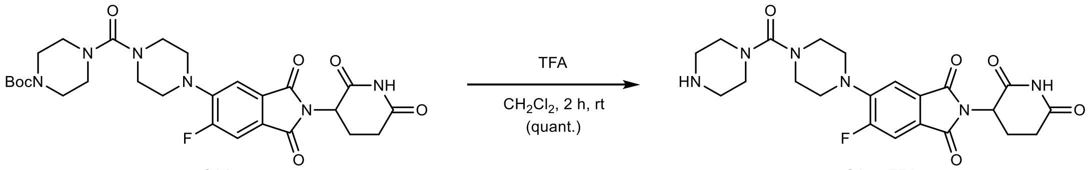

S64

S65 • TFA

**2-(2,6-Dioxopiperidin-3-yl)-5-fluoro-6-(4-(piperazine-1-carbonyl)piperazin-1-yl)isoindoline-1,3-dione (S65 · TFA):** To a solution of **S64** (42 mg, 0.07 mmol, 1.00 equiv.) in $CH_2Cl_2$ (1.0 mL) was added trifluoroacetic acid (0.3 mL, 3.96 mmol, 54.0 equiv.) slowly at 0 °C. After 5 min, the reaction was allowed to warm to rt and stirred for 2 h. The mixture was concentrated *in vacuo* – residual trifluoroacetic acid was removed by co-evaporation with MeOH (3 × 5 mL) and drying *in vacuo*, yielding 43 mg (0.07 mmol, quantitative) of **S65 · TFA** as a yellow solid.

**TLC** $R_f$ = 0.15, streaky (50% MeOH with 0.5% $NH_4OH$ in $CH_2Cl_2$).

**LC/MS** (ESI) *m/z*: $(M+H)^+$ 473.2.

**$^1$H NMR** (400 MHz, $CD_3OD$) δ 7.55 (d, *J* = 11.1 Hz, 1H), 7.47 (d, *J* = 7.3 Hz, 1H), 5.09 (dd, *J* = 12.7, 5.4 Hz, 1H), 3.64 – 3.43 (m, 8H), 3.30 – 3.20 (m, 8H), 2.87 (ddd, *J* = 17.3, 13.8, 5.2 Hz, 1H), 2.80 – 2.63 (m, 2H), 2.13 (dtd, *J* = 13.0, 5.8, 2.7 Hz, 1H) ppm. **Note:** The NH peaks are in exchange with $CD_3OD$ in the spectrum.

**$^{19}$F{$^1$H} NMR** (377 MHz, $CD_3OD$) δ –77.1 (TFA), –113.2 ppm.

**$^{13}$C NMR** (101 MHz, $CD_3OD$) δ 173.9, 170.8, 167.6, 167.2, 163.9, 158.9 (d, *J* = 254.6 Hz), 146.2 (d, *J* = 8.9 Hz), 129.8, 125.4 (d, *J* = 9.5 Hz), 114.3 (d, *J* = 4.4 Hz), 112.1 (d, *J* = 25.3 Hz), 50.2, 50.2, 50.1, 46.9, 44.4, 43.6, 31.5, 23.0 ppm.

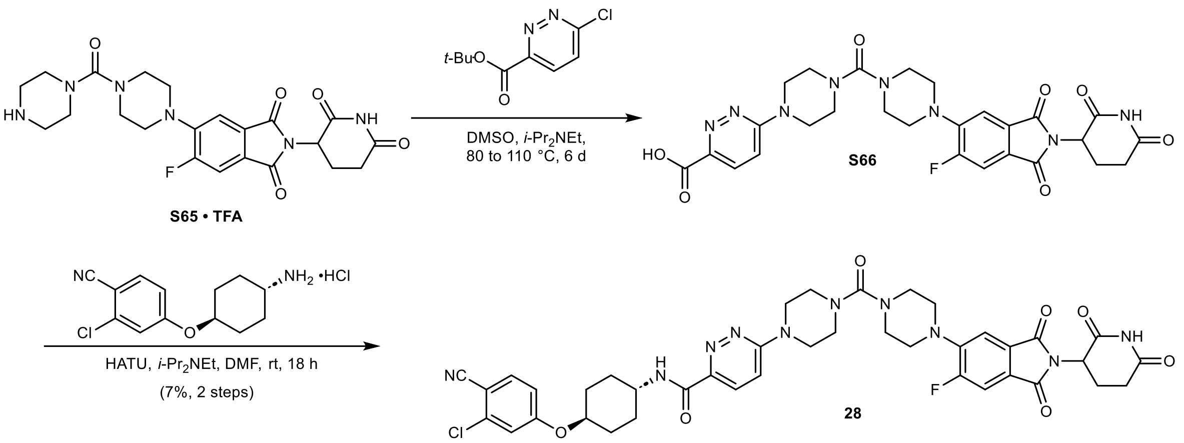

***N*-((1*r*,4*r*)-4-(3-Chloro-4-cyanophenoxy)cyclohexyl)-6-(4-(4-(2-(2,6-dioxopiperidin-3-yl)-6-fluoro-1,3-dioxoisoindolin-5-yl)piperazine-1-carbonyl)piperazin-1-yl)pyridazine-3-carboxamide (28):** To a solution of **S65 · TFA** (44 mg, 0.07 mmol, 1.00 equiv.) in anhydrous DMSO (0.8 mL) was added *tert*-butyl 6-chloropyridazine-3-carboxylate (48 mg, 0.22 mmol, 3.00 equiv.) and *i*-$Pr_2NEt$ (39 µL, 0.22 mmol, 3.00 equiv.) at rt under argon. The reaction was stirred at 90 °C for 16 h. To drive the reaction to completion, the reaction was heated to 110 °C and stirred for 3 h. **Note:** Elevated temperatures resulted in the cleavage of the *tert*-butyl ester of *tert*-butyl 6-chloropyridazine-3-carboxylate and the corresponding ester of **S66**. *Tert*-butyl 6-chloropyridazine-3-carboxylate (16 mg, 0.07 mmol, 1.00 equiv.) and *i*-$Pr_2NEt$ (29 µL, 0.15 mmol, 2.00 equiv.) were added and the mixture was stirred at 80 °C for 5 days. Purification by reversed-phase column chromatography (15.5 g C18, 7 to 70% MeCN in $H_2O$ with 0.05% TFA in 15 min) yielded **S66** as an inseparable mixture with *tert*-butyl 6-hydroxypyridazine-3-carboxylate and was used without further purification. To a solution of **S66** (24 mg, approx. 50% purity according to LC/MS, 0.04 mmol, 1.00 equiv.) in anhydrous DMF (0.5 mL) was added HATU (18 mg, 0.05 mmol, 1.20 equiv.), 4-([trans-4-aminocyclohexyl]oxy)-2-chlorobenzonitrile hydrochloride (12 mg, 0.04 mmol, 1.00 equiv.), and *i*-$Pr_2NEt$ (21 µL, 0.12 mmol, 3.00 equiv.) at rt under argon. The reaction was stirred at rt for 18 h. The mixture was poured into water (150 mL) and extracted with EtOAc (3 × 50 mL). The combined organic layers were washed with 5% aqueous LiCl (2 × 50 mL), water (50 mL), and brine (50 mL), dried with $MgSO_4$, filtered, and concentrated *in vacuo*. Purification by column chromatography (4 g silica, 60 to 80% EtOAc in *n*-heptane in 4 min, then 0 to 9% MeOH in $CH_2Cl_2$ in 10 min) yielded 2.4 mg (0.01 mmol, 7% over two steps) of **28** as a light-yellow powder after lyophilization from MeCN/$H_2O$.

**TLC** $R_f$ = 0.40 (5% MeOH in $CH_2Cl_2$).

**HRMS** (ESI) *m/z*: $(M+Na)^+$ calc. for $C_{40}H_{40}FN_{10}NaO_7Cl$: 849.2646; found: 849.2650.

**$^1$H NMR** (400 MHz, $CDCl_3$) δ 8.05 (d, *J* = 9.5 Hz, 1H), 8.02 (s, 1H), 7.86 (d, *J* = 8.2 Hz, 1H), 7.56 (d, *J* = 8.7 Hz, 1H), 7.51 (d, *J* = 10.8 Hz, 1H), 7.40 (d, *J* = 7.2 Hz, 1H), 7.04 – 6.97 (m, 2H), 6.85 (dd, *J* = 8.8, 2.4 Hz, 1H), 4.94 (dd, *J* = 12.2, 5.3 Hz, 1H), 4.37 – 4.27 (m, 1H), 4.13 – 4.01 (m, 1H), 3.89 – 3.76 (m, 4H), 3.62 – 3.42 (m, 8H), 3.38 – 3.23 (m, 4H), 2.96 – 2.67 (m, 3H), 2.27 – 2.10 (m, 5H), 1.75 – 1.62 (m, 2H), 1.53 – 1.42 (m, 2H) ppm.

**$^{19}$F{$^1$H} NMR** (377 MHz, $CDCl_3$) δ –110.8 ppm.

**$^{13}$C NMR** (151 MHz, $CDCl_3$) δ 170.8, 168.0, 166.9, 166.3, 163.5, 162.7, 161.7, 160.4, 158.5 (d, *J* = 256.2 Hz), 145.7 (d, *J* = 8.9 Hz), 145.1, 138.5, 135.2, 129.1, 127.2, 125.0 (d, *J* = 9.6 Hz), 117.0, 116.6, 114.8, 114.0 (d, *J* = 4.4 Hz), 112.5, 112.5 (d, *J* = 24.6 Hz), 105.0, 75.8, 49.9, 49.9, 49.6, 47.3, 46.7, 46.4, 44.6, 31.6, 30.2, 29.8, 22.8 ppm.

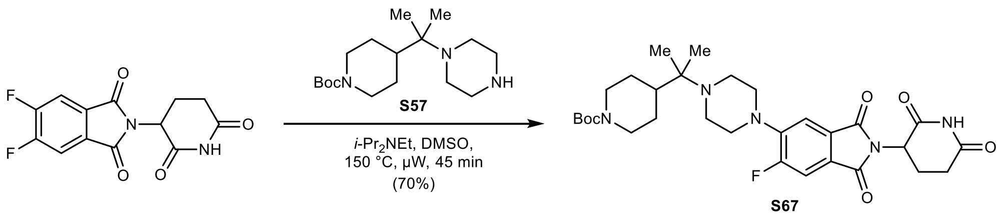

***tert*-Butyl 4-(2-(4-(2-(2,6-dioxopiperidin-3-yl)-6-fluoro-1,3-dioxoisoindolin-5-yl)piperazin-1-yl)propan-2-yl)piperidine-1-carboxylate (S67):** To a solution of 2-(2,6-dioxopiperidin-3-yl)-5,6-difluoroisoindoline-1,3-dione (50 mg, 0.17 mmol, 1.00 equiv.) in anhydrous DMSO (1.0 mL) was added **S57** (53 mg, 0.17 mmol, 1.00 equiv.) and *i*-$Pr_2NEt$ (59 µL, 0.34 mmol, 2.00 equiv.) at rt under argon. The reaction was stirred at 150 °C for 45 min under microwave irradiation. The mixture was diluted with water (100 mL) and extracted with EtOAc (3 × 30 mL). The combined organic layers were washed with water (50 mL) and brine (50 mL), dried with $MgSO_4$, filtered, and concentrated *in vacuo*. Purification by column chromatography (12 g silica, 10 to 65% EtOAc in *n*-heptane in 12 min) yielded 69 mg (0.12 mmol, 70%) of **S67** as a yellow powder.

**TLC** $R_f$ = 0.60 (75% EtOAc in *n*-heptane).

**LC/MS** (ESI) *m/z*: $(M+H)^+$ 586.3.

**$^1$H NMR** (400 MHz, $CDCl_3$) δ 8.19 (s, 1H), 7.45 (d, *J* = 11.2 Hz, 1H), 7.36 (d, *J* = 7.3 Hz, 1H), 4.93 (dd, *J* = 12.2, 5.3 Hz, 1H), 4.33 – 4.04 (m, 2H), 3.32 – 3.15 (m, 4H), 2.96 – 2.67 (m, 7H), 2.67 – 2.51 (m, 2H), 2.13 (ddt, *J* = 9.8, 4.9, 2.2 Hz, 1H), 1.83 – 1.69 (m, 2H), 1.69 – 1.57 (m, 1H), 1.45 (s, 9H), 1.27 – 1.11 (m, 2H), 0.93 (s, 6H) ppm.

**$^{19}$F{$^1$H} NMR** (377 MHz, $CDCl_3$) δ –110.9 ppm.

**$^{13}$C NMR** (101 MHz, $CDCl_3$) δ 171.0, 168.2, 167.1, 166.6 (d, *J* = 2.3 Hz), 158.2 (d, *J* = 255.8 Hz), 155.0, 146.1 (d, *J* = 8.9 Hz), 129.1 (d, *J* = 2.5 Hz), 123.8 (d, *J* = 9.6 Hz), 113.6 (d, *J* = 5.2 Hz), 112.2 (d, *J* = 25.3 Hz), 79.4, 57.9, 51.1, 51.0, 49.5, 45.1, 44.8, 43.2, 31.5, 28.6, 27.0, 22.8, 19.4 ppm.

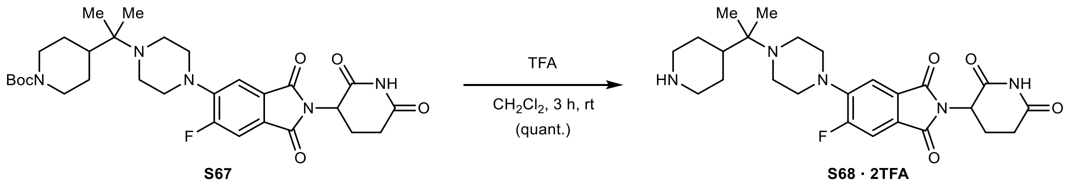

**2-(2,6-Dioxopiperidin-3-yl)-5-fluoro-6-(4-(2-(piperidin-4-yl)propan-2-yl)piperazin-1-yl) isoindoline-1,3-dione (S68 · 2TFA):** To a solution of **S67** (69 mg, 0.12 mmol, 1.00 equiv.) in $CH_2Cl_2$ (1.0 mL) was added trifluoroacetic acid (0.5 mL, 6.54 mmol, 55.2 equiv.) slowly at 0 °C. After 5 min, the reaction was allowed to warm to rt and stirred for 3 h. The mixture was concentrated *in vacuo* – residual trifluoroacetic acid was removed by co-evaporation with MeOH (3 × 5 mL) and drying *in vacuo*, yielding 85 mg (0.12 mmol, quantitative) of **S68 · 2TFA** as a pale-yellow powder.

**TLC** $R_f$ = 0.18, streaky (50% MeOH with 0.5% $NH_4OH$ in $CH_2Cl_2$).

**LC/MS** (ESI) *m/z*: $(M+H)^+$ 486.2.

**$^1$H NMR** (400 MHz, $CD_3OD$) δ 7.63 (d, *J* = 10.9 Hz, 1H), 7.57 (d, *J* = 7.3 Hz, 1H), 5.11 (dd, *J* = 12.5, 5.4 Hz, 1H), 3.97 – 3.36 (m, 10H), 3.09 (td, *J* = 13.1, 2.7 Hz, 2H), 2.95 – 2.64 (m, 3H), 2.38 – 2.27 (m, 1H), 2.13 (dtd, *J* = 10.0, 5.9, 2.8 Hz, 1H), 2.10 – 2.01 (m, 2H), 1.71 (qd, *J* = 13.2, 3.7 Hz, 2H), 1.45 (s, 6H) ppm. **Note:** The NH peaks are in exchange with $CD_3OD$ in the spectrum.

**$^{19}$F{$^1$H} NMR** (377 MHz, $CD_3OD$) δ –77.2 (TFA), –113.2 ppm.

**$^{13}$C NMR** (101 MHz, $CD_3OD$) δ 174.5, 171.4, 168.0, 167.6, 159.8 (d, *J* = 254.7 Hz), 145.0 (d, *J* = 9.7 Hz), 130.5 (d, *J* = 2.9 Hz), 127.5 (d, *J* = 9.8 Hz), 115.1 (d, *J* = 3.9 Hz), 113.0 (d, *J* = 25.1 Hz), 71.2, 50.8, 48.3, 48.2, 47.9, 44.9, 39.9, 32.2, 25.0, 23.6, 18.8 ppm.

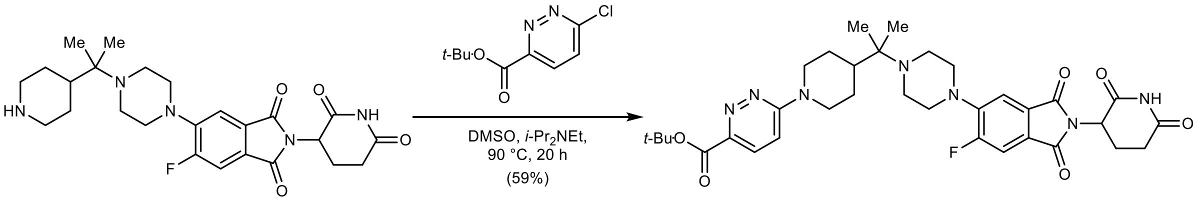

***tert*-Butyl 6-(4-(2-(4-(2-(2,6-dioxopiperidin-3-yl)-6-fluoro-1,3-dioxoisoindolin-5-yl)piperazin-1-yl)propan-2-yl)piperidin-1-yl)pyridazine-3-carboxylate (S69):** To a solution of **S68 · 2TFA** (66 mg, 0.09 mmol, 1.00 equiv.) in anhydrous DMSO (1.0 mL) was added *tert*-butyl 6-chloropyridazine-3-carboxylate (60 mg, 0.28 mmol, 3.00 equiv.) and *i*-$Pr_2NEt$ (64 µL, 0.37 mmol, 4.00 equiv.) at rt under argon. The reaction was stirred at 90 °C for 20 h. The mixture was diluted with water (150 mL) and extracted with EtOAc (4 × 50 mL). The combined organic layers were washed with water (50 mL) and brine (50 mL), dried with $MgSO_4$, filtered, and concentrated *in vacuo*. Purification by column chromatography (12 g silica, 0 to 6% MeOH with 2.5% $NH_4OH$ in $CH_2Cl_2$ in 9 min) yielded 36 mg (0.05 mmol, 59%) of **S69** as a yellow solid.

**TLC** $R_f$ = 0.71 (10% MeOH with 2.5% $NH_4OH$ in $CH_2Cl_2$).

**LC/MS** (ESI) *m/z*: $(M+H)^+$ 664.3.

**$^1$H NMR** (400 MHz, $CDCl_3$) δ 8.40 (s, 1H), 7.69 (d, *J* = 9.6 Hz, 1H), 7.34 (d, *J* = 11.1 Hz, 1H), 7.25 (d, *J* = 7.2 Hz, 1H), 6.73 (d, *J* = 9.6 Hz, 1H), 4.83 (dd, *J* = 12.0, 5.3 Hz, 1H), 4.62 – 4.44 (m, 2H), 3.22 – 3.05 (m, 4H), 2.90 – 2.52 (m, 9H), 2.07 – 1.96 (m, 1H), 1.87 – 1.65 (m, 3H), 1.50 (s, 9H), 1.29 – 1.15 (m, 2H), 0.83 (s, 6H) ppm.

**$^{19}$F{$^1$H} NMR** (377 MHz, $CDCl_3$) δ –110.9 ppm.

**$^{13}$C NMR** (101 MHz, $CDCl_3$) δ 171.2, 168.4, 167.1, 166.5 (d, *J* = 2.4 Hz), 164.0, 159.5, 158.2 (d, *J* = 255.9 Hz), 146.0 (d, *J* = 8.8 Hz), 143.9, 129.1 (d, *J* = 2.6 Hz), 128.5, 123.8 (d, *J* = 9.7 Hz), 113.6 (d, *J* = 5.1 Hz), 112.2 (d, *J* = 25.2 Hz), 110.5, 82.0, 57.9, 51.0, 50.9, 49.5, 45.9, 45.1, 43.1, 31.5, 28.3, 26.7, 22.7 (d, *J* = 9.9 Hz), 19.3 ppm.

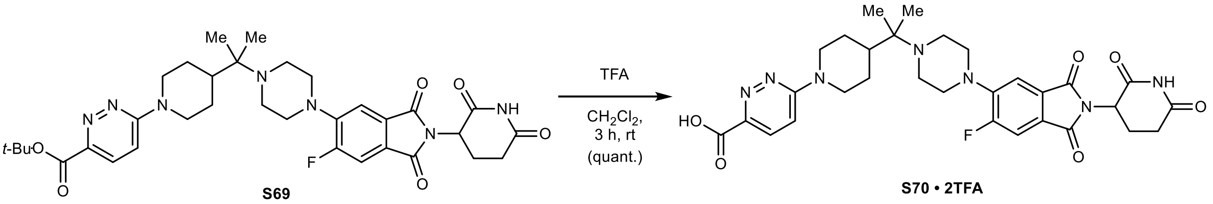

**6-(4-(2-(4-(2-(2,6-Dioxopiperidin-3-yl)-6-fluoro-1,3-dioxoisoindolin-5-yl)piperazin-1-yl)propan-2-yl)piperidin-1-yl)pyridazine-3-carboxylic acid (S70 · TFA):** To a solution of **S69** (36 mg, 0.05 mmol, 1.00 equiv.) in $CH_2Cl_2$ (1.0 mL) was added trifluoroacetic acid (0.25 mL, 3.24 mmol, 60.0 equiv.) slowly at 0 °C. After 5 min, the reaction was allowed to warm to rt and stirred for 3 h. The mixture was concentrated *in vacuo* – residual trifluoroacetic acid was removed by co-evaporation with MeOH (2 × 5 mL) and lyophilization from MeCN/$H_2O$, yielding 39 mg (0.05 mmol, quantitative) of **S70 · 2TFA** as a yellow solid.

**TLC** $R_f$ = 0.13, streaky (20% MeOH in $CH_2Cl_2$).

**LC/MS** (ESI) *m/z*: $(M+H)^+$ 608.3.

**$^1$H NMR** (400 MHz, $CD_3OD$) δ 8.06 (d, *J* = 9.5 Hz, 1H), 7.59 (d, *J* = 10.5 Hz, 2H), 7.53 (d, *J* = 7.0 Hz, 1H), 5.11 (dd, *J* = 12.6, 5.4 Hz, 1H), 4.73 – 4.54 (m, 2H), 4.09 – 3.37 (m, 8H), 3.28 – 3.10 (m, 2H), 2.88 (ddd, *J* = 18.3, 13.7, 5.0 Hz, 1H), 2.81 – 2.64 (m, 2H), 2.47 – 2.30 (m, 1H), 2.23 – 2.09 (m, 1H), 2.09 – 1.91 (m, 2H), 1.71 – 1.54 (m, 2H), 1.46 (s, 6H) ppm. **Note:** The proton corresponding to the carboxylic acid and the NH peaks are in exchange with $CD_3OD$ in the spectrum.

**$^{19}$F{$^1$H} NMR** (377 MHz, $CD_3OD$) δ –77.1 (TFA), –113.0 ppm.

**$^{13}$C NMR** (101 MHz, $CD_3OD$) δ 173.9, 170.8, 167.3, 167.0, 164.3, 158.8, 157.7 (d, *J* = 255.3 Hz), 144.4 (d, *J* = 9.3 Hz), 142.4, 131.1, 129.8, 126.6 (d, *J* = 9.7 Hz), 117.0, 114.4 (d, *J* = 3.7 Hz), 112.3 (d, *J* = 24.8 Hz), 71.0, 50.1, 47.6, 47.6, 47.2, 45.6, 41.3, 31.5, 26.9, 23.0, 18.4 ppm.

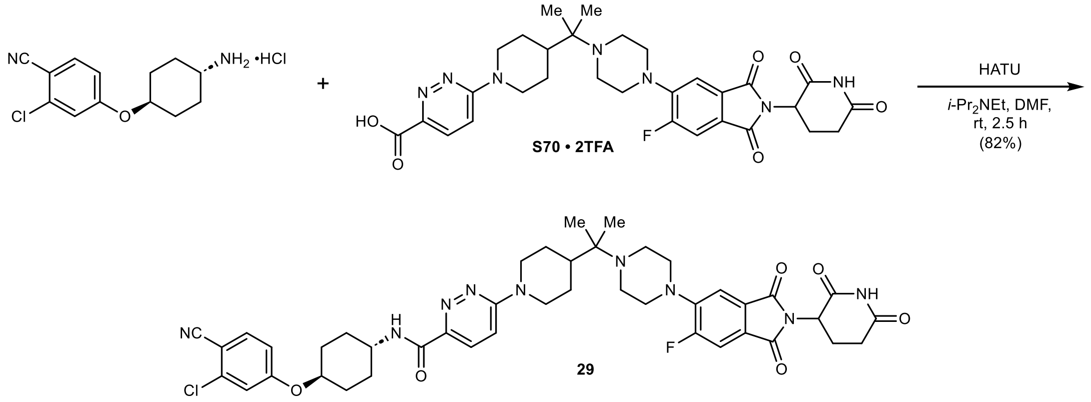

***N*-((1*r*,4*r*)-4-(3-Chloro-4-cyanophenoxy)cyclohexyl)-6-(4-(2-(4-(2-(2,6-dioxopiperidin-3-yl)-6-fluoro-1,3-dioxoisoindolin-5-yl)piperazin-1-yl)propan-2-yl)piperidin-1-yl)pyridazine-3-carboxamide (29):** To a solution of **S70 · 2TFA** (39 mg, 0.05 mmol, 1.00 equiv.) in anhydrous DMF (0.5 mL) was added HATU (25 mg, 0.07 mmol, 1.20 equiv.), 4-([trans-4-aminocyclohexyl] oxy)-2-chlorobenzonitrile hydrochloride (17 mg, 0.06 mmol, 1.10 equiv.), and *i*-$Pr_2NEt$ (47 µL, 0.27 mmol, 5.00 equiv.) at rt under argon. The reaction was stirred at rt for 2.5 h. The mixture was poured into water (200 mL) and extracted with EtOAc (3 × 50 mL). The combined organic layers were washed with 5% aqueous LiCl (50 mL), water (50 mL), and brine (50 mL), dried with $MgSO_4$, filtered, and concentrated *in vacuo*. Purification by column chromatography (12 g silica, 0 to 4% MeOH in $CH_2Cl_2$ in 8 min) yielded 39 mg (0.05 mmol, 82%) of **29** as a pale-yellow powder after lyophilization from $MeCN/H_2O$.

**TLC** $R_f$ = 0.51 (5% MeOH in $CH_2Cl_2$).

**HRMS** (ESI) *m/z*: $(M+Na)^+$ calc. for $C_{43}H_{47}FN_9NaO_6Cl$: 862.3214; found: 862.3215.

**$^1$H NMR** (400 MHz, $CDCl_3$) δ 8.18 (s, 1H), 7.96 (d, *J* = 9.6 Hz, 1H), 7.88 (d, *J* = 8.2 Hz, 1H), 7.55 (d, *J* = 8.7 Hz, 1H), 7.46 (d, *J* = 11.1 Hz, 1H), 7.37 (d, *J* = 7.3 Hz, 1H), 7.00 (d, *J* = 2.4 Hz, 1H), 6.97 (d, *J* = 9.6 Hz, 1H), 6.85 (dd, *J* = 8.8, 2.4 Hz, 1H), 4.94 (dd, *J* = 12.3, 5.3 Hz, 1H), 4.66 – 4.54 (m, 2H), 4.31 (td, *J* = 10.0, 5.0 Hz, 1H), 4.12 – 3.99 (m, 1H), 3.32 – 3.19 (m, 4H), 3.02 – 2.86 (m, 3H), 2.86 – 2.67 (m, 6H), 2.26 – 2.07 (m, 5H), 1.99 – 1.81 (m, 3H), 1.75 – 1.64 (m, 2H), 1.52 – 1.41 (m, 2H), 1.41 – 1.30 (m, 2H), 0.97 (s, 6H) ppm.

**$^{19}$F{$^1$H} NMR** (377 MHz, $CDCl_3$) δ –110.9 ppm.

**$^{13}$C NMR** (101 MHz, $CDCl_3$) δ 170.9, 168.2, 167.1, 166.5 (d, *J* = 2.4 Hz), 163.0, 161.8, 160.2, 158.2 (d, *J* = 255.9 Hz), 146.0 (d, *J* = 8.9 Hz), 144.1, 138.4, 135.2, 129.1 (d, *J* = 2.7 Hz), 126.9, 123.9 (d, *J* = 9.6 Hz), 117.0, 116.6, 114.8, 113.6 (d, *J* = 5.1 Hz), 112.4, 112.1, 104.9, 75.8, 57.9, 51.0, 51.0, 49.5, 47.2, 46.1, 45.1, 43.1, 31.6, 30.2, 29.8, 26.7, 22.8, 19.4 ppm.

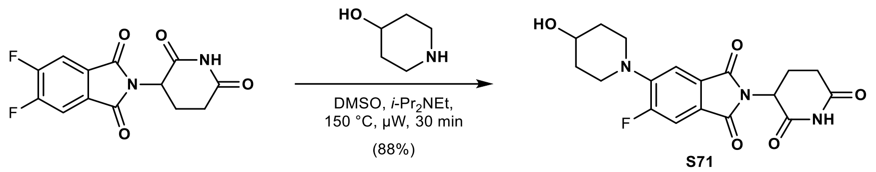

**2-(2,6-Dioxopiperidin-3-yl)-5-fluoro-6-(4-hydroxypiperidin-1-yl)isoindoline-1,3-dione (S71):** To a solution of 2-(2,6-dioxopiperidin-3-yl)-5,6-difluoroisoindoline-1,3-dione (500 mg, 1.70 mmol, 1.00 equiv.) in anhydrous DMSO (5.0 mL) was added piperidin-4-ol (189 mg, 1.87 mmol, 1.10 equiv.) and *i*-$Pr_2NEt$ (592 µL, 3.40 mmol, 2.00 equiv.) at rt under argon. The reaction was stirred at 150 °C for 30 min under microwave irradiation. The mixture was diluted with water (200 mL), extracted with EtOAc (2 × 50 mL) and 10% MeOH in $CH_2Cl_2$ (2 × 50 mL). The combined organic layers were washed with brine (2 × 50 mL), dried with $MgSO_4$, filtered, and concentrated *in vacuo*. Purification by column chromatography (40 g silica, 0 to 6% MeOH in $CH_2Cl_2$ in 10 min) yielded 561 mg (1.49 mmol, 88%) of **S71** as a yellow solid.

**TLC** $R_f$ = 0.50 (10% MeOH in $CH_2Cl_2$).

**LC/MS** (ESI) *m/z*: $(M+H)^+$ 376.1.

**$^1$H NMR** (400 MHz, $CDCl_3$) δ 8.41 (s, 1H), 7.45 (d, *J* = 11.1 Hz, 1H), 7.39 (d, *J* = 7.3 Hz, 1H), 4.97 – 4.87 (m, 1H), 3.99 – 3.88 (m, 1H), 3.53 (dt, *J* = 11.1, 4.6 Hz, 2H), 3.05 (ddd, *J* = 12.4, 9.0, 3.2 Hz, 2H), 2.93 – 2.67 (m, 3H), 2.19 – 2.09 (m, 1H), 2.09 – 1.99 (m, 2H), 1.81 (d, *J* = 3.6 Hz, 1H), 1.74 (dtd, *J* = 12.9, 8.7, 3.7 Hz, 2H) ppm.

**$^{19}$F{$^1$H} NMR** (377 MHz, $CDCl_3$) δ –110.9 ppm.

**$^{13}$C NMR** (101 MHz, $CDCl_3$) δ 171.1, 168.3, 167.1, 166.5 (d, *J* = 2.6 Hz), 158.1 (d, *J* = 255.6 Hz), 146.1 (d, *J* = 9.2 Hz), 129.0 (d, *J* = 2.8 Hz), 123.7 (d, *J* = 9.7 Hz), 113.9 (d, *J* = 4.9 Hz), 112.2 (d, *J* = 25.3 Hz), 66.8, 49.4, 47.7, 47.6, 34.1, 34.1, 31.4, 22.7 ppm.

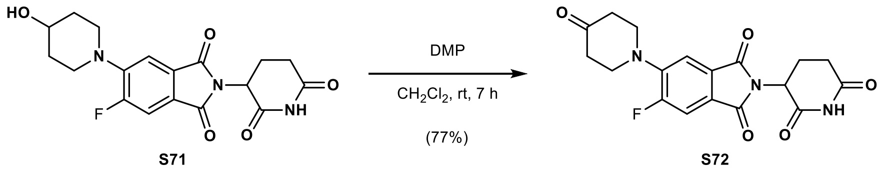

**2-(2,6-Dioxopiperidin-3-yl)-5-fluoro-6-(4-oxopiperidin-1-yl)isoindoline-1,3-dione (S72):** To a solution of **S71** (549 mg, 1.46 mmol, 1.00 equiv.) in anhydrous $CH_2Cl_2$ (10 mL) was added Dess–Martin periodinane (744 mg, 1.76 mmol, 1.20 equiv.) at 0 °C under argon. After 5 min, the reaction was allowed to warm to rt and stirred for 7 h. The mixture was poured into saturated aqueous $NaHCO_3$ (200 mL) and extracted with $CH_2Cl_2$ (3 × 50 mL). The combined organic layers were washed with brine (50 mL), dried with $MgSO_4$, filtered, and concentrated *in vacuo*. Purification by column chromatography (40 g silica, 0 to 10% MeOH in $CH_2Cl_2$ in 13 min), followed by reversed-phase column chromatography (50 g C18, 10 to 60% MeCN in $H_2O$ in 15 min) yielded 420 mg (1.12 mmol, 77%) of **S72** as a yellow powder. **Note:** A reductive work-up procedure using $Na_2S_2O_3$ is recommended to avoid iodine-containing side products.

**TLC** $R_f$ = 0.45 (10% MeOH in $CH_2Cl_2$).

**LC/MS** (ESI) *m/z*: $(M+H)^+$ 374.1.

**$^1$H NMR** (400 MHz, DMSO-$d_6$) δ 11.11 (s, 1H), 7.77 (d, *J* = 11.5 Hz, 1H), 7.54 (d, *J* = 7.6 Hz, 1H), 5.11 (dd, *J* = 12.8, 5.4 Hz, 1H), 3.64 (t, *J* = 6.1 Hz, 4H), 2.89 (ddd, *J* = 17.0, 13.9, 5.4 Hz, 1H), 2.64 – 2.51 (m, 6H), 2.07 – 1.99 (m, 1H) ppm.

**$^{19}$F{$^1$H} NMR** (377 MHz, $CDCl_3$) δ –111.3 ppm.

$^{13}$**C NMR** (101 MHz, DMSO-$d_6$) δ 207.0, 172.8, 169.9, 166.6, 166.2, 156.9 (d, *J* = 252.2 Hz), 144.4 (d, *J* = 8.9 Hz), 128.8, 123.1 (d, *J* = 9.7 Hz), 113.9 (d, *J* = 4.7 Hz), 112.1 (d, *J* = 25.3 Hz), 49.1, 48.6, 48.6, 40.5, 40.5, 30.9, 22.1 ppm.

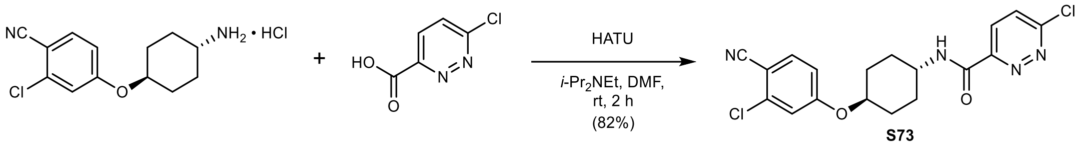

**6-Chloro-*N*-((1*r*,4*r*)-4-(3-chloro-4-cyanophenoxy)cyclohexyl)pyridazine-3-carboxamide (S73):** To a solution of 4-([trans-4-aminocyclohexyl]oxy)-2-chlorobenzonitrile hydrochloride (500 mg, 1.74 mmol, 1.00 equiv.) in anhydrous DMF (8.0 mL) was added 6-chloropyridazine-3-carboxylic acid (290 mg, 1.83 mmol, 1.05 equiv.), HATU (795 mg, 2.09 mmol, 1.20 equiv.), and *i*-$Pr_2NEt$ (0.91 mL, 5.22 mmol, 3.00 equiv.) at rt under argon. The reaction was stirred at rt for 2 h. The mixture was poured into water (200 mL) and extracted with EtOAc (3 × 50 mL). The combined organic layers were washed with 5% aqueous LiCl (3 × 50 mL), water (2 × 50 mL), and brine (50 mL), dried with $MgSO_4$, filtered, and concentrated *in vacuo*. Purification by column chromatography (80 g silica, 30 to 100% EtOAc in *n*-heptane in 25 min) yielded 561 mg (1.43 mmol, 82%) of **S73** as an off-white solid.

**TLC** $R_f$ = 0.70 (75% EtOAc in *n*-heptane).

**LC/MS** (ESI) *m/z*: $(M+H)^+$ 391.1.

$^{1}$**H NMR** (400 MHz, DMSO-$d_6$) δ 9.14 (d, *J* = 8.2 Hz, 1H), 8.22 (d, *J* = 8.9 Hz, 1H), 8.10 (d, *J* = 8.9 Hz, 1H), 7.85 (d, *J* = 8.8 Hz, 1H), 7.39 (d, *J* = 2.4 Hz, 1H), 7.14 (dd, *J* = 8.8, 2.5 Hz, 1H), 4.54 (td, *J* = 10.4, 5.2 Hz, 1H), 3.91 (tdt, *J* = 11.7, 8.1, 4.0 Hz, 1H), 2.12 (d, *J* = 12.1 Hz, 2H), 1.90 (d, *J* = 12.7 Hz, 2H), 1.70 (qd, *J* = 13.3, 3.2 Hz, 2H), 1.51 (qd, *J* = 13.2, 3.5 Hz, 2H) ppm.

$^{13}$**C NMR** (101 MHz, DMSO-$d_6$) δ 161.8, 161.2, 158.1, 152.8, 137.0, 135.7, 130.2, 129.0, 116.8, 116.4, 115.5, 103.2, 75.5, 47.7, 29.9, 29.2 ppm.

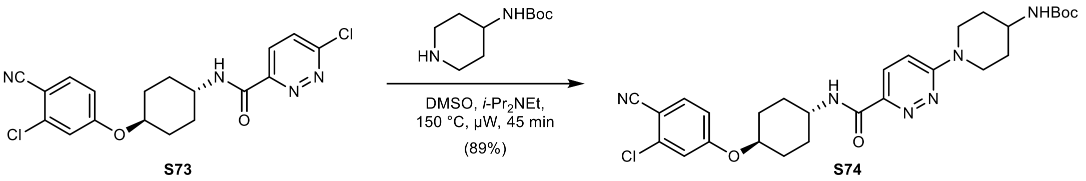

***tert*-Butyl (1-(6-(((1*r*,4*r*)-4-(3-chloro-4-cyanophenoxy)cyclohexyl)carbamoyl)pyridazin-3-yl) piperidin-4-yl)carbamate (S74):** To a solution of **S73** (250 mg, 0.64 mmol, 1.00 equiv.) in anhydrous DMSO (3.0 mL) was added *tert*-butyl piperidin-4-ylcarbamate (141 mg, 0.70 mmol, 1.10 equiv.) and *i*-$Pr_2NEt$ (223 µL, 1.28 mmol, 2.00 equiv.) at rt under argon. The reaction was stirred at 150 °C for 45 min under microwave irradiation. The mixture was diluted with water (200 mL) and extracted with EtOAc (3 × 50 mL). The combined organic layers were washed with water (50 mL) and brine (50 mL), dried with $MgSO_4$, filtered, and concentrated *in vacuo*. Purification by column chromatography (24 g silica, 0 to 5% MeOH in $CH_2Cl_2$ in 11 min) yielded 317 mg (0.57 mmol, 89%) of **S74** as an off-white solid.

**TLC** $R_f$ = 0.47 (5% MeOH in $CH_2Cl_2$).

**LC/MS** (ESI) *m/z*: $(M+H)^+$ 555.2.

$^{1}$**H NMR** (400 MHz, $CDCl_3$) δ 7.98 (d, *J* = 9.6 Hz, 1H), 7.85 (d, *J* = 8.2 Hz, 1H), 7.55 (d, *J* = 8.7 Hz, 1H), 7.03 – 6.94 (m, 2H), 6.85 (dd, *J* = 8.8, 2.4 Hz, 1H), 4.55 – 4.45 (m, 1H), 4.41 (dt, *J* = 13.5, 3.3 Hz,

2H), 4.32 (tt, $J$ = 9.9, 3.8 Hz, 1H), 4.12 – 3.98 (m, 1H), 3.89 – 3.68 (m, 1H), 3.18 (ddd, $J$ = 14.0, 11.7, 2.8 Hz, 2H), 2.28 – 2.03 (m, 6H), 1.76 – 1.64 (m, 2H), 1.45 (s, 13H) ppm.

**$^{13}$C NMR** (101 MHz, $CDCl_3$) δ 162.8, 161.8, 160.1, 155.2, 144.4, 138.5, 135.2, 127.0, 117.0, 116.6, 114.8, 112.3, 104.9, 79.8, 75.8, 48.0, 47.2, 44.1, 32.1, 30.2, 29.8, 28.5 ppm.

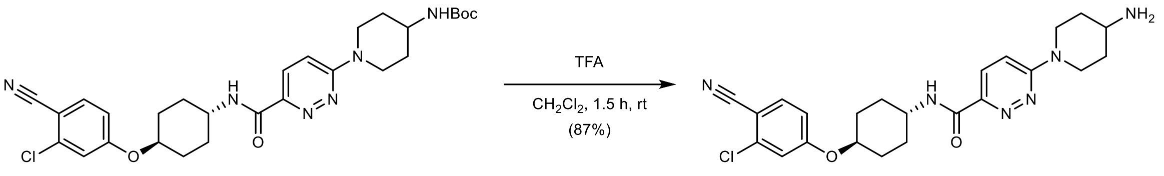

**S74** **S75**

**6-(4-Aminopiperidin-1-yl)-*N*-((1*r*,4*r*)-4-(3-chloro-4-cyanophenoxy)cyclohexyl)pyridazine-3-carboxamide (S75):** To a solution of **S74** (310 mg, 0.56 mmol, 1.00 equiv.) in $CH_2Cl_2$ (4.5 mL) was added trifluoroacetic acid (1.50 mL, 19.6 mmol, 35.0 equiv.) slowly at 0 °C. After 5 min, the reaction was allowed to warm to rt and stirred for 1.5 h. The mixture was concentrated *in vacuo* – residual trifluoroacetic acid was removed by co-evaporation with MeOH (4 × 5 mL), yielding **S75 · 2TFA** as a hygroscopic salt. The crude product was dissolved in 10% MeOH in $CH_2Cl_2$ (10 mL), poured into saturated aqueous $Na_2CO_3$ (50 mL), and extracted with 10% MeOH in $CH_2Cl_2$ (3 × 25 mL). The combined organic layers were dried with $MgSO_4$, filtered, and concentrated *in vacuo*, yielding 220 mg (0.48 mmol, 87%) of **S75** as an off-white solid.

**TLC** $R_f$ = 0.19, streaky (20% MeOH with 0.5% $NH_4OH$ in $CH_2Cl_2$).

**LC/MS** (ESI) *m/z*: $(M+H)^+$ 455.2.

**$^1$H NMR** (400 MHz, DMSO-$d_6$) δ 8.58 (d, $J$ = 8.2 Hz, 1H), 7.85 (d, $J$ = 8.8 Hz, 1H), 7.79 (d, $J$ = 9.5 Hz, 1H), 7.39 (d, $J$ = 2.4 Hz, 1H), 7.34 (d, $J$ = 9.7 Hz, 1H), 7.13 (dd, $J$ = 8.8, 2.5 Hz, 1H), 4.53 (tt, $J$ = 9.4, 4.2 Hz, 1H), 4.34 (dt, $J$ = 13.4, 3.3 Hz, 2H), 3.92 – 3.78 (m, 1H), 3.11 (ddd, $J$ = 13.8, 11.4, 2.8 Hz, 2H), 2.88 (td, $J$ = 9.8, 4.7 Hz, 1H), 2.16 – 2.05 (m, 2H), 1.97 – 1.85 (m, 2H), 1.83 – 1.57 (m, 4H), 1.57 – 1.43 (m, 2H), 1.28 – 1.14 (m, 2H) ppm. **Note:** The $NH_2$ peak is in exchange with residual water in the spectrum.

**$^{13}$C NMR** (101 MHz, DMSO-$d_6$) δ 162.4, 161.8, 159.8, 144.9, 137.1, 135.8, 126.6, 116.8, 116.5, 115.5, 113.0, 103.3, 75.6, 47.6, 47.2, 42.9, 29.9, 29.4, 28.9 ppm.

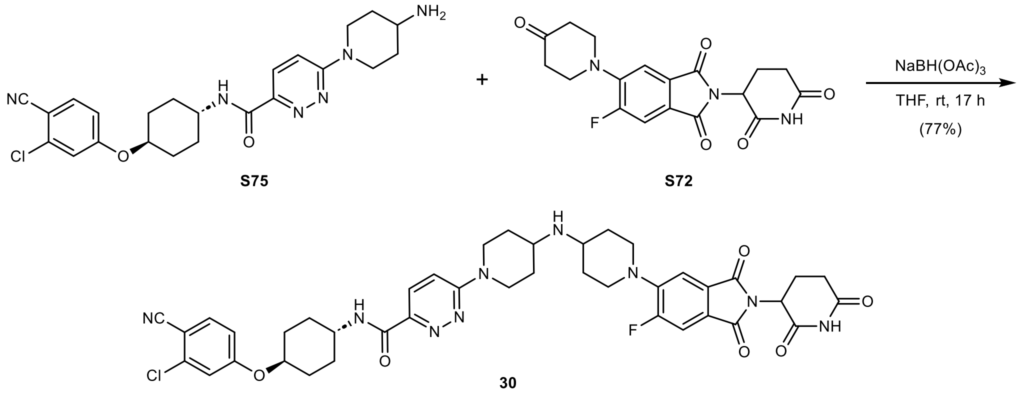

**S75** **S72**

**30**

***N*-((1*r*,4*r*)-4-(3-Chloro-4-cyanophenoxy)cyclohexyl)-6-(4-((1-(2-(2,6-dioxopiperidin-3-yl)-6-fluoro-1,3-dioxoisoindolin-5-yl)piperidin-4-yl)amino)piperidin-1-yl)pyridazine-3-carboxamide (30):** To a solution of **S75** (49 mg, 0.11 mmol, 1.00 equiv.) in anhydrous THF (1.5 mL) was added **S72** (40 mg, 0.11 mmol, 1.00 equiv.) and $NaBH(OAc)_3$ (45 mg, 0.21 mmol, 2.00 equiv.) at rt under argon. The reaction was stirred at rt for 17 h. The mixture was poured into aqueous $NaHCO_3$ (150 mL) and extracted with EtOAc (3 × 50 mL). The combined organic layers were washed with brine (50 mL), dried with $MgSO_4$, filtered, and concentrated *in vacuo*. Purification by column chromatography (12 g silica, 0 to 9% MeOH with 2.5% $NH_4OH$ in $CH_2Cl_2$ in 17 min) yielded 67 mg (0.08 mmol, 77%) of **30** as a yellow solid.

**TLC** $R_f$ = 0.52 (10% MeOH with 0.5% $NH_4OH$ in $CH_2Cl_2$).

**HRMS** (ESI) *m/z*: $(M+H)^+$ calc. for $C_{41}H_{44}FN_9O_6Cl$: 812.3082; found: 812.3082.

**$^1$H NMR** (400 MHz, $CDCl_3$) δ 7.98 (d, *J* = 9.6 Hz, 1H), 7.87 (d, *J* = 8.2 Hz, 1H), 7.55 (d, *J* = 8.7 Hz, 1H), 7.46 (d, *J* = 11.0 Hz, 1H), 7.38 (d, *J* = 7.4 Hz, 1H), 7.03 – 6.96 (m, 2H), 6.85 (dd, *J* = 8.8, 2.4 Hz, 1H), 4.93 (dd, *J* = 12.3, 5.3 Hz, 1H), 4.43 (dt, *J* = 13.4, 3.3 Hz, 2H), 4.32 (td, *J* = 10.0, 5.0 Hz, 1H), 4.11 – 3.98 (m, 1H), 3.64 (dt, *J* = 12.4, 3.1 Hz, 2H), 3.24 –3.12 (m, 2H), 3.08 – 2.68 (m, 7H), 2.25 – 2.10 (m, 5H), 2.10 – 1.97 (m, 4H), 1.75 – 1.62 (m, 2H), 1.61 – 1.37 (m, 6H) ppm.

**$^{19}$F{$^1$H} NMR** (377 MHz, $CDCl_3$) δ –110.9 ppm.

**$^{13}$C NMR** (101 MHz, $CDCl_3$) δ 171.0, 168.2, 167.2, 166.5, 162.9, 161.8, 160.2, 158.2 (d, *J* = 255.5 Hz), 146.2 (d, *J* = 9.0 Hz), 144.3, 138.4, 135.2, 129.1 (d, *J* = 2.5 Hz), 126.9, 123.8 (d, *J* = 9.4 Hz), 117.0, 116.6, 114.8, 114.0 (d, *J* = 4.9 Hz), 112.3 (d, *J* = 25.2 Hz), 112.3, 104.9, 75.8, 51.1, 50.8, 49.6, 49.4, 49.4, 47.2, 44.0, 33.2, 32.8, 31.6, 30.2, 29.8, 22.8 ppm.

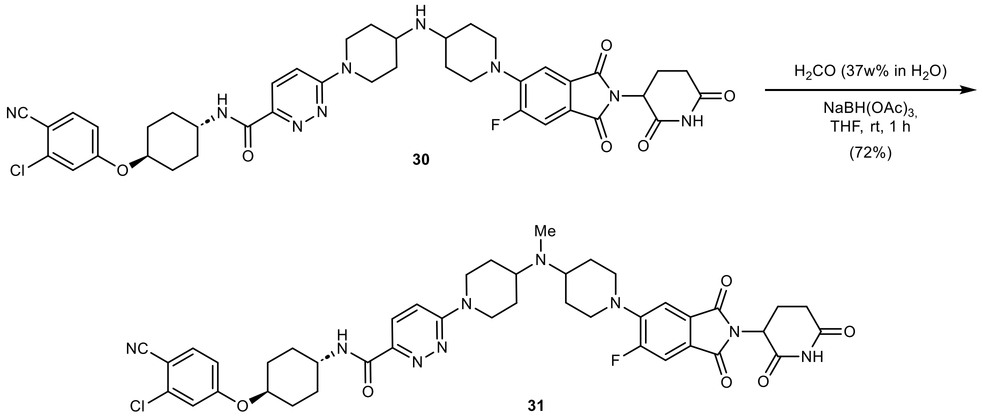

***N*-((1*r*,4*r*)-4-(3-Chloro-4-cyanophenoxy)cyclohexyl)-6-(4-((1-(2-(2,6-dioxopiperidin-3-yl)-6-fluoro-1,3-dioxoisoindolin-5-yl)piperidin-4-yl)(methyl)amino)piperidin-1-yl)pyridazine-3-carboxamide (31):** To a solution of **30** (20 mg, 0.02 mmol, 1.00 equiv.) in THF (0.5 mL) was added formaldehyde (18.3 µL, 37% in $H_2O$, 0.25 mmol, 10.0 equiv.) and $NaBH(OAc)_3$ (10 mg, 0.05 mmol, 2.50 equiv.) at rt. The reaction was stirred at rt for 1 h. The mixture was poured into aqueous $NaHCO_3$ (50 mL) and extracted with EtOAc (3 × 25 mL). The combined organic layers were washed with brine (25 mL), dried with $MgSO_4$, filtered, and concentrated *in vacuo*. The crude reaction mixture was subjected to column chromatography (4 g silica, 0 to 12% MeOH in $CH_2Cl_2$ in 12 min), but $^1$H NMR spectroscopy of the apparently pure material revealed 15% of the addition product of formaldehyde and **31** across the imide N-H bond (*N*-((1*r*,4*r*)-4-(3-chloro-4-cyanophenoxy)cyclohexyl)-6-(4-((1-(6-fluoro-2-(1-(hydroxymethyl)-2,6-dioxopiperidin-3-yl)-1,3-dioxoisoindolin-5-yl)piperidin-4-yl)(methyl)amino)piperidin-1-yl)pyridazine-3-carboxamide). To hydrolyze the side product, the mixture was suspended in MeCN (1.0 mL) and $H_2O$ with 0.01% formic acid (1.0 mL). After 1 h, two drops of formic acid were added and the mixture was stirred at rt for 60 h. The mixture was poured into aqueous $NaHCO_3$ (50 mL) and extracted with $CH_2Cl_2$ (3 × 25 mL). The combined organic layers were dried with $MgSO_4$, filtered, and concentrated *in vacuo*, yielding 15 mg (0.02 mmol, 72%) of **31** as a yellow solid after lyophilization from MeCN/$H_2O$.

**TLC** $R_f$ = 0.59 (10% MeOH in $CH_2Cl_2$).

**HRMS** (ESI) *m/z*: $(M+H)^+$ calc. for $C_{42}H_{46}FN_9O_6Cl$: 826.3238; found: 826.3239.

**$^1$H NMR** (400 MHz, $CDCl_3$) δ 8.21 (s, 1H), 7.98 (d, *J* = 9.6 Hz, 1H), 7.87 (d, *J* = 8.2 Hz, 1H), 7.55 (d, *J* = 8.7 Hz, 1H), 7.46 (d, *J* = 11.0 Hz, 1H), 7.38 (d, *J* = 7.3 Hz, 1H), 7.04 – 6.97 (m, 2H), 6.85 (dd, *J* = 8.8, 2.4 Hz, 1H), 4.93 (dd, *J* = 12.3, 5.3 Hz, 1H), 4.62 – 4.50 (m, 2H), 4.32 (td, *J* = 10.0, 5.0 Hz, 1H), 4.05 (td, *J* = 9.1, 5.5 Hz, 1H), 3.78 – 3.63 (m, 2H), 3.07 (td, *J* = 13.4, 1.9 Hz, 2H), 3.00 – 2.68 (m, 7H), 2.30 (s, 3H), 2.23 – 2.09 (m, 5H), 1.97 – 1.77 (m, 6H), 1.77 – 1.60 (m, 4H), 1.53 – 1.38 (m, 2H) ppm.

**$^{19}$F{$^1$H} NMR** (377 MHz, $CDCl_3$) δ –110.9 ppm.

**$^{13}$C NMR** (101 MHz, $CDCl_3$) δ 170.9, 168.2, 167.2, 166.5, 162.9, 161.8, 160.1, 158.2 (d, *J* = 255.7 Hz), 146.1 (d, *J* = 8.9 Hz), 144.3, 138.5, 135.2, 129.1, 127.0, 123.8 (d, *J* = 9.6 Hz), 117.0, 116.6, 114.8, 114.0 (d, *J* = 4.8 Hz), 112.3 (d, *J* = 25.1 Hz), 112.2, 104.9, 75.8, 57.8, 57.6, 50.3, 50.3, 49.6, 47.2, 44.9, 32.8, 31.6, 30.2, 29.8, 29.6, 29.3, 22.8 ppm.

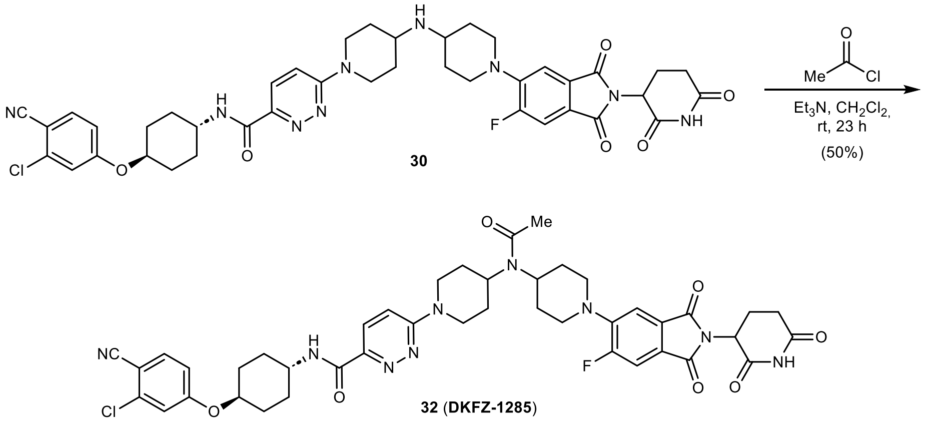

***N*-((1*r*,4*r*)-4-(3-Chloro-4-cyanophenoxy)cyclohexyl)-6-(4-(*N*-(1-(2-(2,6-dioxopiperidin-3-yl)-6-fluoro-1,3-dioxoisoindolin-5-yl)piperidin-4-yl)acetamido)piperidin-1-yl)pyridazine-3-carboxamide (32, DKFZ-1285):** To a solution of **30** (12 mg, 0.01 mmol, 1.00 equiv.) in anhydrous $CH_2Cl_2$ (0.5 mL) was added $Et_3N$ (7.2 µL, 0.05 mmol, 3.50 equiv.) and acetyl chloride (3.2 µL, 0.04 mmol, 3.00 equiv.) at rt under argon. The reaction was stirred at rt for 16 h. To drive the reaction to completion, $Et_3N$ (7.2 µL, 0.05 mmol, 3.50 equiv.) and acetyl chloride (3.2 µL, 0.04 mmol, 3.00 equiv.) were added. The reaction was stirred at rt for another 7 h. The mixture was poured into aqueous $NaHCO_3$ (25 mL) and extracted with $CH_2Cl_2$ (3 × 10 mL). The combined organic layers were dried with $MgSO_4$, filtered, and concentrated *in vacuo*. Purification by column chromatography (4 g silica, 0 to 9% MeOH in $CH_2Cl_2$ in 14 min) yielded 6 mg (0.01 mmol, 50%) of **32** (**DKFZ-1285**) as a yellow solid after lyophilization from MeCN/$H_2O$.

**TLC** $R_f$ = 0.30 (5% MeOH in $CH_2Cl_2$).

**HRMS** (ESI) *m/z*: $(M+H)^+$ calc. for $C_{43}H_{46}FN_9O_7Cl$: 854.3187; found: 854.3192.

**$^1$H NMR** (400 MHz, $CDCl_3$, 328.1 K) δ 8.03 (s, 1H), 8.00 (d, *J* = 9.4 Hz, 1H), 7.84 (d, *J* = 8.1 Hz, 1H), 7.55 (d, *J* = 8.7 Hz, 1H), 7.47 (d, *J* = 10.9 Hz, 1H), 7.38 (d, *J* = 7.2 Hz, 1H), 7.01 (d, *J* = 2.4 Hz, 1H), 7.00 – 6.94 (m, 1H), 6.85 (dd, *J* = 8.7, 2.4 Hz, 1H), 4.92 (dd, *J* = 12.2, 5.3 Hz, 1H), 4.78 – 4.50 (m, 2H), 4.33 (td, *J* = 9.9, 5.0 Hz, 1H), 4.13 – 4.00 (m, 1H), 3.90 – 3.30 (m, 4H), 3.08 (t, *J* = 12.8 Hz, 2H), 3.01 – 2.63 (m, 6H), 2.26 – 2.09 (m, 9H), 1.96 – 1.61 (m, 6H), 1.58 – 1.42 (m, 4H) ppm.

**$^{19}$F{$^1$H} NMR** (377 MHz, $CDCl_3$, 328.2 K) δ –111.1 ppm.

**$^{13}$C NMR** (101 MHz, $CDCl_3$, 328.1 K) δ 170.6, 170.0, 168.0, 167.0, 166.4, 163.0, 161.9, 160.2, 158.4 (d, *J* = 256.5 Hz), 145.8 (d, *J* = 8.7 Hz), 138.6, 135.2, 129.4 (d, *J* = 2.4 Hz), 127.1, 117.3, 116.4, 114.9, 114.1, 112.5, 112.3, 105.3, 76.1, 50.6, 50.6, 49.8, 47.4, 45.3, 31.6, 30.8, 30.2, 29.9, 29.4, 24.0, 22.9 ppm. **Note:** The $^{1}$H and $^{13}$C NMR show broadened signals at 298.1 K due to the presence of rotamers. Therefore, shifts are reported at 328.1 K. Nonetheless, four carbons cannot be detected due to signal broadening.

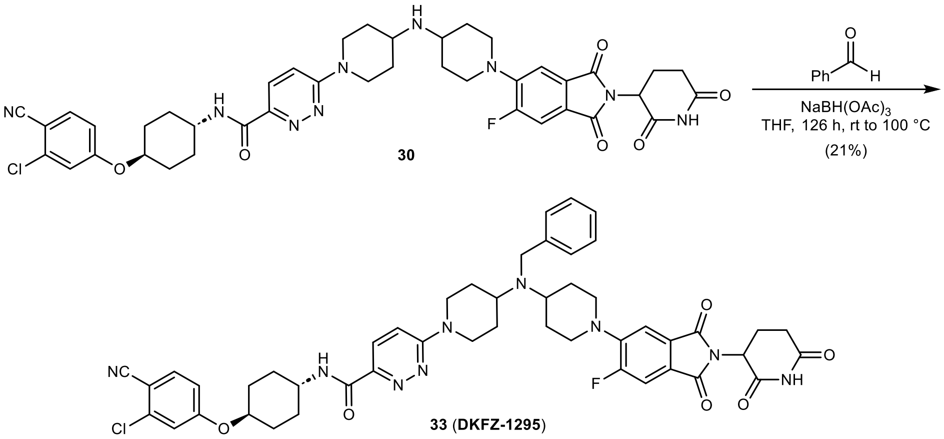

**6-(4-(Benzyl(1-(2-(2,6-dioxopiperidin-3-yl)-6-fluoro-1,3-dioxoisoindolin-5-yl)piperidin-4-yl) amino)piperidin-1-yl)-*N*-((1*r*,4*r*)-4-(3-chloro-4-cyanophenoxy)cyclohexyl)pyridazine-3-carboxamide (33, DKFZ-1295):** To a solution of **30** (17 mg, 0.02 mmol, 1.00 equiv.) in THF (1.0 mL) was added benzaldehyde (4.3 µL, 0.04 mmol, 2.00 equiv.) and $NaBH(OAc)_3$ (9 mg, 0.04 mmol, 2.00 equiv.) at rt. After 5 h at rt, benzaldehyde (21.3 µL, 0.21 mmol,10.0 equiv.) and $NaBH(OAc)_3$ (44 mg, 0.21 mmol, 10.0 equiv.) were added. The mixture was stirred for 15 h at rt, then for 28 h at 60 °C. To enhance solubility, $CH_2Cl_2$ (0.5 mL) was added and the mixture was continued stirring for 16 h at 40 °C. To drive the reaction to completion, DMF (0.2 mL), benzaldehyde (21.3 µL, 0.21 mmol, 10.0 equiv.), and $NaBH(OAc)_3$ (44 mg, 0.21 mmol, 10.0 equiv.) were added and the mixture was stirred for another 60 h at 45 °C, 1 h at 80 °C, and 1.5 h at 100 °C. The reaction was aborted. The mixture was poured into aqueous $NaHCO_3$ (25 mL) and extracted with EtOAc (3 × 10 mL). The combined organic layers were dried with $MgSO_4$, filtered, and concentrated *in vacuo*. Purification by column chromatography (4 g silica, 0 to 6% MeOH with 2.5% $NH_4OH$ in $CH_2Cl_2$ in 12 min) yielded 4 mg (0.01 mmol, 21%) of **33** (**DKFZ-1295**) as a pale-yellow solid after lyophilization from MeCN/$H_2O$.

**TLC** $R_f$ = 0.48 (5% MeOH with 2.5% $NH_4OH$ in $CH_2Cl_2$).

**HRMS** (ESI) *m/z*: $(M+H)^+$ calc. for $C_{48}H_{50}FN_9O_6Cl$: 902.3551; found: 902.3548.

**$^{1}$H NMR** (400 MHz, $CDCl_3$) δ 8.01 (s, 1H), 7.96 (d, *J* = 9.6 Hz, 1H), 7.86 (d, *J* = 8.2 Hz, 1H), 7.55 (d, *J* = 8.7 Hz, 1H), 7.45 (d, *J* = 11.0 Hz, 1H), 7.38 – 7.28 (m, 5H), 7.25 – 7.20 (m, 1H), 7.00 (d, *J* = 2.4 Hz, 1H), 6.96 (d, *J* = 9.6 Hz, 1H), 6.85 (dd, *J* = 8.7, 2.4 Hz, 1H), 4.92 (dd, *J* = 12.3, 5.3 Hz, 1H), 4.56 (d, *J* = 13.2 Hz, 2H), 4.36 – 4.26 (m, 1H), 4.13 – 3.98 (m, 1H), 3.80 (s, 2H), 3.66 (d, *J* = 12.0 Hz, 2H), 3.07 – 2.94 (m, 3H), 2.94 – 2.68 (m, 6H), 2.26 – 2.10 (m, 5H), 1.97 – 1.79 (m, 6H), 1.74 – 1.62 (m, 4H), 1.52 – 1.39 (m, 2H) ppm.

**$^{19}$F{$^{1}$H} NMR** (377 MHz, $CDCl_3$) δ –111.0 ppm.

**$^{13}$C NMR** (101 MHz, $CDCl_3$) δ 170.8, 168.1, 167.2, 166.5 (d, *J* = 2.5 Hz), 162.9, 161.8, 160.0, 158.2 (d, *J* = 255.7 Hz), 146.2 (d, *J* = 9.0 Hz), 144.2, 141.8, 138.5, 135.2, 129.1 (d, *J* = 2.0 Hz), 128.4, 127.7, 127.0, 126.9, 123.8 (d, *J* = 9.1 Hz), 117.0, 116.6, 114.8, 114.0 (d, *J* = 5.1 Hz), 112.3 (d,

$J$ = 25.2 Hz), 112.2, 104.9, 75.8, 56.2, 55.8, 50.7, 50.6, 50.2, 49.5, 47.2, 45.2, 31.6, 30.8, 30.5, 30.2, 29.8, 22.8 ppm.

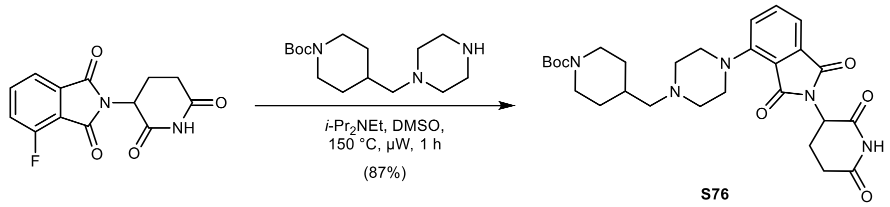

***tert*-Butyl 4-((4-(2-(2,6-dioxopiperidin-3-yl)-1,3-dioxoisoindolin-4-yl)piperazin-1-yl)methyl) piperidine-1-carboxylate (S76):** To a solution of 2-(2,6-dioxopiperidin-3-yl)-4-fluoroisoindoline-1,3-dione (250 mg, 0.91 mmol, 1.00 equiv.) in anhydrous DMSO (4.5 mL) was added *tert*-butyl 4-(piperazin-1-ylmethyl)piperidine-1-carboxylate (282 mg, 0.99 mmol, 1.10 equiv.) and *i*-$Pr_2NEt$ (480 µL, 2.76 mmol, 3.04 equiv.) at rt under argon. The reaction was stirred at 150 °C for 1 h under microwave irradiation. The mixture was diluted with water (200 mL) and extracted with EtOAc (3 × 50 mL). The combined organic layers were washed with water (50 mL) and brine (50 mL), dried with $MgSO_4$, filtered, and concentrated *in vacuo*. Purification by column chromatography (80 g silica, 0 to 10% MeOH in $CH_2Cl_2$ in 15 min) yielded 426 mg (0.79 mmol, 87%) of **S76** as a yellow solid.

**TLC** $R_f$ = 0.51 (10% MeOH in $CH_2Cl_2$).

**LC/MS** (ESI) *m/z*: $(M+H)^+$ 540.2.

**$^1$H NMR** (400 MHz, DMSO-$d_6$) δ 11.08 (s, 1H), 7.70 (dd, $J$ = 8.4, 7.2 Hz, 1H), 7.35 (dd, $J$ = 7.2, 0.7 Hz, 1H), 7.33 (dd, $J$ = 8.5, 0.8 Hz, 1H), 5.09 (dd, $J$ = 12.8, 5.4 Hz, 1H), 4.04 – 3.84 (m, 2H), 3.31 – 3.22 (m, 4H), 2.87 (ddd, $J$ = 17.4, 14.1, 5.5 Hz, 1H), 2.79 – 2.52 (m, 9H), 2.22 – 2.16 (m, 2H), 2.06 – 1.96 (m, 1H), 1.80 – 1.64 (m, 2H), 1.39 (s, 9H), 1.03 – 0.88 (m, 2H) ppm.

**$^{13}$C NMR** (101 MHz, DMSO-$d_6$) δ 172.8, 170.0, 167.0, 166.3, 153.9, 149.7, 135.9, 133.6, 123.7, 116.5, 114.8, 78.4, 63.7, 53.0, 50.5, 48.8, 43.4, 32.6, 30.9, 30.3, 28.1, 22.0 ppm.

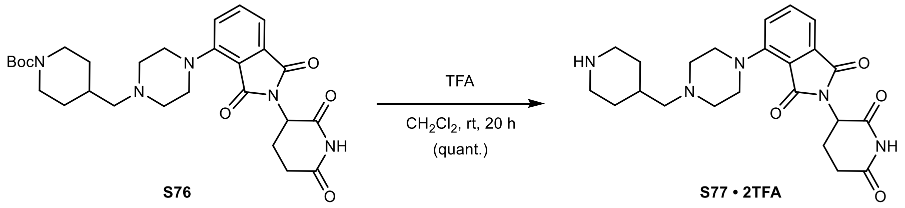

**2-(2,6-Dioxopiperidin-3-yl)-4-(4-(piperidin-4-ylmethyl)piperazin-1-yl)isoindoline-1,3-dione (S77 · 2TFA):** To a solution of **S76** (424 mg, 0.79 mmol, 1.00 equiv.) in $CH_2Cl_2$ (12 mL) was added trifluoroacetic acid (0.6 mL, 7.84 mmol, 9.9 equiv.) slowly at 0 °C. After 5 min, the reaction was allowed to warm to rt and stirred for 20 h. The mixture was concentrated *in vacuo* – residual trifluoroacetic acid was removed by co-evaporation with MeOH (2 × 5 mL), toluene (2 × 5 mL), and drying *in vacuo*, yielding 525 mg (0.79 mmol, quantitative) of **S77 · 2TFA** as a yellow solid.

**TLC** $R_f$ = 0.08 (50% MeOH + 0.5% $NH_4OH$ in $CH_2Cl_2$).

**LC/MS** (ESI) *m/z*: $(M+H)^+$ 440.2.

**$^1$H NMR** (400 MHz, DMSO-$d_6$) δ 11.10 (s, 1H), 9.86 (bs, 1H), 8.65 (d, $J$ = 8.8 Hz, 1H), 8.41 (d, $J$ = 8.4 Hz, 1H), 7.77 (dd, $J$ = 8.4, 7.2 Hz, 1H), 7.45 (dd, $J$ = 18.3, 7.8 Hz, 2H), 5.11 (dd, $J$ = 12.7, 5.5 Hz, 1H), 3.94 – 3.58 (m, 6H), 3.39 – 3.08 (m, 8H), 2.94 – 2.82 (m, 3H), 2.65 – 2.52 (m, 2H), 2.26 – 2.10 (m, 1H), 2.09 – 1.98 (m, 1H), 1.96 – 1.83 (m, 2H), 1.47 – 1.29 (m, 2H) ppm.

**$^{13}$C NMR** (101 MHz, DMSO-$d_6$) δ 172.8, 169.9, 166.9, 166.5, 148.1, 136.1, 133.5, 124.0, 117.4, 116.0, 59.9, 51.5, 48.9, 47.3, 42.6, 30.9, 28.3, 26.2, 22.1 ppm.

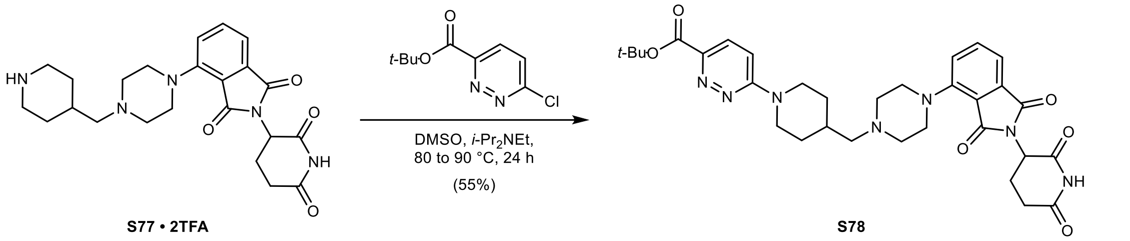

***tert*-Butyl 6-(4-((4-(2-(2,6-dioxopiperidin-3-yl)-1,3-dioxoisoindolin-4-yl)piperazin-1-yl) methyl)piperidin-1-yl)pyridazine-3-carboxylate (S78):** To a solution of **S77 · 2TFA** (117 mg, 0.18 mmol, 1.00 equiv.) in anhydrous DMSO (1.0 mL) was added *tert*-butyl 6-chloropyridazine-3-carboxylate (60 mg, 0.26 mmol, 1.50 equiv.) and *i*-$Pr_2NEt$ (137 µL, 0.79 mmol, 4.50 equiv.) at rt under argon. The reaction was stirred at 80 °C for 18 h, then at 90 °C for 6 h. The mixture was diluted with water (150 mL) and extracted with 10% MeOH in $CH_2Cl_2$ (4 × 50 mL). The combined organic layers were washed with water (50 mL) and brine (50 mL), dried with $MgSO_4$, filtered, and concentrated *in vacuo*. Purification by column chromatography (24 g silica, 0 to 7.5% MeOH with 2.5% $NH_4OH$ in $CH_2Cl_2$ in 15 min) yielded 59 mg (0.10 mmol, 55%) of **S78** as a yellow solid.

**TLC** $R_f$ = 0.28 (5% MeOH with 0.5% $NH_4OH$ in $CH_2Cl_2$).

**LC/MS** (ESI) *m/z*: $(M+H)^+$ 618.3.

**$^1$H NMR** (400 MHz, DMSO-$d_6$) δ 11.09 (s, 1H), 7.77 – 7.66 (m, 2H), 7.39 – 7.31 (m, 2H), 7.25 (d, *J* = 9.7 Hz, 1H), 5.09 (dd, *J* = 12.9, 5.4 Hz, 1H), 4.50 (d, *J* = 13.1 Hz, 2H), 3.32 – 3.25 (m, 4H), 3.03 (t, *J* = 12.5 Hz, 2H), 2.88 (ddd, *J* = 17.2, 13.9, 5.3 Hz, 1H), 2.65 – 2.52 (m, 6H), 2.22 (d, *J* = 7.1 Hz, 2H), 2.09 – 1.99 (m, 1H), 1.99 – 1.89 (m, 1H), 1.85 (d, *J* = 13.1 Hz, 2H), 1.55 (s, 9H), 1.12 (q, *J* = 11.8 Hz, 2H) ppm.

**$^{13}$C NMR** (101 MHz, DMSO-$d_6$) δ 172.8, 170.0, 167.0, 166.3, 163.1, 159.5, 149.7, 143.0, 135.9, 133.7, 128.2, 123.7, 116.5, 114.8, 111.1, 81.0, 63.6, 53.0, 50.5, 48.8, 44.4, 32.7, 30.9, 29.8, 27.8, 22.0 ppm.

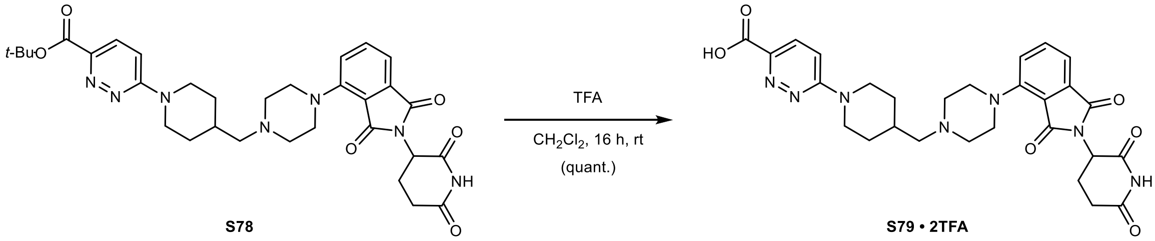

**6-(4-((4-(2-(2,6-Dioxopiperidin-3-yl)-1,3-dioxoisoindolin-4-yl)piperazin-1-yl)methyl)piperi-din-1-yl)pyridazine-3-carboxylic acid (S79 · 2TFA):** To a solution of **S78** (59 mg, 0.10 mmol, 1.00 equiv.) in $CH_2Cl_2$ (1.0 mL) was added trifluoroacetic acid (0.25 mL, 3.24 mmol, 60.0 equiv.) slowly at 0 °C. After 5 min, the reaction was allowed to warm to rt and stirred for 16 h. The mixture was concentrated *in vacuo* – residual trifluoroacetic acid was removed by co-evaporation with MeOH (4 × 5 mL), *n*-heptane (2 × 5 mL), and drying *in vacuo*, yielding 75 mg (0.10 mmol, quantitative) of **S79 · 2TFA** as a yellow solid.

**TLC** $R_f$ = 0.05, streaky (30% MeOH in $CH_2Cl_2$).

**LC/MS** (ESI) *m/z*: $(M+H)^+$ 562.2.

**$^{1}$H NMR** (400 MHz, DMSO-$d_6$) δ 11.10 (s, 1H), 9.51 (bs, 1H), 7.84 (d, *J* = 9.6 Hz, 1H), 7.78 (t, *J* = 7.8 Hz, 1H), 7.48 (d, *J* = 7.2 Hz, 1H), 7.43 (d, *J* = 8.4 Hz, 1H), 7.35 (d, *J* = 9.7 Hz, 1H), 5.12 (dd, *J* = 12.6, 5.5 Hz, 1H), 4.65 – 4.48 (m, 2H), 3.95 – 3.81 (m, 2H), 3.81 – 3.62 (m, 2H), 3.42 – 3.21 (m, 4H), 3.21 – 3.14 (m, 2H), 3.14 – 3.01 (m, 2H), 2.89 (ddd, *J* = 17.6, 14.3, 5.4 Hz, 1H), 2.71 – 2.53 (m, 2H), 2.36 – 2.18 (m, 1H), 2.10 – 1.98 (m, 1H), 1.96 – 1.82 (m, 2H), 1.31 – 1.17 (m, 2H) ppm.

**$^{13}$C NMR** (101 MHz, DMSO-$d_6$) δ 172.8, 169.9, 166.9, 166.5, 165.3, 159.5, 148.0, 142.6, 136.1, 133.5, 128.5, 124.0, 117.4, 116.0, 111.7, 60.6, 51.6, 48.8, 47.3, 47.1, 43.8, 30.9, 30.4, 28.8, 22.0 ppm.

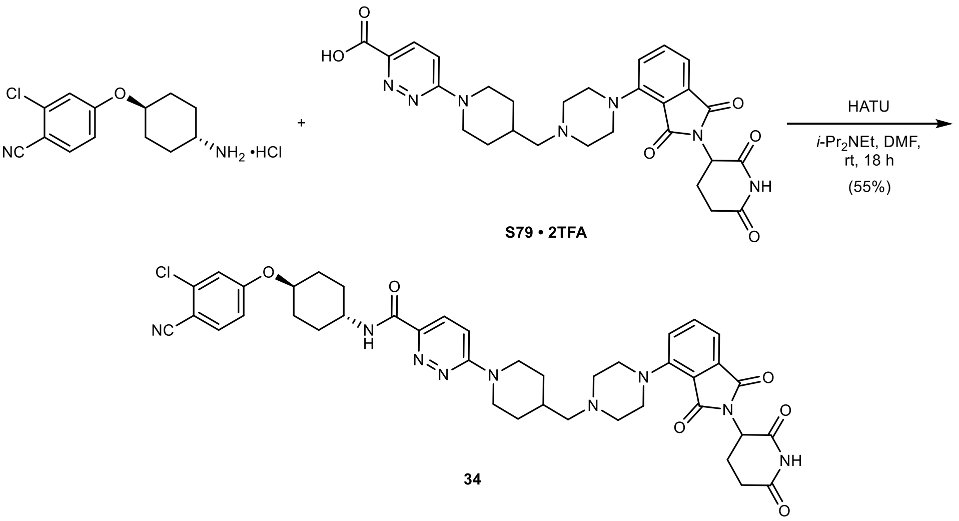

***N*-((1*r*,4*r*)-4-(3-Chloro-4-cyanophenoxy)cyclohexyl)-6-(4-((4-(2-(2,6-dioxopiperidin-3-yl)-1,3-dioxoisoindolin-4-yl)piperazin-1-yl)methyl)piperidin-1-yl)pyridazine-3-carboxamide (34):** To a solution of **S79 · 2TFA** (75 mg, 0.09 mmol, 1.00 equiv.) in anhydrous DMF (1.0 mL) was added HATU (42 mg, 0.11 mmol, 1.20 equiv.), 4-(((1*r*,4*r*)-4-aminocyclohexyl) oxy)-2-chlorobenzo-nitrile hydrochloride (31 mg, 0.11 mmol, 1.20 equiv.), and *i*-Pr$_2$NEt (71 µL, 0.41 mmol, 4.50 equiv.) at rt under argon. The reaction was stirred at rt for 18 h. The mixture was poured into water (150 mL) and extracted with 10% MeOH in $CH_2Cl_2$ (3 × 50 mL). The combined organic layers were washed with 5% aqueous LiCl (2 × 50 mL), water (50 mL), and brine (50 mL), dried with $MgSO_4$, filtered, and concentrated *in vacuo*. Purification by column chromatography (12 g silica, 0 to 9% MeOH with 2.5% $NH_4OH$ in $CH_2Cl_2$ in 13 min) yielded 40 mg (0.05 mmol, 55%) of **34** (**DKFZ-1270**) as a yellow powder.

**TLC** $R_f$ = 0.45 (10% MeOH with 0.5% $NH_4OH$ in $CH_2Cl_2$).

**HRMS** (ESI) *m/z*: $(M+H)^+$ calc. for $C_{41}H_{45}N_9O_6Cl$: 794.3176; found: 794.3178.

**$^{1}$H NMR** (400 MHz, DMSO-$d_6$) δ 11.09 (s, 1H), 8.58 (d, *J* = 8.3 Hz, 1H), 7.85 (d, *J* = 8.8 Hz, 1H), 7.80 (d, *J* = 9.6 Hz, 1H), 7.71 (dd, *J* = 8.4, 7.2 Hz, 1H), 7.42 – 7.30 (m, 4H), 7.13 (dd, *J* = 8.8, 2.4 Hz, 1H), 5.09 (dd, *J* = 12.8, 5.4 Hz, 1H), 4.61 – 4.41 (m, 3H), 3.91 – 3.79 (m, 1H), 3.31 – 3.22 (m, 4H), 3.02 (t, *J* = 12.4 Hz, 2H), 2.88 (ddd, *J* = 17.3, 14.0, 5.5 Hz, 1H), 2.58 (d, *J* = 23.6 Hz, 6H), 2.22 (d, *J* = 7.1 Hz, 2H), 2.16 – 2.06 (m, 2H), 2.06 – 1.99 (m, 1H), 1.99 – 1.80 (m, 5H), 1.64 (q, *J* = 12.2 Hz, 2H), 1.51 (q, *J* = 10.2 Hz, 2H), 1.13 (q, *J* = 11.9 Hz, 2H) ppm.

**$^{13}$C NMR** (101 MHz, DMSO-$d_6$) δ 172.8, 170.0, 167.0, 166.3, 162.5, 161.8, 160.0, 149.7, 144.2, 137.0, 135.9, 135.7, 133.7, 126.2, 123.7, 116.8, 116.5, 116.4, 115.5, 114.8, 112.4, 103.2, 75.6, 63.7, 53.0, 50.6, 48.8, 47.0, 44.5, 32.7, 30.9, 29.9, 29.8, 29.4, 22.0 ppm.

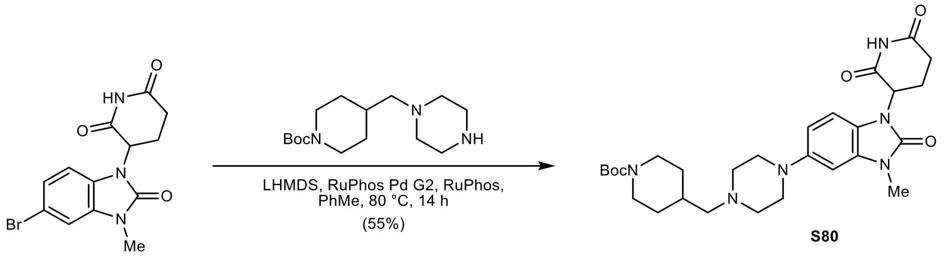

**S80**

***tert*-Butyl 4-((4-(1-(2,6-dioxopiperidin-3-yl)-3-methyl-2-oxo-2,3-dihydro-1*H*-benzo[*d*]imidazol-5-yl)piperazin-1-yl)methyl)piperidine-1-carboxylate (S80):** To a solution of 3-(5-bromo-3-methyl-2-oxo-2,3-dihydro-1*H*-benzo[*d*]imidazol-1-yl)piperidine-2,6-dione (150 mg, 0.44 mmol, 1.00 equiv.) and *tert*-butyl 4-(piperazin-1-ylmethyl)piperidine-1-carboxylate (277 mg, 0.98 mmol, 2.20 equiv.) in degassed, anhydrous toluene (1.5 mL) was added LHMDS (4.0 mL, 1.0 M in THF, 3.99 mmol, 9.00 equiv.) dropwise at 0 °C under argon. The mixture was degassed again, then RuPhos (62 mg, 0.13 mmol, 0.30 equiv.) and RuPhos Pd G2 (52 mg, 0.07 mmol, 0.15 equiv.) were added. The reaction was heated to 80 °C and stirred for 14 h. The mixture was filtered through Celite on a sintered glass funnel, rinsed with EtOAc (50 mL), washed with saturated aqueous $NH_4Cl$ solution (2 × 50 mL) and brine (2 × 25 mL), dried with $MgSO_4$, filtered, and concentrated *in vacuo*. Purification by column chromatography (12 g silica, 0 to 9% MeOH in $CH_2Cl_2$ in 12 min) yielded 136 mg (0.24 mmol, 55%) of **S80** as an off-white solid.

**TLC** $R_f$ = 0.63 (10% MeOH in $CH_2Cl_2$).

**LC/MS** (ESI) *m/z*: $(M+H)^+$ 541.3.

**$^1$H NMR** (400 MHz, $CDCl_3$) δ 8.43 (s, 1H), 6.72 – 6.61 (m, 3H), 5.18 (dd, *J* = 12.7, 5.4 Hz, 1H), 4.26 – 3.91 (m, 2H), 3.39 (s, 3H), 3.17 – 3.08 (m, 4H), 2.98 – 2.87 (m, 1H), 2.87 – 2.62 (m, 4H), 2.30 – 2.15 (m, 3H), 1.81 – 1.62 (m, 3H), 1.45 (s, 9H), 1.09 (qd, *J* = 12.4, 4.4 Hz, 2H), 2.62 – 2.50 (m, 4H) ppm.

**$^{13}$C NMR** (101 MHz, $CDCl_3$) δ 171.2, 168.5, 155.1, 154.4, 148.1, 131.2, 121.5, 110.7, 108.8, 98.4, 79.4, 64.6, 53.8, 52.3, 51.0, 44.0, 33.7, 31.5, 30.9, 28.6, 27.4, 23.2 ppm.

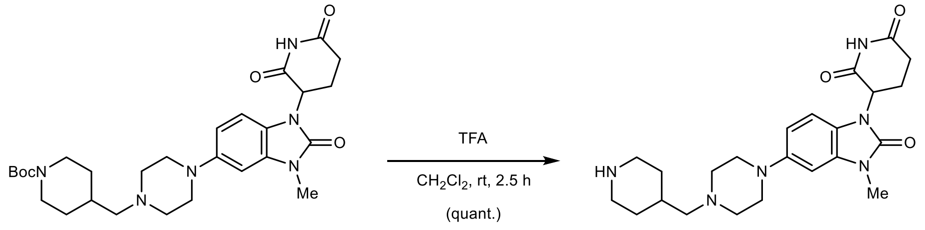

**S80** **S81 · 2TFA**

**3-(3-Methyl-2-oxo-5-(4-(piperidin-4-ylmethyl)piperazin-1-yl)-2,3-dihydro-1*H*-benzo[*d*]imidazol-1-yl)piperidine-2,6-dione (S81 · 2TFA):** To a solution of **S80** (130 mg, 0.24 mmol, 1.00 equiv.) in $CH_2Cl_2$ (2.0 mL) was added trifluoroacetic acid (1.0 mL, 13.2 mmol, 55.0 equiv.) slowly at 0 °C. After 5 min, the reaction was allowed to warm to rt and stirred for 2.5 h. The mixture was concentrated *in vacuo* – residual trifluoroacetic acid was removed by co-evaporation with MeOH (3 × 5 mL) and drying *in vacuo*, yielding 161 mg (0.24 mmol, quantitative) of **S81 · 2TFA** as a yellow solid.

**TLC** $R_f$ = 0.05, streaky (50% MeOH + 0.5% $NH_4OH$ in $CH_2Cl_2$).

**LC/MS** (ESI) *m/z*: $(M+H)^+$ 441.2.

**$^{1}$H NMR** (400 MHz, DMSO-$d_6$) δ 11.07 (s, 1H), 9.87 (bs, 1H), 8.73 (bs, 1H), 8.49 (bs, 1H), 7.00 (d, $J$ = 8.6 Hz, 1H), 6.93 (d, $J$ = 2.3 Hz, 1H), 6.70 (dd, $J$ = 8.6, 2.3 Hz, 1H), 5.31 (dd, $J$ = 12.9, 5.3 Hz, 1H), 3.87 – 3.69 (m, 2H), 3.69 – 3.56 (m, 2H), 3.39 – 3.28 (m, 5H), 3.23 – 3.11 (m, 4H), 3.11 – 2.99 (m, 2H), 2.97 – 2.81 (m, 3H), 2.76 – 2.57 (m, 2H), 2.26 – 2.08 (m, 1H), 2.08 – 1.84 (m, 3H), 1.45 – 1.31 (m, 2H) ppm.

**$^{13}$C NMR** (101 MHz, DMSO-$d_6$) δ 172.8, 170.0, 153.6, 145.2, 130.7, 122.5, 109.5, 108.9, 98.0, 59.8, 51.6, 51.3, 46.7, 42.5, 31.2, 28.1, 27.0, 26.2, 22.1 ppm.

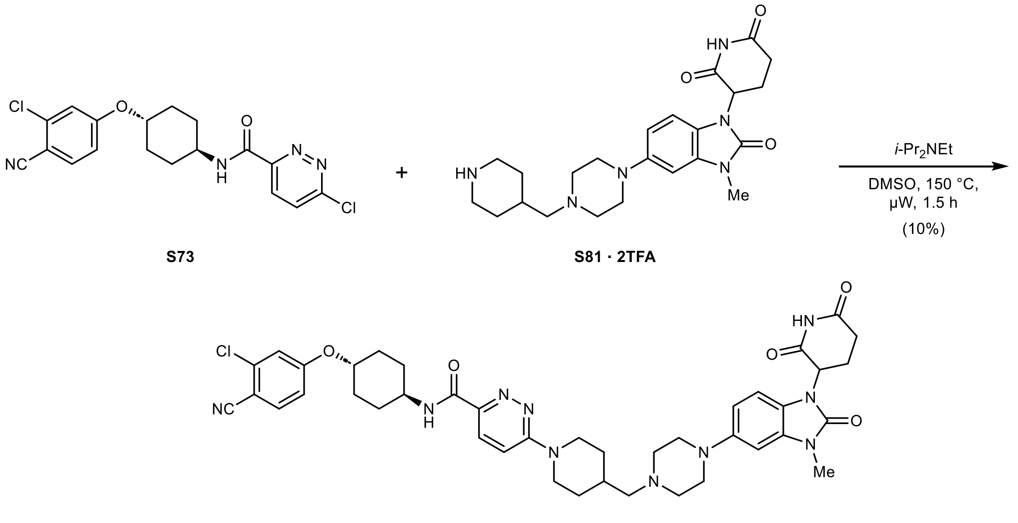

***N*-((1*r*,4*r*)-4-(3-Chloro-4-cyanophenoxy)cyclohexyl)-6-(4-((4-(1-(2,6-dioxopiperidin-3-yl)-3-methyl-2-oxo-2,3-dihydro-1*H*-benzo[*d*]imidazol-5-yl)piperazin-1-yl)methyl)piperidin-1-yl) pyridazine-3-carboxamide (35 · TFA, DKFZ-1300):** To a solution of **S73** (30 mg, 0.08 mmol, 1.00 equiv.) in anhydrous DMSO (0.75 mL) was added **S81 · 2TFA** (54 mg, 0.08 mmol, 1.05 equiv.) and *i*-$Pr_2NEt$ (53 µL, 0.31 mmol, 4.00 equiv.) at rt under argon. The reaction was stirred at 150 °C for 1.5 h under microwave irradiation. The mixture was diluted with water (100 mL) and extracted with EtOAc (3 × 25 mL). The combined organic layers were washed with water (25 mL) and brine (25 mL), dried with $MgSO_4$, filtered, and concentrated *in vacuo*. Purification by column chromatography (12 g silica, 0 to 12% MeOH in $CH_2Cl_2$ in 12 min), followed by reversed-phase column chromatography (15.5 g C18, 0 to 70% MeCN in $H_2O$ in 15 min) and preparative HPLC (1 to 60% MeCN in $H_2O$ with 0.05% TFA in 9 min) yielded 7 mg (0.01 mmol, 10%) of **35 · TFA** (**DKFZ-1300**) as a colorless solid.

**TLC** $R_f$ = 0.29 (5% MeOH in $CH_2Cl_2$).

**HRMS** (ESI) *m/z*: $(M+H)^+$ calc. for $C_{41}H_{48}N_{10}O_5Cl$: 795.3492; found: 795.3476.

**$^{1}$H NMR** (400 MHz, DMSO-$d_6$) δ 11.08 (s, 1H), 9.68 (s, 1H), 8.59 (d, $J$ = 8.2 Hz, 1H), 7.89 – 7.80 (m, 2H), 7.46 – 7.36 (m, 2H), 7.13 (dd, $J$ = 8.8, 2.5 Hz, 1H), 7.01 (d, $J$ = 8.5 Hz, 1H), 6.93 (d, $J$ = 2.2 Hz, 1H), 6.70 (dd, $J$ = 8.7, 2.2 Hz, 1H), 5.32 (dd, $J$ = 12.9, 5.3 Hz, 1H), 4.61 – 4.45 (m, 3H), 3.89 – 3.67 (m, 3H), 3.63 (d, $J$ = 11.6 Hz, 2H), 3.33 (s, 3H), 3.25 – 2.97 (m, 8H), 2.97 – 2.83 (m, 1H), 2.73 – 2.58 (m, 2H), 2.31 – 2.16 (m, 1H), 2.16 – 2.04 (m, 2H), 2.04 – 1.95 (m, 1H), 1.94 – 1.80 (m, 4H), 1.64 (q, $J$ = 12.2 Hz, 2H), 1.52 (q, $J$ = 11.8 Hz, 2H), 1.35 – 1.17 (m, 2H) ppm.

**$^{13}$C NMR** (101 MHz, DMSO-$d_6$) δ 172.8, 170.0, 162.5, 161.8, 159.9, 153.6, 145.3, 144.5, 137.0, 135.8, 130.7, 126.4, 122.4, 116.8, 116.4, 115.5, 112.7, 109.5, 108.9, 103.2, 98.0, 75.6, 60.6, 51.6, 51.4, 47.1, 46.7, 44.0, 31.2, 30.4, 29.9, 29.4, 28.8, 27.0, 22.1 ppm.

### References

1. Woehrmann, M. H. *et al.* Large-scale cytological profiling for functional analysis of bioactive compounds. *Mol. Biosyst.* **9**, 2604–2617 (2013).
2. Grigalunas, M. *et al.* Natural product fragment combination to performance-diverse pseudo-natural products. *Nat. Commun.* **12**, 1883 (2021).
3. Pahl, A. *et al.* Morphological subprofile analysis for bioactivity annotation of small molecules. *Cell Chem. Biol.* **30**, 839–853 e837 (2023).
4. Rezaei Adariani, S. *et al.* Detection of a Mitochondrial Fragmentation and Integrated Stress Response Using the Cell Painting Assay. *J. Med. Chem.* **67**, 13252–13270 (2024).
5. Adam, W., Bialas, J. & Hadjiarapoglou, L. Kurzmitteilung / Short Communication A Convenient Preparation of Acetone Solutions of Dimethyldioxirane. *Chem. Ber.* **124**, 2377–2377 (1991).
6. Marker, T. *et al.* Site-specific activation of the proton pump inhibitor rabeprazole by tetrathiolate zinc centres. *Nat. Chem. Biol.* **17**, 507–517 (2025).
7. Han, X. *et al.* Strategies toward Discovery of Potent and Orally Bioavailable Proteolysis Targeting Chimera Degraders of Androgen Receptor for the Treatment of Prostate Cancer. *J. Med. Chem.* **64**, 12831–12854 (2021).
8. Rédl, S., Klecén, O. & Havlíček, J. Synthetic studies connected with the preparation of $H^+/K^+$-ATPase inhibitors rabeprazole and lansoprazole. *J. Het. Chem.* **43**, 1447–1453 (2006).
9. Giarrusso, M. A. *et al.* Fluorescent Angiotensin AT1 Receptor Antagonists. *Asian J. Org. Chem.* **1**, 274–279 (2012).

### Spectra

NR-3-078-A Instrument 1  PDA - Total Absorbance Chromatogram

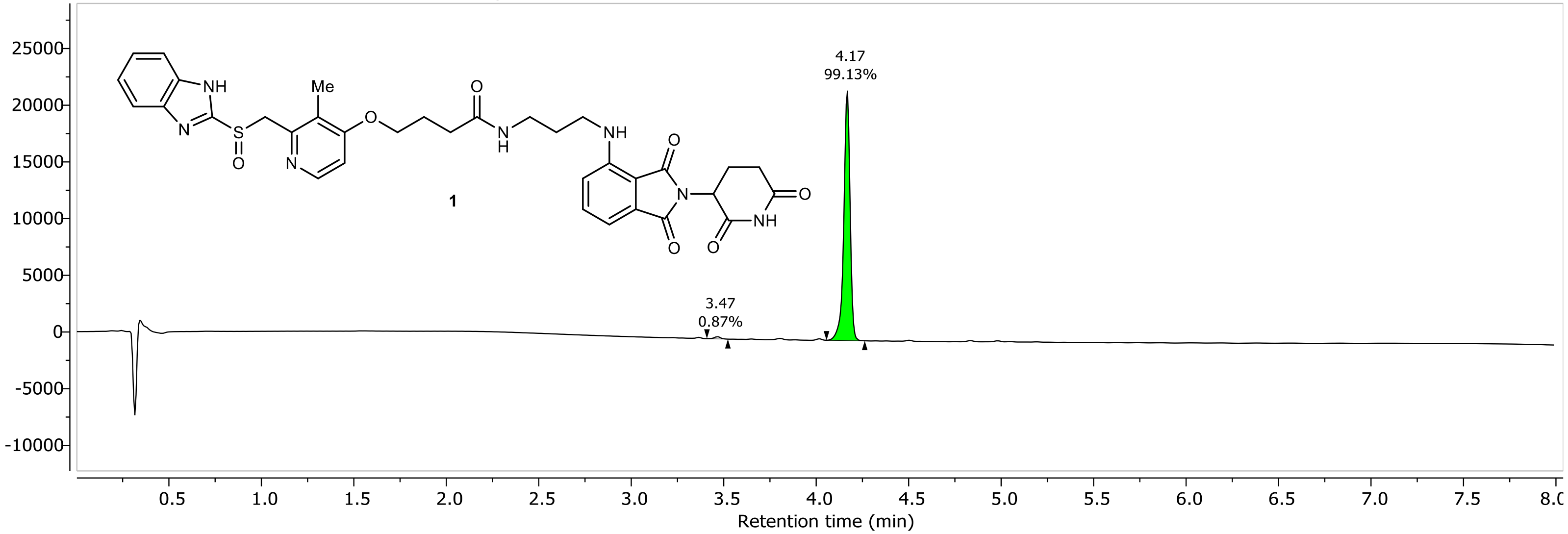

NR-3-078-A Instrument 1 DAD1A, Sig=254...Ref=360.0,100.0 Chromatogram

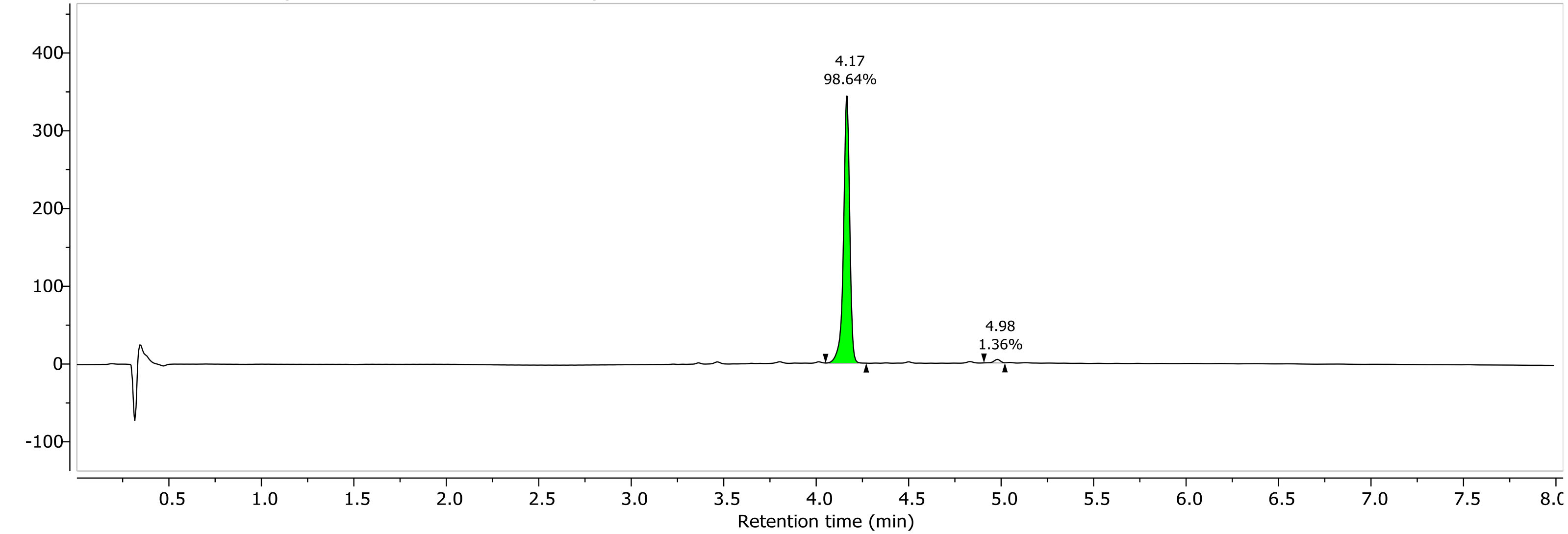

1H NMR (400.26 MHz, MeOD, 298.1 K)

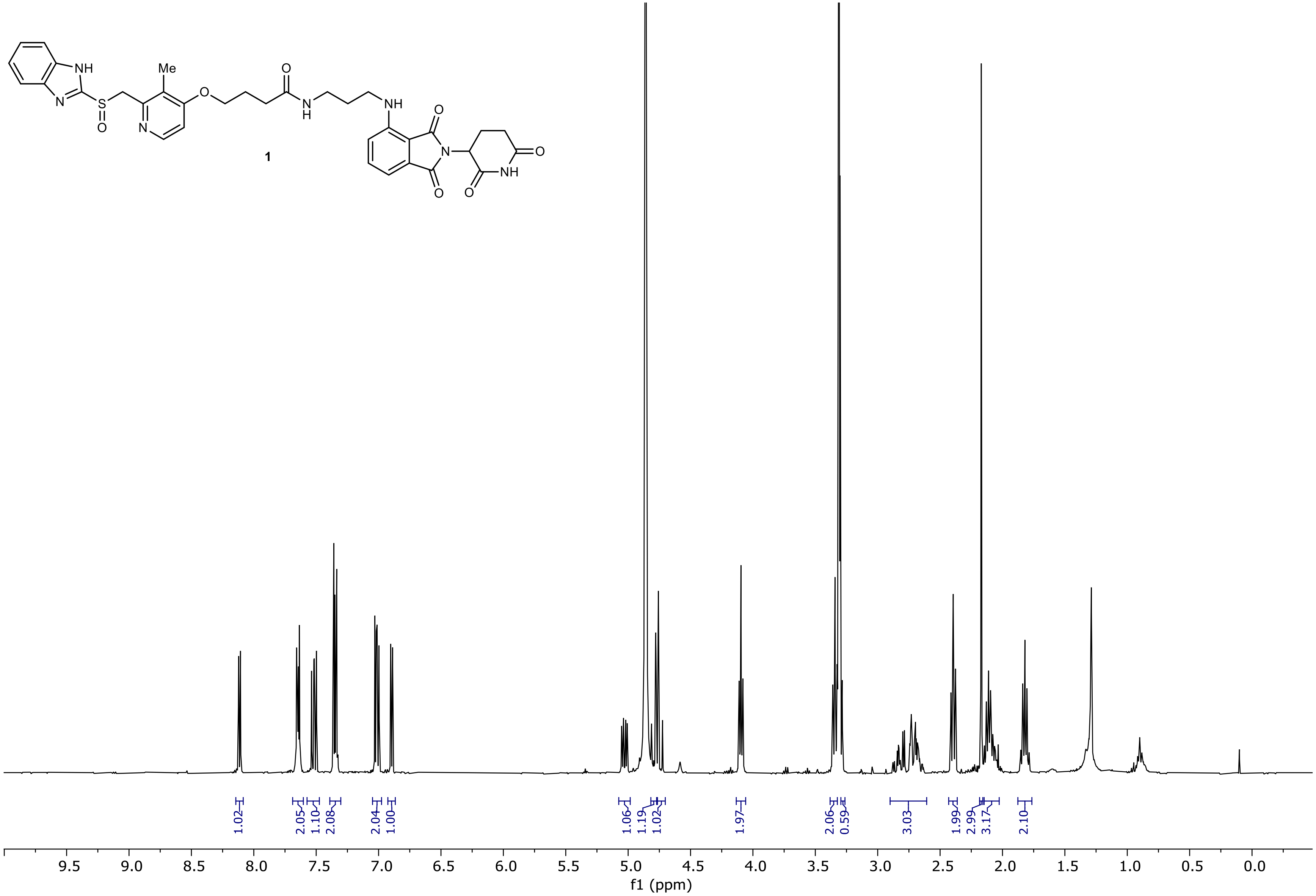

13C NMR (100.66 MHz, MeOD, 298.1 K)

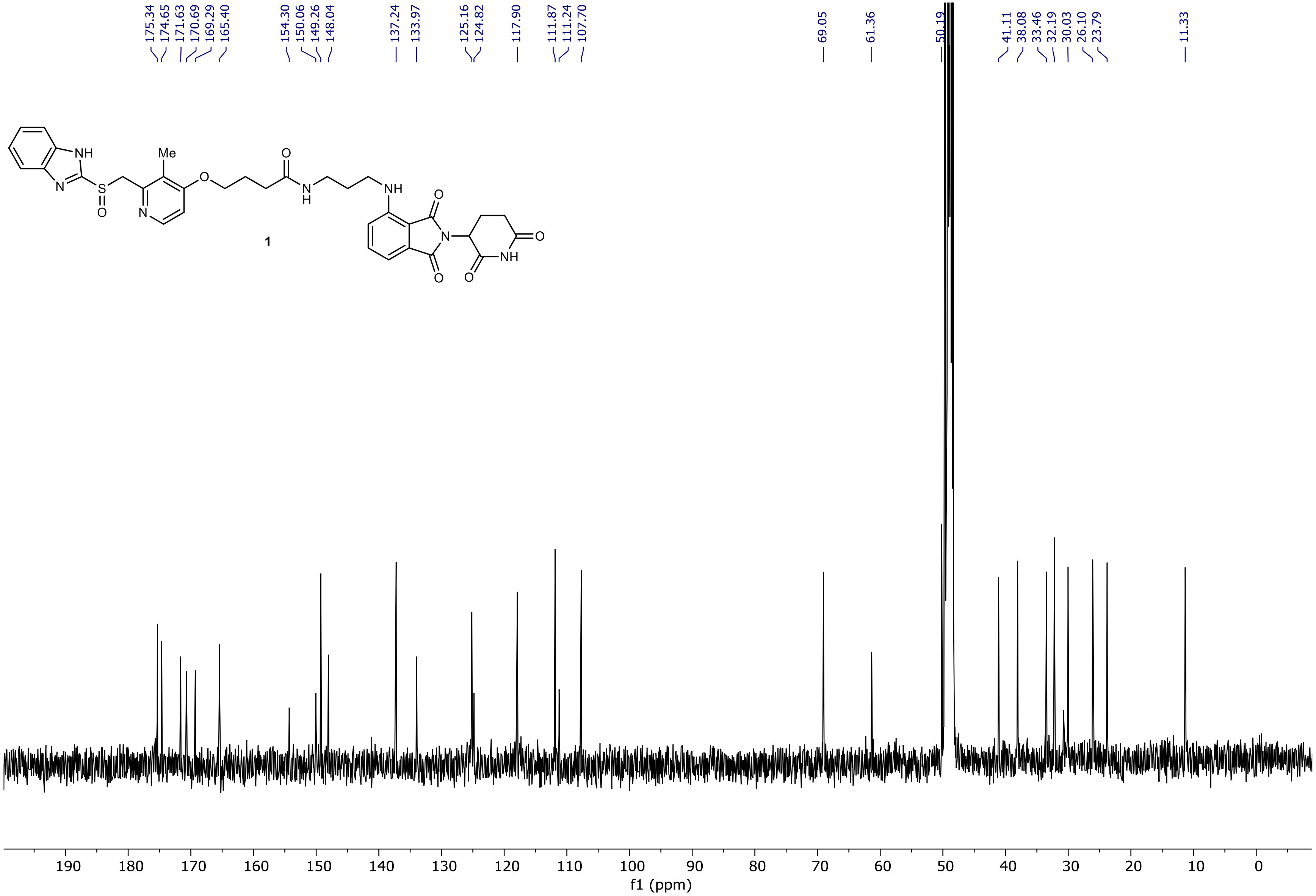

NR-3-150-A Instrument 1  PDA - Total Absorbance Chromatogram

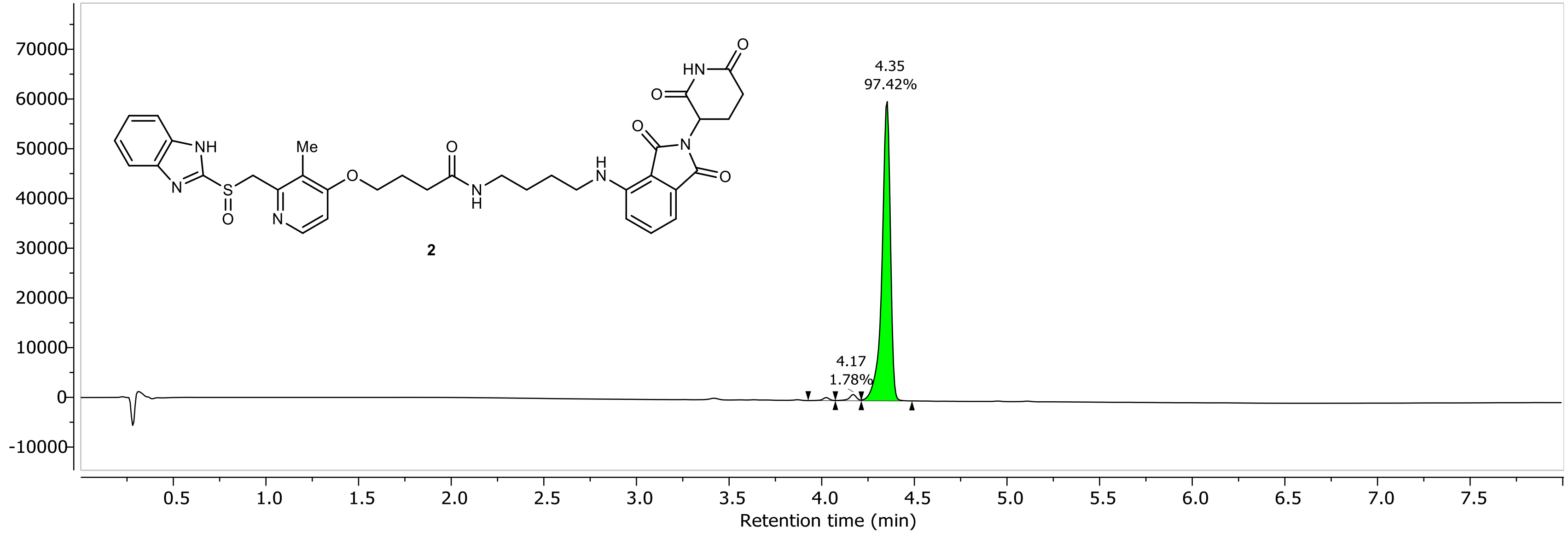

NR-3-150-A Instrument 1 DAD1A, Sig=254...Ref=360.0,100.0 Chromatogram

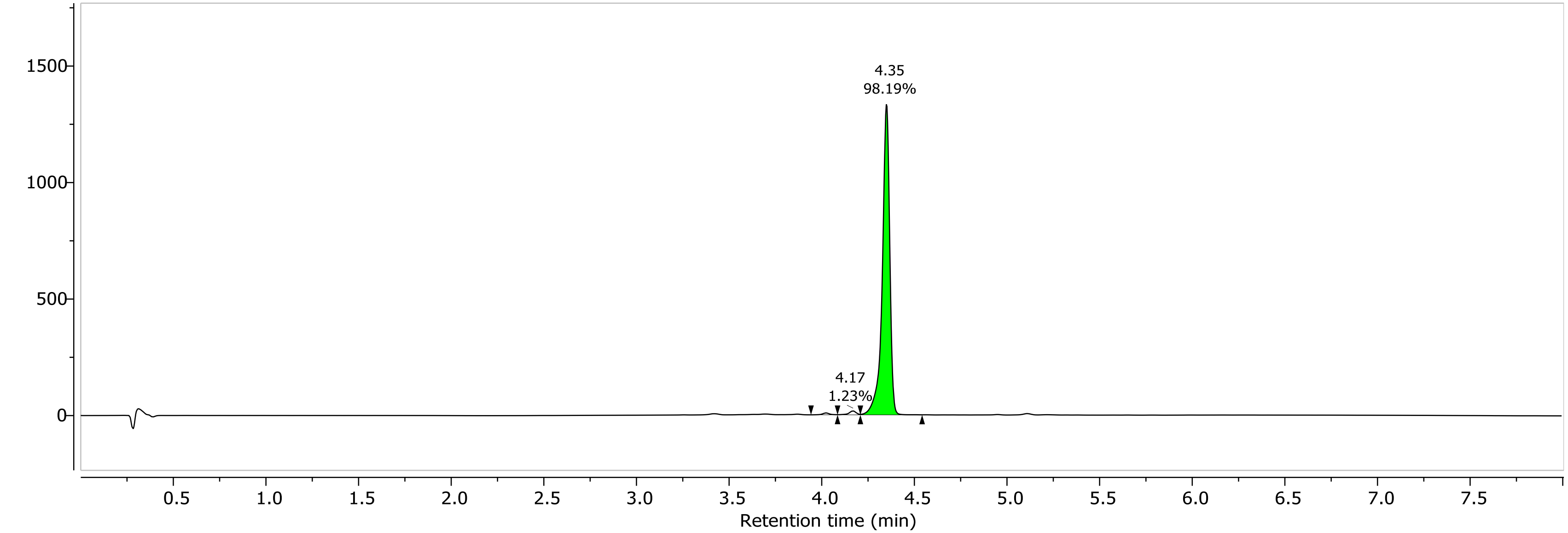

1H NMR (400.26 MHz, DMSO, 298.1 K)

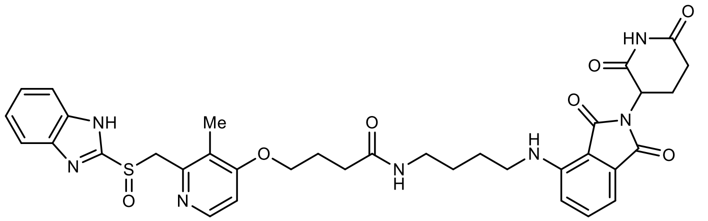

**2**

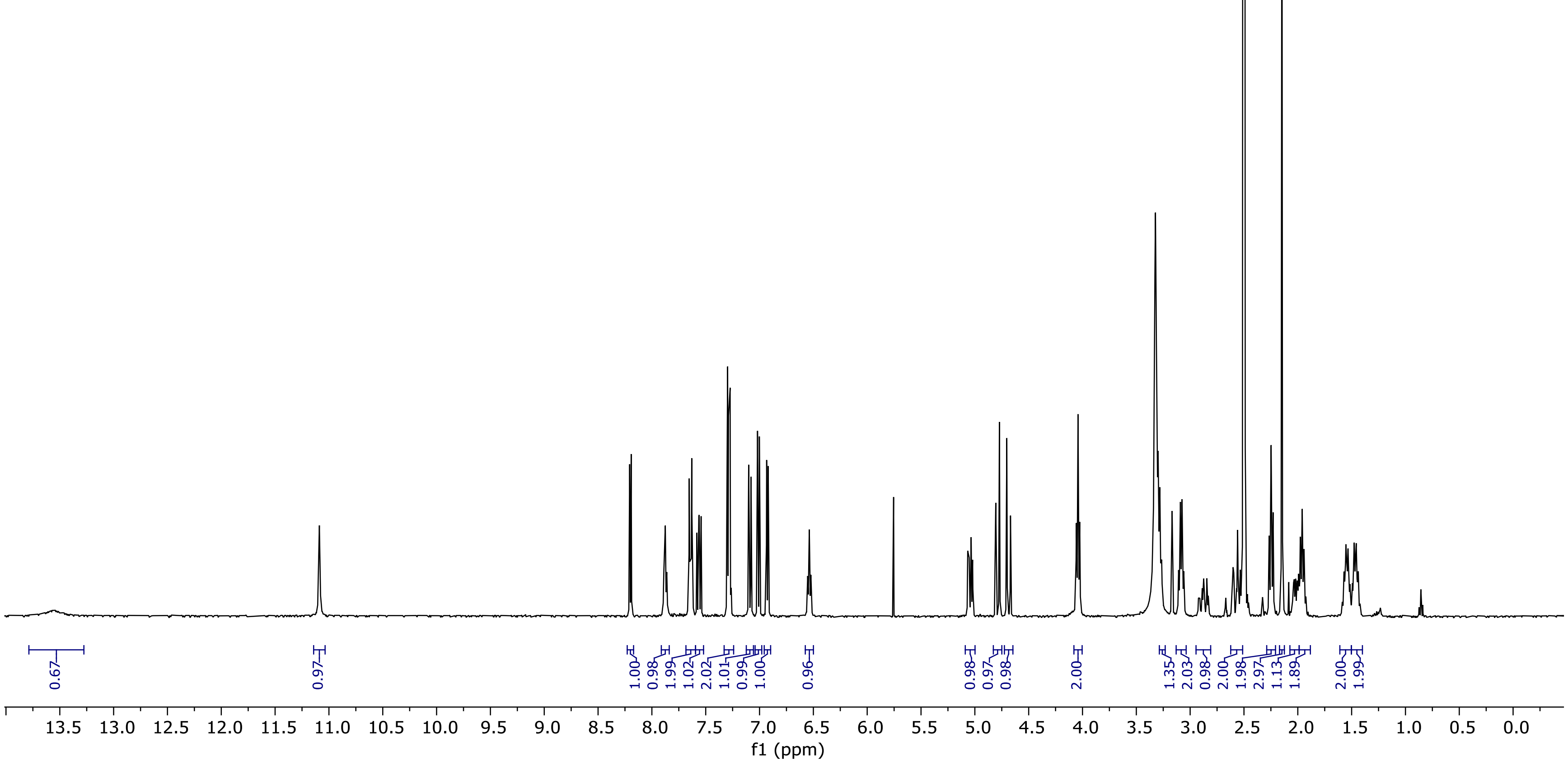

13C NMR (100.66 MHz, DMSO, 298.1 K)

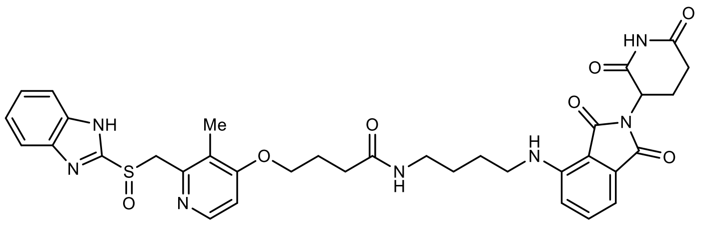

2

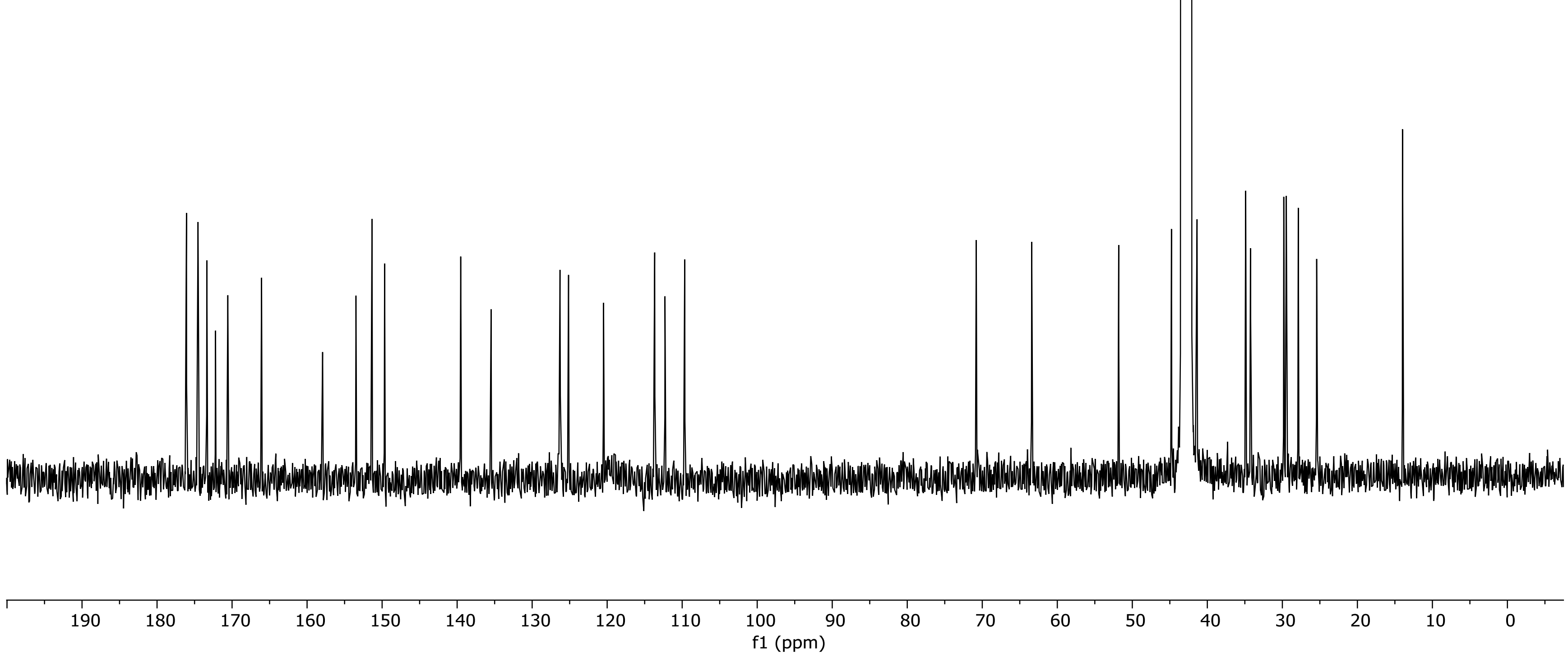

NR-4-035-A-acidic Instrument 1 PDA - Total Absorbance Chromatogram

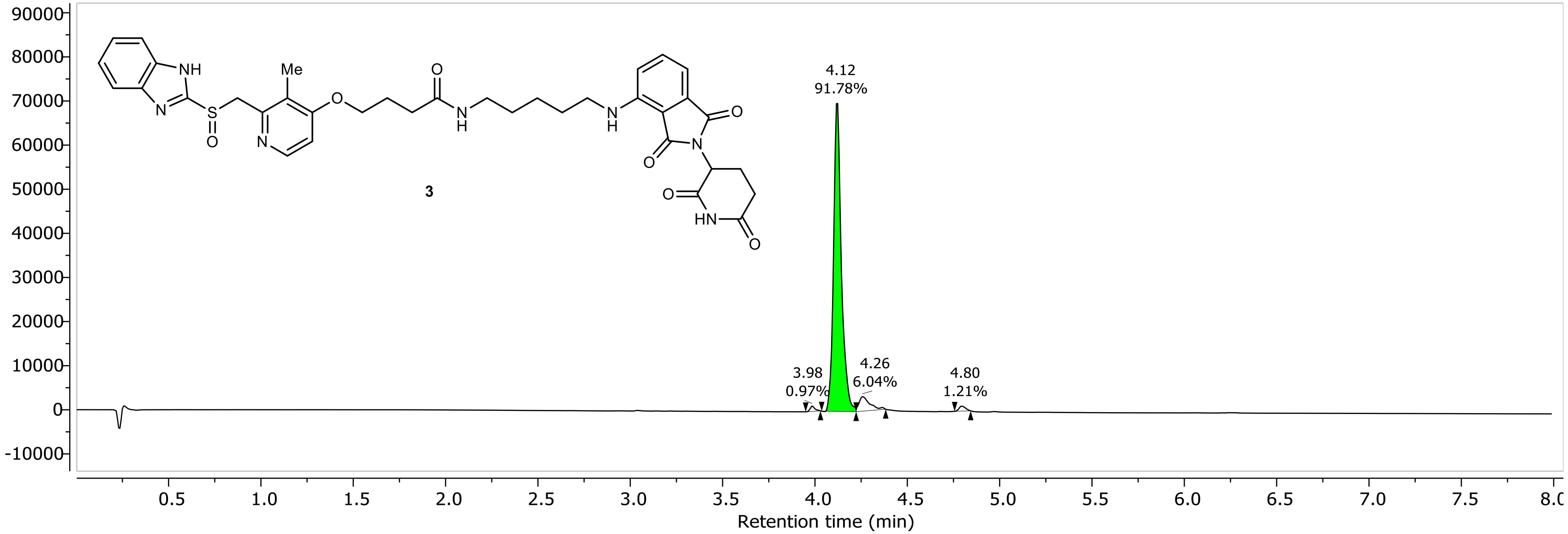

NR-4-035-A-acidic Instrument 1 DAD1A, Sig=254...Ref=360.0,100.0 Chromatogram

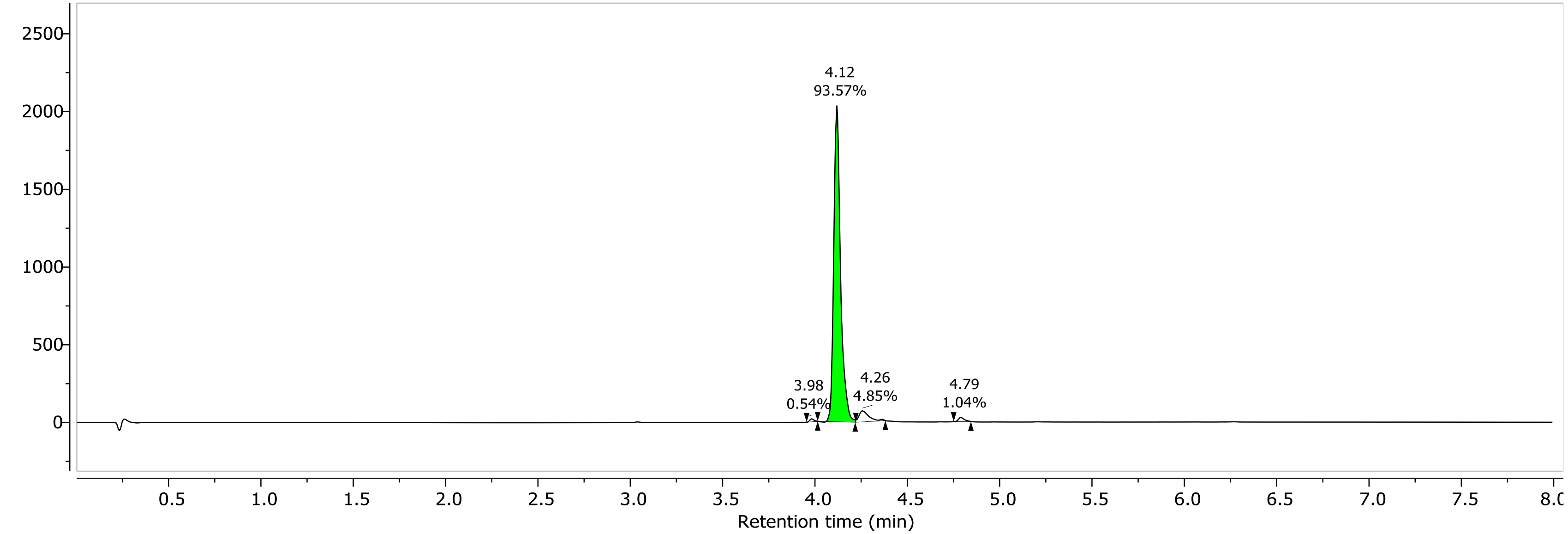

1H NMR (400.26 MHz, DMSO, 298.1 K)

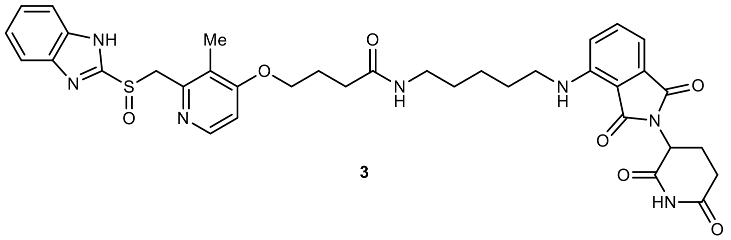

**3**

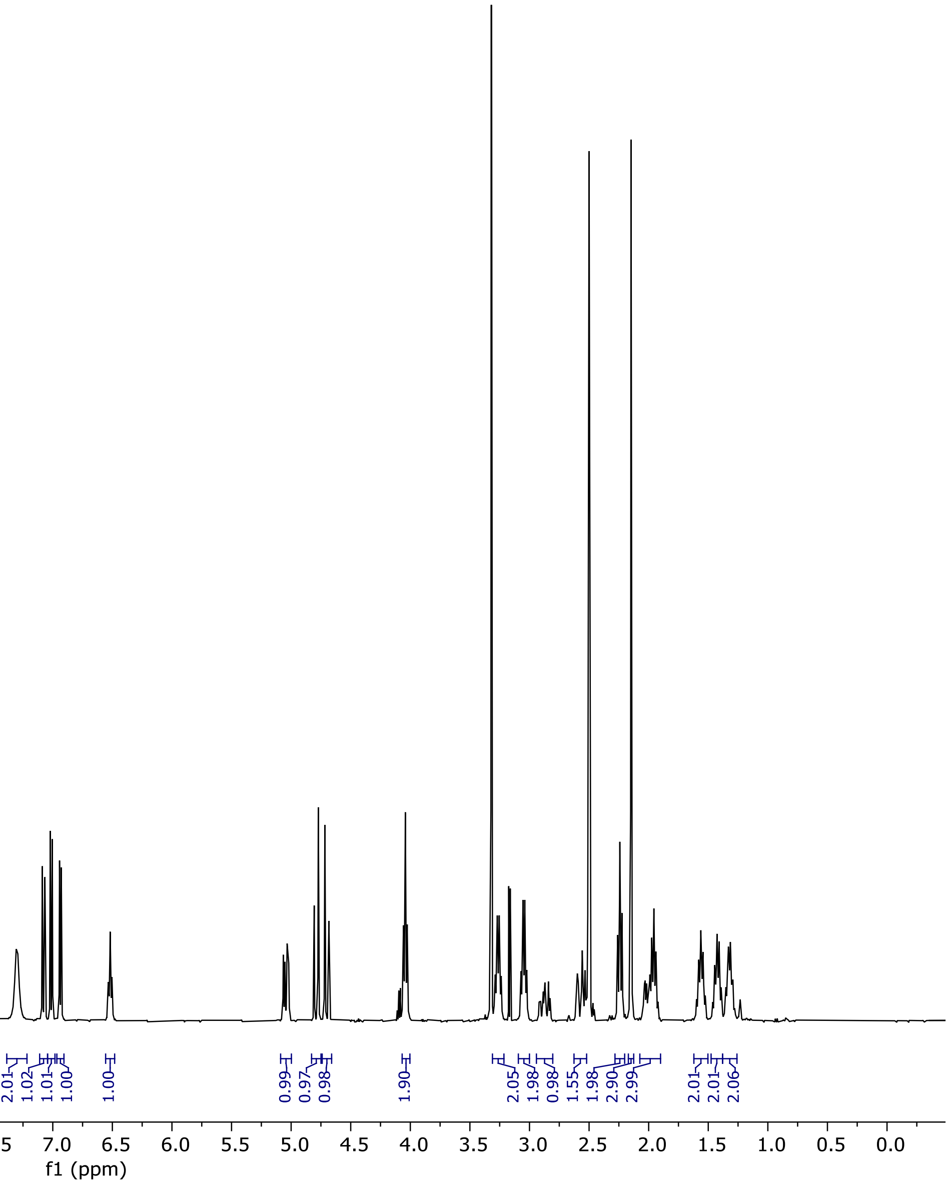

13C NMR (100.66 MHz, DMSO, 298.1 K)

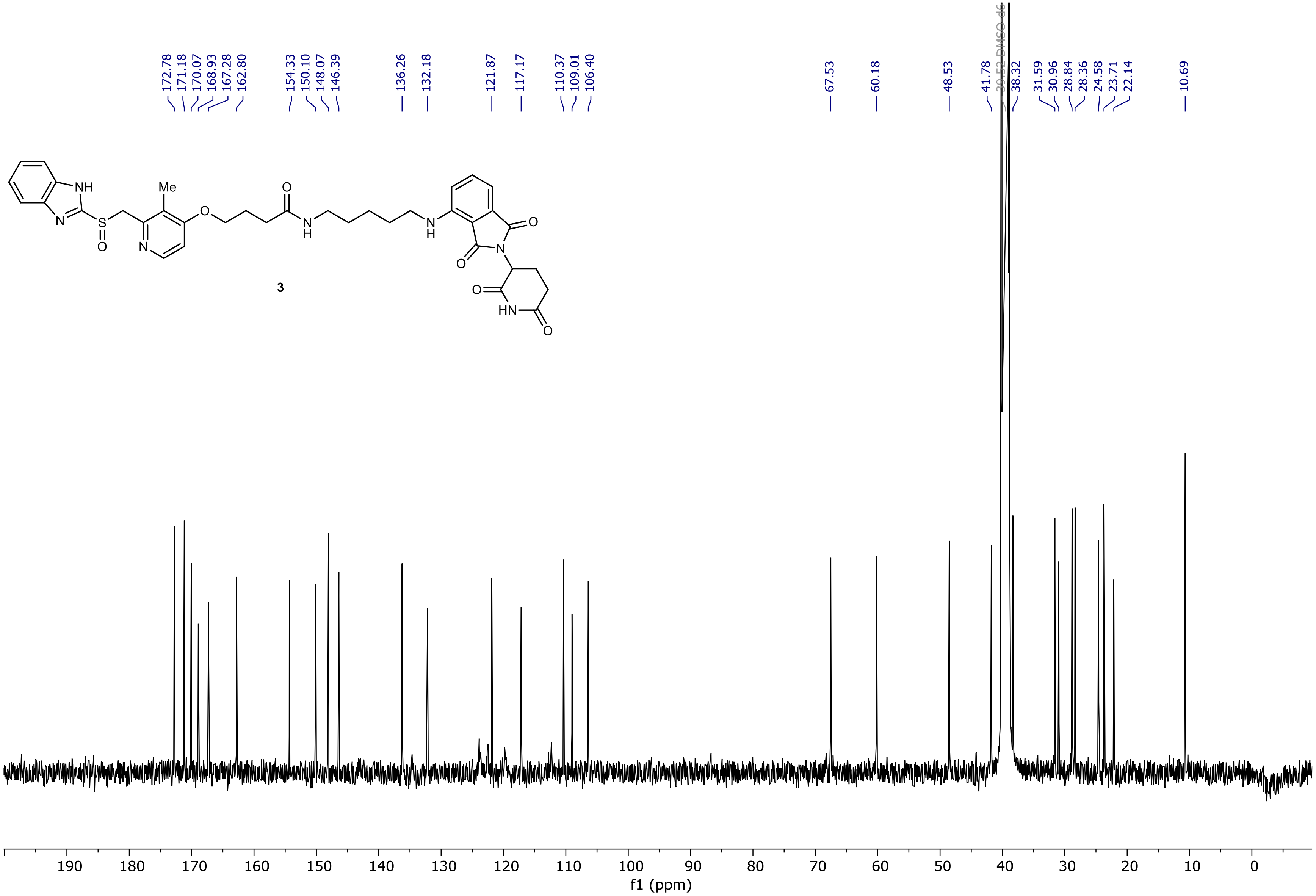

NR-4-016-B Instrument 1  PDA - Total Absorbance Chromatogram

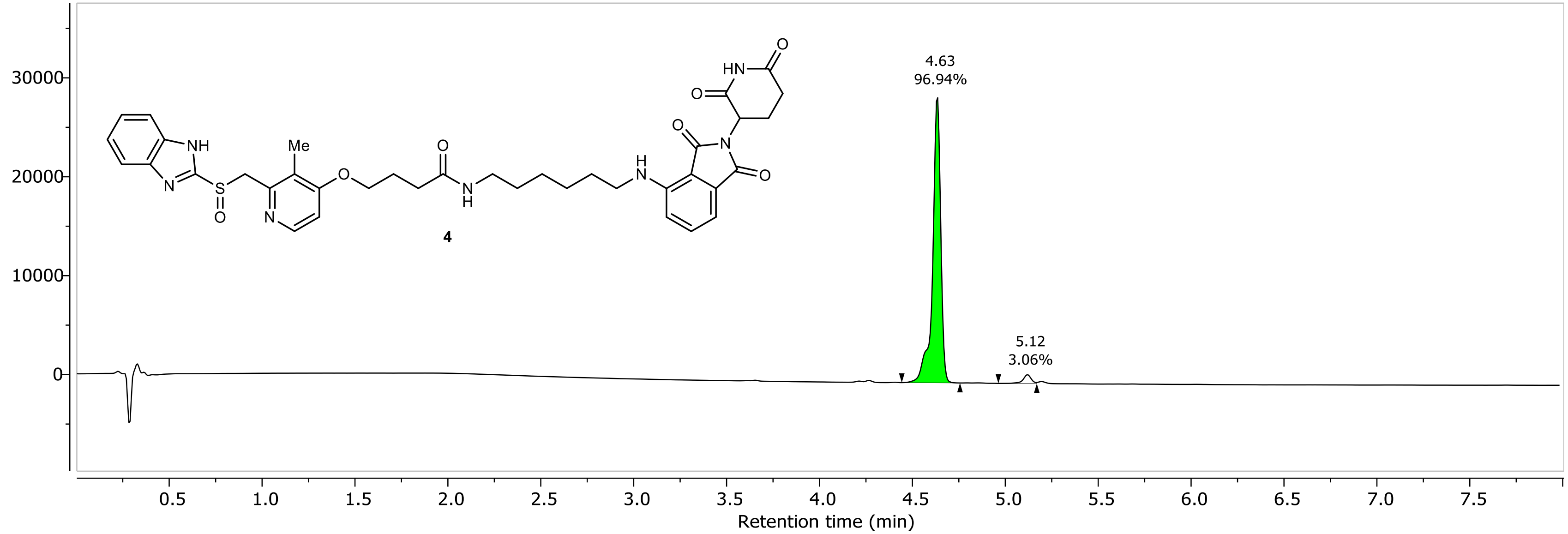

NR-4-016-B Instrument 1 DAD1A, Sig=254...Ref=360.0,100.0 Chromatogram

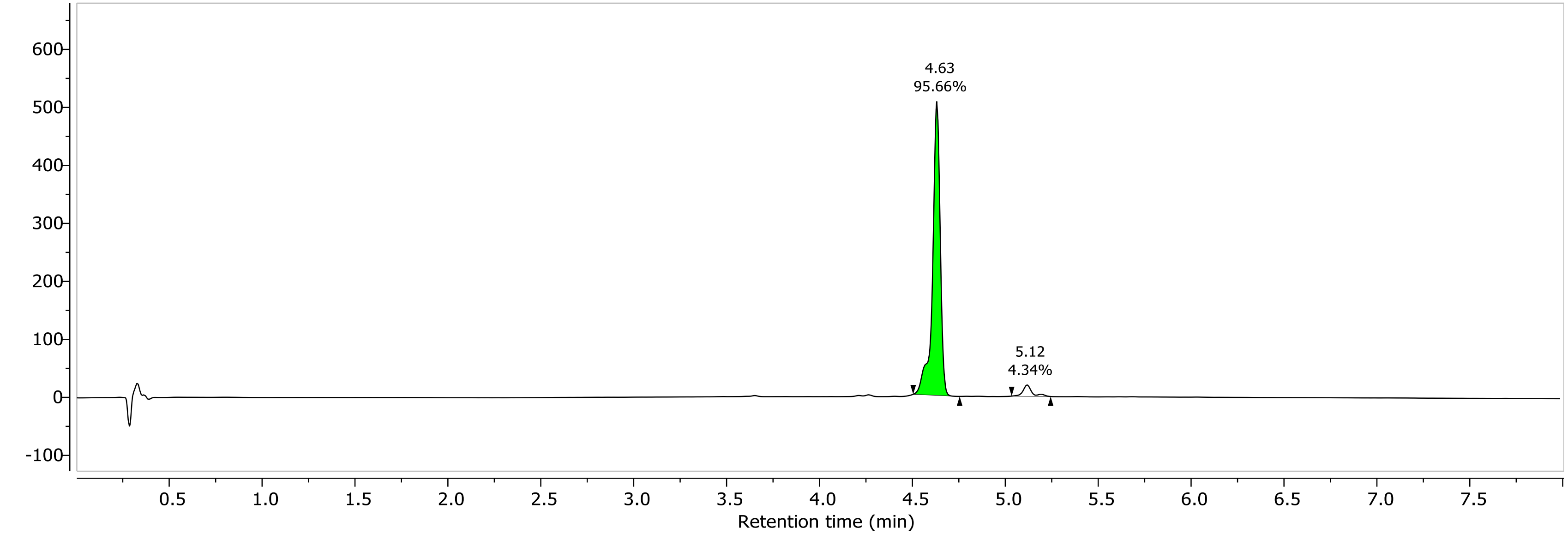

1H NMR (400.26 MHz, DMSO, 298.1 K)

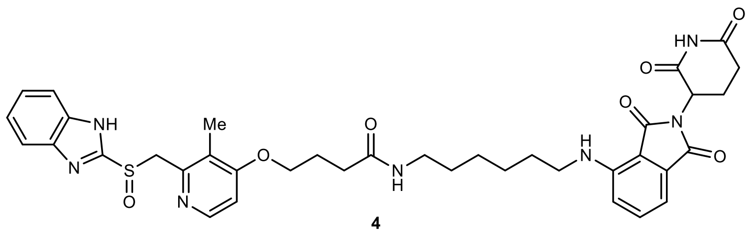

4

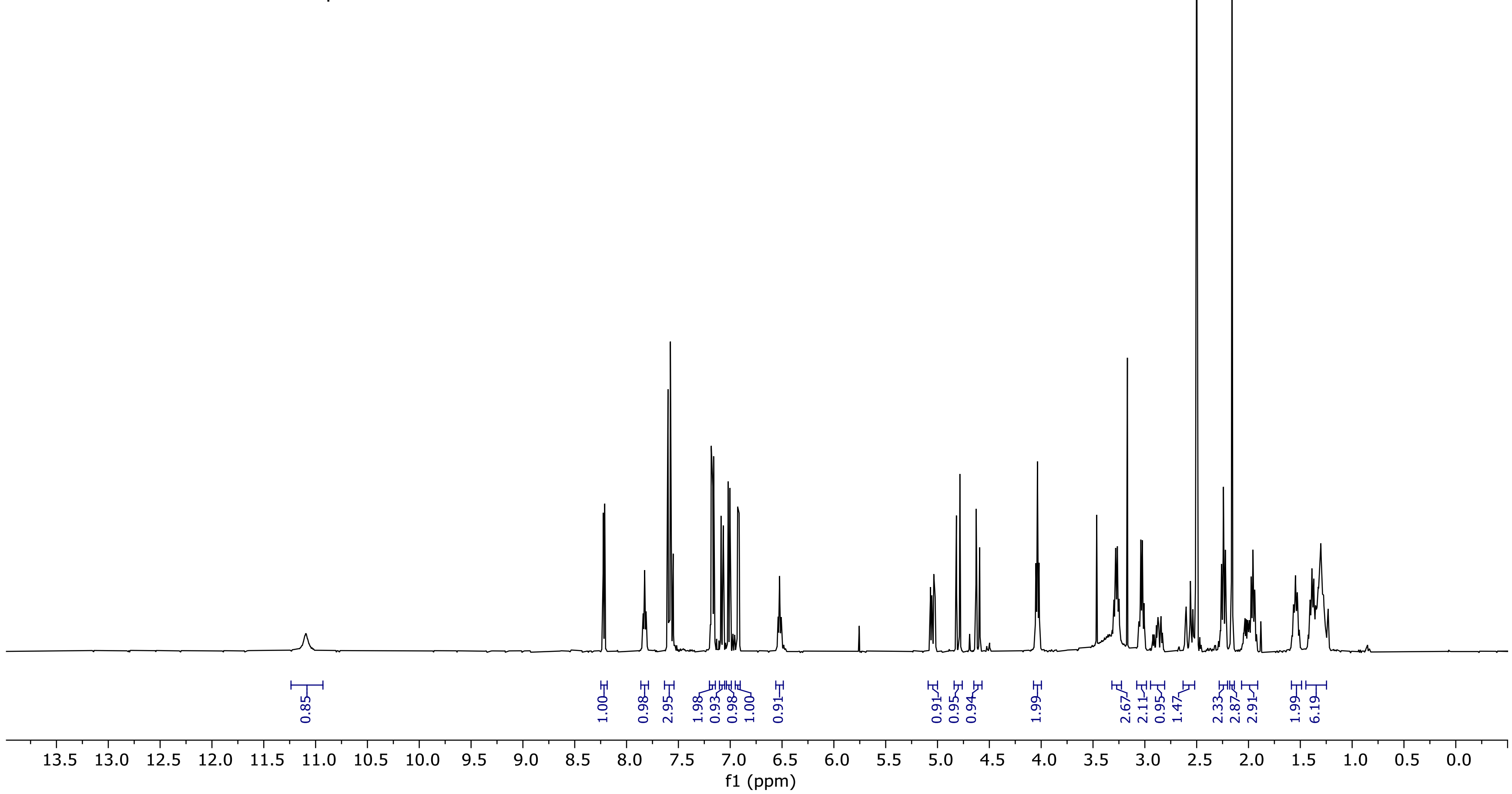

13C NMR (100.66 MHz, DMSO, 298.1 K)

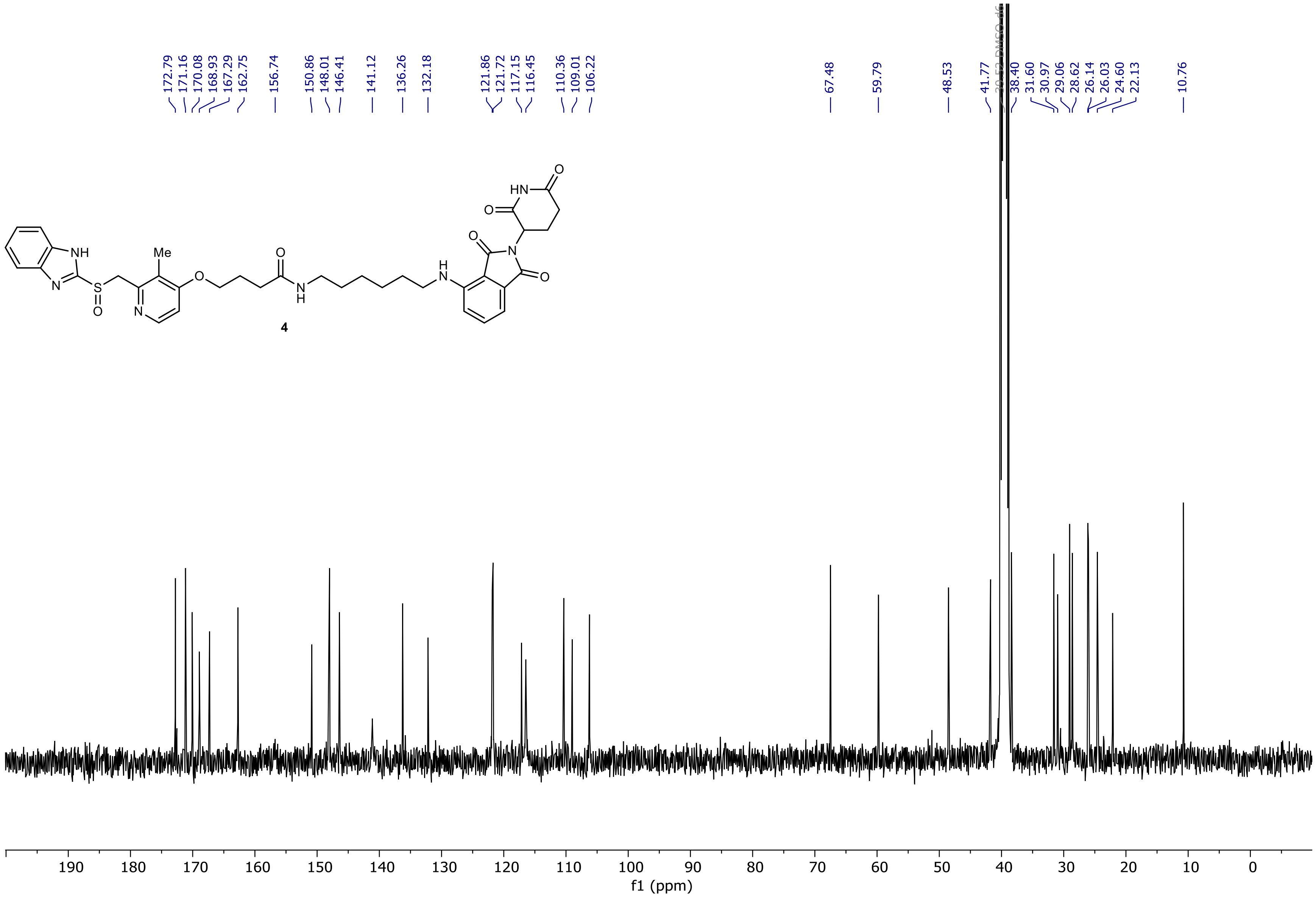

NR-3-062-B-basic Instrument 1  PDA - Total Absorbance Chromatogram

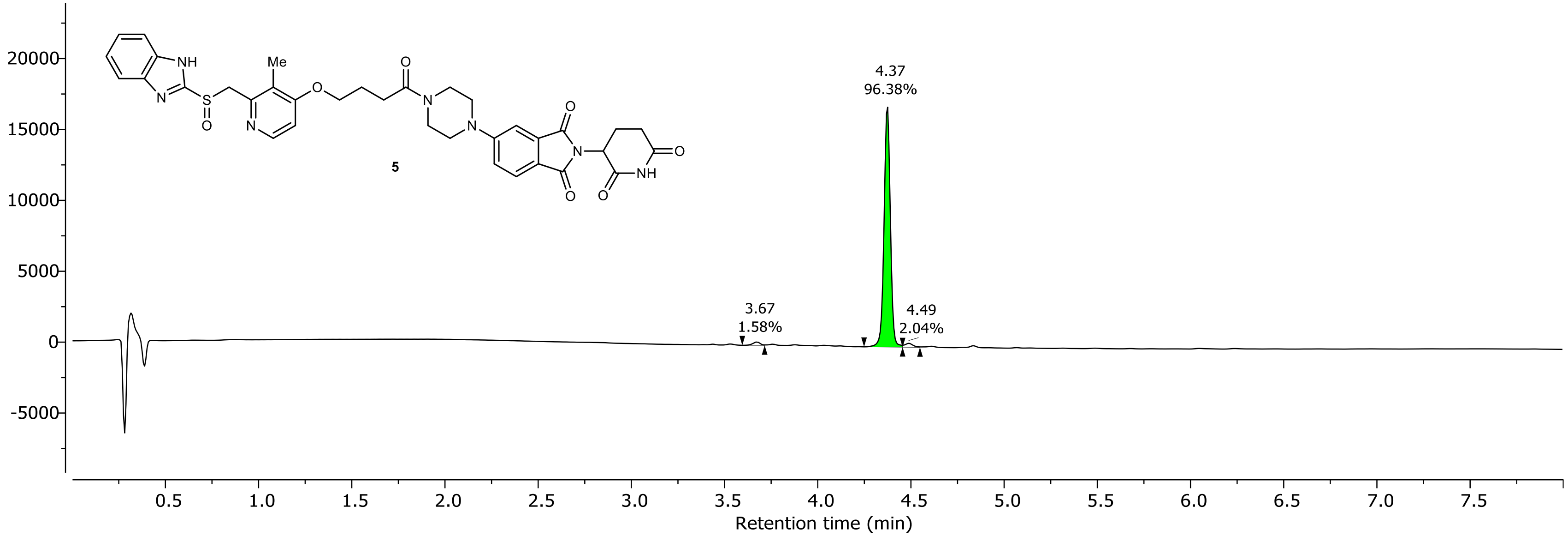

NR-3-062-B-basic Instrument 1 DAD1A, Sig=254...Ref=360.0,100.0 Chromatogram

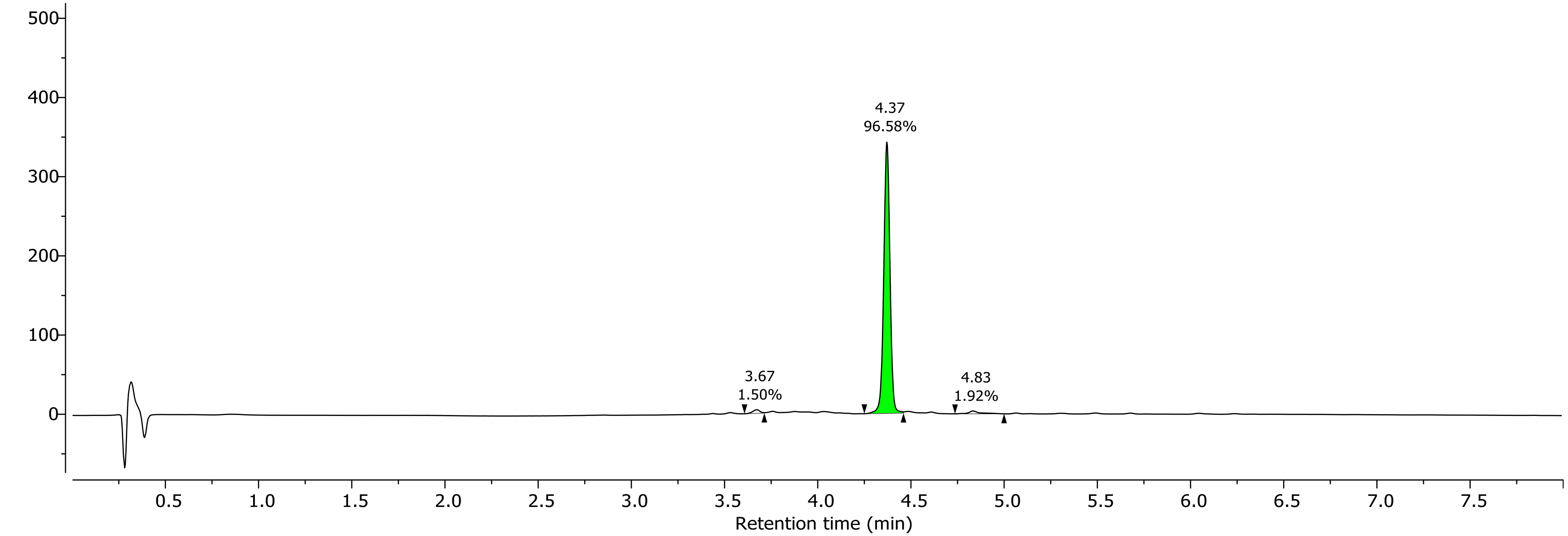

1H NMR (400.26 MHz, DMSO, 298.1 K)

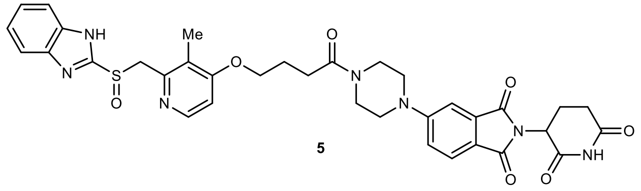

**5**

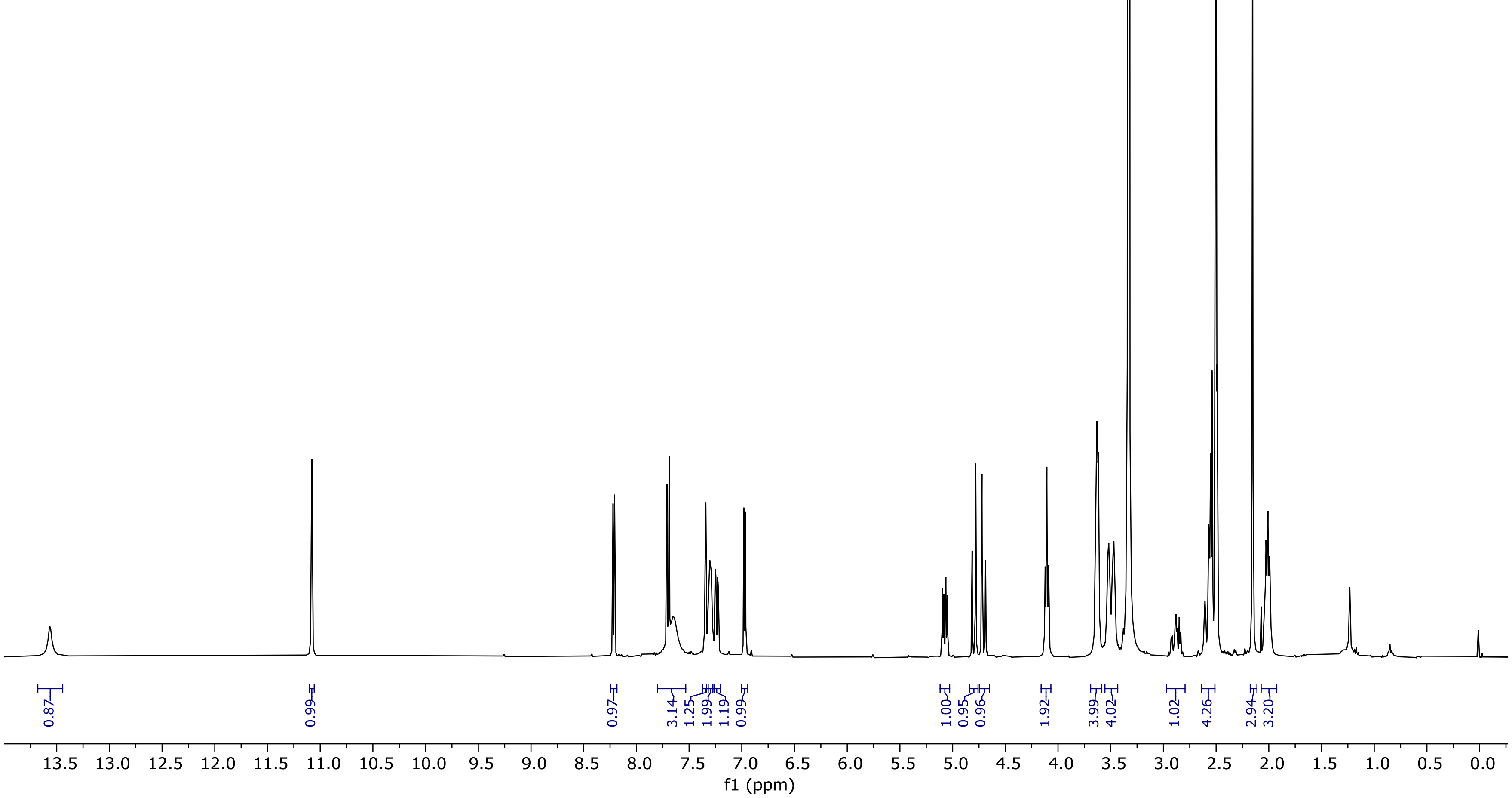

13C NMR (100.66 MHz, DMSO, 298.1 K)

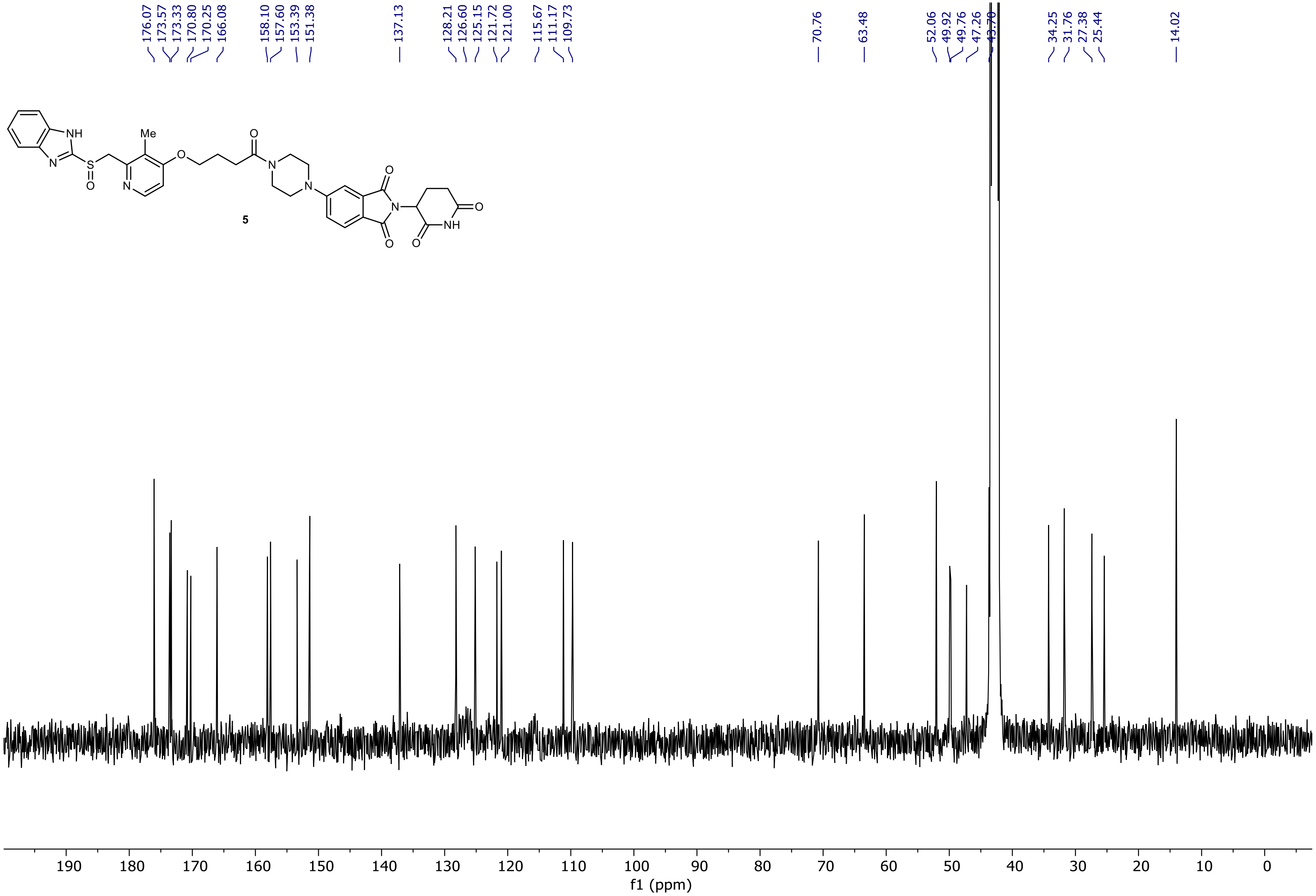

NR-6-003-A LC-MS_ELS  PDA - Total Absorbance Chromatogram

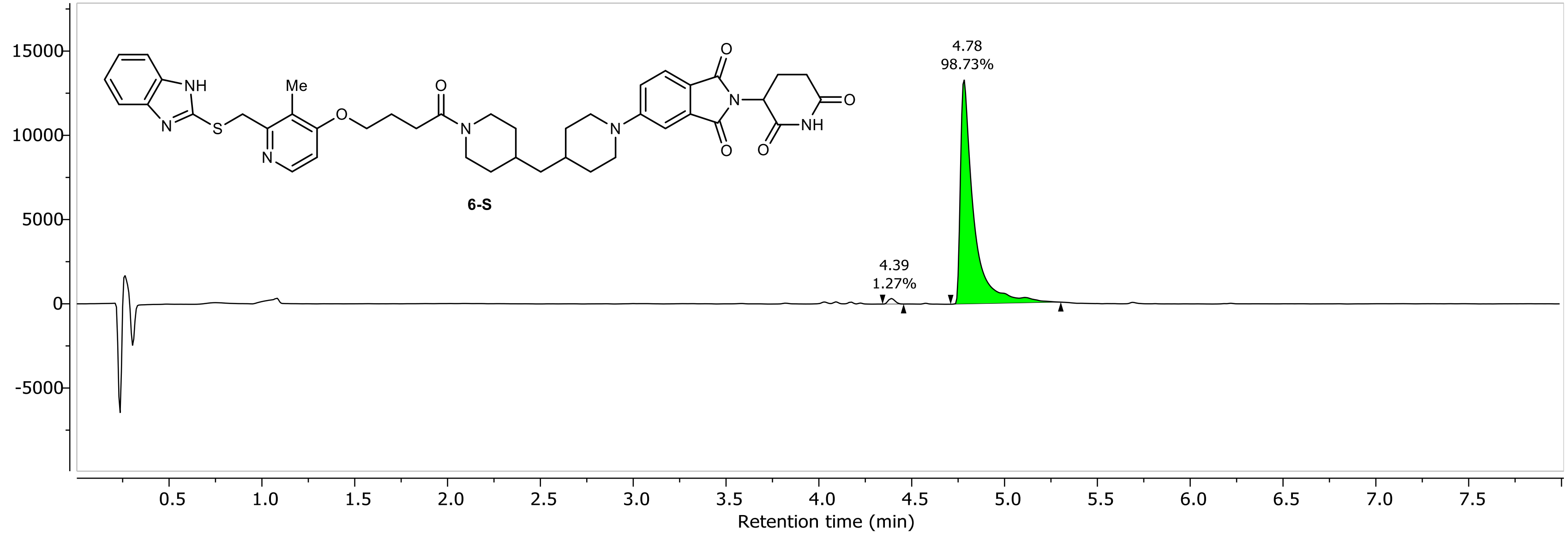

NR-6-003-A LC-MS_ELS DAD1A, Sig=254...Ref=360.0,100.0 Chromatogram

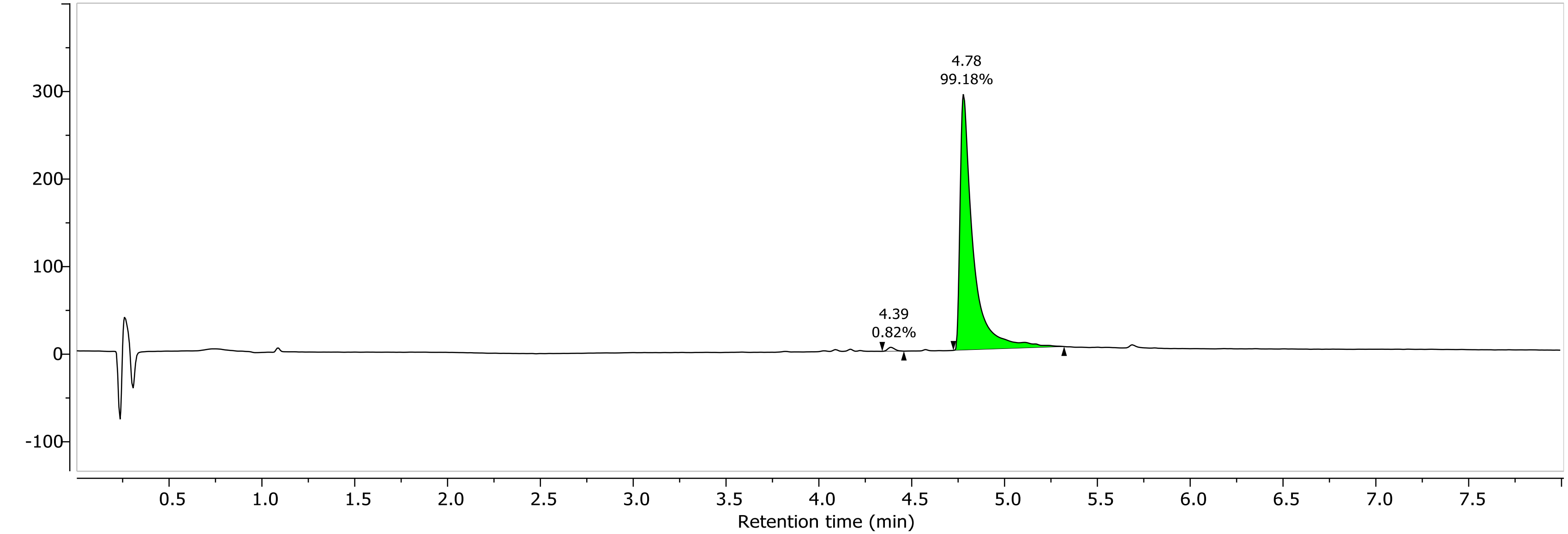

1H NMR (400.26 MHz, CDCl3, 298.1 K)

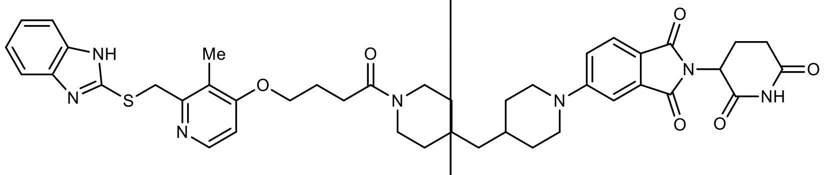

**6-S**

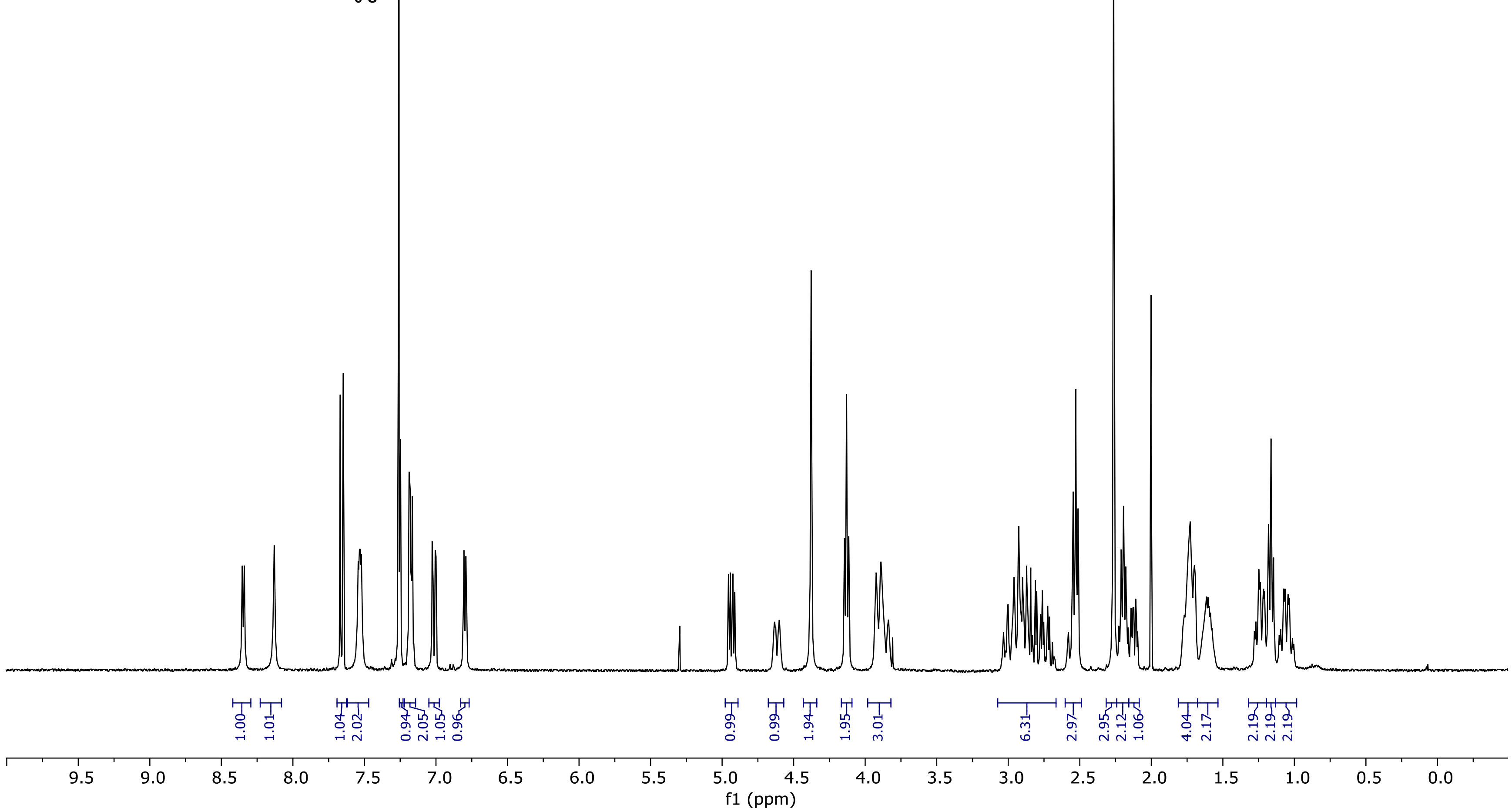

13C NMR (100.66 MHz, CDCl3, 298.1 K)

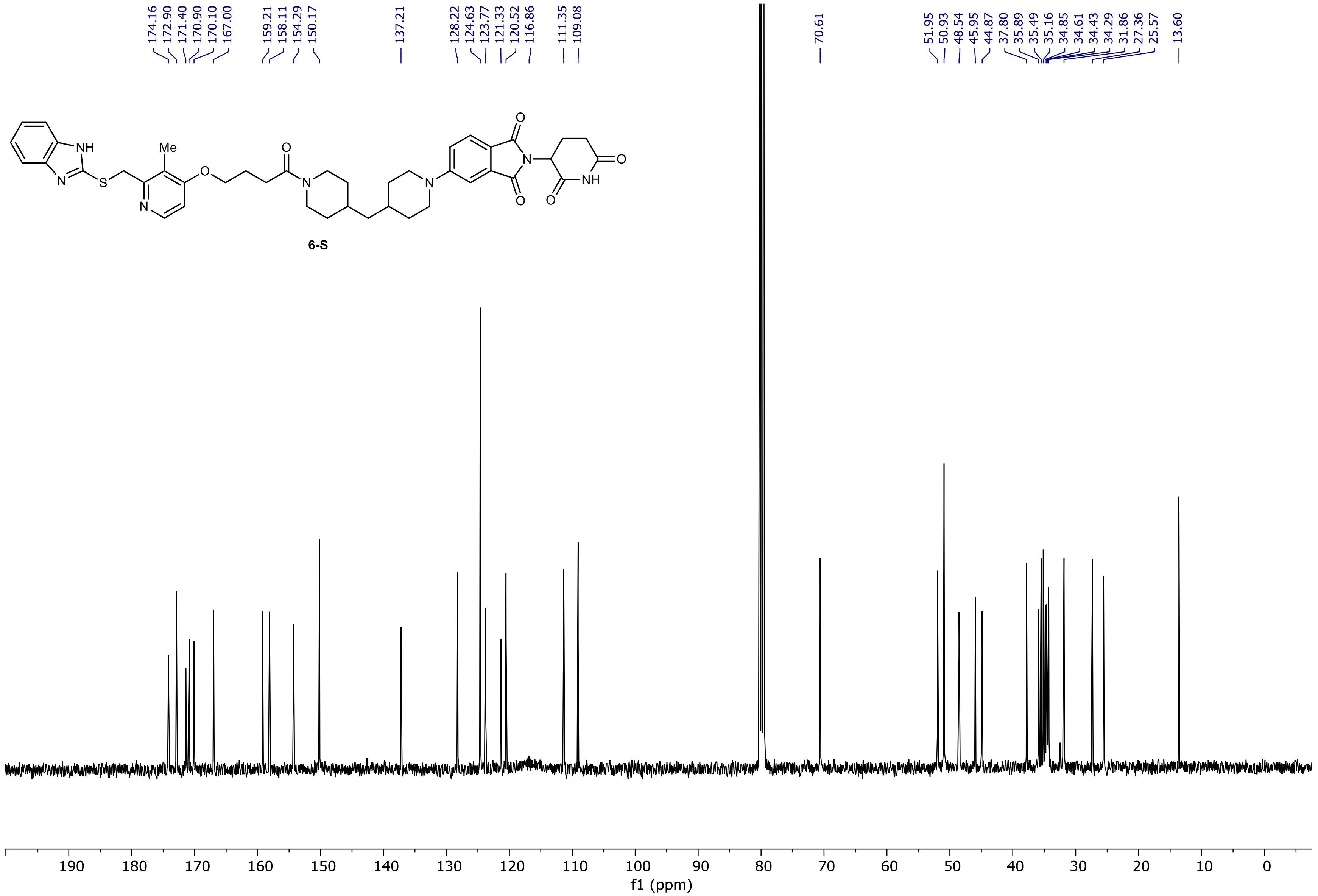

NR-3-143-A Instrument 1 PDA - Total Absorbance Chromatogram

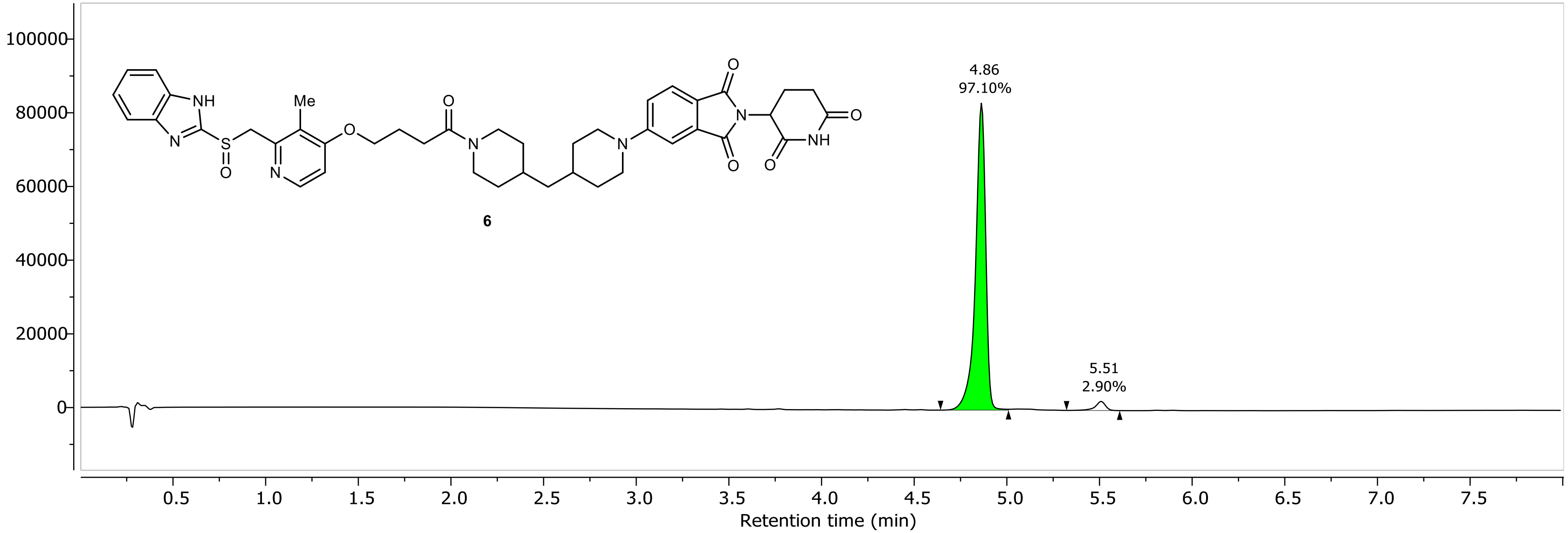

NR-3-143-A Instrument 1 DAD1A, Sig=254...Ref=360.0,100.0 Chromatogram

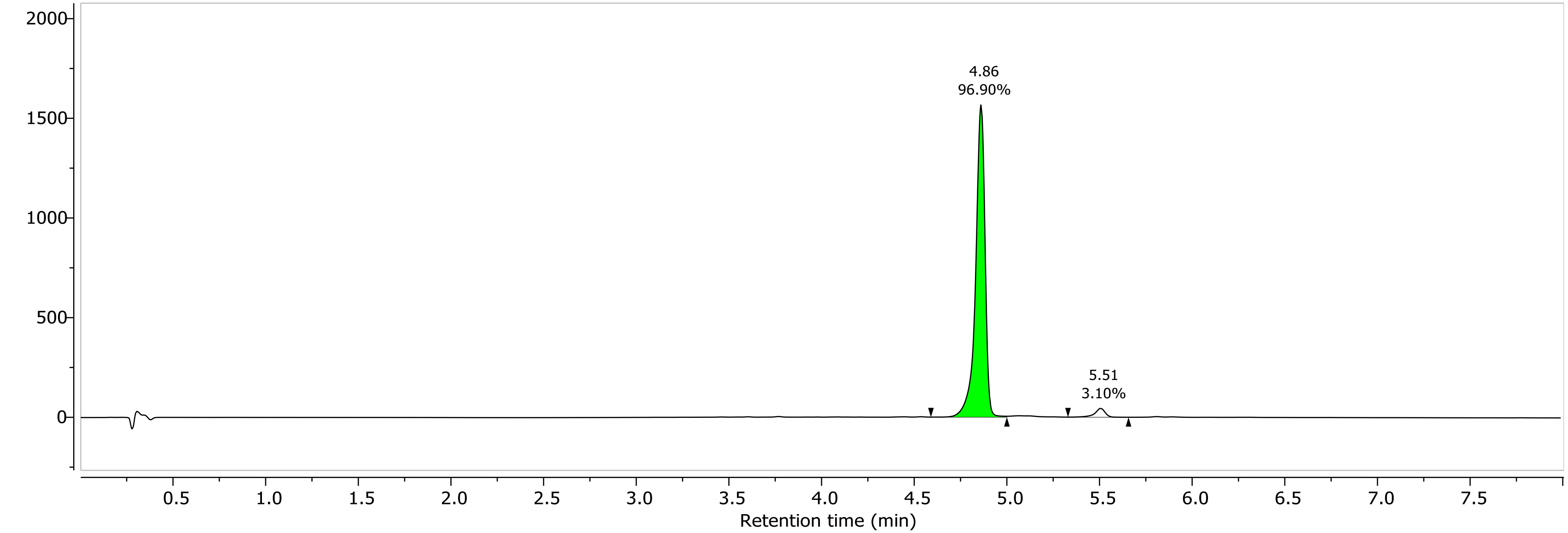

1H NMR (400.26 MHz, DMSO, 298.1 K)

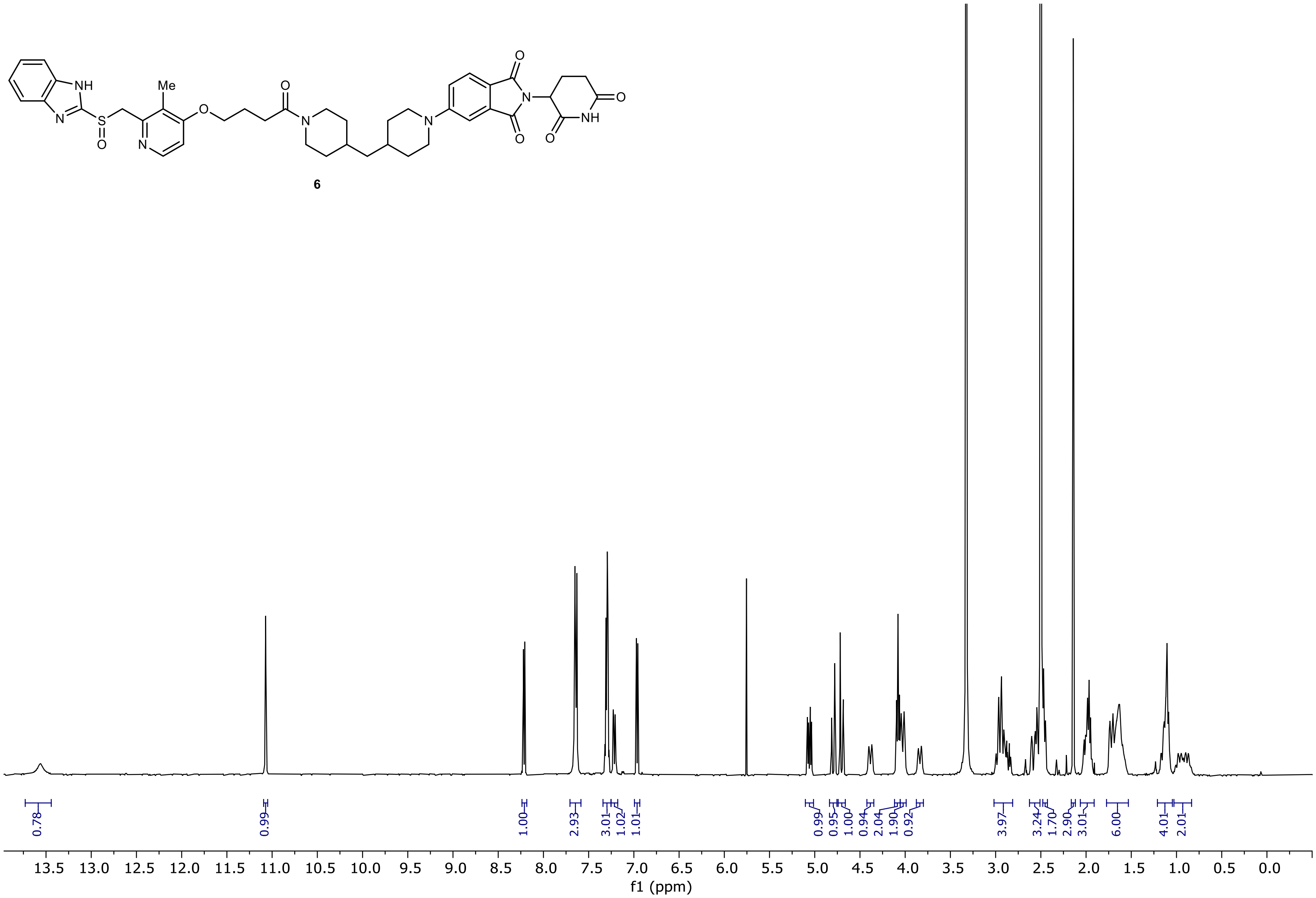

13C NMR (100.66 MHz, DMSO, 298.1 K)

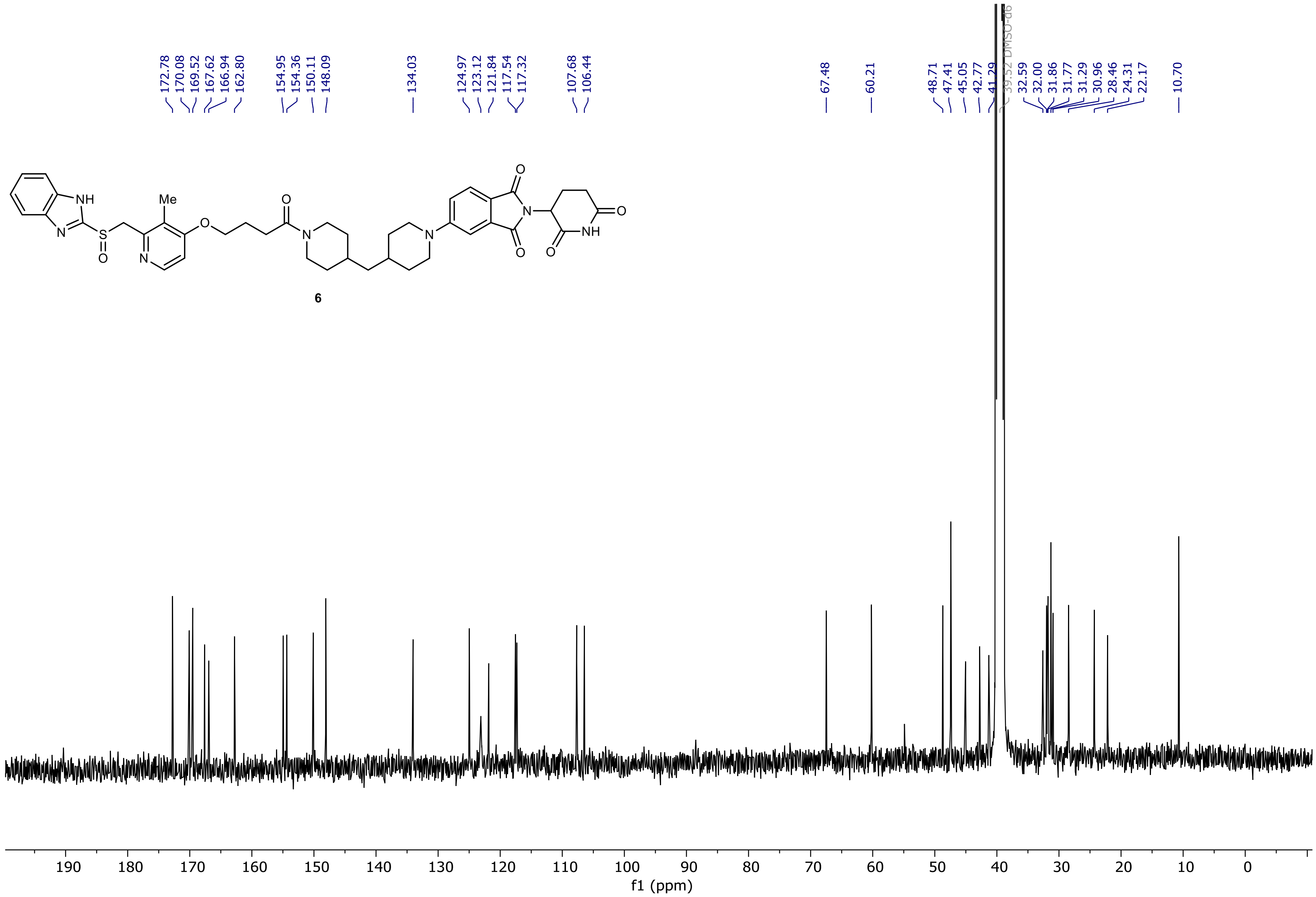

NR-5-142-A LC-MS_ELS  PDA - Total Absorbance Chromatogram

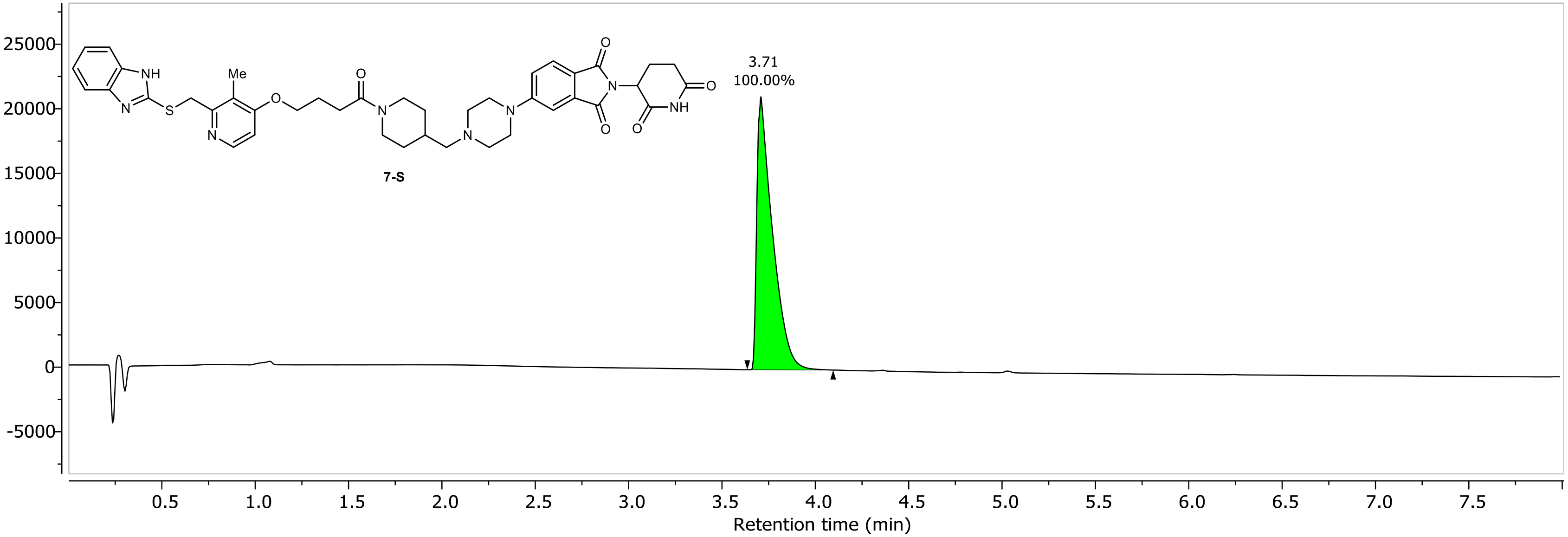

NR-5-142-A LC-MS_ELS DAD1A, Sig=254...Ref=360.0,100.0 Chromatogram

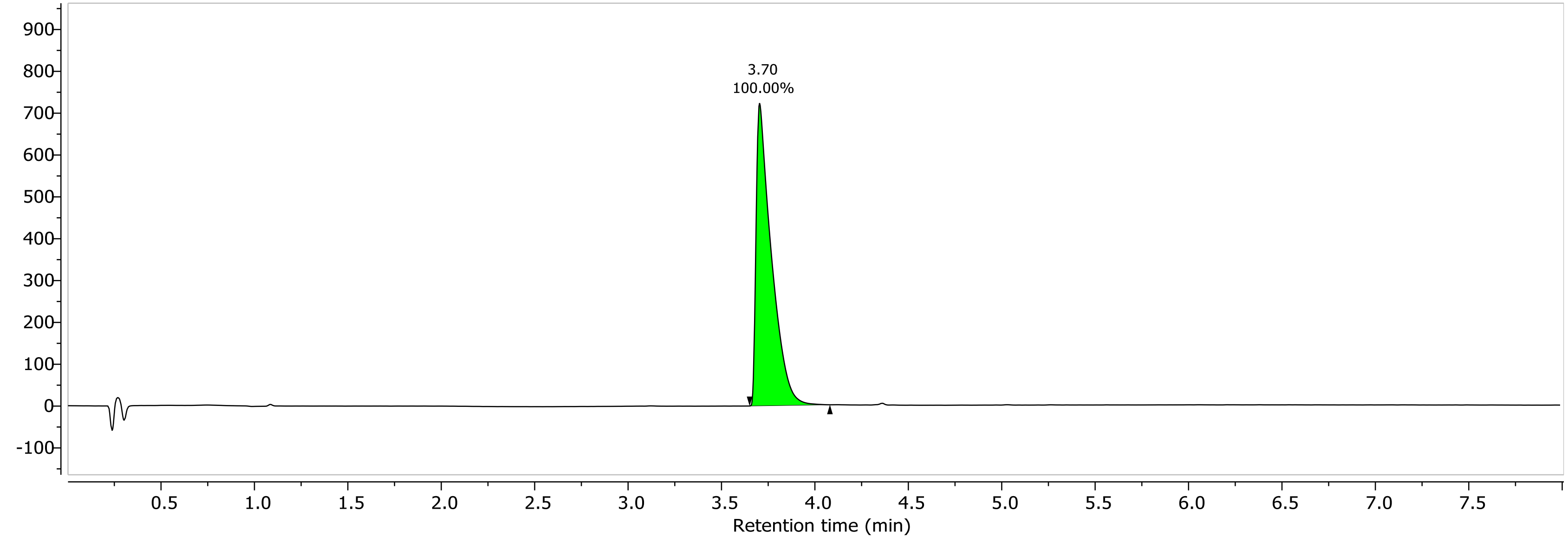

1H NMR (400.26 MHz, DMSO, 298.1 K)

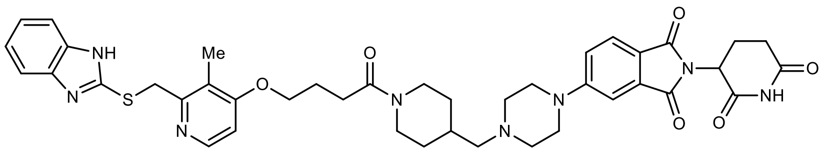

**7-S**

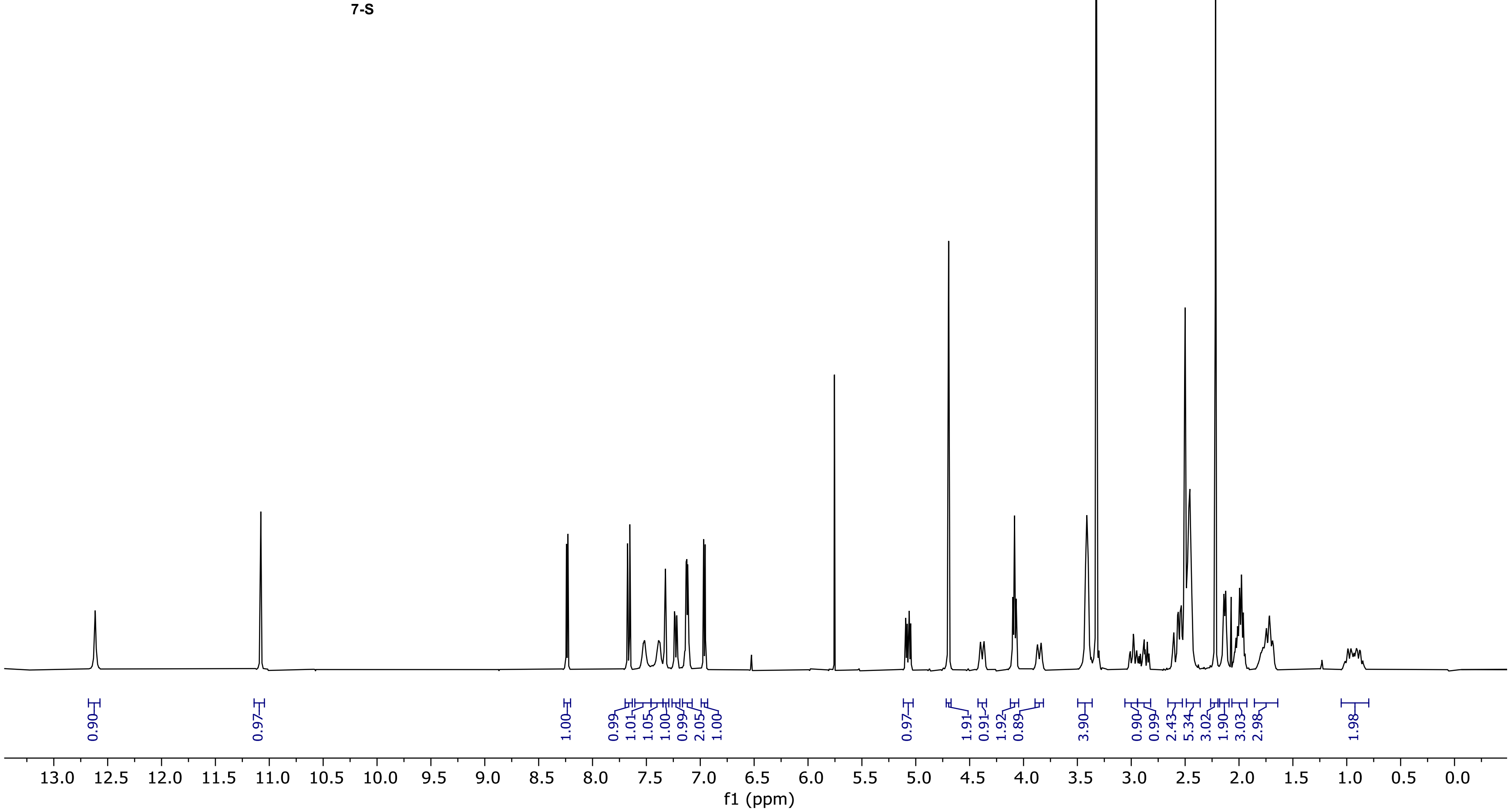

13C NMR (100.66 MHz, DMSO, 298.1 K)

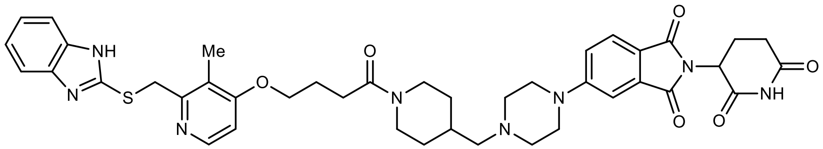

**7-S**

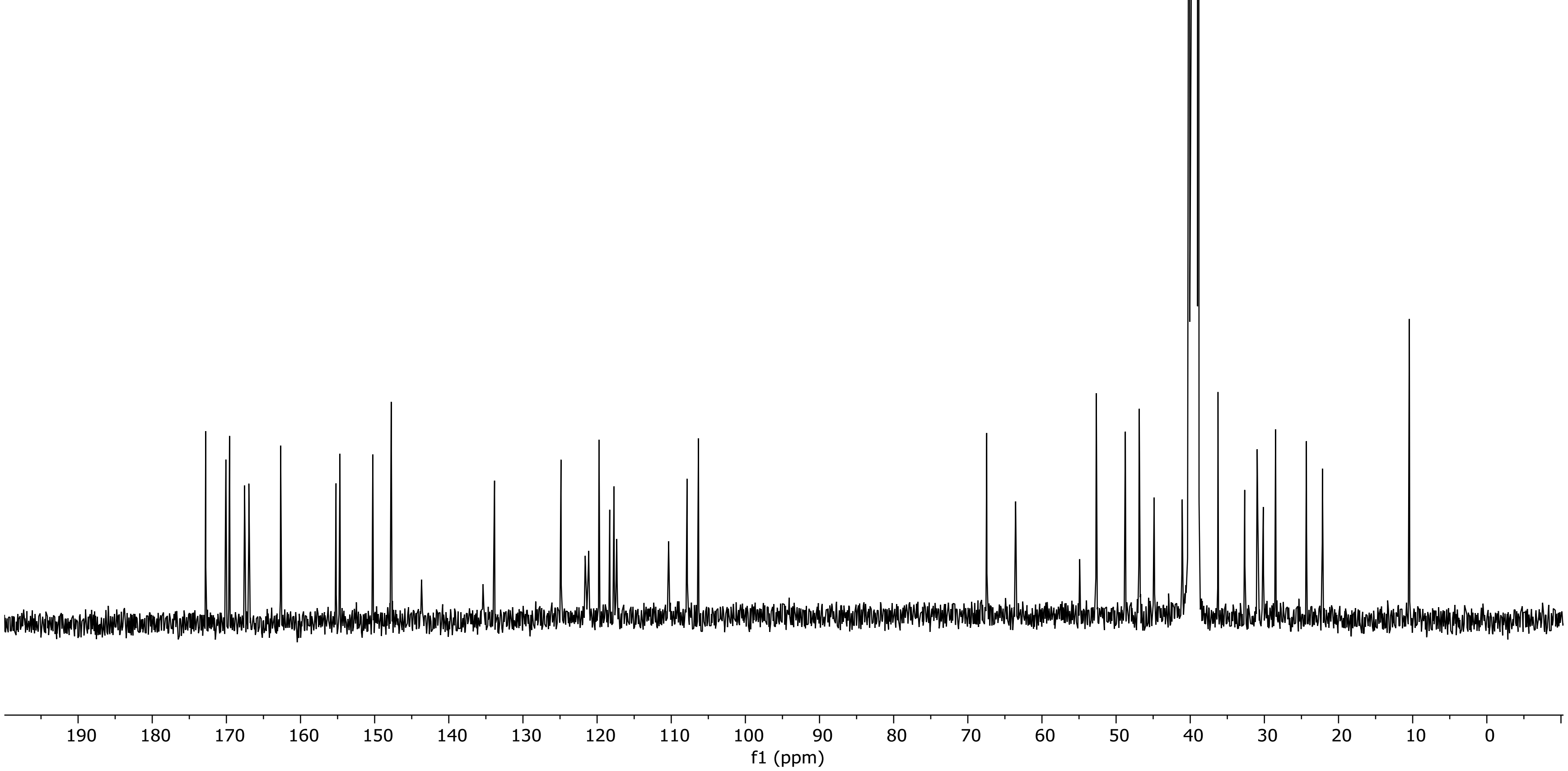

NR-3-149-A Instrument 1 PDA - Total Absorbance Chromatogram

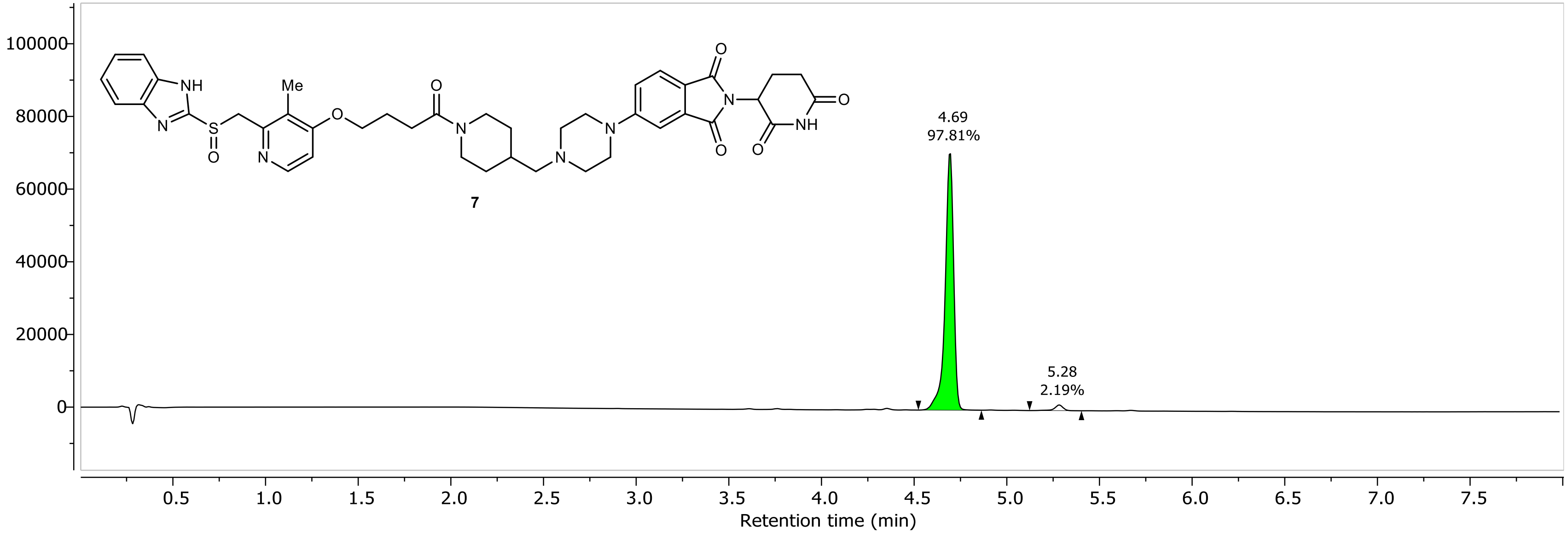

NR-3-149-A Instrument 1 DAD1A, Sig=254...Ref=360.0,100.0 Chromatogram

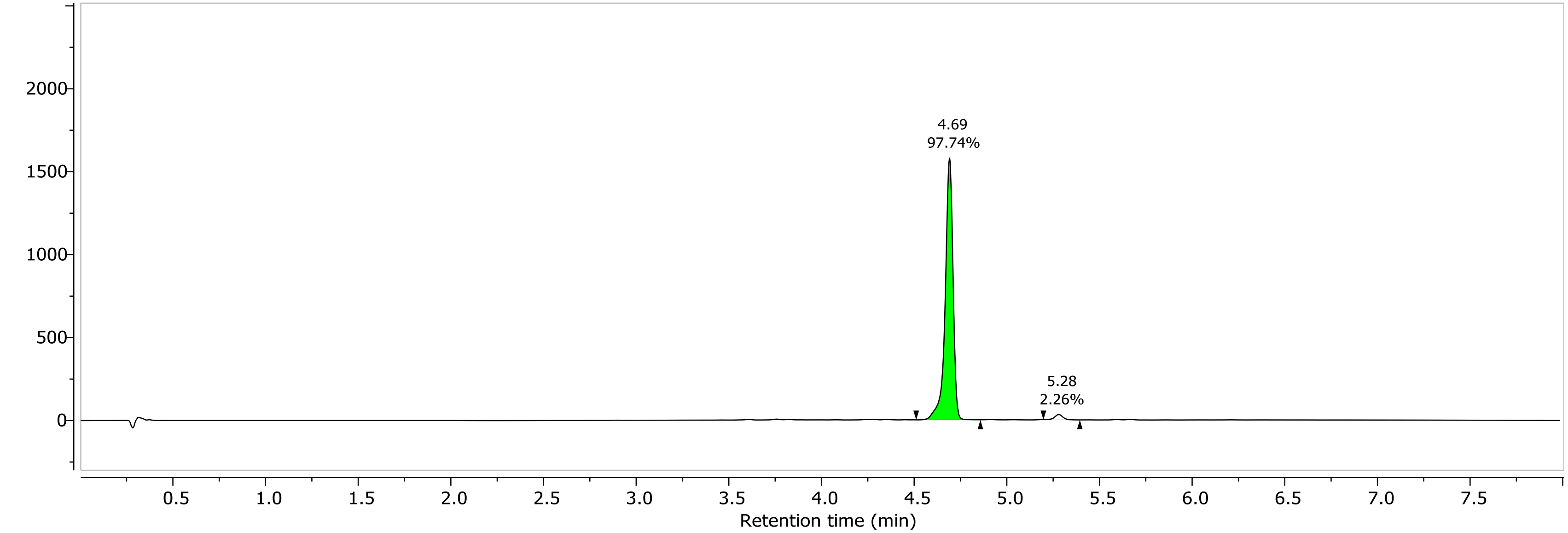

1H NMR (400.26 MHz, DMSO, 298.2 K)

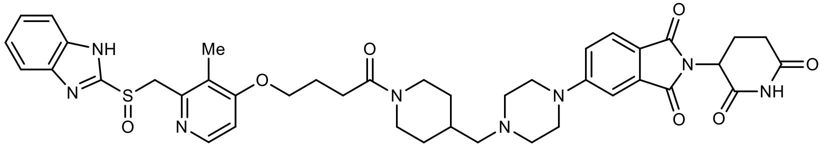

7

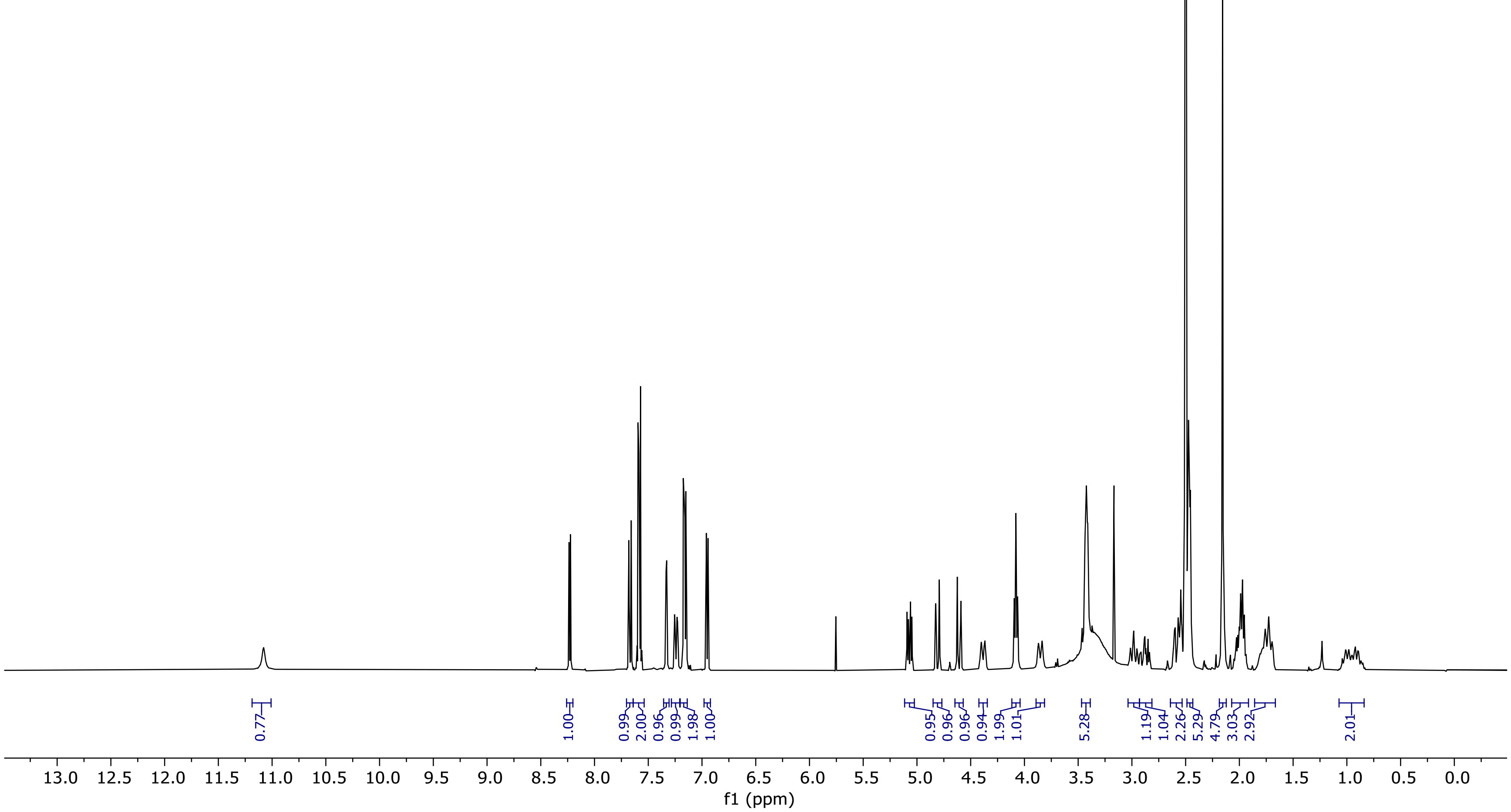

13C NMR (100.66 MHz, DMSO, 298.1 K)

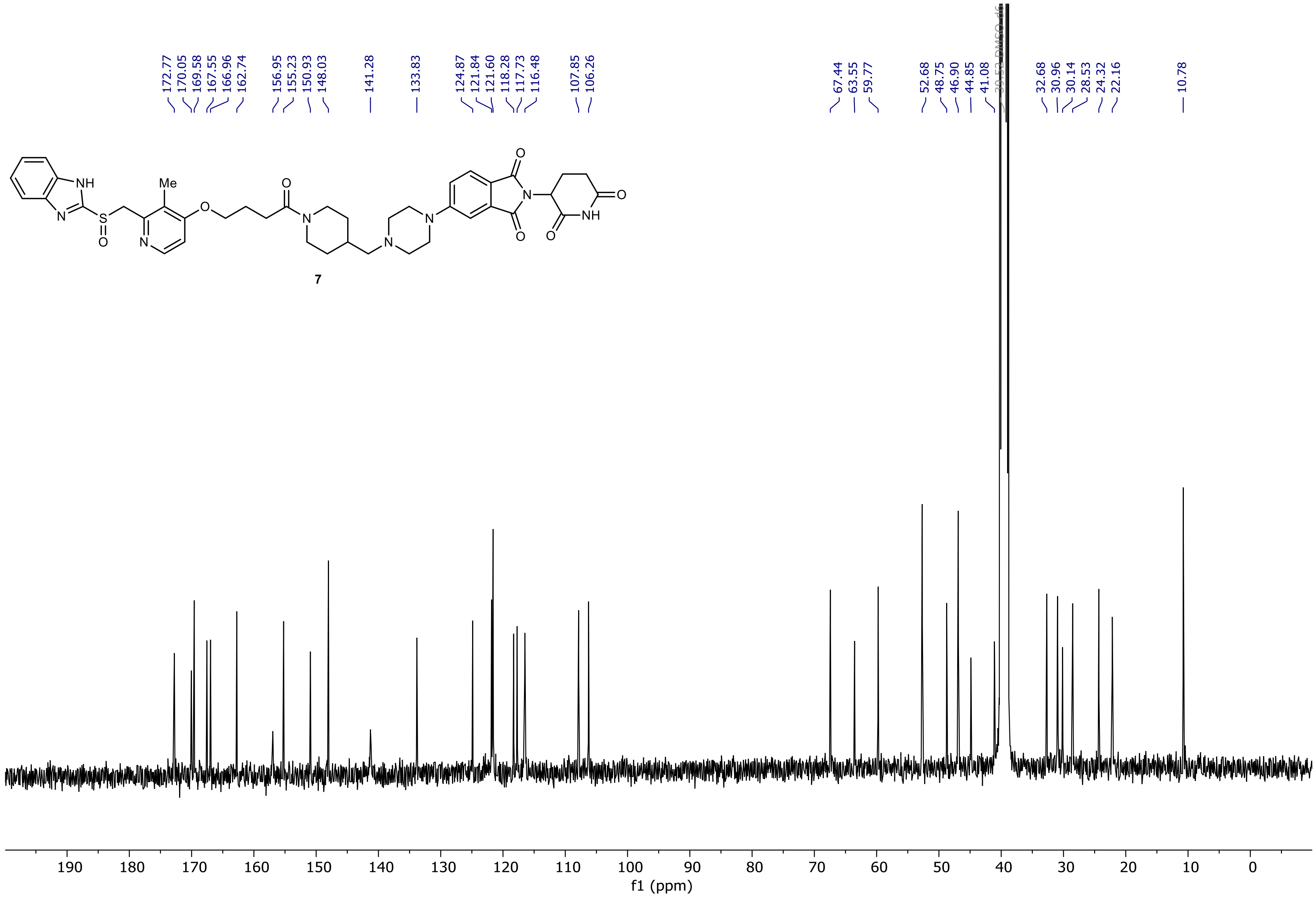

7

NR-5-156-A LC-MS_ELS  PDA - Total Absorbance Chromatogram

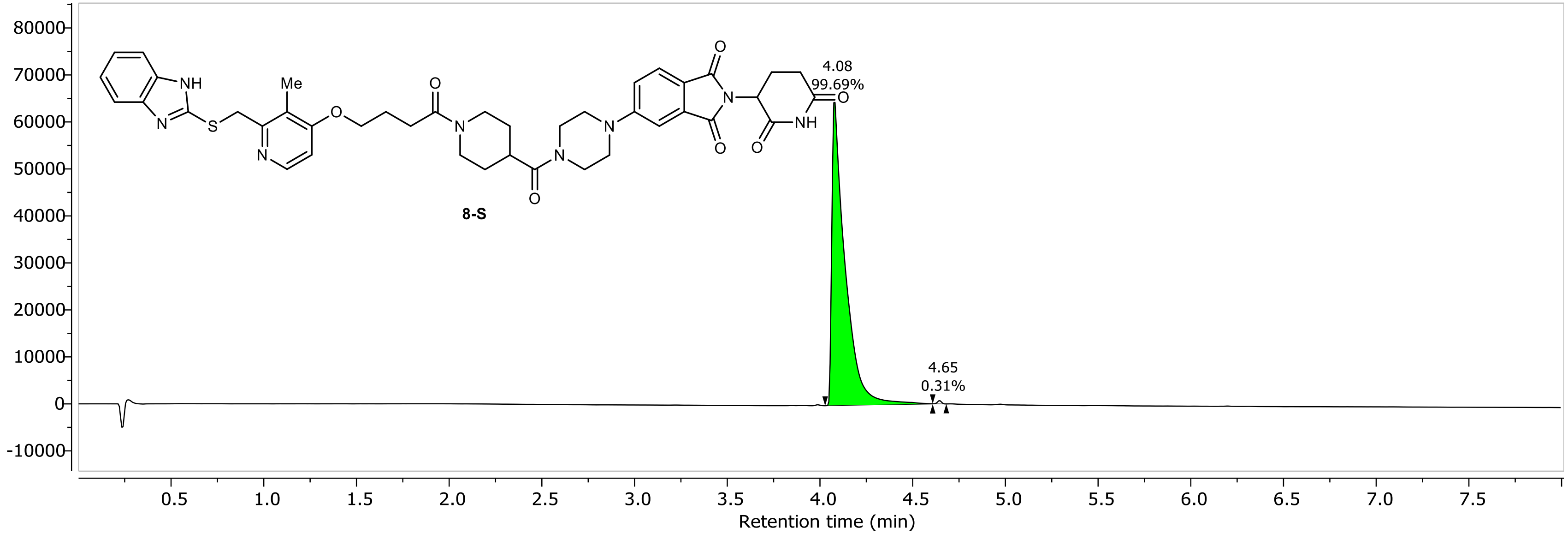

NR-5-156-A LC-MS_ELS DAD1A, Sig=254...Ref=360.0,100.0 Chromatogram

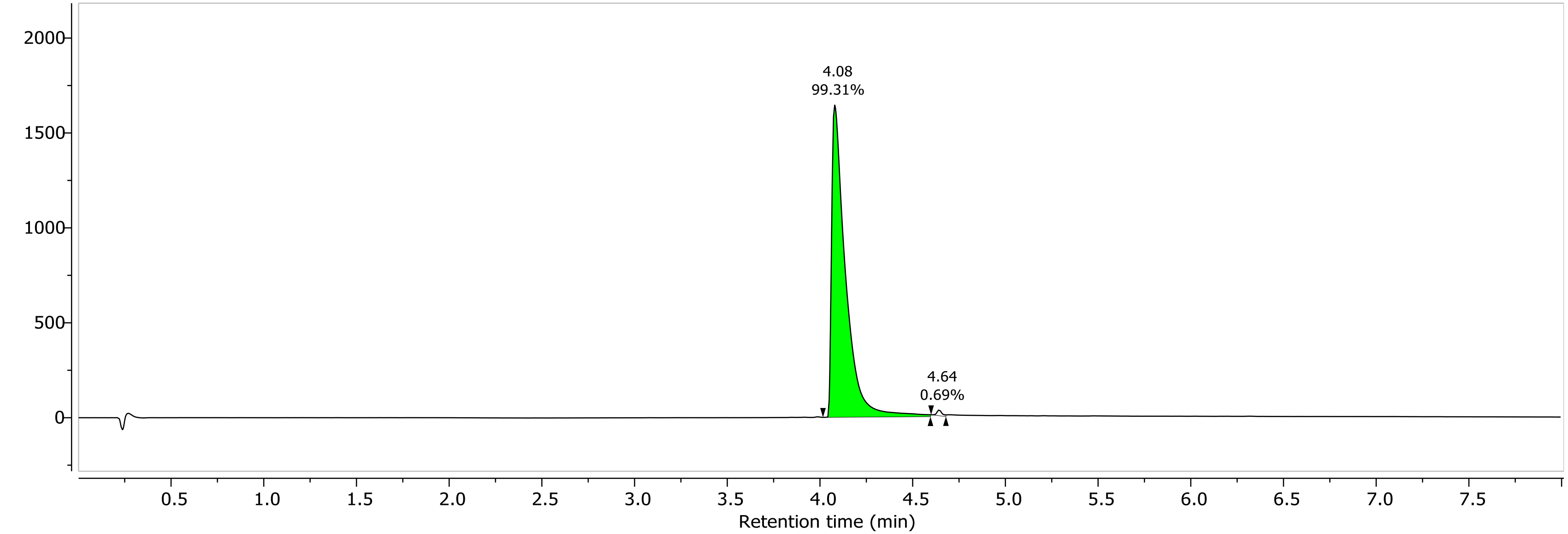

1H NMR (400.26 MHz, CDCl3, 298.1 K)

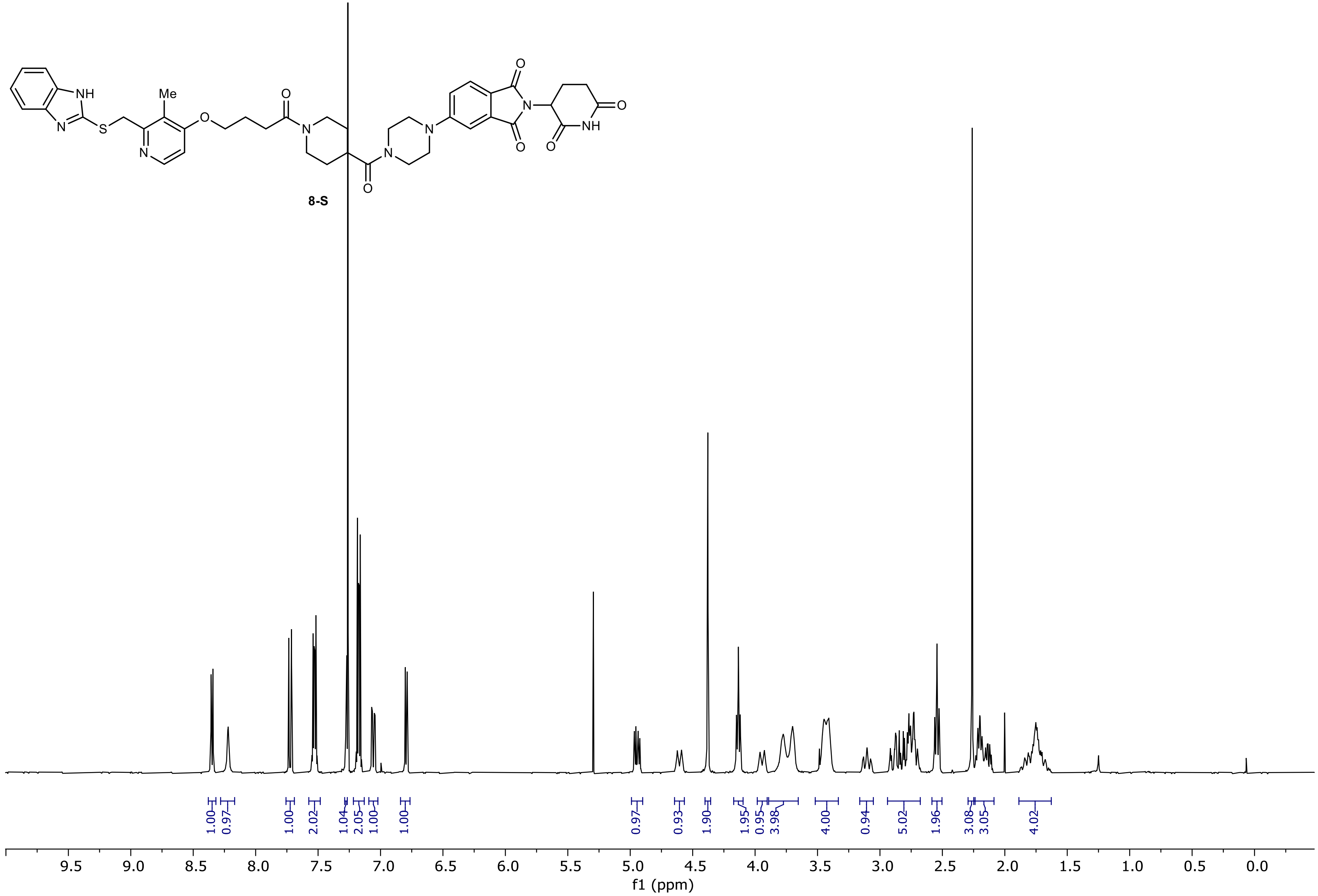

13C NMR (100.66 MHz, CDCl3, 298.1 K)

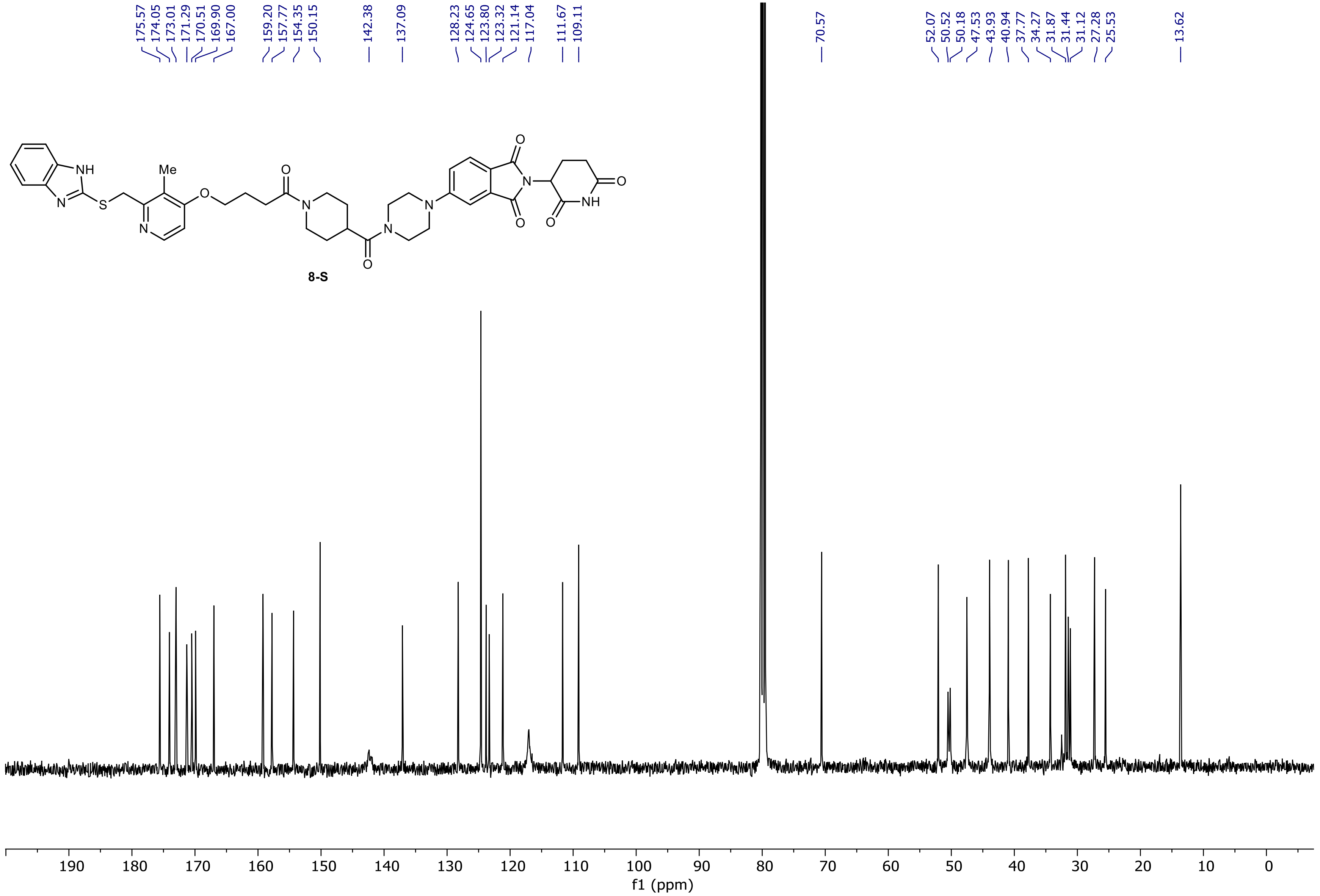

PDA - Total Absorbance Chromatogram

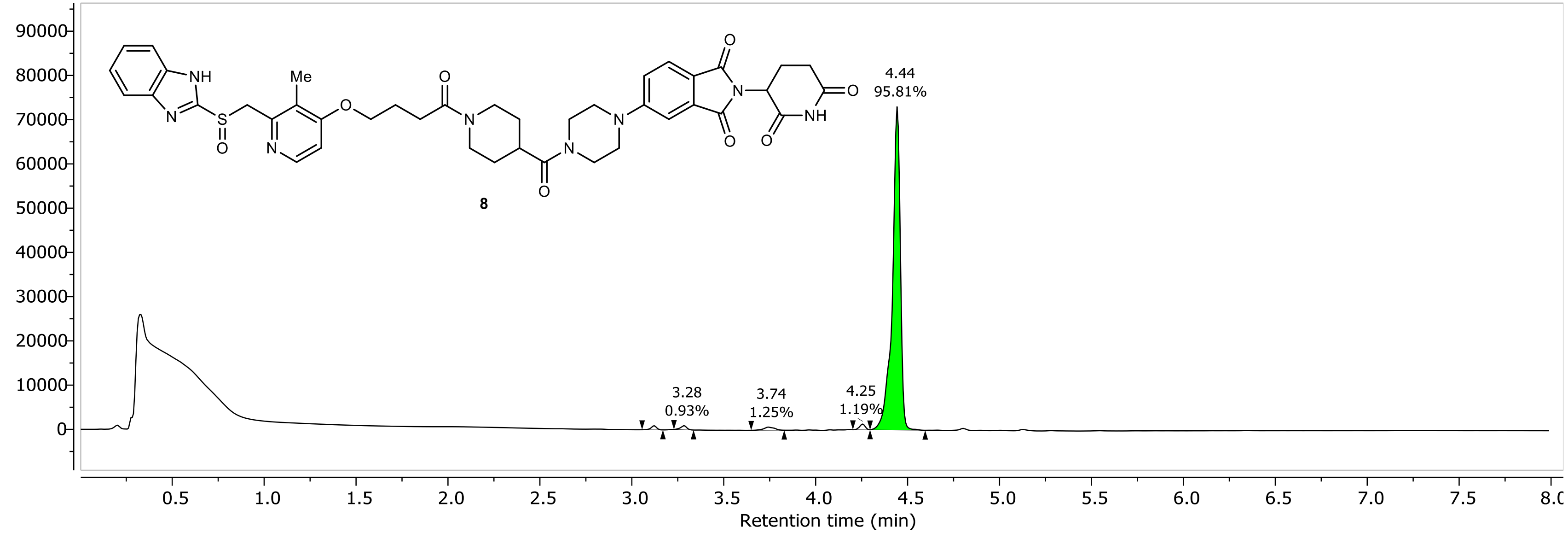

DAD1A, Sig=254...Ref=360.0,100.0 Chromatogram

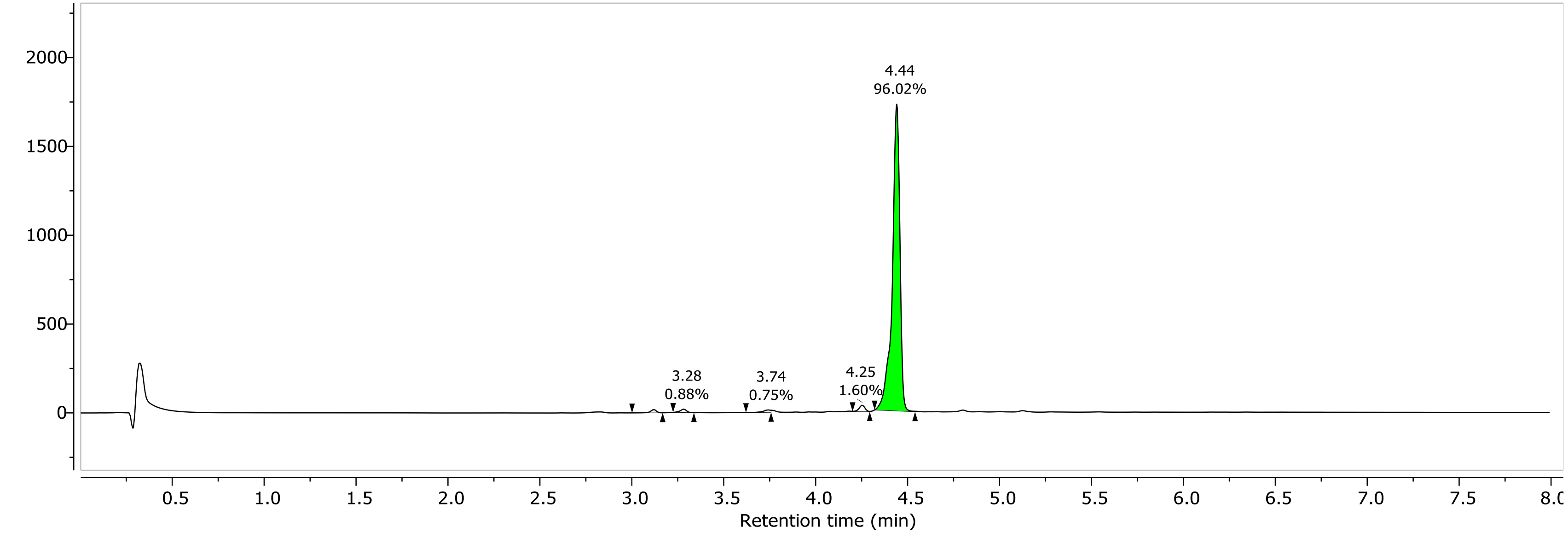

1H NMR (400.26 MHz, DMSO, 298.1 K)

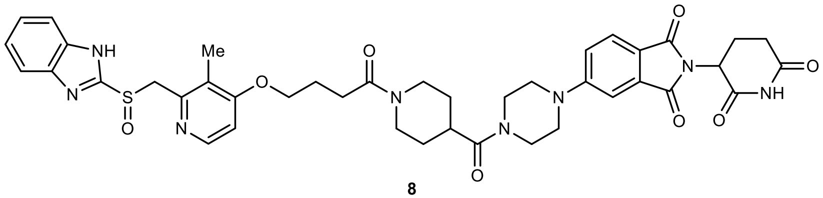

**8**

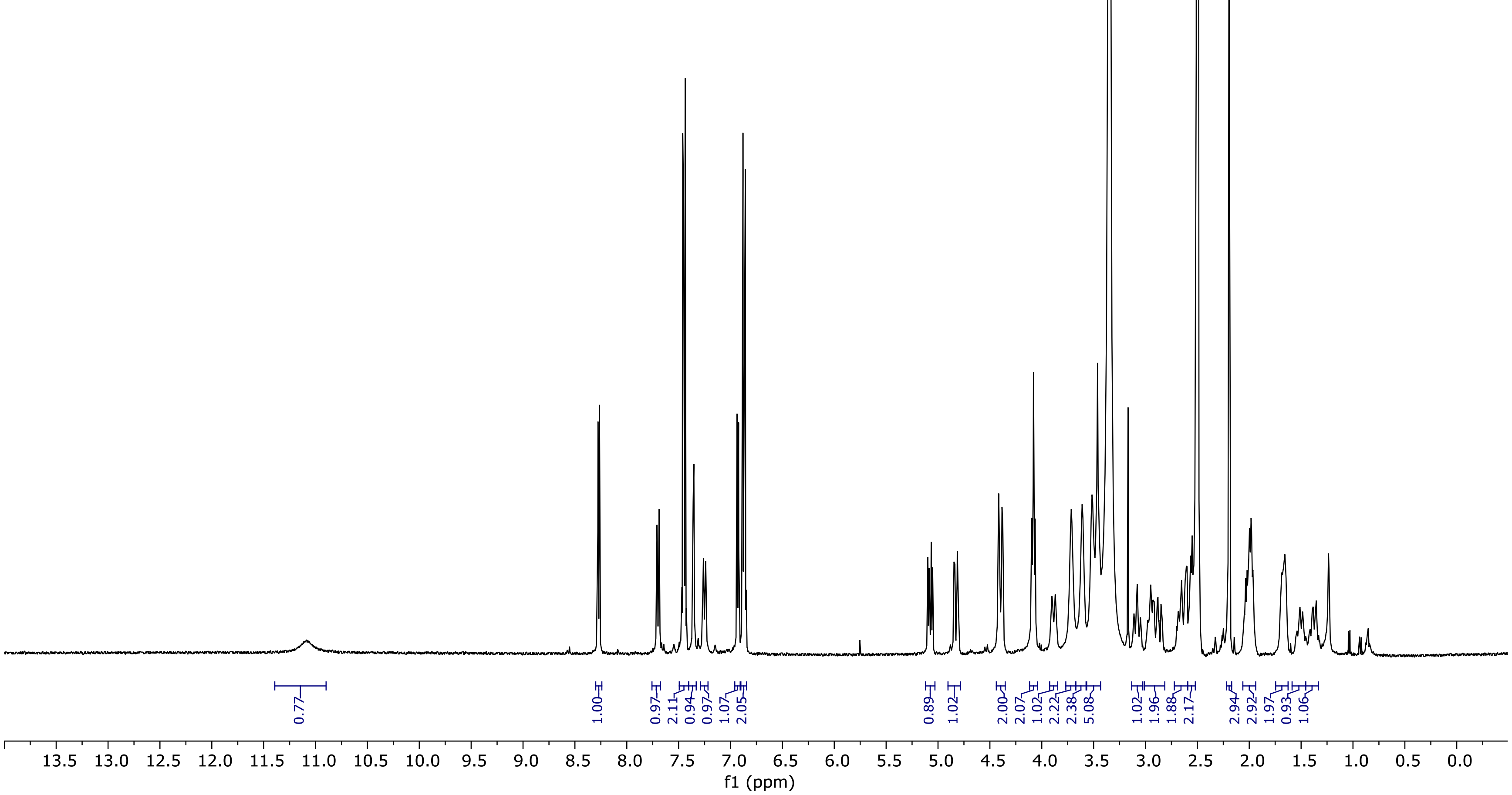

13C NMR (100.66 MHz, DMSO, 298.1 K)

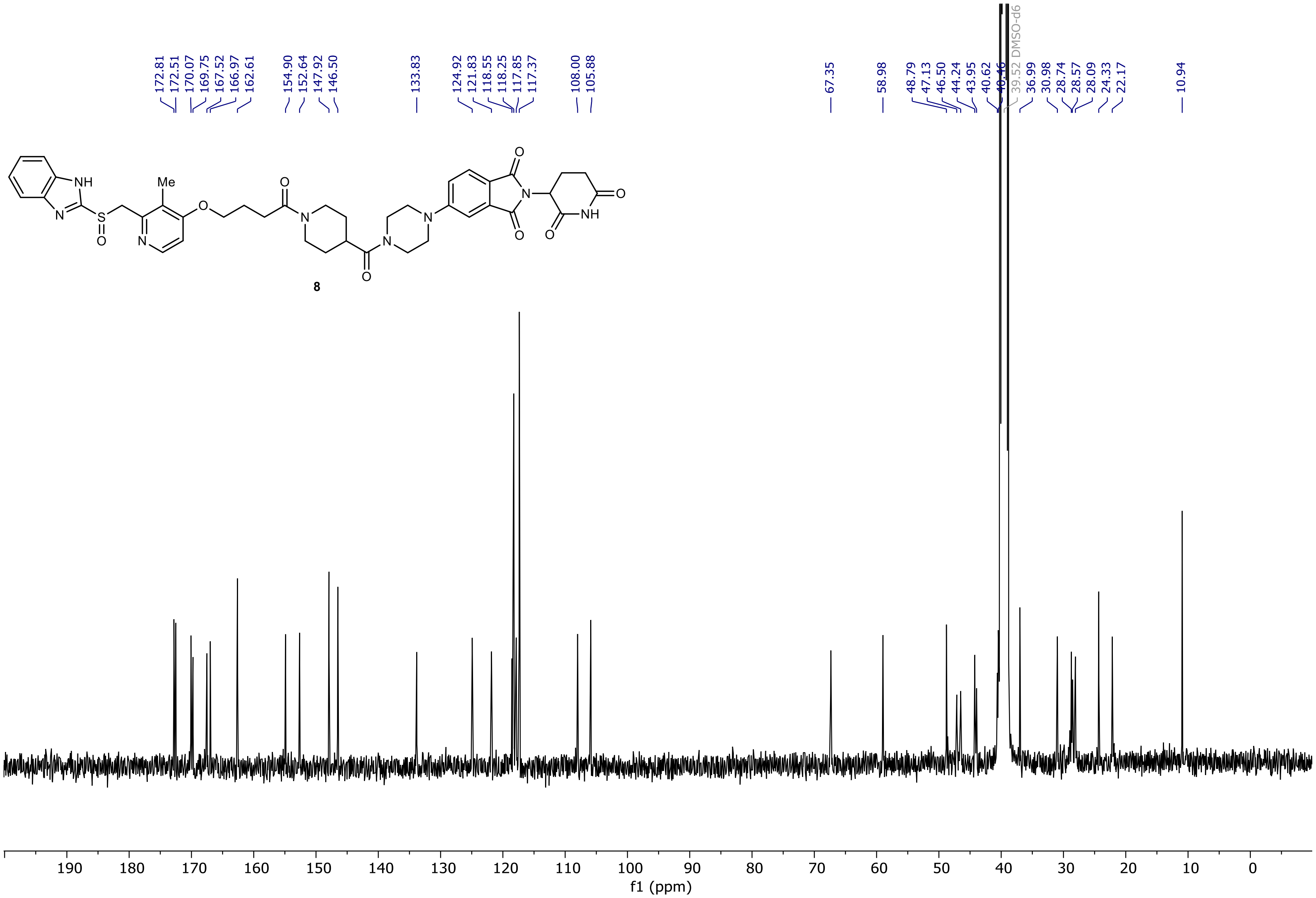

NR-3-152-A Instrument 1  PDA - Total Absorbance Chromatogram

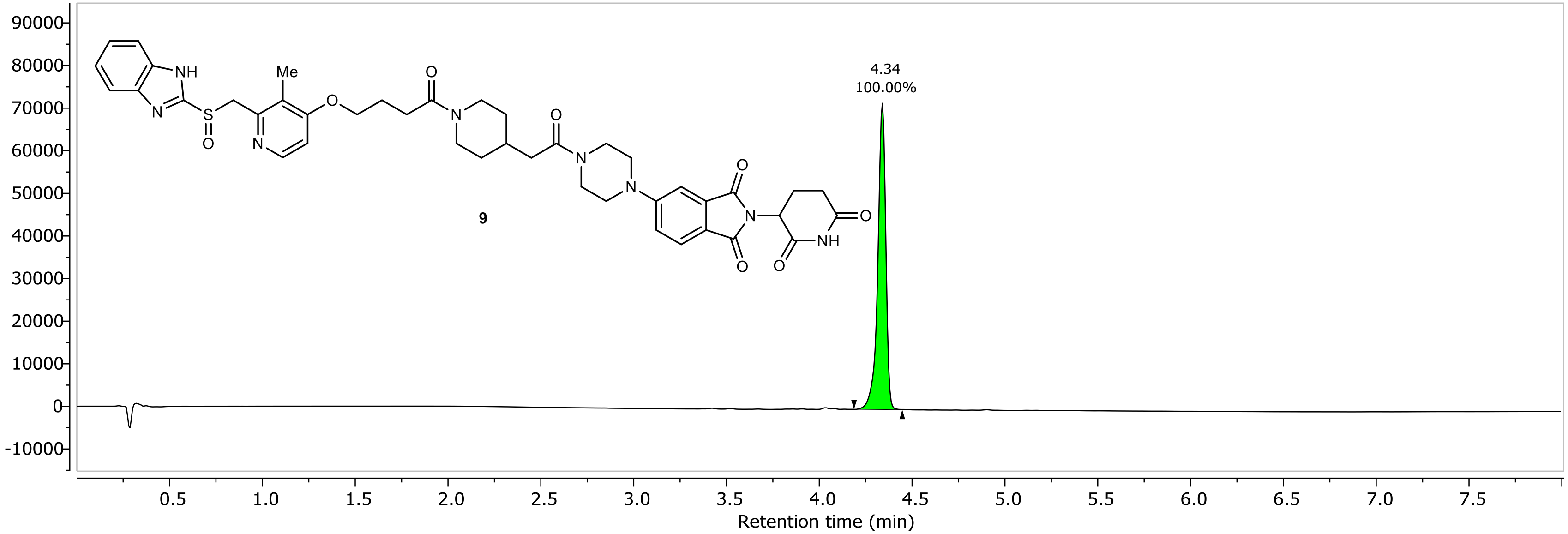

NR-3-152-A Instrument 1 DAD1A, Sig=254...Ref=360.0,100.0 Chromatogram

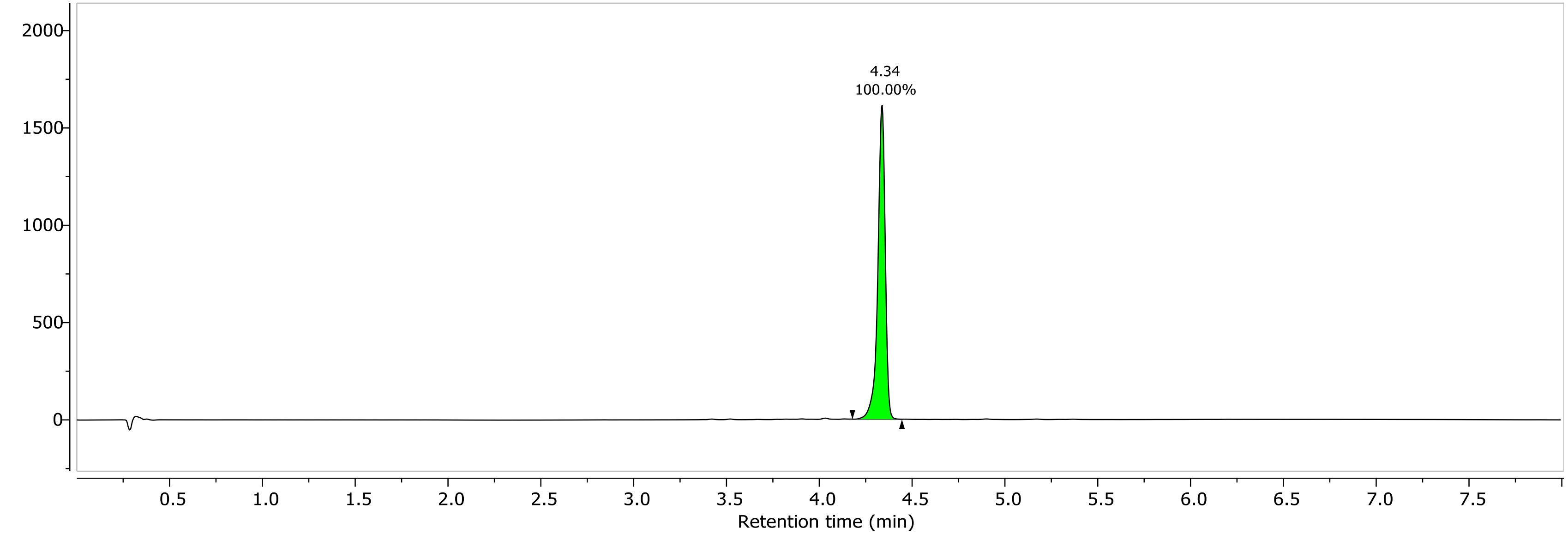

1H NMR (400.26 MHz, DMSO, 298.1 K)

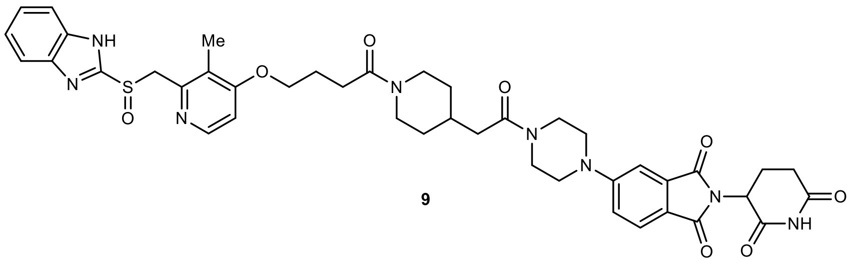

**9**

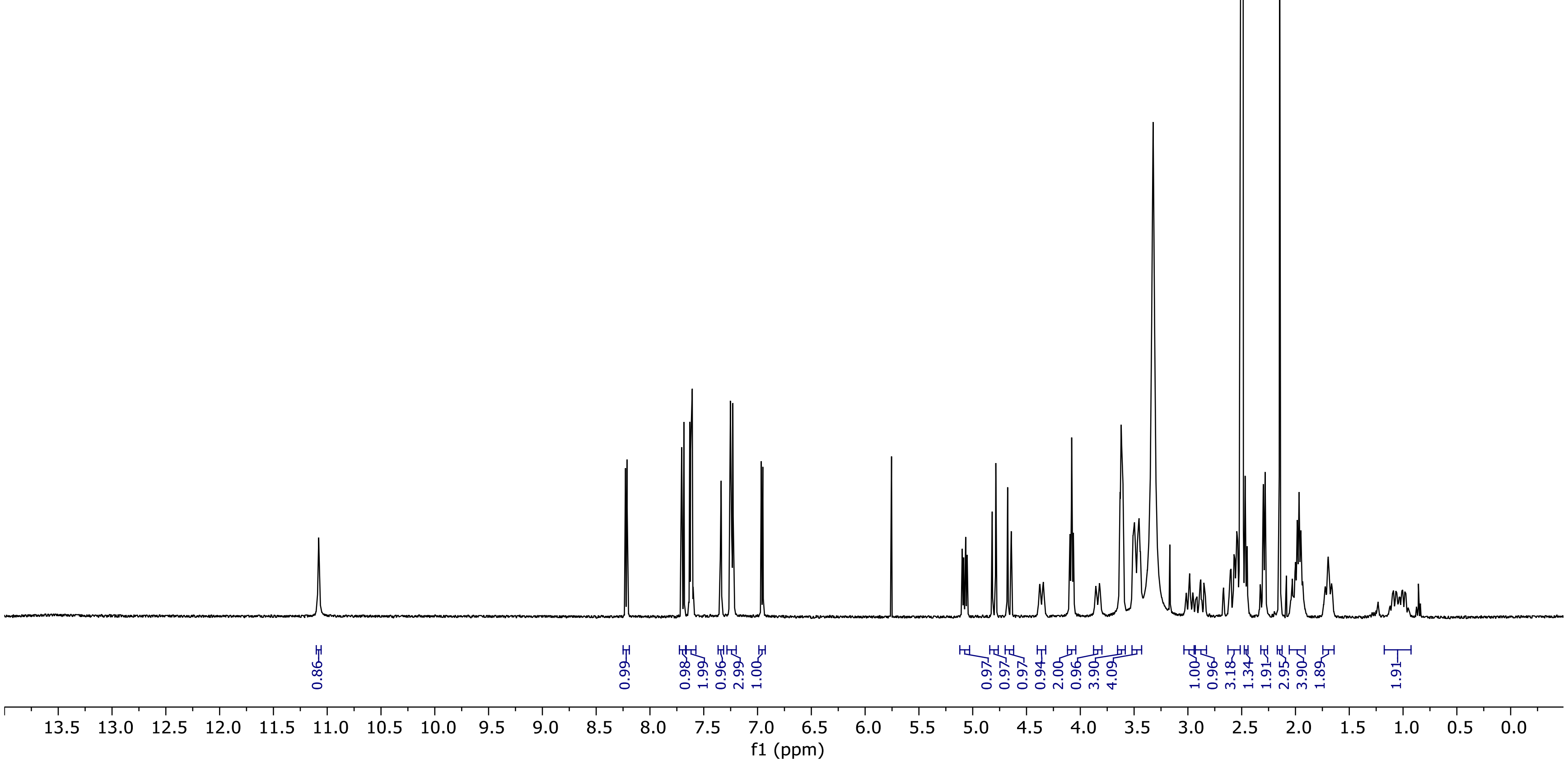

13C NMR (100.66 MHz, DMSO, 298.1 K)

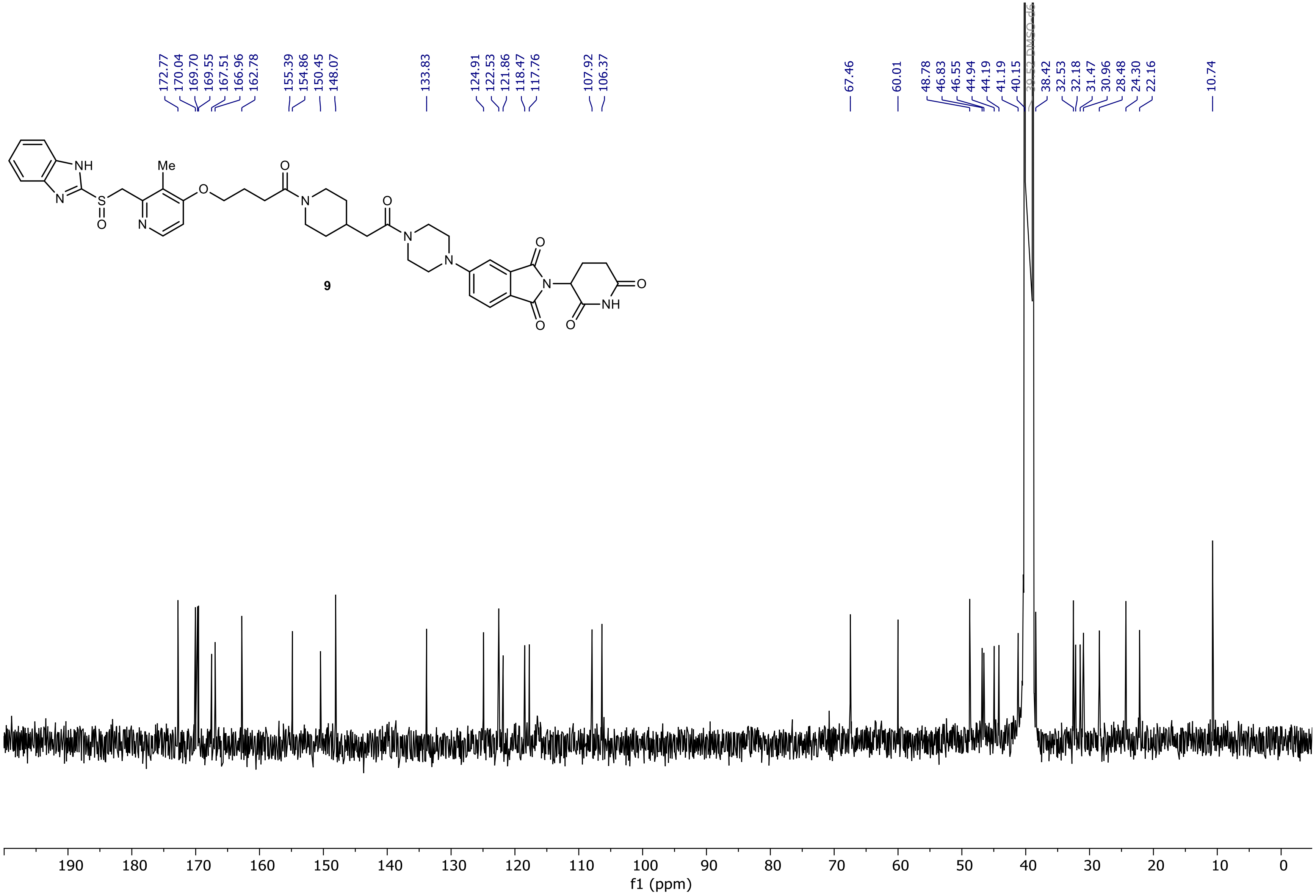

NR-5-075-A LC-MS_ELS  PDA - Total Absorbance Chromatogram

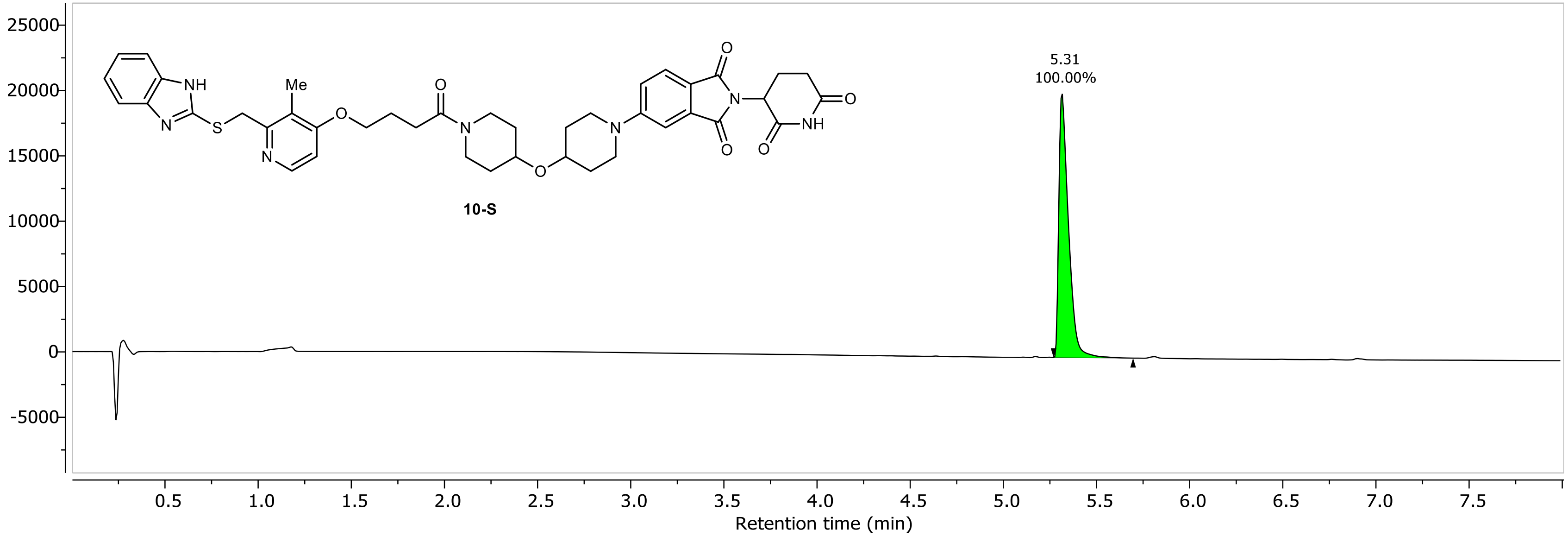

NR-5-075-A LC-MS_ELS DAD1A, Sig=254...Ref=360.0,100.0 Chromatogram

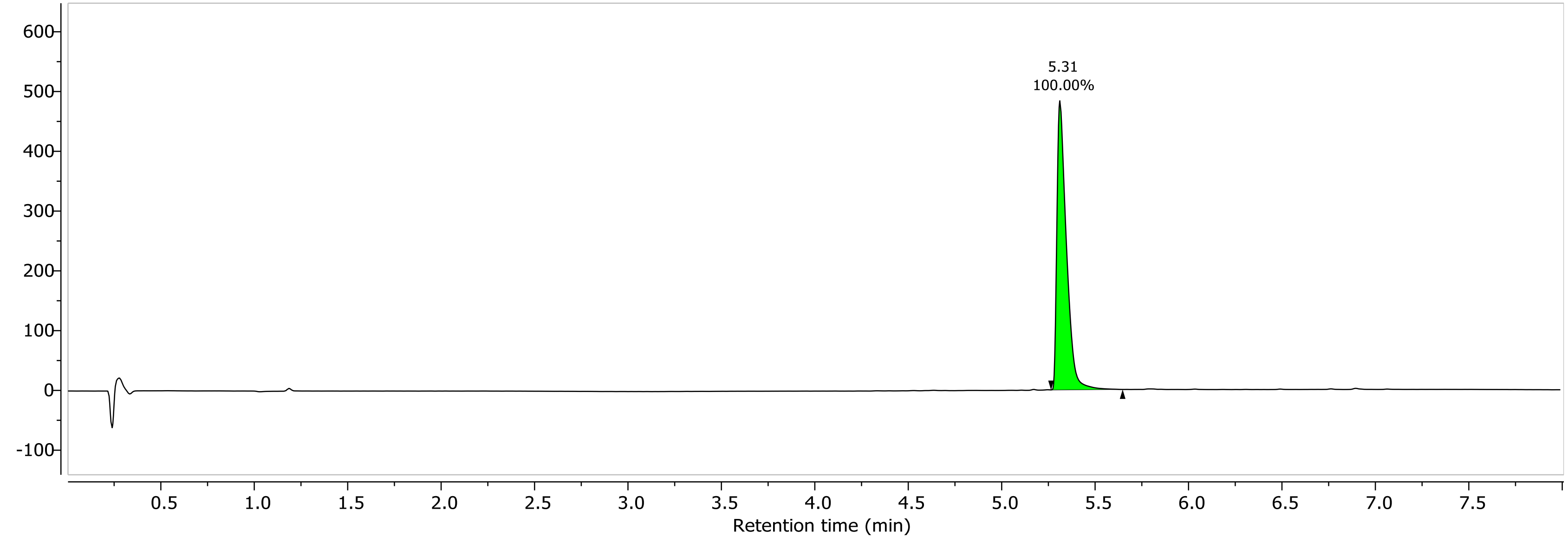

1H NMR (400.26 MHz, CDCl3, 298.1 K)

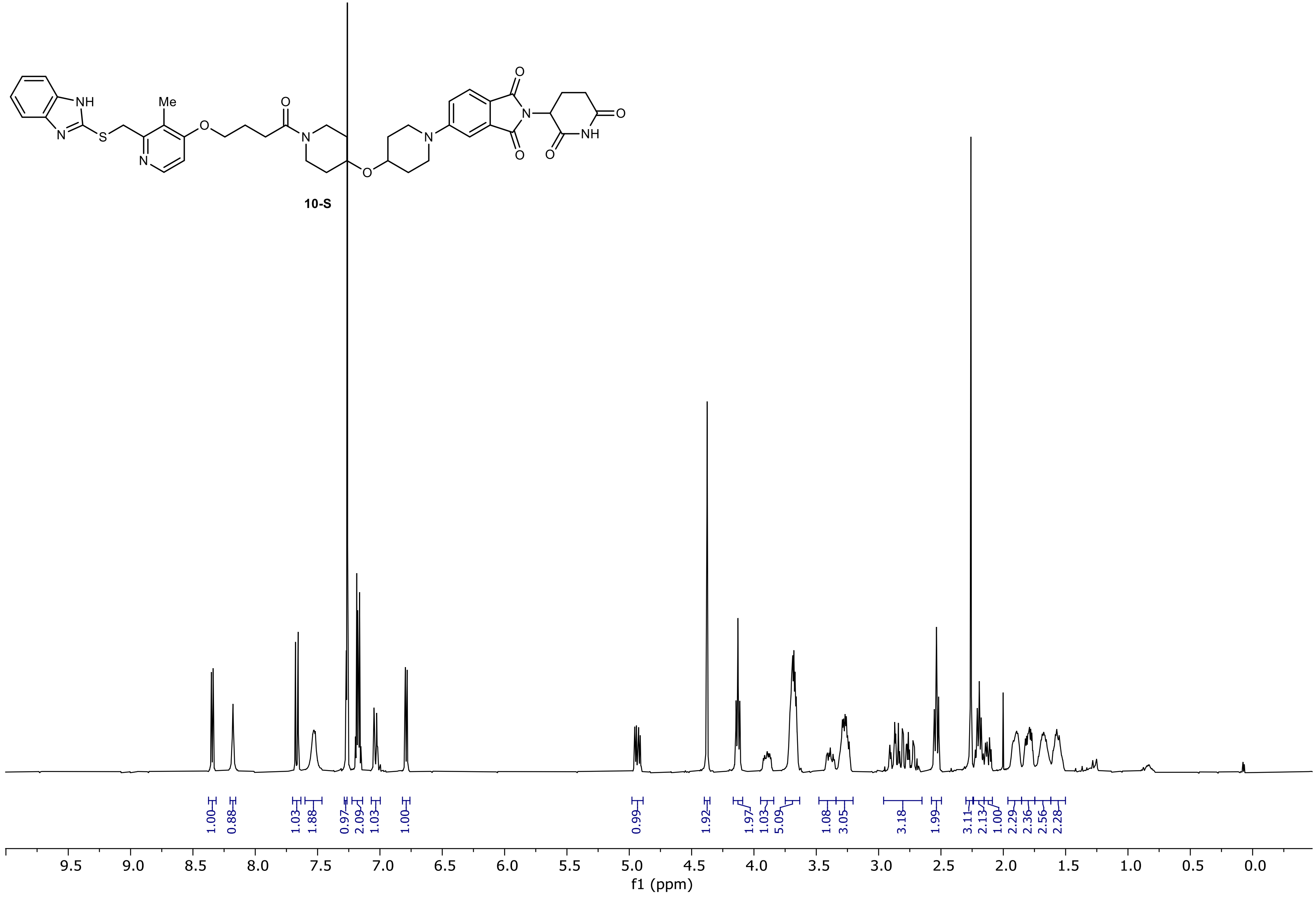

13C NMR (100.66 MHz, CDCl3, 298.1 K)

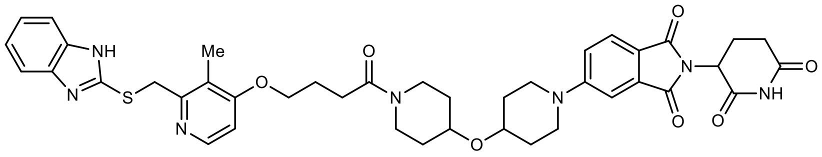

10-S

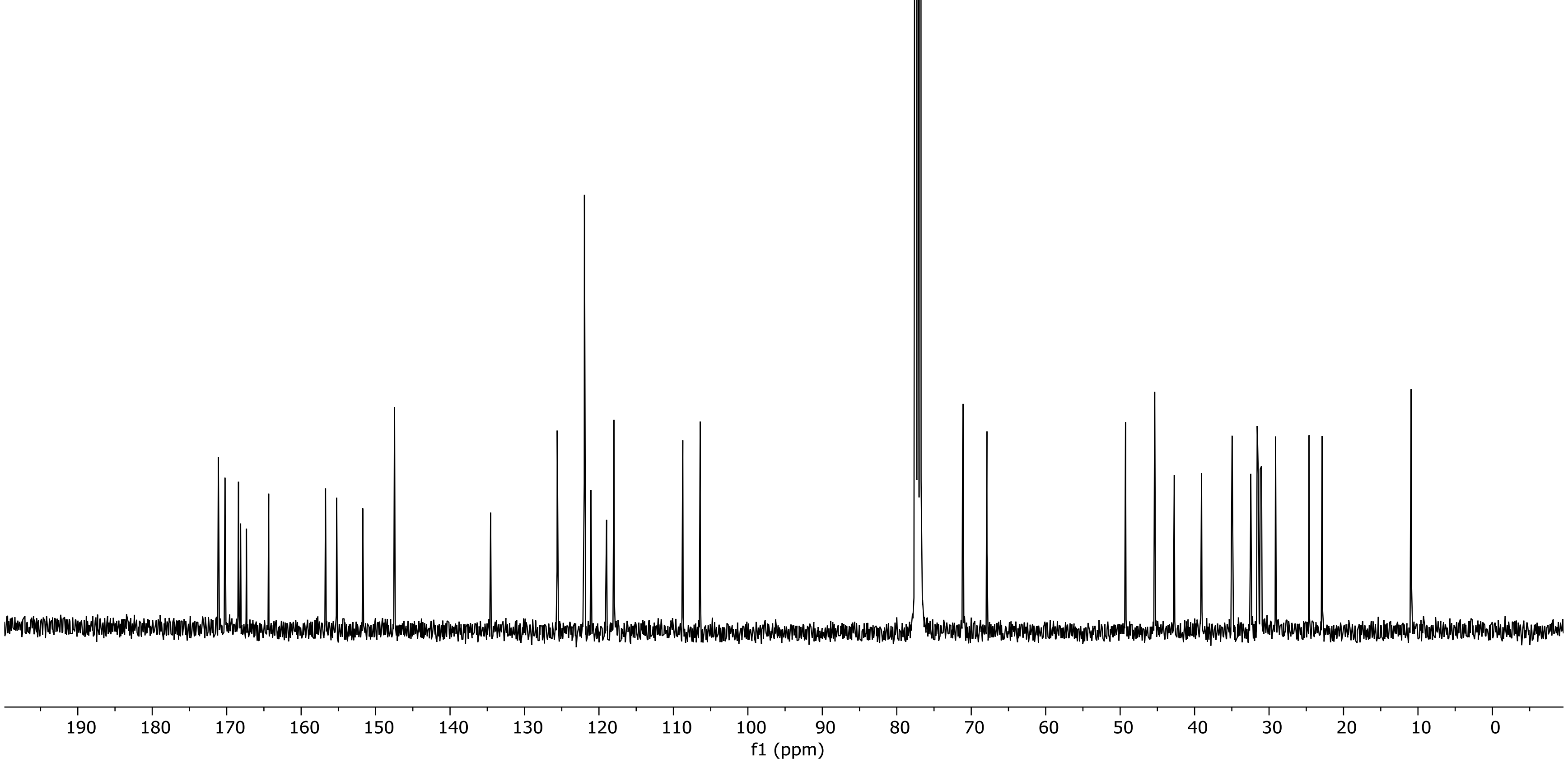

PDA - Total Absorbance Chromatogram

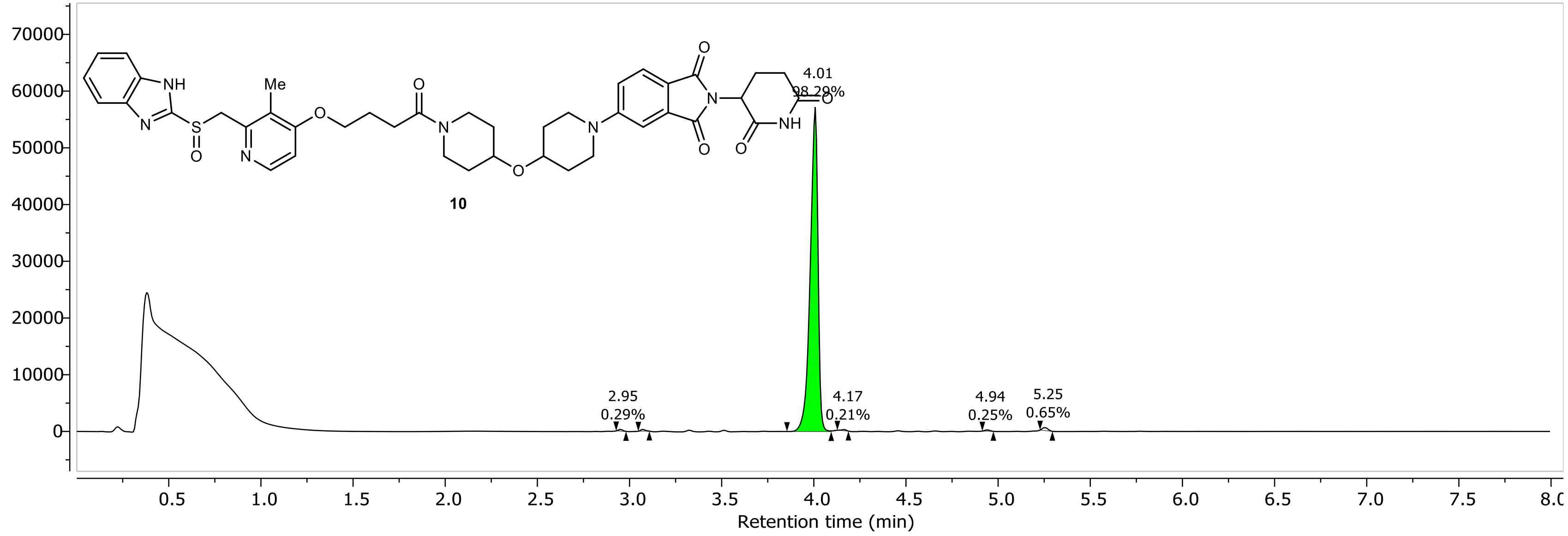

DAD1A, Sig=254...Ref=360.0,100.0 Chromatogram

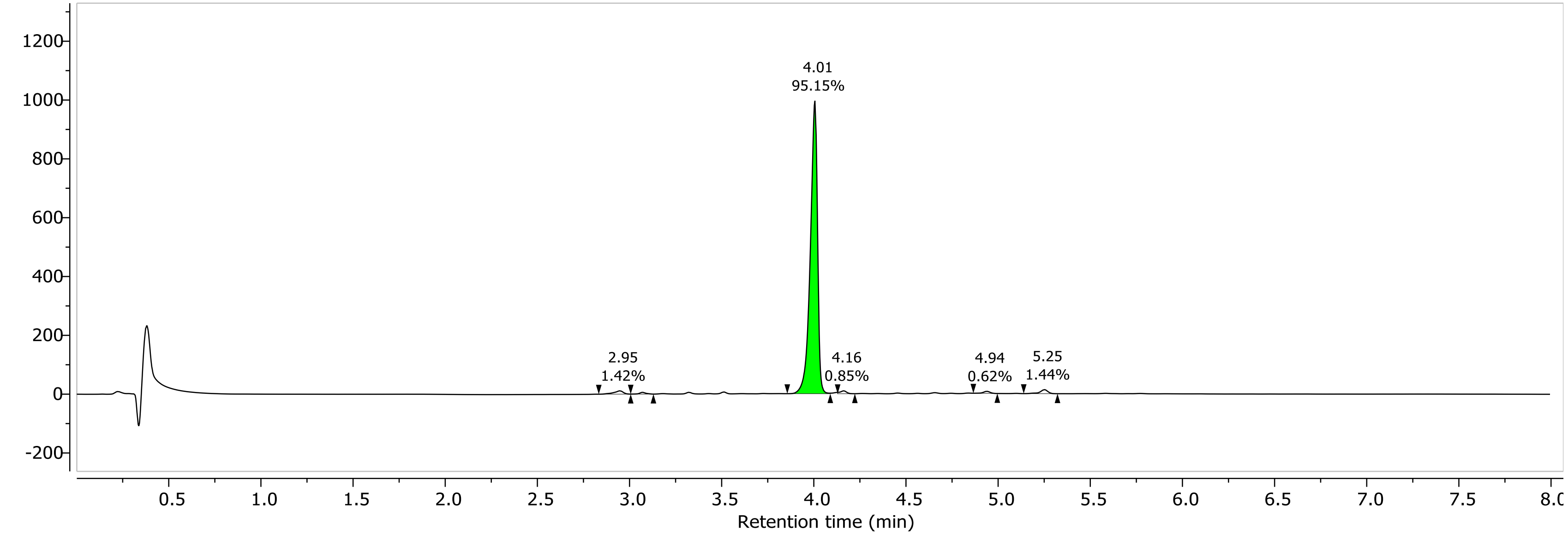

1H NMR (400.26 MHz, DMSO, 298.1 K)

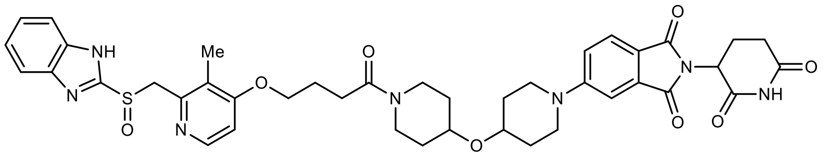

**10**

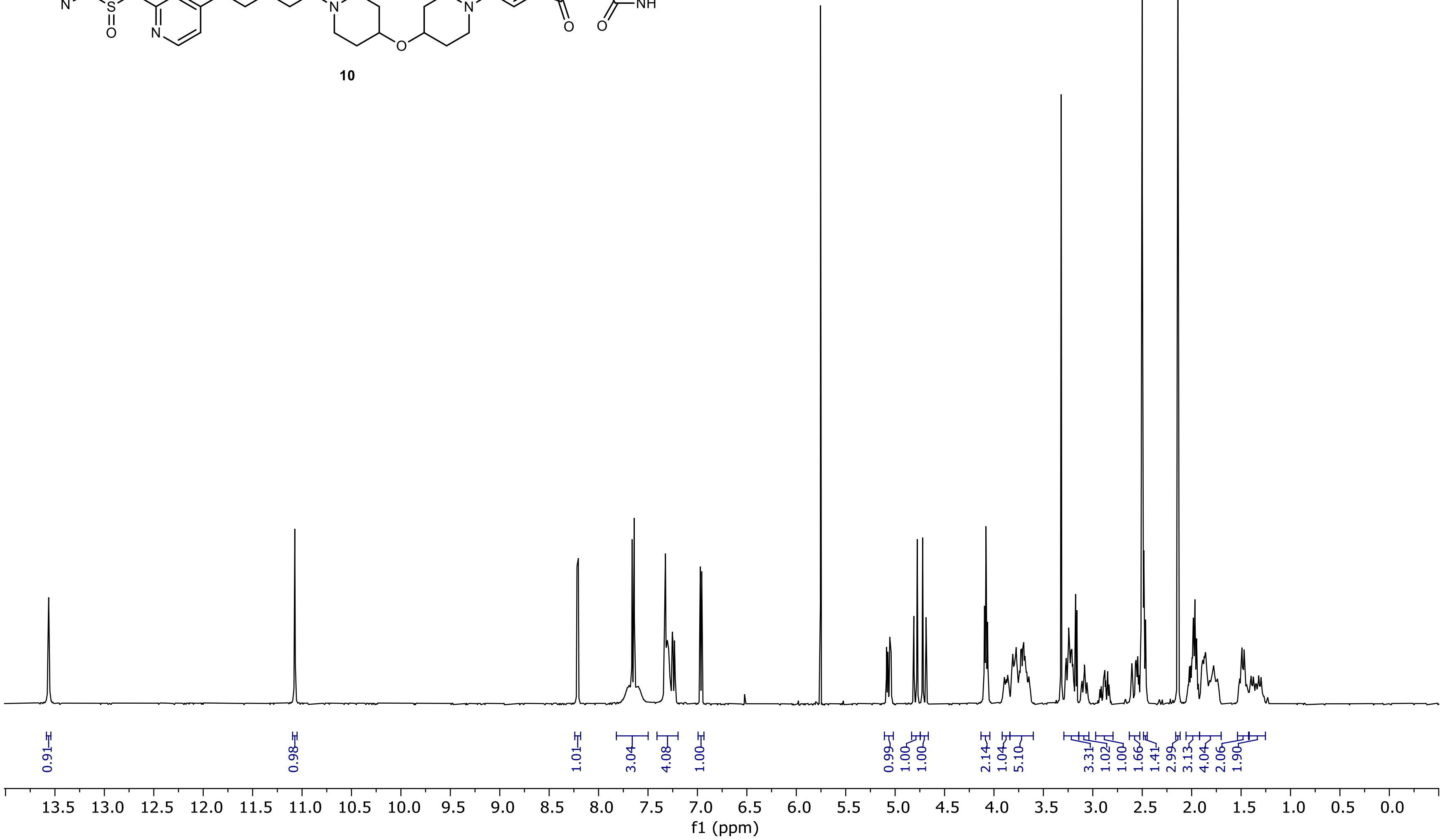

13C NMR (100.66 MHz, DMSO, 298.1 K)

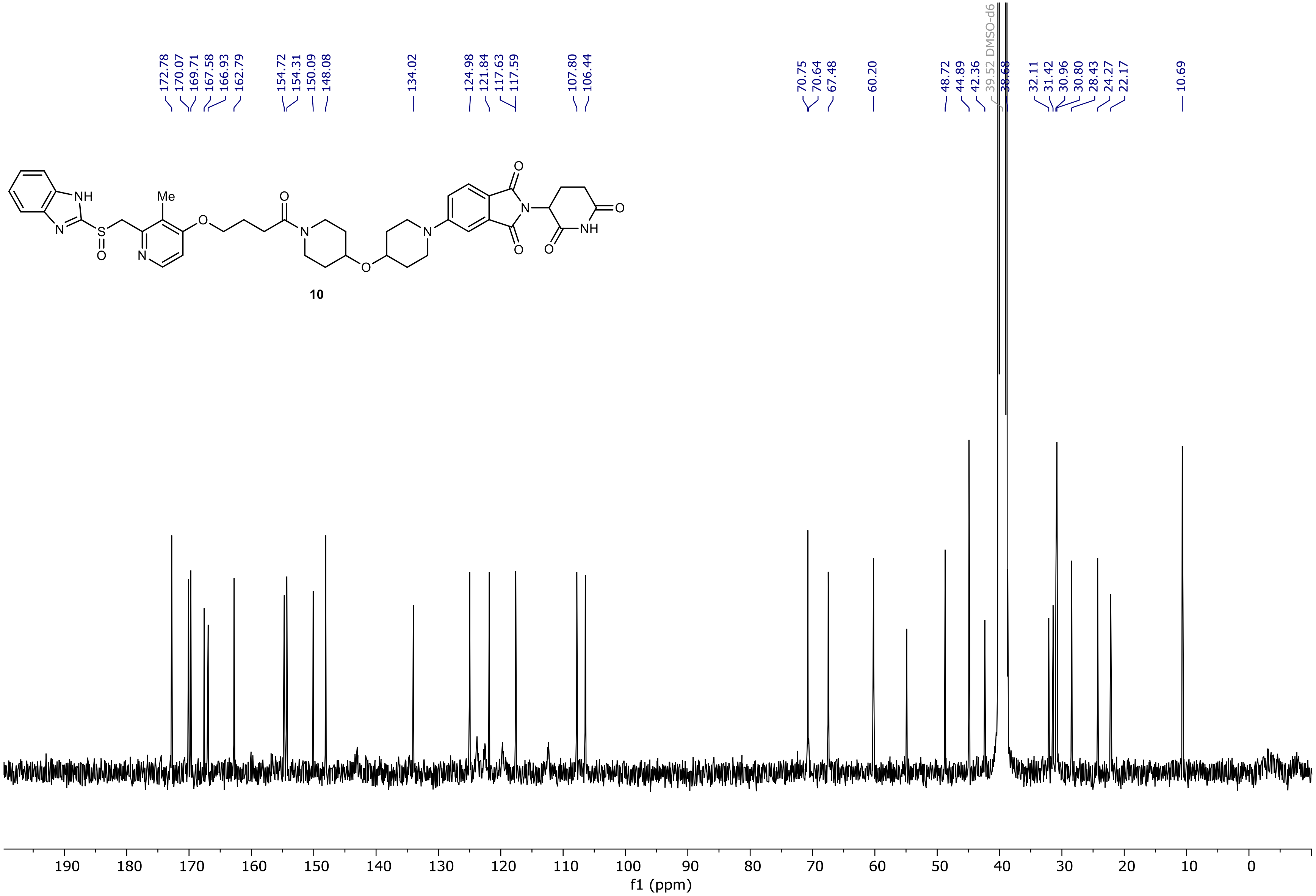

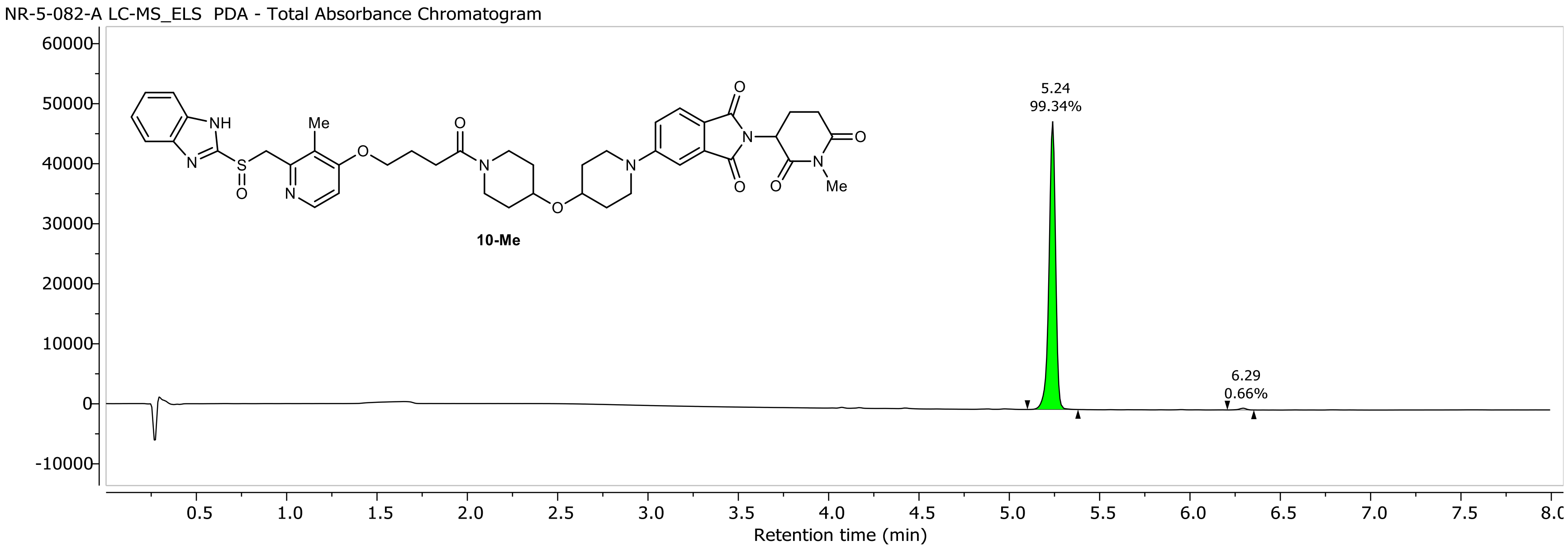
NR-5-082-A LC-MS_ELS PDA - Total Absorbance Chromatogram
10-Me
5.24
99.34%
6.29
0.66%
Retention time (min)

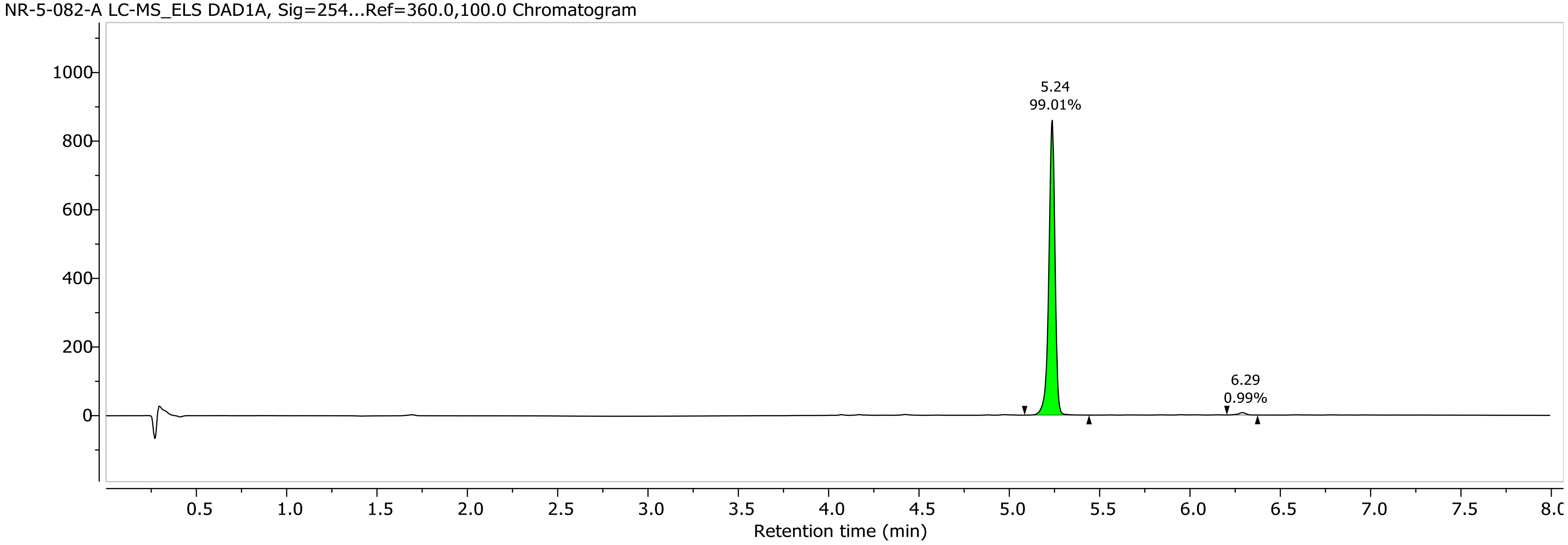
NR-5-082-A LC-MS_ELS DAD1A, Sig=254...Ref=360.0,100.0 Chromatogram
5.24
99.01%
6.29
0.99%
Retention time (min)

1H NMR (400.26 MHz, DMSO, 298.1 K)

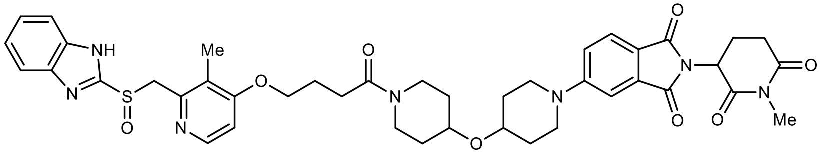

**10-Me**

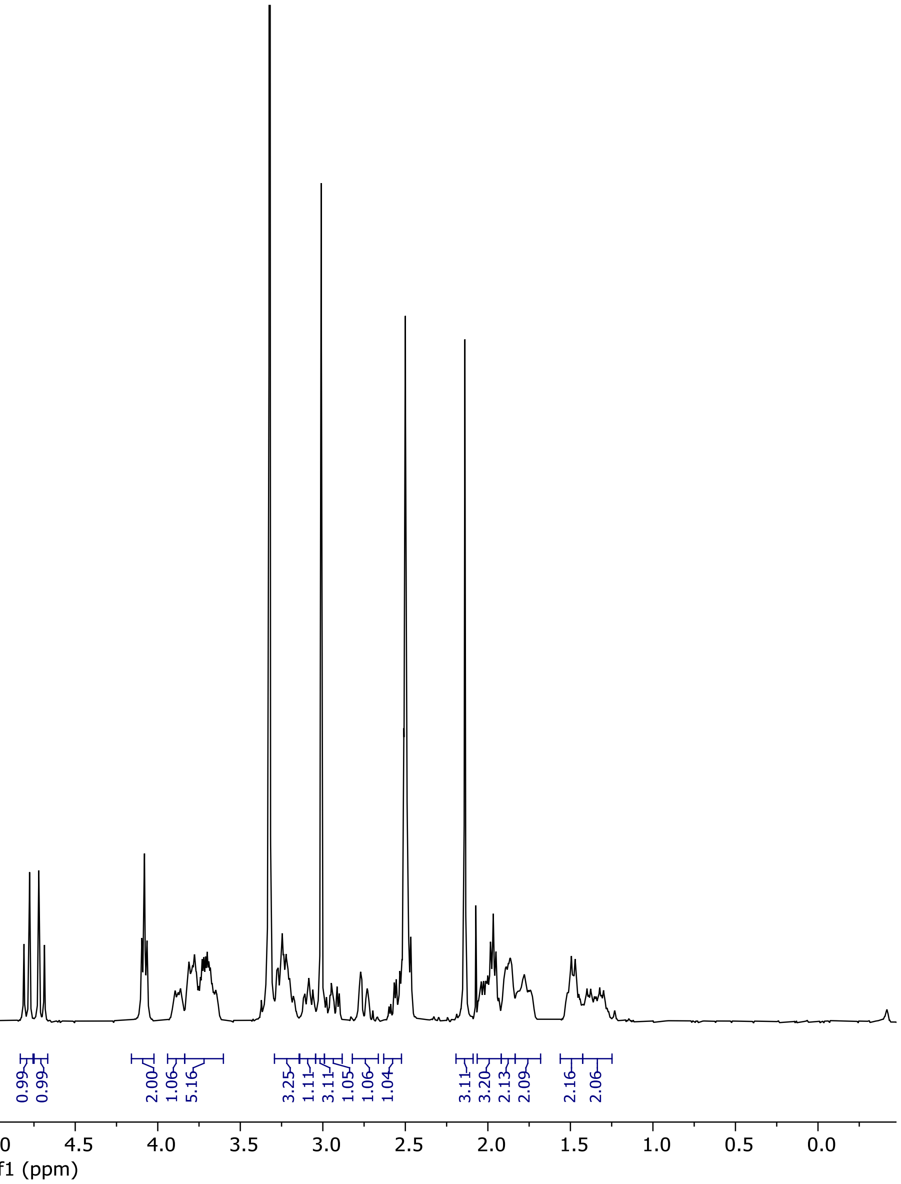

13C NMR (100.66 MHz, DMSO, 298.1 K)

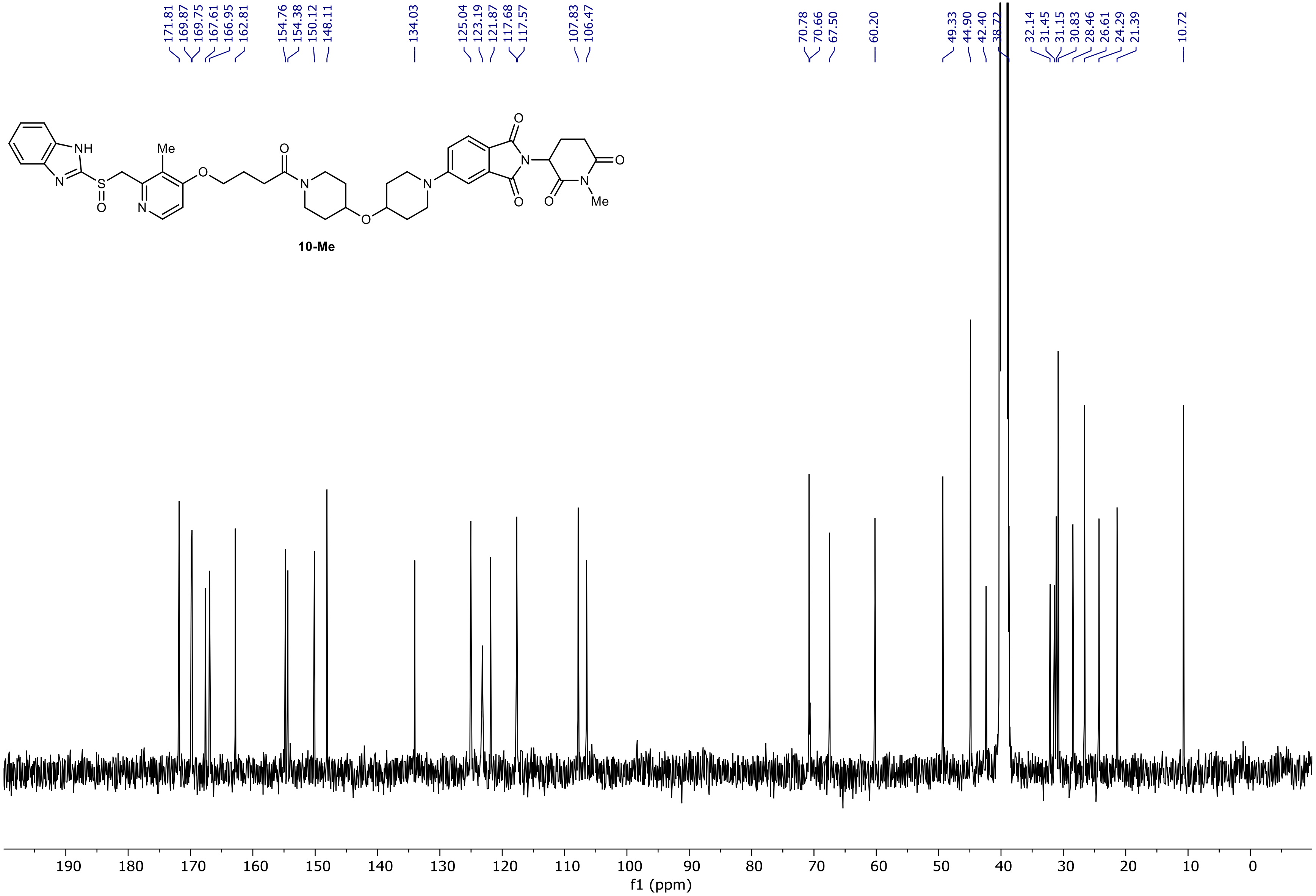

NR-5-124-A LC-MS_ELS  PDA - Total Absorbance Chromatogram

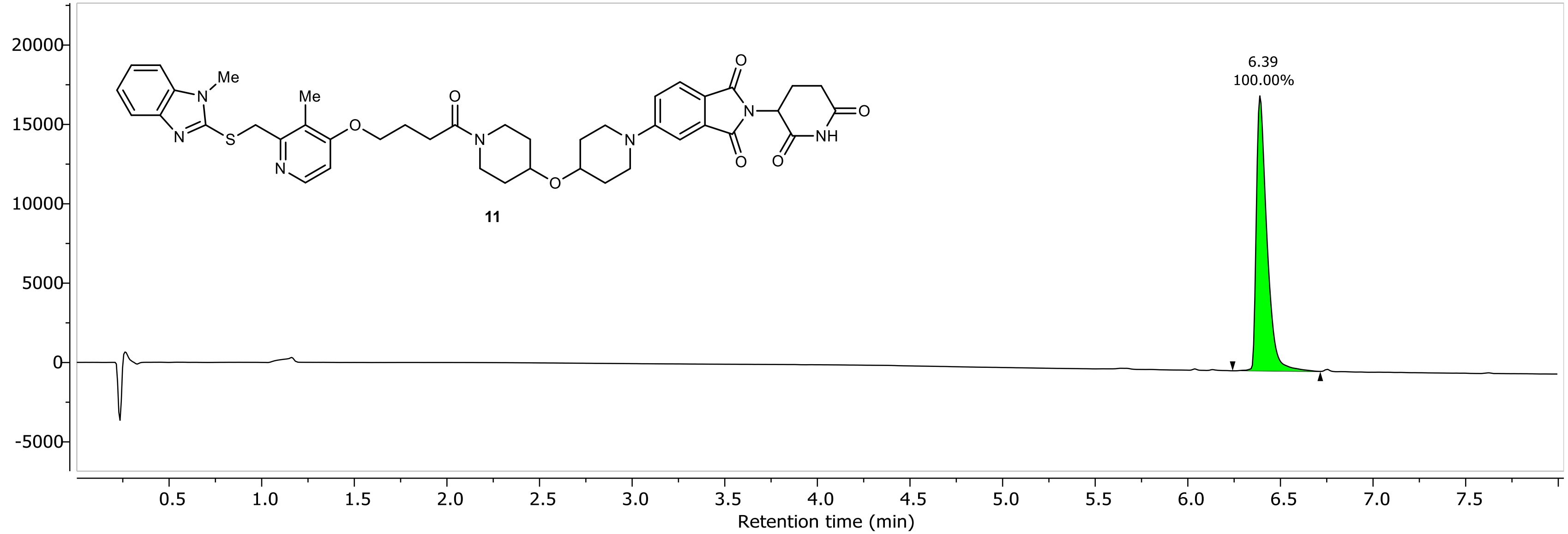

NR-5-124-A LC-MS_ELS DAD1A, Sig=254...Ref=360.0,100.0 Chromatogram

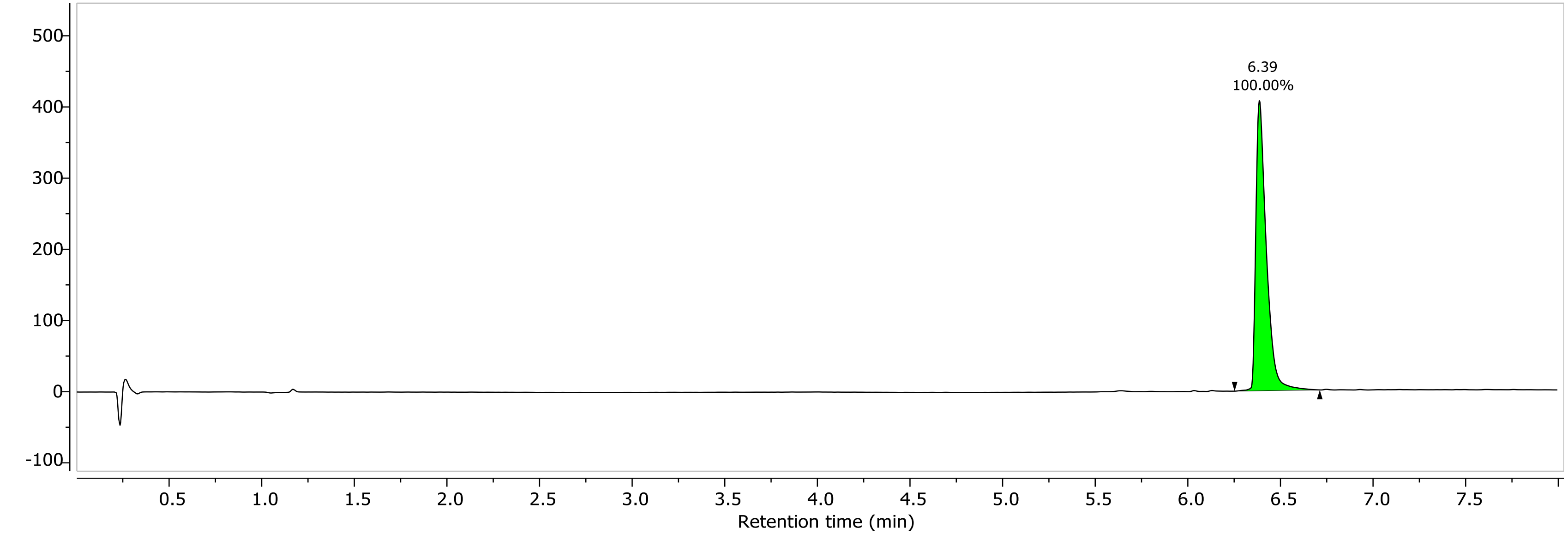

1H NMR (400.26 MHz, CDCl3, 298.1 K)

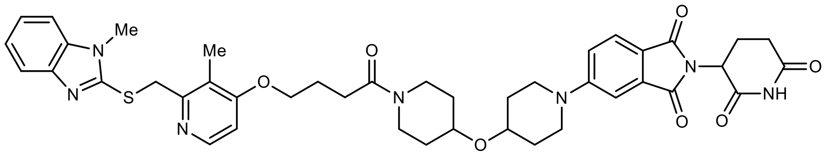

**11**

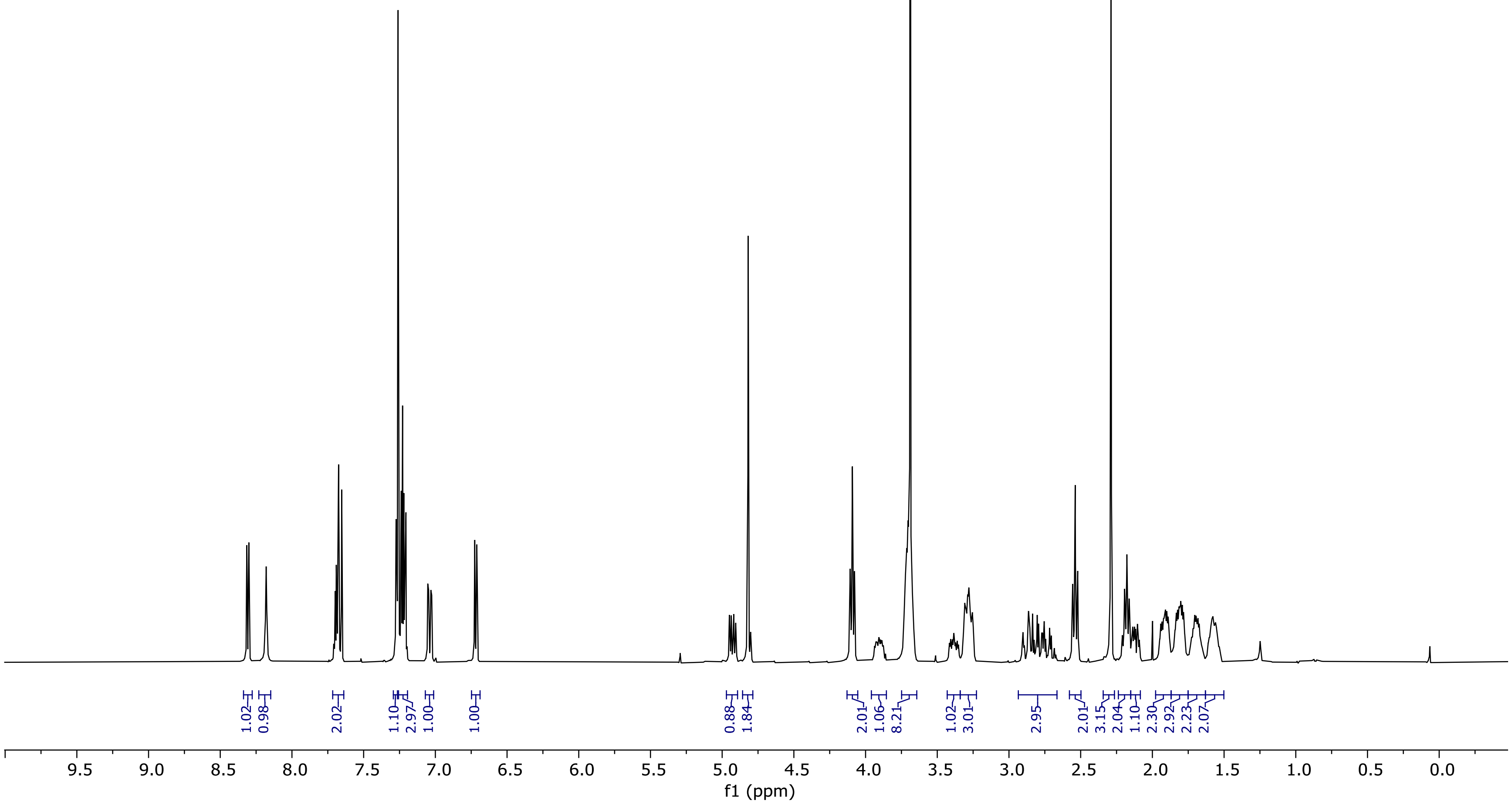

13C NMR (100.66 MHz, CDCl3, 298.1 K)

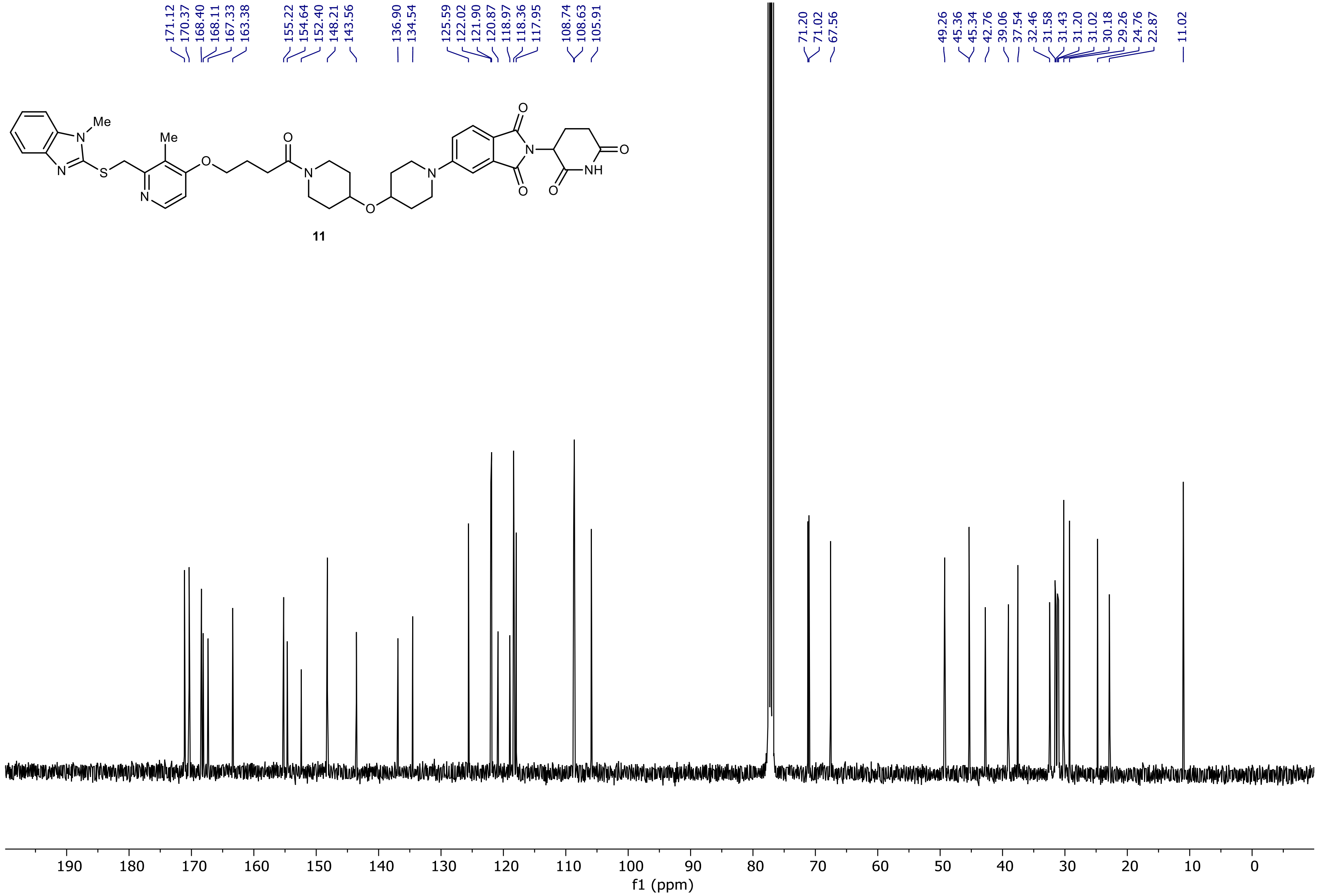

NR-6-050-B LC-MS_ELS PDA - Total Absorbance Chromatogram

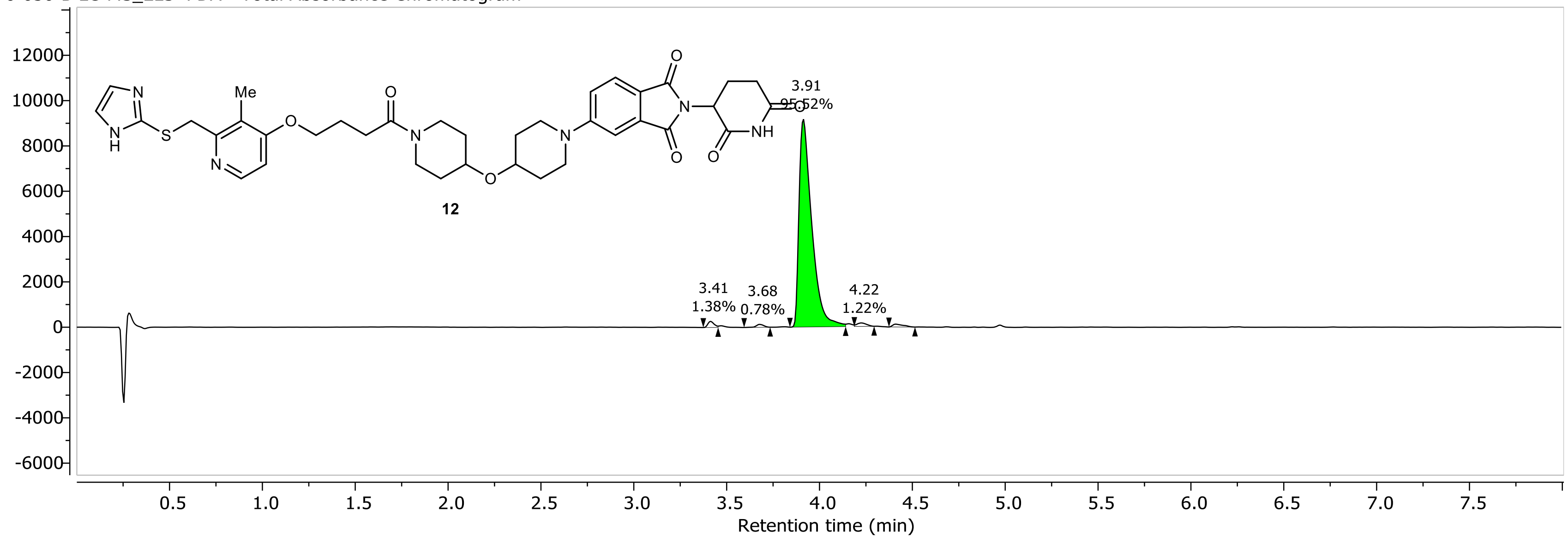

NR-6-050-B LC-MS_ELS DAD1A, Sig=254...Ref=360.0,100.0 Chromatogram

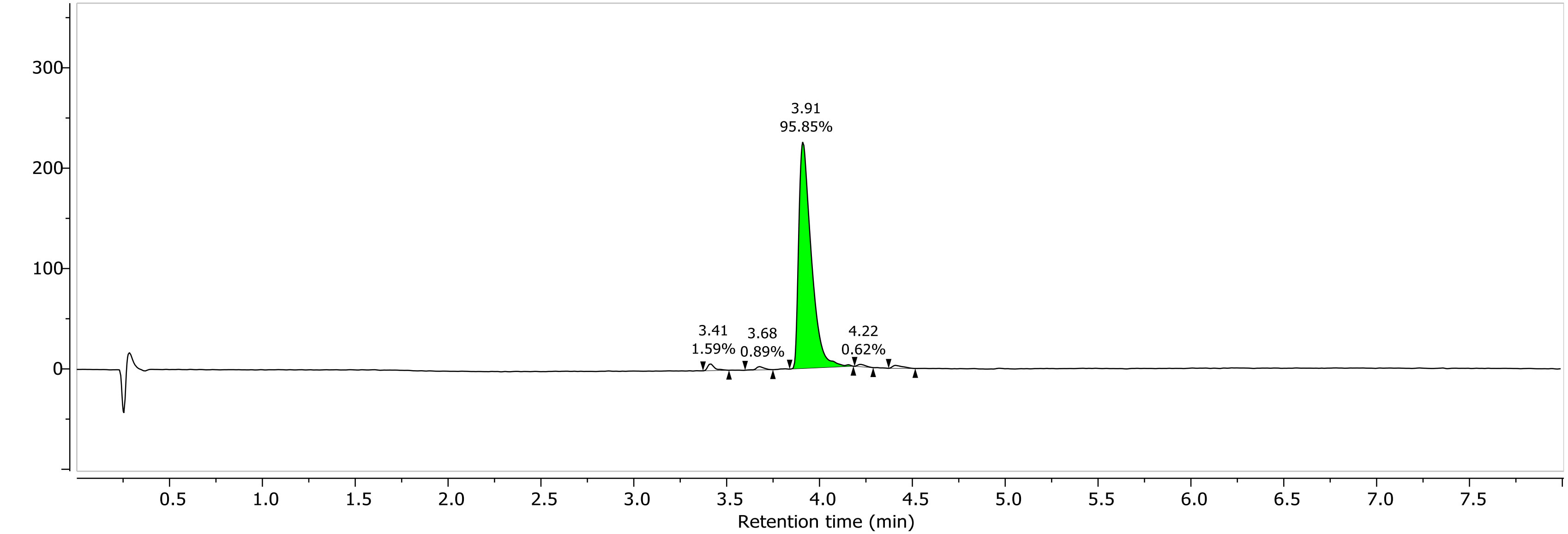

1H NMR (400.26 MHz, CDCl3, 298.1 K)

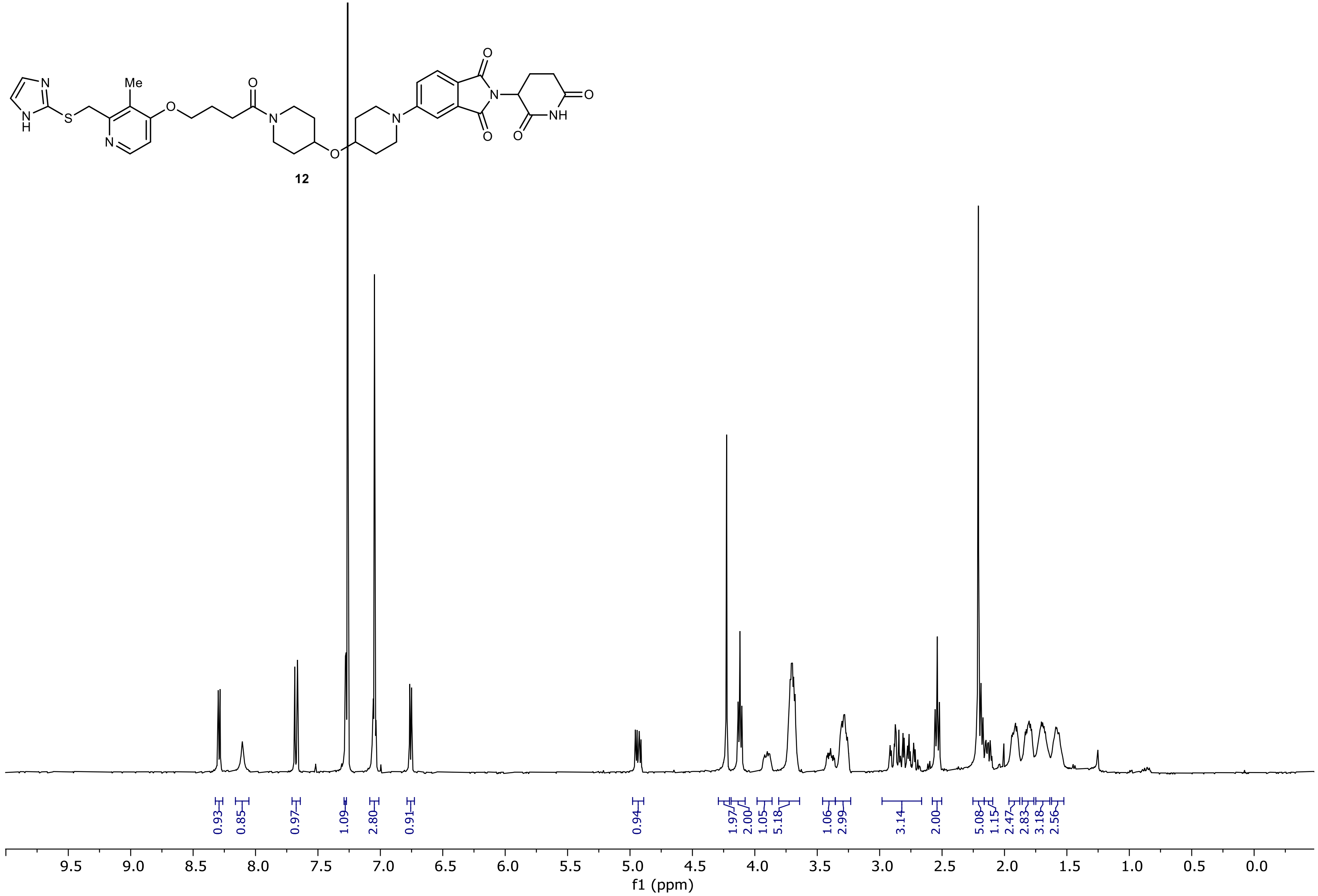

13C NMR (100.66 MHz, CDCl3, 298.1 K)

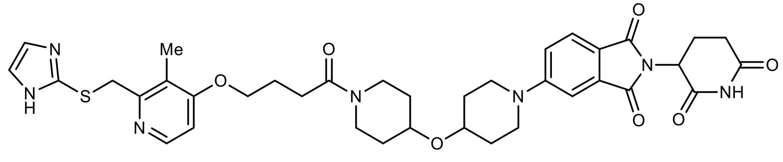

12

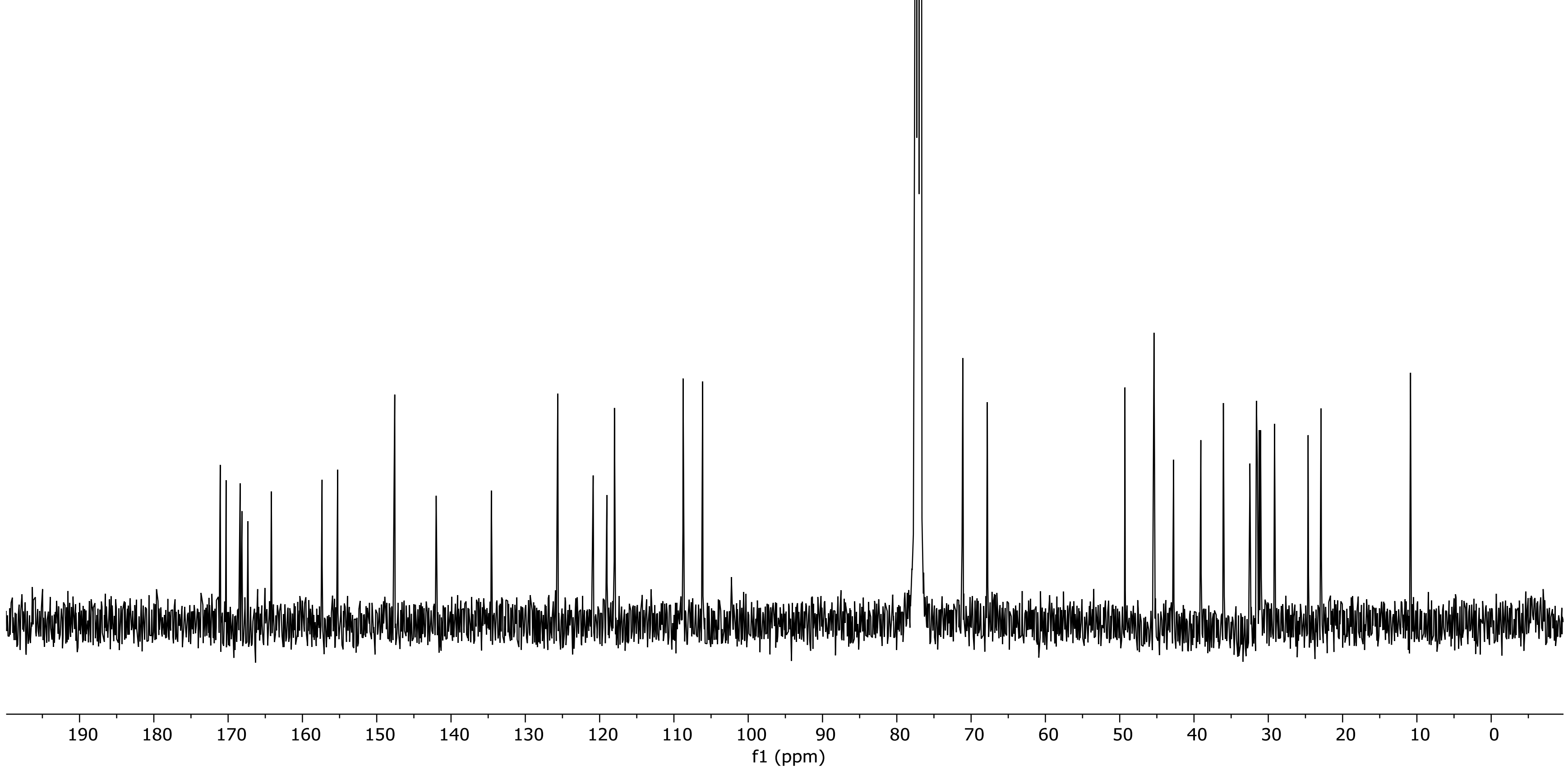

NR-6-053-A LC-MS_ELS  PDA - Total Absorbance Chromatogram

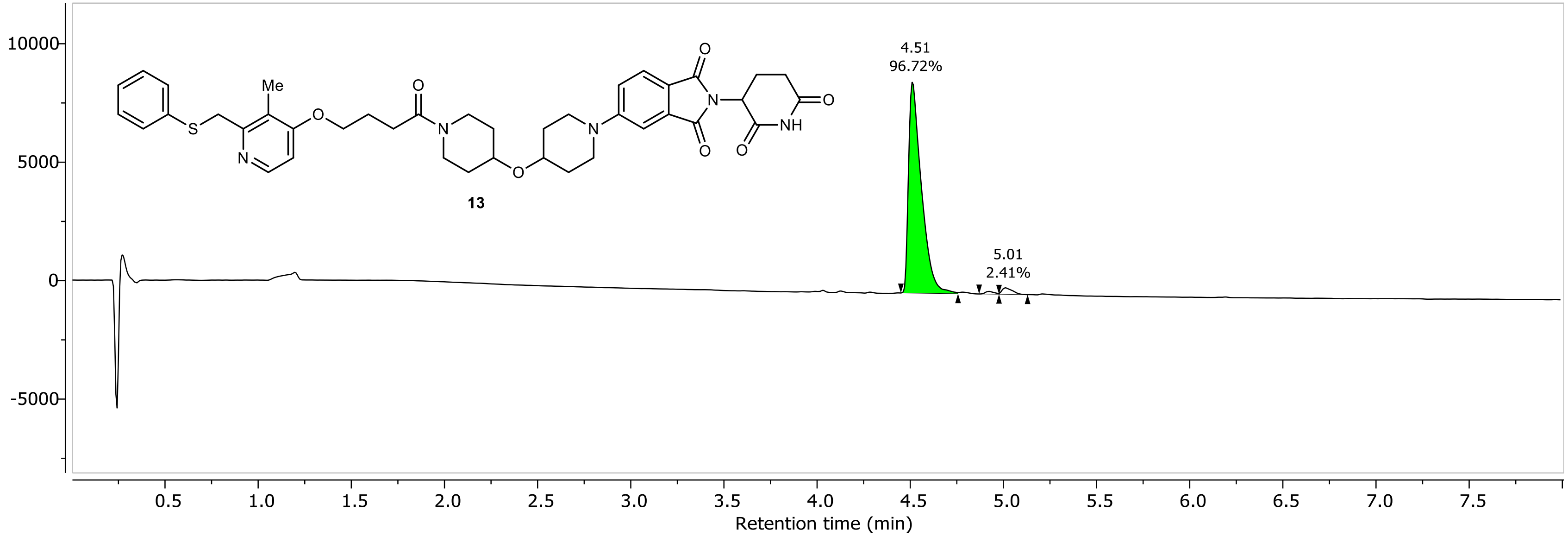

NR-6-053-A LC-MS_ELS DAD1A, Sig=254...Ref=360.0,100.0 Chromatogram

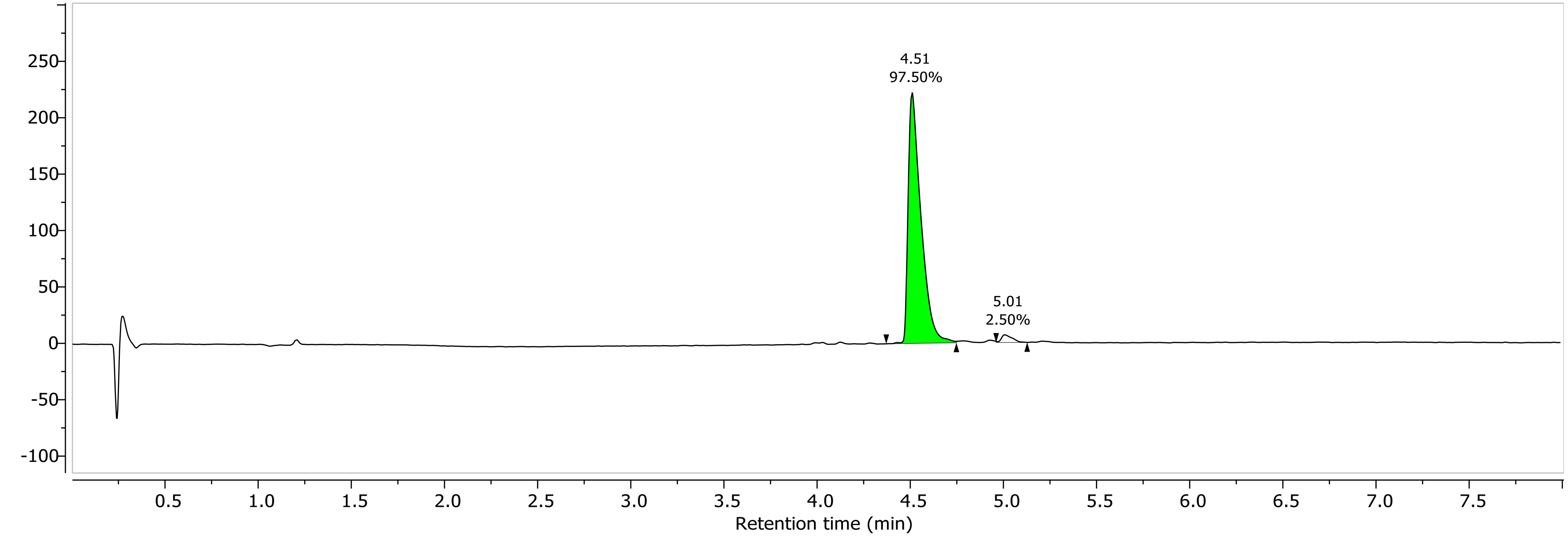

1H NMR (400.26 MHz, CDCl3, 298.1 K)

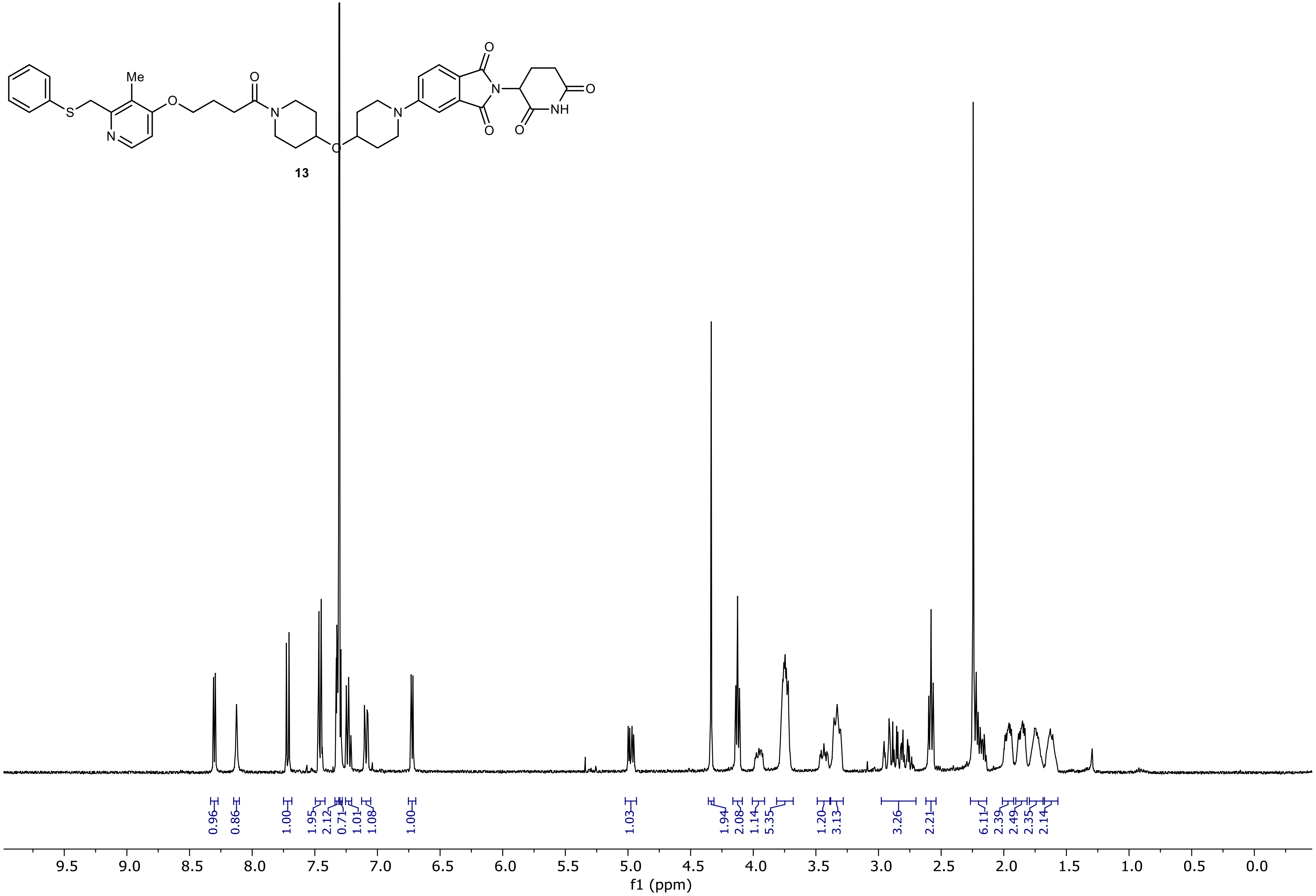

13C NMR (100.66 MHz, CDCl3, 298.1 K)

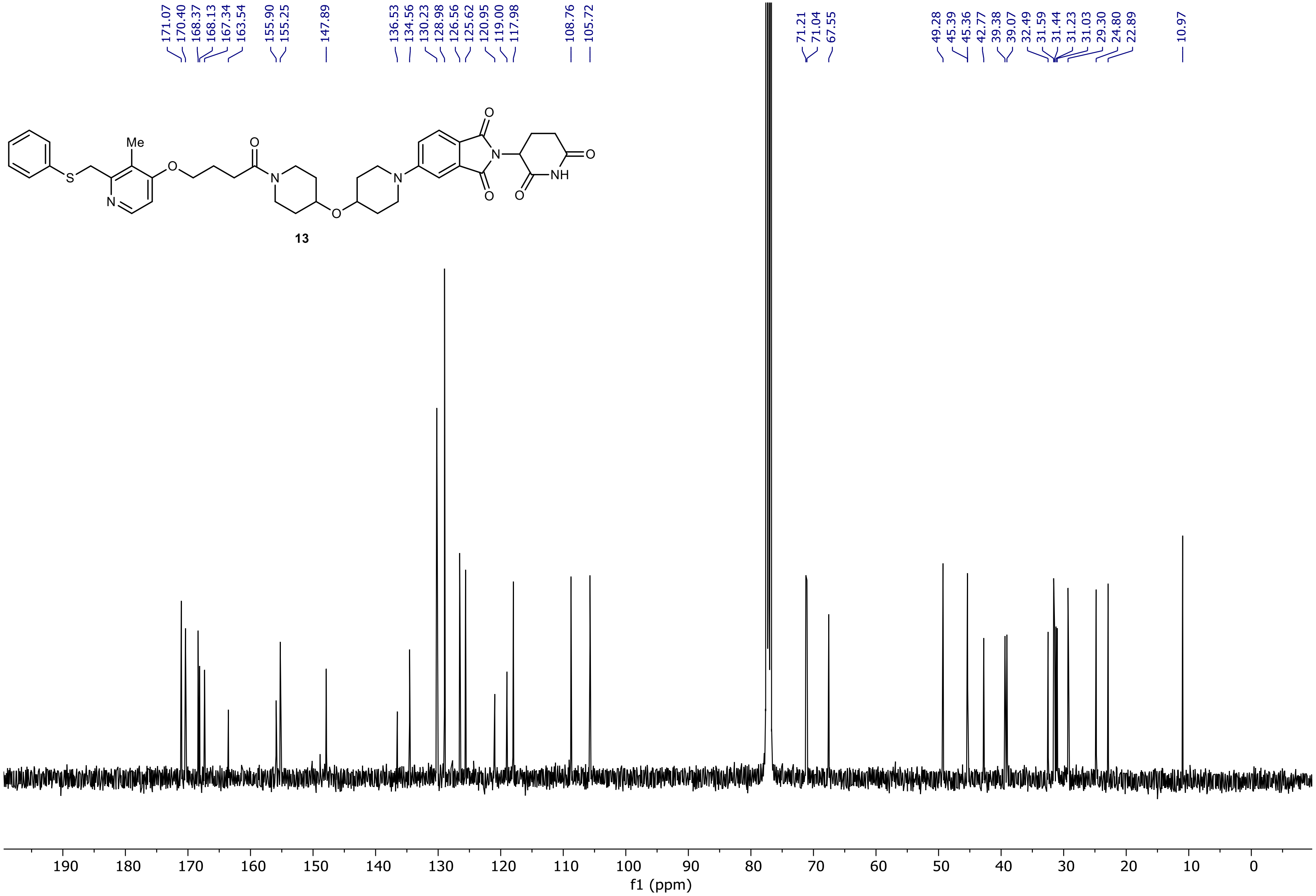

NR-6-002-A LC-MS_ELS  PDA - Total Absorbance Chromatogram

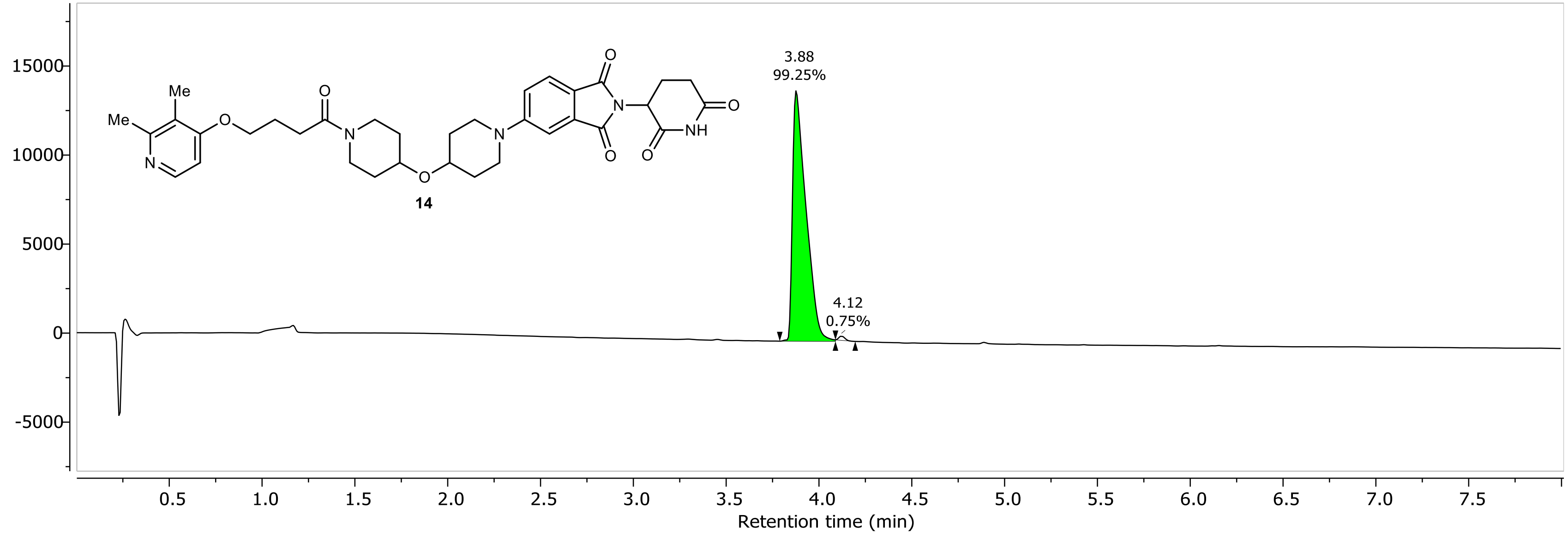

NR-6-002-A LC-MS_ELS DAD1A, Sig=254...Ref=360.0,100.0 Chromatogram

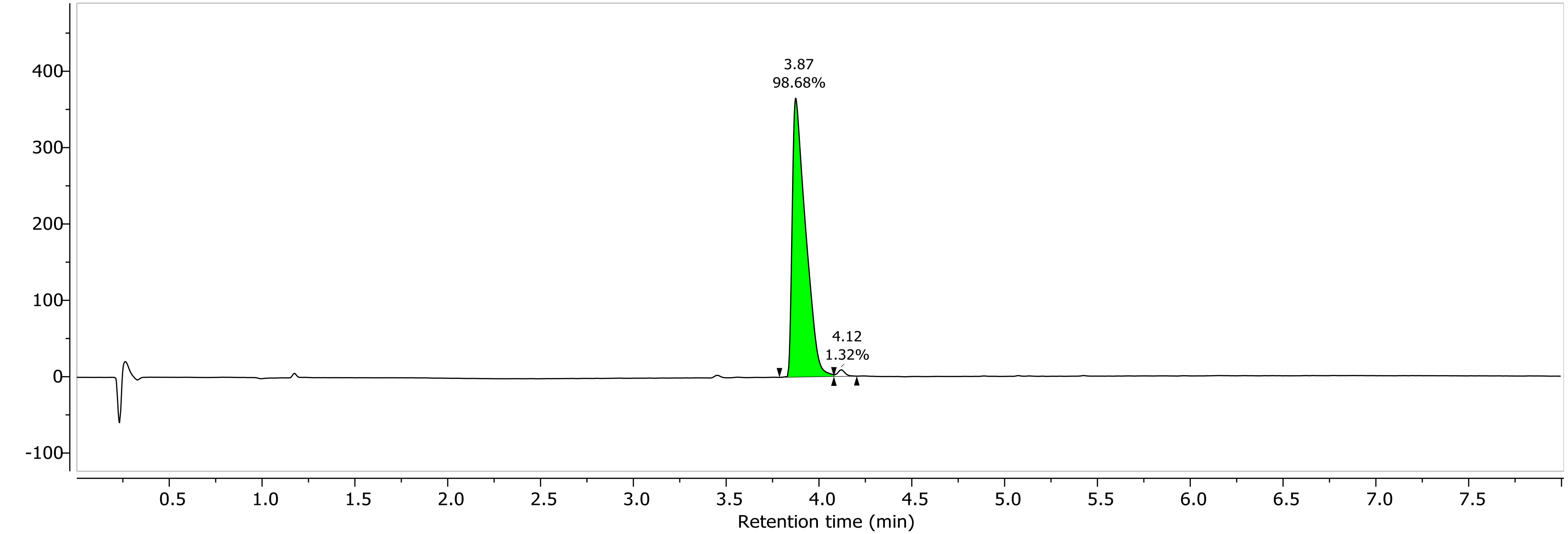

1H NMR (400.26 MHz, CDCl3, 298.1 K)

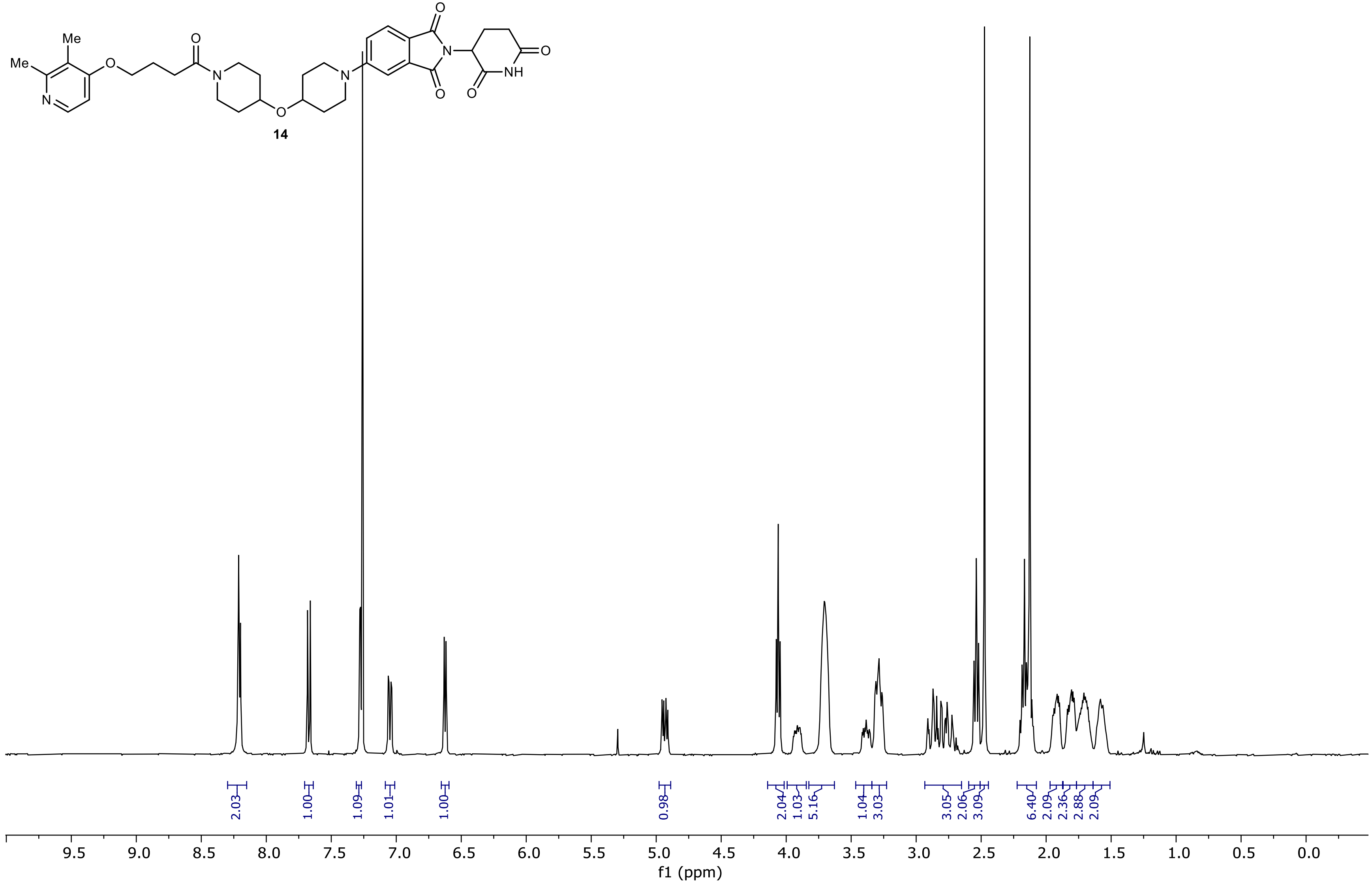

13C NMR (100.66 MHz, CDCl3, 298.1 K)

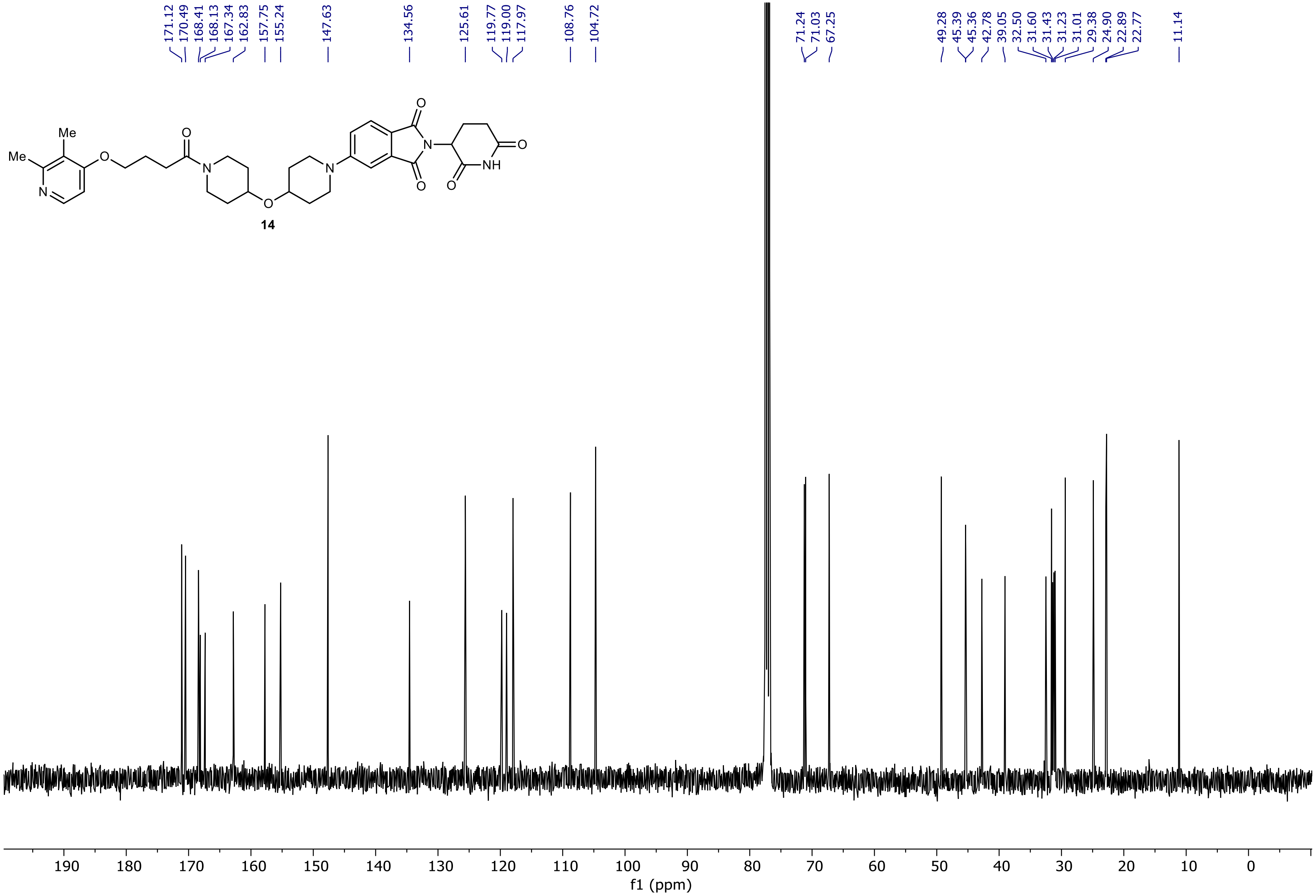

NR-5-119-A LC-MS_ELS  PDA - Total Absorbance Chromatogram

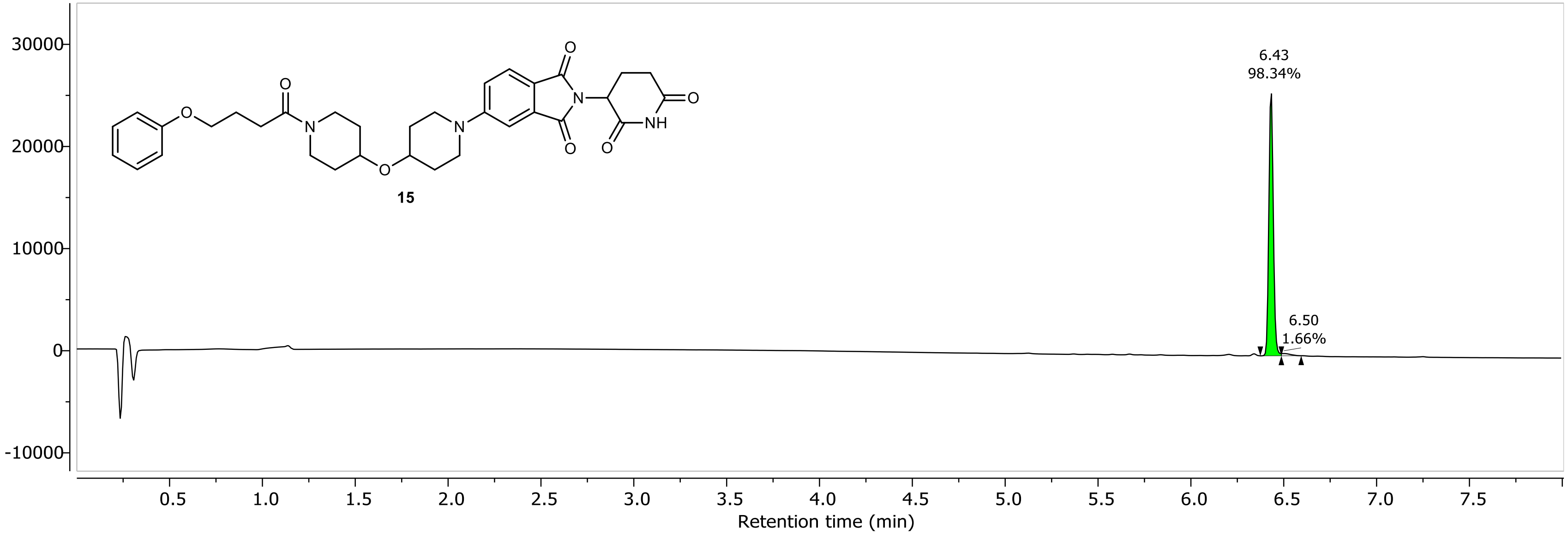

NR-5-119-A LC-MS_ELS DAD1A, Sig=254...Ref=360.0,100.0 Chromatogram

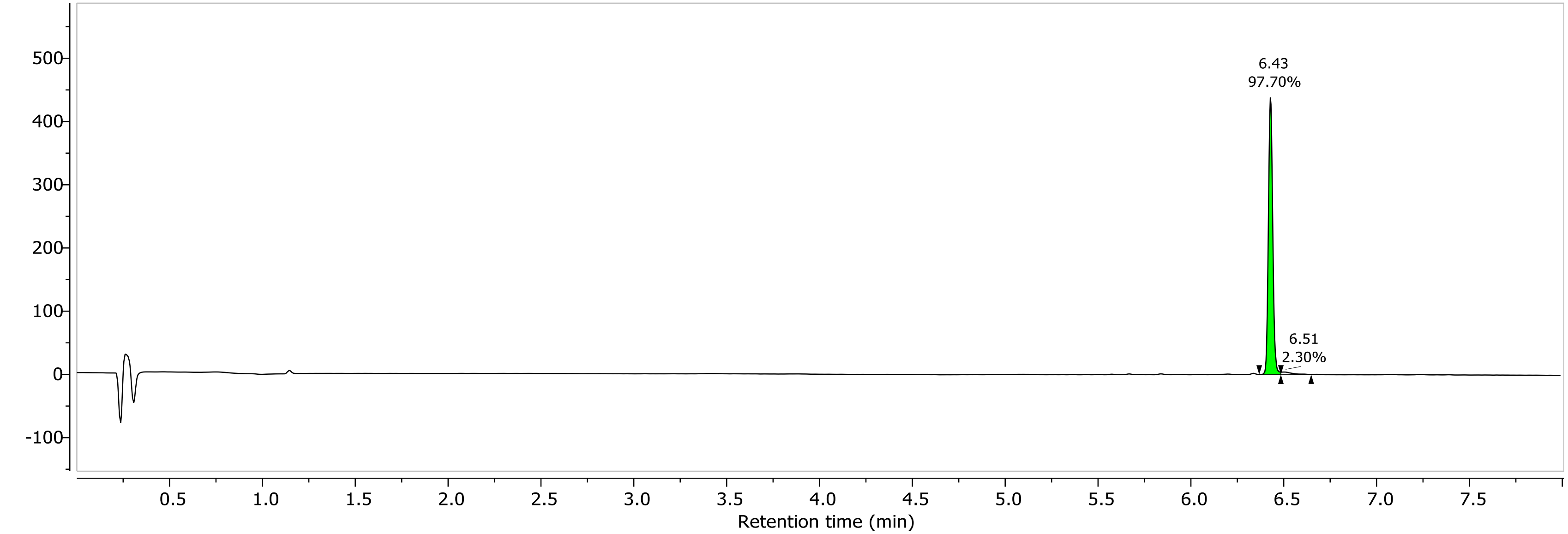

1H NMR (400.26 MHz, CDCl3, 298.1 K)

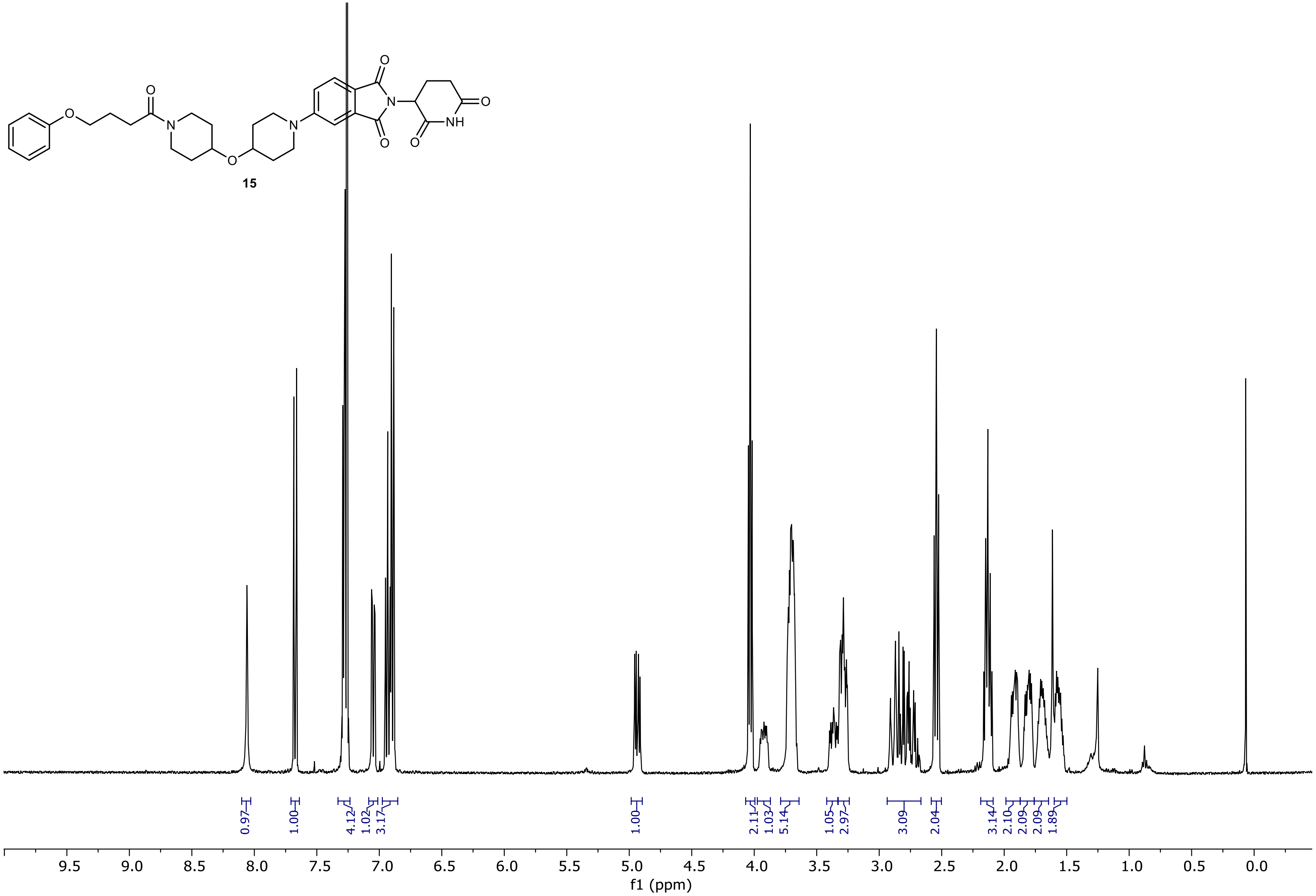

13C NMR (100.66 MHz, CDCl3, 298.1 K)

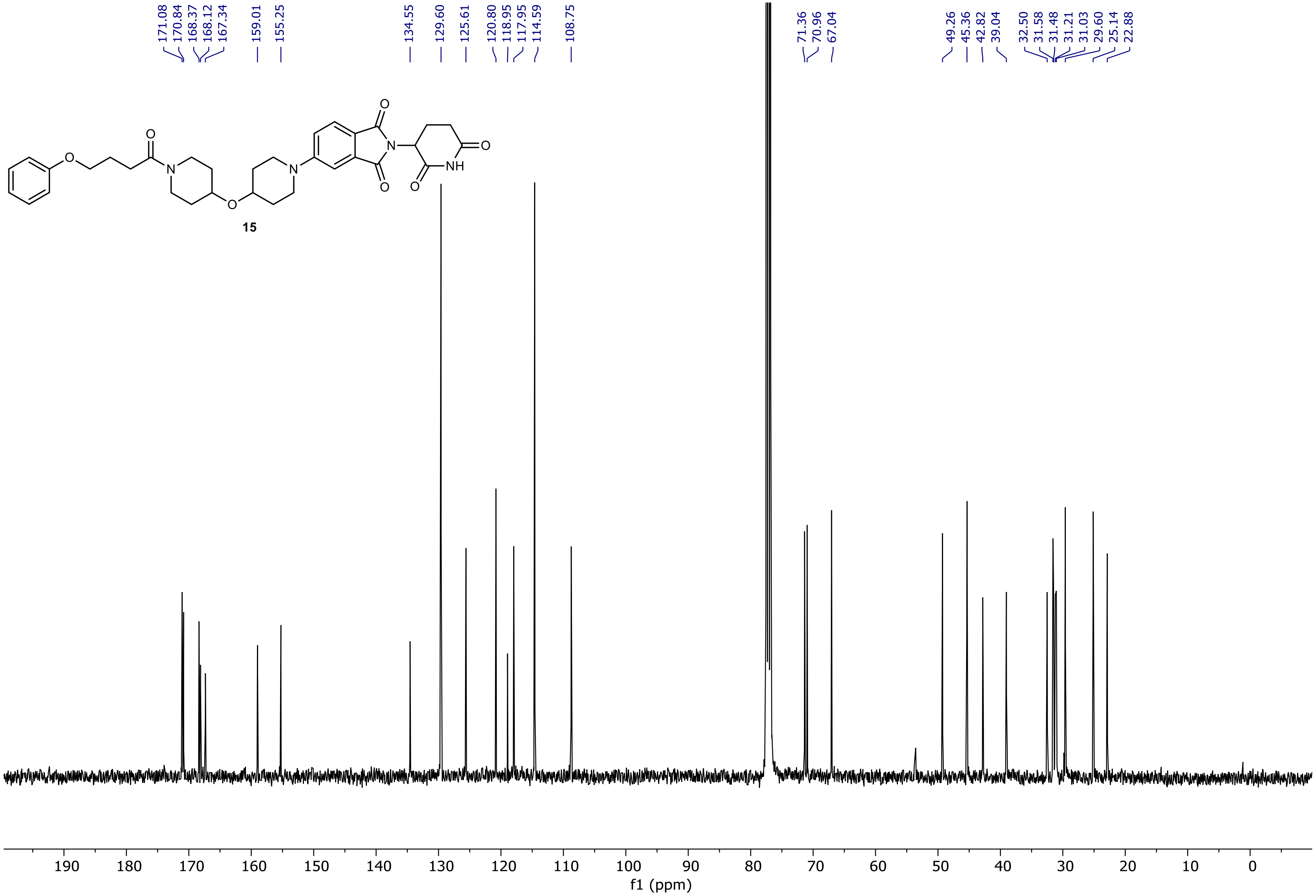

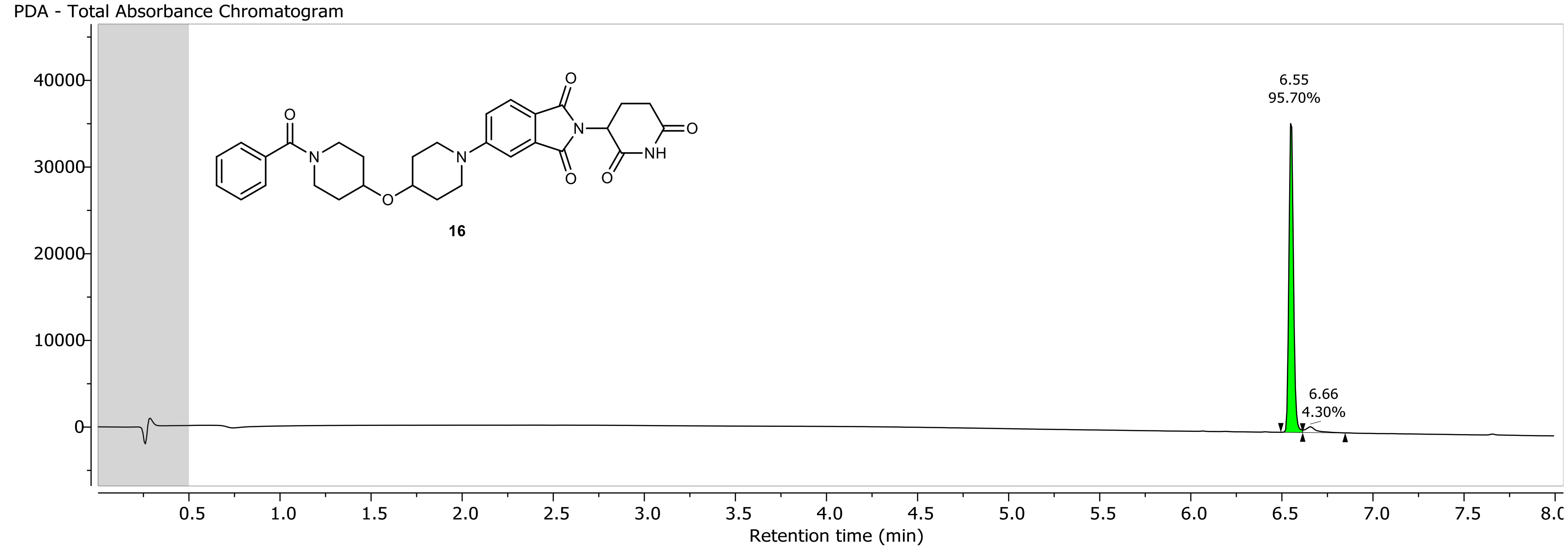
PDA - Total Absorbance Chromatogram
16
6.55
95.70%
6.66
4.30%
40000
30000
20000
10000
0
0.5 1.0 1.5 2.0 2.5 3.0 3.5 4.0 4.5 5.0 5.5 6.0 6.5 7.0 7.5 8.0
Retention time (min)

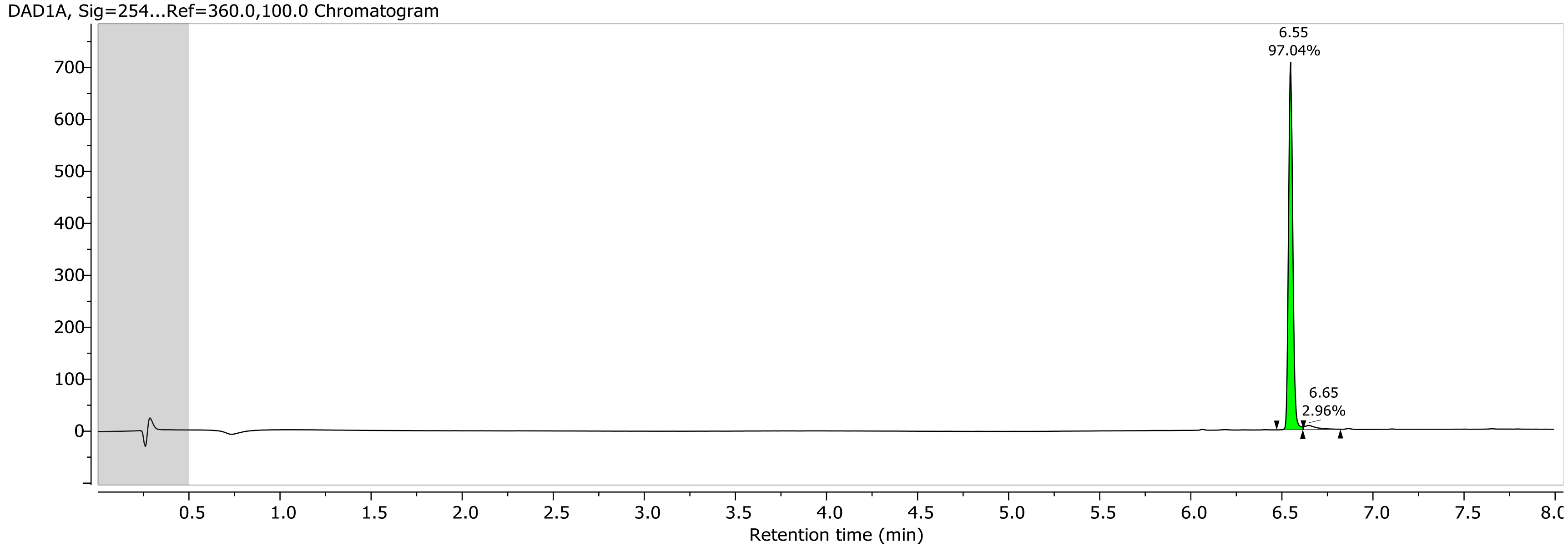
DAD1A, Sig=254...Ref=360.0,100.0 Chromatogram
6.55
97.04%
6.65
2.96%
700
600
500
400
300
200
100
0
0.5 1.0 1.5 2.0 2.5 3.0 3.5 4.0 4.5 5.0 5.5 6.0 6.5 7.0 7.5 8.0
Retention time (min)

1H NMR (400.26 MHz, CDCl3, 298.1 K)

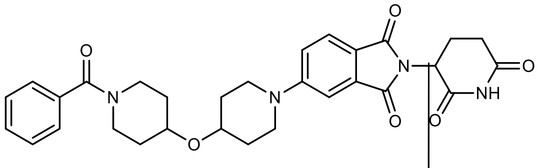

16

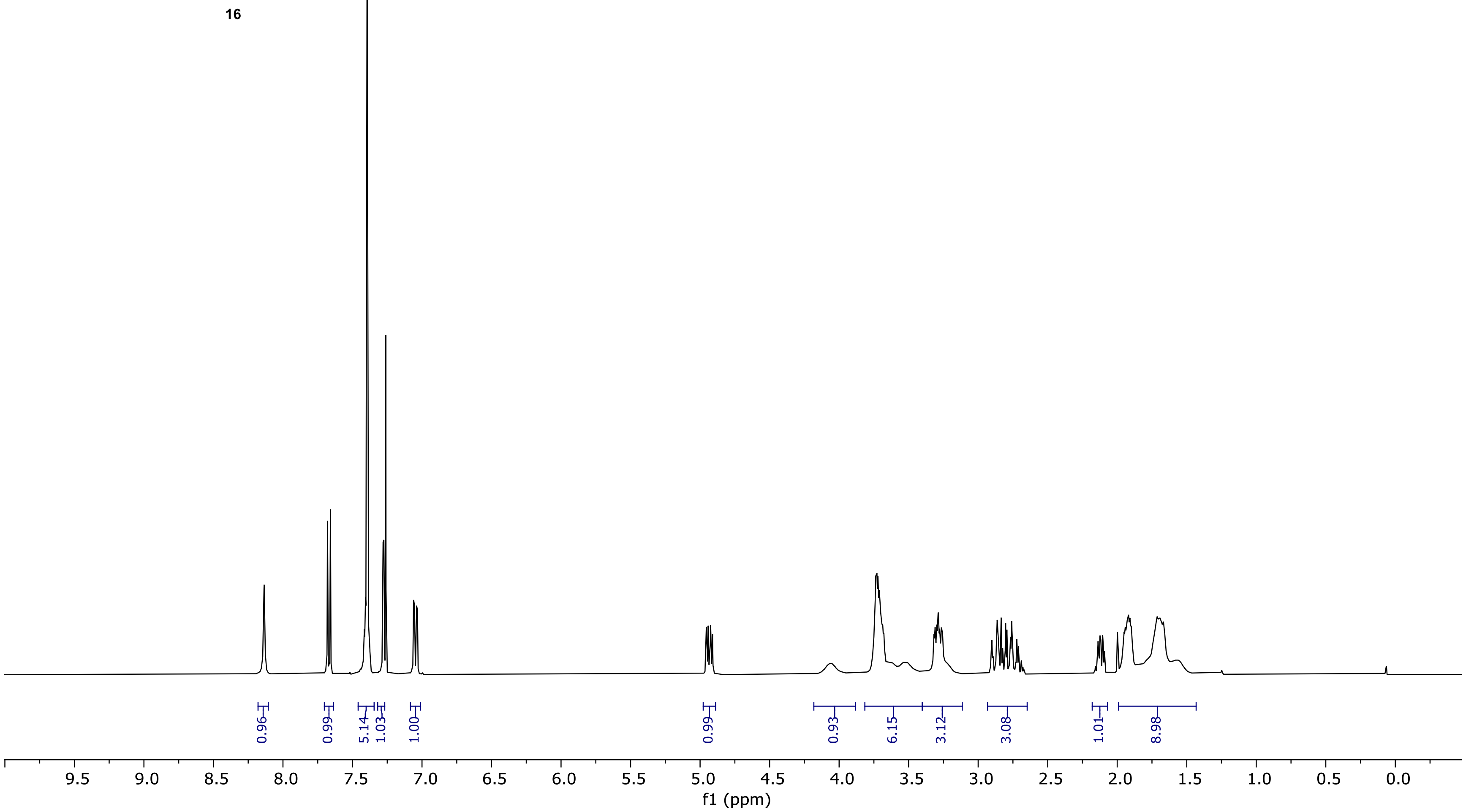

13C NMR (100.66 MHz, CDCl3, 298.1 K)

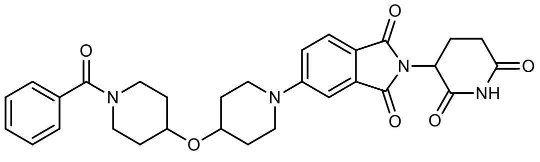

16

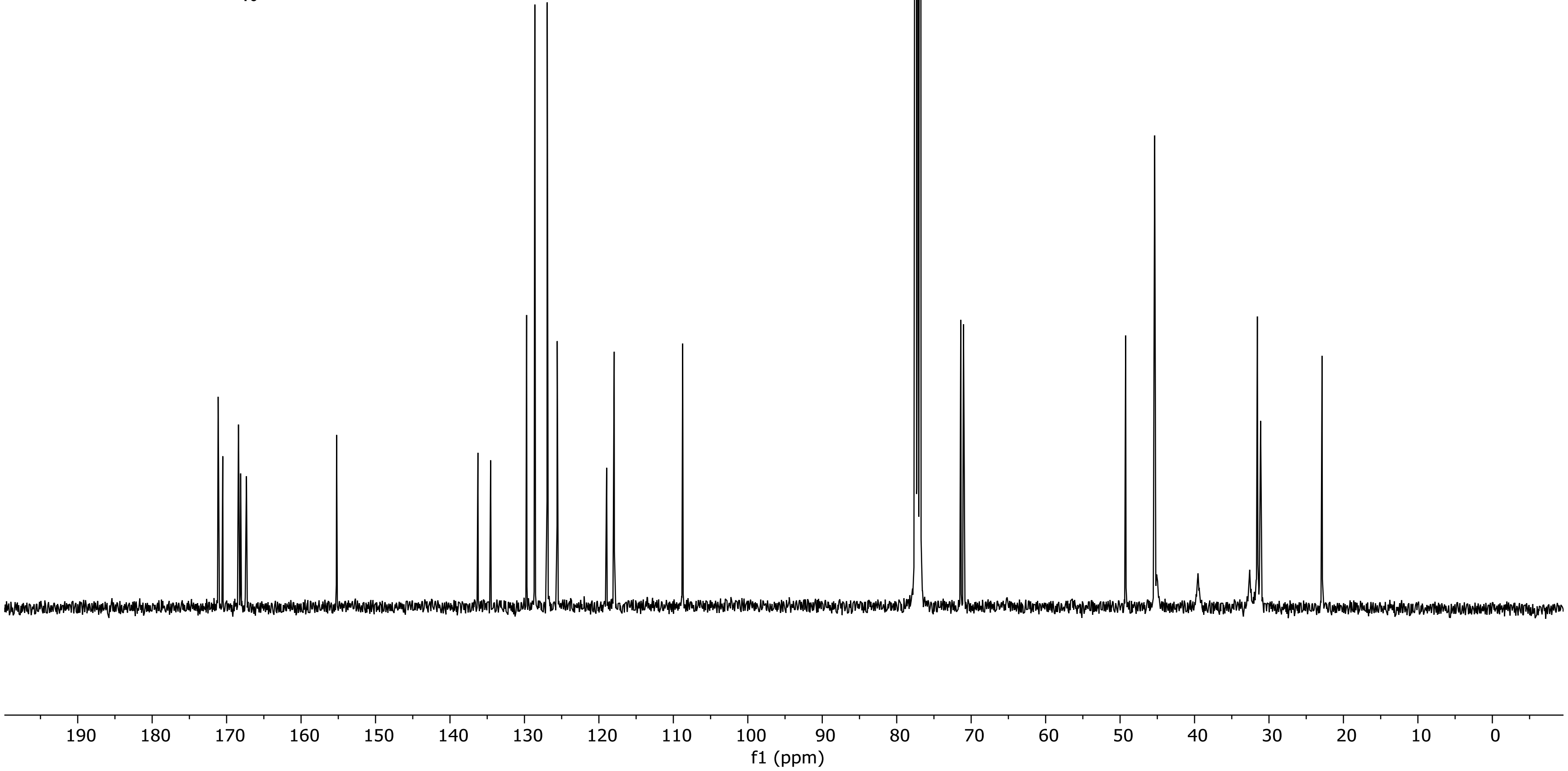

NR-5-131-A LC-MS_ELS  PDA - Total Absorbance Chromatogram

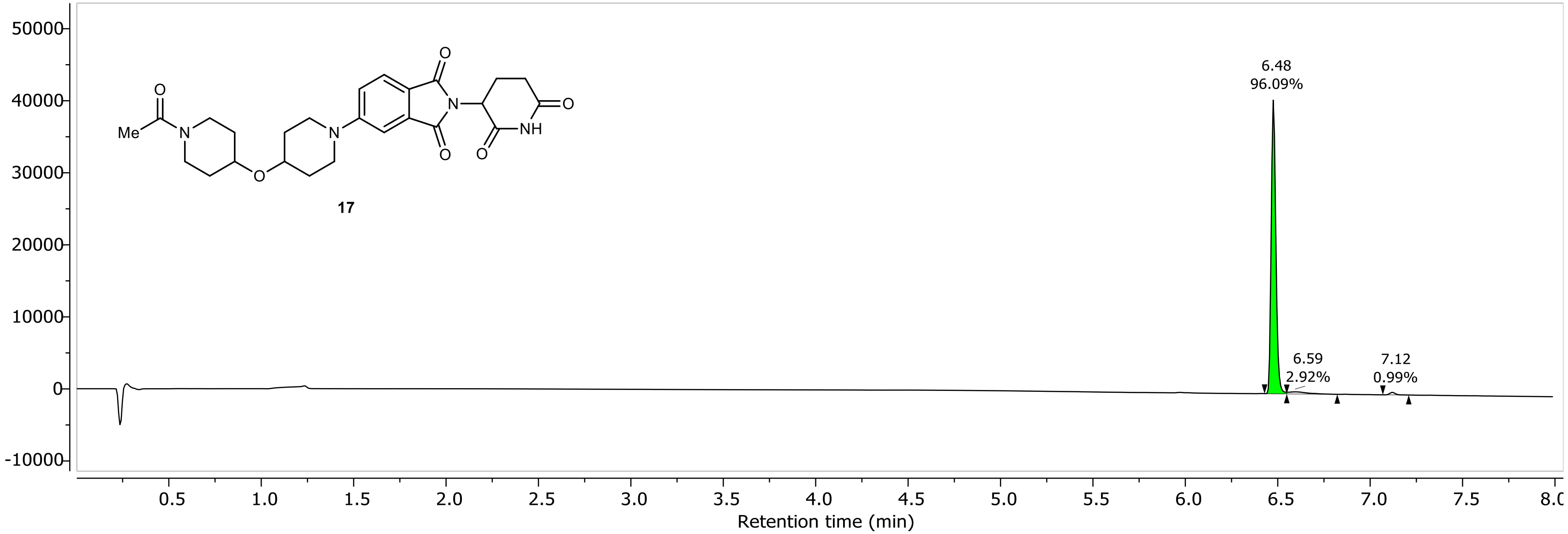

NR-5-131-A LC-MS_ELS DAD1A, Sig=254...Ref=360.0,100.0 Chromatogram

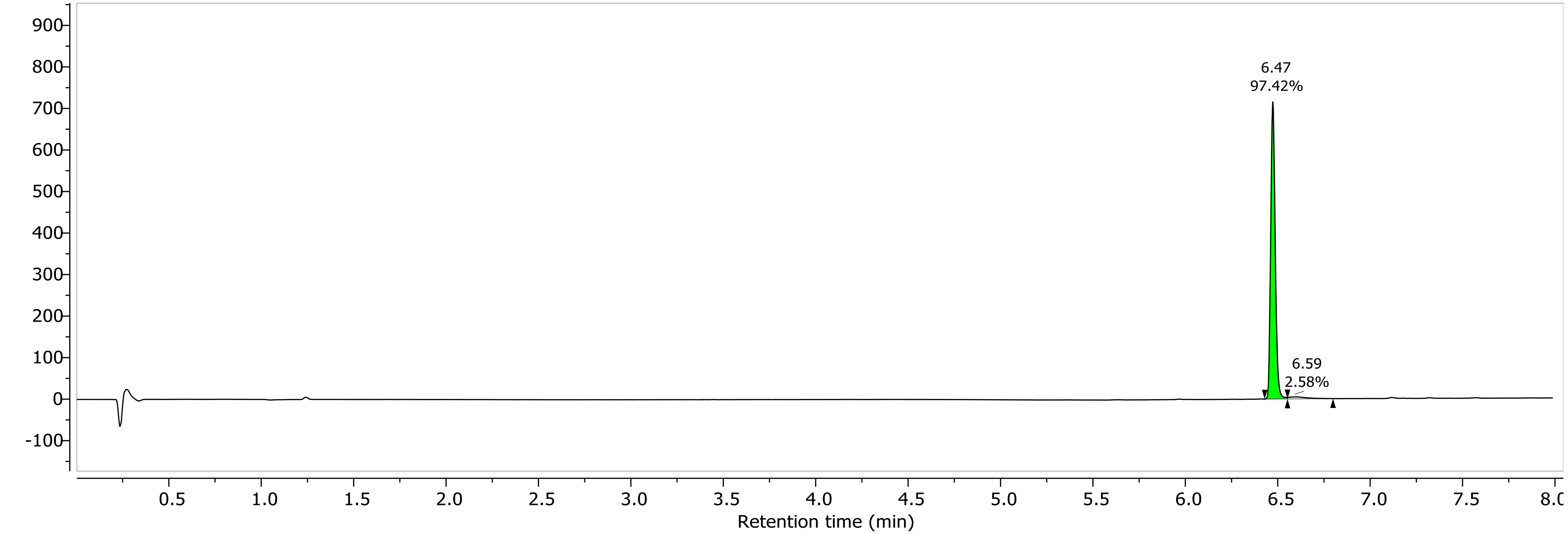

1H NMR (400.26 MHz, CDCl3, 298.1 K)

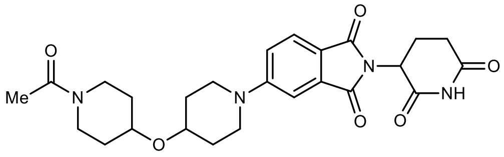

**17**

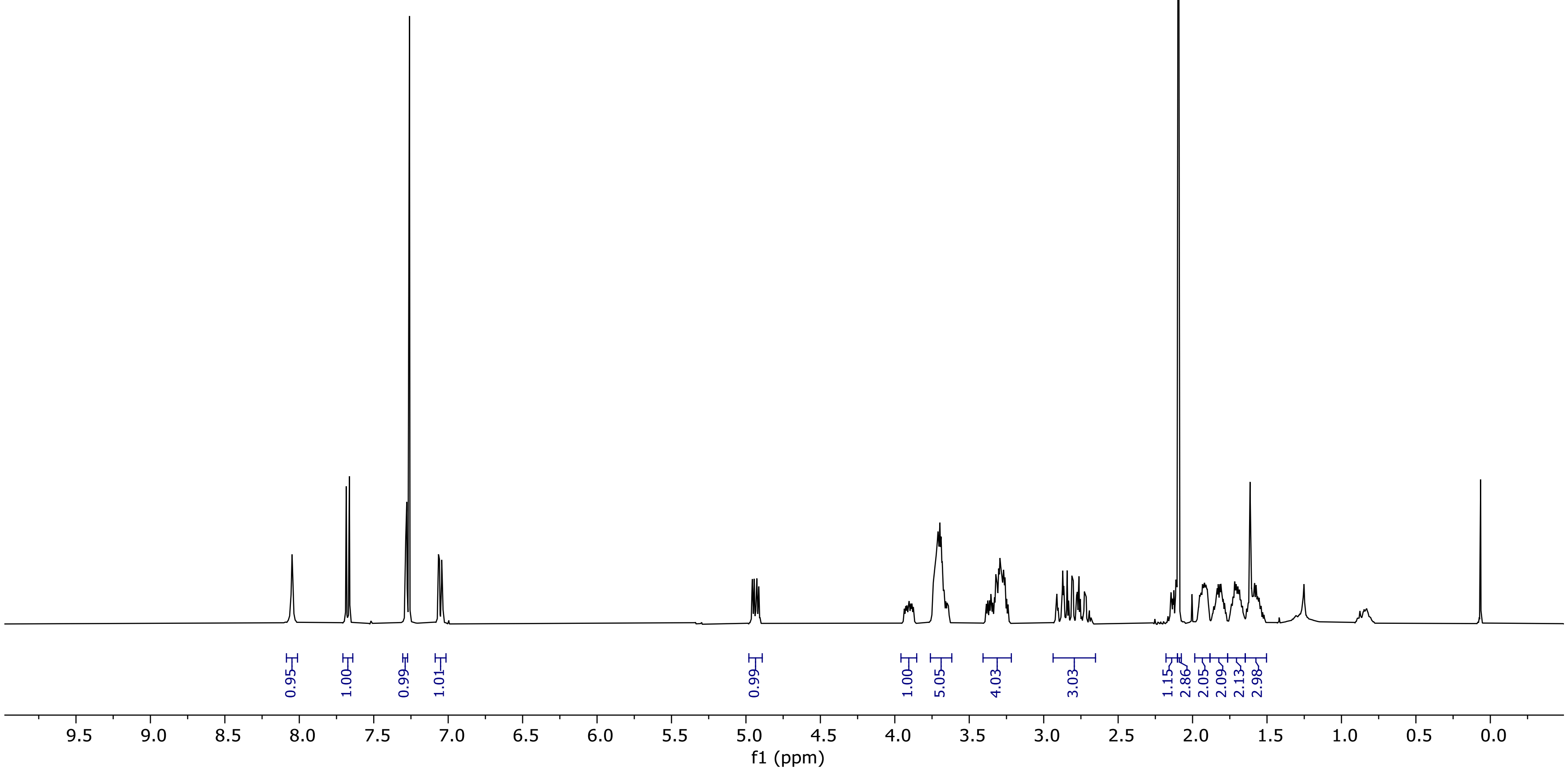

13C NMR (100.66 MHz, CDCl3, 298.1 K)

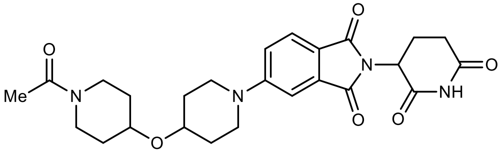

**17**

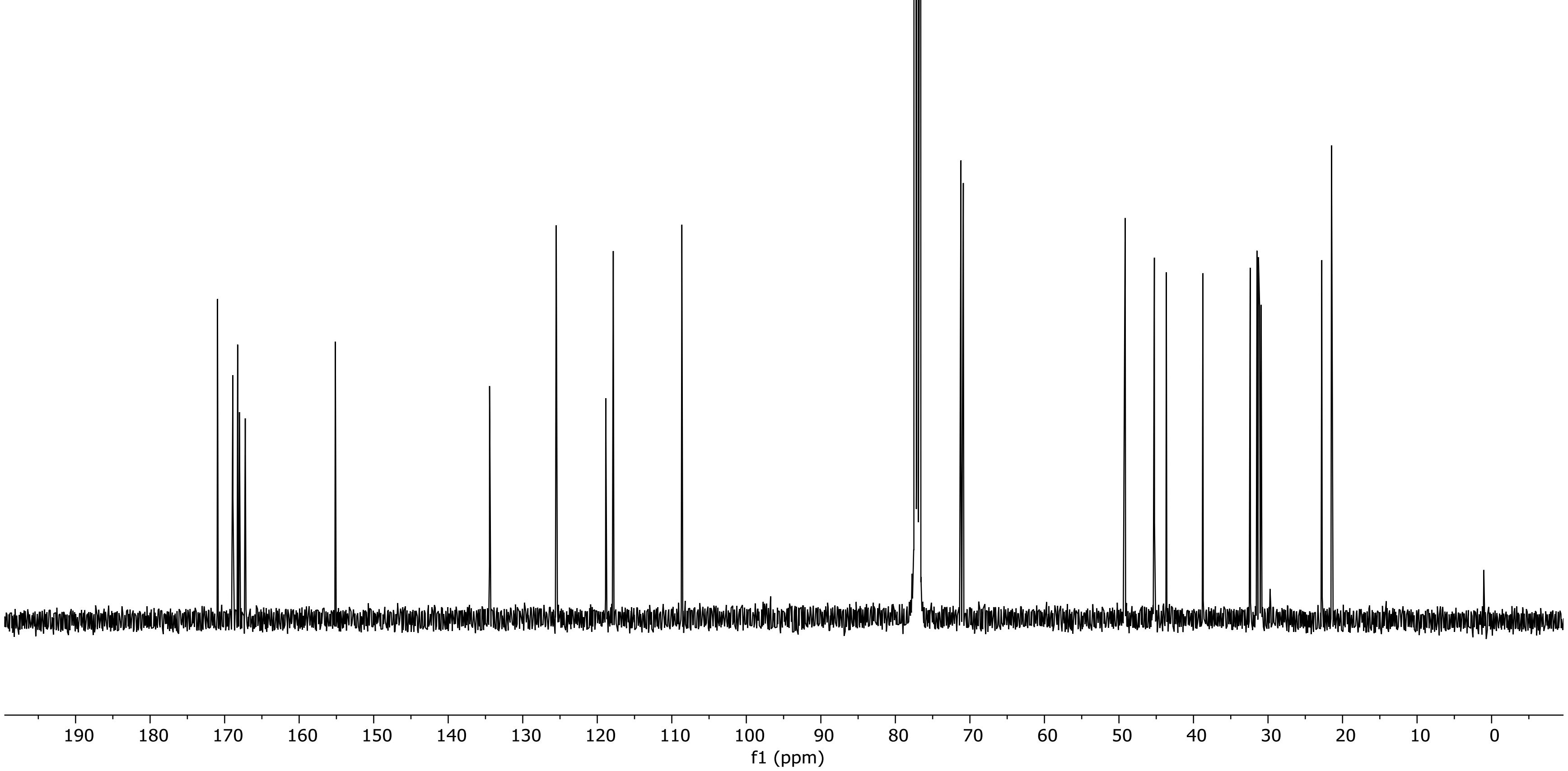

NR-6-071-A LC-MS_ELS  PDA - Total Absorbance Chromatogram

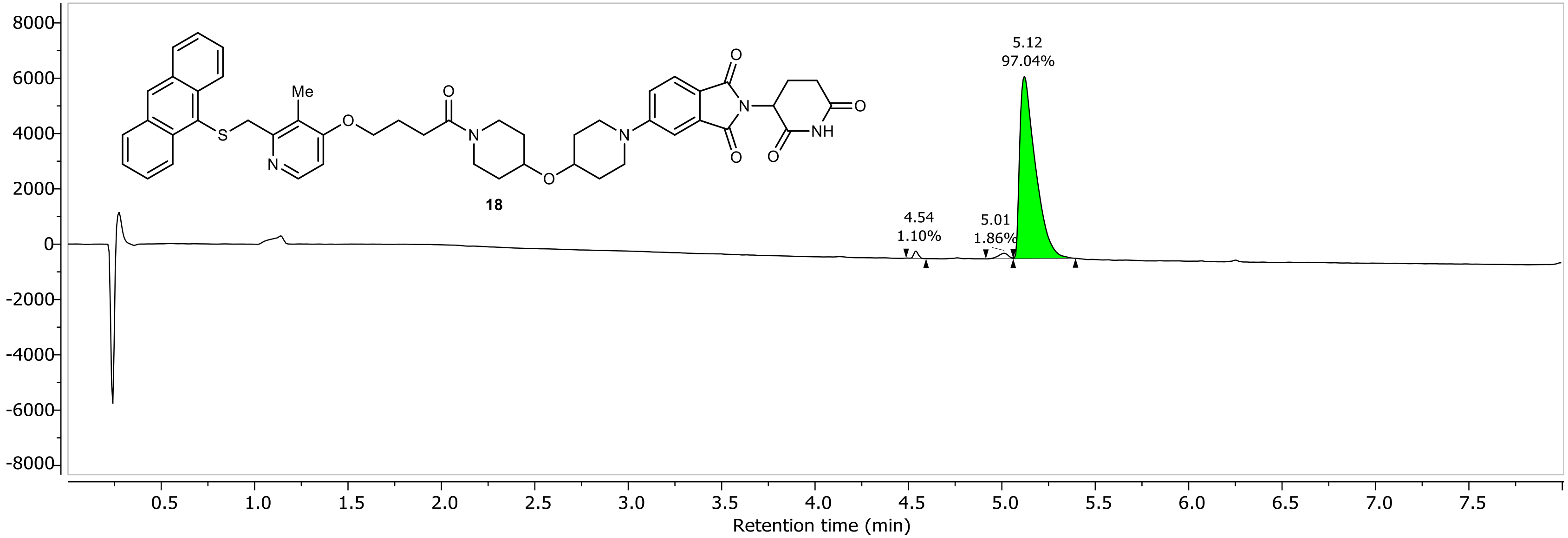

NR-6-071-A LC-MS_ELS DAD1A, Sig=254...Ref=360.0,100.0 Chromatogram

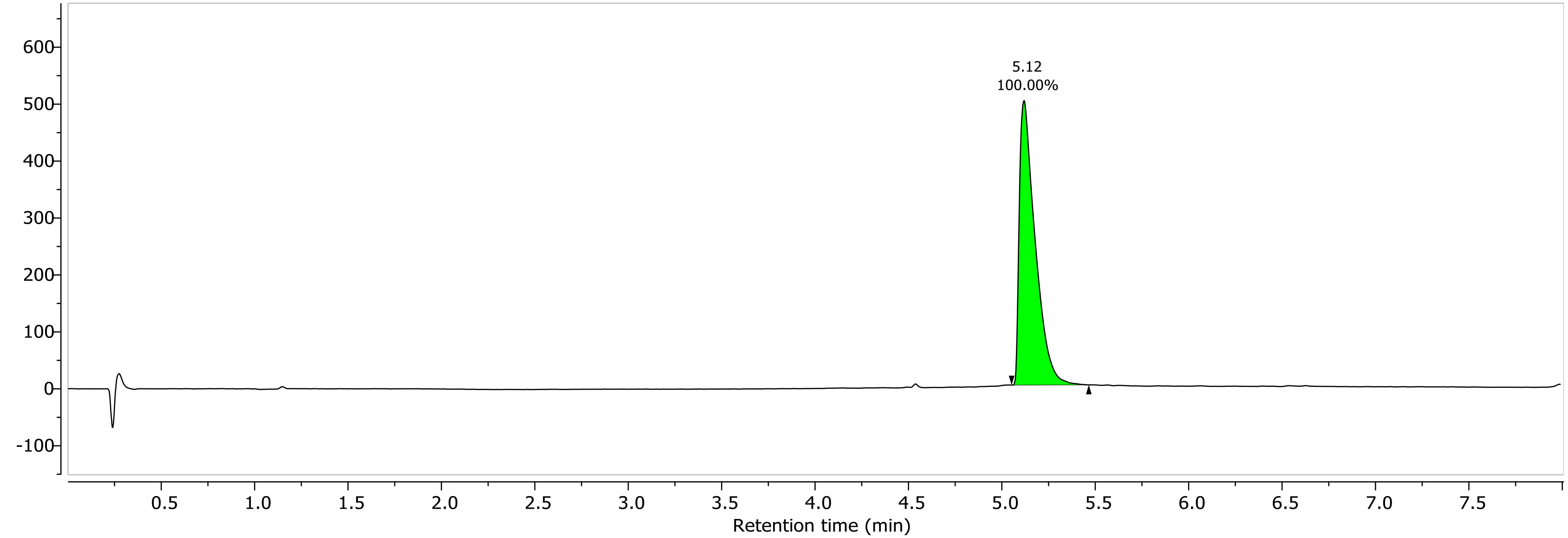

1H NMR (400.26 MHz, CDCl3, 298.2 K)

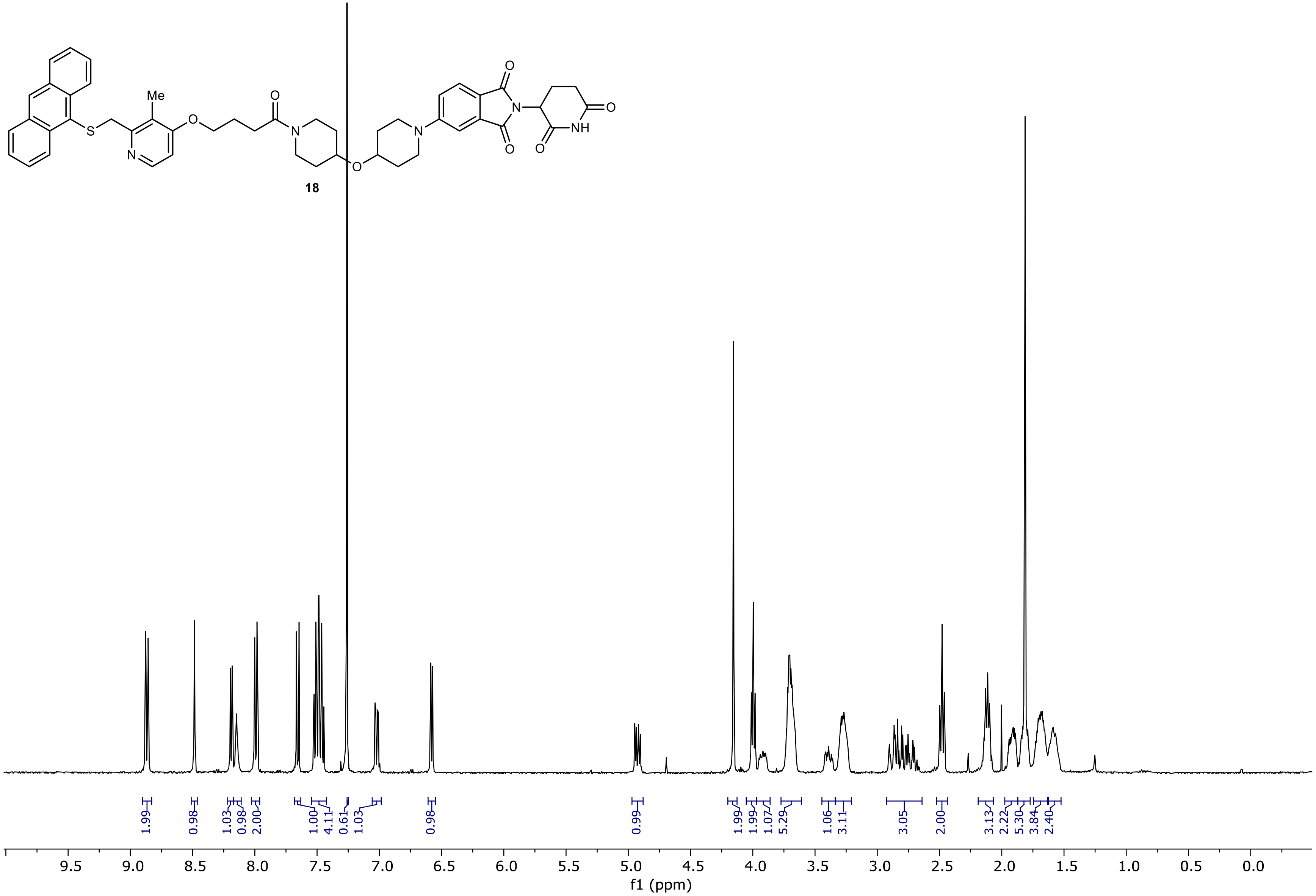

13C NMR (100.66 MHz, CDCl3, 298.1 K)

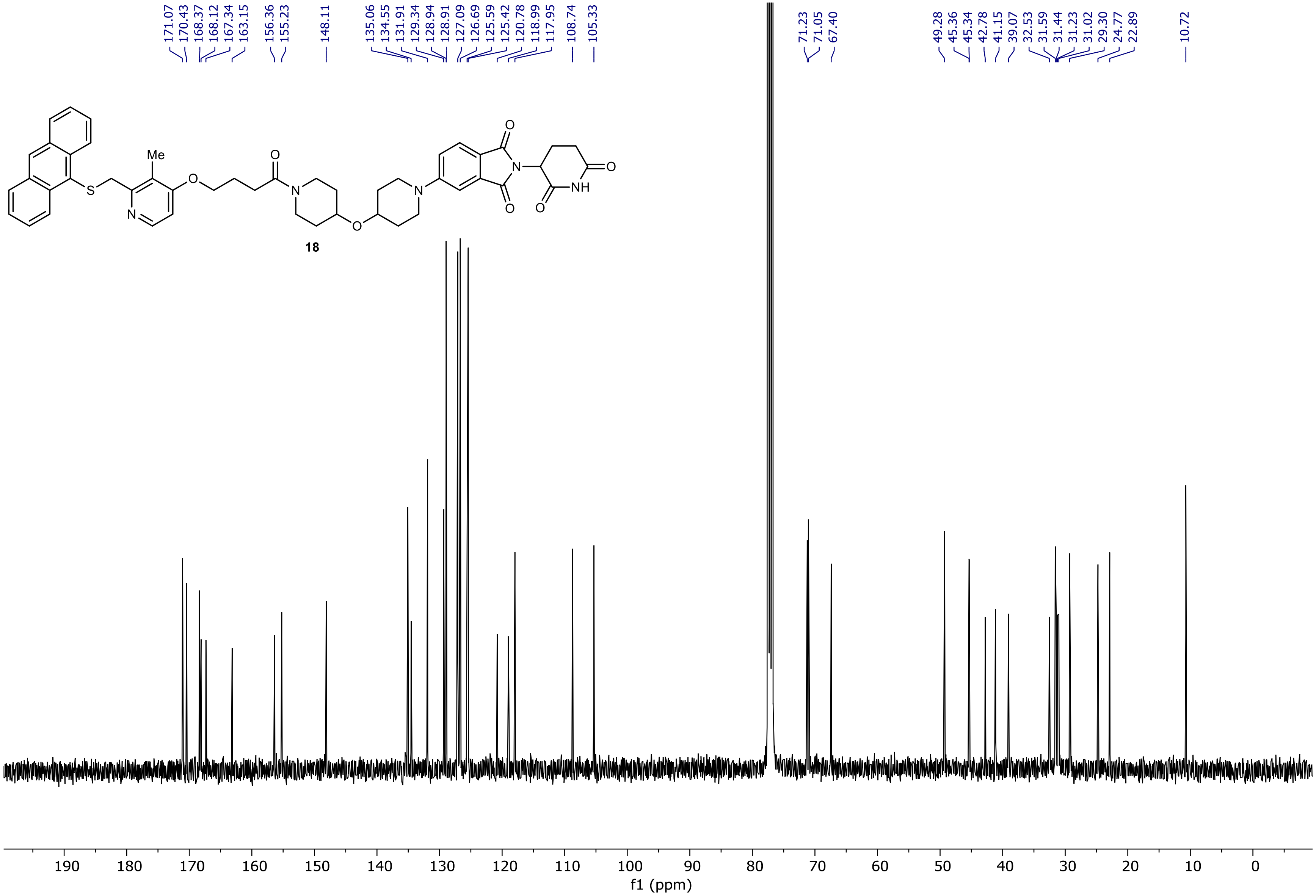

NR-5-125-A LC-MS_ELS  PDA - Total Absorbance Chromatogram

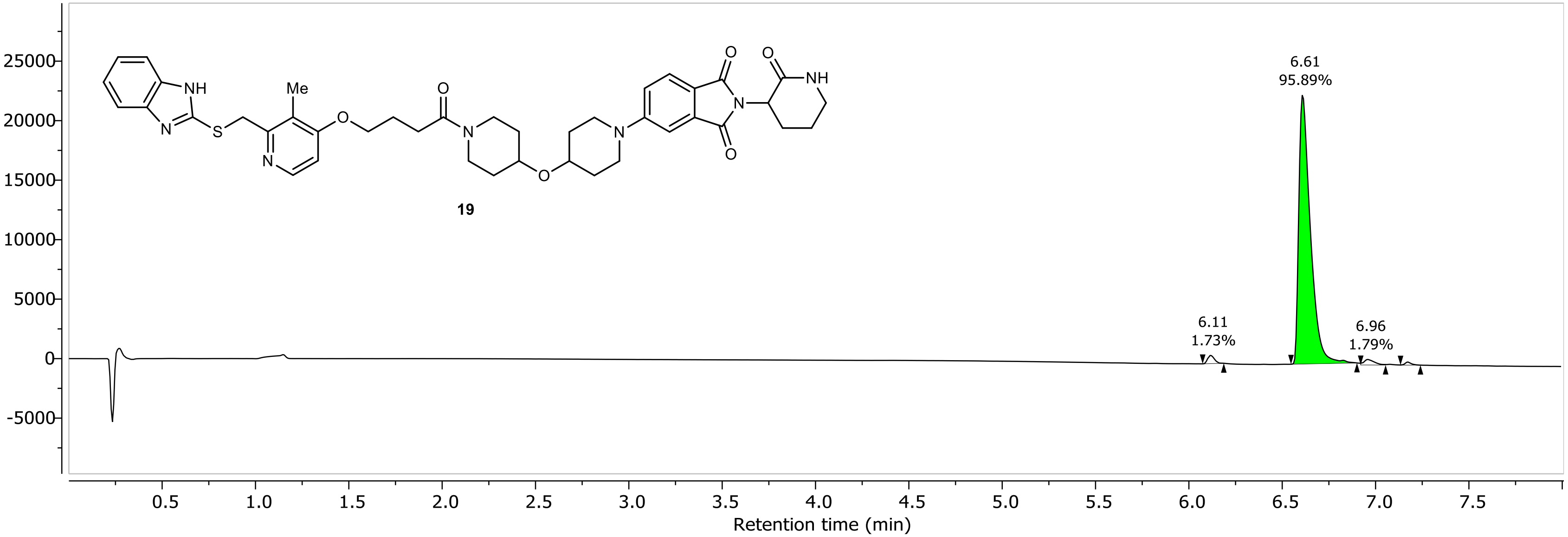

NR-5-125-A LC-MS_ELS DAD1A, Sig=254...Ref=360.0,100.0 Chromatogram

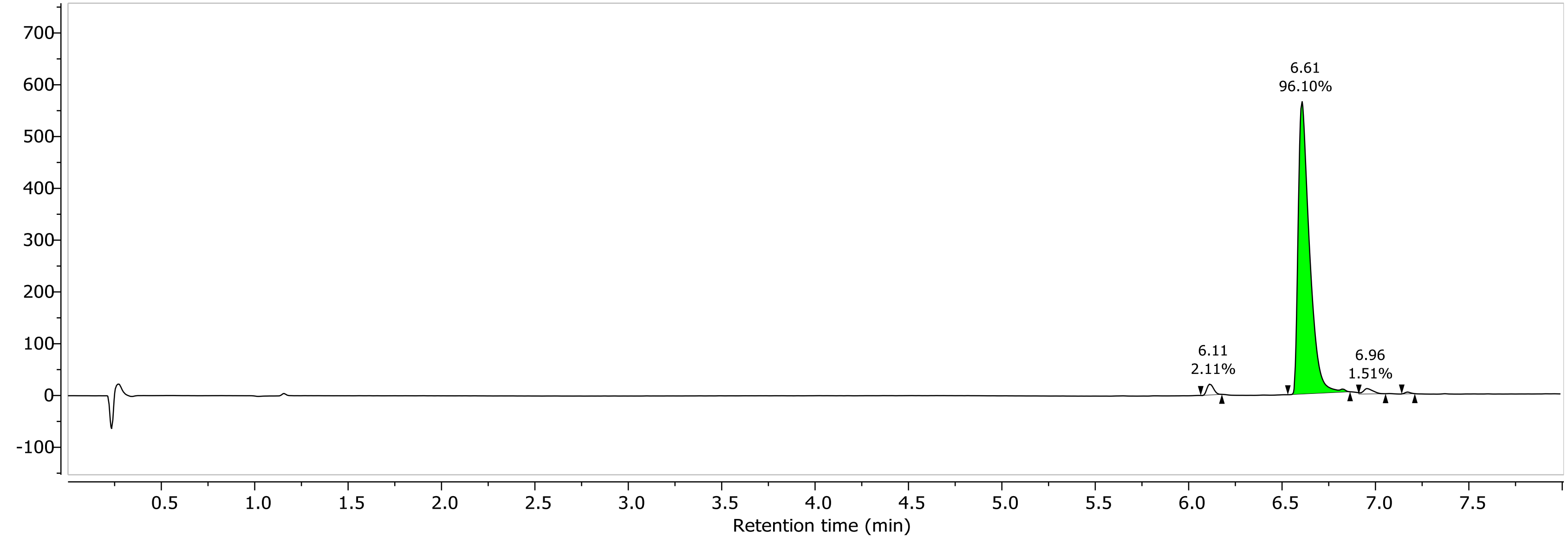

1H NMR (400.26 MHz, CDCl3, 298.1 K)

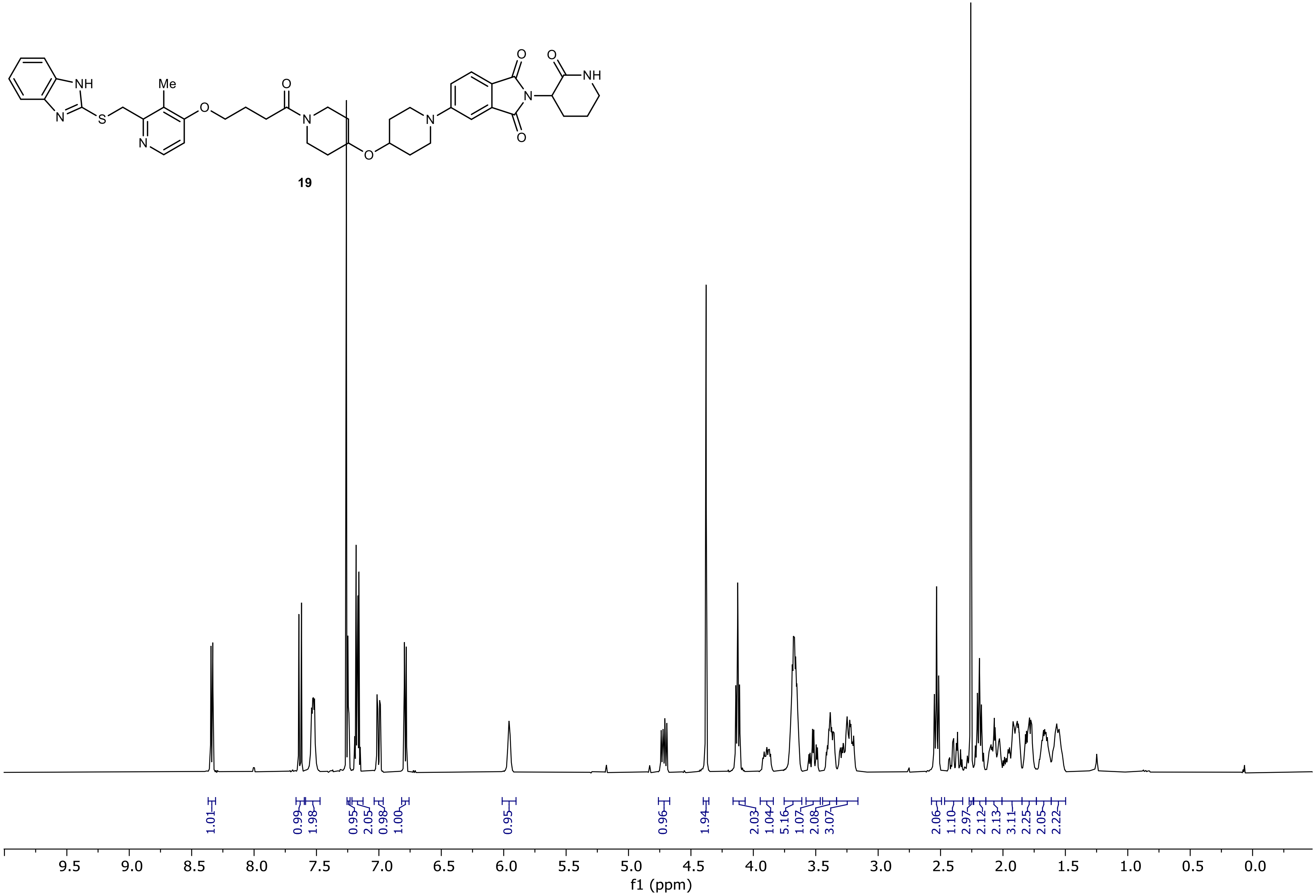

13C NMR (100.66 MHz, CDCl3, 298.1 K)

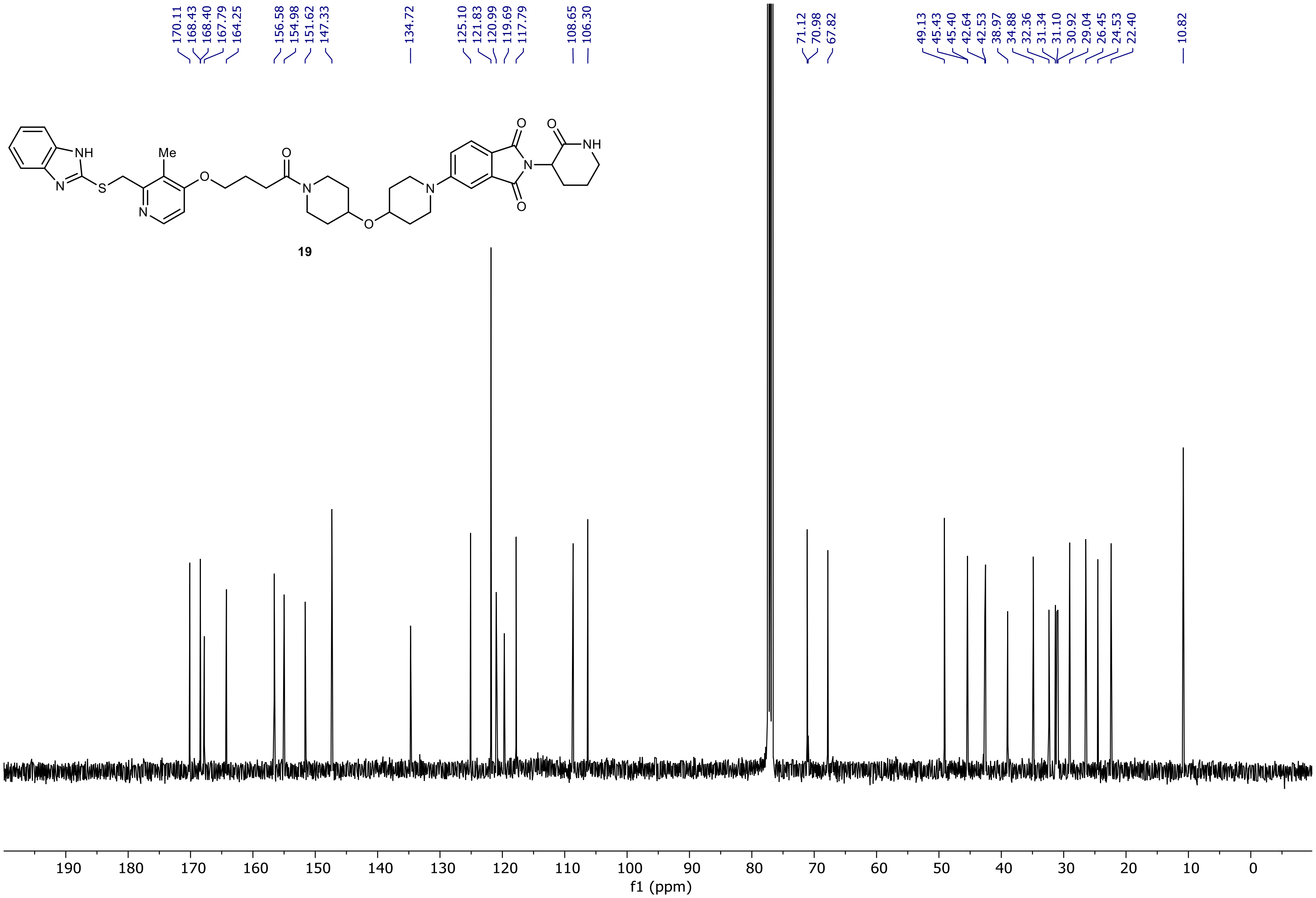

NR-5-114-A LC-MS_ELS  PDA - Total Absorbance Chromatogram

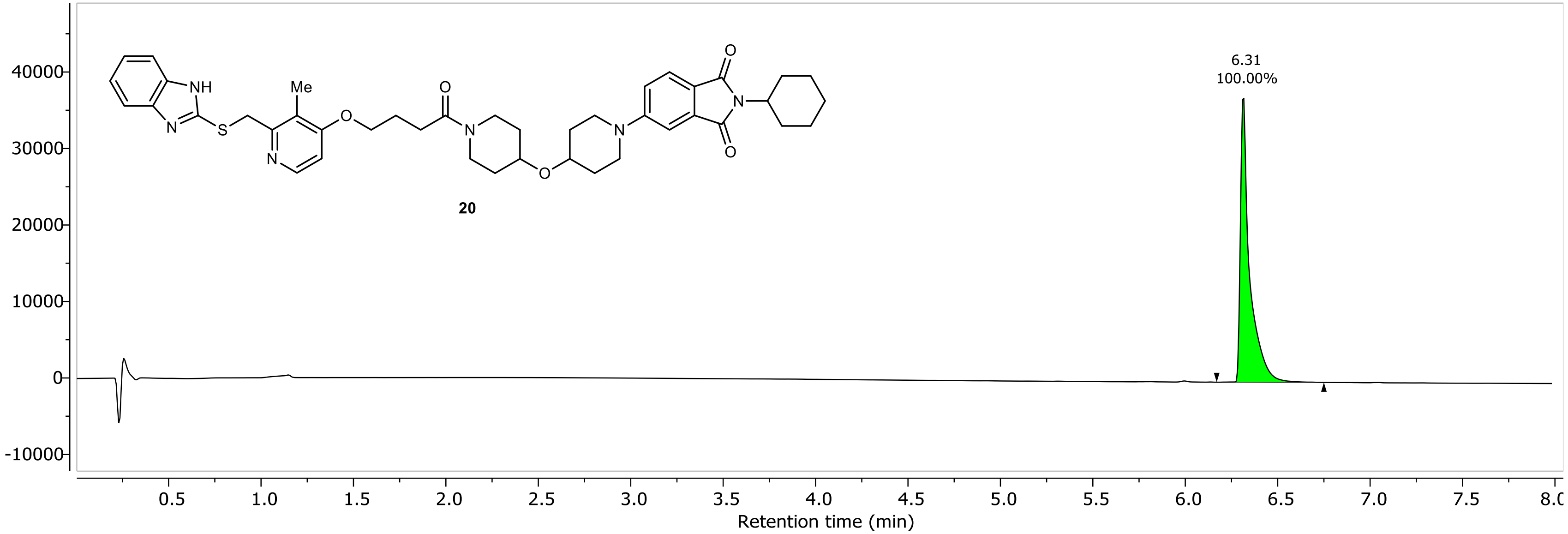

NR-5-114-A LC-MS_ELS DAD1A, Sig=254...Ref=360.0,100.0 Chromatogram

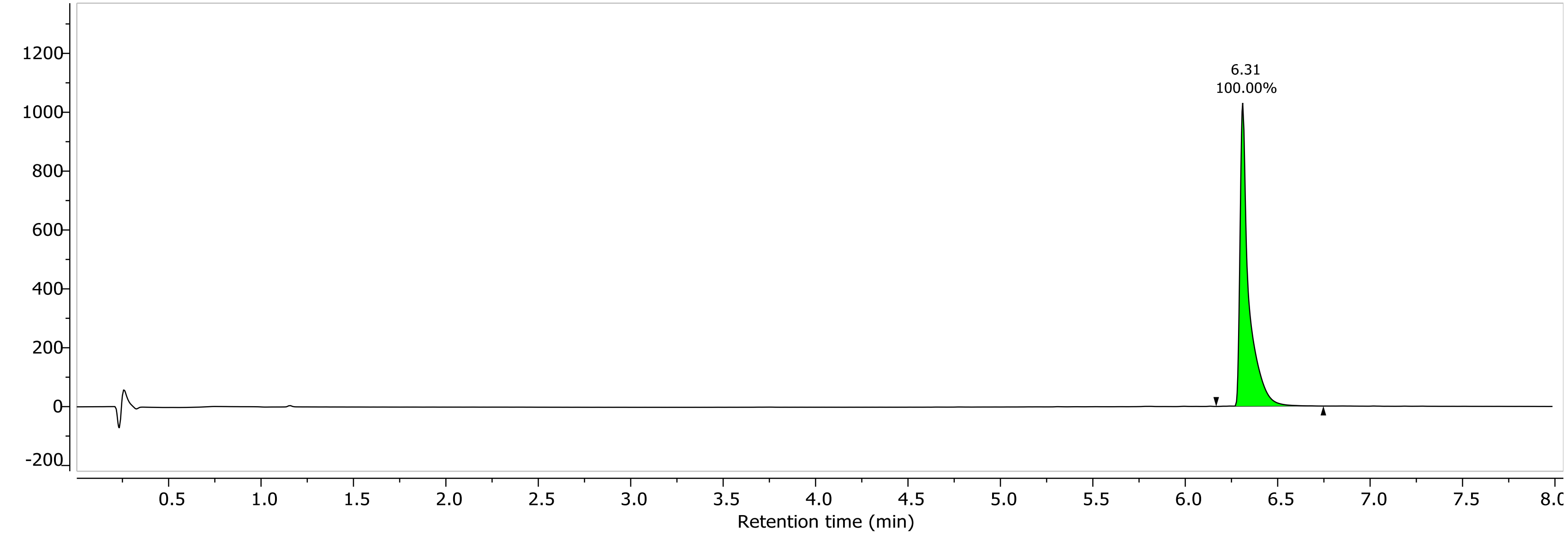

1H NMR (400.26 MHz, CDCl3, 298.1 K)

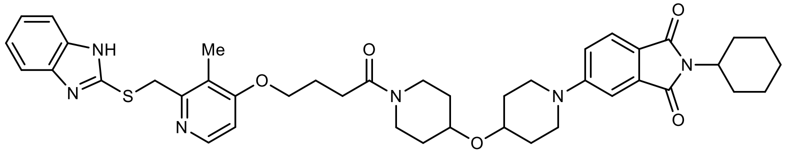

**20**

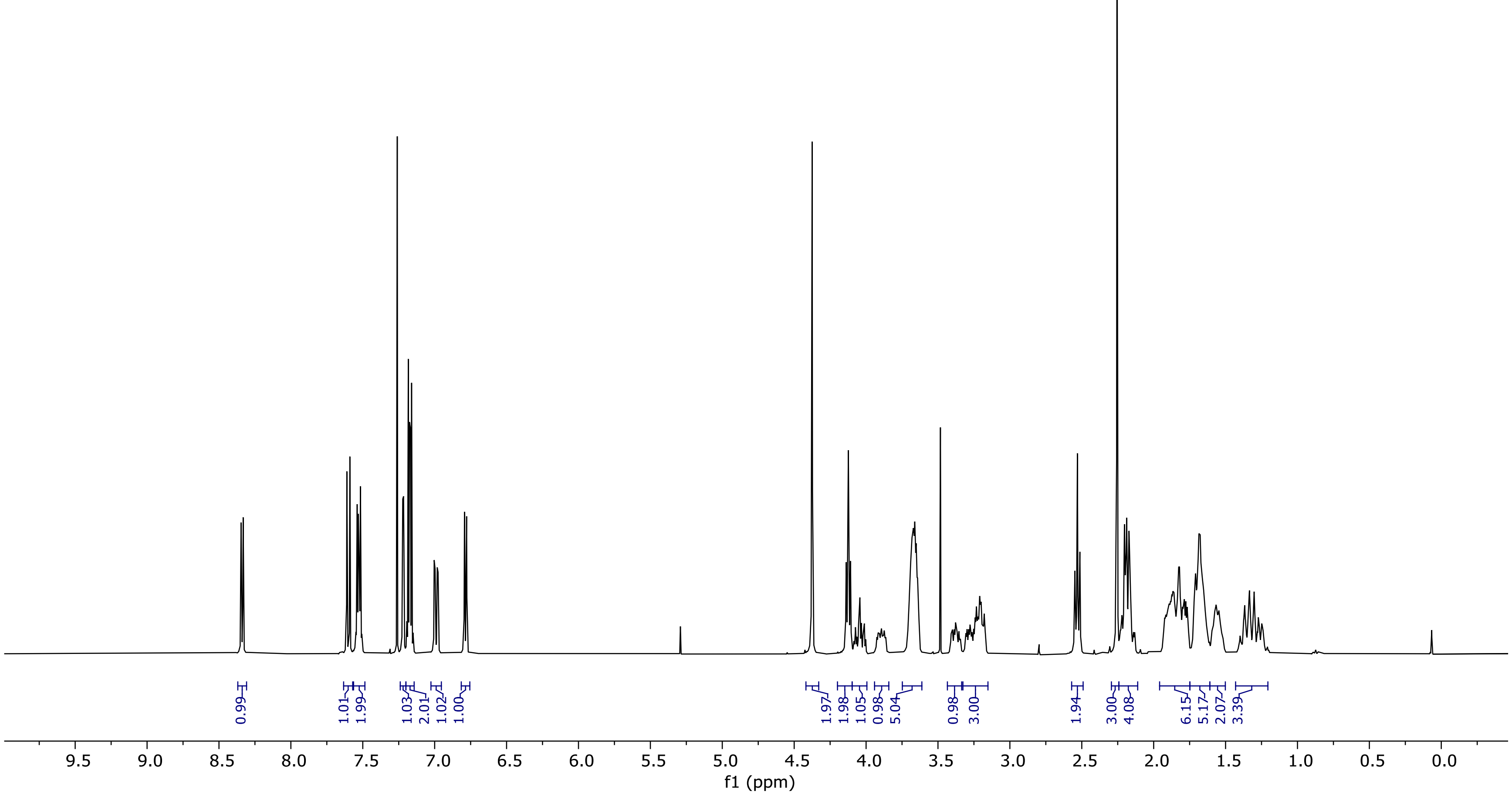

13C NMR (100.66 MHz, CDCl3, 298.1 K)

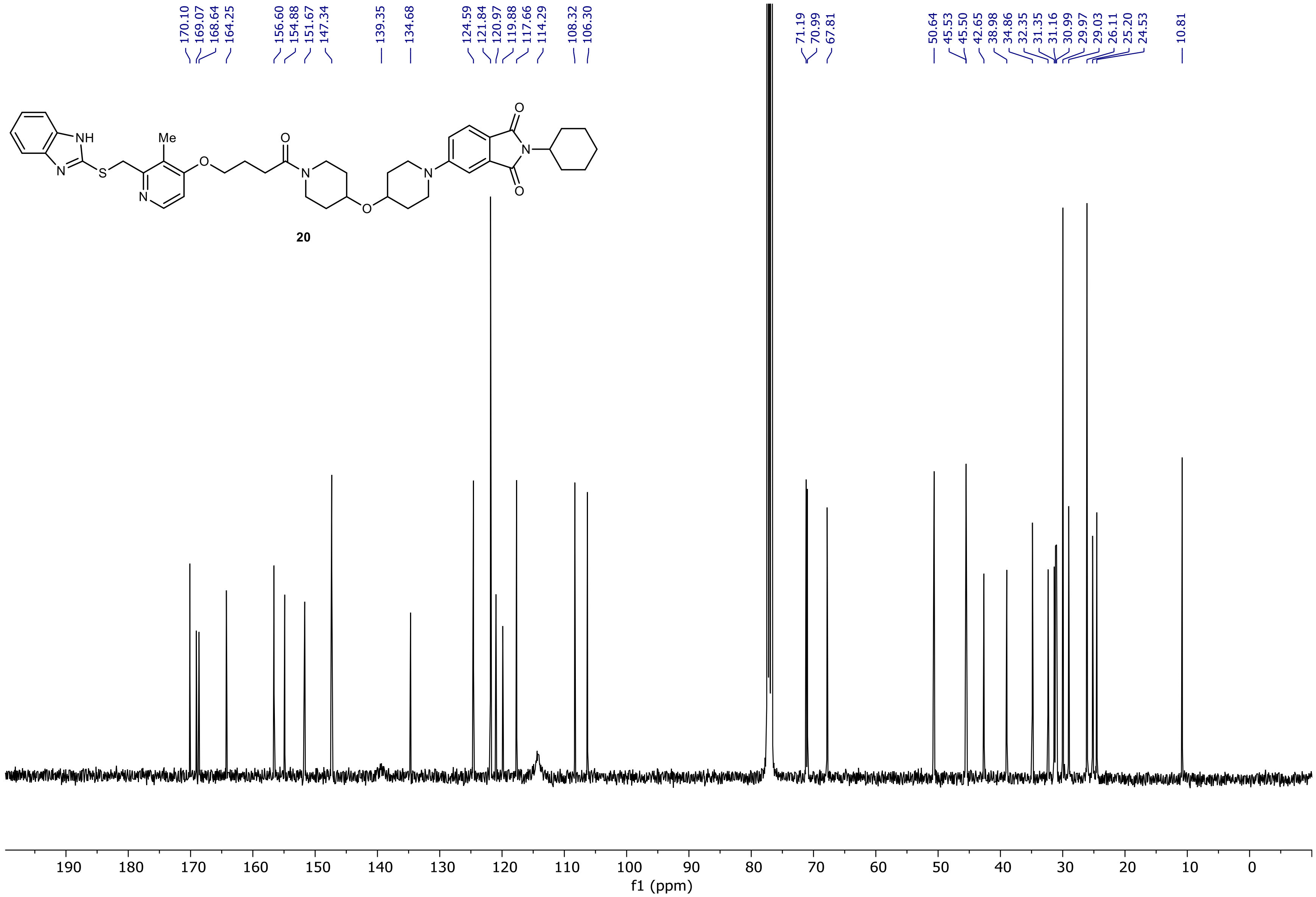

NR-5-121-A LC-MS_ELS  PDA - Total Absorbance Chromatogram

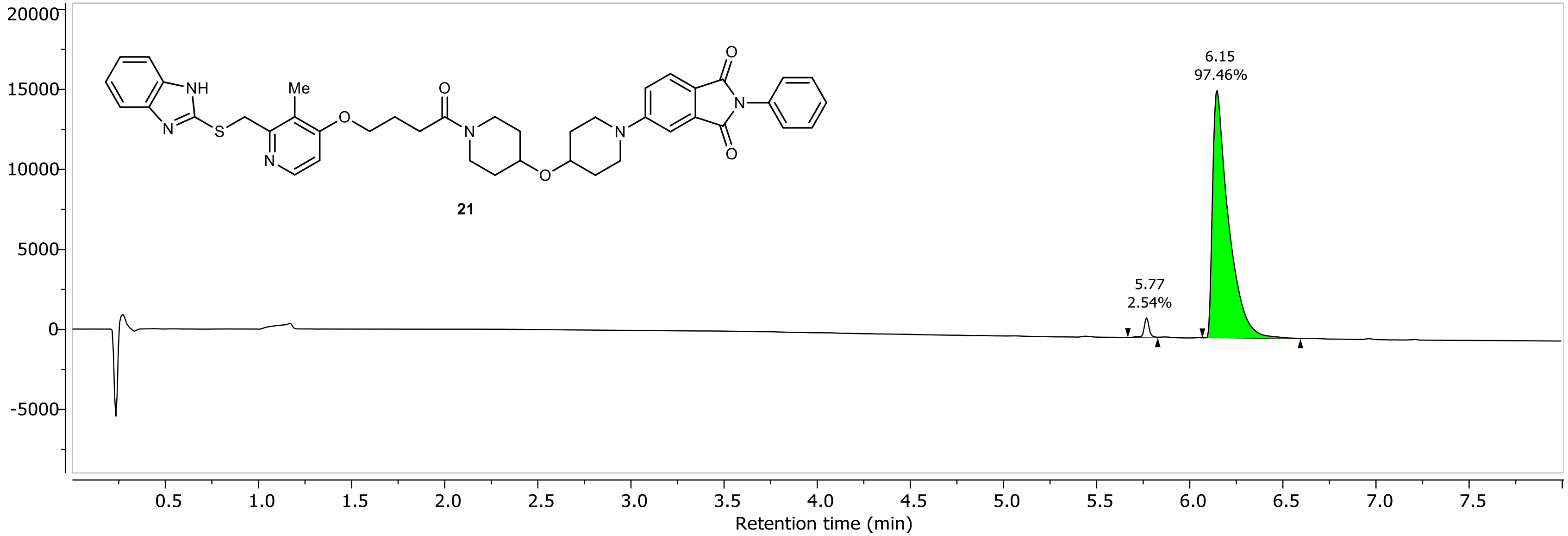

NR-5-121-A LC-MS_ELS DAD1A, Sig=254...Ref=360.0,100.0 Chromatogram

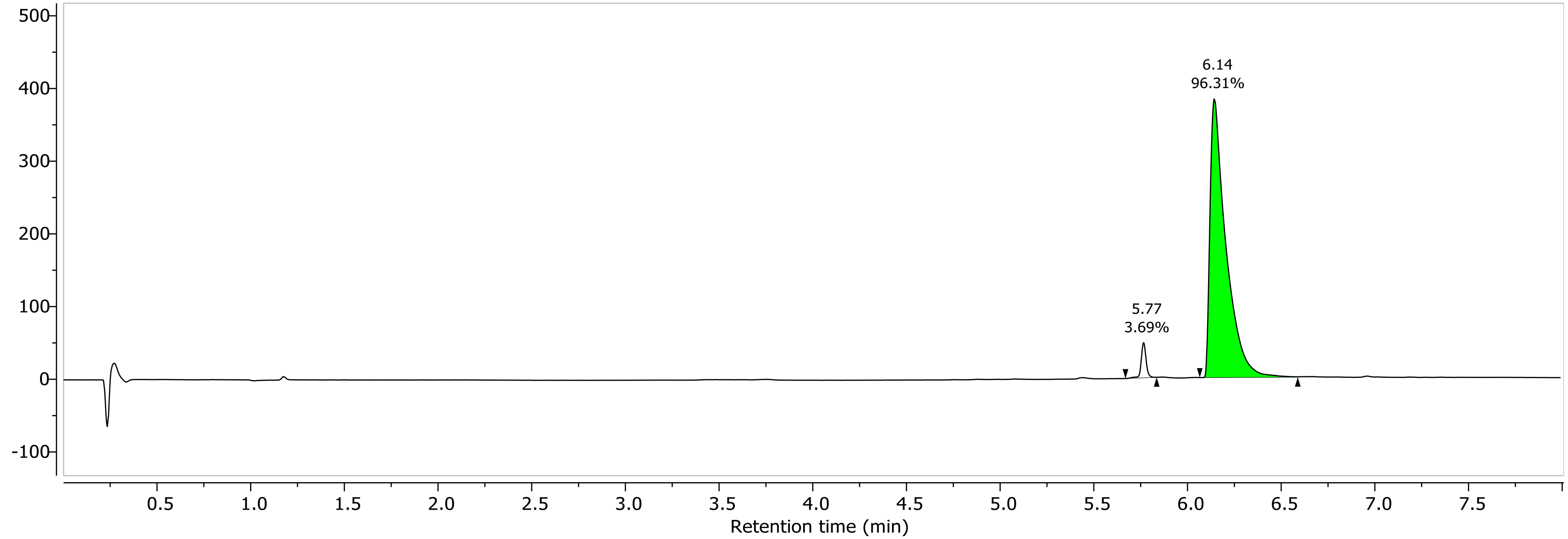

1H NMR (400.26 MHz, CDCl3, 298.1 K)

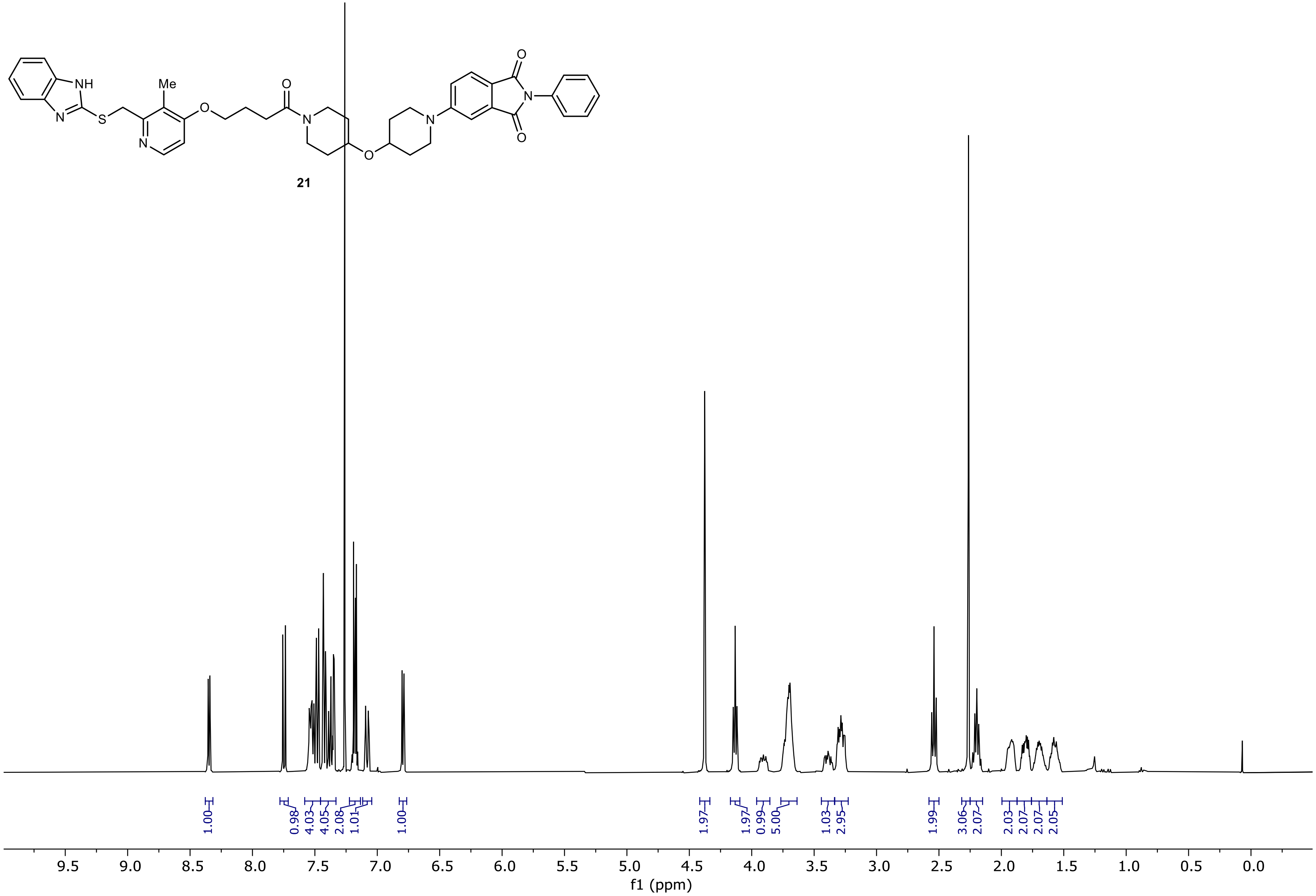

13C NMR (100.66 MHz, CDCl3, 298.1 K)

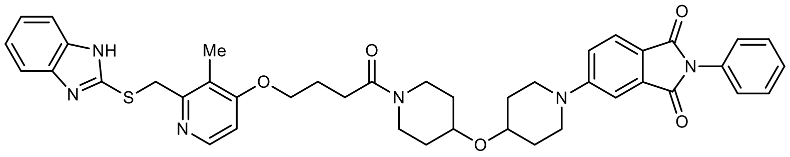

21

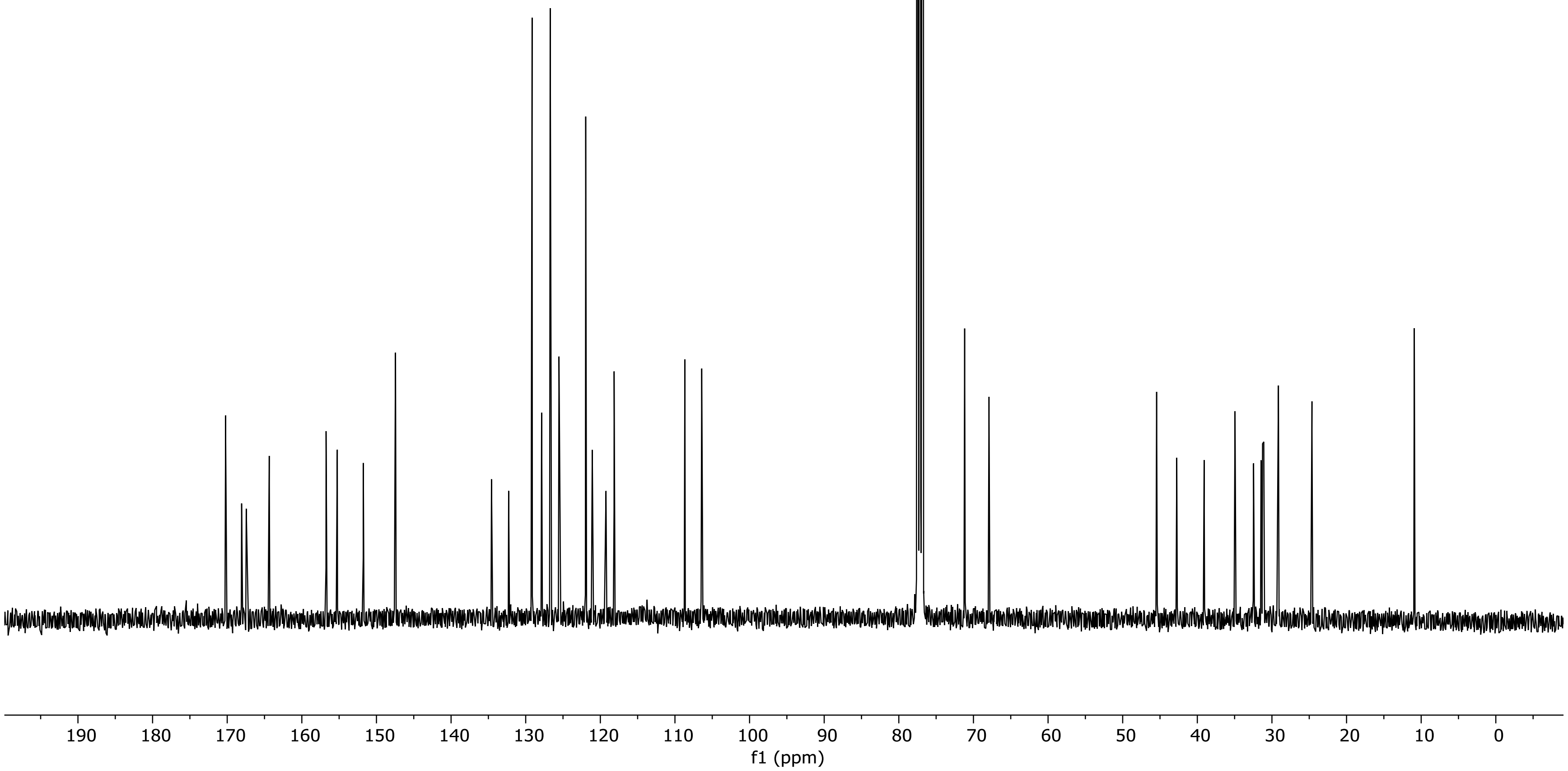

NR-5-126-A LC-MS_ELS  PDA - Total Absorbance Chromatogram

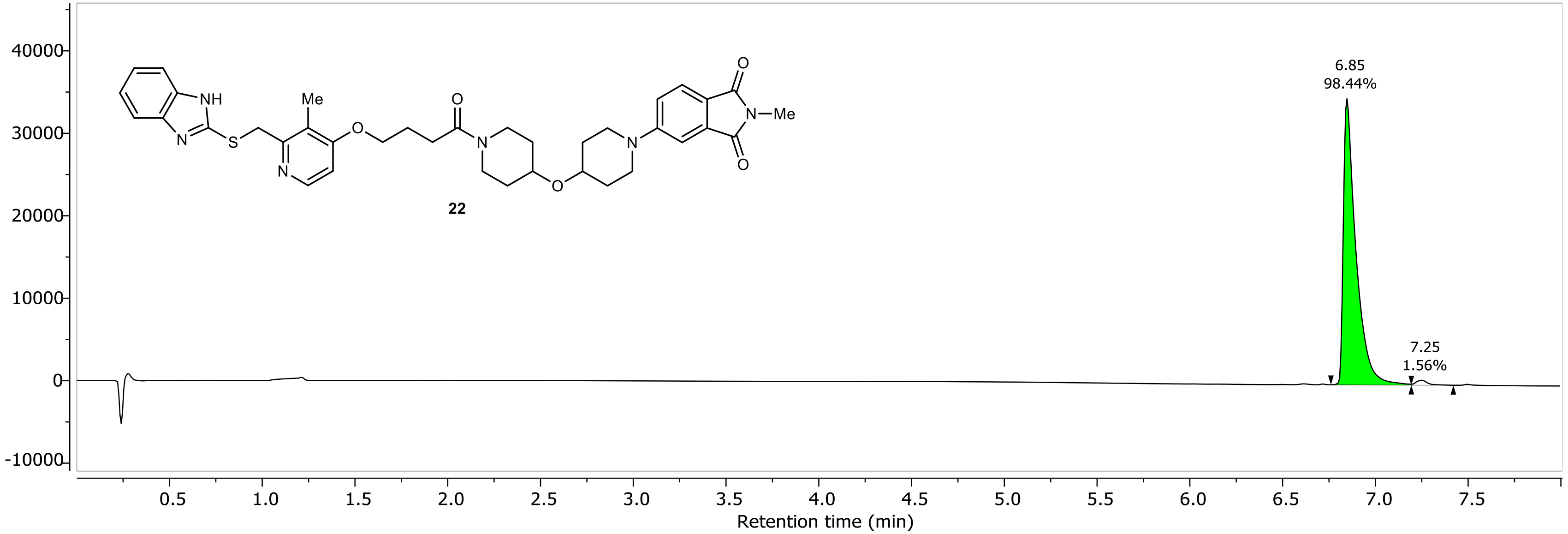

NR-5-126-A LC-MS_ELS DAD1A, Sig=254...Ref=360.0,100.0 Chromatogram

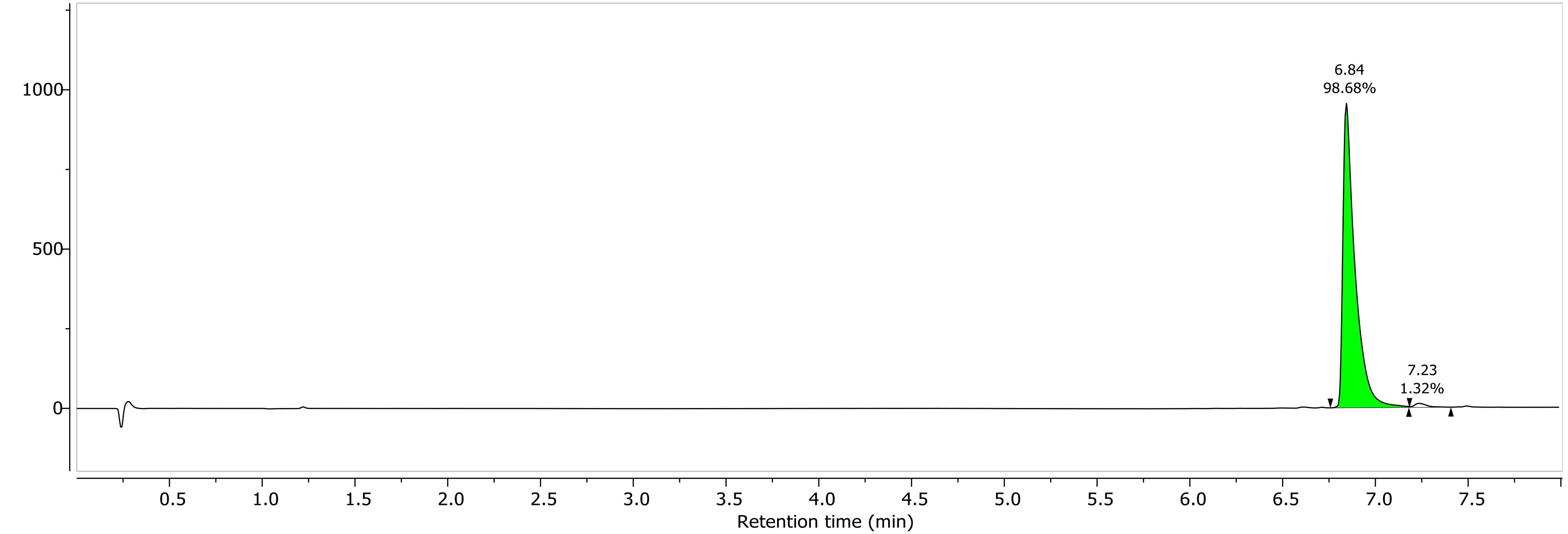

1H NMR (400.26 MHz, CDCl3, 298.1 K)

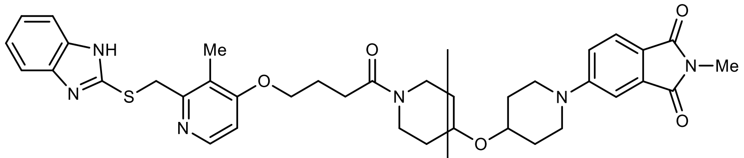

22

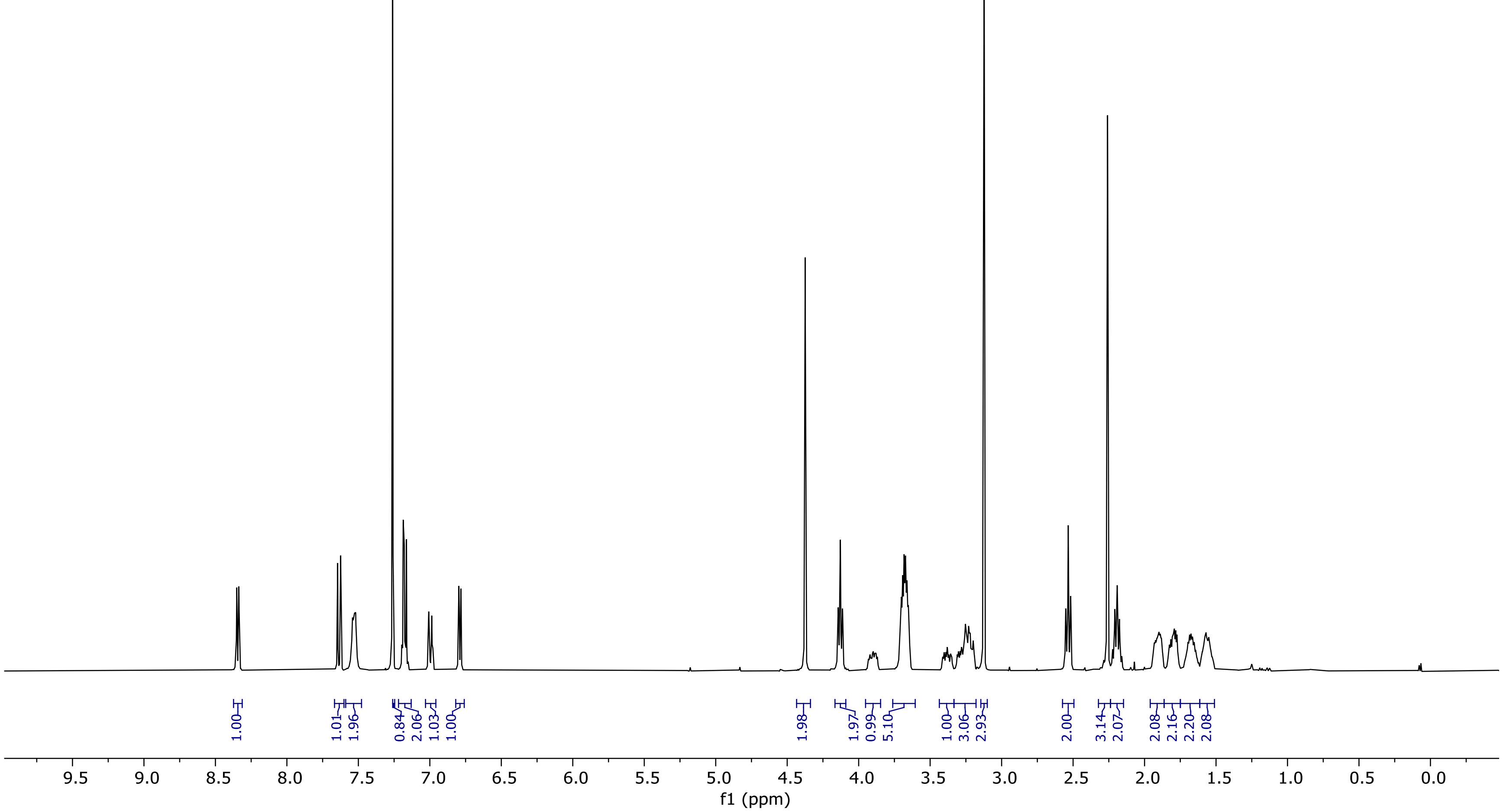

13C NMR (100.66 MHz, CDCl3, 298.1 K)

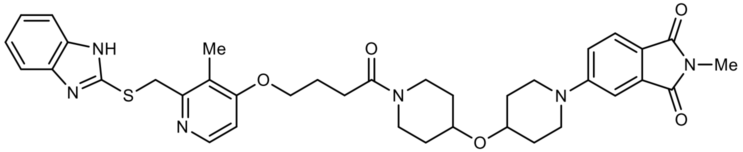

22

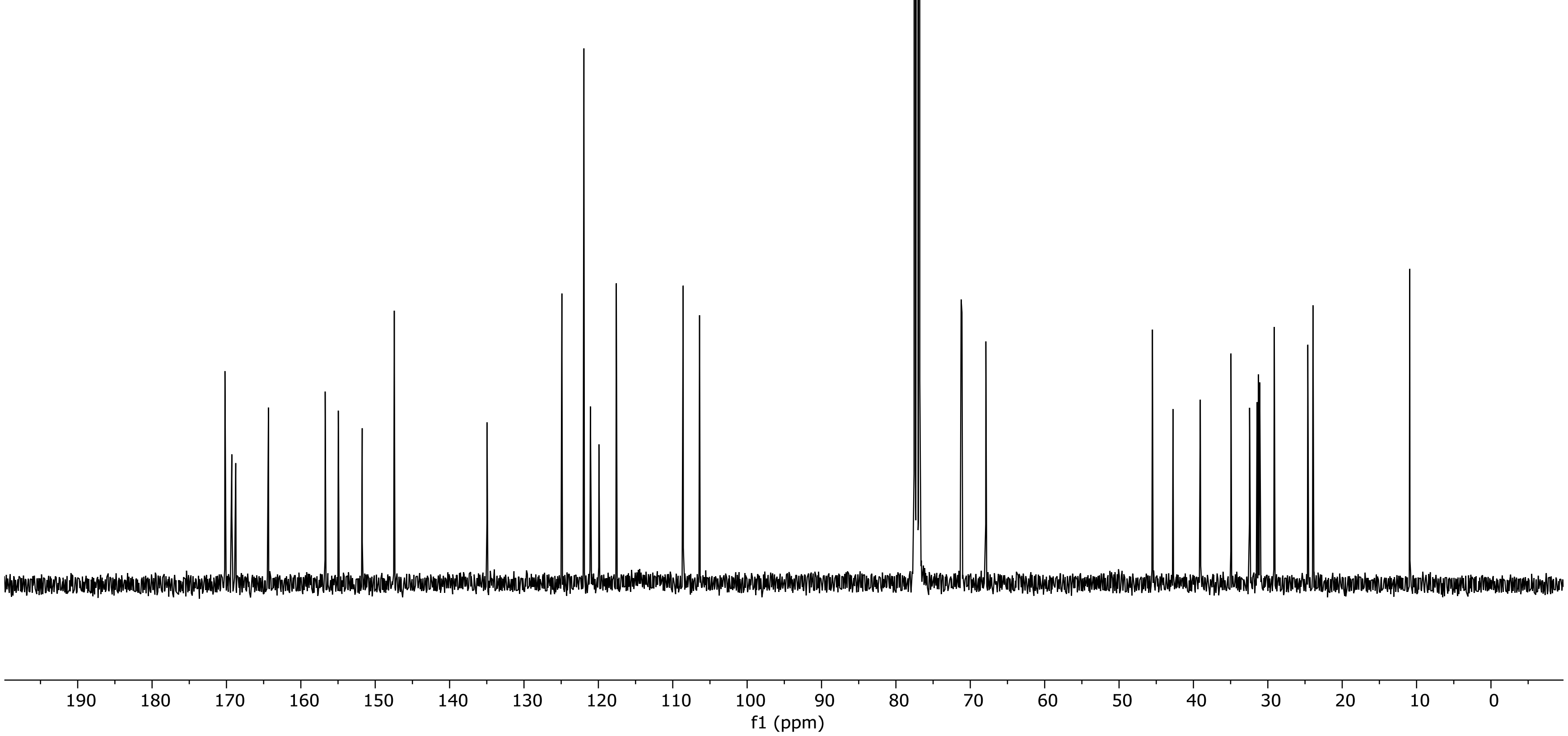

NR-5-113-A LC-MS_ELS  PDA - Total Absorbance Chromatogram

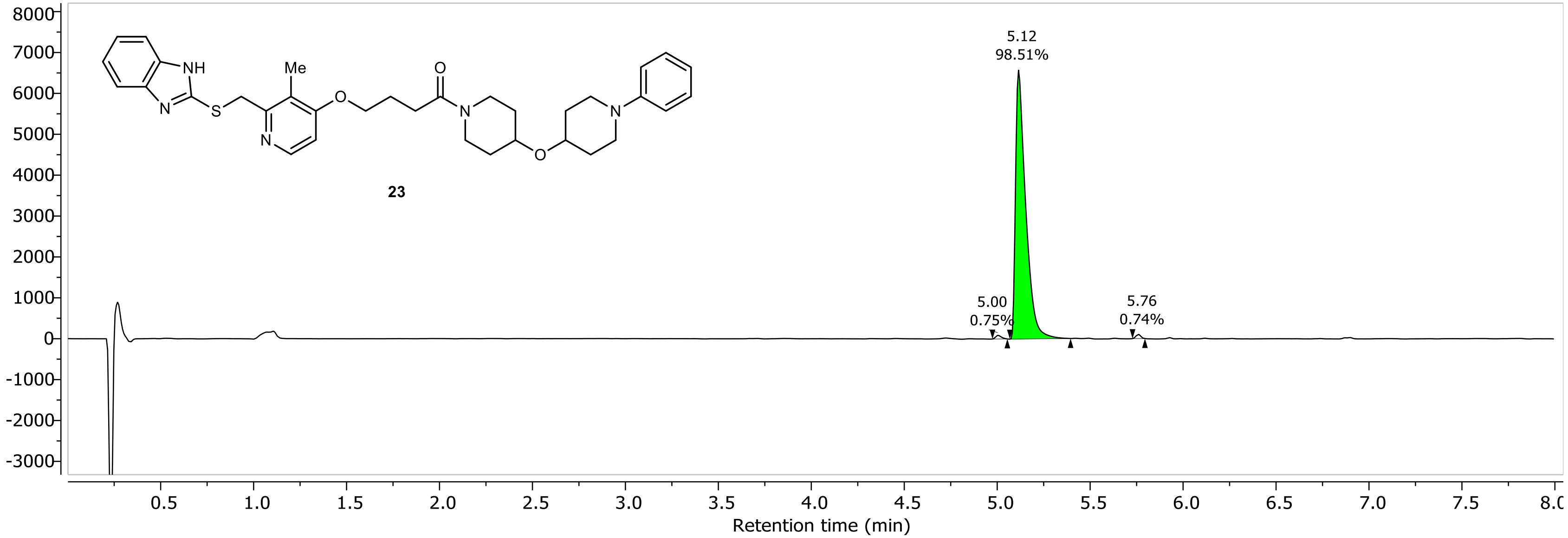

NR-5-113-A LC-MS_ELS DAD1A, Sig=254...Ref=360.0,100.0 Chromatogram

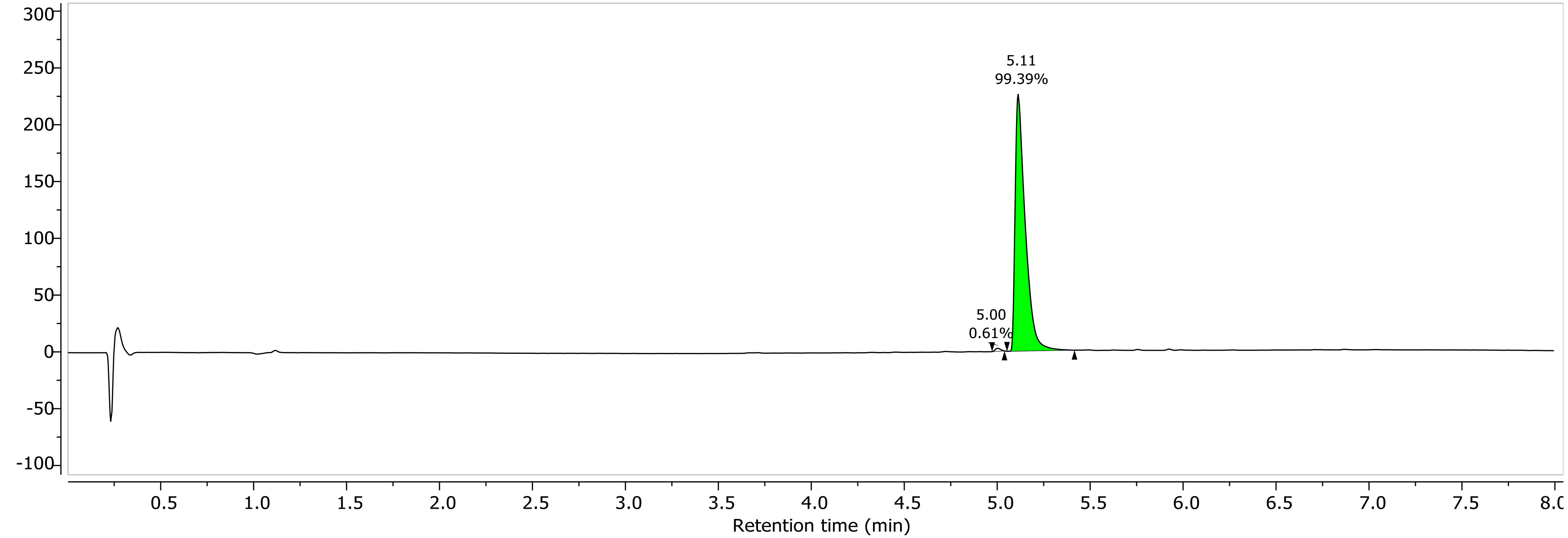

1H NMR (400.26 MHz, CDCl3, 298.1 K)

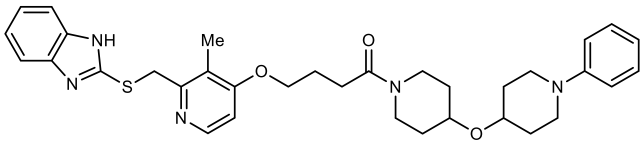

**23**

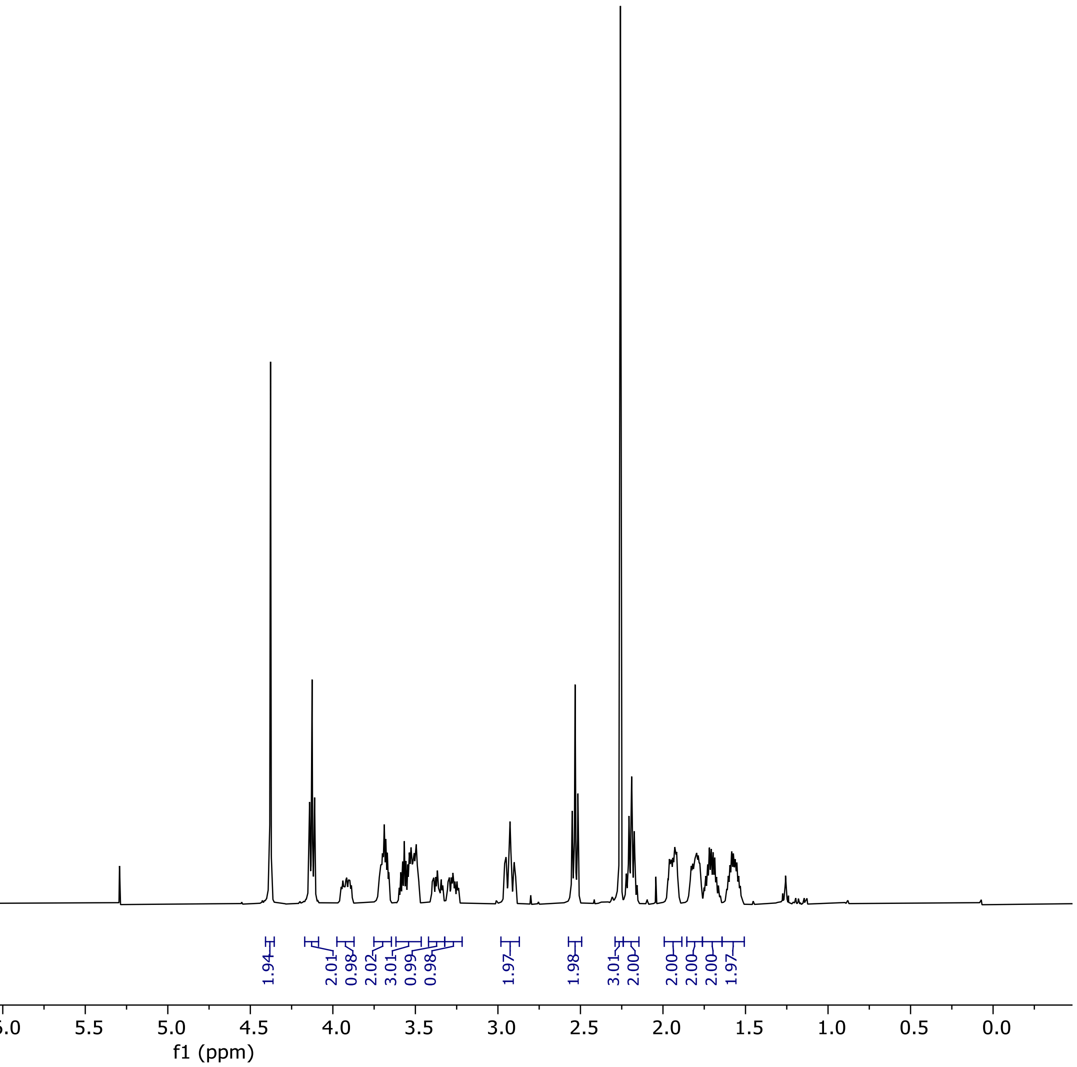

13C NMR (100.66 MHz, CDCl3, 298.1 K)

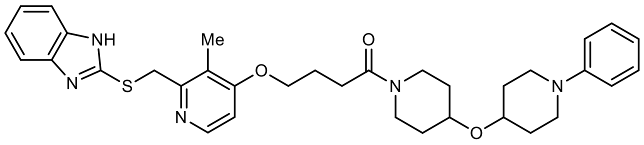

23

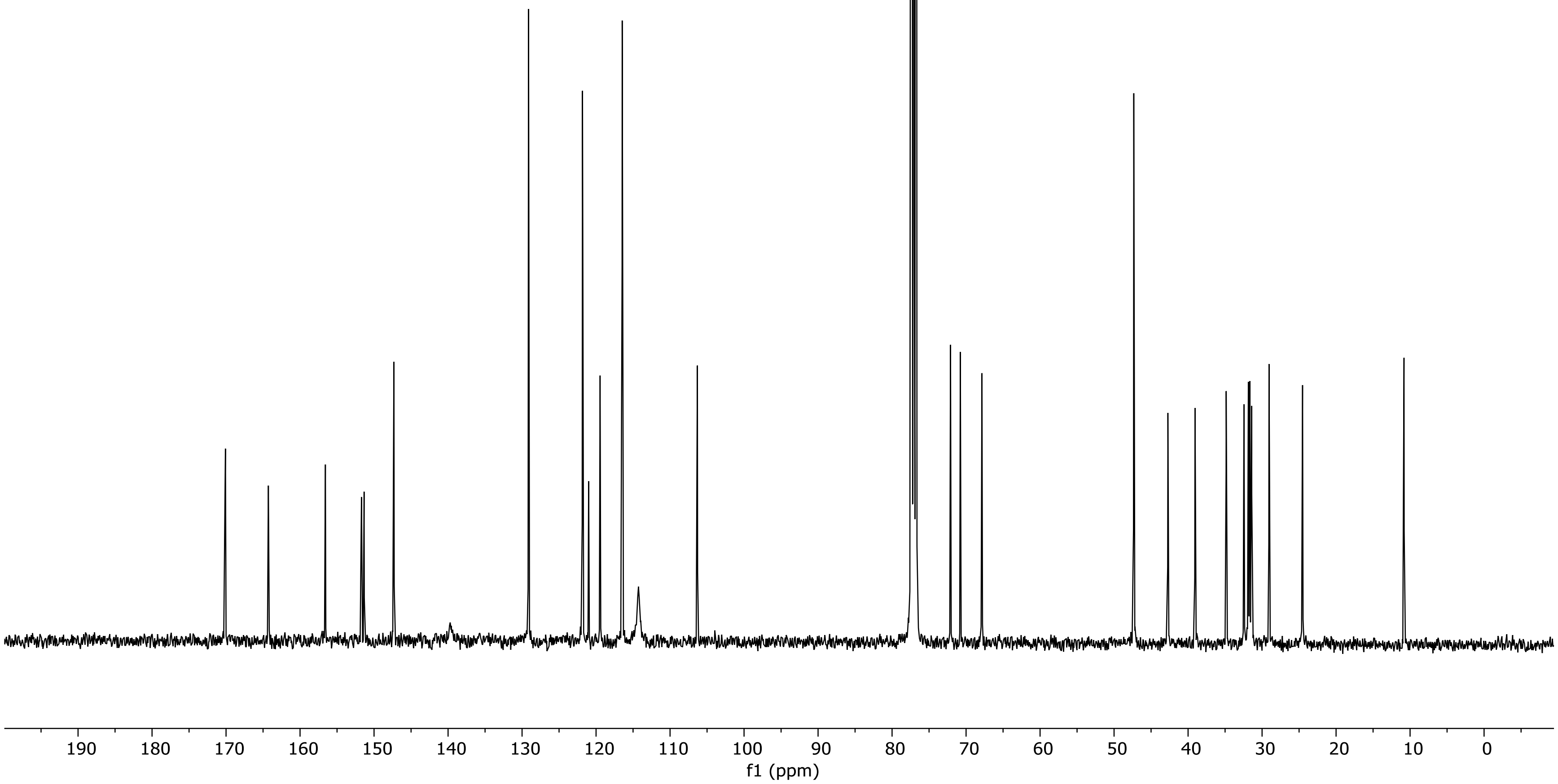

NR-6-129-A LC-MS_ELS  PDA - Total Absorbance Chromatogram

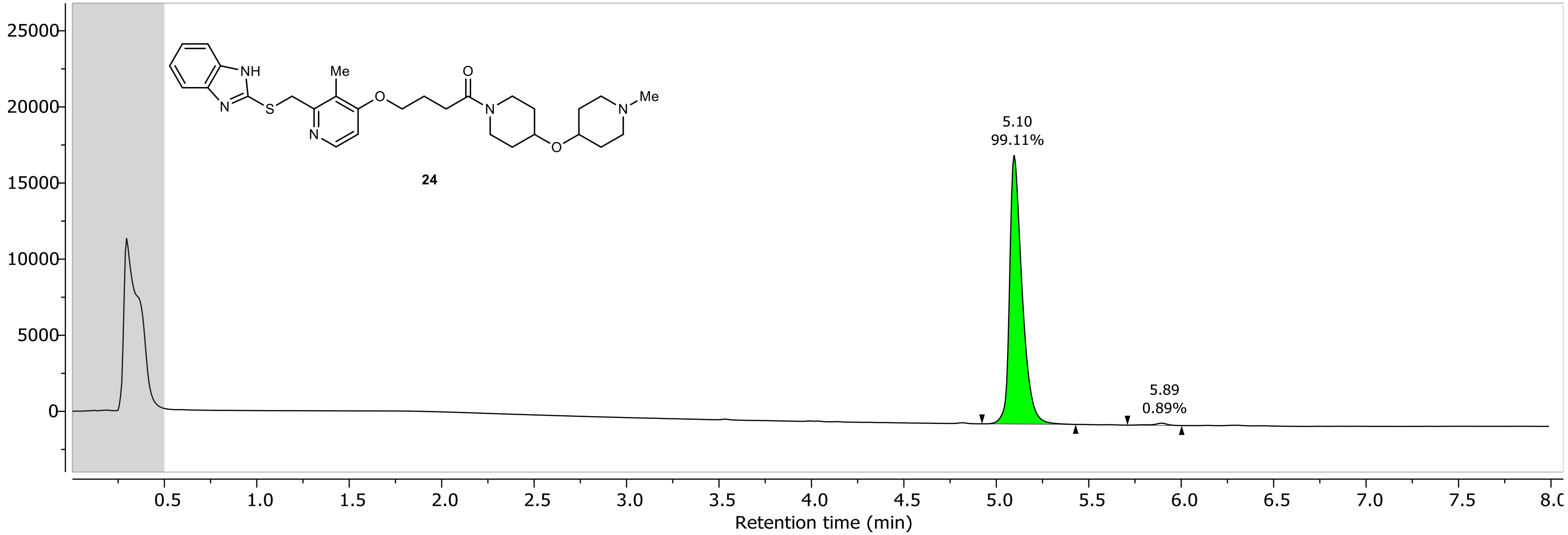

NR-6-129-A LC-MS_ELS DAD1A, Sig=254...Ref=360.0,100.0 Chromatogram

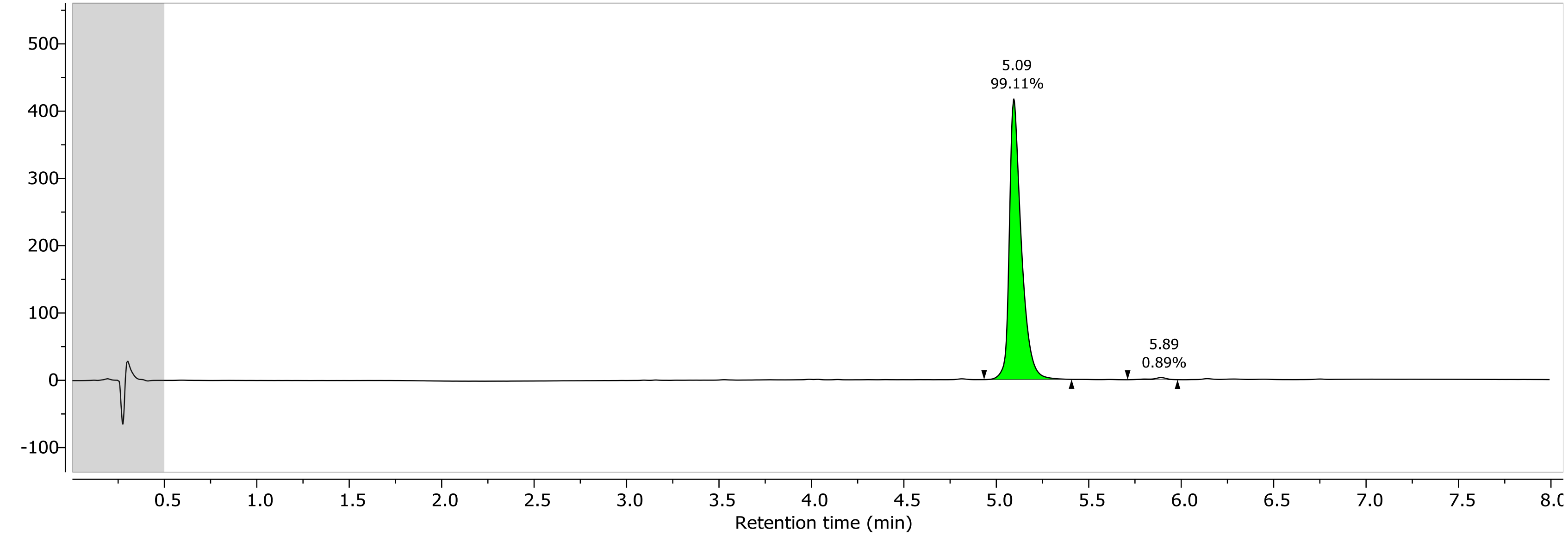

1H NMR (600.15 MHz, DMSO, 298.0 K)

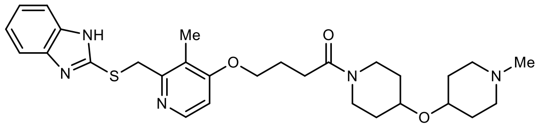

**24**

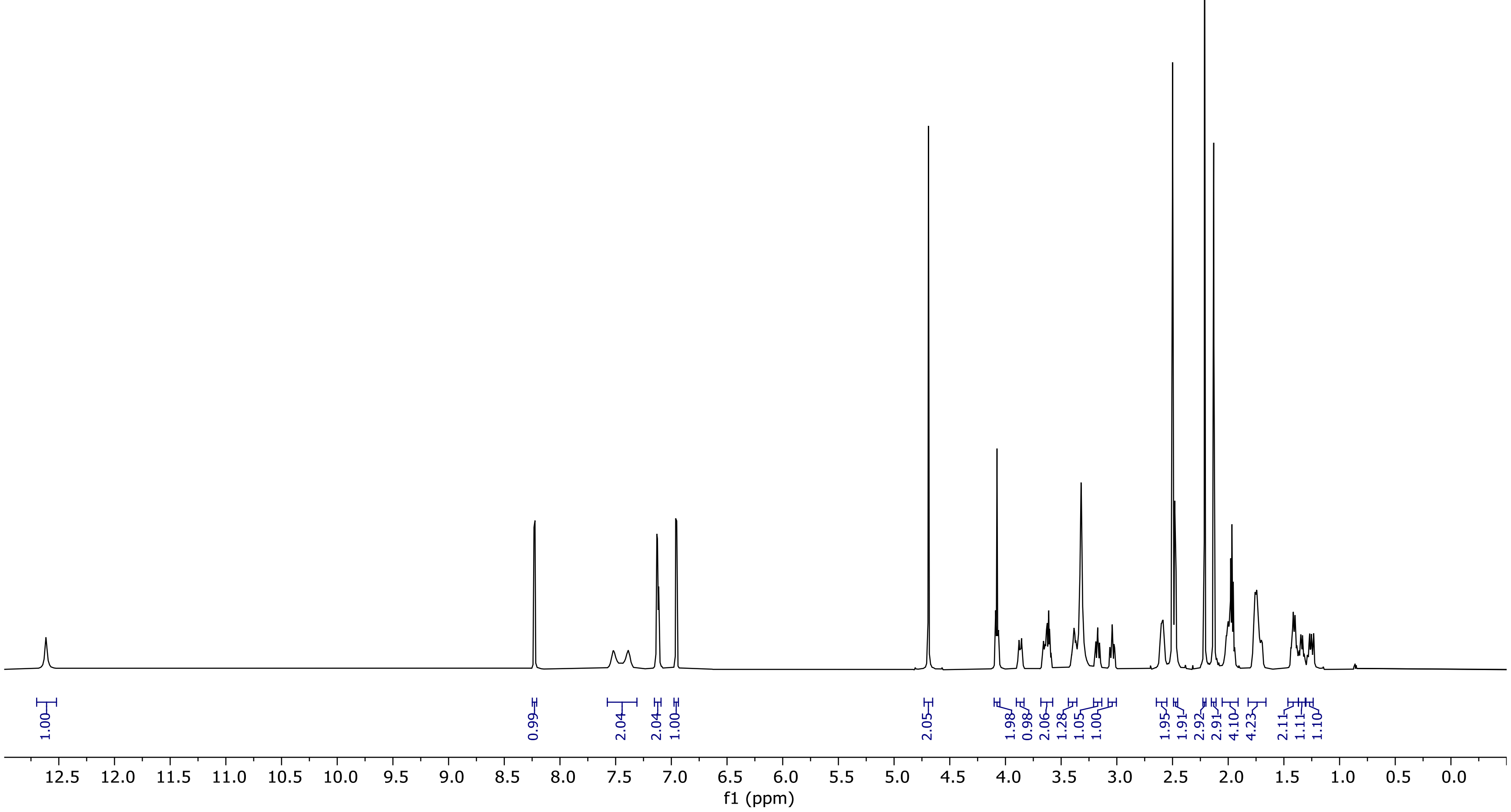

13C NMR (100.66 MHz, DMSO, 298.1 K)

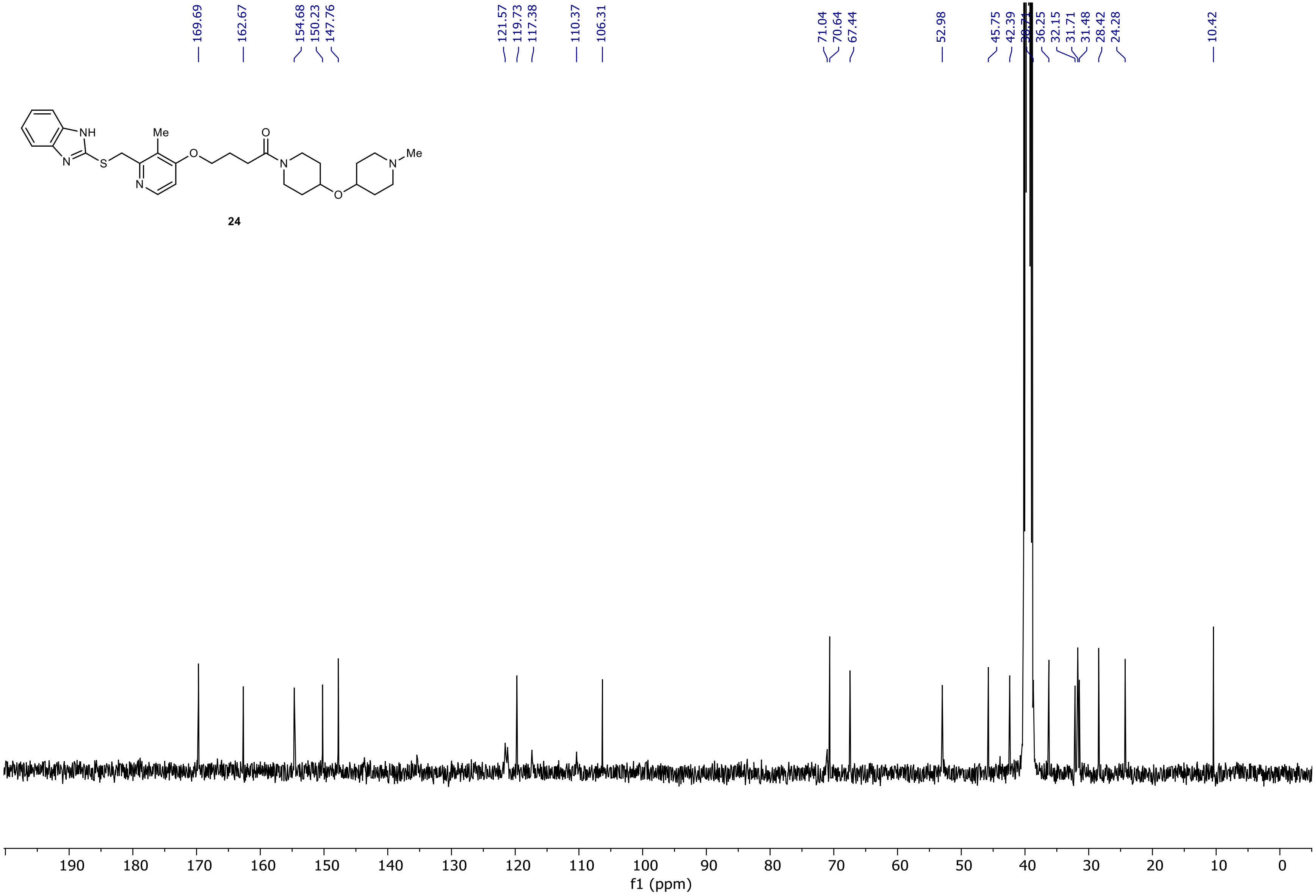

NR-6-064-A LC-MS_ELS  PDA - Total Absorbance Chromatogram

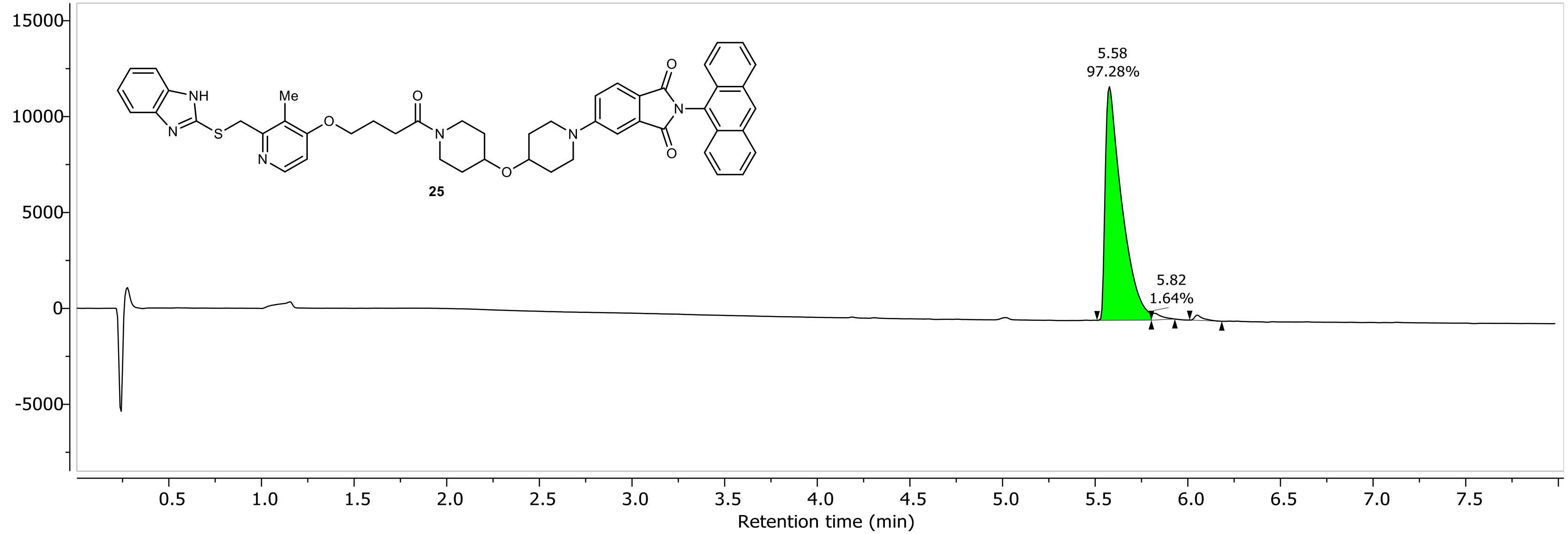

NR-6-064-A LC-MS_ELS DAD1A, Sig=254...Ref=360.0,100.0 Chromatogram

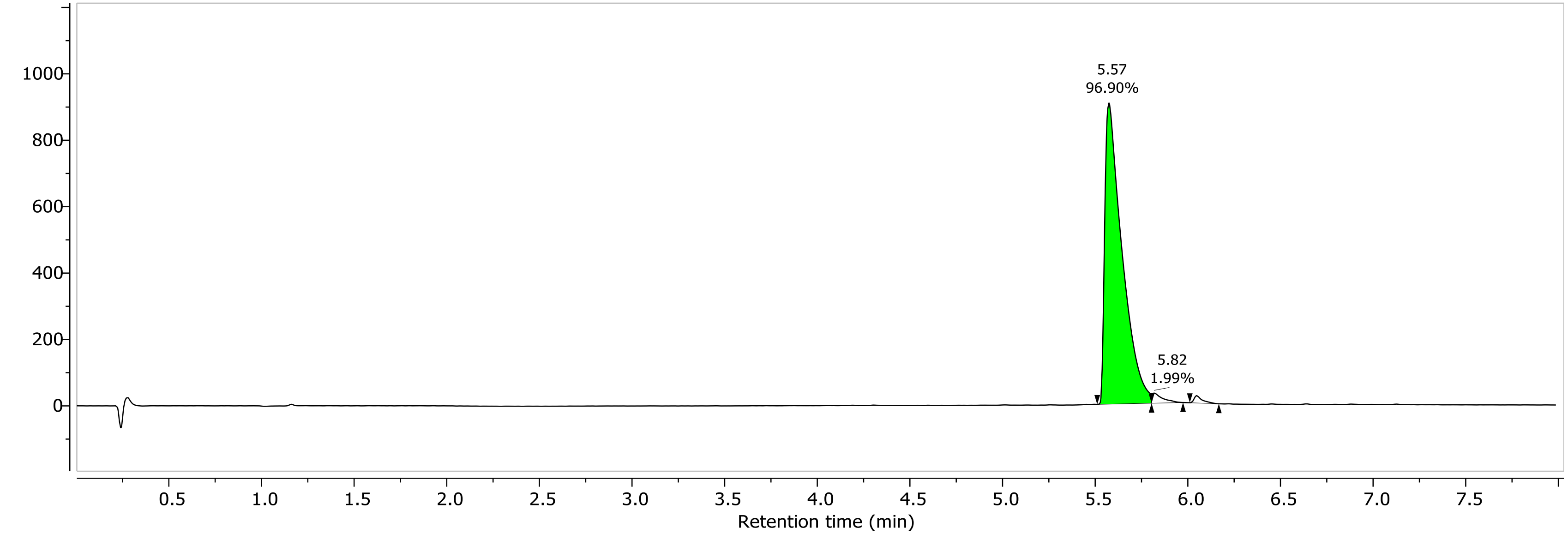

1H NMR (400.26 MHz, CDCl3, 298.0 K)

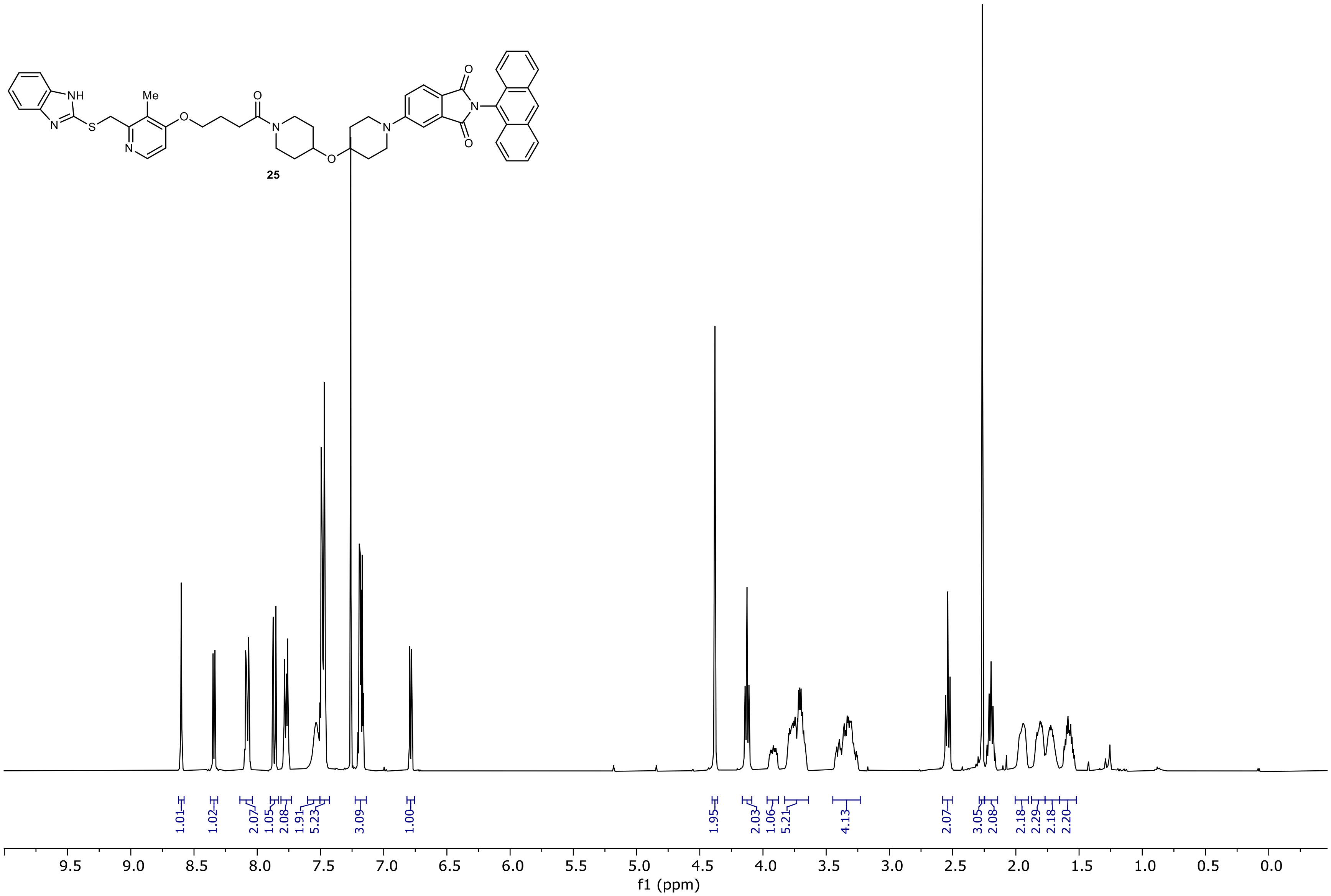

13C NMR (100.66 MHz, CDCl3, 298.1 K)

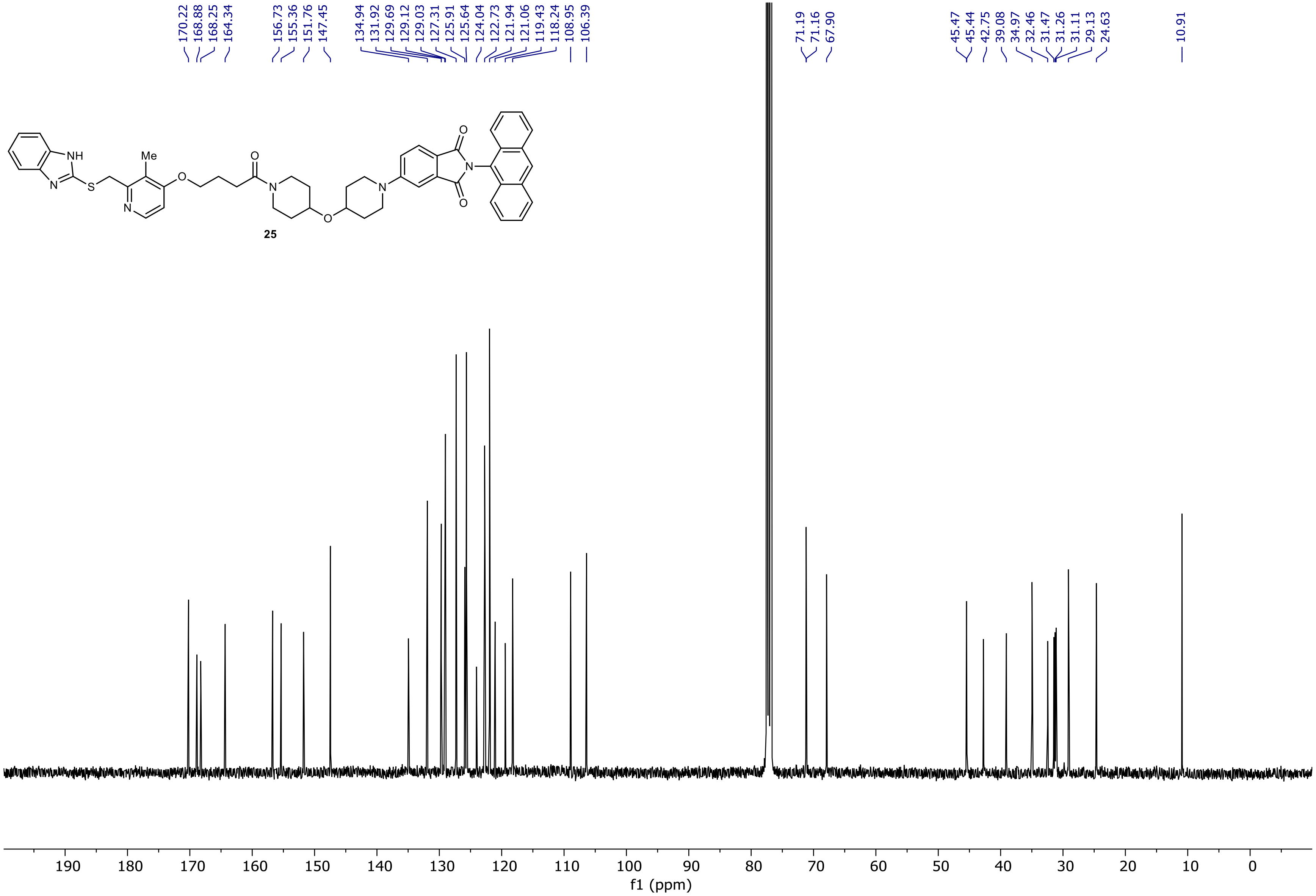

NR-6-022-A LC-MS_ELS PDA - Total Absorbance Chromatogram

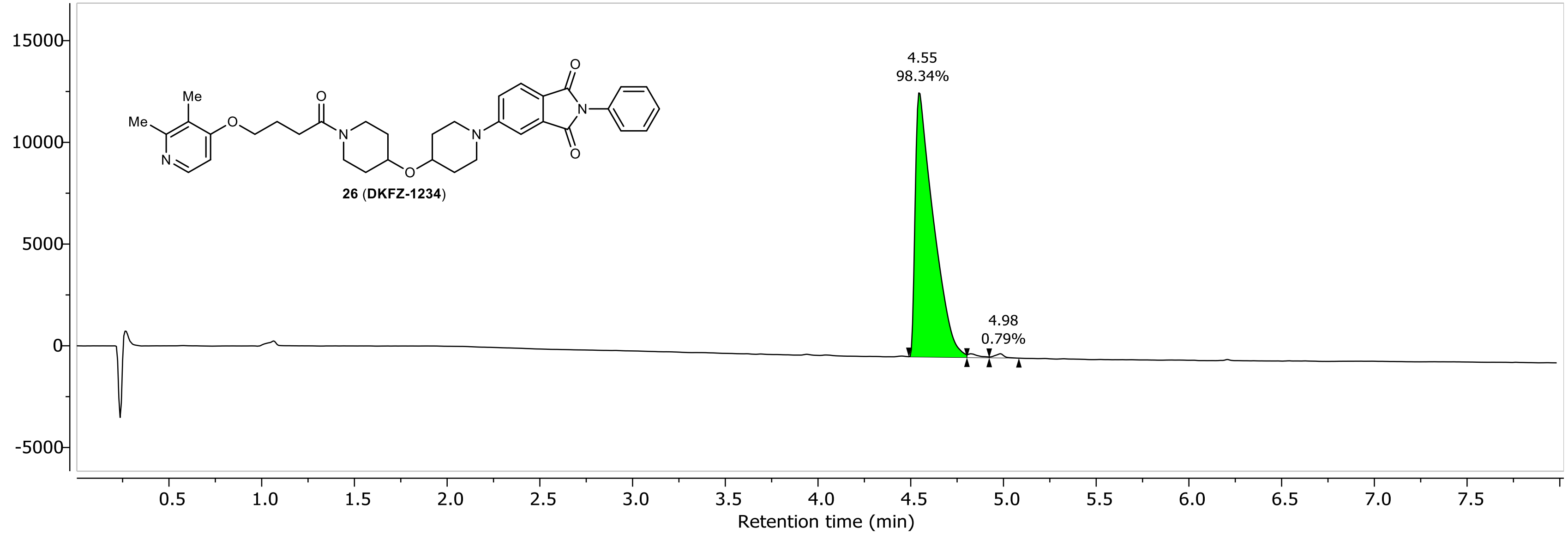

NR-6-022-A LC-MS_ELS DAD1A, Sig=254...Ref=360.0,100.0 Chromatogram

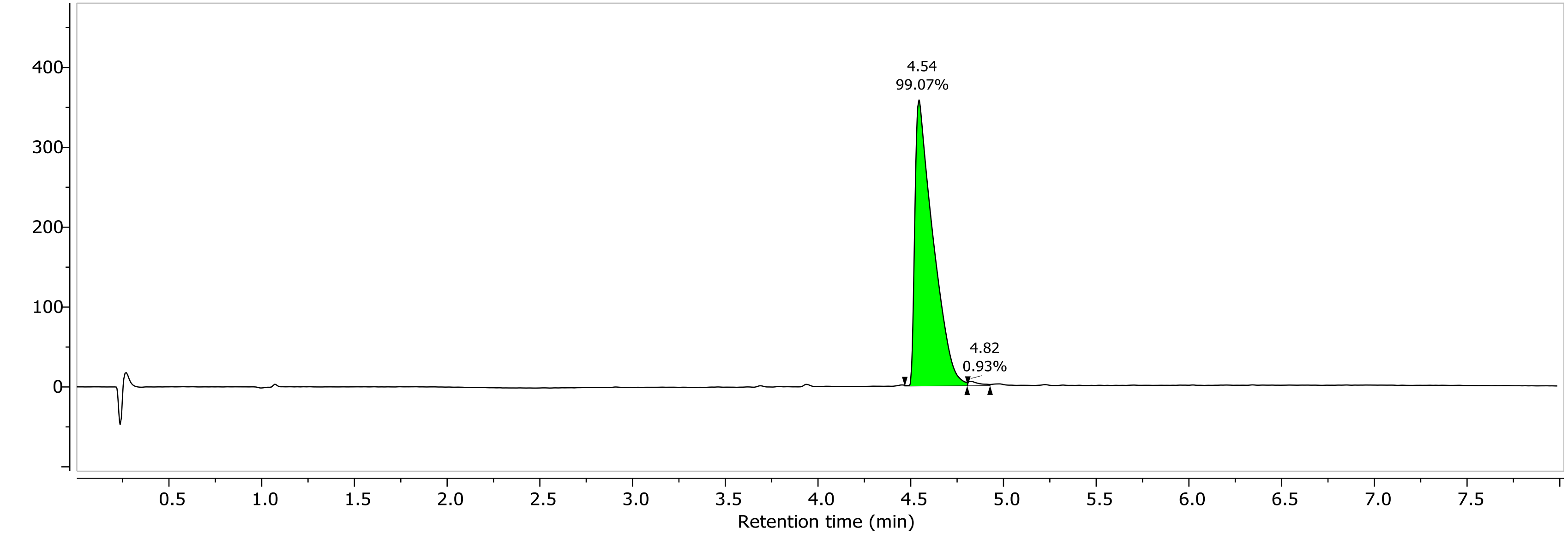

1H NMR (400.26 MHz, CDCl3, 298.1 K)

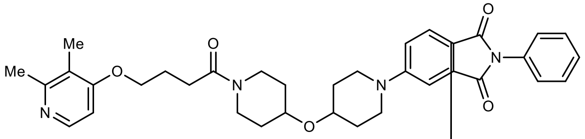

**26** (**DKFZ-1234**)

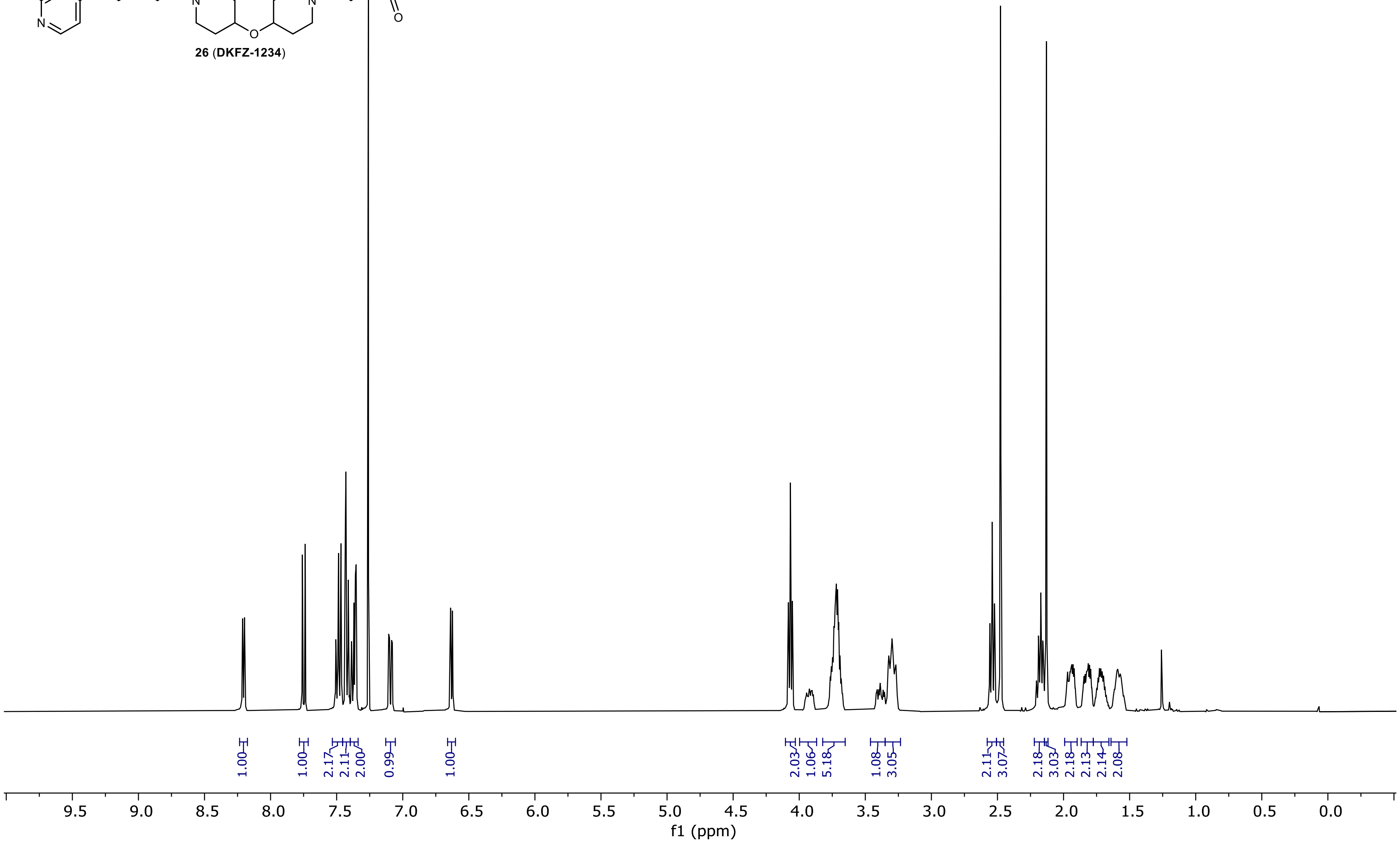

13C NMR (100.66 MHz, CDCl3, 298.1 K)

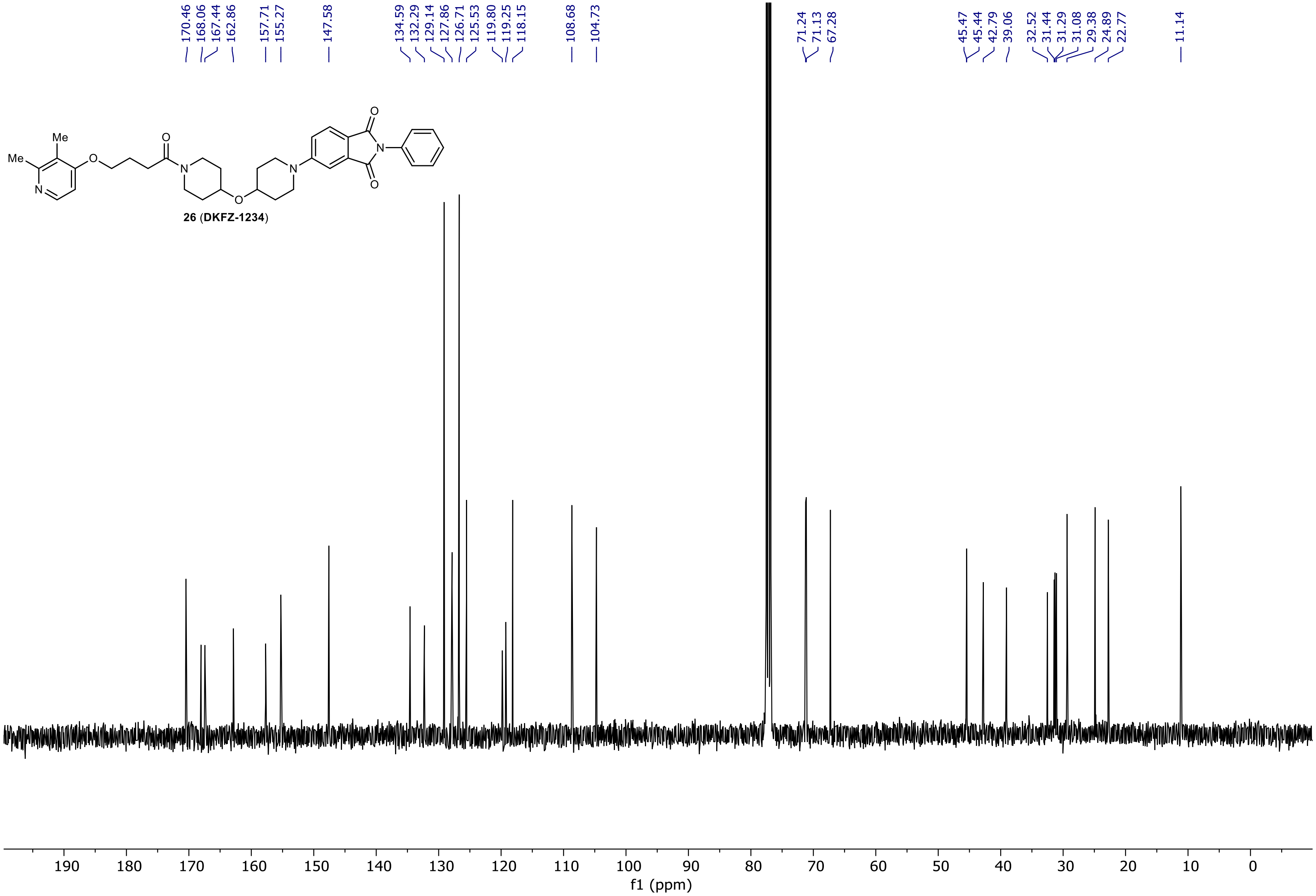

NR-6-026-A LC-MS_ELS  PDA - Total Absorbance Chromatogram

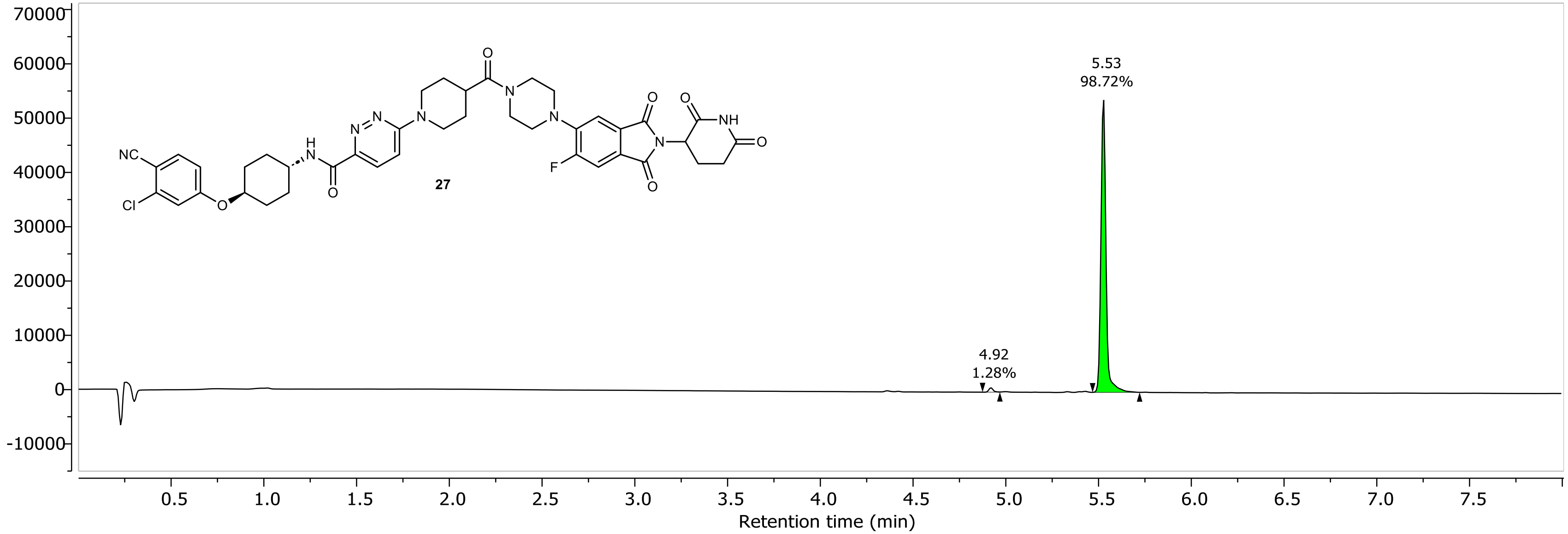

NR-6-026-A LC-MS_ELS DAD1A, Sig=254...Ref=360.0,100.0 Chromatogram

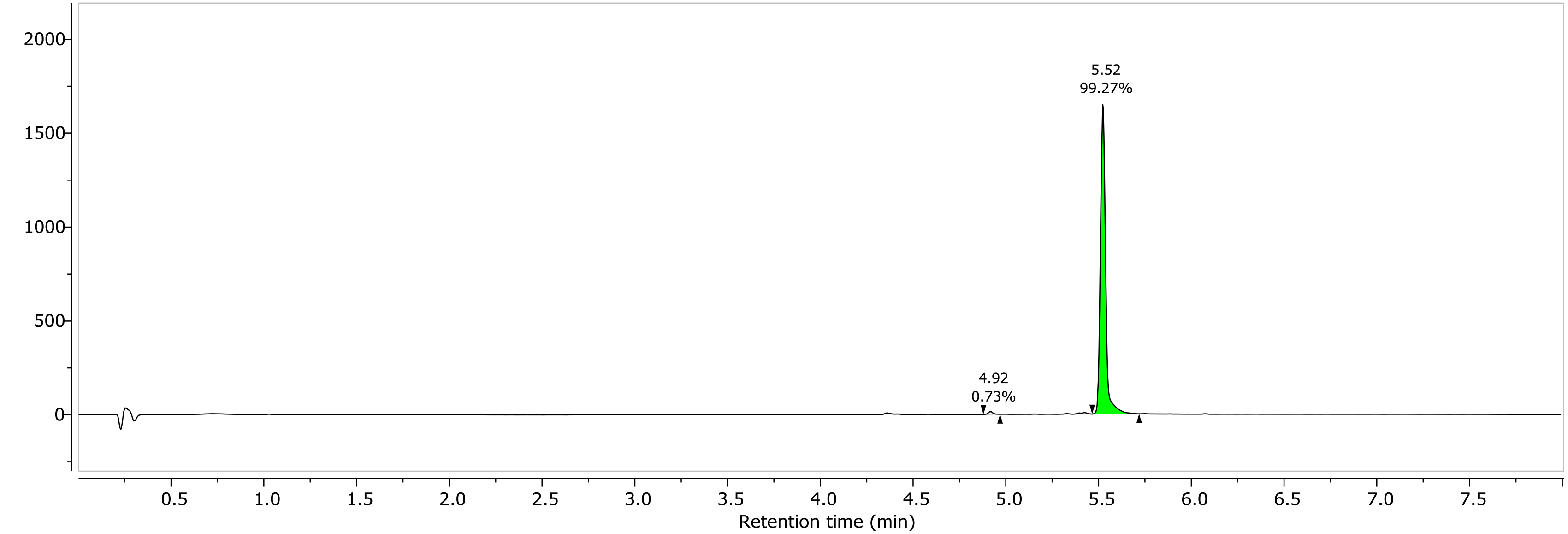

1H NMR (400.26 MHz, CDCl3, 298.1 K)

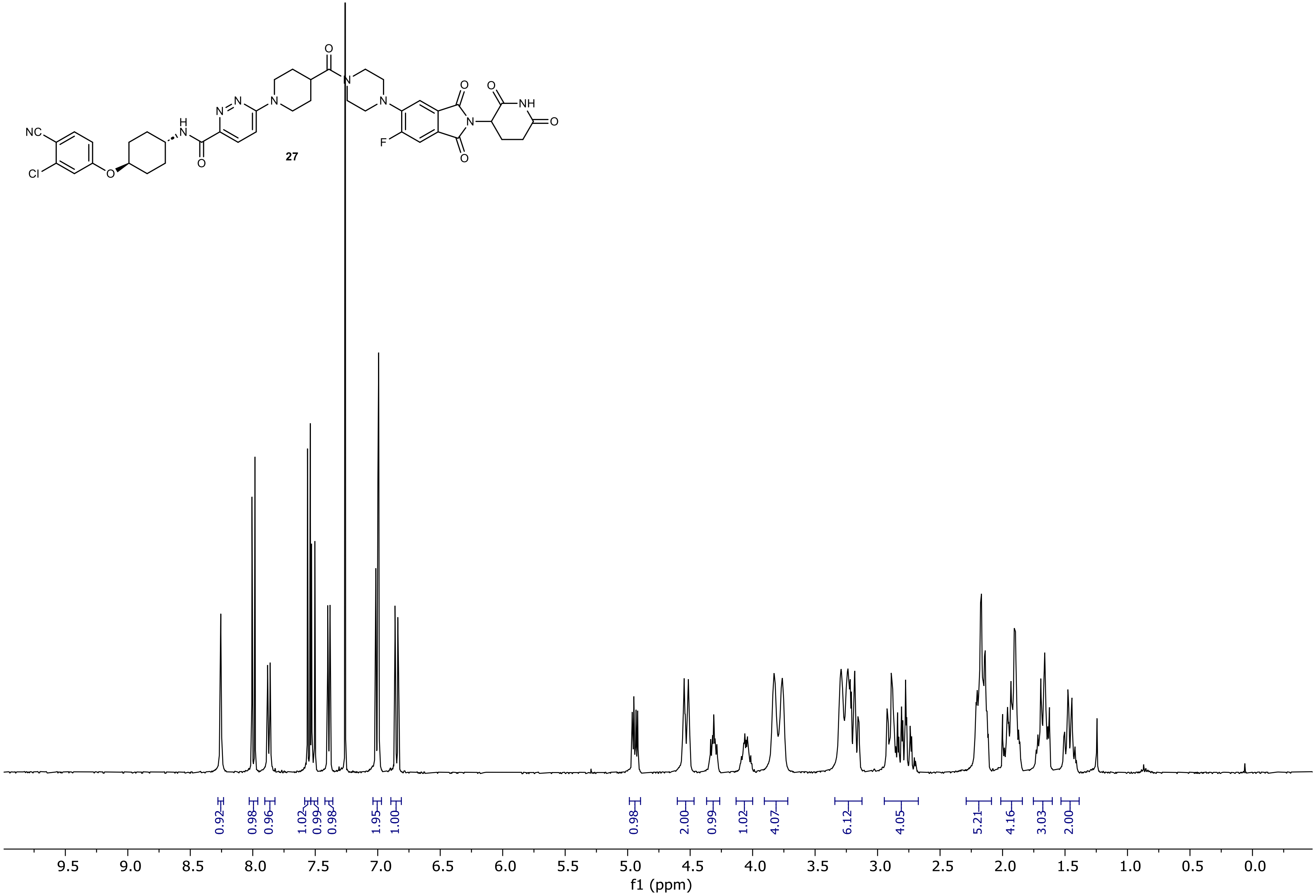

13C NMR (100.66 MHz, CDCl3, 298.1 K)

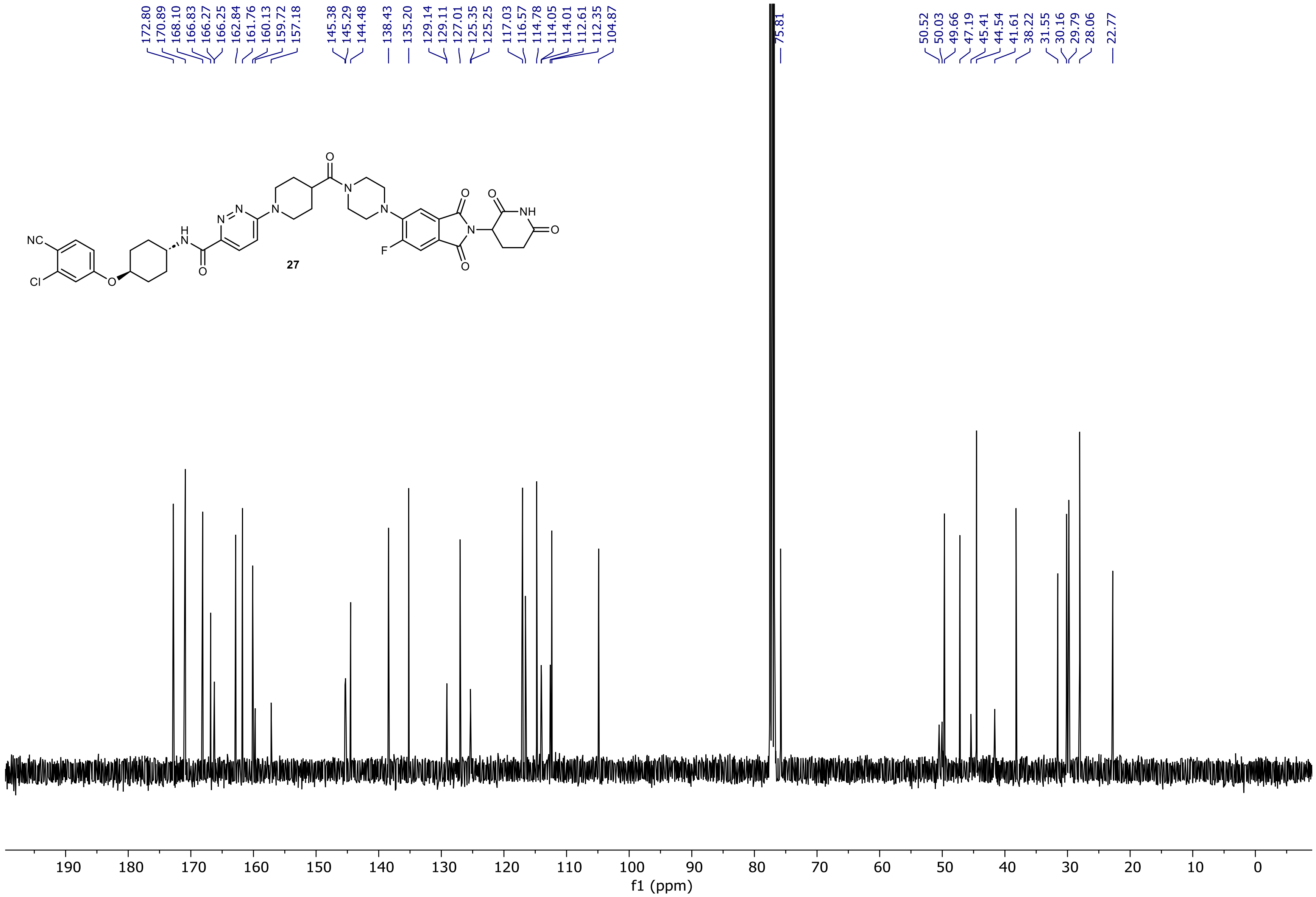

19F NMR (376.58 MHz, CDCl3, 298.2 K)

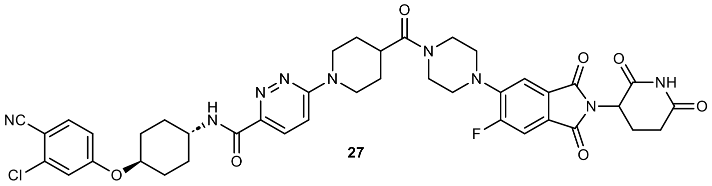

27

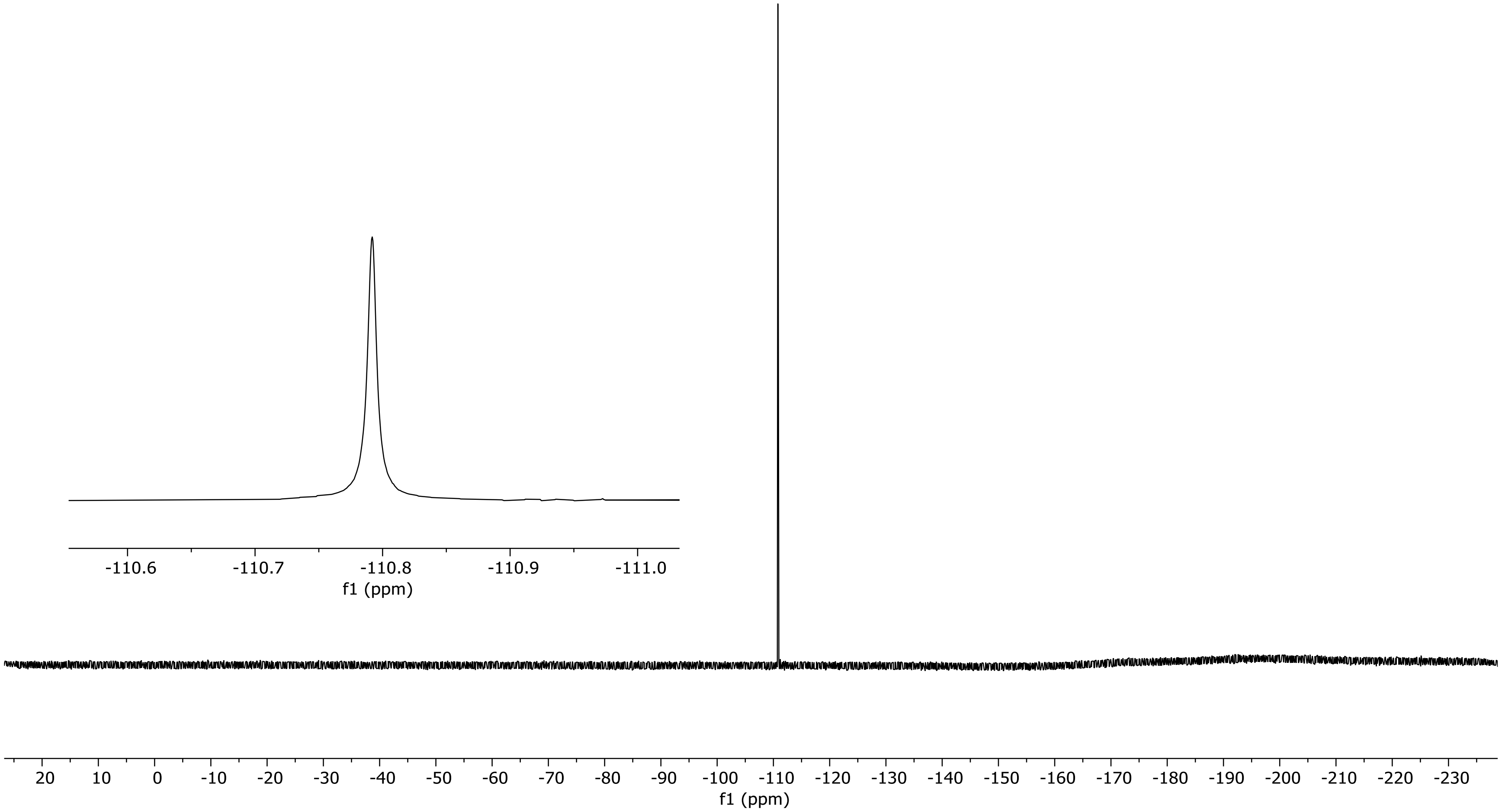

NR-6-054-B LC-MS_ELS PDA - Total Absorbance Chromatogram

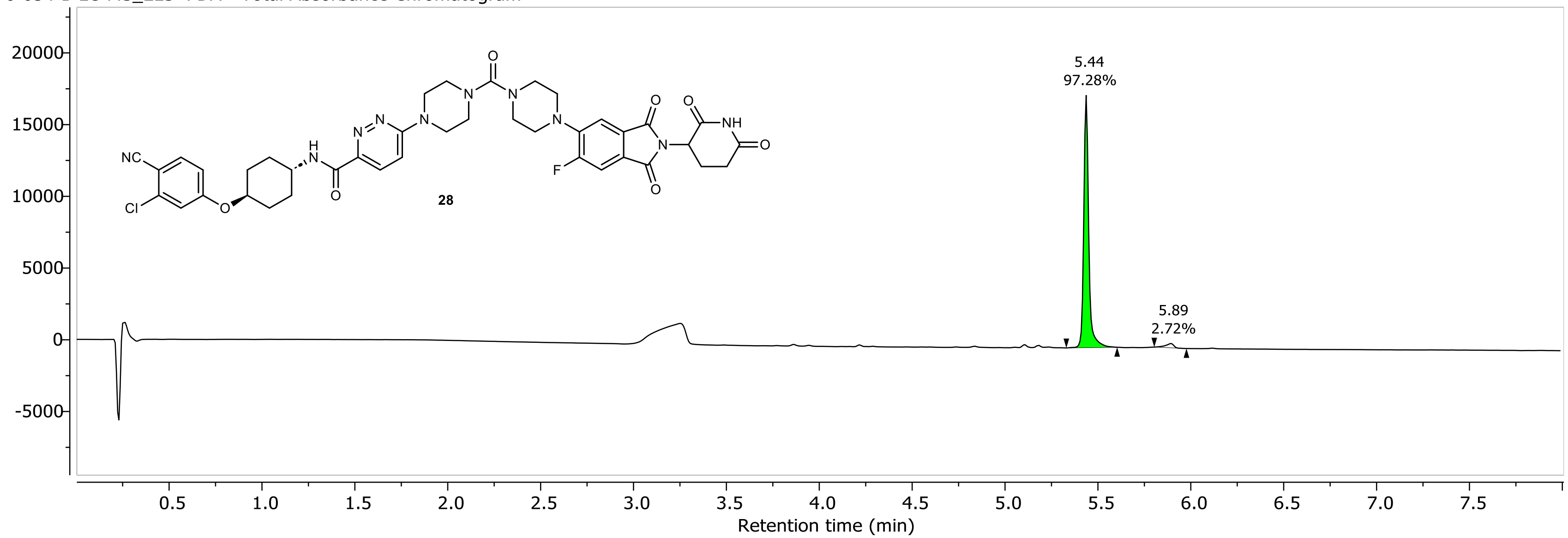

NR-6-054-B LC-MS_ELS DAD1A, Sig=254...Ref=360.0,100.0 Chromatogram

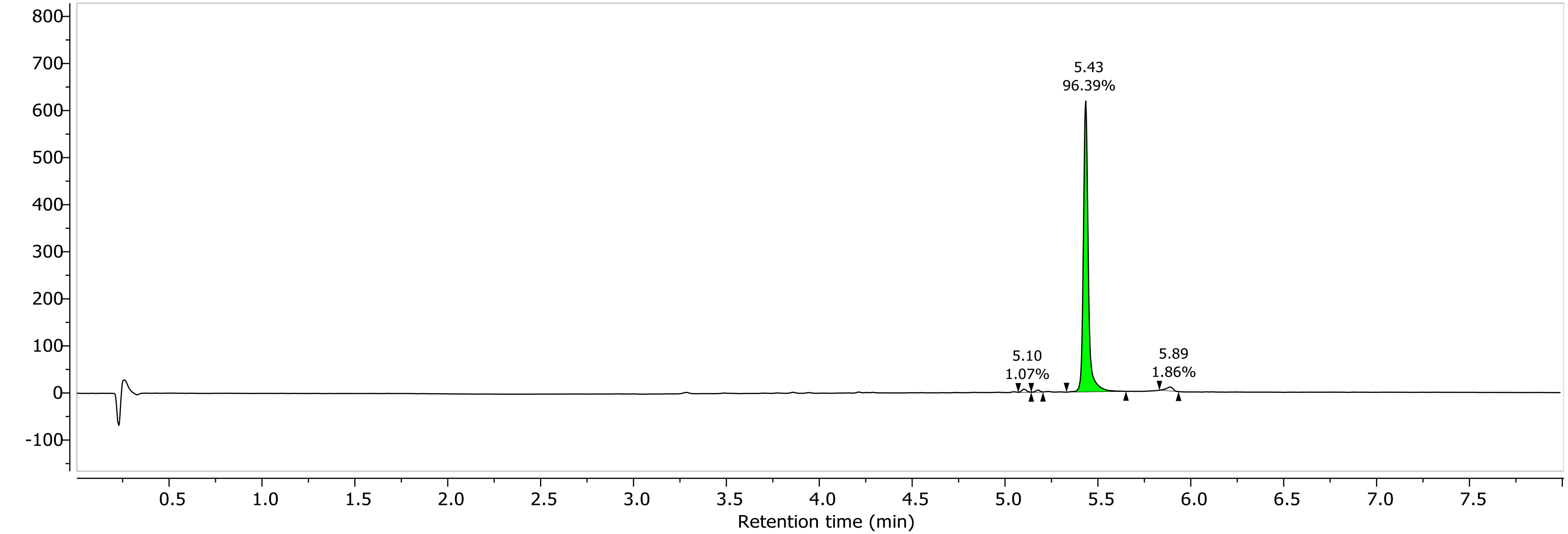

1H NMR (400.26 MHz, CDCl3, 298.2 K)

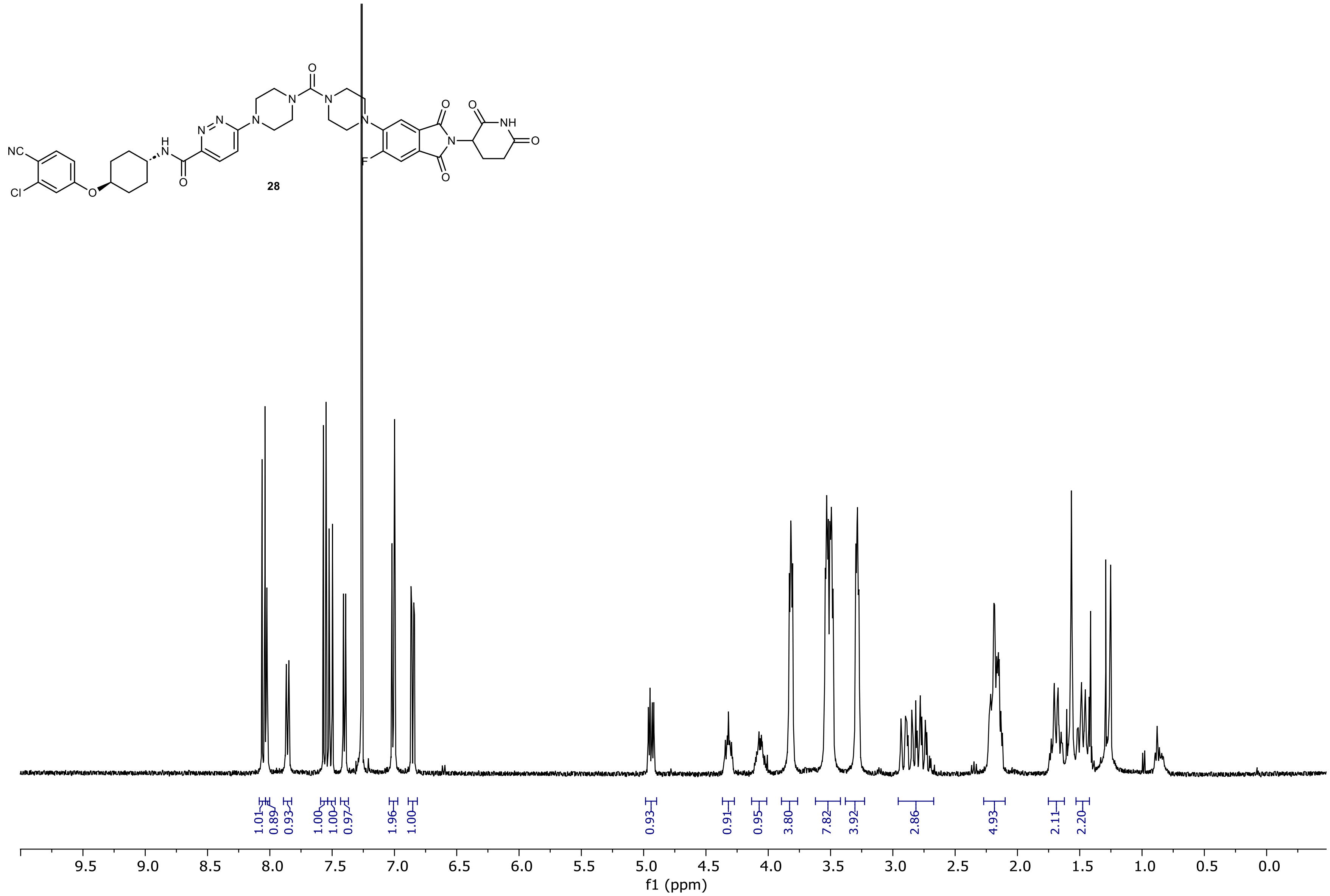

13C NMR (150.92 MHz, CDCl3, 298.0 K)

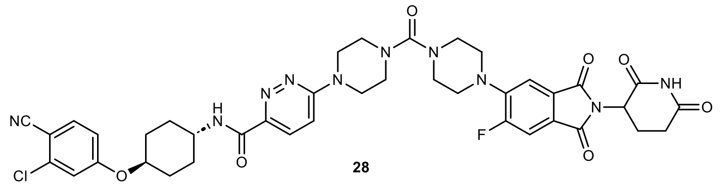

28

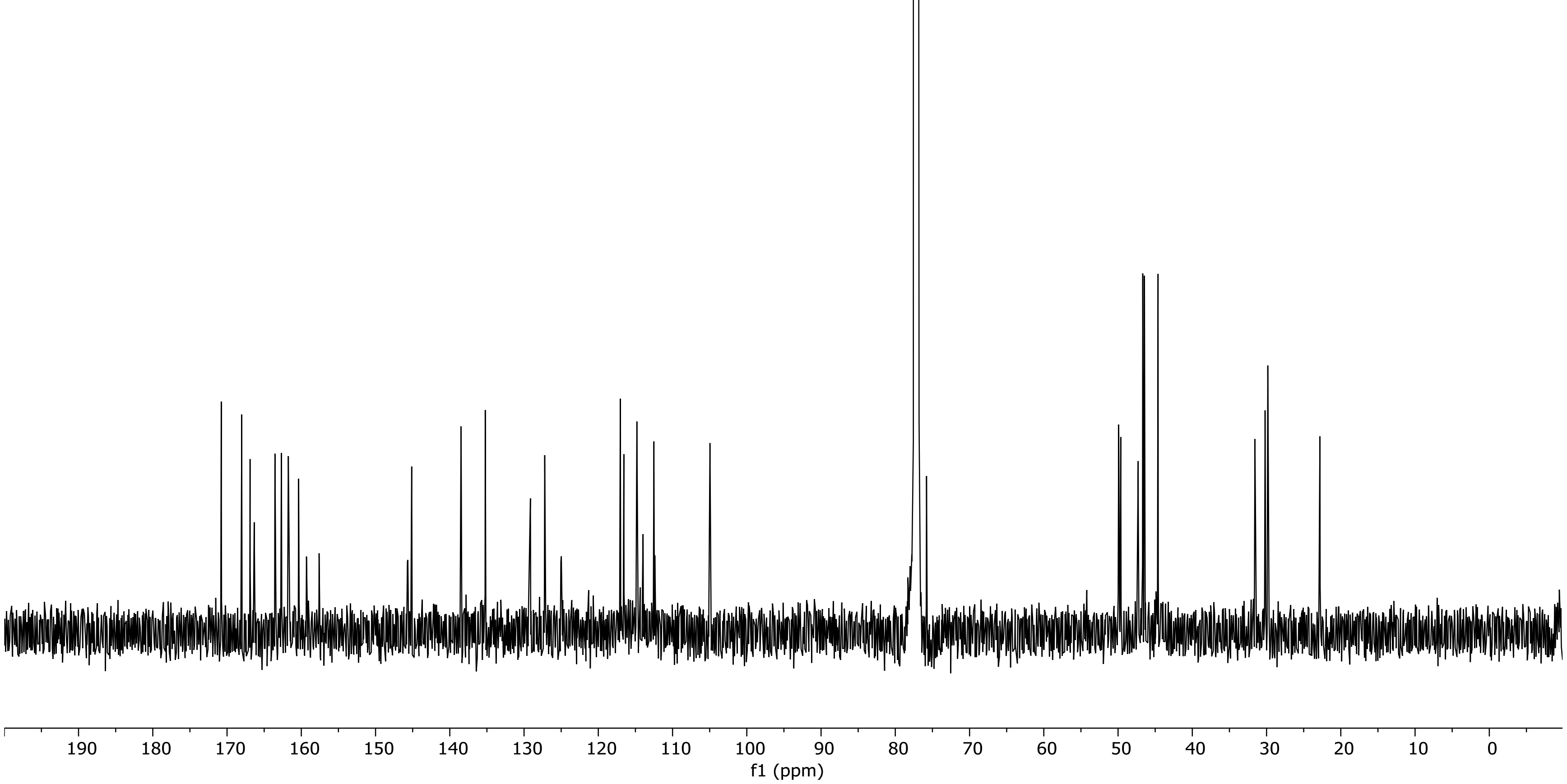

19F NMR (376.58 MHz, CDCl3, 298.0 K)

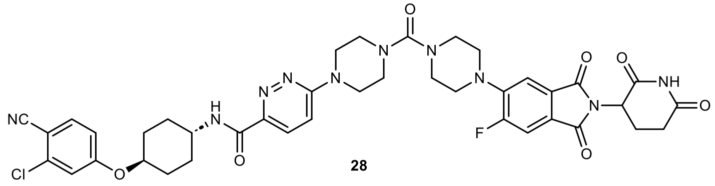

28

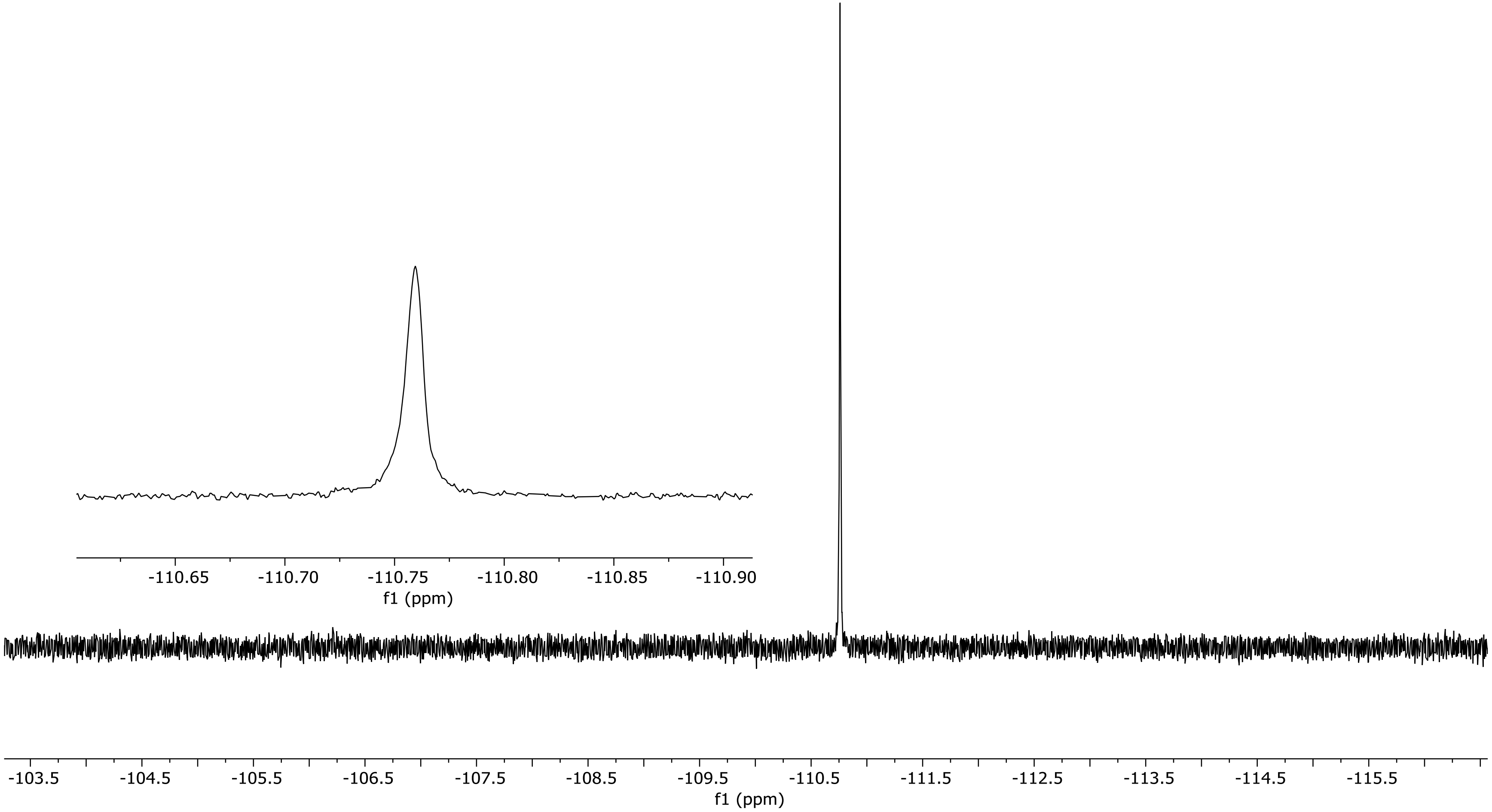

NR-6-042-A LC-MS_ELS PDA - Total Absorbance Chromatogram

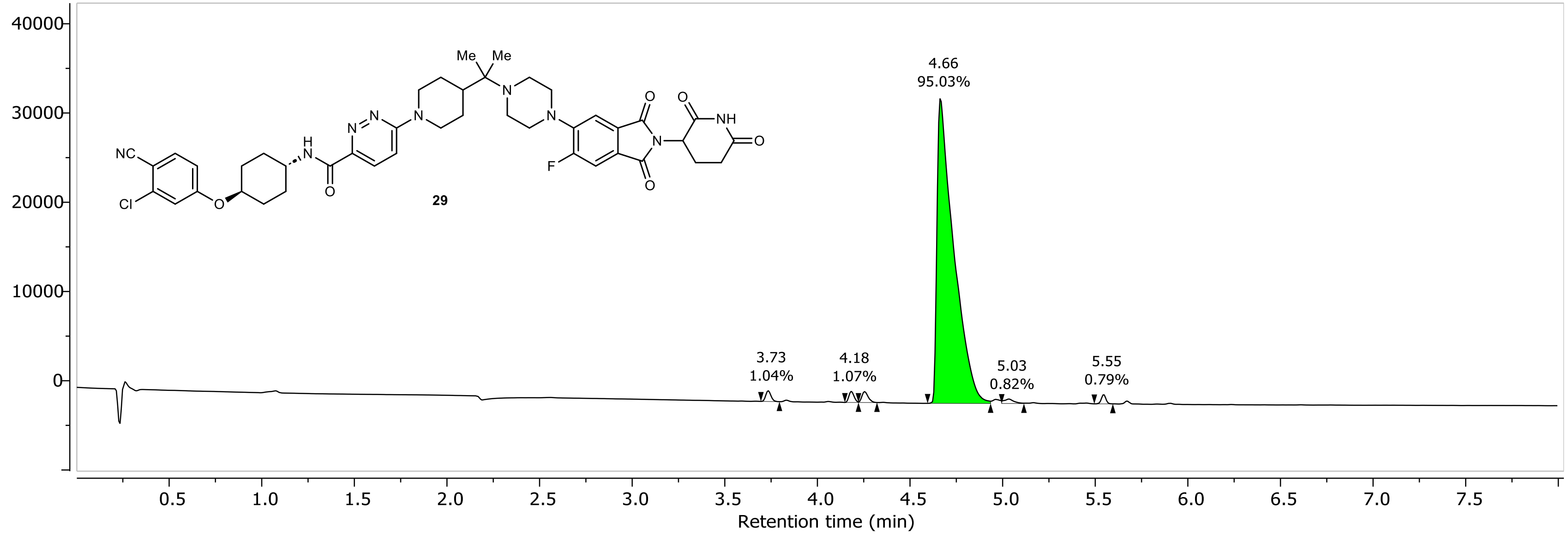

NR-6-042-A LC-MS_ELS DAD1A, Sig=254...Ref=360.0,100.0 Chromatogram

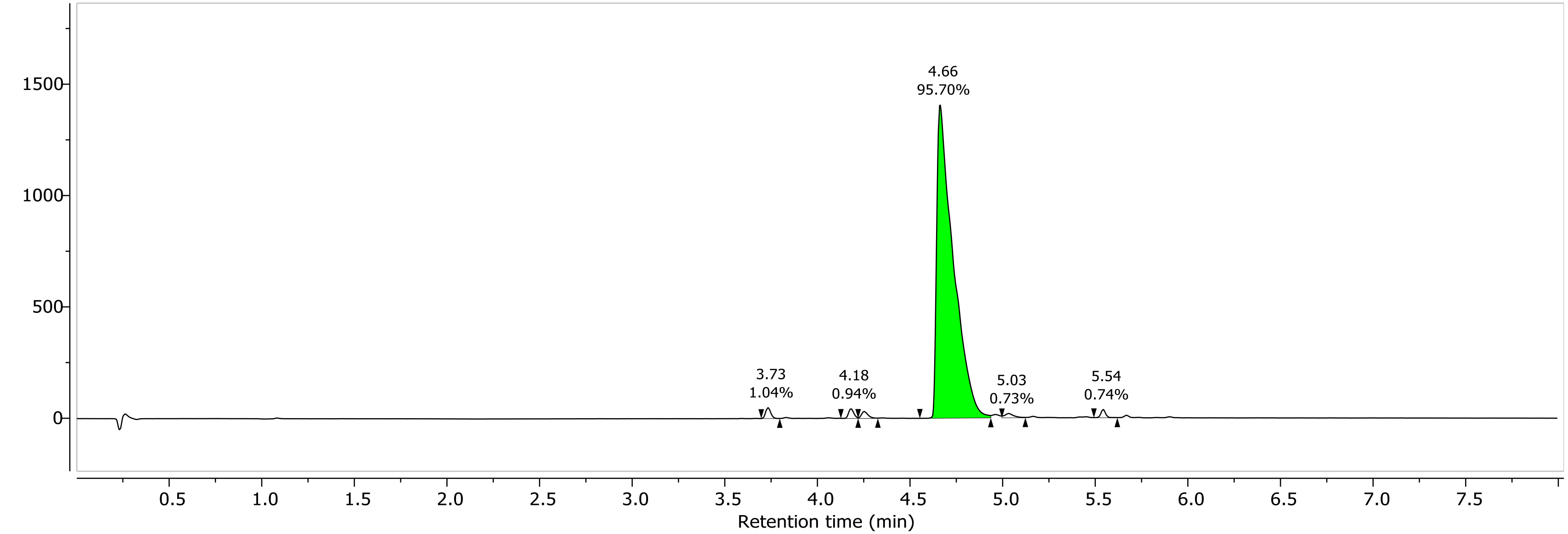

1H NMR (400.26 MHz, CDCl3, 298.1 K)

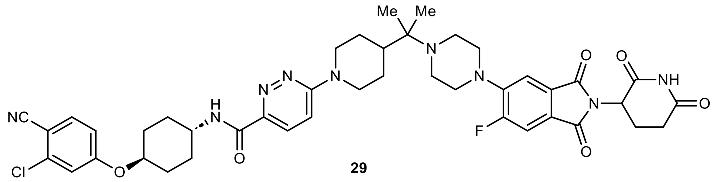

29

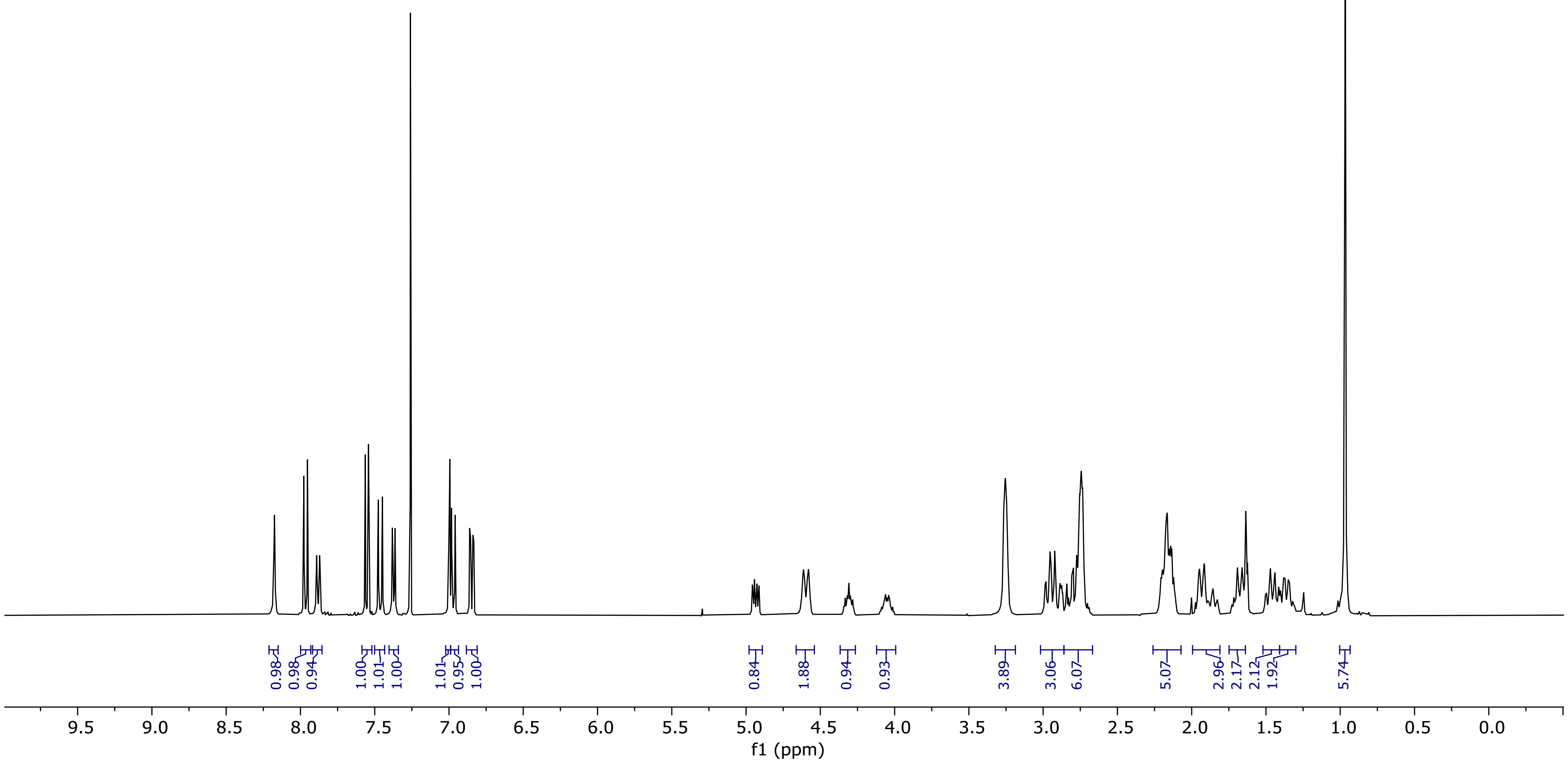

13C NMR (100.66 MHz, CDCl3, 298.1 K)

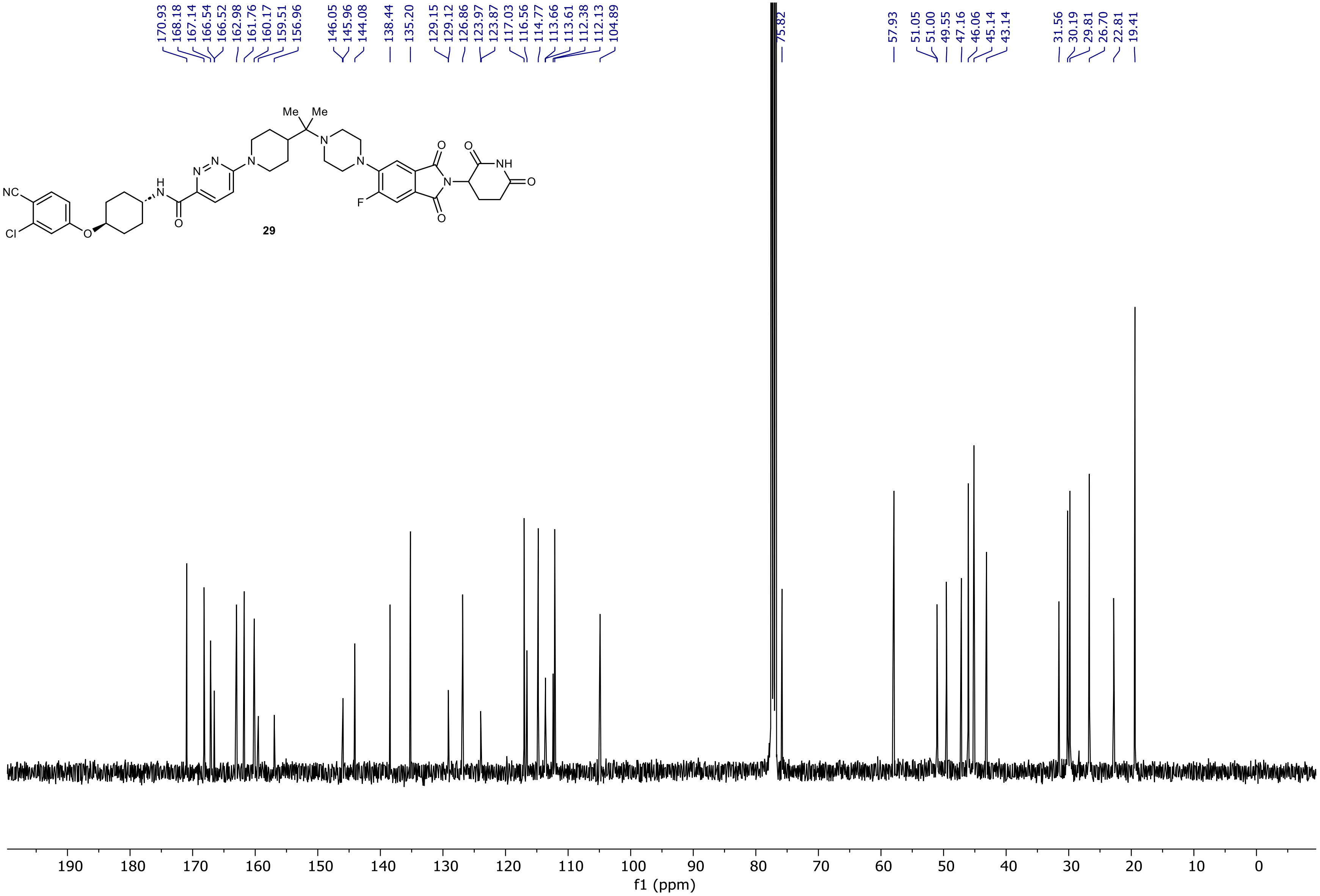

29

19F NMR (376.58 MHz, CDCl3, 298.1 K)

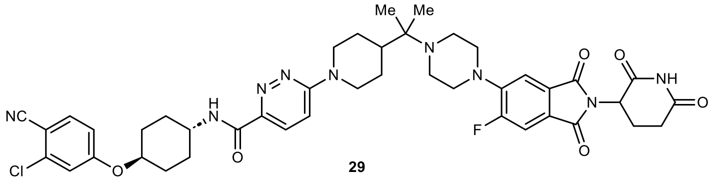

29

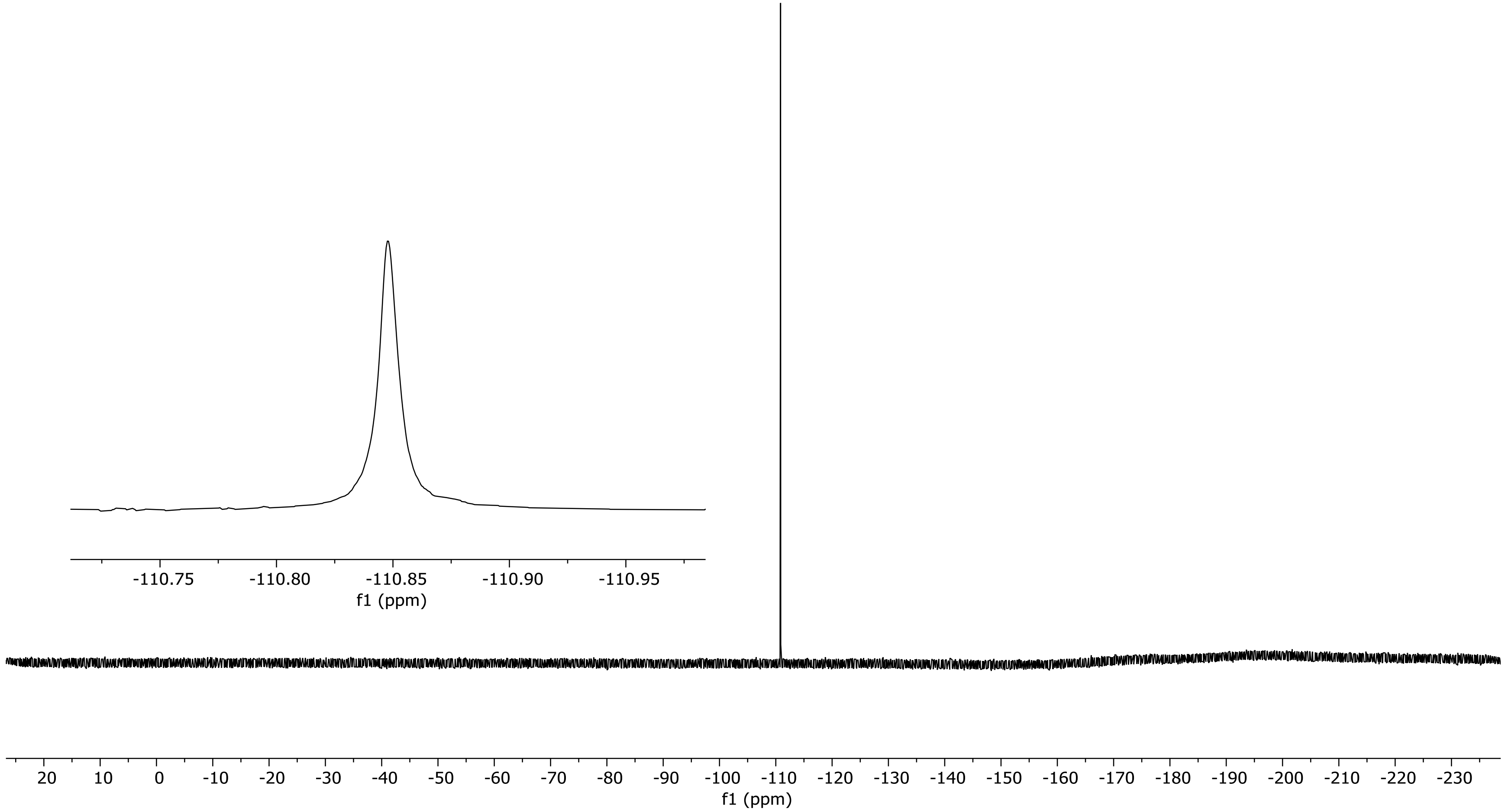

PDA - Total Absorbance Chromatogram

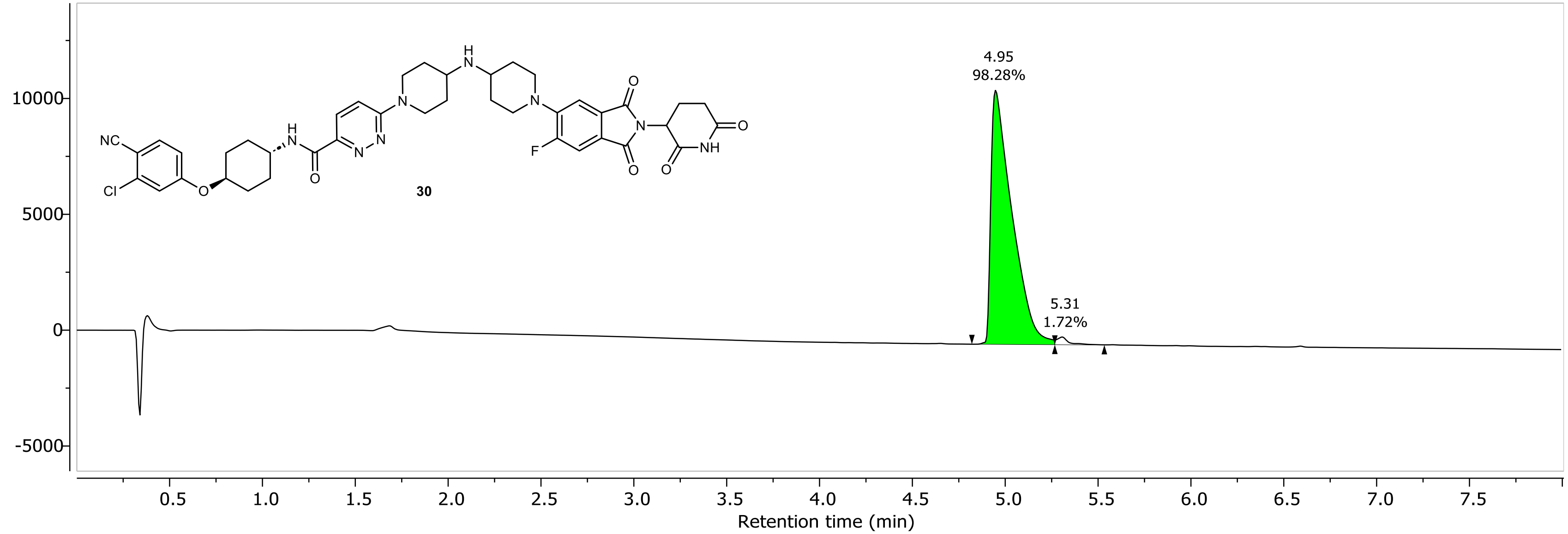

DAD1A, Sig=254...Ref=360.0,100.0 Chromatogram

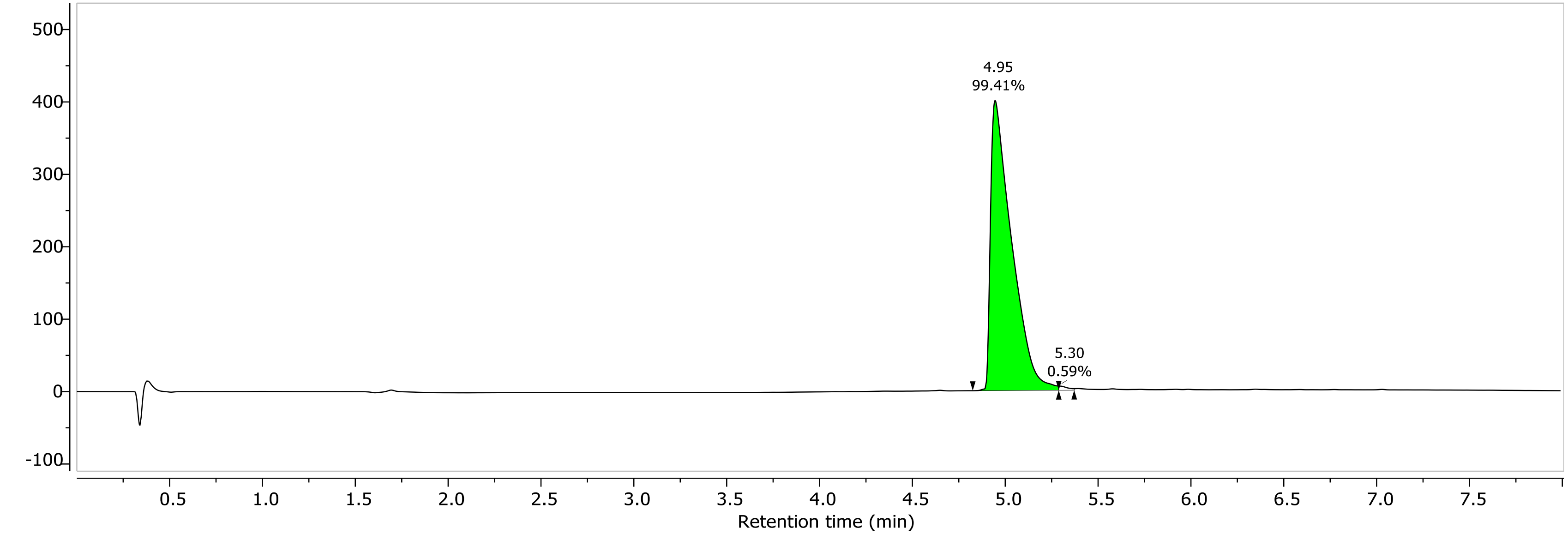

1H NMR (400.26 MHz, CDCl3, 298.1 K)

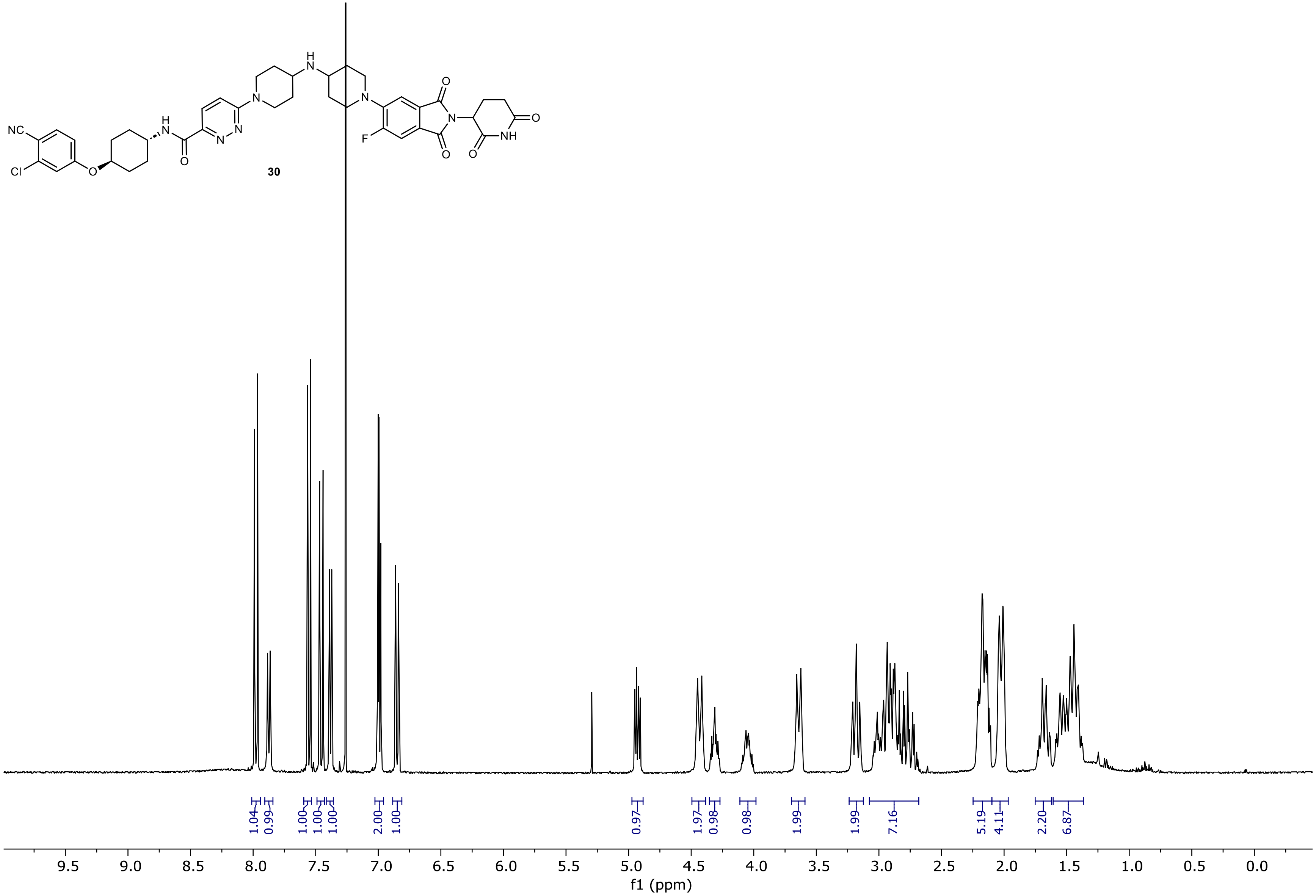

13C NMR (100.66 MHz, CDCl3, 298.1 K)

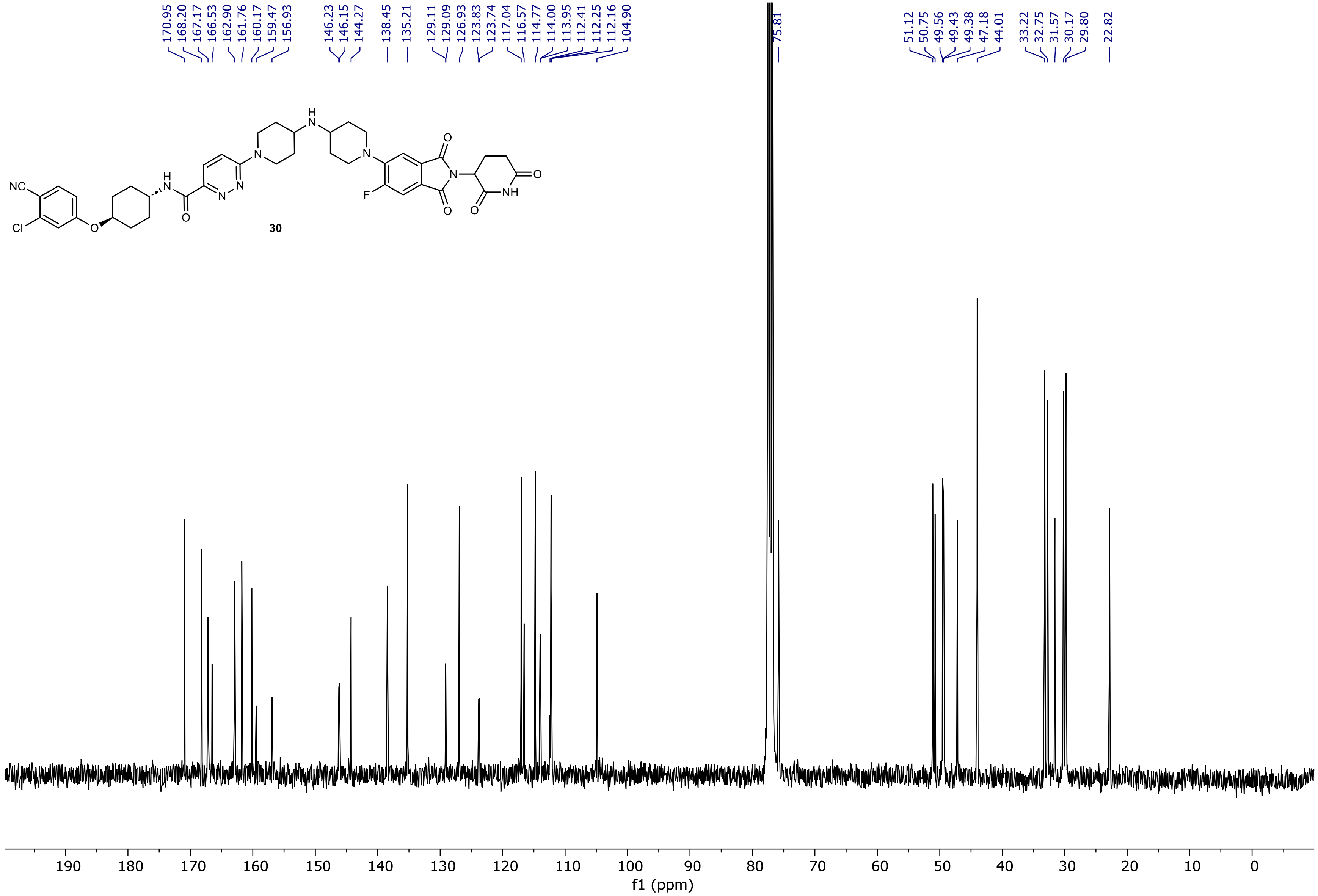

19F NMR (376.58 MHz, CDCl3, 298.1 K)

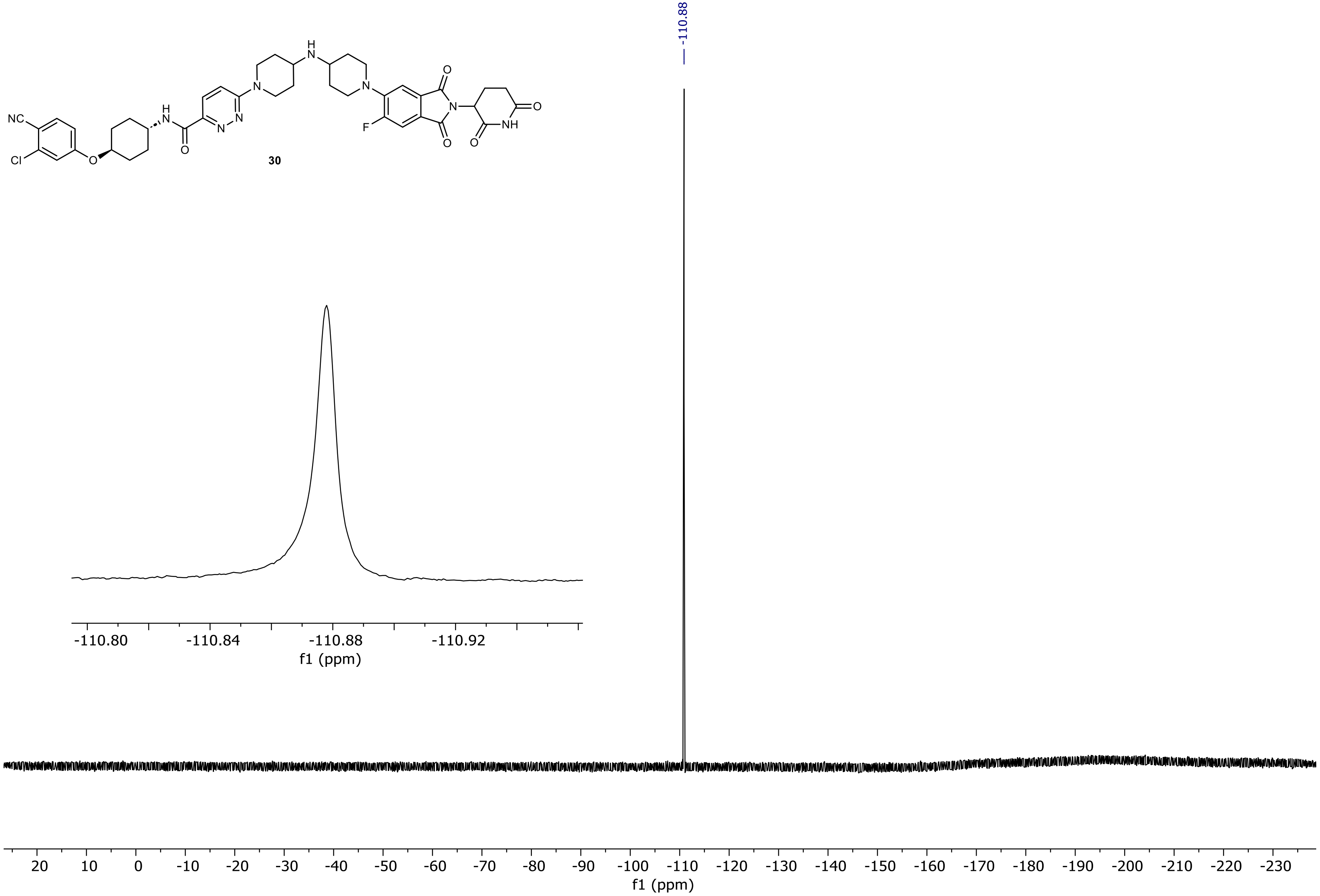

PDA - Total Absorbance Chromatogram

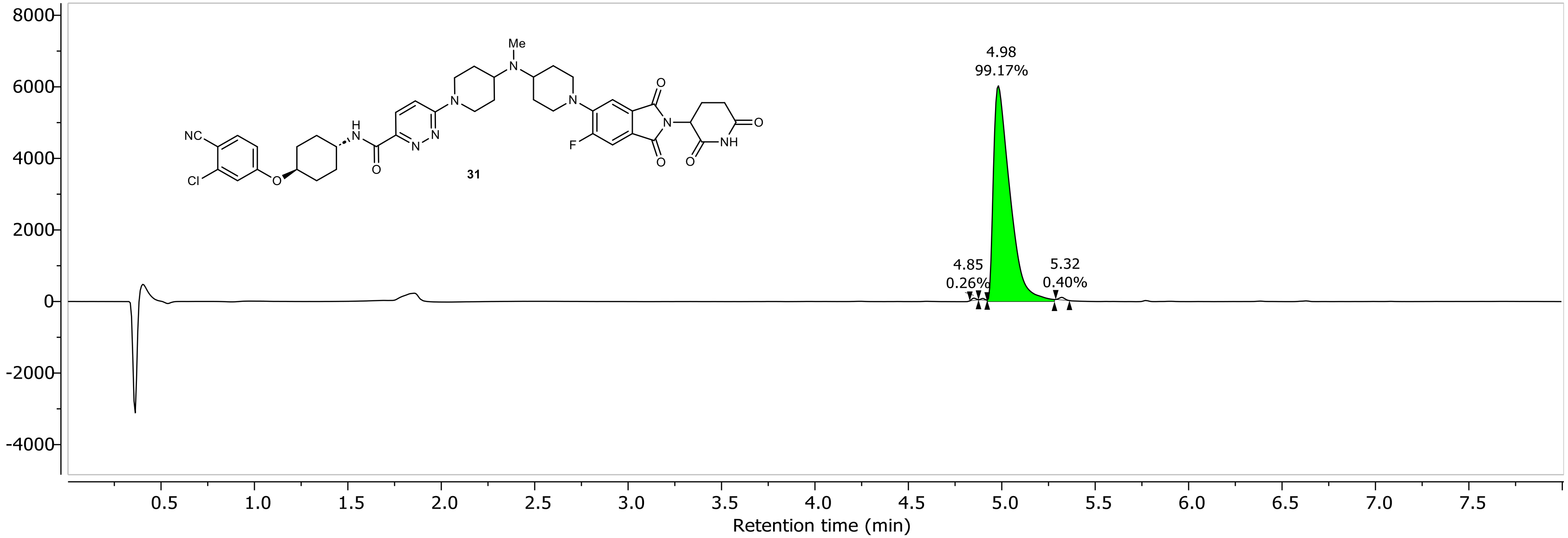

DAD1A, Sig=254...Ref=360.0,100.0 Chromatogram

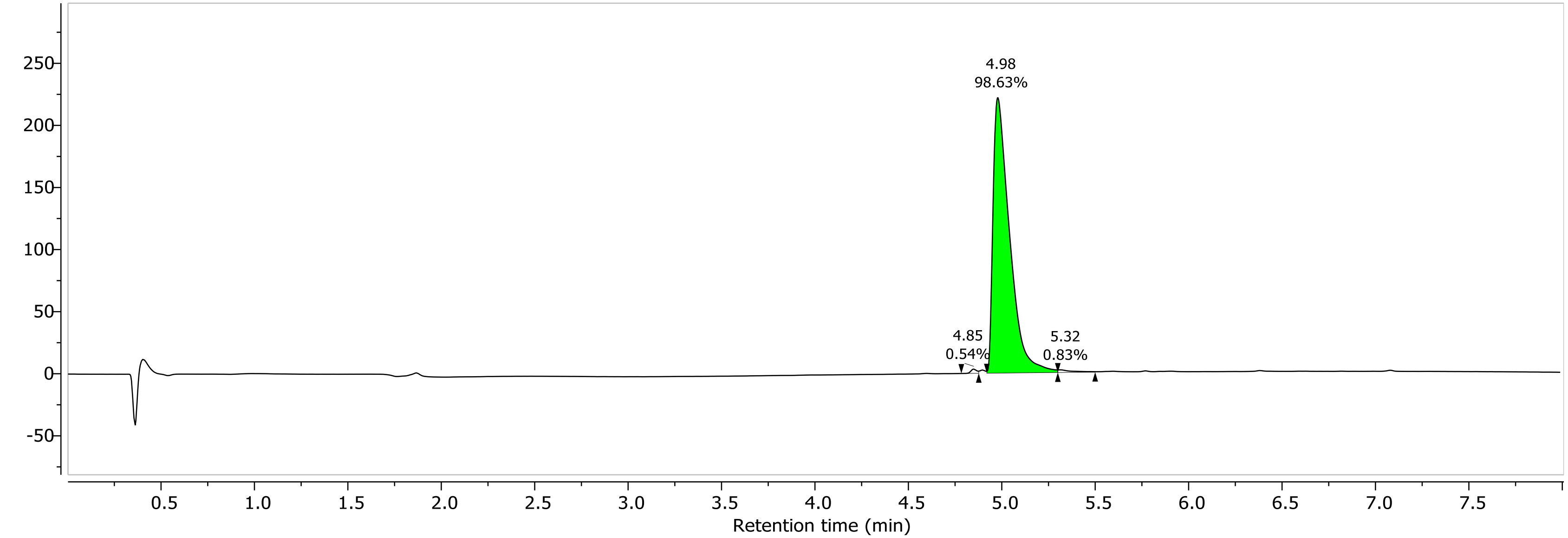

1H NMR (400.26 MHz, CDCl3, 298.1 K)

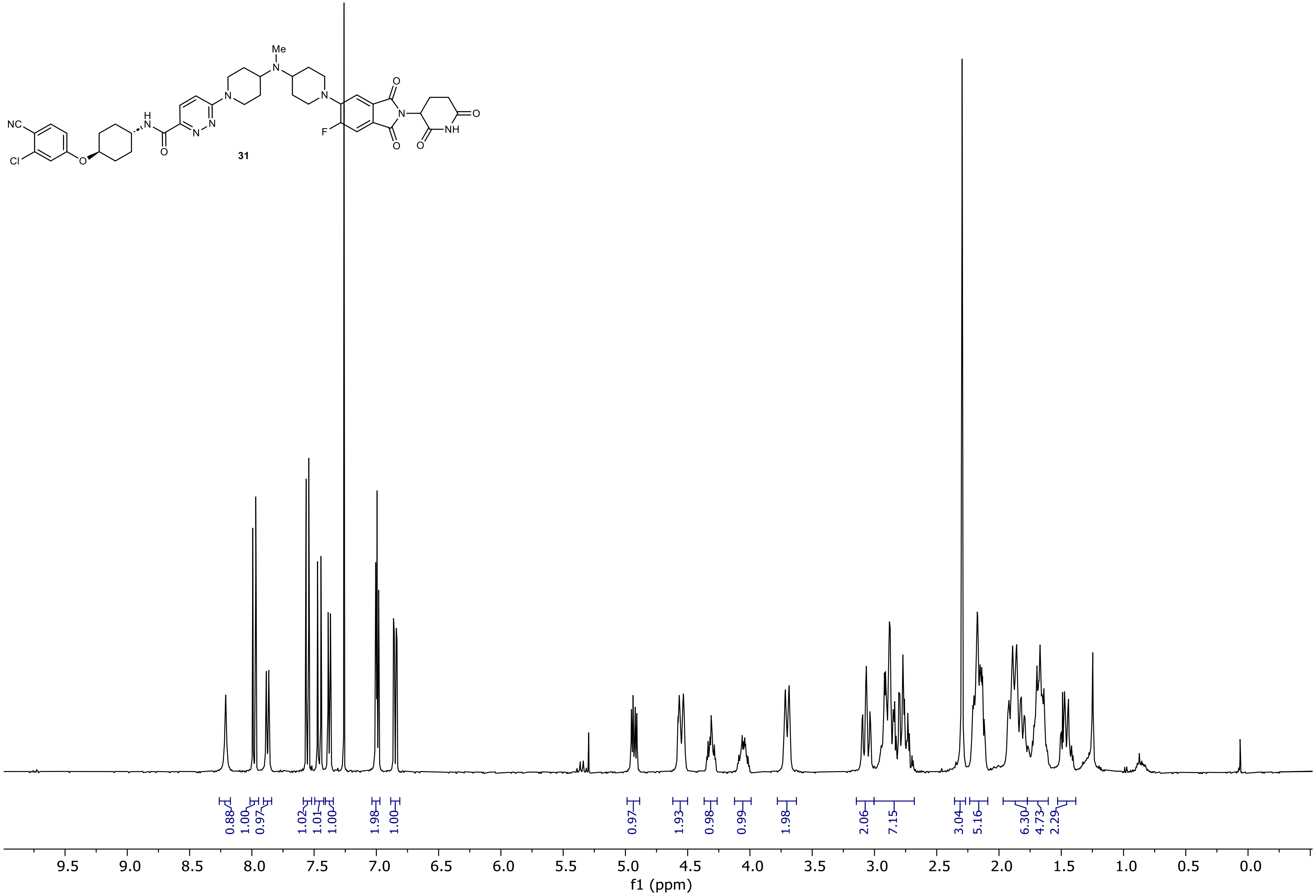

13C NMR (100.66 MHz, CDCl3, 298.1 K)

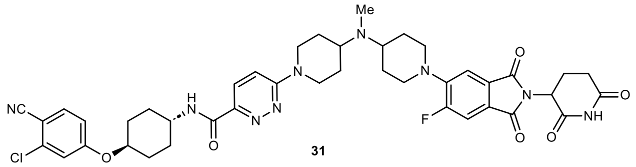

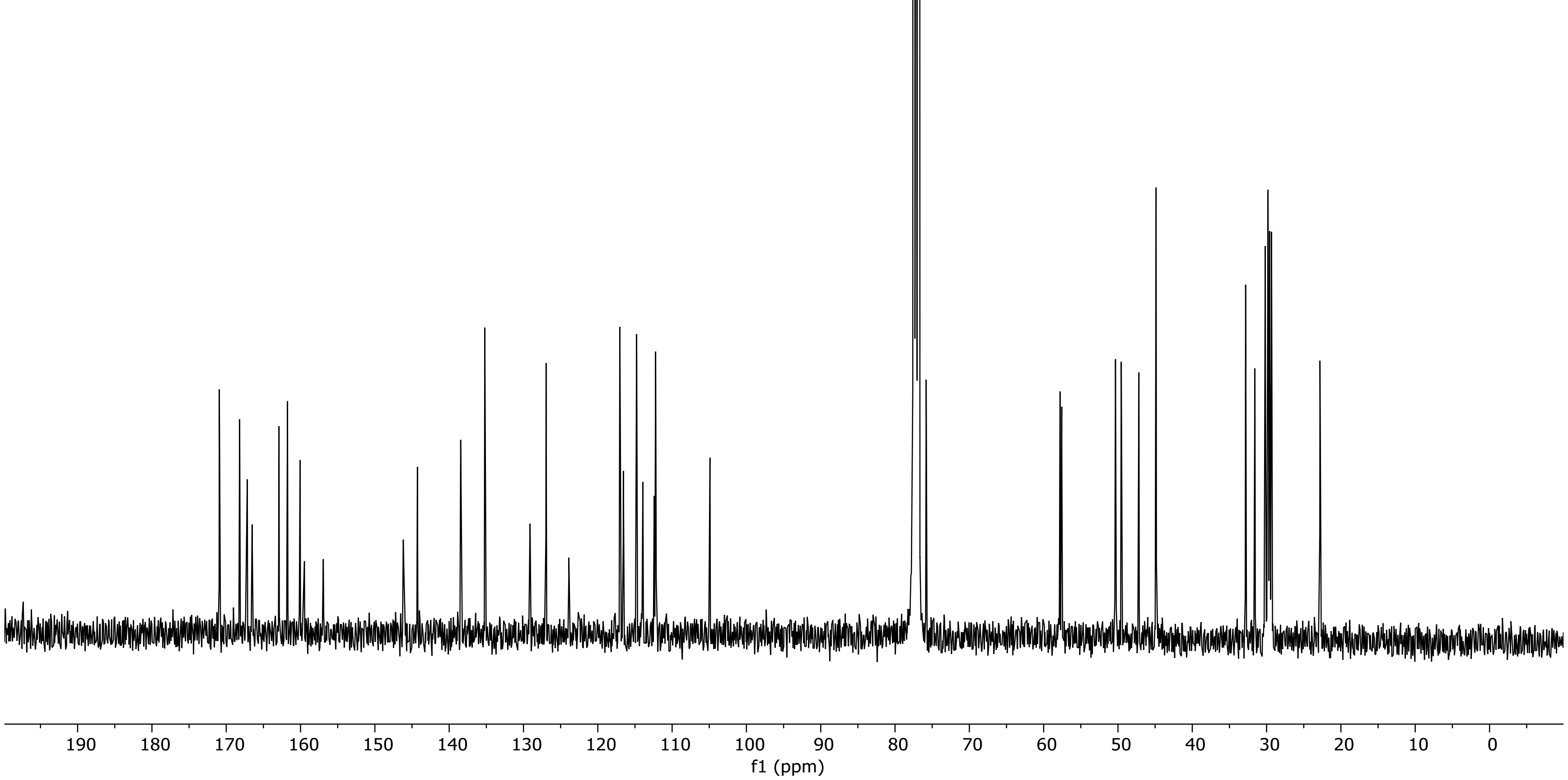

19F NMR (376.58 MHz, CDCl3, 298.2 K)

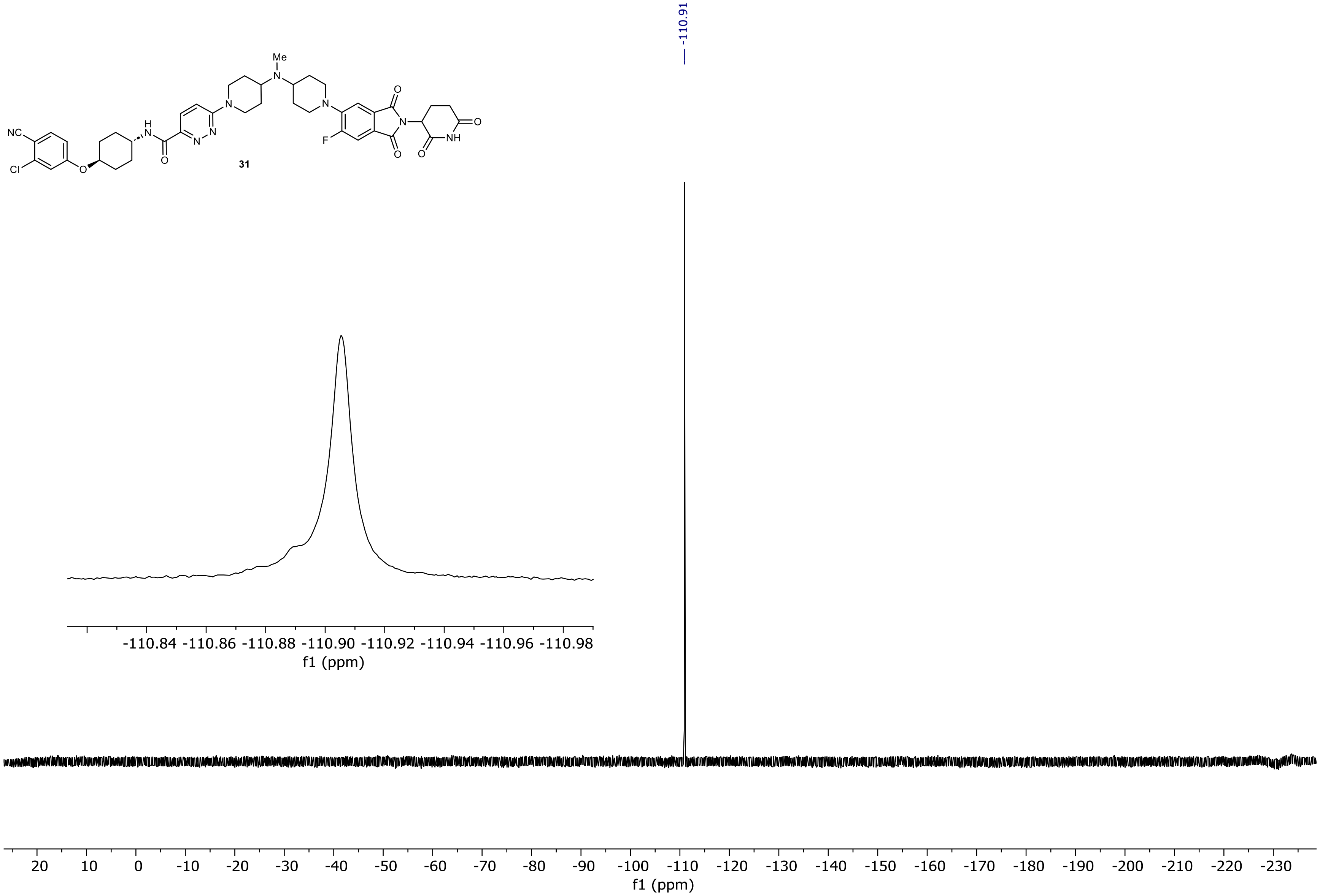

PDA - Total Absorbance Chromatogram

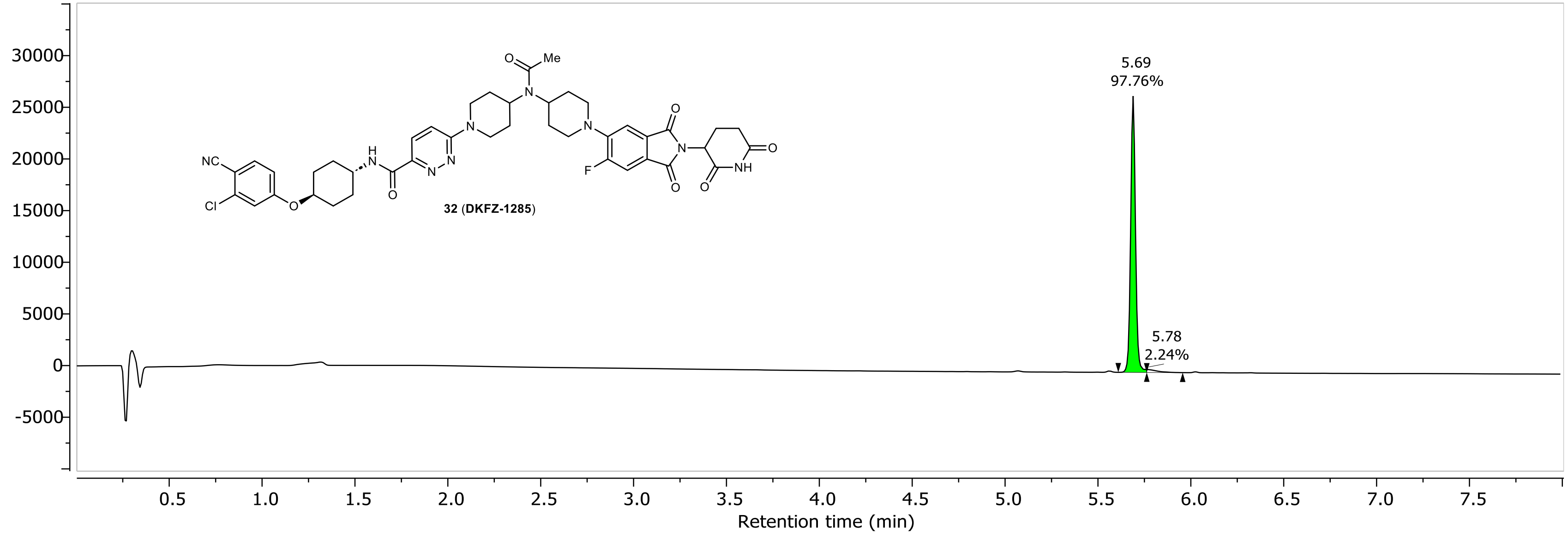

DAD1A, Sig=254...Ref=360.0,100.0 Chromatogram

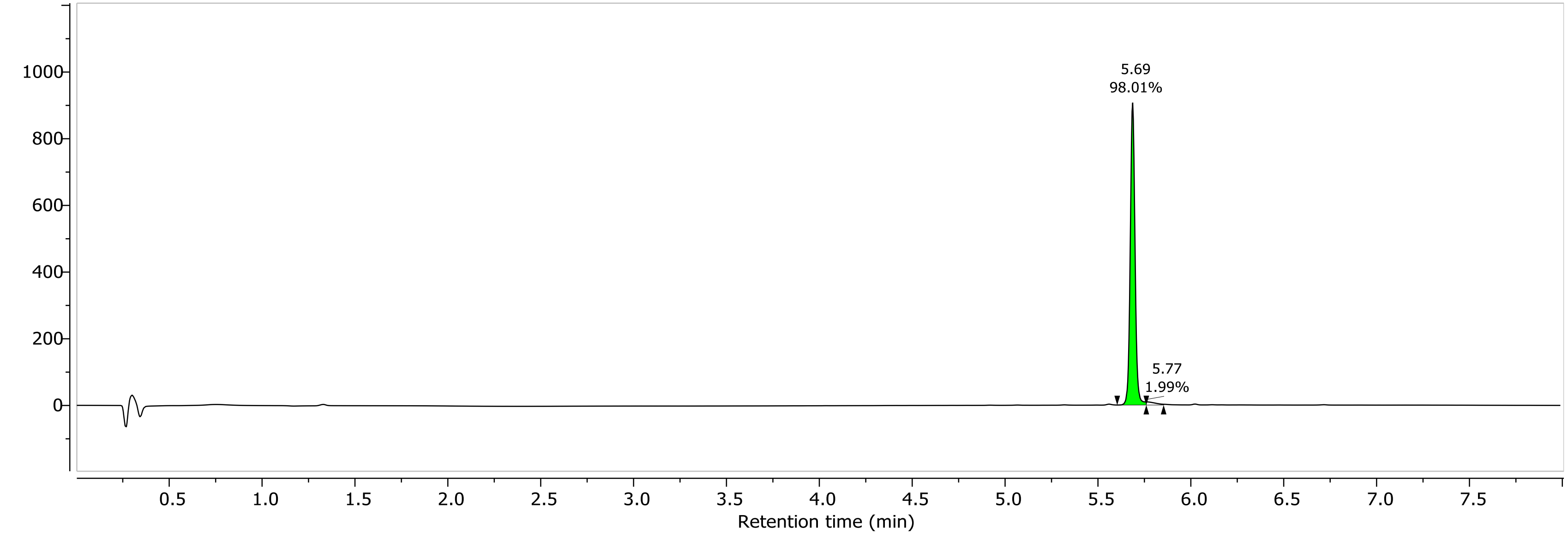

1H NMR (400.26 MHz, CDCl3, 328.1 K)

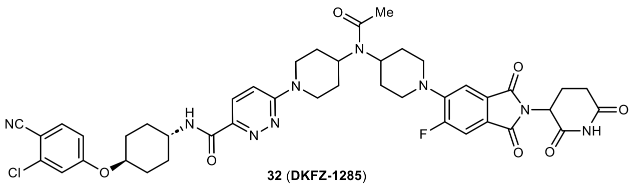

**32** (**DKFZ-1285**)

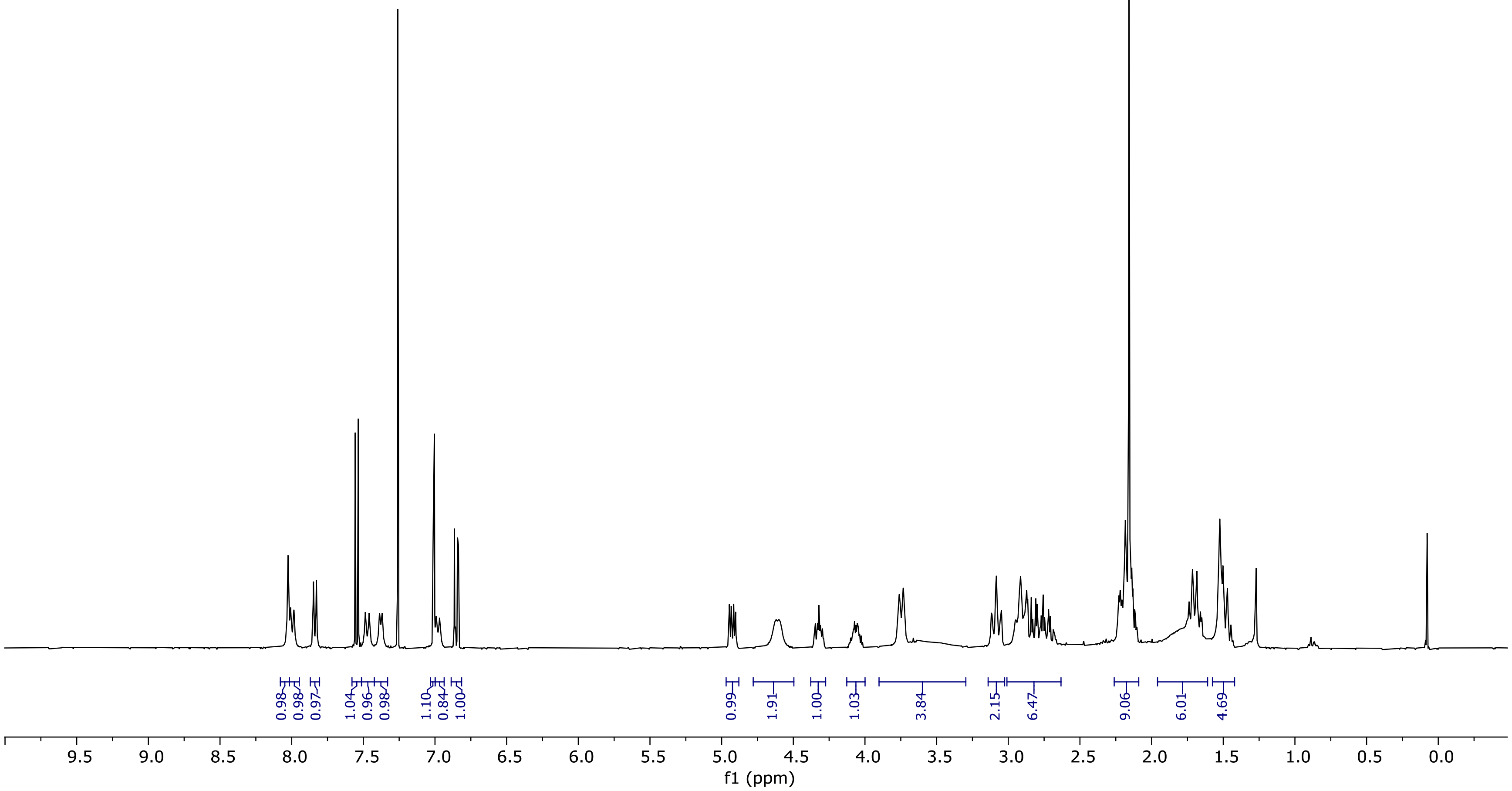

13C NMR (100.66 MHz, CDCl3, 328.1 K)

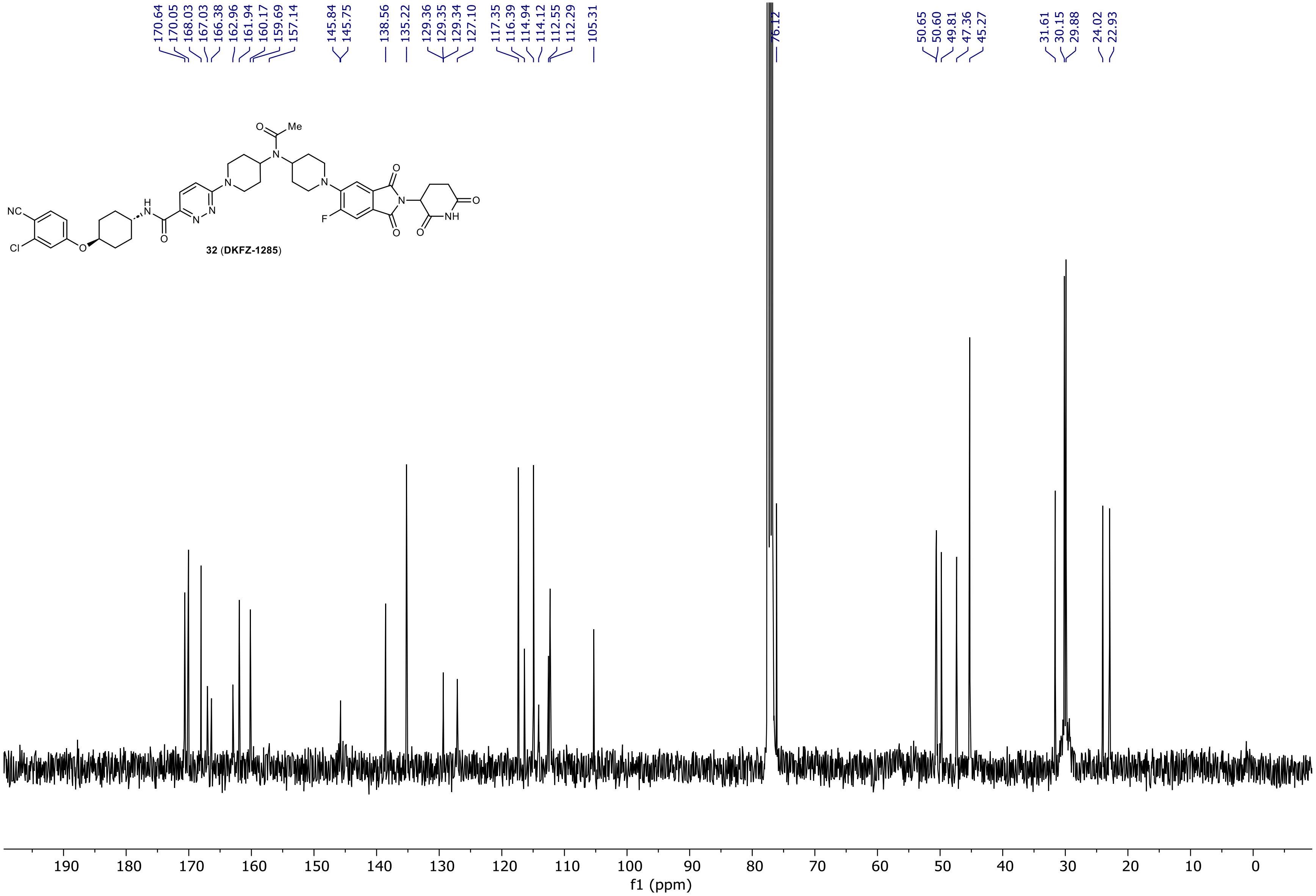

19F NMR (376.58 MHz, CDCl3, 328.2 K)

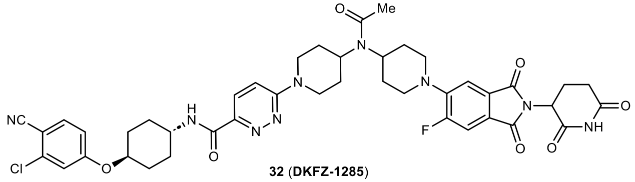

32 (DKFZ-1285)

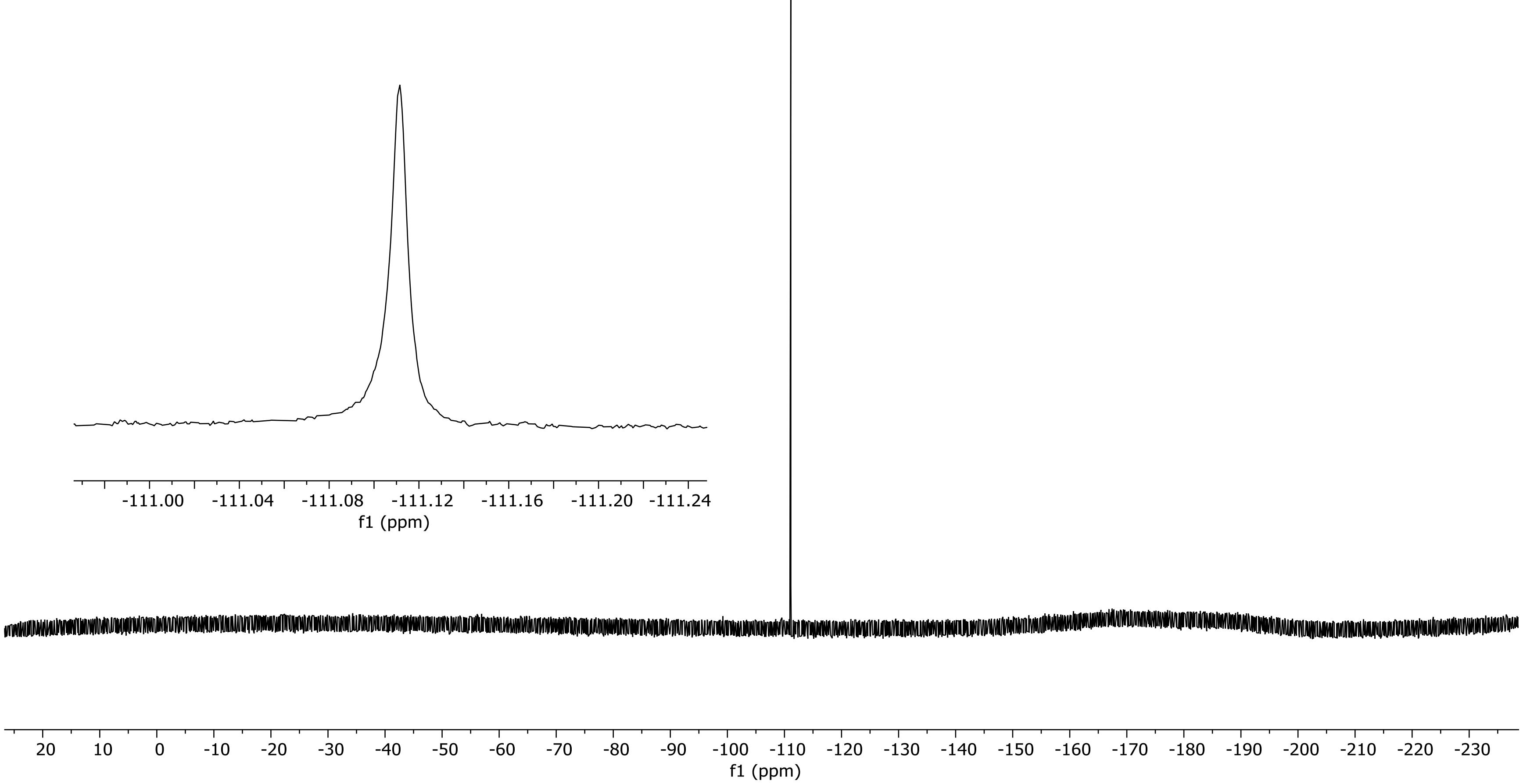

NR-6-090-A-corr LC-MS_ELS  PDA - Total Absorbance Chromatogram

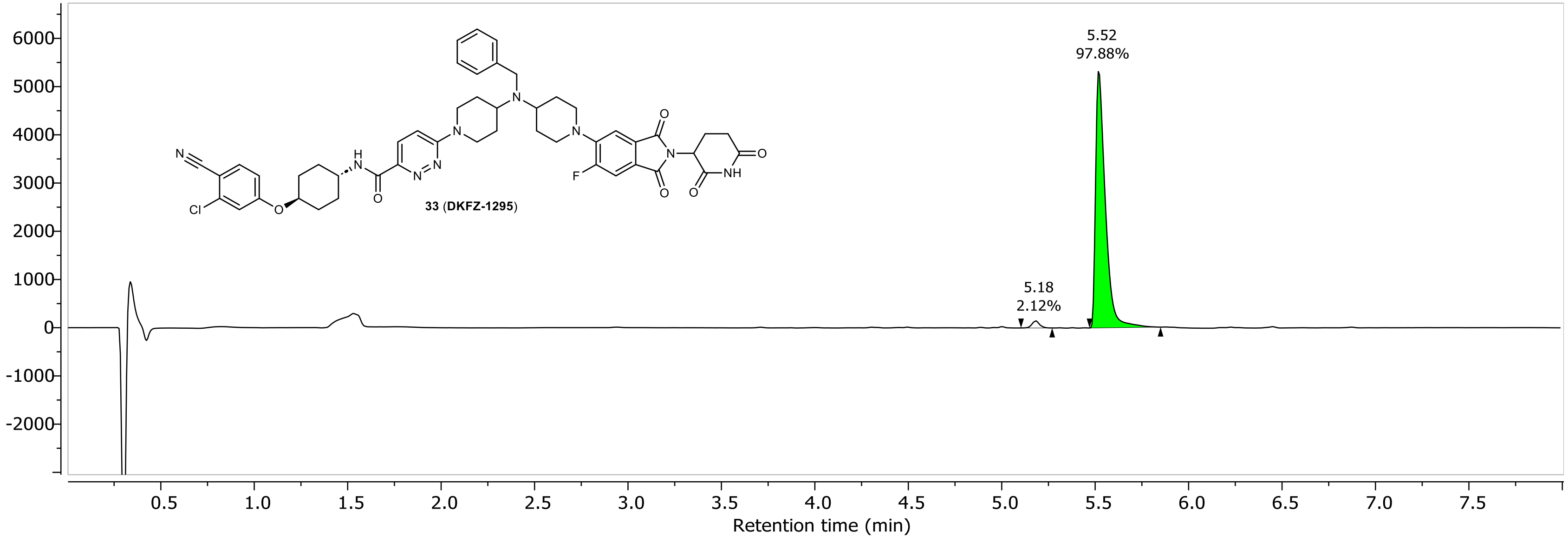

NR-6-090-A-corr LC-MS_ELS DAD1A, Sig=254...Ref=360.0,100.0 Chromatogram

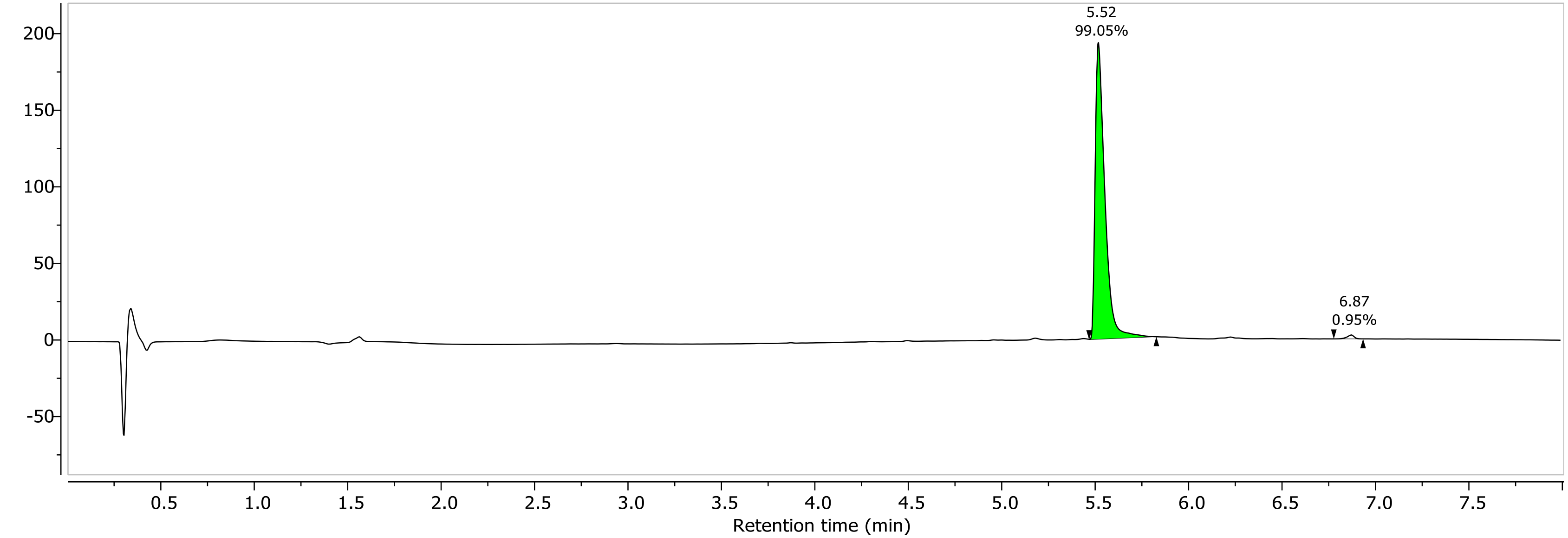

1H NMR (400.26 MHz, CDCl3, 298.1 K)

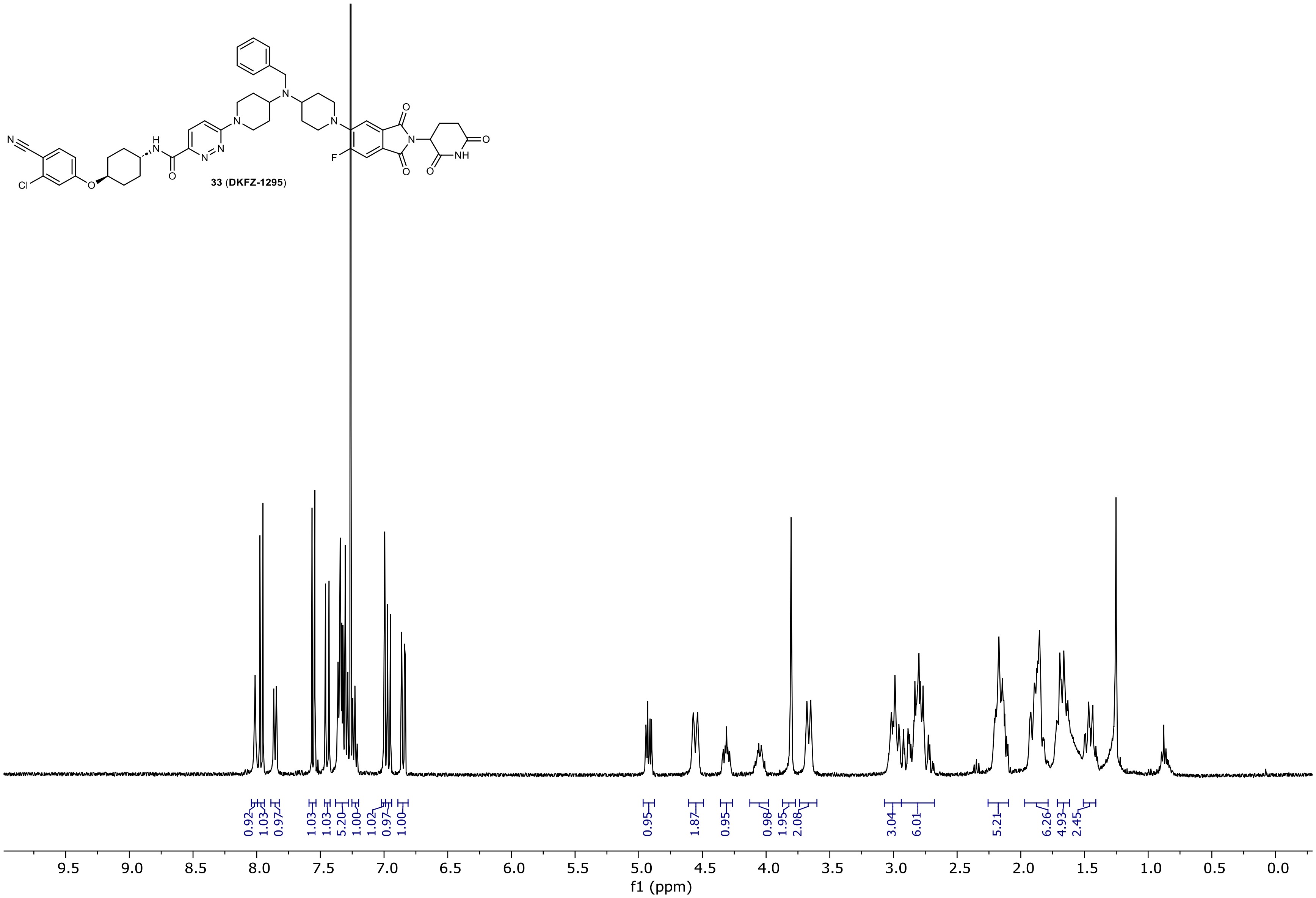

13C NMR (100.66 MHz, CDCl3, 298.1 K)

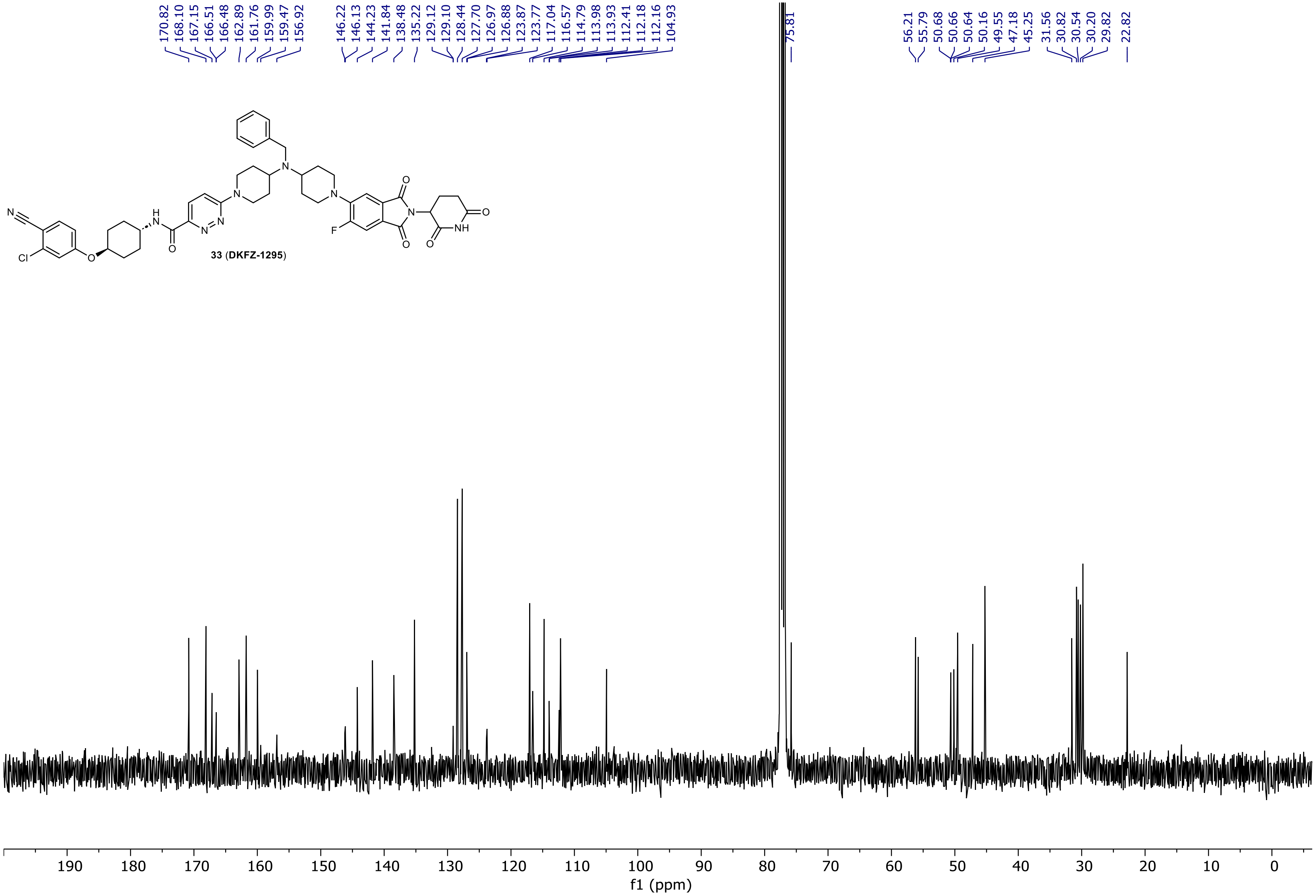

19F NMR (376.58 MHz, CDCl3, 298.2 K)

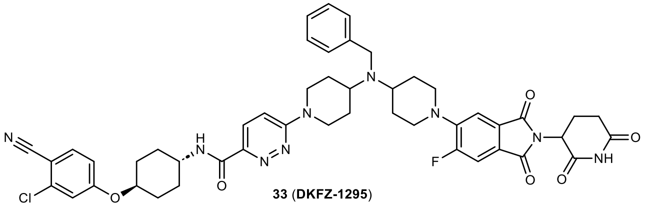

33 (DKFZ-1295)

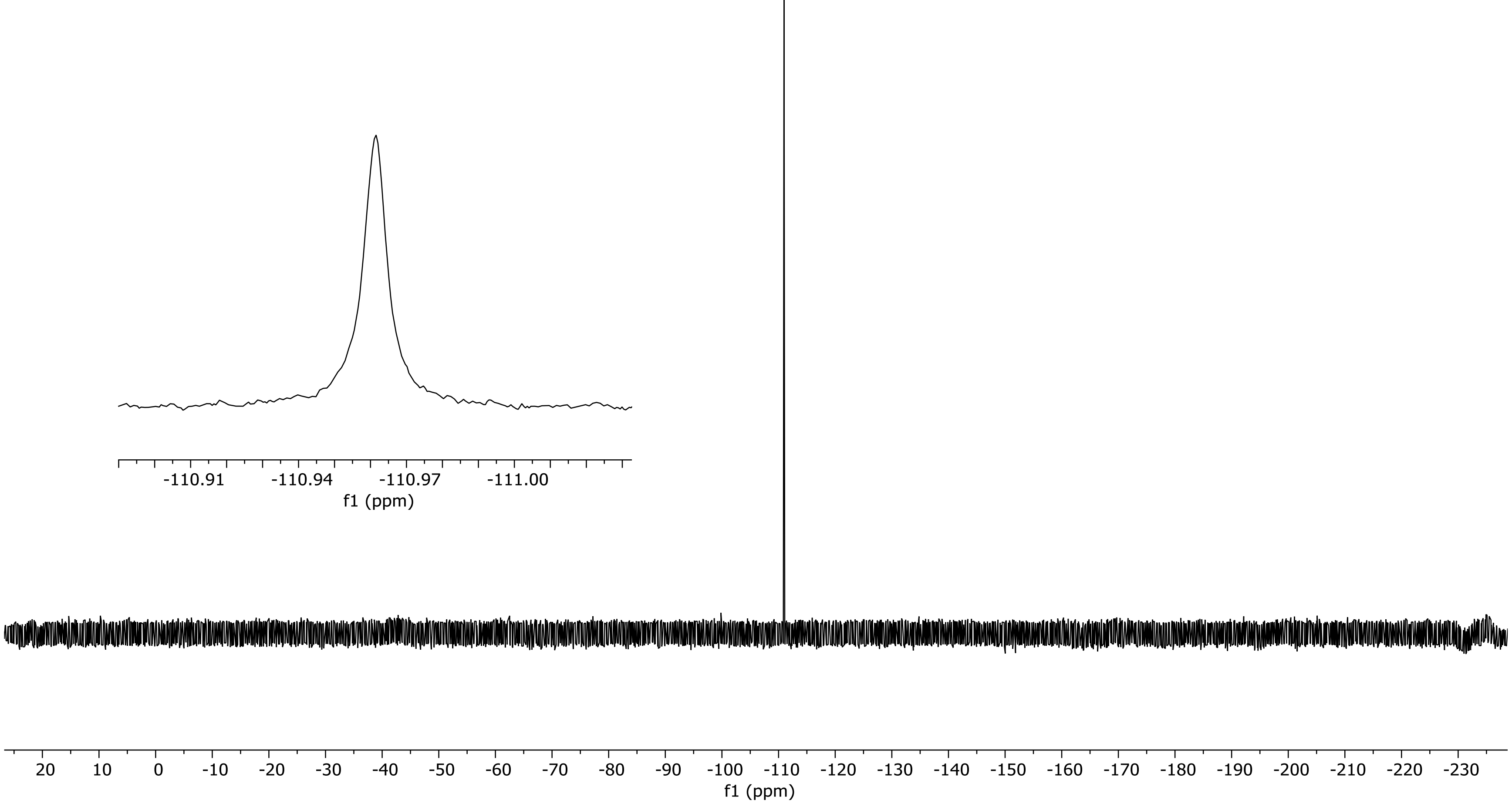

NR-6-069-A LC-MS_ELS  PDA - Total Absorbance Chromatogram

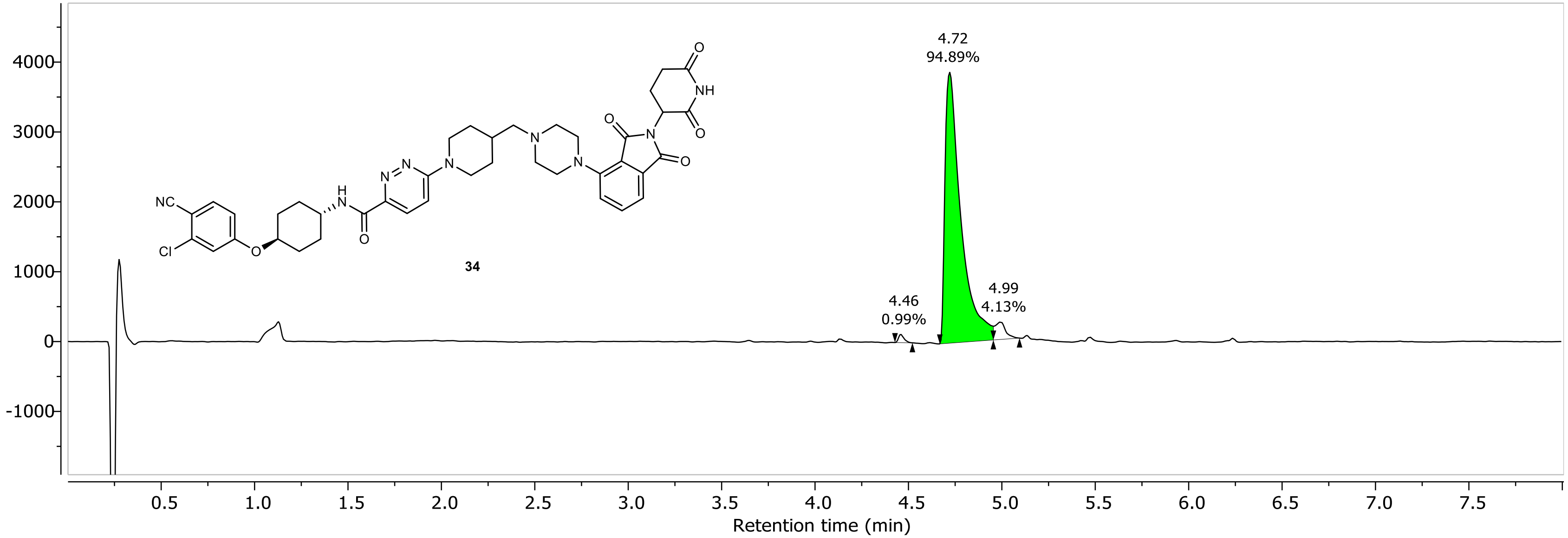

NR-6-069-A LC-MS_ELS DAD1A, Sig=254...Ref=360.0,100.0 Chromatogram

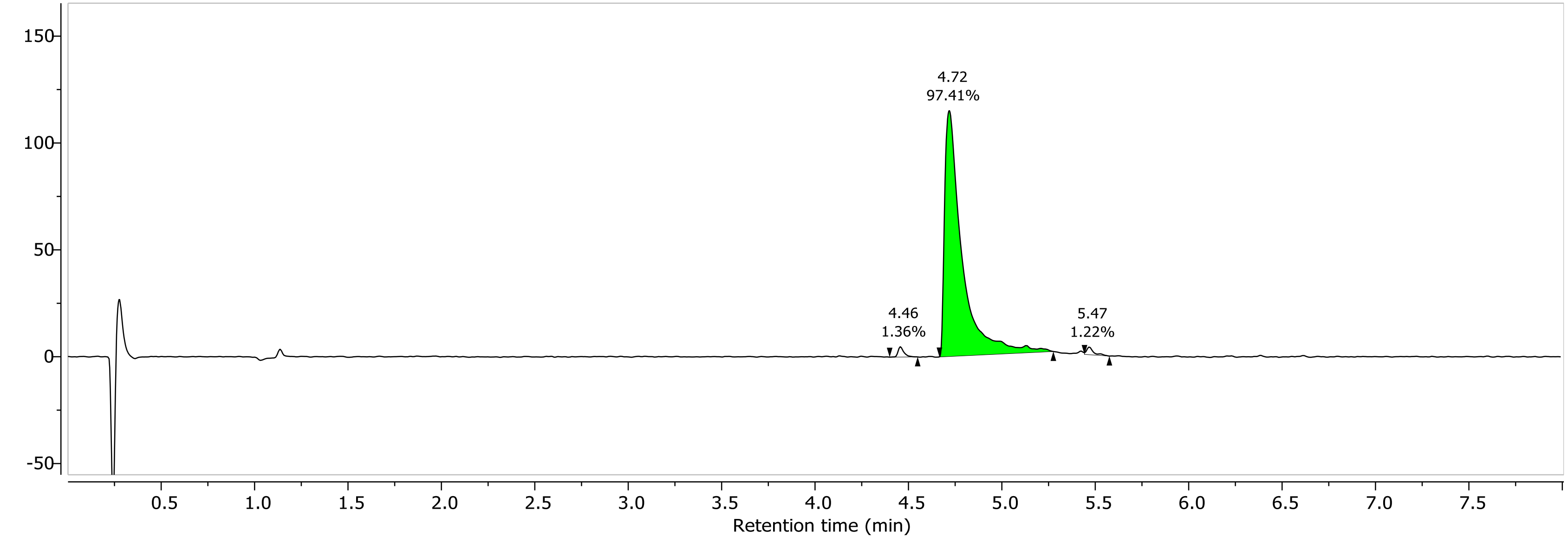

1H NMR (400.26 MHz, DMSO, 298.1 K)

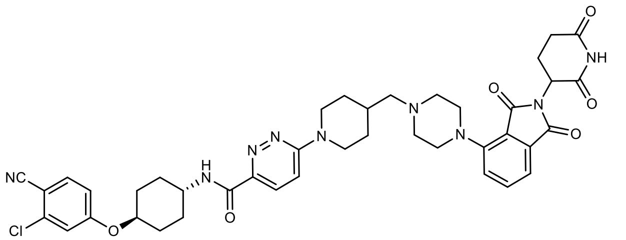

**34**

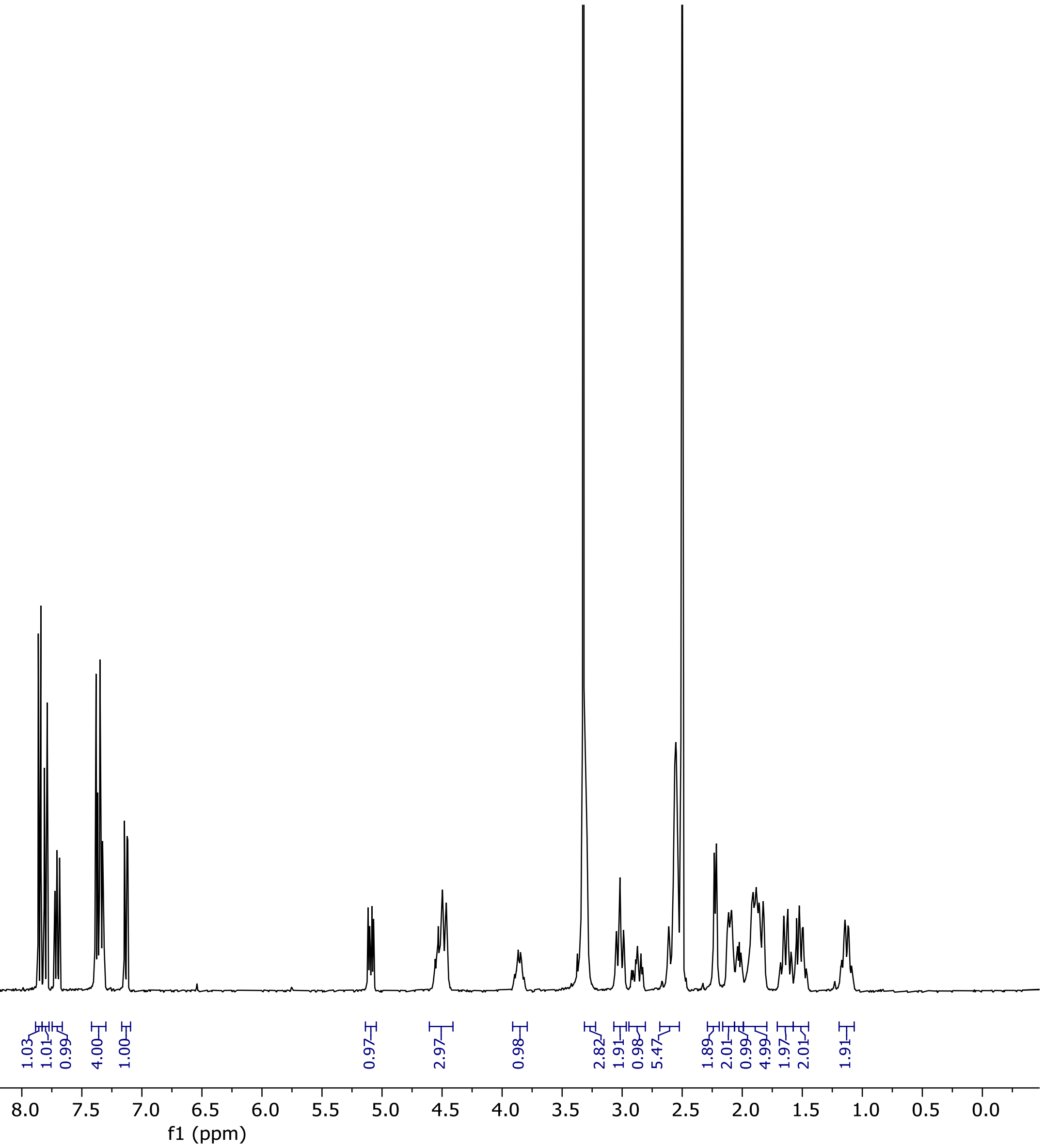

13C NMR (100.66 MHz, DMSO, 298.1 K)

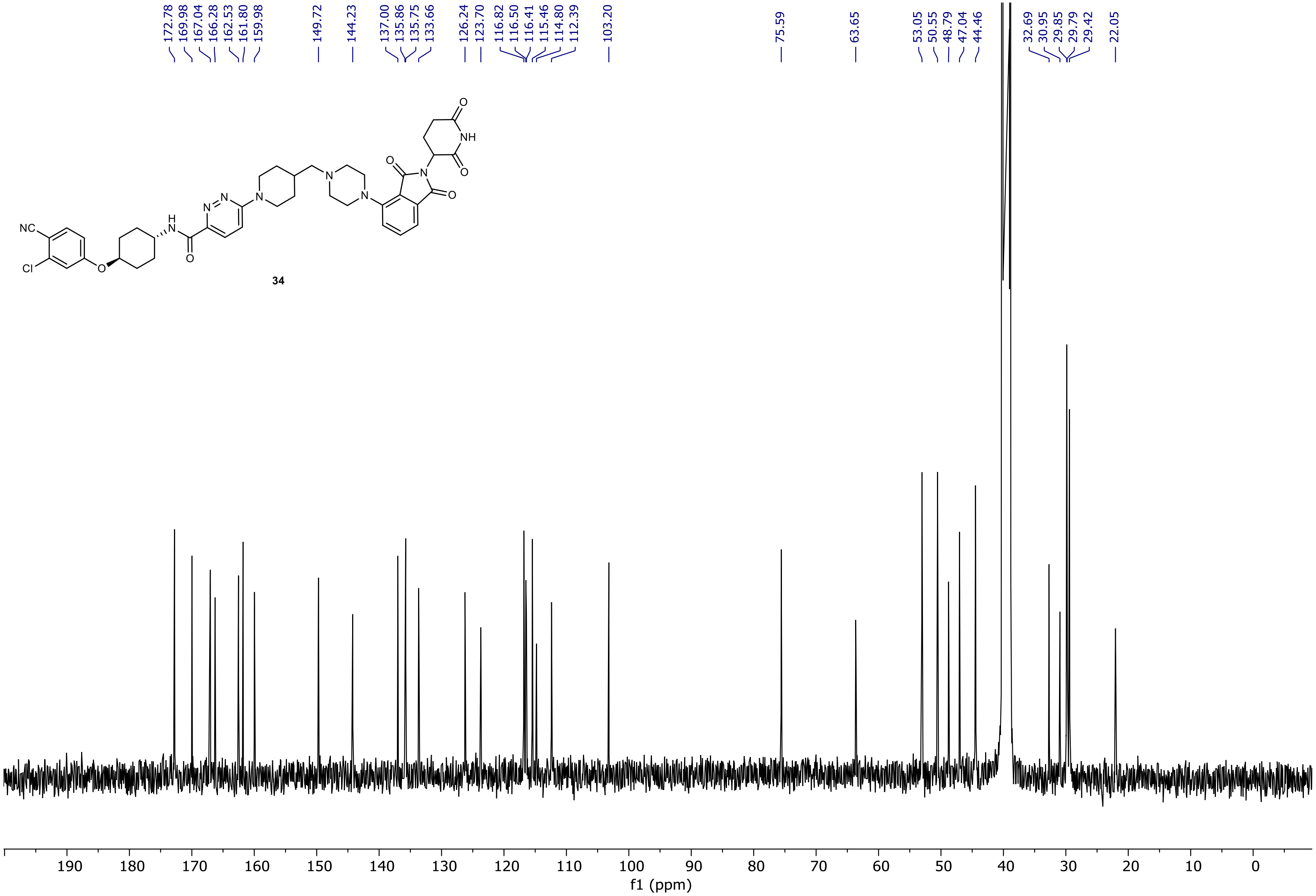

NR-6-103-C-f4 LC-MS_ELS  PDA - Total Absorbance Chromatogram

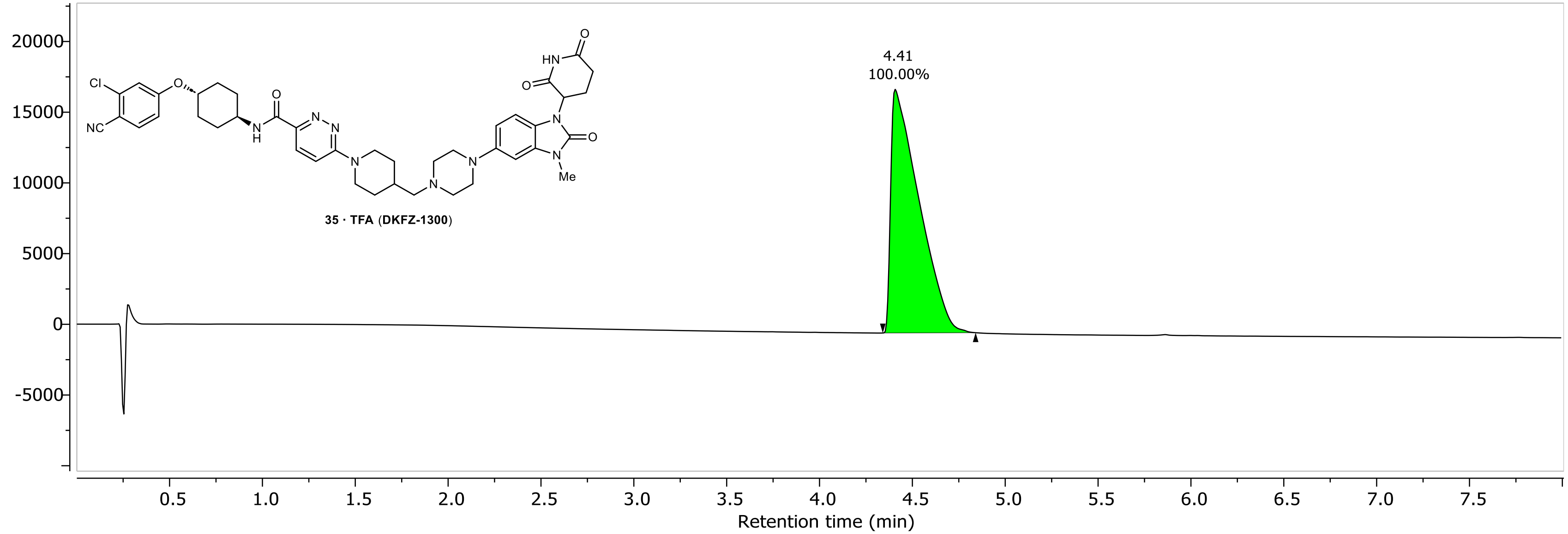

NR-6-103-C-f4 LC-MS_ELS DAD1A, Sig=254...Ref=360.0,100.0 Chromatogram

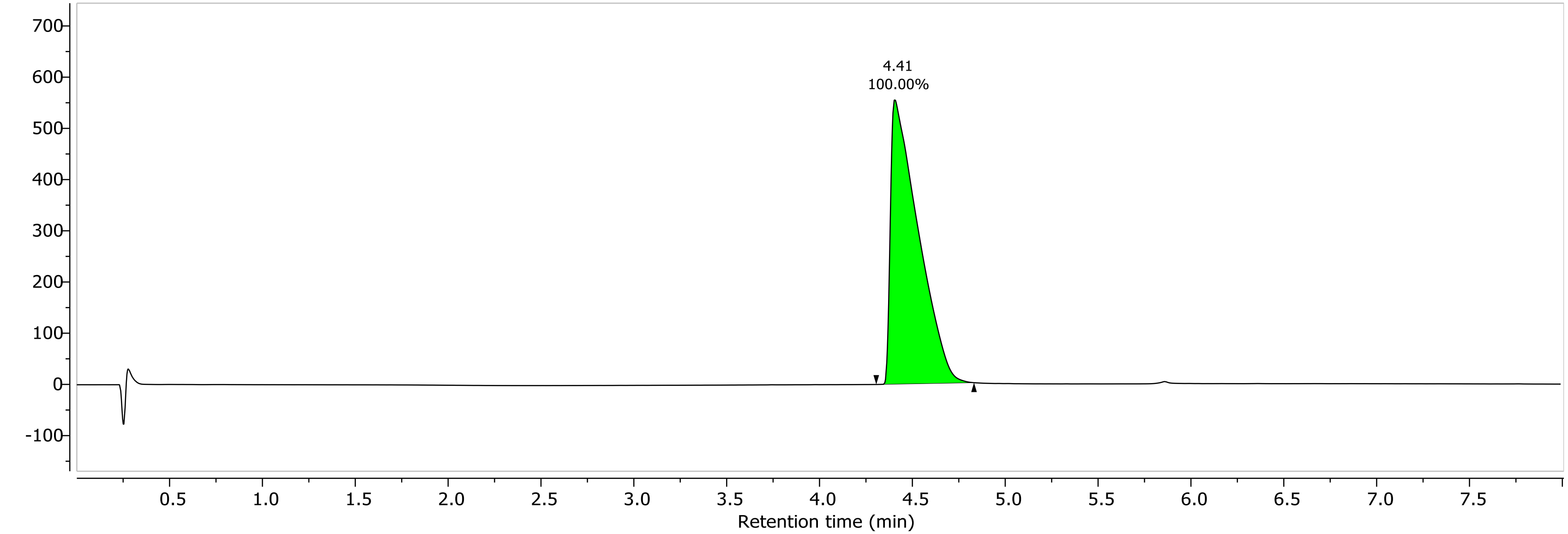

1H NMR (400.26 MHz, DMSO, 298.1 K)

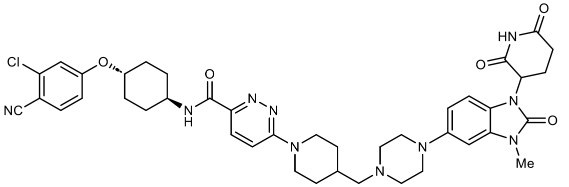

**35 · TFA (DKFZ-1300)**

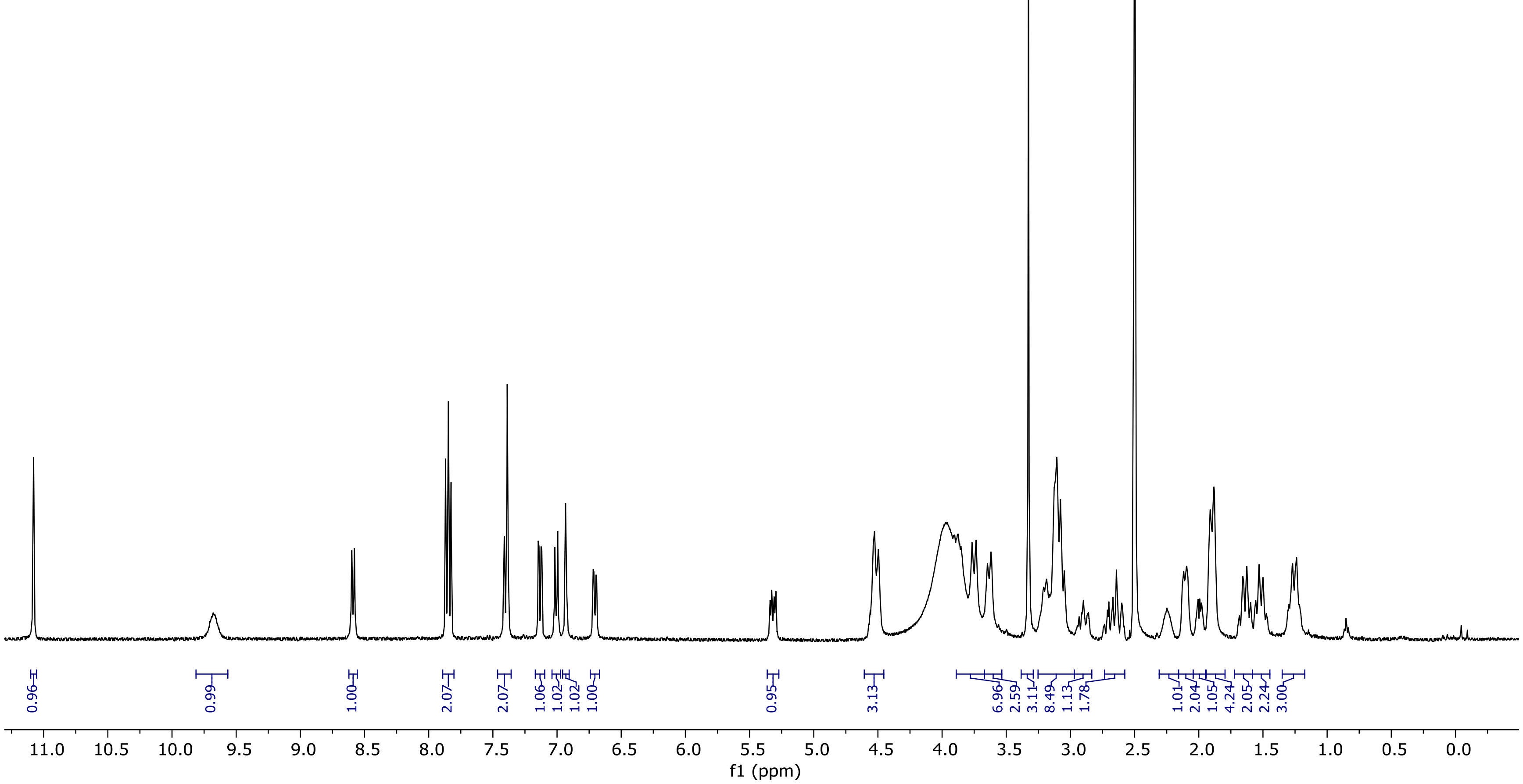

13C NMR (100.66 MHz, DMSO, 298.1 K)

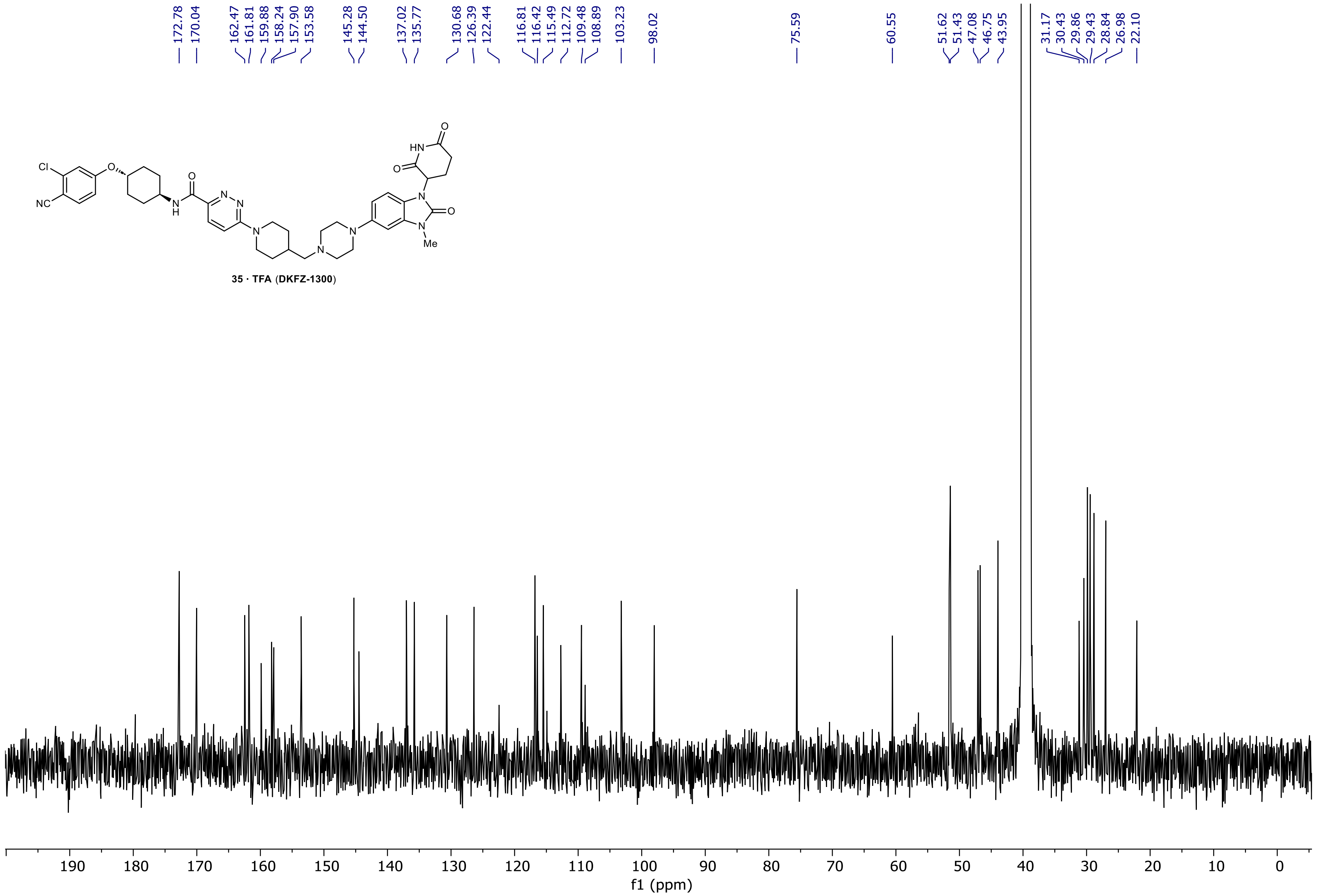
